## Supplementary material for "A novel eukaryotic RdRP-dependent small RNA pathway represses antiviral immunity by controlling an ERK pathway component in the black-legged tick": CDS_Relative.html

 


### CDS mapping table (Relative level compared to GFP control)

| ID | GFP | Ago-16 | Ago-30 | Ago-96 | Ago-78 | Aub | AGO3-1 | AGO3-2 | RdRP1 | RdRP3 | SUM\_KD |
| --- | --- | --- | --- | --- | --- | --- | --- | --- | --- | --- | --- |
| ISCI012234.RA | 1 | 0.42342196 | 1.42218036 | 0.87947184 | 1.9295286 | 1.51873925 | 1.36087145 | 1.45611576 | 1.15789739 | 0.28251798 | 6437.86105 |
| ISCI005507.RA | 1 | 1.36093553 | 0.7550073 | 1.24797125 | 1.02713914 | 0.43273448 | 0.52918475 | 0.60834739 | 1.59080199 | 1.12014133 | 3624.28861 |
| ISCI011233.RA | 1 | 1.3420158 | 1.18565991 | 0.94078484 | 1.06655343 | 0.14903487 | 0.36870337 | 0.42571768 | 1.91498092 | 1.57375048 | 2638.36889 |
| ISCI013597.RA | 1 | 1.18280005 | 1.02396499 | 1.09105796 | 1.21966699 | 0.817011 | 0.89475289 | 1.07697389 | 1.75069076 | 1.34382898 | 2584.51701 |
| ISCI009240.RA | 1 | 0.93010923 | 1.18977372 | 1.52927103 | 1.43322493 | 1.79573247 | 1.58856255 | 1.33401054 | 0.10252305 | 1.154665 | 2436.48548 |
| ISCI017315.RA | 1 | 0.66627823 | 0.95277939 | 1.05567753 | 1.09767857 | 0.77417672 | 0.52033636 | 0.62694015 | 1.48830685 | 0.79295692 | 1973.45298 |
| ISCI007738.RA | 1 | 1.36745049 | 1.0520388 | 1.09592545 | 1.4007239 | 0.60923367 | 0.18499885 | 0.20212685 | 2.04038035 | 1.60563204 | 1574.67285 |
| ISCI024113.RA | 1 | 1.10989339 | 0.95040506 | 1.06765744 | 0.94685957 | 0.86427168 | 0.50986376 | 0.5786106 | 1.40600087 | 1.1231206 | 1515.60895 |
| ISCI024064.RA | 1 | 1.00205487 | 0.9863126 | 1.18671181 | 1.65343587 | 0.93443159 | 0.80170654 | 1.05148292 | 1.62632229 | 1.01380339 | 1305.92335 |
| ISCI024649.RA | 1 | 1.09266976 | 1.0995202 | 0.94850118 | 0.94818661 | 1.11701215 | 0.55114066 | 0.71952334 | 1.5016125 | 1.16754922 | 1305.61972 |
| ISCI010315.RA | 1 | 1.31911424 | 1.19499245 | 1.00797738 | 0.91591494 | 0.33293701 | 0.32443461 | 0.30616597 | 1.67773277 | 1.32344294 | 1002.29043 |
| ISCI024767.RA | 1 | 1.19903982 | 1.08031561 | 0.89548956 | 1.01018224 | 0.48331819 | 0.41038743 | 0.48830264 | 1.42944309 | 1.11466525 | 928.143144 |
| ISCI017059.RA | 1 | 0.54770171 | 1.07862756 | 0.89875261 | 1.01790806 | 2.34814585 | 0.78128853 | 1.11129414 | 1.73502707 | 0.91113701 | 875.468496 |
| ISCI003449.RA | 1 | 1.14855157 | 0.98549288 | 0.91587585 | 0.54785985 | 1.1682167 | 0.37507589 | 0.33633141 | 1.39257374 | 0.93441614 | 852.163311 |
| ISCI024070.RA | 1 | 0.88000988 | 0.9688217 | 0.9298142 | 1.06346148 | 1.5633218 | 0.6626857 | 1.00828043 | 1.52756791 | 1.14825695 | 851.914112 |
| ISCI009768.RA | 1 | 0.2692515 | 0.9027875 | 1.12520153 | 0.67510223 | 1.01580428 | 1.03086667 | 1.2922099 | 1.56734257 | 0.24780344 | 805.852331 |
| ISCI002834.RA | 1 | 1.18601878 | 1.11372338 | 0.93764298 | 1.0846214 | 0.50469128 | 0.36918691 | 0.37488545 | 1.16321758 | 0.93846402 | 795.331658 |
| ISCI005745.RA | 1 | 0.59222336 | 1.06178774 | 0.81608761 | 0.96480569 | 0.88903502 | 0.97474661 | 1.10071127 | 1.53332819 | 0.87600492 | 773.690497 |
| ISCI019388.RA | 1 | 1.08533716 | 1.20433564 | 0.96518242 | 1.04327477 | 0.21626145 | 0.33340944 | 0.28286933 | 1.49689332 | 1.28831246 | 745.981703 |
| ISCI009034.RA | 1 | 1.06255085 | 1.28843356 | 1.47492414 | 1.51200876 | 1.8890358 | 1.68384108 | 1.30152321 | 0.23502856 | 1.44975838 | 731.14182 |
| ISCI021730.RA | 1 | 0.67190707 | 6.48894037 | 0.72988038 | 2.79525559 | 1.89391615 | 1.14124499 | 1.20239643 | 0.8084677 | 0.76074592 | 694.676419 |
| ISCI022167.RA | 1 | 1.0853487 | 1.01113084 | 1.05825381 | 1.65618546 | 1.6888114 | 1.16404365 | 1.27142399 | 0.29033307 | 0.75346422 | 689.127205 |
| ISCI010273.RA | 1 | 1.01261209 | 1.27727848 | 1.36817232 | 1.38117975 | 1.6068121 | 1.47102266 | 1.14327297 | 0.18996269 | 1.30873244 | 659.818586 |
| ISCI003367.RA | 1 | 0.82441332 | 1.15604142 | 0.97459227 | 1.31868628 | 13.739794 | 3.5432371 | 6.11463446 | 1.51757815 | 1.18645247 | 655.784848 |
| ISCI004588.RA | 1 | 1.31429759 | 1.02710508 | 0.87464769 | 0.94579357 | 0.29774858 | 0.45256813 | 0.42452786 | 1.52460337 | 1.34098014 | 648.031407 |
| ISCI017737.RA | 1 | 1.0522722 | 1.01645405 | 0.85230436 | 0.67089216 | 0.72473946 | 0.6461504 | 0.92729173 | 0.93125348 | 0.77792725 | 607.22761 |
| ISCI011355.RA | 1 | 0.8899317 | 1.05404667 | 1.15167585 | 1.4607084 | 0.47942147 | 0.40927712 | 0.52744169 | 1.31521151 | 1.05447755 | 525.10613 |
| ISCI006668.RA | 1 | 1.12219307 | 1.19034981 | 0.91589779 | 0.88795381 | 0.16842028 | 0.35589447 | 0.27048823 | 1.46113868 | 1.19882602 | 523.660869 |
| ISCI015077.RA | 1 | 1.59820355 | 1.1647996 | 1.18964071 | 1.96574386 | 0.1840426 | 0.39551749 | 0.48397638 | 2.00181324 | 1.75635063 | 485.477834 |
| ISCI012349.RA | 1 | 1.13305235 | 1.22132699 | 0.96206628 | 1.00539879 | 0.24447924 | 0.33099745 | 0.33942888 | 1.19458733 | 0.97747372 | 483.458157 |
| ISCI017738.RA | 1 | 1.1141172 | 1.26621667 | 1.14458925 | 1.22762375 | 0.54482402 | 0.45780021 | 0.48380134 | 1.47945321 | 1.18119998 | 482.586191 |
| ISCI024452.RA | 1 | 0.79812161 | 1.20470618 | 0.96848694 | 1.21379824 | 0.860082 | 0.98659318 | 0.903483 | 1.70650373 | 1.34771656 | 449.491754 |
| ISCI018019.RA | 1 | 1.17981451 | 0.88148605 | 0.97197535 | 0.99018339 | 1.06996479 | 0.48471973 | 0.71010819 | 1.29526363 | 1.09473606 | 442.54089 |
| ISCI009817.RA | 1 | 1.13254187 | 1.16897775 | 1.09098164 | 1.07434602 | 1.72504351 | 0.87007622 | 0.89235993 | 1.70657855 | 1.41124079 | 434.770077 |
| ISCI011625.RA | 1 | 1.20291042 | 1.22818539 | 0.97373619 | 0.98627391 | 2.23995661 | 0.67364933 | 1.05295616 | 1.58330442 | 1.43210888 | 422.531951 |
| ISCI013345.RA | 1 | 1.10650148 | 1.07199218 | 0.80470435 | 0.62330346 | 0.0673543 | 0.17518363 | 0.11318085 | 1.59755554 | 1.18428709 | 375.016615 |
| ISCI007332.RA | 1 | 0.65117463 | 1.2467961 | 1.56547209 | 1.07195398 | 1.49220775 | 1.20892071 | 1.00134735 | 0.04848109 | 0.70720863 | 372.122398 |
| ISCI007053.RA | 1 | 0.27678184 | 1.34605825 | 1.15052821 | 1.12790415 | 1.25644108 | 1.25150575 | 1.49582595 | 1.05389842 | 0.28690078 | 366.031762 |
| ISCI001342.RA | 1 | 1.14232385 | 1.25876046 | 1.25717293 | 1.27682731 | 1.01846106 | 0.88571536 | 0.87342713 | 1.66031182 | 1.15143065 | 355.011236 |
