## Supplementary material for "A novel eukaryotic RdRP-dependent small RNA pathway represses antiviral immunity by controlling an ERK pathway component in the black-legged tick": CDS_RPM.html

 

### CDS mapping table (RPM)

| ID | GFP | Ago-16 | Ago-30 | Ago-96 | Ago-78 | Aub | AGO3-1 | AGO3-2 | RdRP1 | RdRP3 | SUM\_KD |
| --- | --- | --- | --- | --- | --- | --- | --- | --- | --- | --- | --- |
| ISCI012234.RA | 563.205736 | 238.473618 | 800.980179 | 495.323572 | 1086.72166 | 855.362706 | 766.45064 | 820.092792 | 652.134465 | 159.115678 | 6437.86105 |
| ISCI005507.RA | 374.709474 | 509.955474 | 282.908362 | 467.626675 | 384.878769 | 162.149651 | 198.290491 | 227.953493 | 596.088637 | 419.72758 | 3624.28861 |
| ISCI011233.RA | 264.705088 | 355.238444 | 313.850228 | 249.030528 | 282.322125 | 39.4502024 | 97.5975936 | 112.689577 | 506.905284 | 416.579815 | 2638.36889 |
| ISCI013597.RA | 226.697142 | 268.137409 | 232.129938 | 247.339729 | 276.495043 | 185.214041 | 202.837912 | 244.14691 | 396.876666 | 304.642224 | 2584.51701 |
| ISCI009240.RA | 202.065934 | 187.943385 | 240.412757 | 309.013633 | 289.605977 | 362.856439 | 320.994435 | 269.558119 | 20.7163258 | 233.318477 | 2436.48548 |
| ISCI017315.RA | 219.88015 | 146.501323 | 209.497269 | 232.122539 | 241.357738 | 170.22607 | 114.411588 | 137.851657 | 327.249181 | 174.355466 | 1973.45298 |
| ISCI007738.RA | 149.137779 | 203.938566 | 156.898736 | 163.443897 | 208.900891 | 90.8597181 | 27.5902359 | 30.1446695 | 304.297898 | 239.460458 | 1574.67285 |
| ISCI024113.RA | 158.591532 | 176.019704 | 150.726189 | 169.321436 | 150.163904 | 137.066156 | 80.8600266 | 91.7626992 | 222.979872 | 178.117429 | 1515.60895 |
| ISCI024064.RA | 116.017488 | 116.255889 | 114.429508 | 137.679341 | 191.827541 | 108.410399 | 93.0119588 | 121.990412 | 188.681889 | 117.618923 | 1305.92335 |
| ISCI024649.RA | 128.686803 | 140.612188 | 141.493749 | 122.05958 | 122.019098 | 143.744734 | 70.9244846 | 92.5931308 | 193.237761 | 150.248194 | 1305.61972 |
| ISCI010315.RA | 106.59589 | 140.612188 | 127.381304 | 107.446246 | 97.6327596 | 35.4896506 | 34.583329 | 32.6359645 | 178.839485 | 141.07361 | 1002.29043 |
| ISCI024767.RA | 101.869013 | 122.145024 | 110.050693 | 91.222628 | 102.906269 | 49.2350951 | 41.8057038 | 49.7428568 | 145.616 | 113.549861 | 928.143144 |
| ISCI017059.RA | 76.5946938 | 41.9509992 | 82.6171553 | 68.8396707 | 77.9663577 | 179.855647 | 59.8425339 | 85.1192458 | 132.893941 | 69.7882514 | 875.468496 |
| ISCI003449.RA | 96.788425 | 111.166513 | 95.3843019 | 88.6461725 | 53.0264466 | 113.069871 | 36.302942 | 32.5529213 | 134.785058 | 90.4406602 | 852.163311 |
| ISCI024070.RA | 79.2314549 | 69.724451 | 76.7611501 | 73.6705248 | 84.2596063 | 123.864317 | 52.5055182 | 79.8875263 | 121.03148 | 90.9780835 | 851.914112 |
| ISCI009768.RA | 88.2993406 | 23.7746564 | 79.7155311 | 99.3545657 | 59.6110493 | 89.69485 | 91.0248504 | 114.101311 | 138.395372 | 21.8808049 | 805.852331 |
| ISCI002834.RA | 91.7078366 | 108.767235 | 102.137173 | 85.9892027 | 99.4682904 | 46.2840957 | 33.8572701 | 34.379871 | 106.676184 | 86.0644992 | 795.331658 |
| ISCI005745.RA | 78.877743 | 46.713201 | 83.7514266 | 64.3711307 | 76.1016914 | 70.1250645 | 76.8858098 | 86.8216308 | 120.945521 | 69.0972786 | 773.690497 |
| ISCI019388.RA | 83.6689309 | 90.8090087 | 100.765496 | 80.7557775 | 87.289689 | 18.0942858 | 27.8959449 | 23.6673025 | 125.243514 | 107.791755 | 745.981703 |
| ISCI009034.RA | 56.6903632 | 60.2364001 | 73.0417954 | 83.6140328 | 85.7163768 | 107.090215 | 95.4576307 | 73.7838536 | 13.323778 | 82.1873742 | 731.14182 |
| ISCI021730.RA | 39.7121943 | 26.6828712 | 257.69061 | 28.9851245 | 111.005913 | 75.2116556 | 45.3213571 | 47.7498208 | 32.1060071 | 30.2108657 | 694.676419 |
| ISCI022167.RA | 62.7677759 | 68.1249328 | 63.4664354 | 66.4242437 | 103.955144 | 106.003005 | 73.0644475 | 79.8044832 | 18.22349 | 47.2932485 | 689.127205 |
| ISCI010273.RA | 56.111562 | 56.8192476 | 71.6701185 | 76.7703228 | 77.5001911 | 90.1607972 | 82.5414261 | 64.1508463 | 10.6590224 | 73.4350522 | 659.818586 |
| ISCI003367.RA | 20.9011549 | 17.231173 | 24.1626163 | 20.3701014 | 27.562098 | 287.178836 | 74.0580017 | 127.803434 | 31.7191877 | 24.7982455 | 655.784848 |
| ISCI004588.RA | 70.4208142 | 92.5539376 | 72.3295785 | 61.5933896 | 66.6035477 | 20.9676273 | 31.8701617 | 29.89554 | 107.363863 | 94.4329474 | 648.031407 |
| ISCI017737.RA | 70.6137479 | 74.3048894 | 71.7756321 | 60.1843905 | 47.374177 | 51.1765421 | 45.6270661 | 65.4795369 | 65.7592916 | 54.9323365 | 607.22761 |
| ISCI011355.RA | 56.2080289 | 50.0212954 | 59.2458911 | 64.7334447 | 82.1035859 | 26.9472839 | 23.0046011 | 29.6464105 | 73.9254781 | 59.2701101 | 525.10613 |
| ISCI006668.RA | 61.0956836 | 68.5611651 | 72.7252545 | 55.9573932 | 54.2501338 | 10.2896689 | 21.7435516 | 16.5255902 | 89.2693128 | 73.2431153 | 523.660869 |
| ISCI015077.RA | 41.3521311 | 66.0891824 | 48.1669622 | 49.1941974 | 81.2877945 | 7.61047212 | 16.3554307 | 20.0134032 | 82.7793435 | 72.6289173 | 485.477834 |
| ISCI012349.RA | 57.4942538 | 65.1440126 | 70.2193063 | 55.3132793 | 57.8046538 | 14.0560761 | 19.0303843 | 19.5151442 | 68.6819268 | 56.1991199 | 483.458157 |
| ISCI017738.RA | 48.7479244 | 54.3109123 | 61.7254609 | 55.7963647 | 59.8441325 | 26.5589945 | 22.3167559 | 23.5842593 | 72.1203211 | 57.5810655 | 482.586191 |
| ISCI024452.RA | 40.9019524 | 32.6447117 | 49.2748551 | 39.6130035 | 49.646739 | 35.1790191 | 40.3535861 | 36.9542092 | 69.7994049 | 55.1242734 | 449.491754 |
| ISCI018019.RA | 45.7252958 | 53.9473854 | 40.3061984 | 44.4438576 | 45.2764274 | 48.9244636 | 22.1639014 | 32.4698782 | 59.2263424 | 50.0571396 | 442.54089 |
| ISCI009817.RA | 36.0142977 | 40.7877133 | 42.0999297 | 39.2909466 | 38.6918247 | 62.126303 | 31.335171 | 32.1377055 | 61.4612987 | 50.8248872 | 434.770077 |
| ISCI011625.RA | 34.1492715 | 41.0785347 | 41.9416593 | 33.252379 | 33.6805342 | 76.4930106 | 23.0046011 | 35.9576912 | 54.0687509 | 48.9055183 | 422.531951 |
| ISCI013345.RA | 48.4263681 | 53.5838586 | 51.9126953 | 38.9688896 | 30.184285 | 3.26163091 | 8.48342434 | 5.480849 | 77.3638725 | 57.3507412 | 375.016615 |
| ISCI007332.RA | 37.2362113 | 24.2472413 | 46.4259877 | 58.292306 | 39.915512 | 55.5642123 | 45.0156481 | 37.2863818 | 1.80515702 | 26.3337406 | 372.122398 |
| ISCI007053.RA | 35.7248971 | 9.88793049 | 48.087827 | 41.1025168 | 40.2942723 | 44.8862539 | 44.7099391 | 53.4382778 | 37.6504179 | 10.2494297 | 366.031762 |
| ISCI001342.RA | 30.8050868 | 35.1893997 | 38.7762511 | 38.7273469 | 39.3328038 | 31.373783 | 27.2845269 | 26.905986 | 51.1461157 | 35.4699363 | 355.011236 |
