## Supplementary material for "A novel eukaryotic RdRP-dependent small RNA pathway represses antiviral immunity by controlling an ERK pathway component in the black-legged tick": 25-30_TE_readcount_libs.pdf

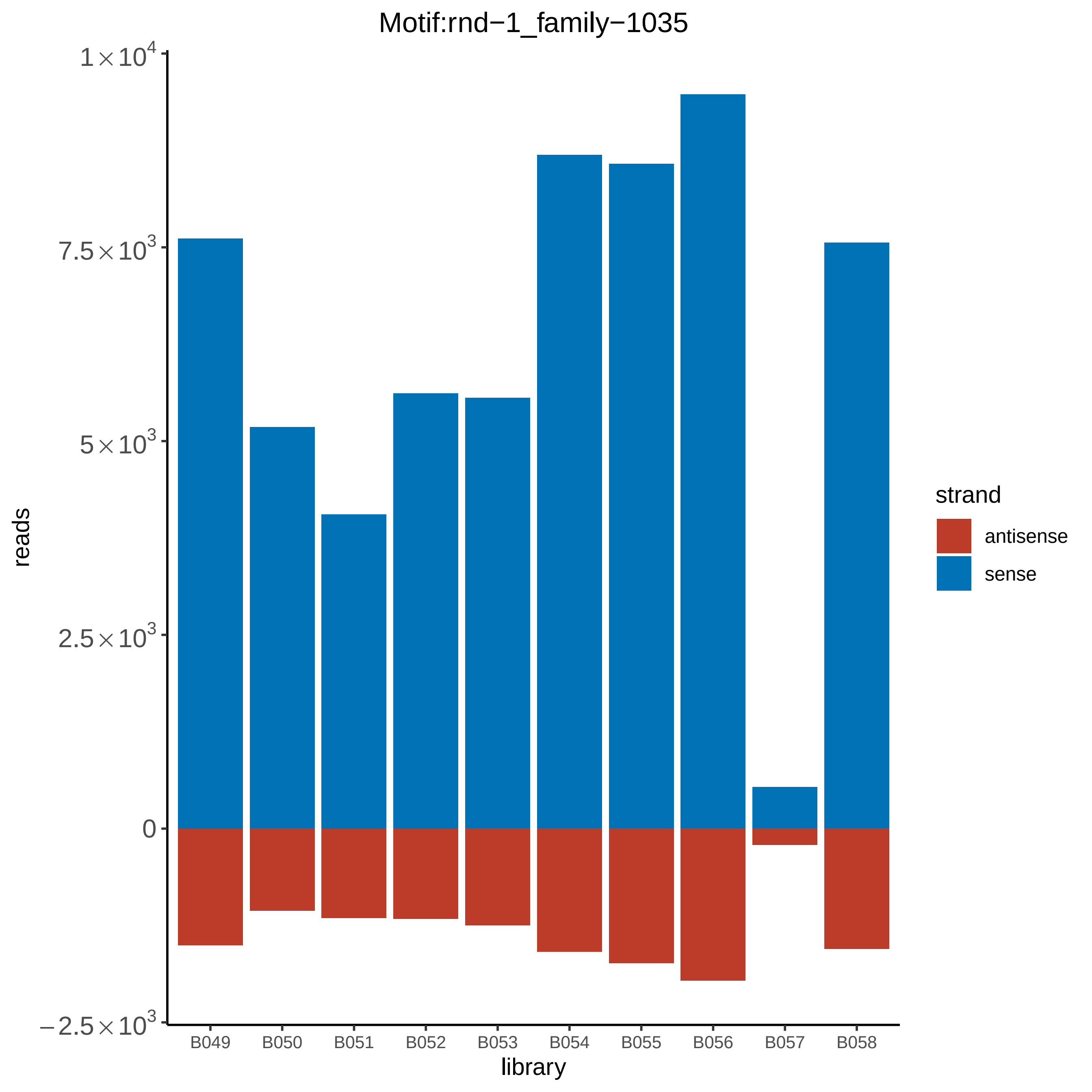

Motif:rnd−1_family−1035
1×10^4
7.5×10^3
5×10^3
2.5×10^3
0
−2.5×10^3
reads
B049
B050
B051
B052
B053
B054
B055
B056
B057
B058
library
strand
antisense
sense

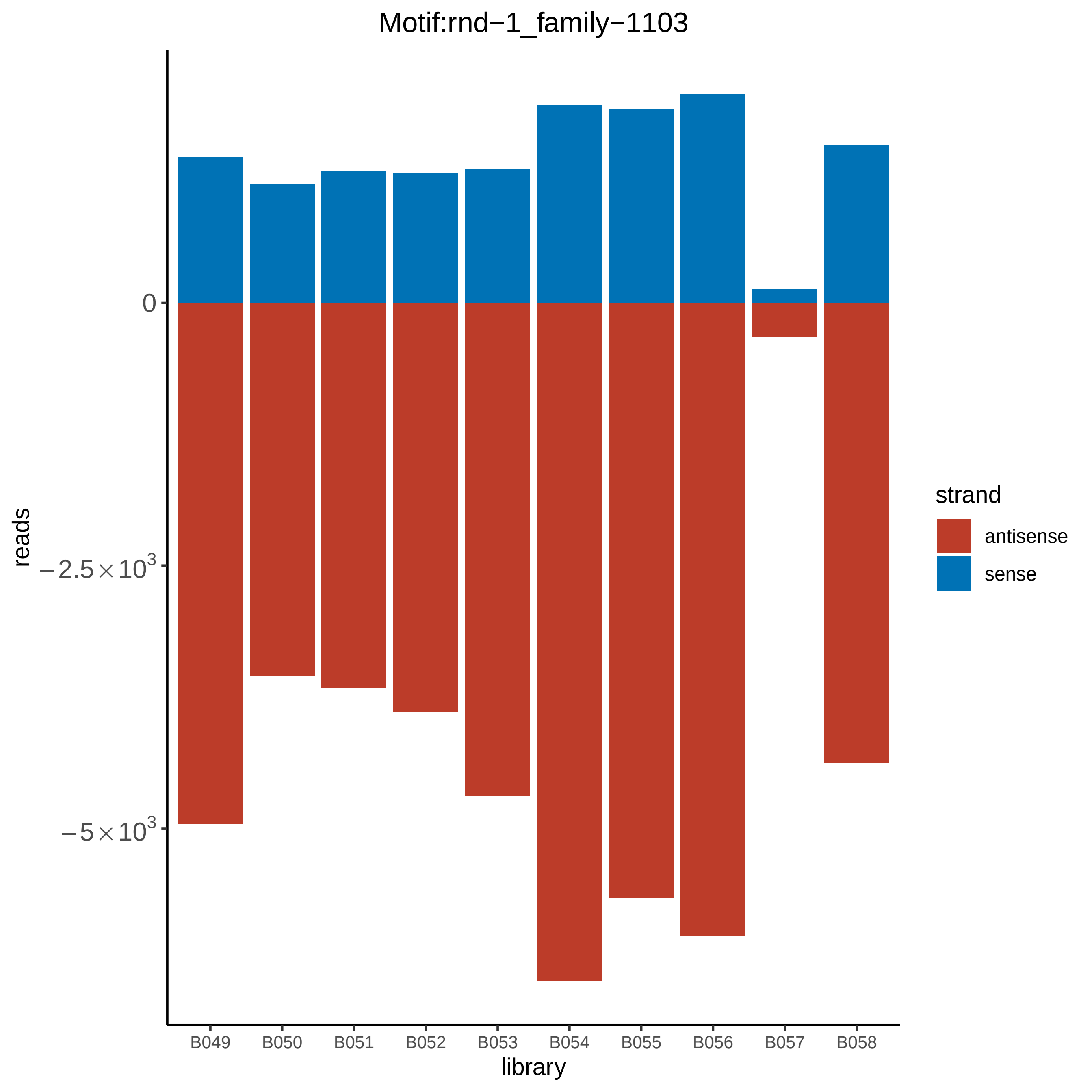

Motif:rnd−1_family−1103
reads
0
−2.5×10^3
−5×10^3
library
B049
B050
B051
B052
B053
B054
B055
B056
B057
B058
strand
antisense
sense

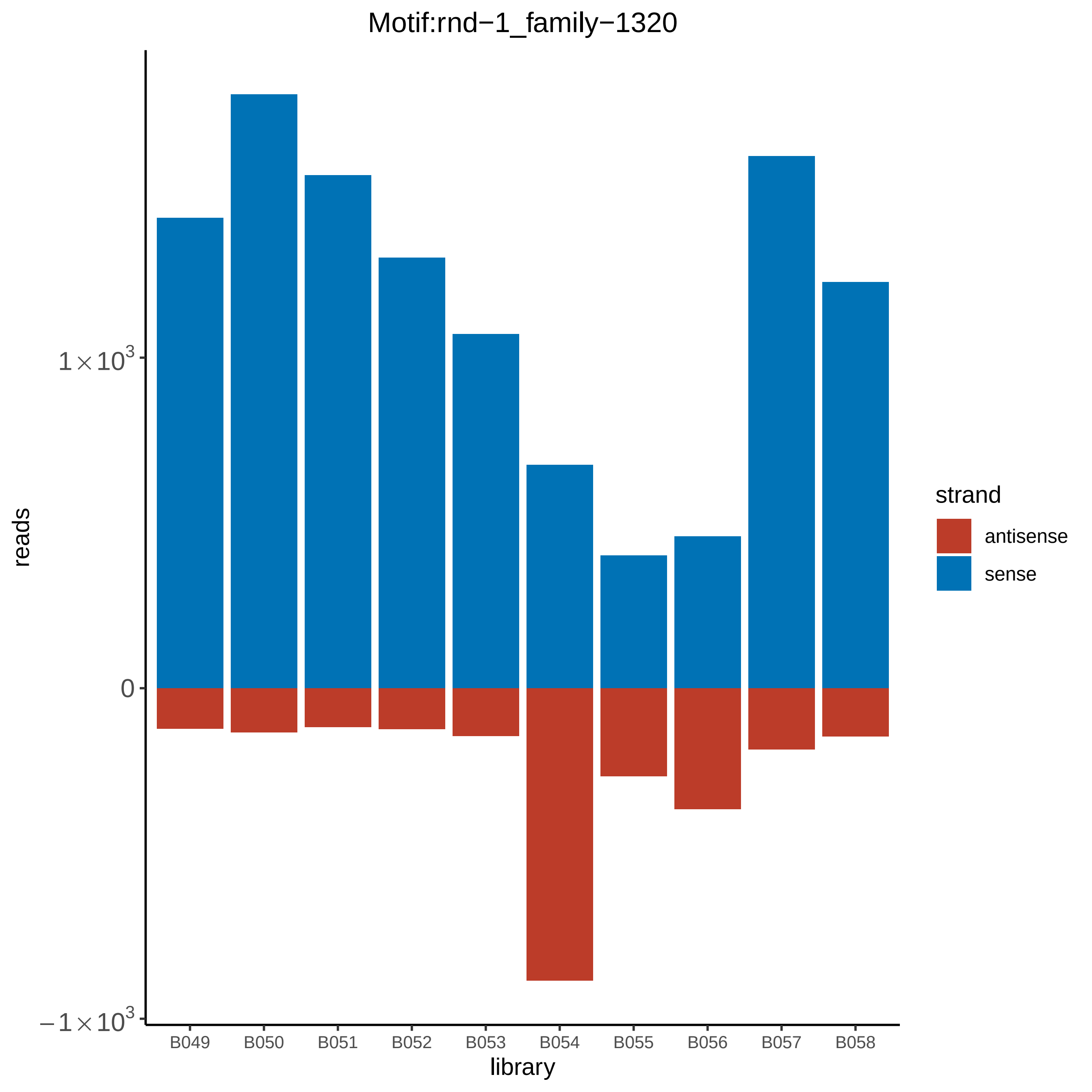
Motif:rnd−1_family−1320
reads
library
1 × 10^3
0
−1 × 10^3
B049
B050
B051
B052
B053
B054
B055
B056
B057
B058
strand
antisense
sense

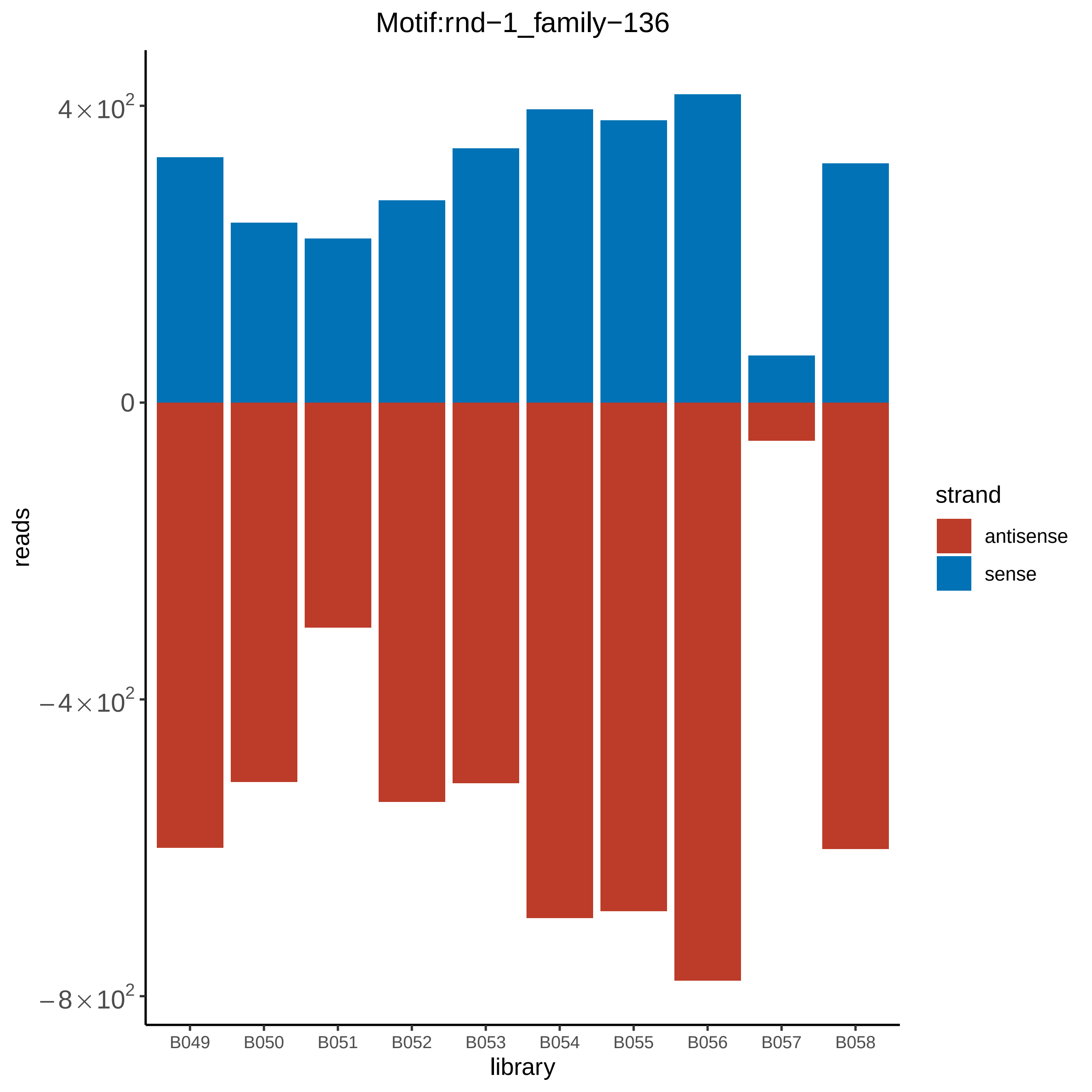
Motif:rnd−1_family−136
4×10^2
0
−4×10^2
−8×10^2
reads
B049
B050
B051
B052
B053
B054
B055
B056
B057
B058
library
strand
antisense
sense

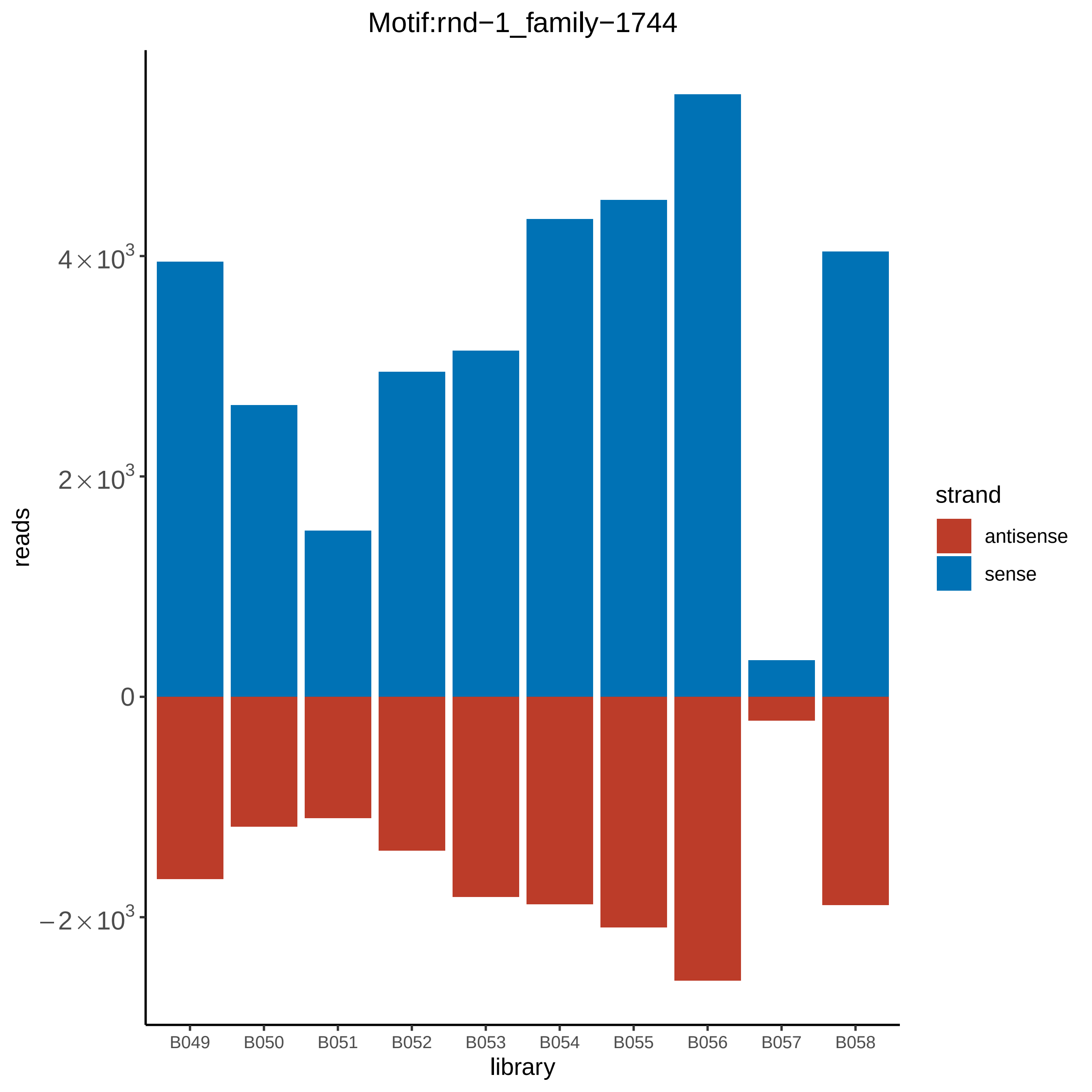

Motif:rnd−1_family−1744
reads
4×10^3
2×10^3
0
−2×10^3
library
B049
B050
B051
B052
B053
B054
B055
B056
B057
B058
strand
antisense
sense

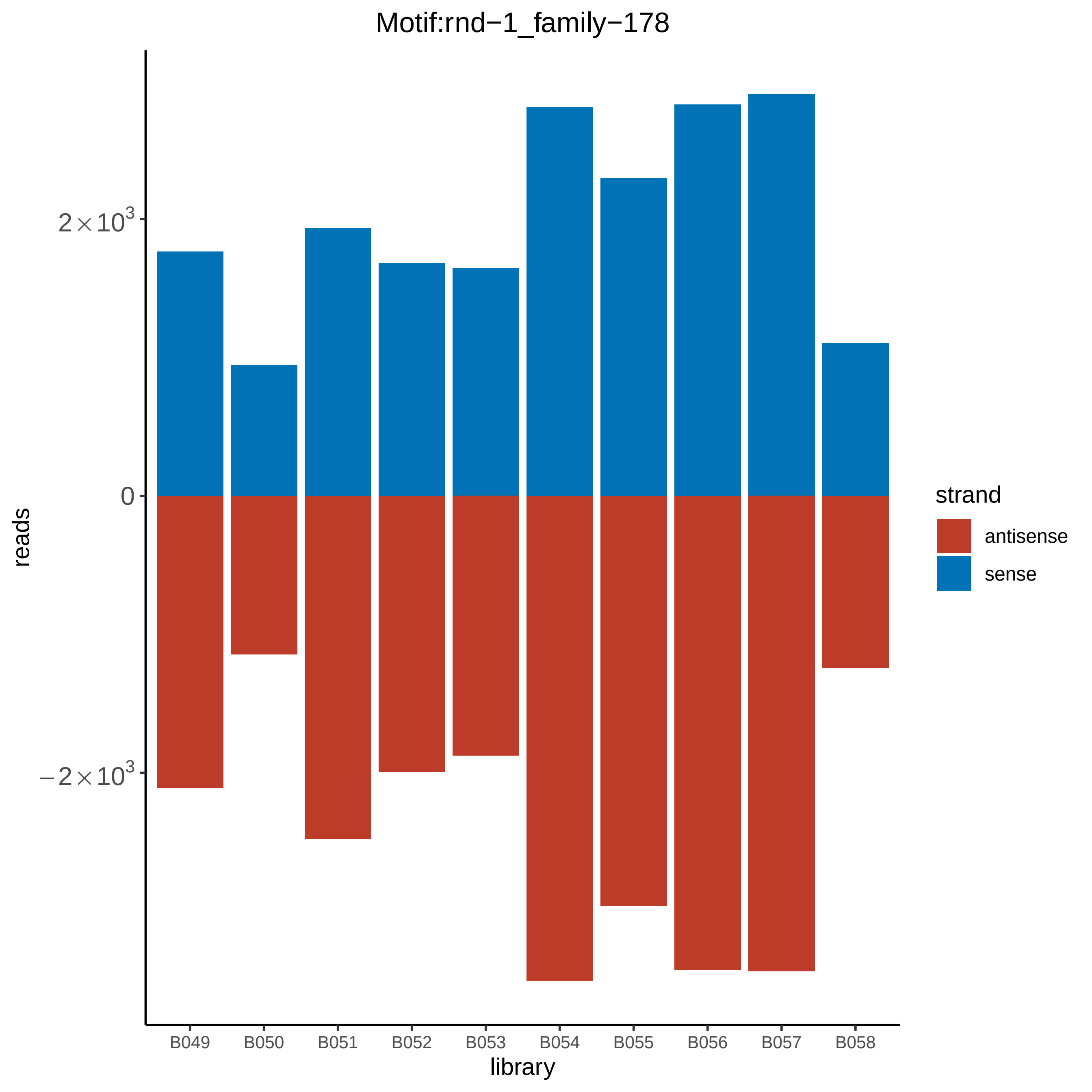

Motif:rnd−1_family−178
reads
$2\times10^3$
0
$-2\times10^3$
library
B049
B050
B051
B052
B053
B054
B055
B056
B057
B058
strand
antisense
sense

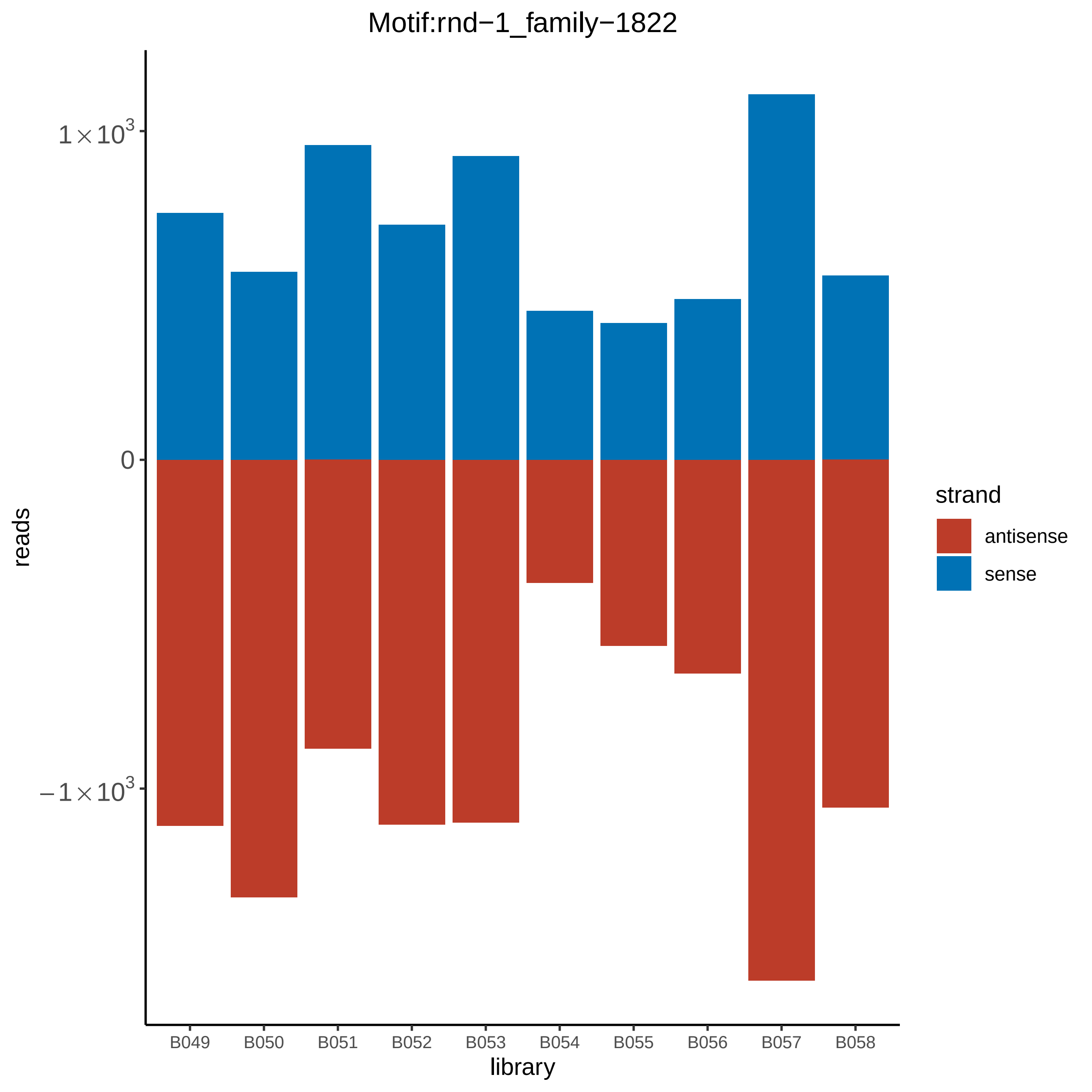

Motif:rnd−1_family−1822
$1\times10^3$
0
$-1\times10^3$
reads
strand
antisense
sense
B049
B050
B051
B052
B053
B054
B055
B056
B057
B058
library

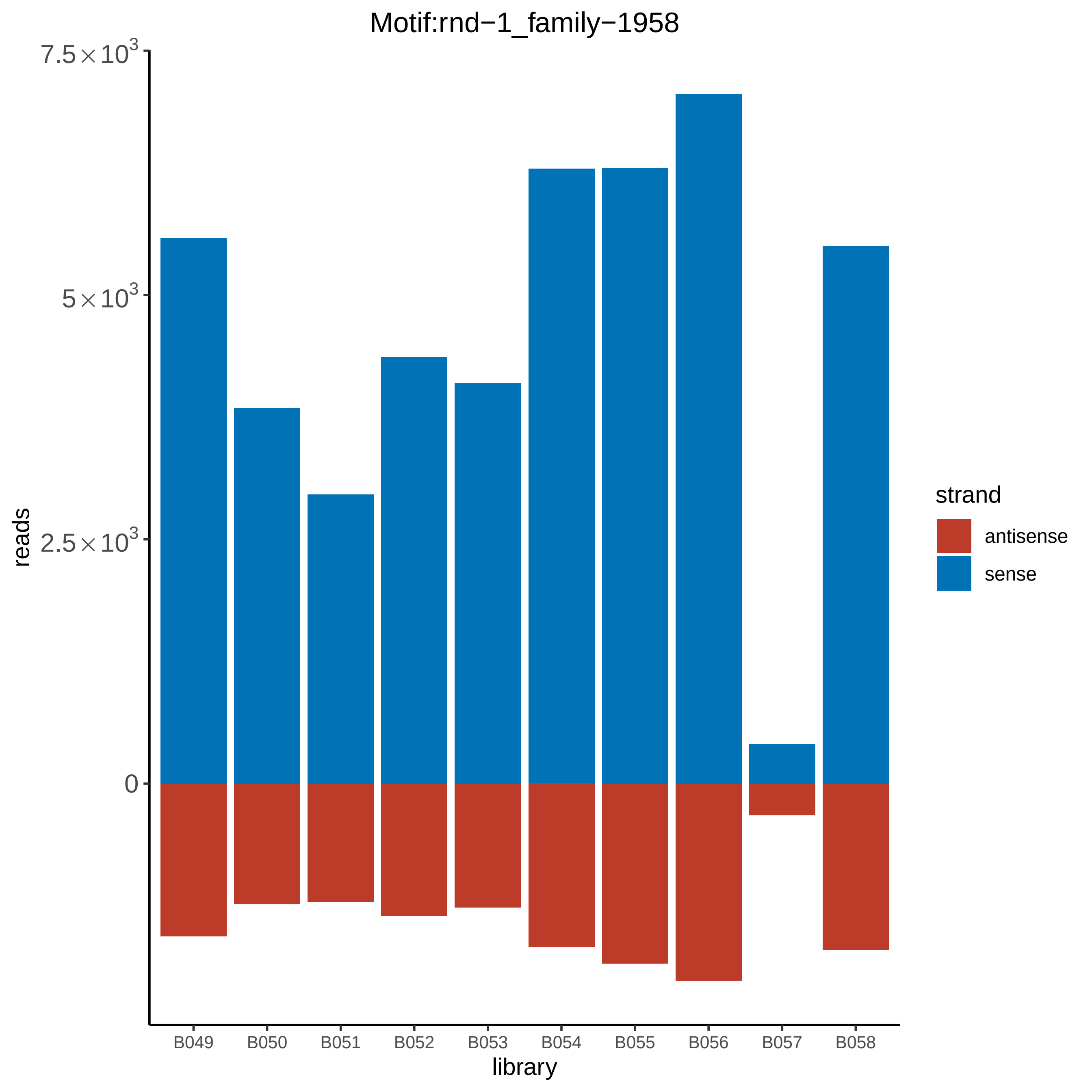
Motif:rnd−1_family−1958
reads
$7.5\times10^3$
$5\times10^3$
$2.5\times10^3$
0
B049
B050
B051
B052
B053
B054
B055
B056
B057
B058
library
strand
antisense
sense

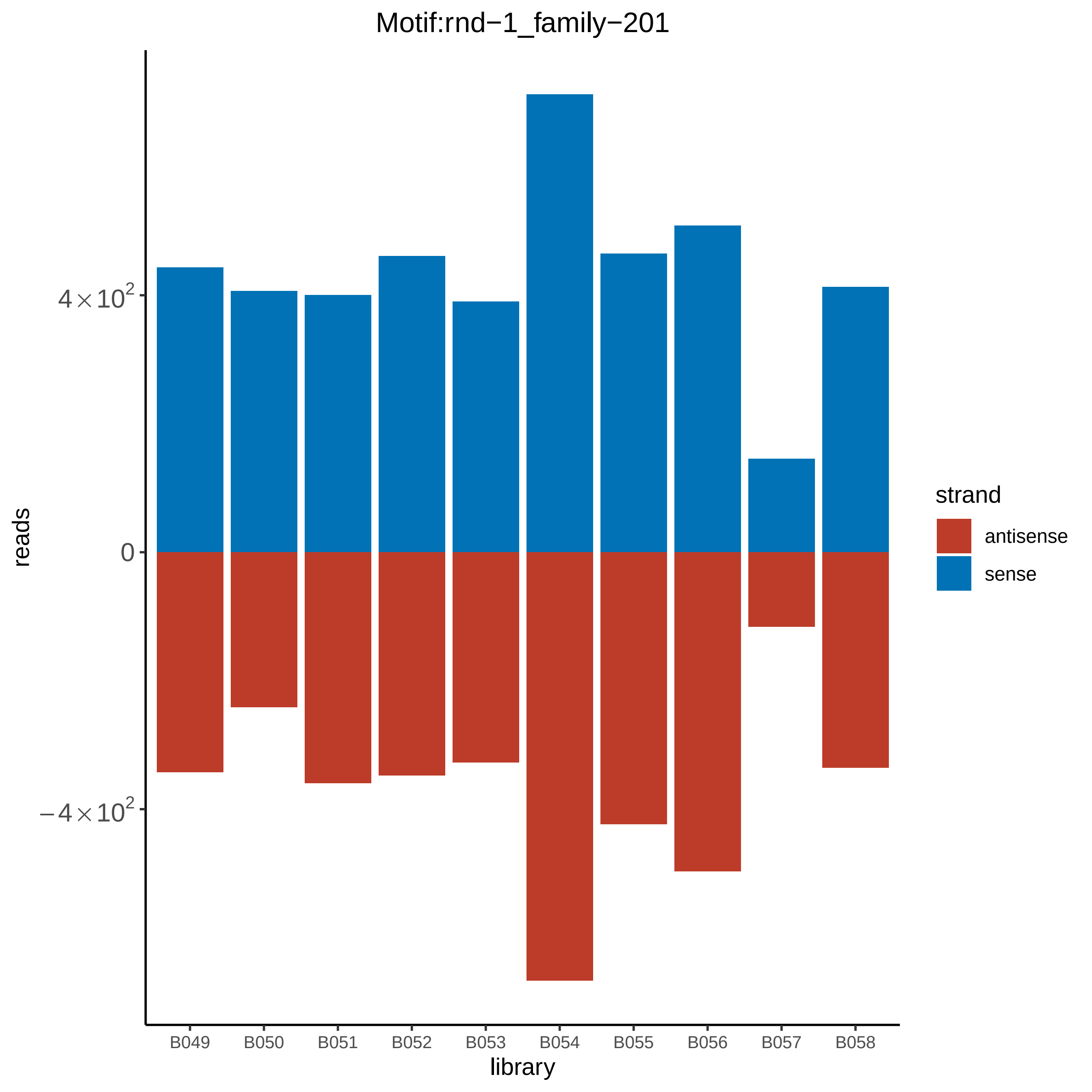

Motif:rnd−1_family−201
reads
4×10^2
0
−4×10^2
library
B049
B050
B051
B052
B053
B054
B055
B056
B057
B058
strand
antisense
sense

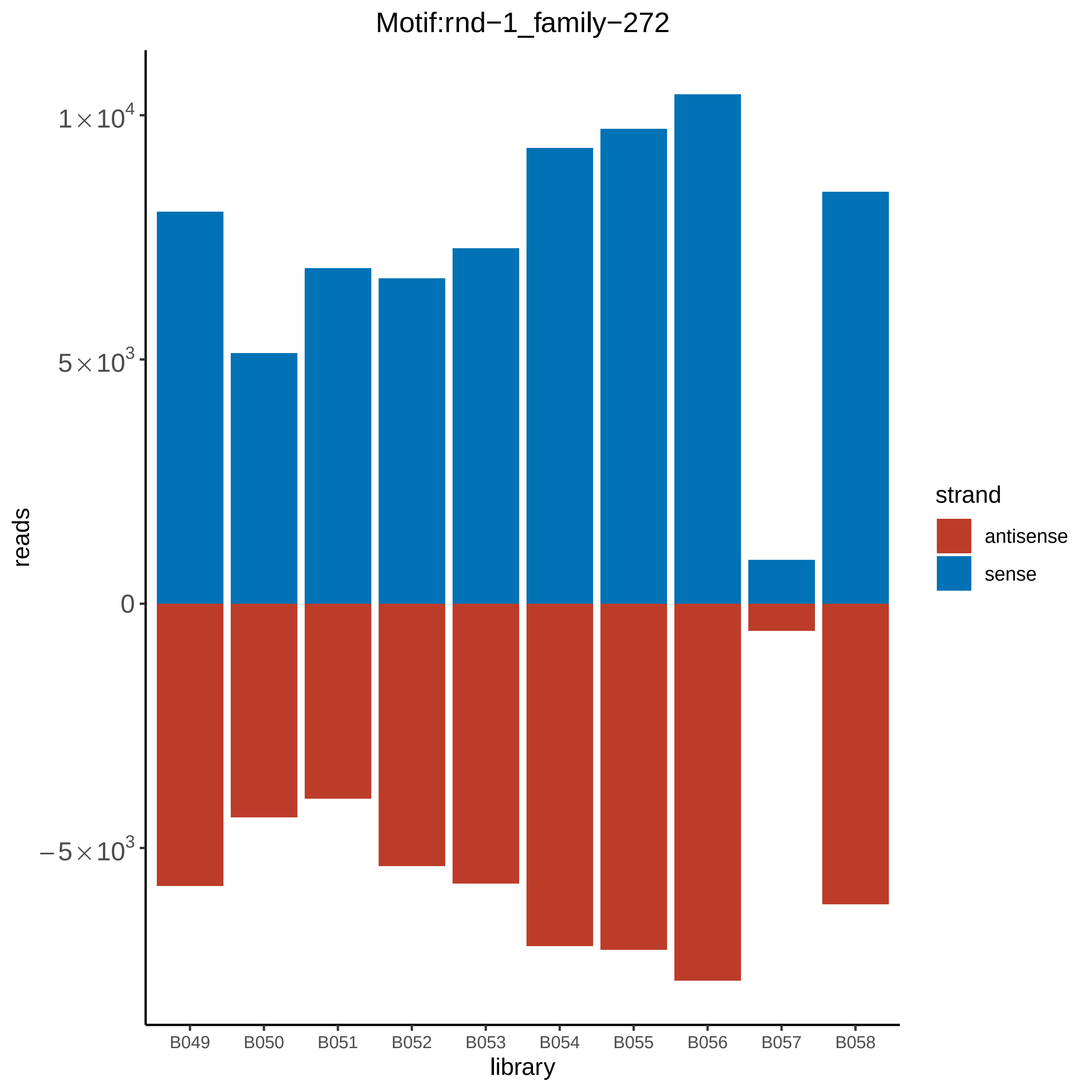
Motif:rnd−1_family−272
$1\times10^4$
$5\times10^3$
0
$-5\times10^3$
reads
B049
B050
B051
B052
B053
B054
B055
B056
B057
B058
library
strand
antisense
sense

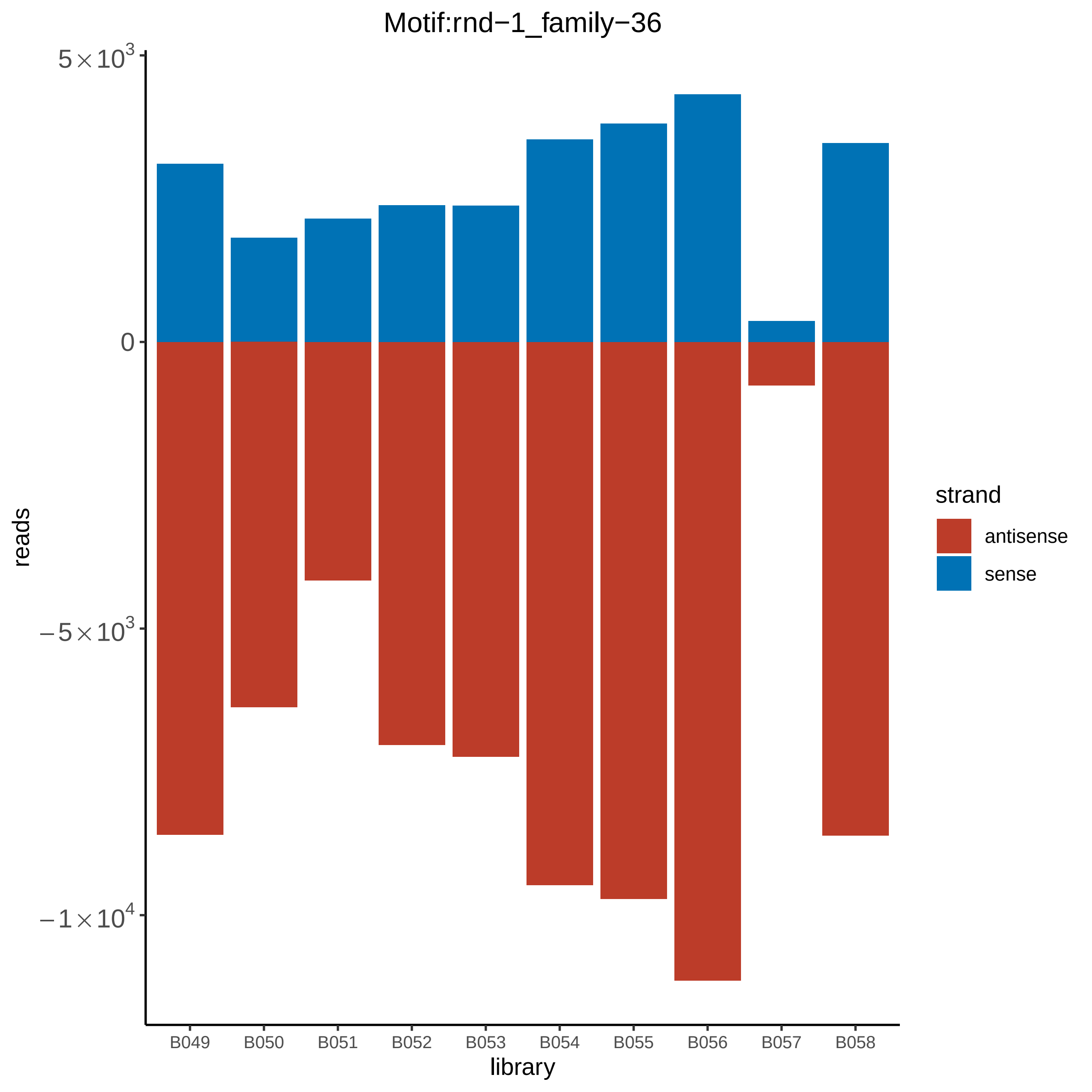

Motif:rnd−1_family−36
5×10^3
0
−5×10^3
−1×10^4
reads
B049
B050
B051
B052
B053
B054
B055
B056
B057
B058
library
strand
antisense
sense

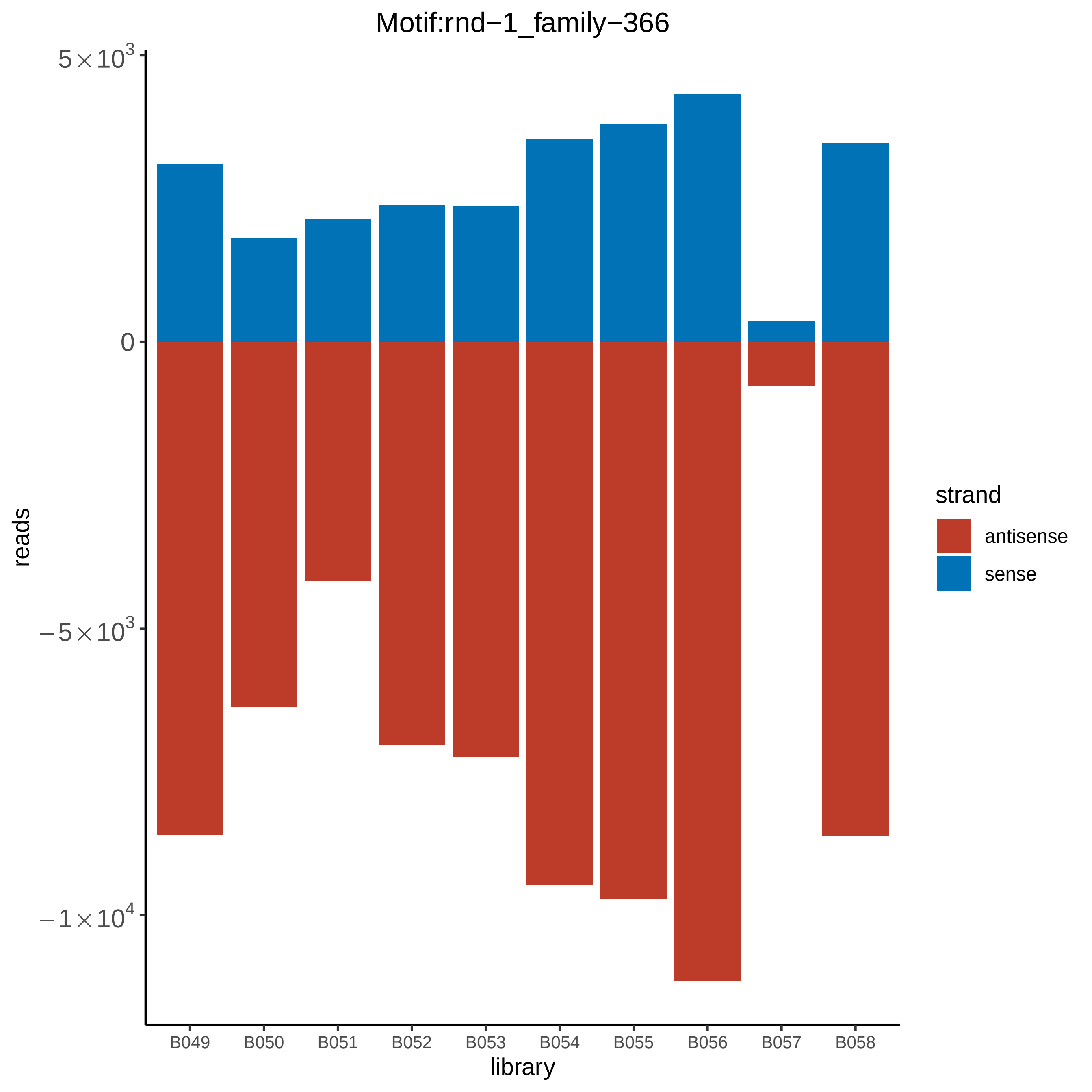

Motif:rnd−1_family−366
5×10^3
0
−5×10^3
−1×10^4
reads
B049
B050
B051
B052
B053
B054
B055
B056
B057
B058
library
strand
antisense
sense

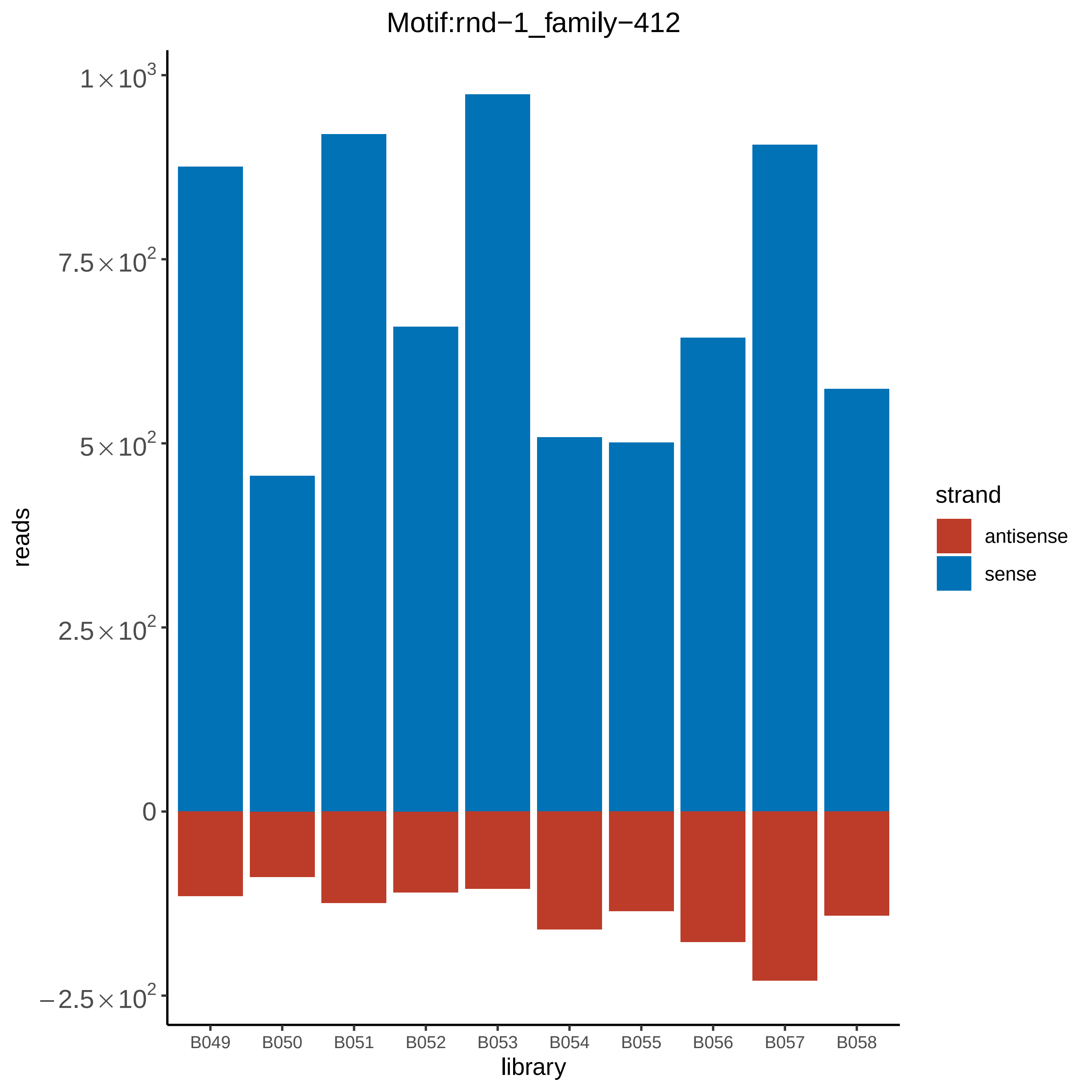

Motif:rnd−1_family−412
reads
$1\times10^3$
$7.5\times10^2$
$5\times10^2$
$2.5\times10^2$
0
$-2.5\times10^2$
B049
B050
B051
B052
B053
B054
B055
B056
B057
B058
library
strand
antisense
sense

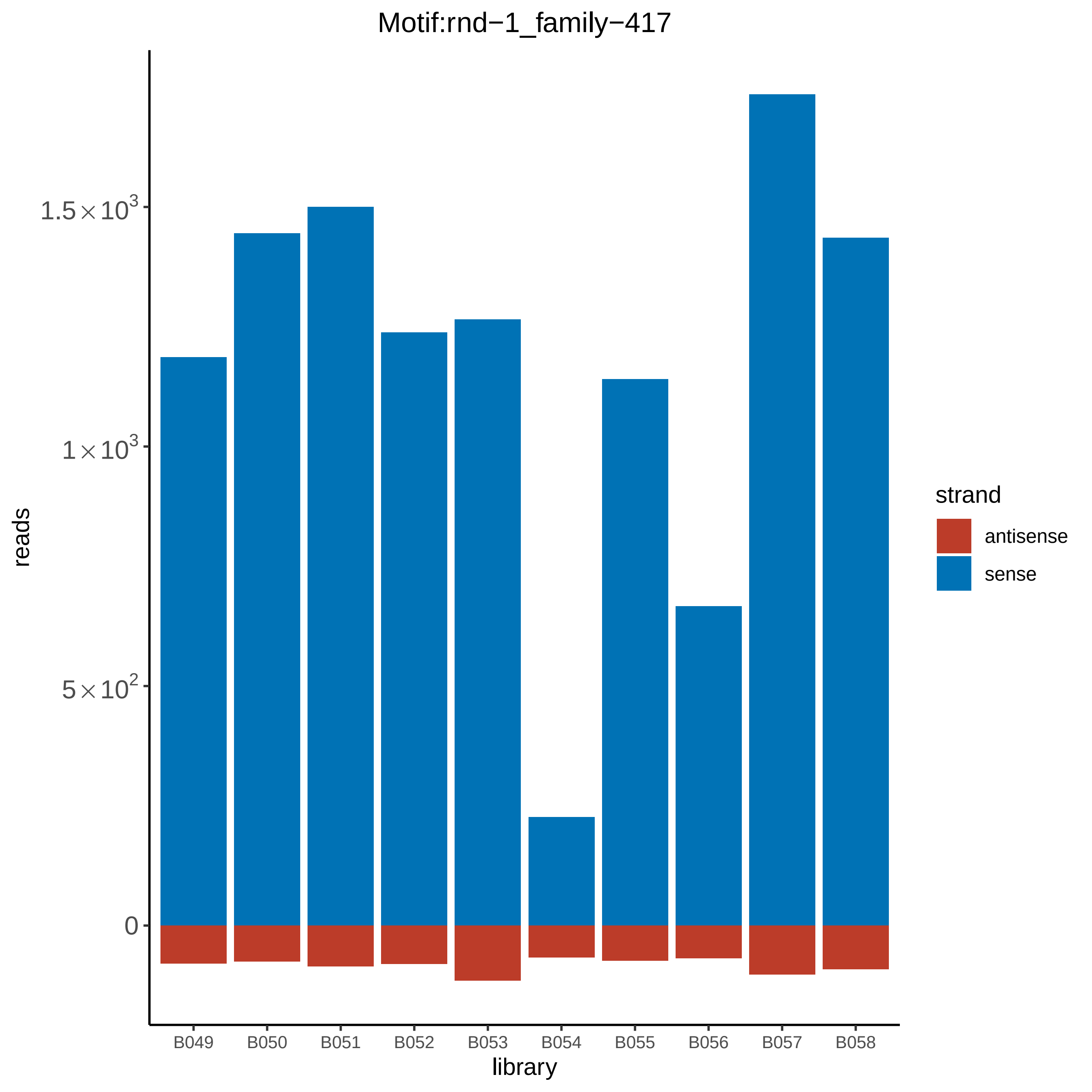

Motif:rnd−1_family−417
reads
1.5×10^3
1×10^3
5×10^2
0
B049
B050
B051
B052
B053
B054
B055
B056
B057
B058
library
strand
antisense
sense

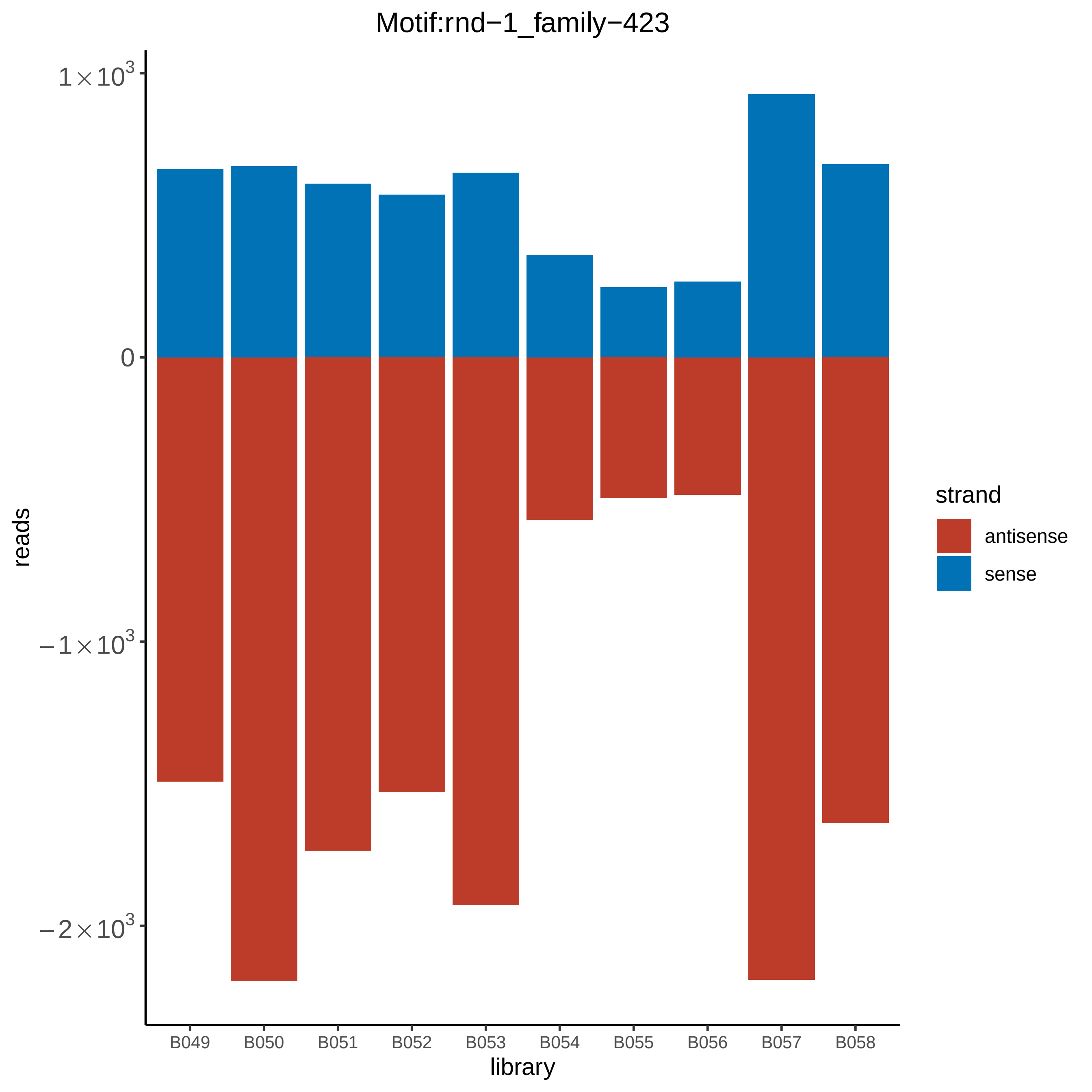

Motif:rnd−1_family−423
reads
1×10^3
0
−1×10^3
−2×10^3
library
B049
B050
B051
B052
B053
B054
B055
B056
B057
B058
strand
antisense
sense

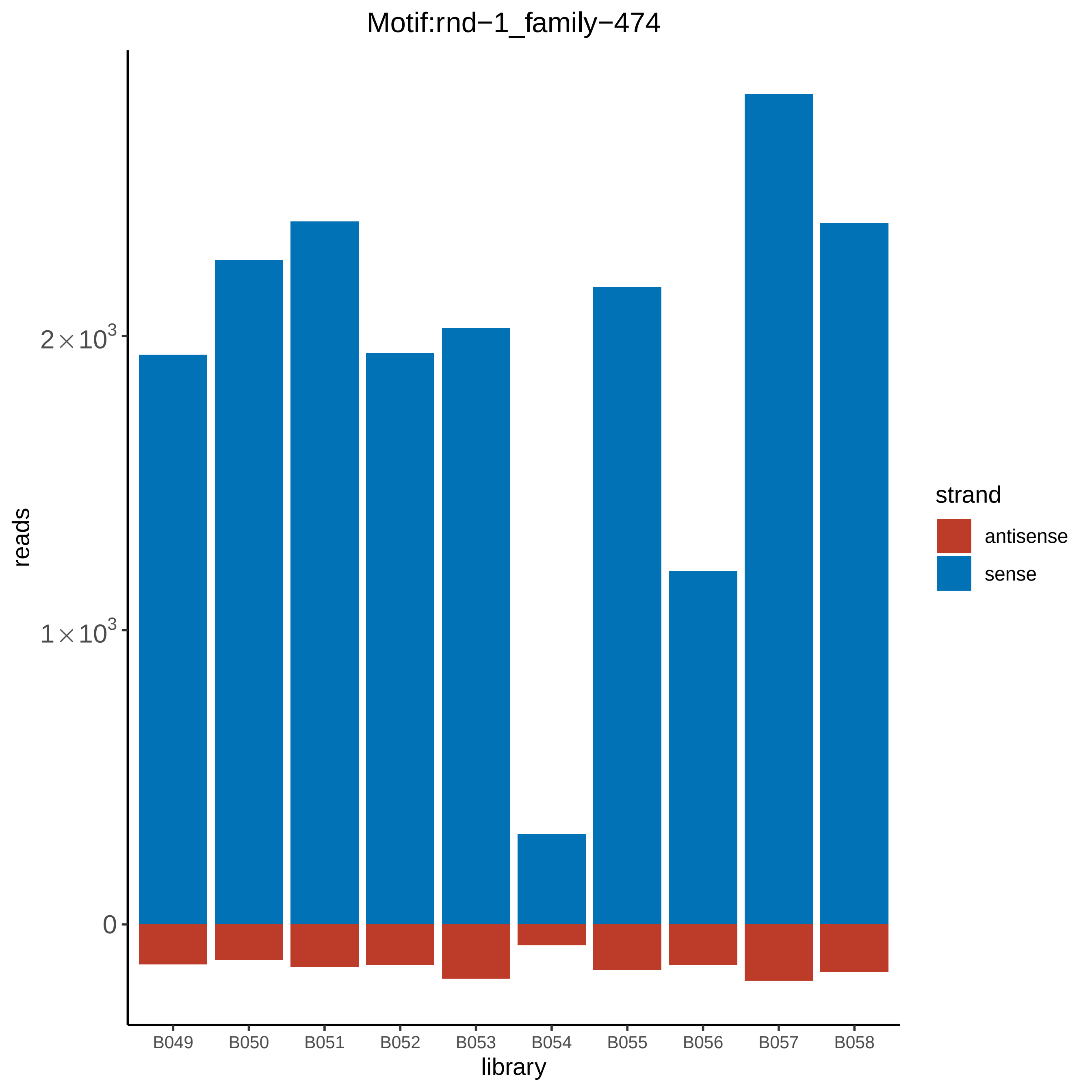
Motif:rnd−1_family−474
reads
2×10^3
1×10^3
0
library
B049
B050
B051
B052
B053
B054
B055
B056
B057
B058
strand
antisense
sense

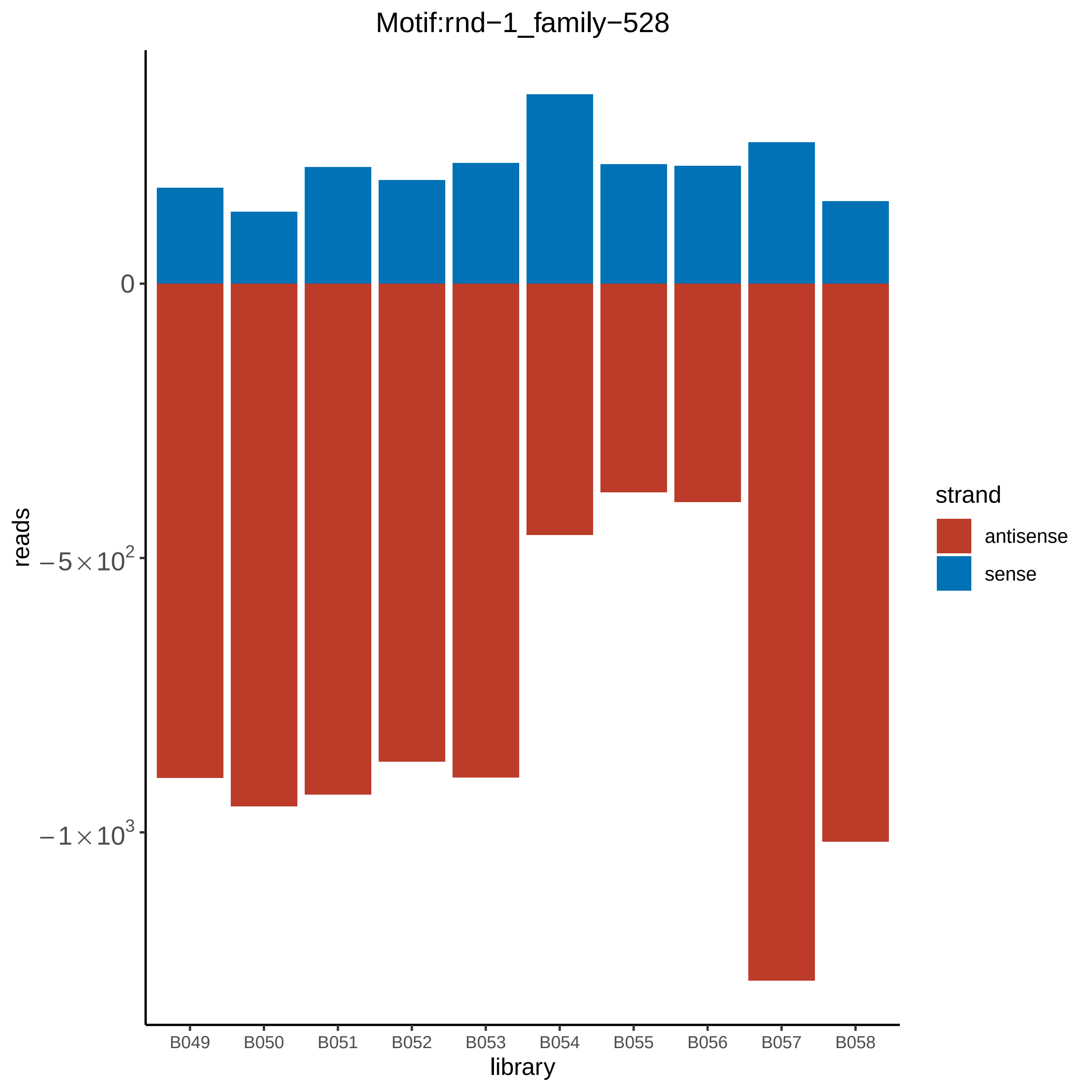
Motif:rnd−1_family−528
reads
0
$-5\times10^2$
$-1\times10^3$
library
B049
B050
B051
B052
B053
B054
B055
B056
B057
B058
strand
antisense
sense

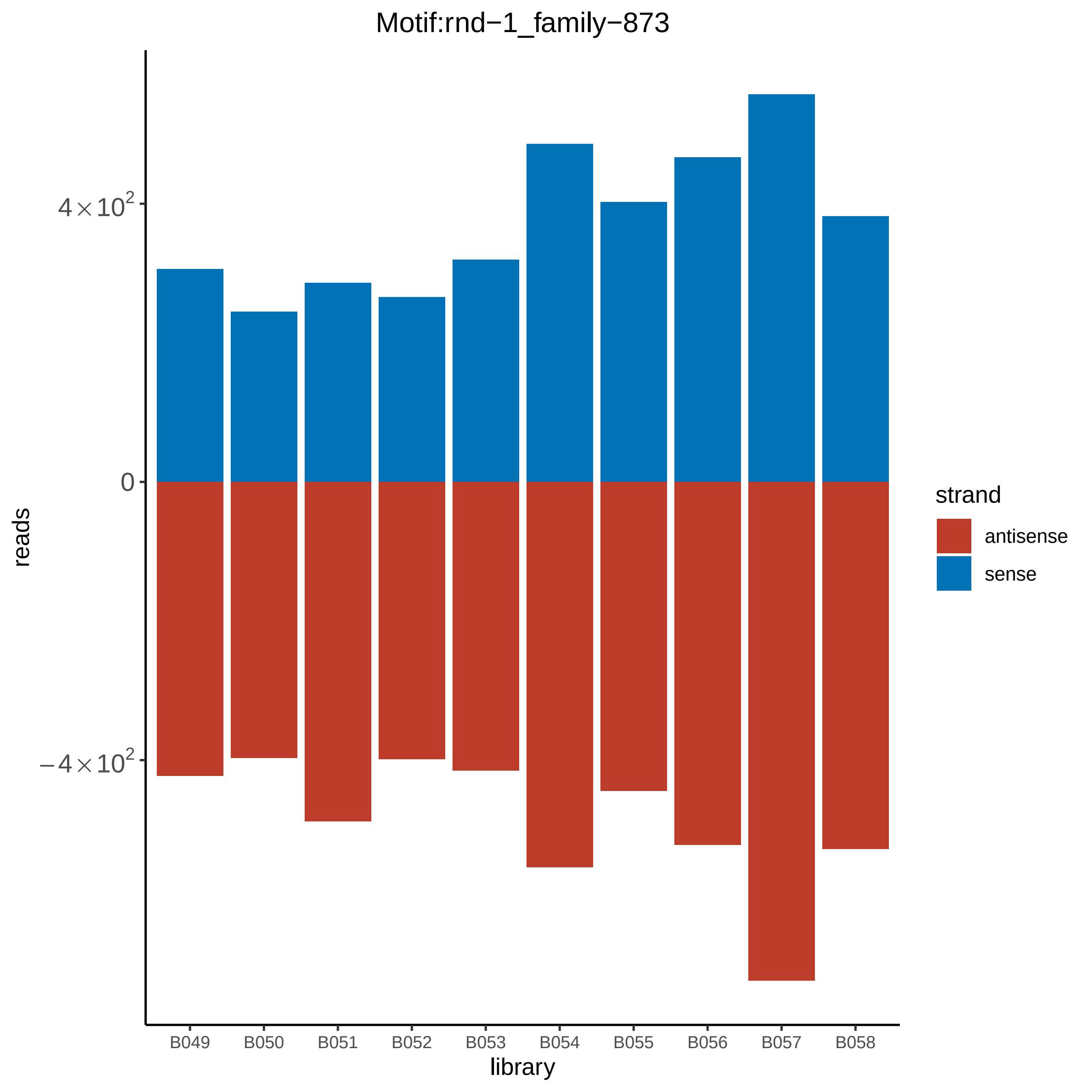

Motif:rnd−1_family−873
reads
4×10^2
0
−4×10^2
library
B049
B050
B051
B052
B053
B054
B055
B056
B057
B058
strand
antisense
sense

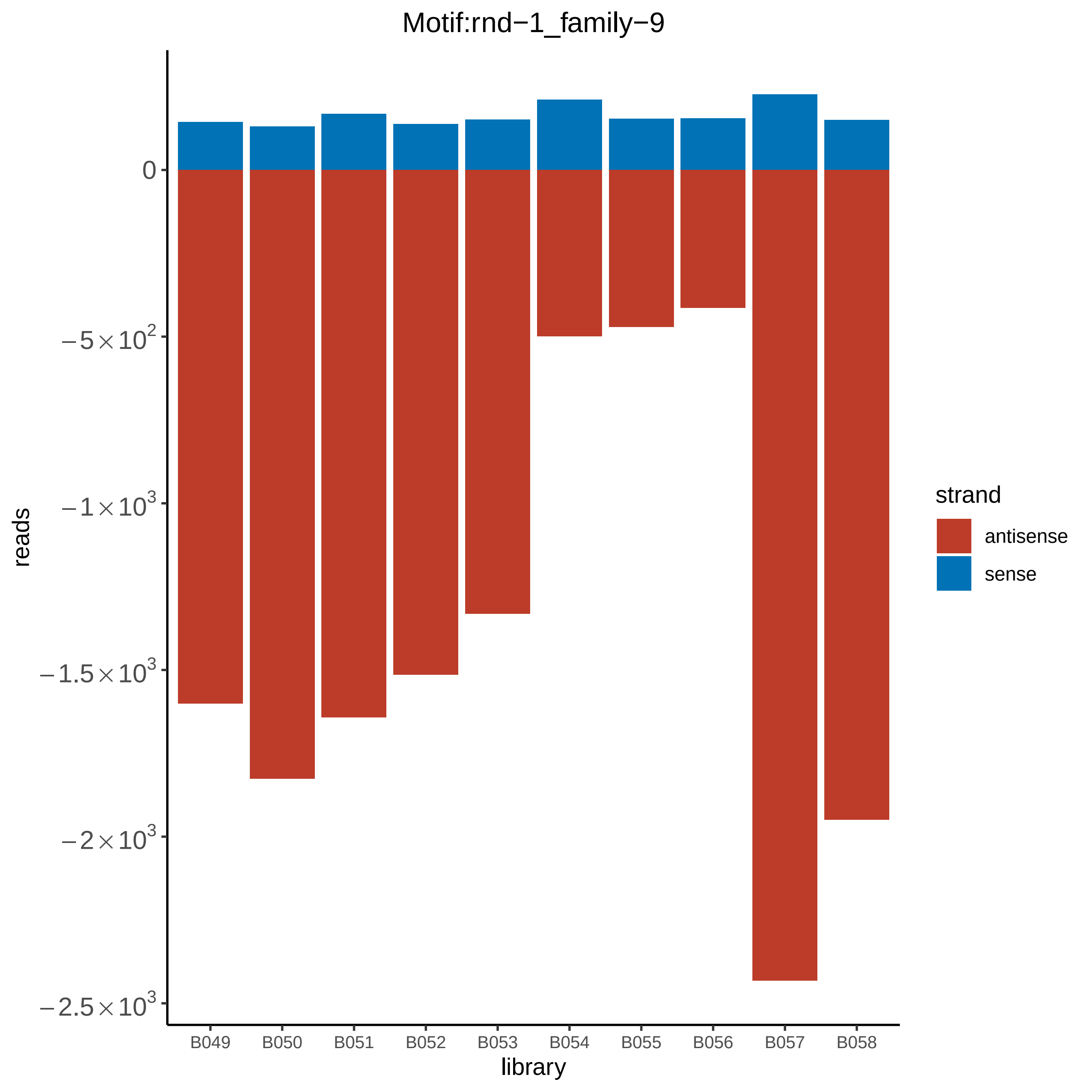

Motif:rnd−1_family−9
0
$-5\times10^2$
$-1\times10^3$
$-1.5\times10^3$
$-2\times10^3$
$-2.5\times10^3$
reads
B049
B050
B051
B052
B053
B054
B055
B056
B057
B058
library
strand
antisense
sense

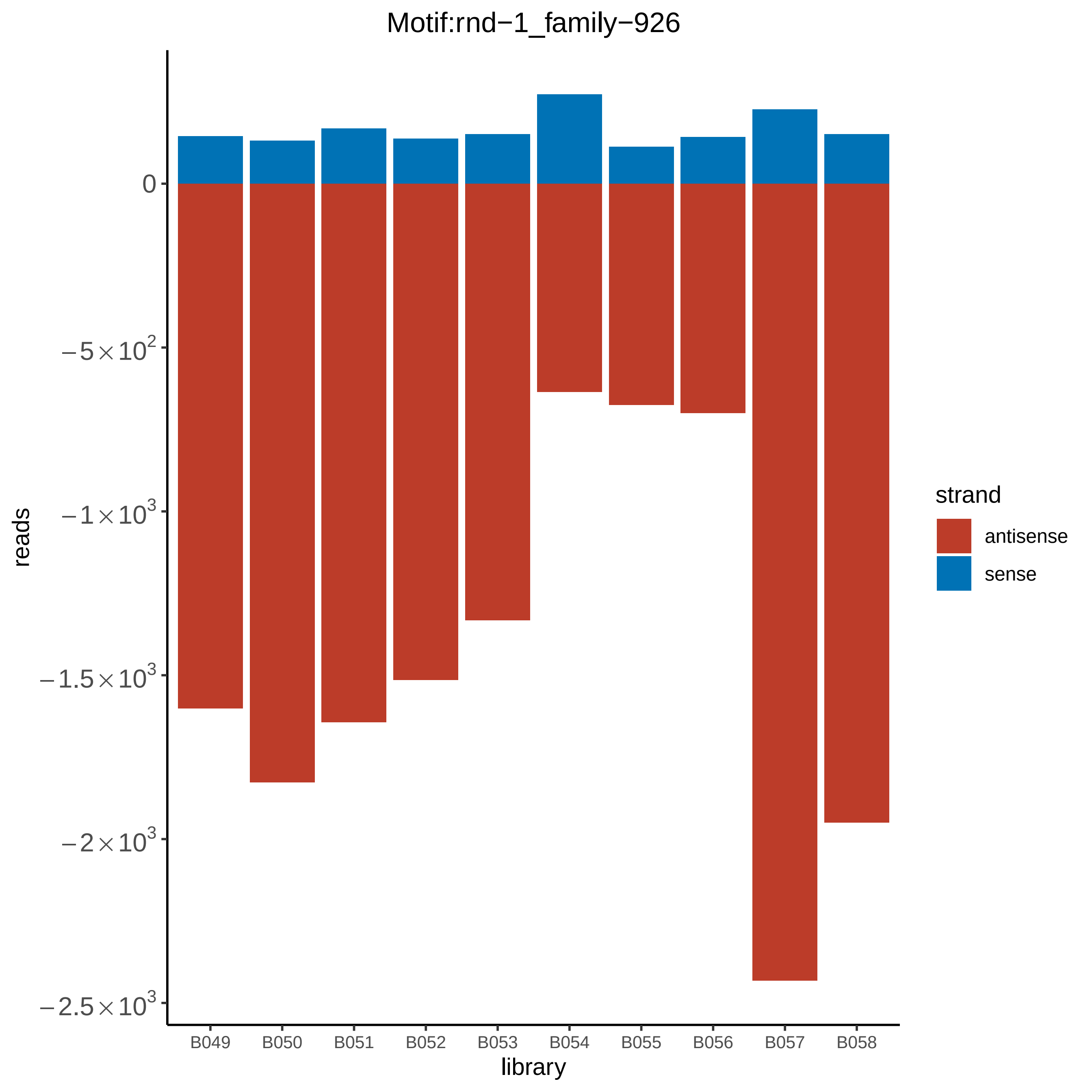

Motif:rnd−1_family−926
reads
0
−5×10^2
−1×10^3
−1.5×10^3
−2×10^3
−2.5×10^3
B049
B050
B051
B052
B053
B054
B055
B056
B057
B058
library
strand
antisense
sense

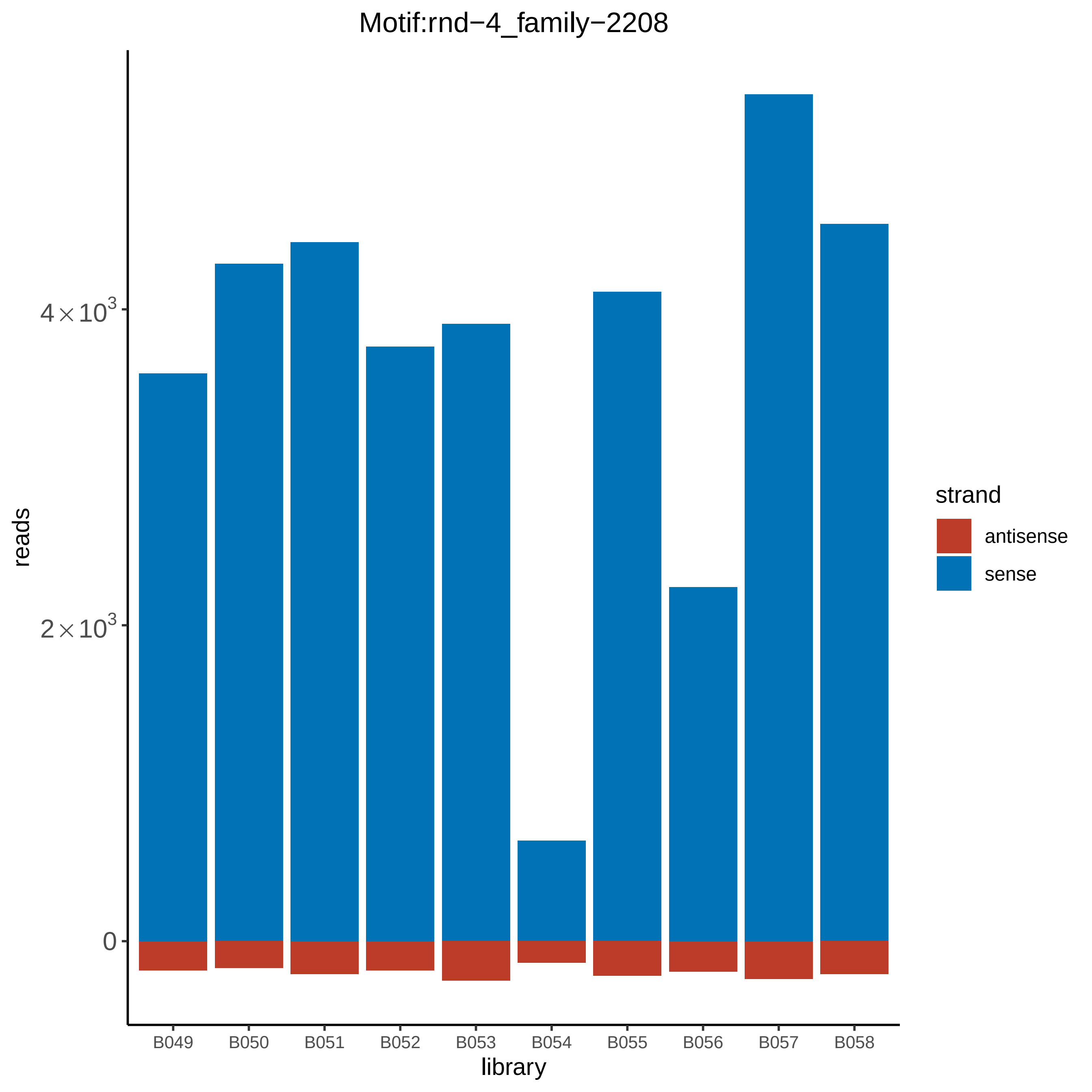

Motif:rnd−4_family−2208
reads
$4\times10^3$
$2\times10^3$
0
library
B049
B050
B051
B052
B053
B054
B055
B056
B057
B058
strand
antisense
sense

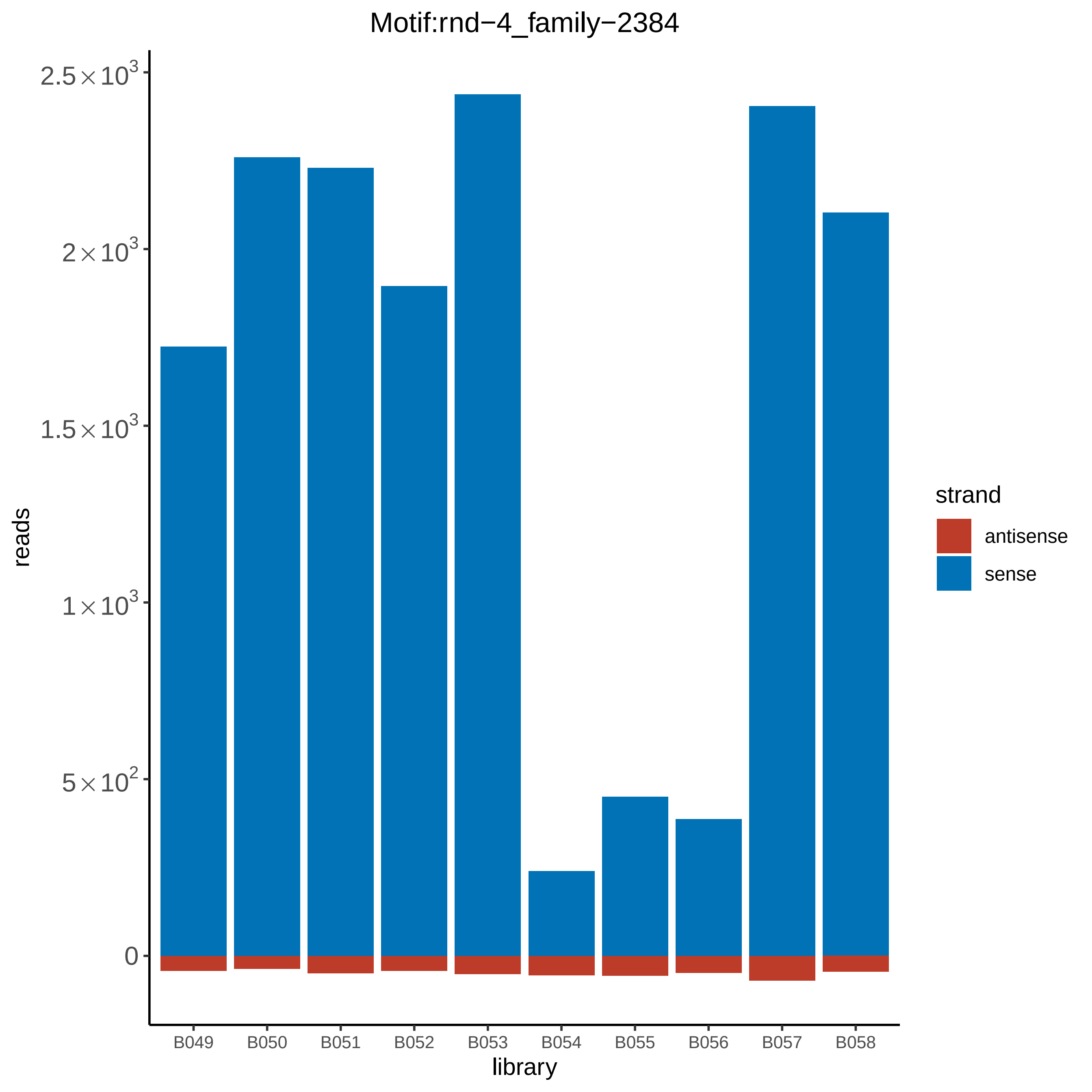
Motif:rnd−4_family−2384
reads
$2.5\times10^3$
$2\times10^3$
$1.5\times10^3$
$1\times10^3$
$5\times10^2$
0
B049
B050
B051
B052
B053
B054
B055
B056
B057
B058
library
strand
antisense
sense

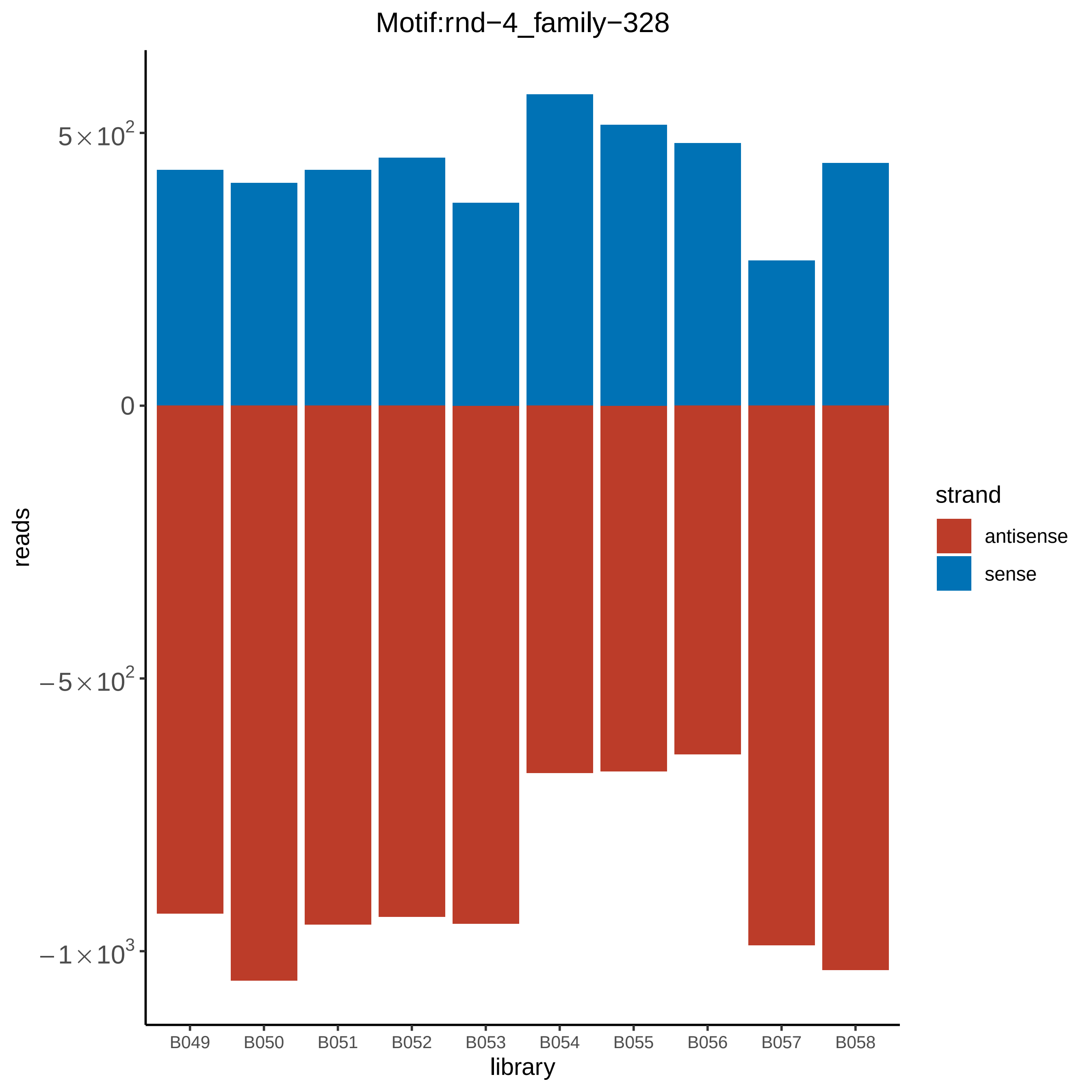

Motif:rnd−4_family−328
5×10^2
0
−5×10^2
−1×10^3
reads
B049
B050
B051
B052
B053
B054
B055
B056
B057
B058
library
strand
antisense
sense

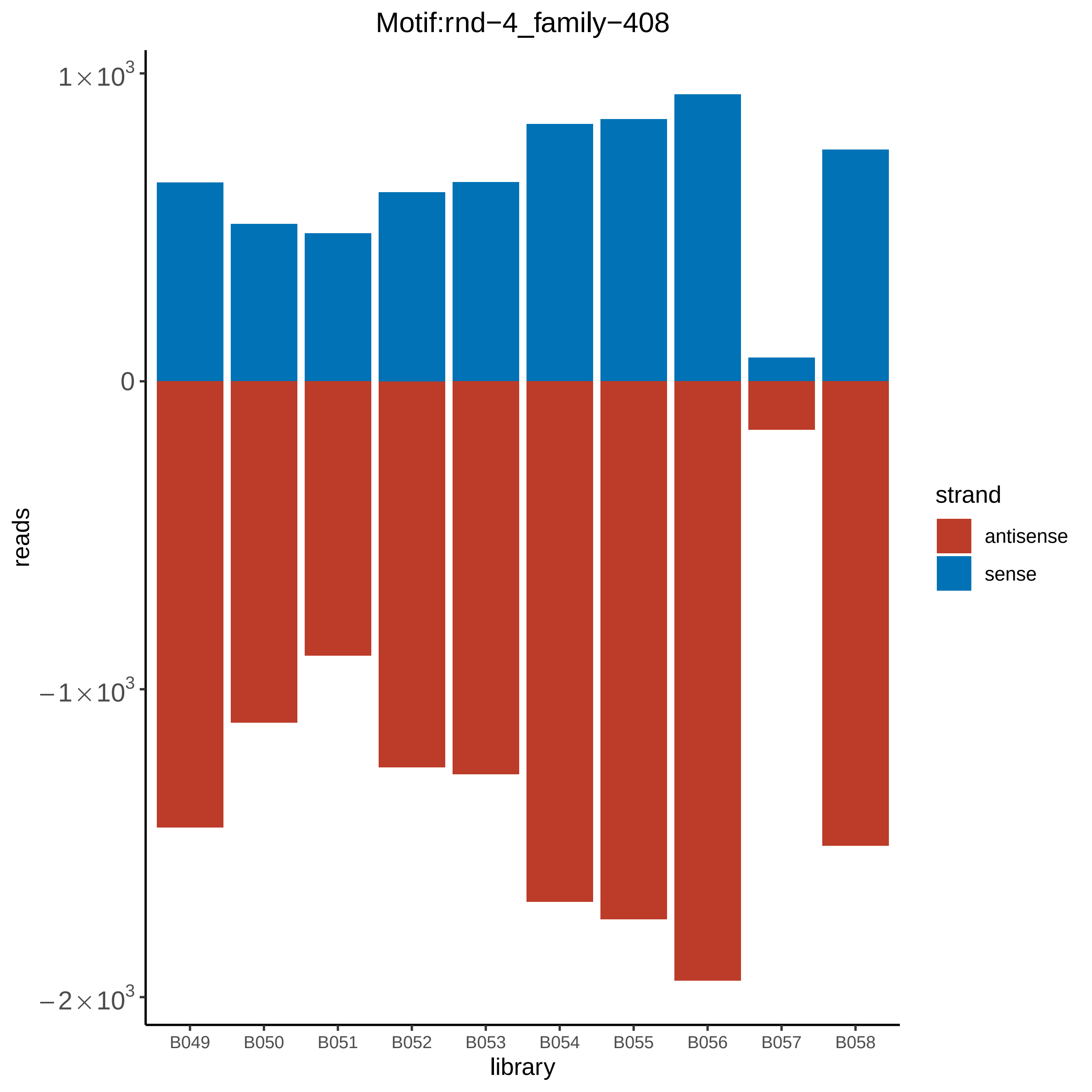

Motif:rnd−4_family−408
$1\times10^3$
0
$-1\times10^3$
$-2\times10^3$
reads
B049
B050
B051
B052
B053
B054
B055
B056
B057
B058
library
strand
antisense
sense

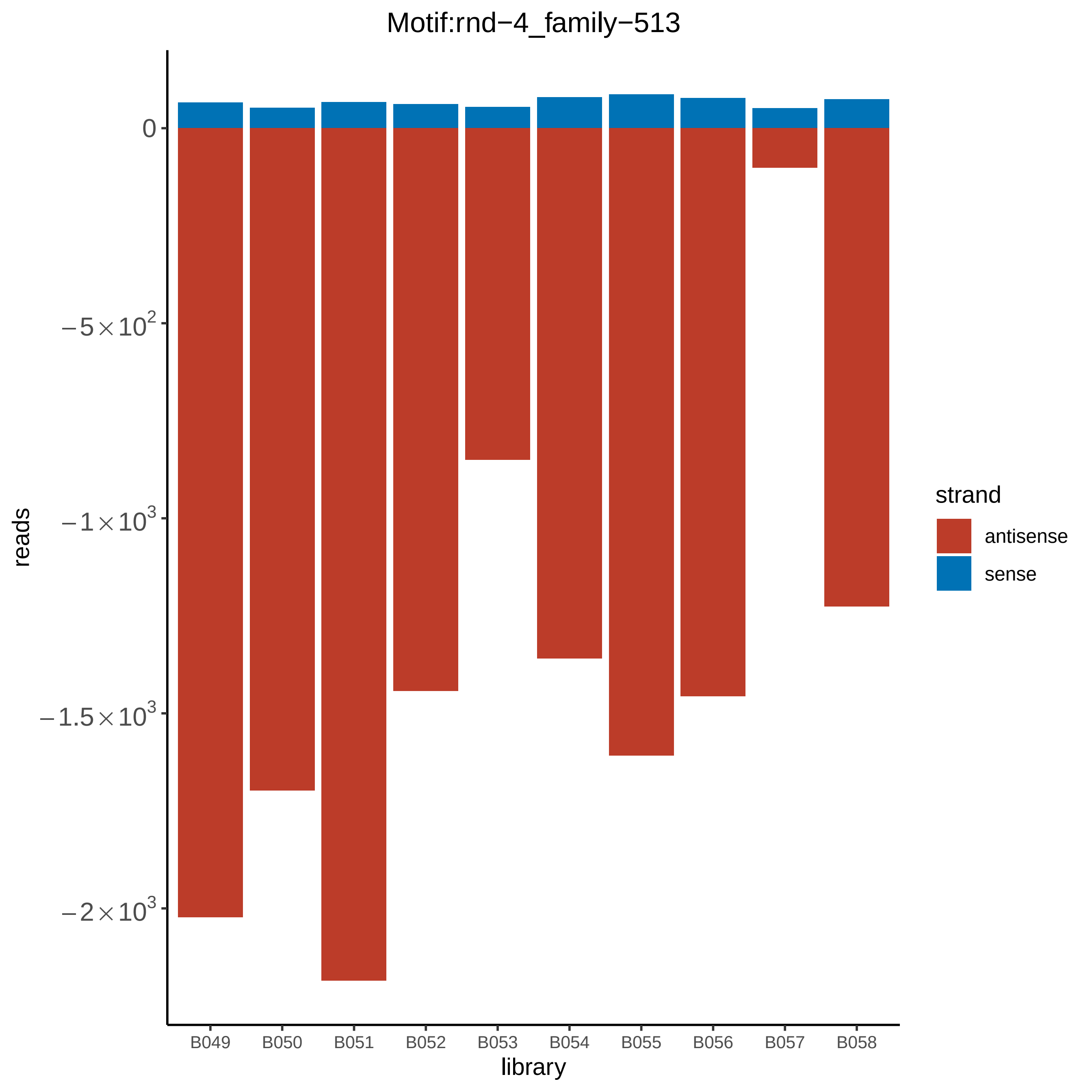

Motif:rnd−4_family−513
reads
0
−5×10^2
−1×10^3
−1.5×10^3
−2×10^3
B049
B050
B051
B052
B053
B054
B055
B056
B057
B058
library
strand
antisense
sense

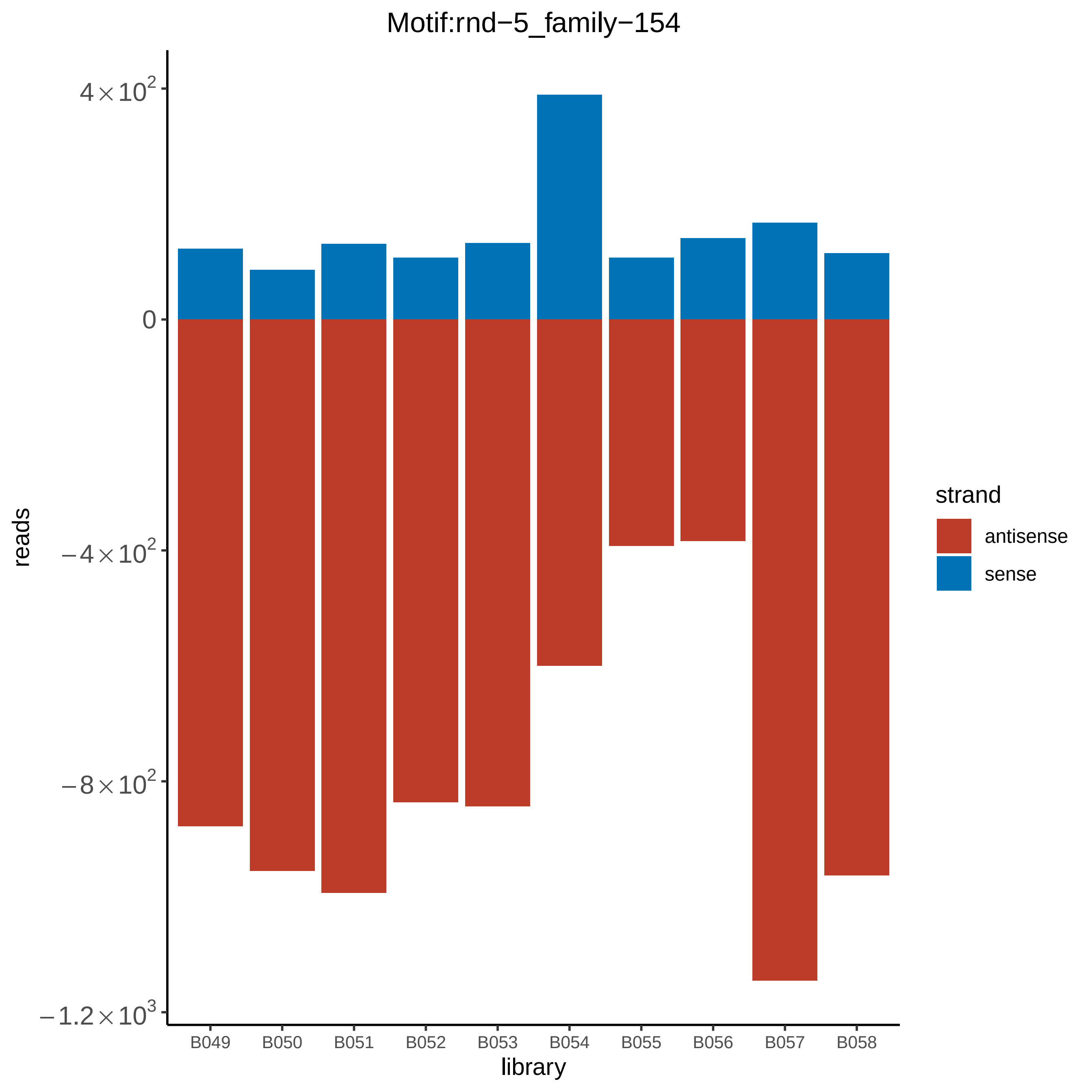

Motif:rnd−5_family−154
reads
$4\times10^2$
0
$-4\times10^2$
$-8\times10^2$
$-1.2\times10^3$
library
B049
B050
B051
B052
B053
B054
B055
B056
B057
B058
strand
antisense
sense

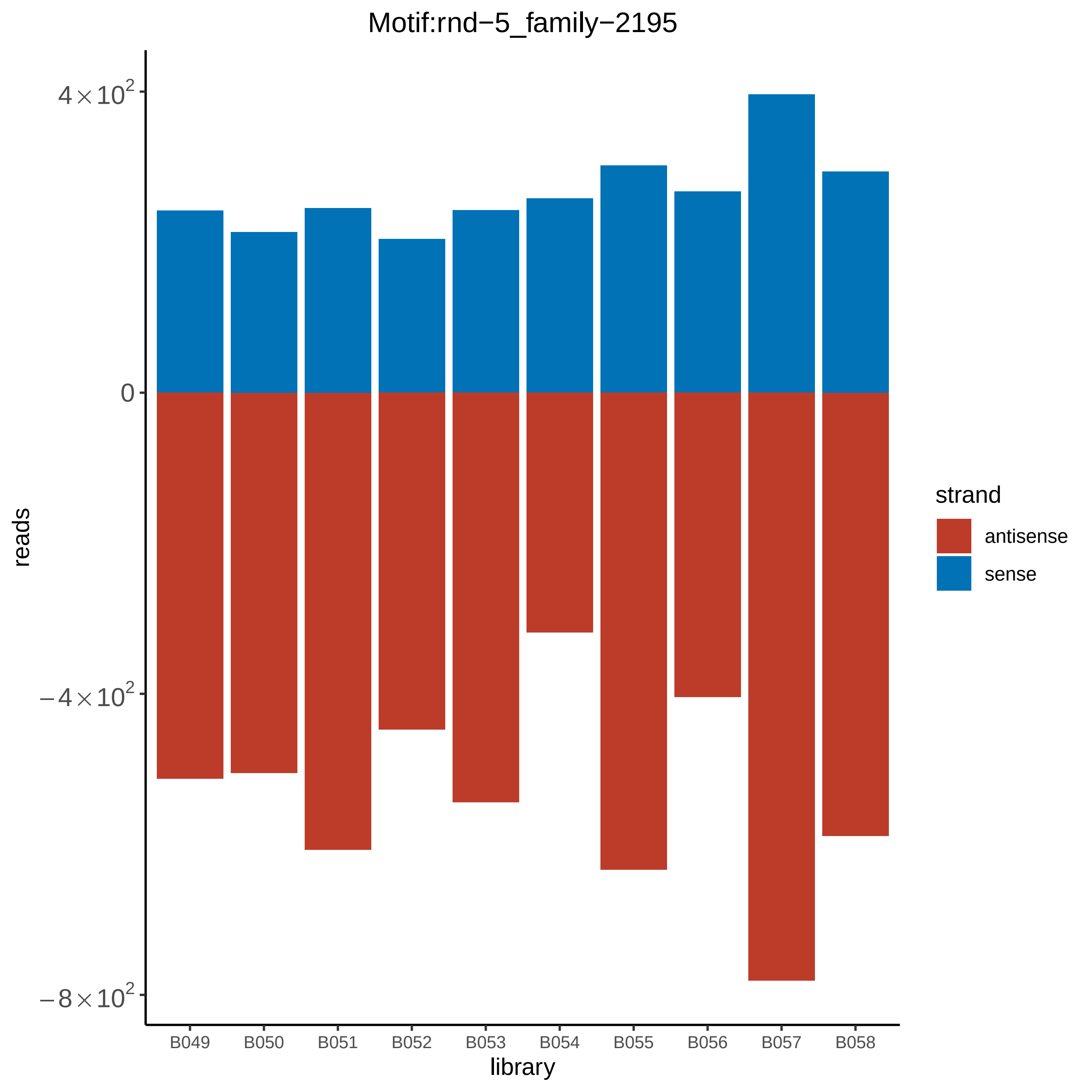

Motif:rnd−5_family−2195
4×10^2
0
−4×10^2
−8×10^2
reads
B049
B050
B051
B052
B053
B054
B055
B056
B057
B058
library
strand
antisense
sense

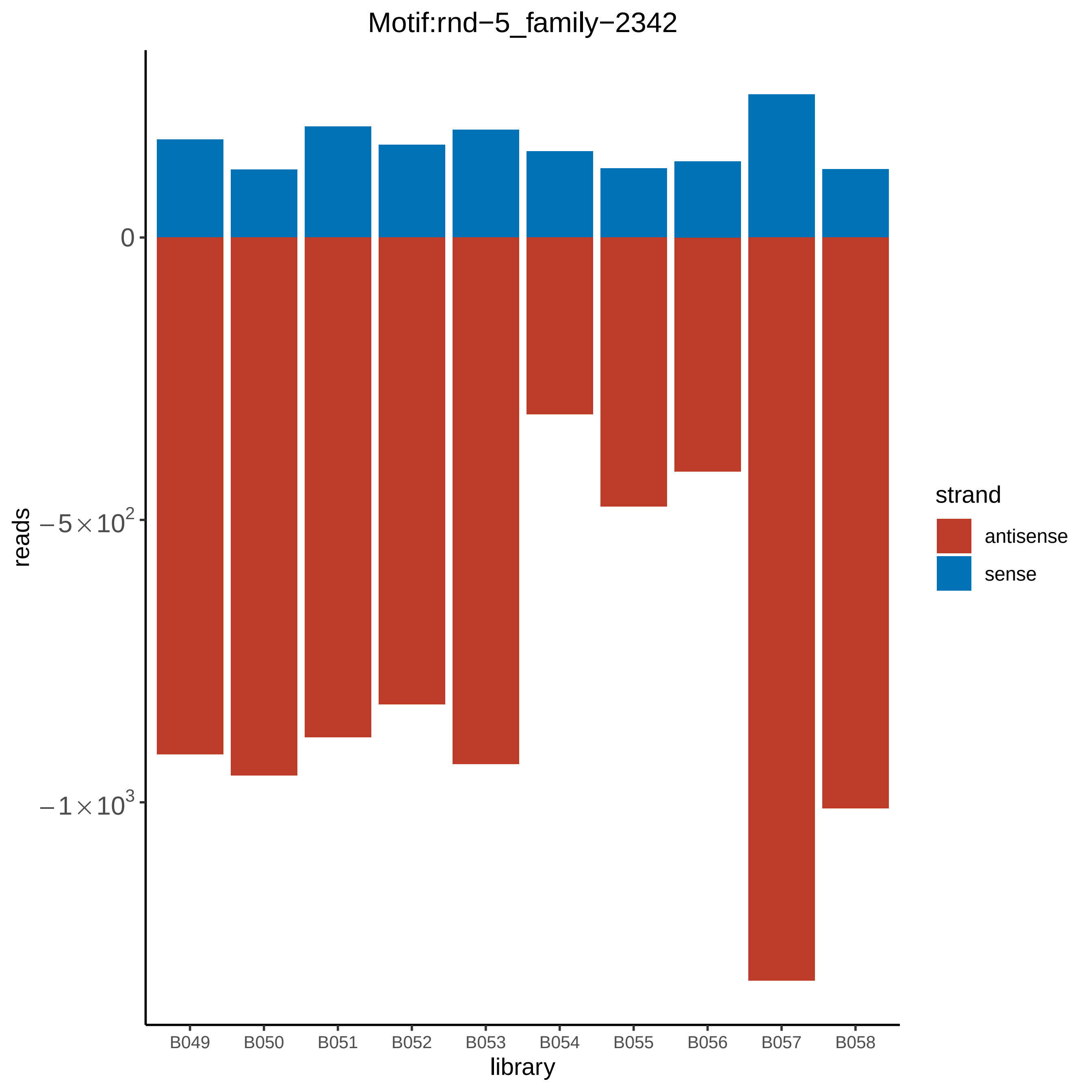

Motif:rnd−5_family−2342
0
−5 × 10^2
−1 × 10^3
reads
B049
B050
B051
B052
B053
B054
B055
B056
B057
B058
library
strand
antisense
sense

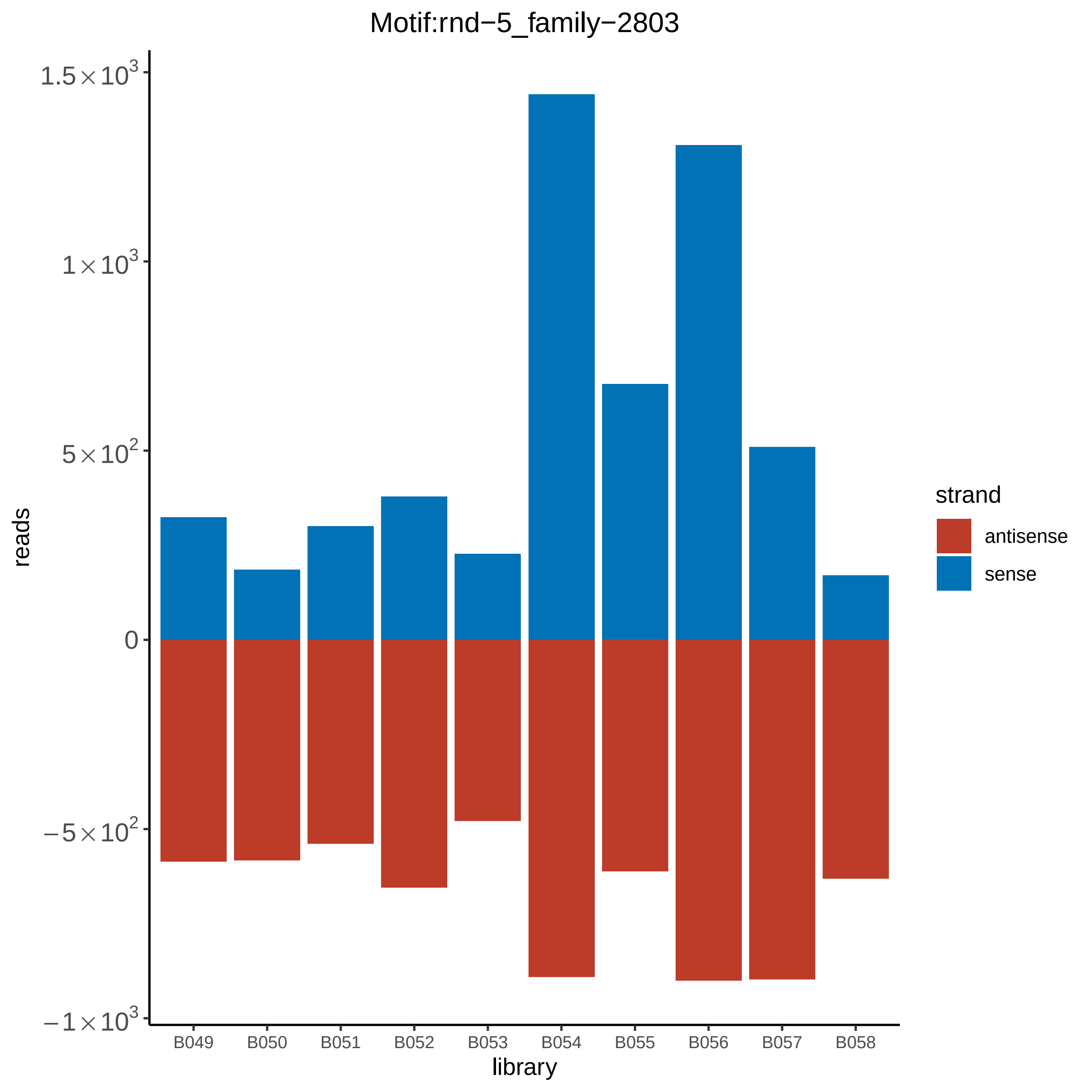
Motif:rnd−5_family−2803
reads
1.5×10^3
1×10^3
5×10^2
0
−5×10^2
−1×10^3
B049
B050
B051
B052
B053
B054
B055
B056
B057
B058
library
strand
antisense
sense

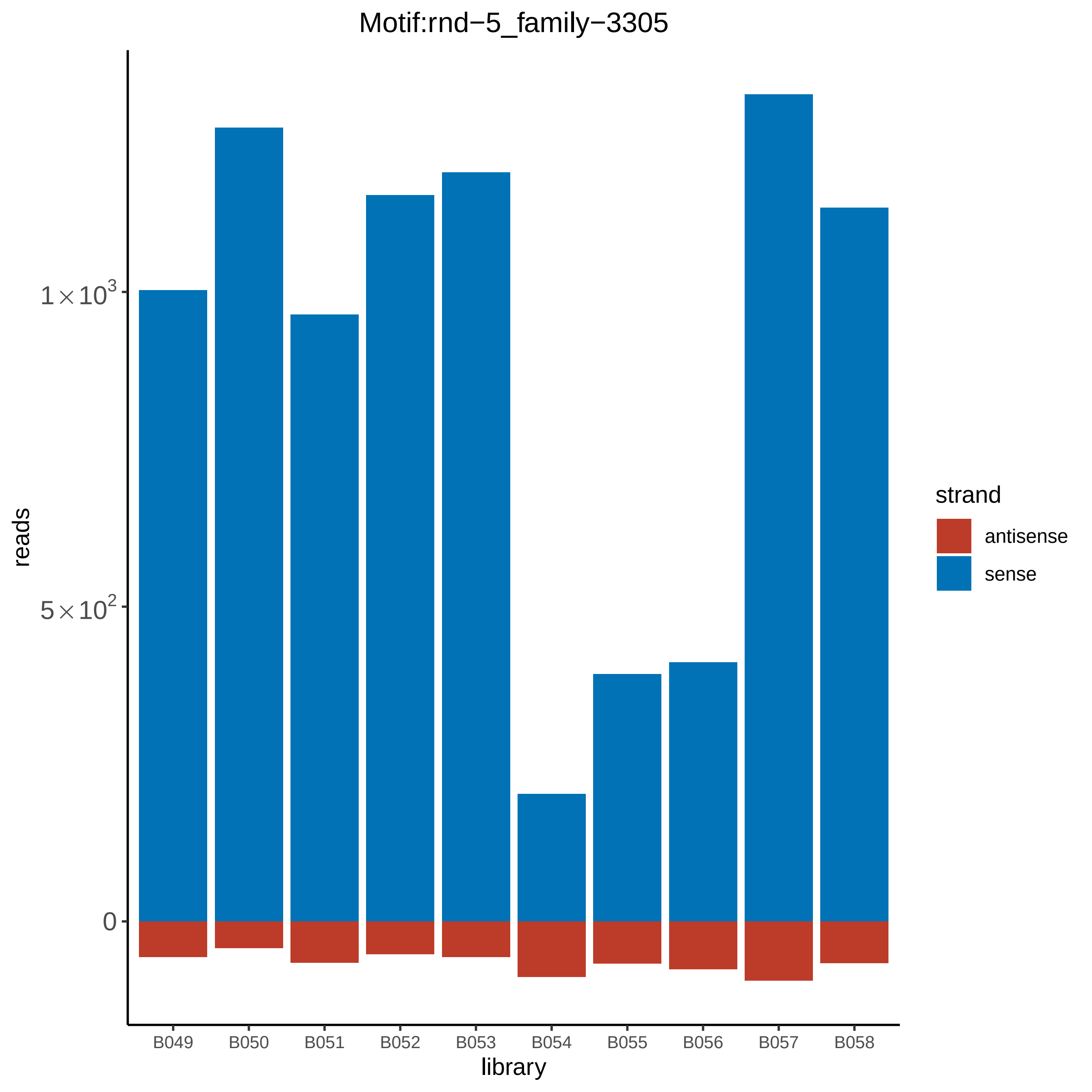

Motif:rnd−5_family−3305
reads
$1\times10^3$
$5\times10^2$
0
B049
B050
B051
B052
B053
B054
B055
B056
B057
B058
library
strand
antisense
sense

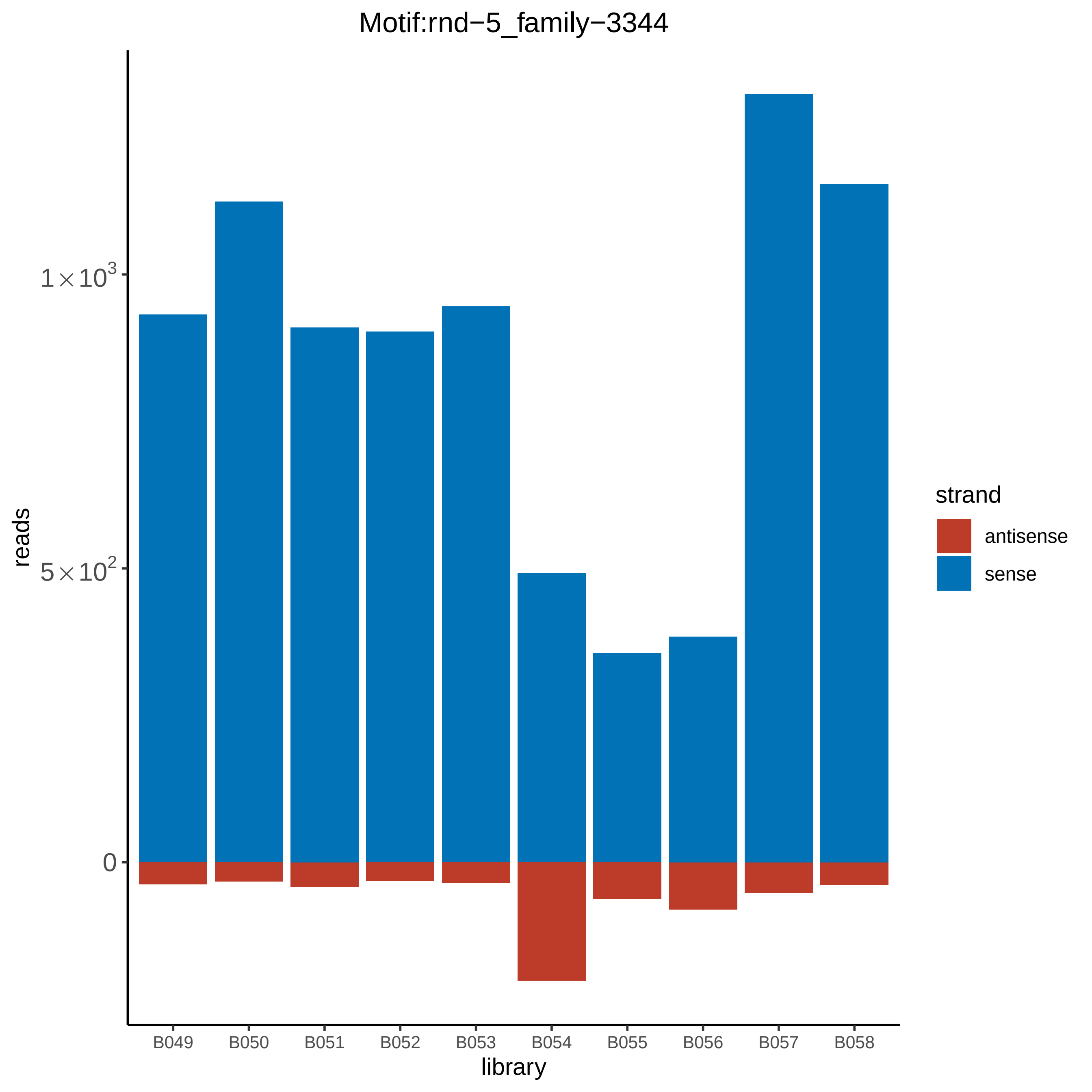

Motif:rnd−5_family−3344
reads
1 × 10^3
5 × 10^2
0
library
B049
B050
B051
B052
B053
B054
B055
B056
B057
B058
strand
antisense
sense

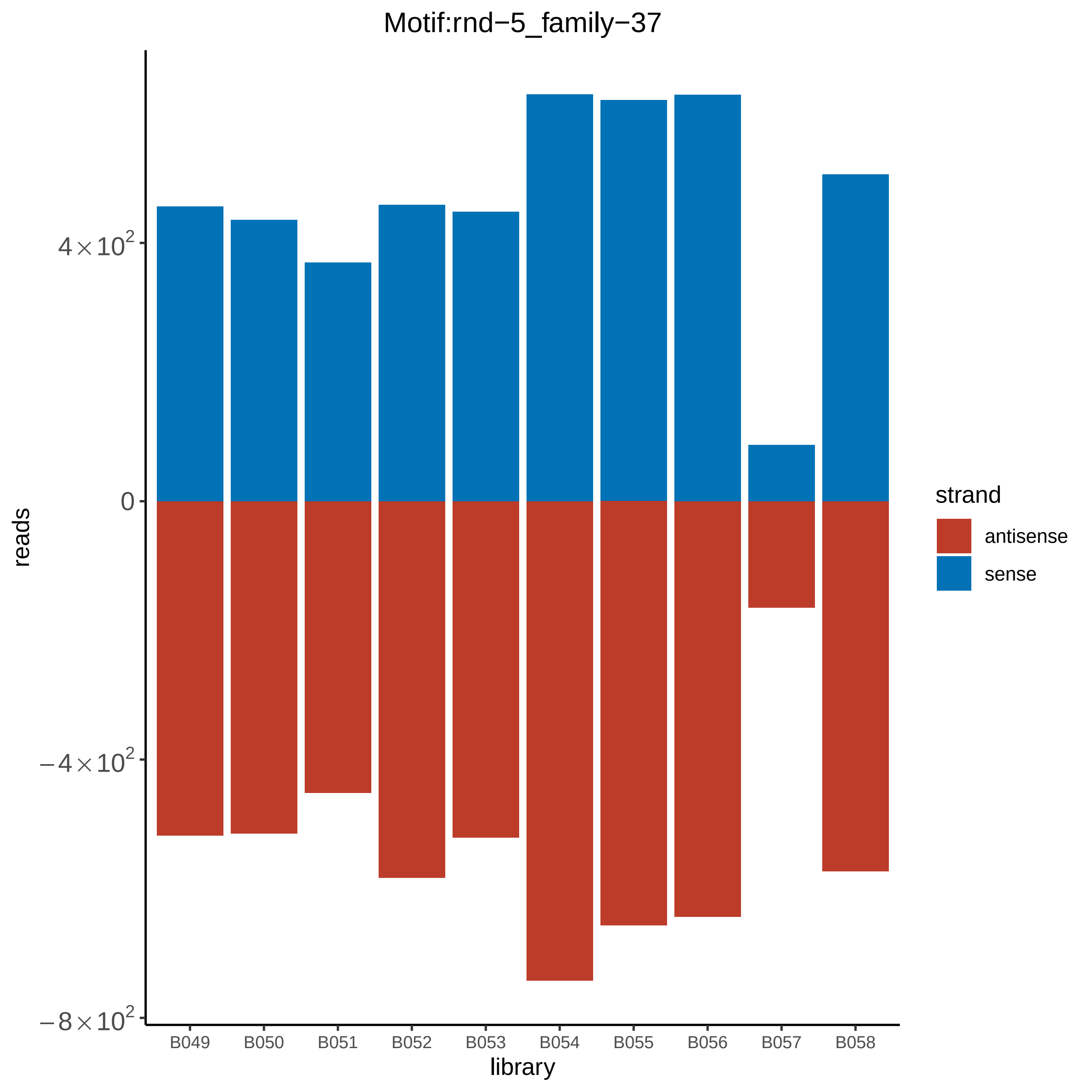

Motif:rnd−5_family−37
reads
4×10^2
0
−4×10^2
−8×10^2
B049
B050
B051
B052
B053
B054
B055
B056
B057
B058
library
strand
antisense
sense

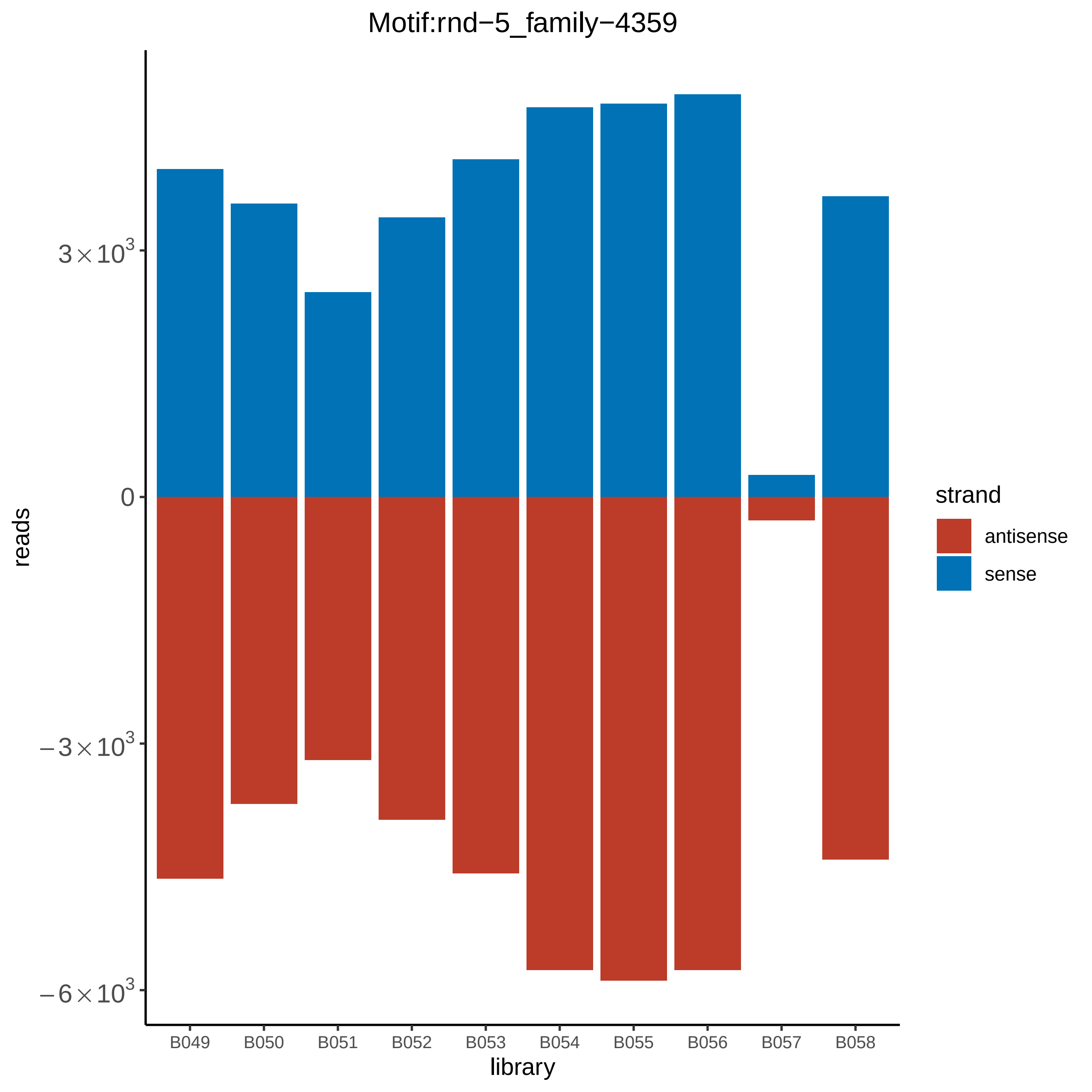
Motif:rnd−5_family−4359
reads
3×10^3
0
−3×10^3
−6×10^3
library
B049
B050
B051
B052
B053
B054
B055
B056
B057
B058
strand
antisense
sense

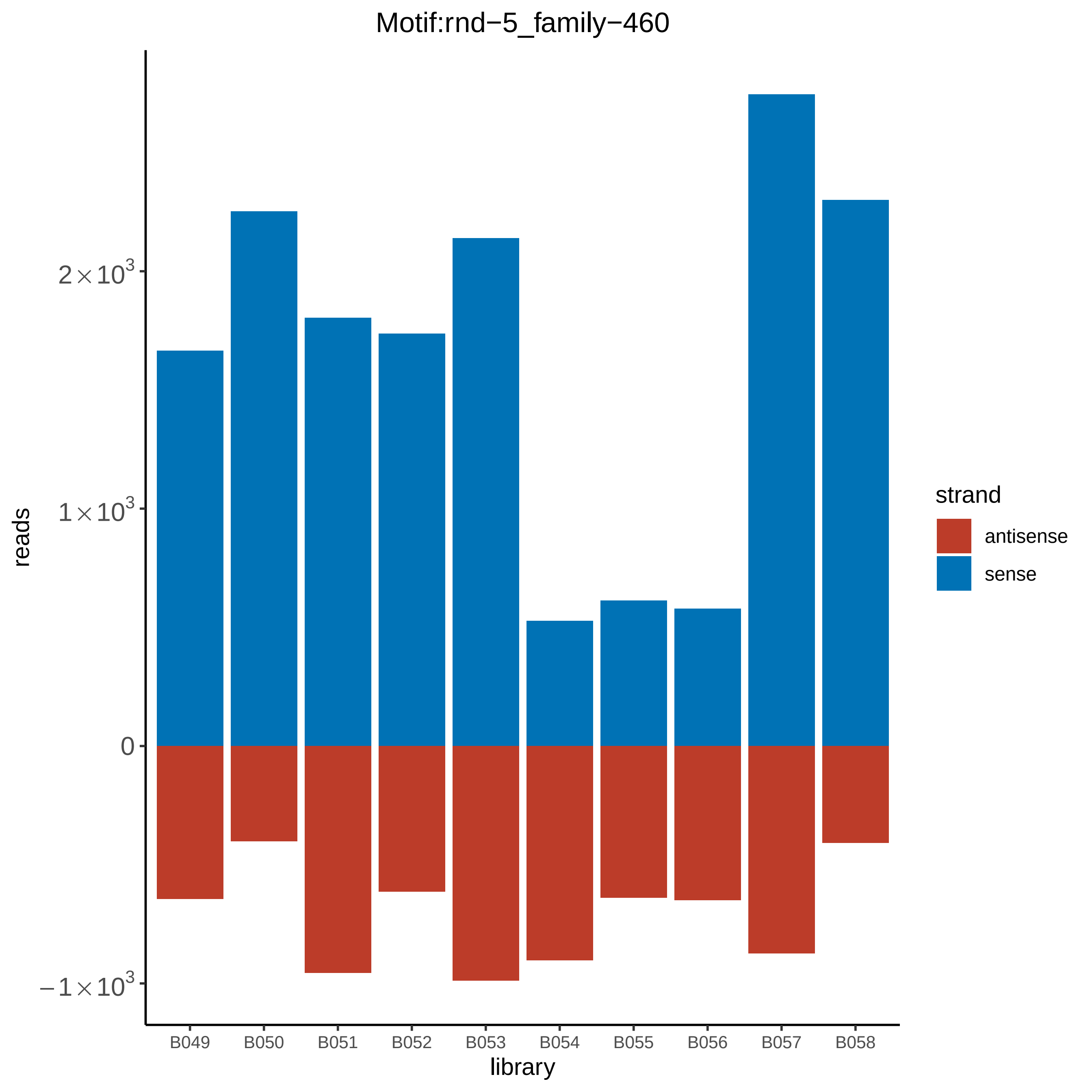

Motif:rnd−5_family−460
reads
$2\times10^3$
$1\times10^3$
0
$-1\times10^3$
library
B049
B050
B051
B052
B053
B054
B055
B056
B057
B058
strand
antisense
sense

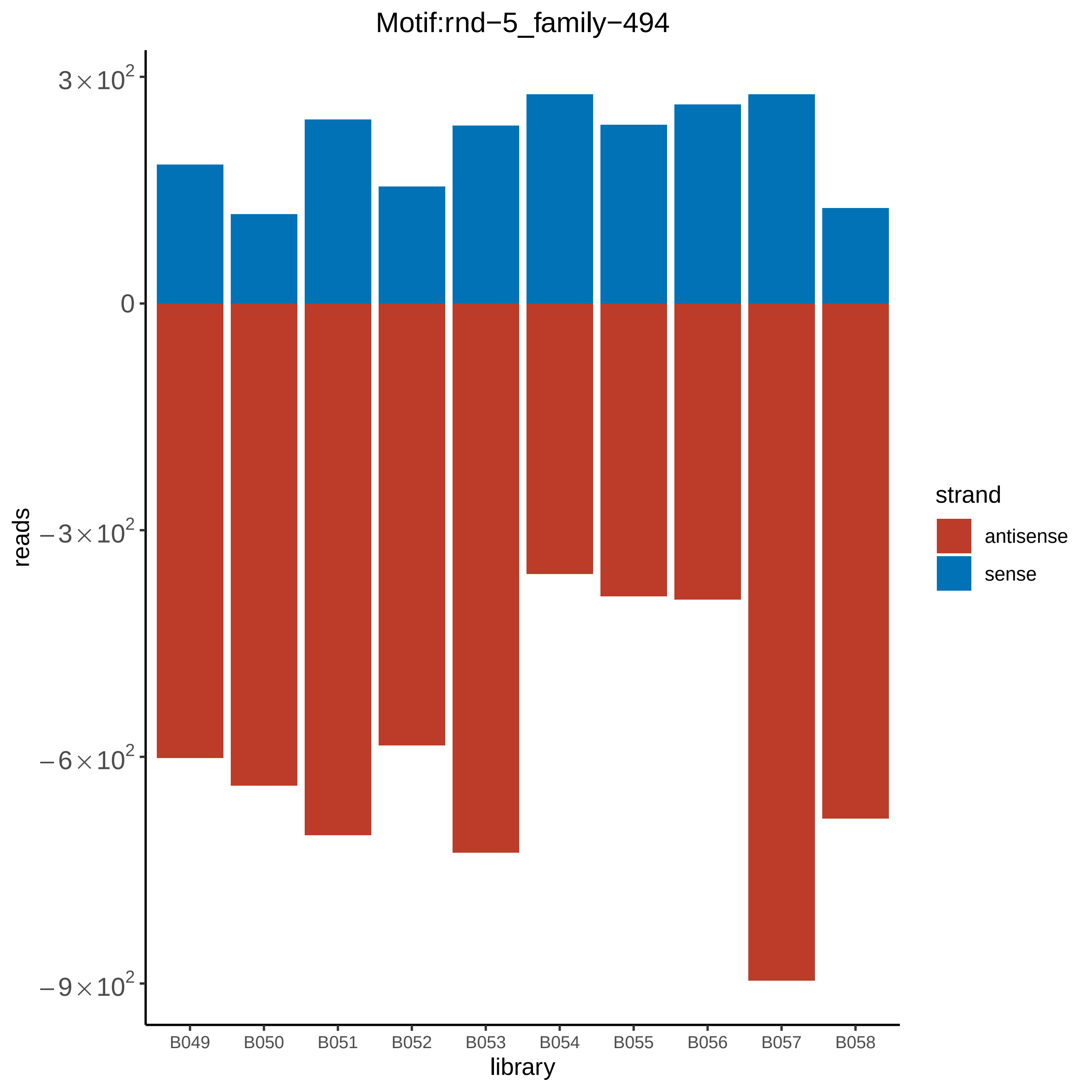

Motif:rnd−5_family−494
3×10^2
0
−3×10^2
−6×10^2
−9×10^2
reads
B049
B050
B051
B052
B053
B054
B055
B056
B057
B058
library
strand
antisense
sense

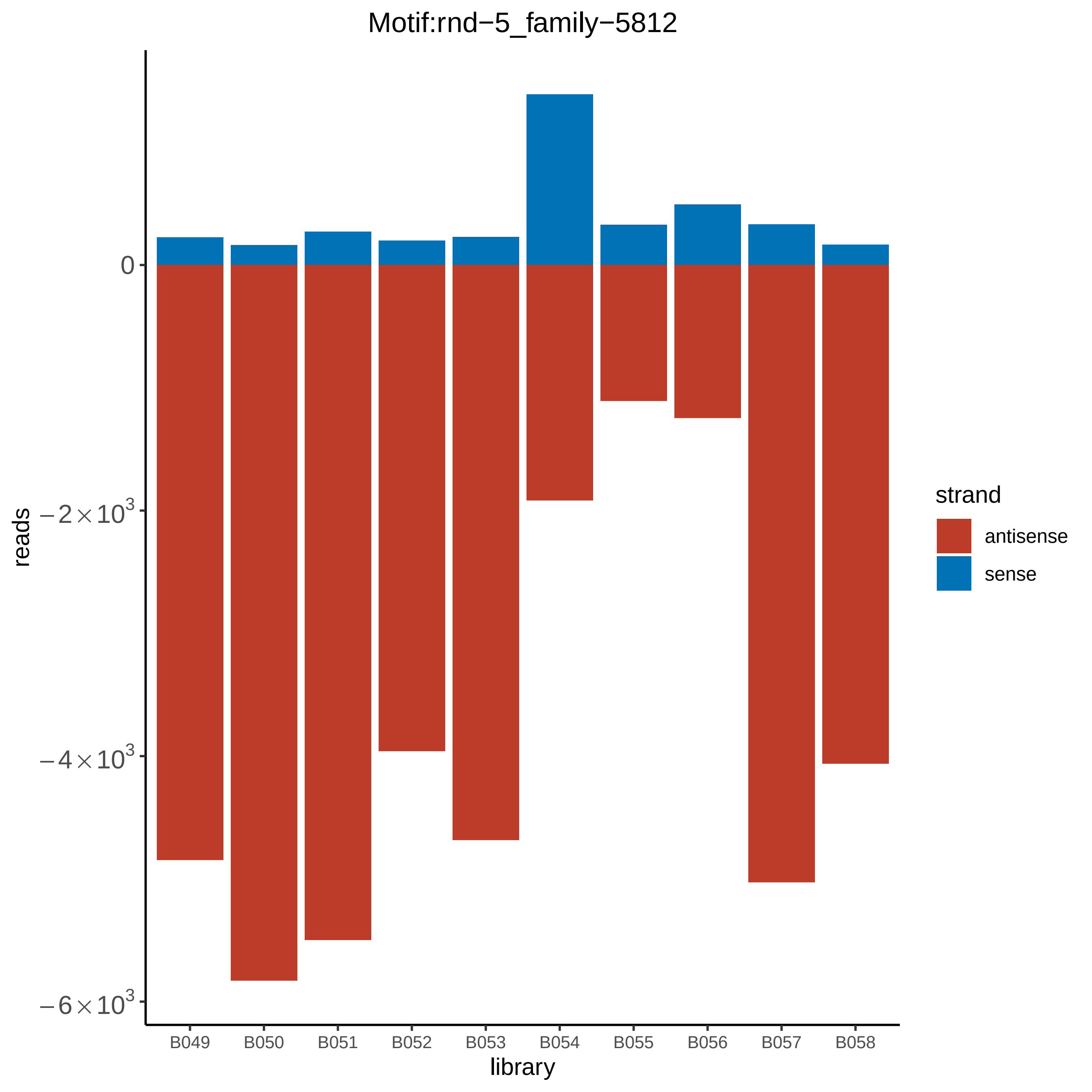

Motif:rnd−5_family−5812
0
−2×10^3
−4×10^3
−6×10^3
reads
B049
B050
B051
B052
B053
B054
B055
B056
B057
B058
library
strand
antisense
sense

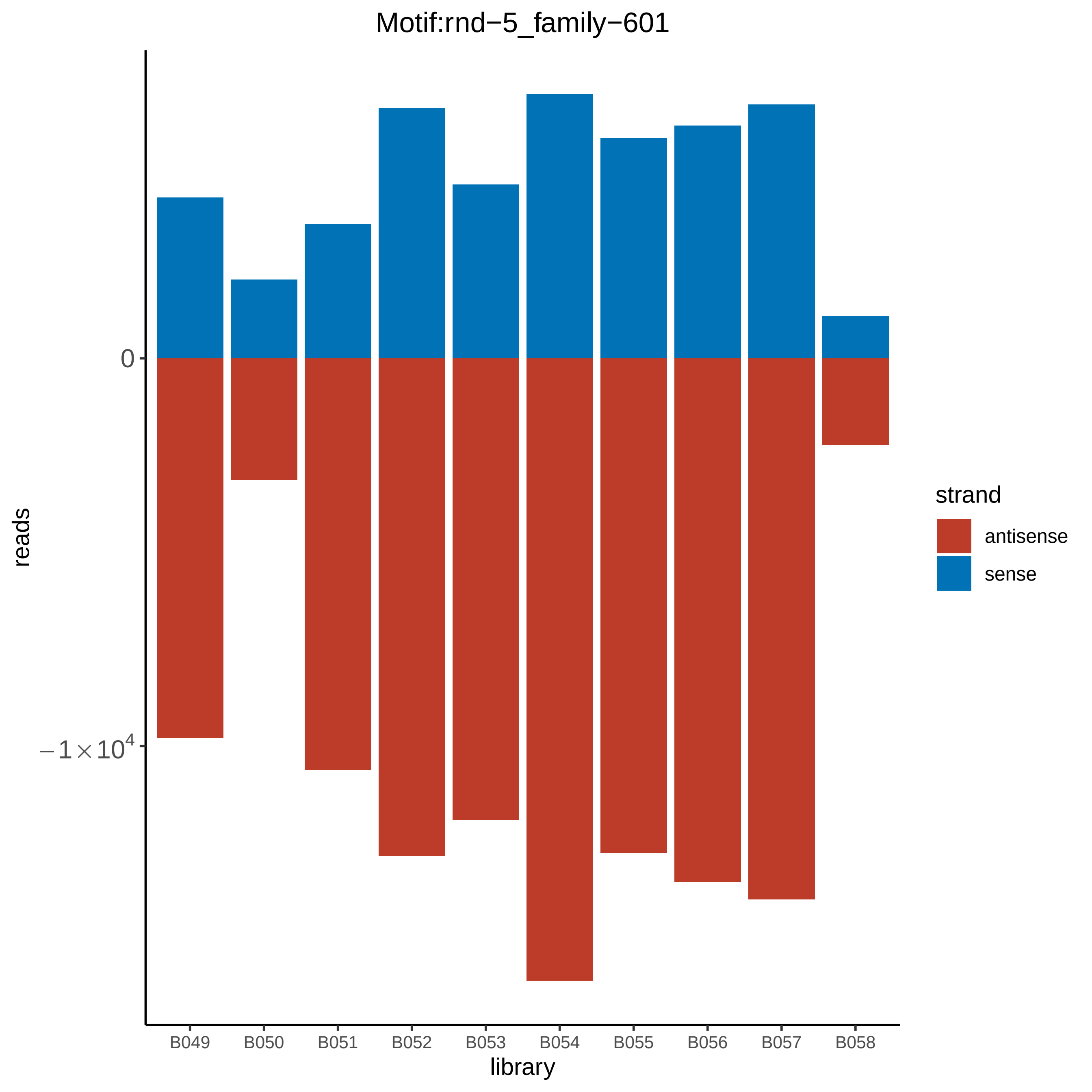

Motif:rnd−5_family−601
0
$-1\times10^4$
reads
B049
B050
B051
B052
B053
B054
B055
B056
B057
B058
library
strand
antisense
sense

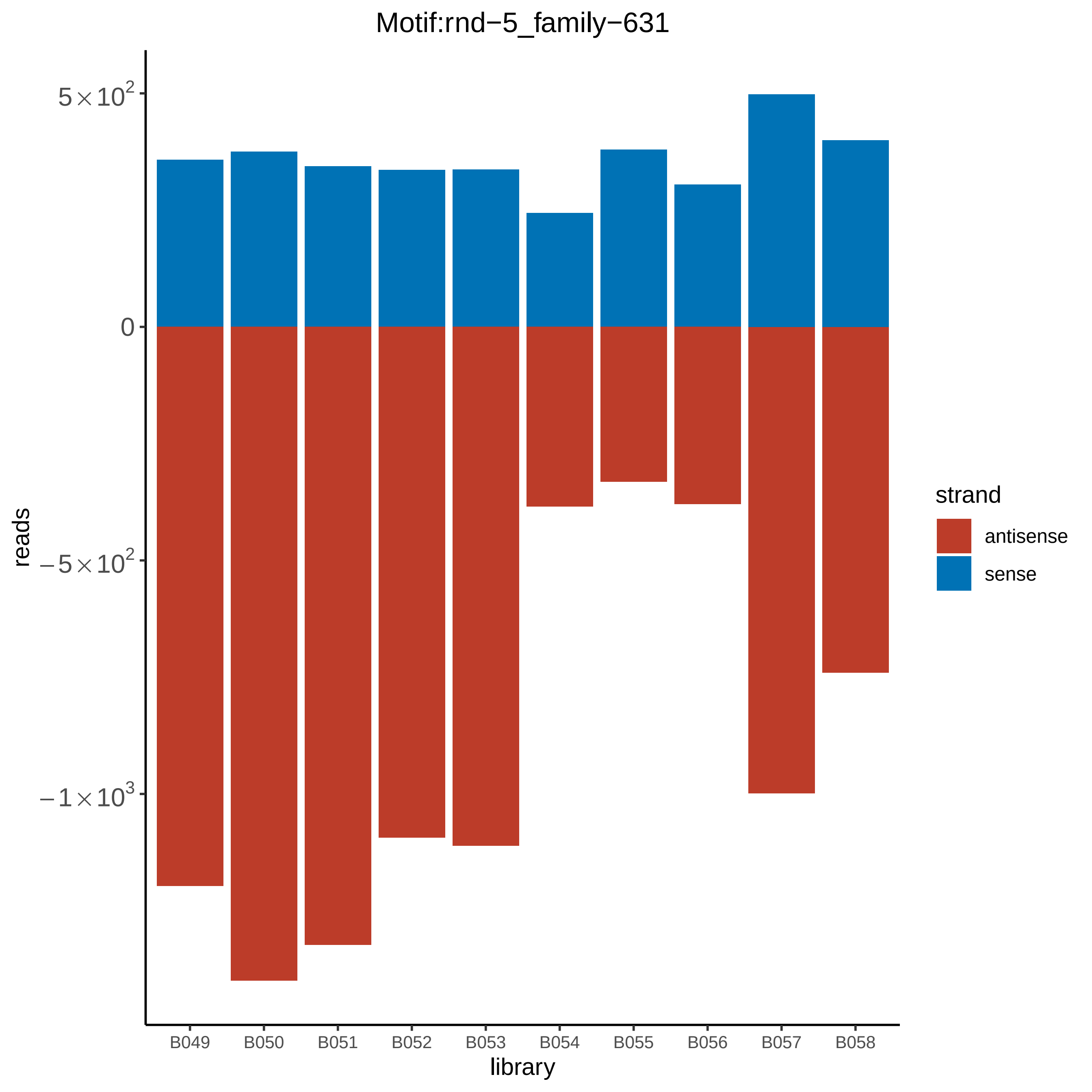

Motif:rnd−5_family−631
5×10^2
0
−5×10^2
−1×10^3
reads
B049
B050
B051
B052
B053
B054
B055
B056
B057
B058
library
strand
antisense
sense

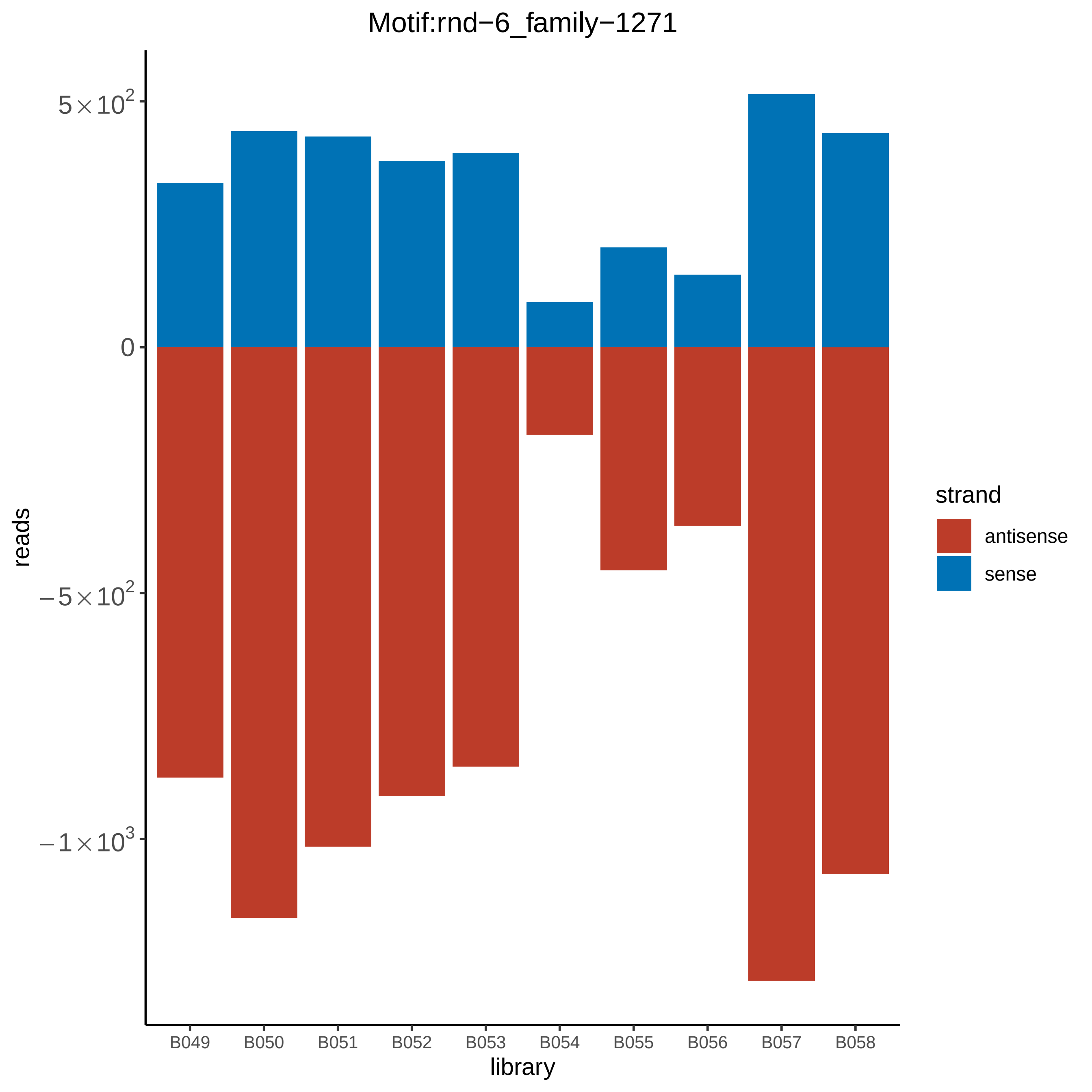

Motif:rnd−6_family−1271
5×10^2
0
−5×10^2
−1×10^3
reads
B049
B050
B051
B052
B053
B054
B055
B056
B057
B058
library
strand
antisense
sense

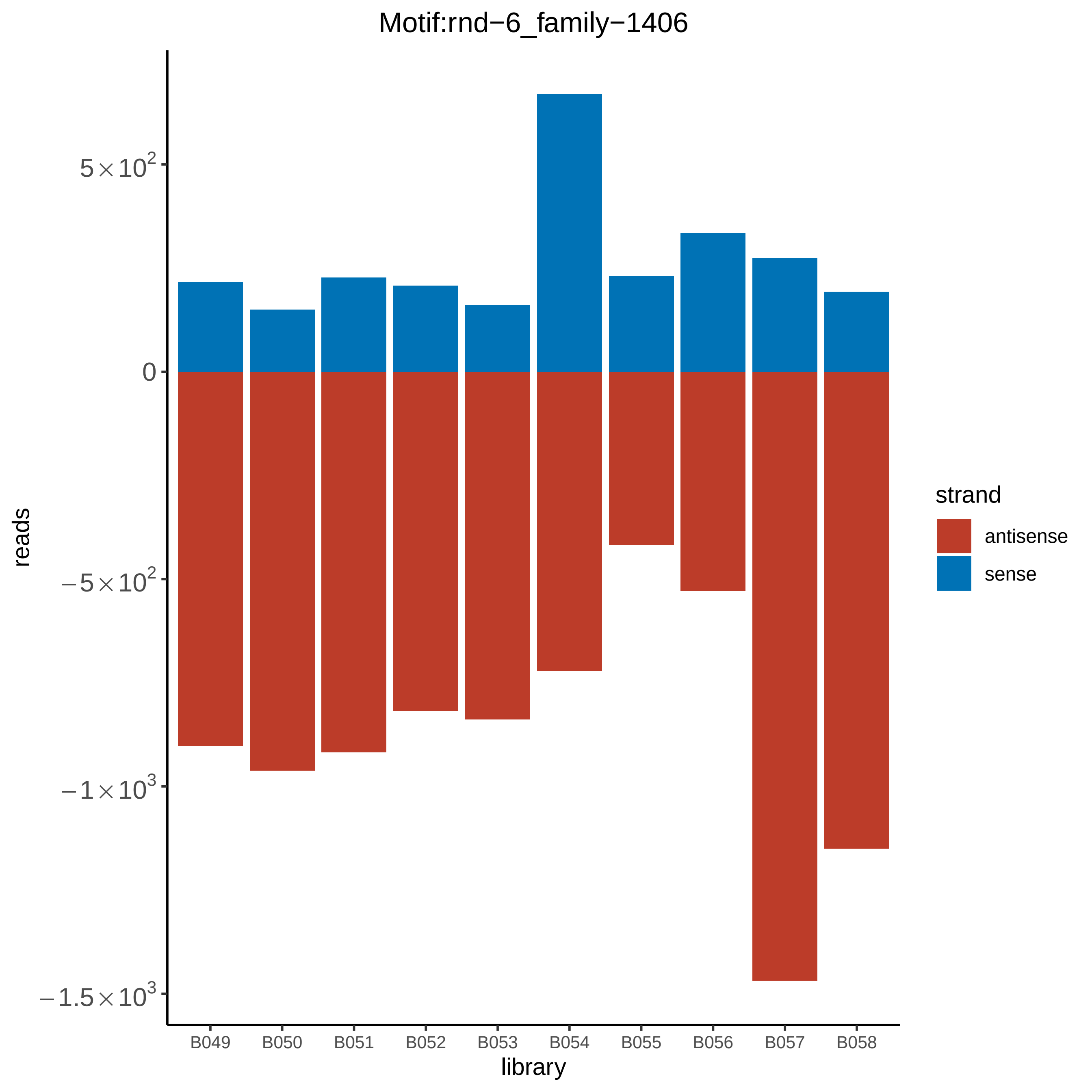

Motif:rnd−6_family−1406
reads
5×10^2
0
−5×10^2
−1×10^3
−1.5×10^3
B049
B050
B051
B052
B053
B054
B055
B056
B057
B058
library
strand
antisense
sense

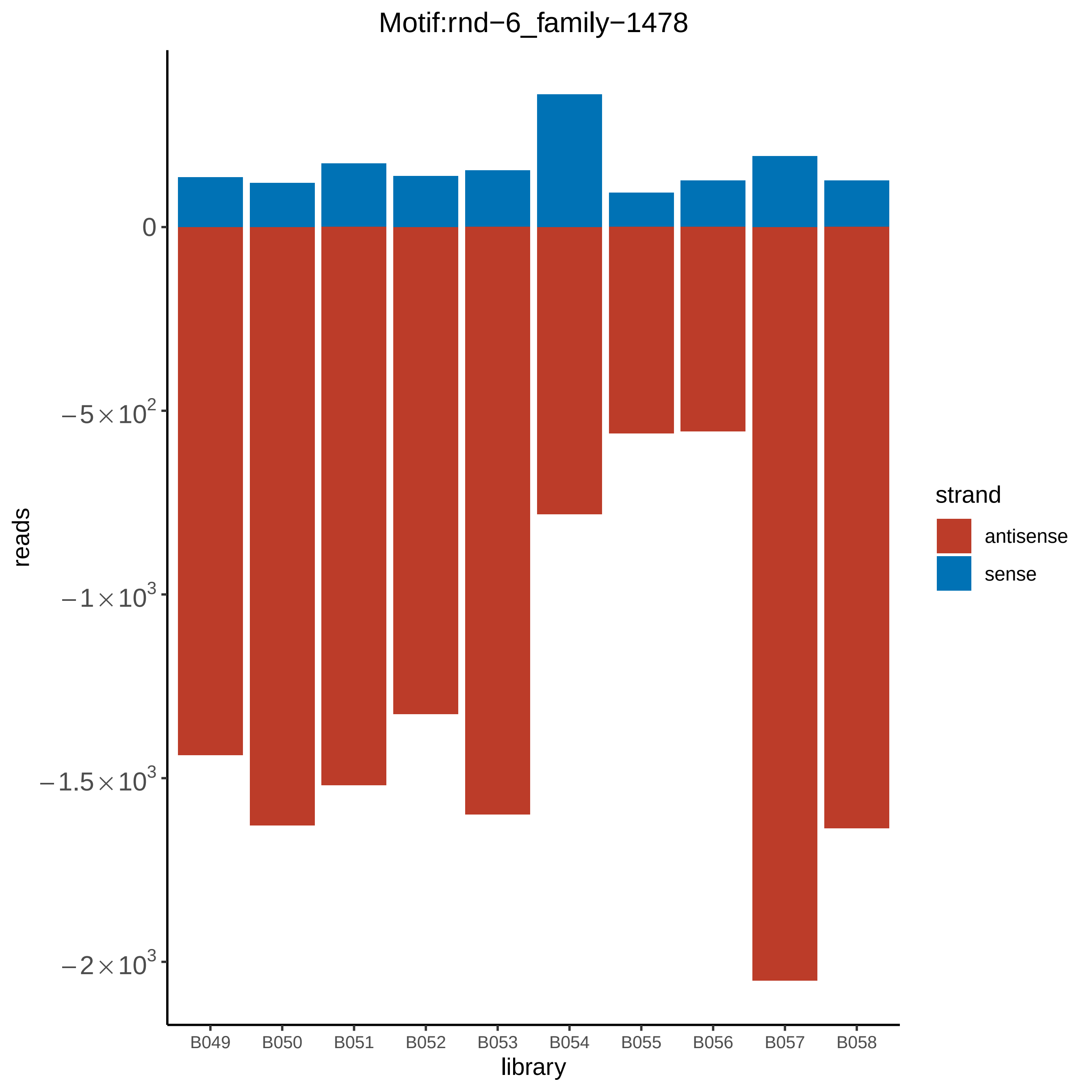

Motif:rnd−6_family−1478
0
$-5\times10^2$
$-1\times10^3$
$-1.5\times10^3$
$-2\times10^3$
reads
B049
B050
B051
B052
B053
B054
B055
B056
B057
B058
library
strand
antisense
sense

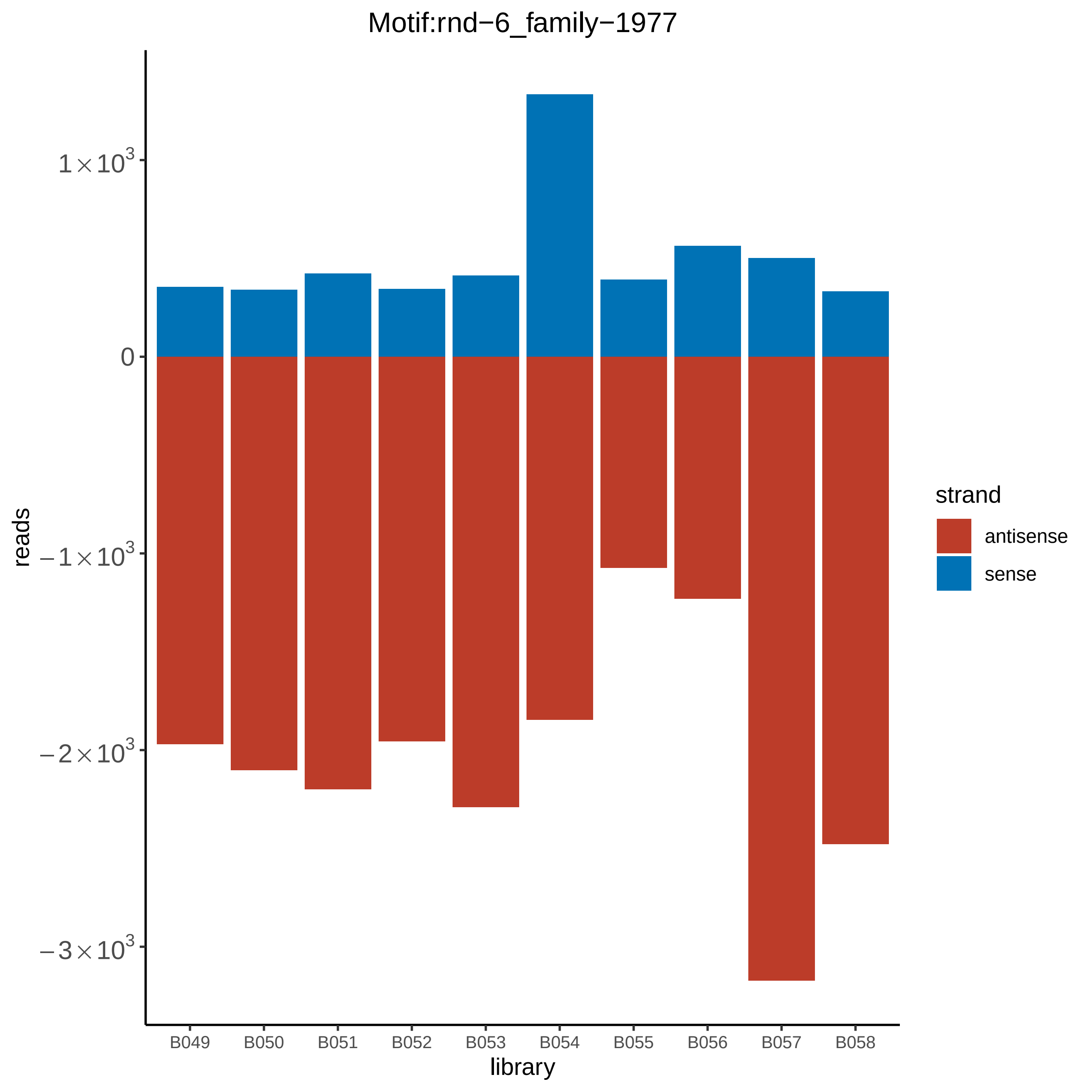
Motif:rnd−6_family−1977
1×10^3
0
−1×10^3
−2×10^3
−3×10^3
reads
B049
B050
B051
B052
B053
B054
B055
B056
B057
B058
library
strand
antisense
sense

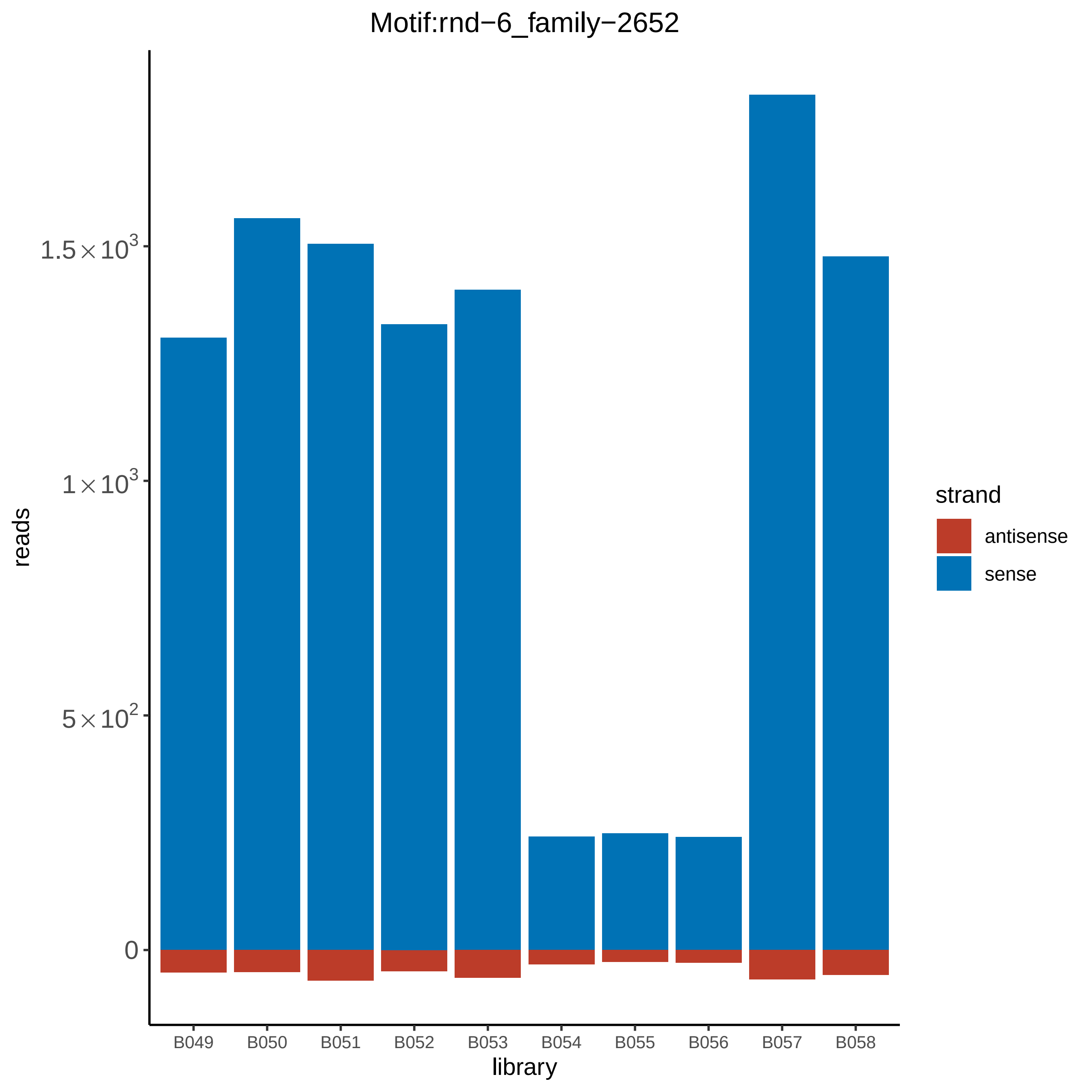

Motif:rnd−6_family−2652
reads
1.5×10^3
1×10^3
5×10^2
0
library
B049
B050
B051
B052
B053
B054
B055
B056
B057
B058
strand
antisense
sense

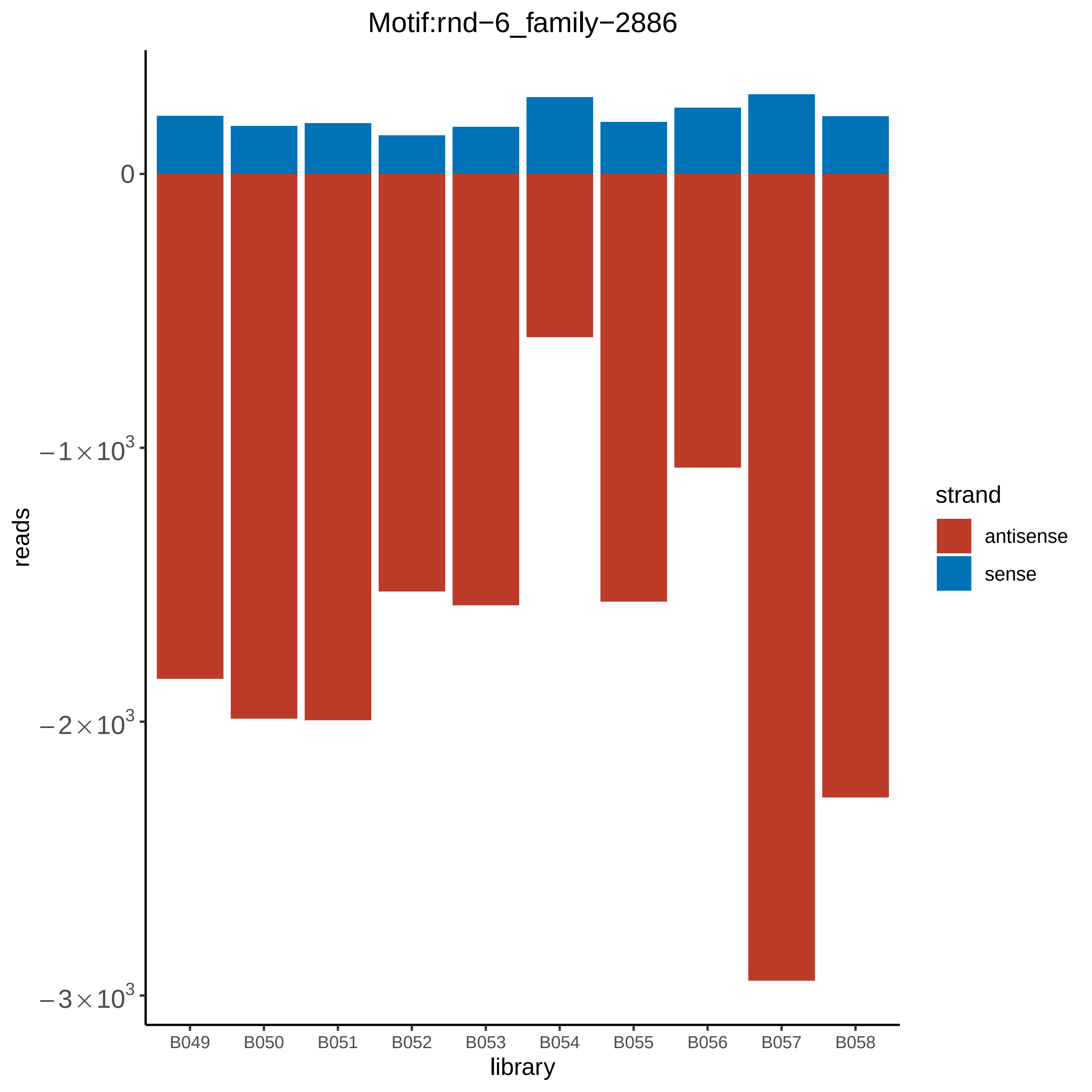

Motif:rnd−6_family−2886
reads
0
$-1\times10^3$
$-2\times10^3$
$-3\times10^3$
library
B049
B050
B051
B052
B053
B054
B055
B056
B057
B058
strand
antisense
sense

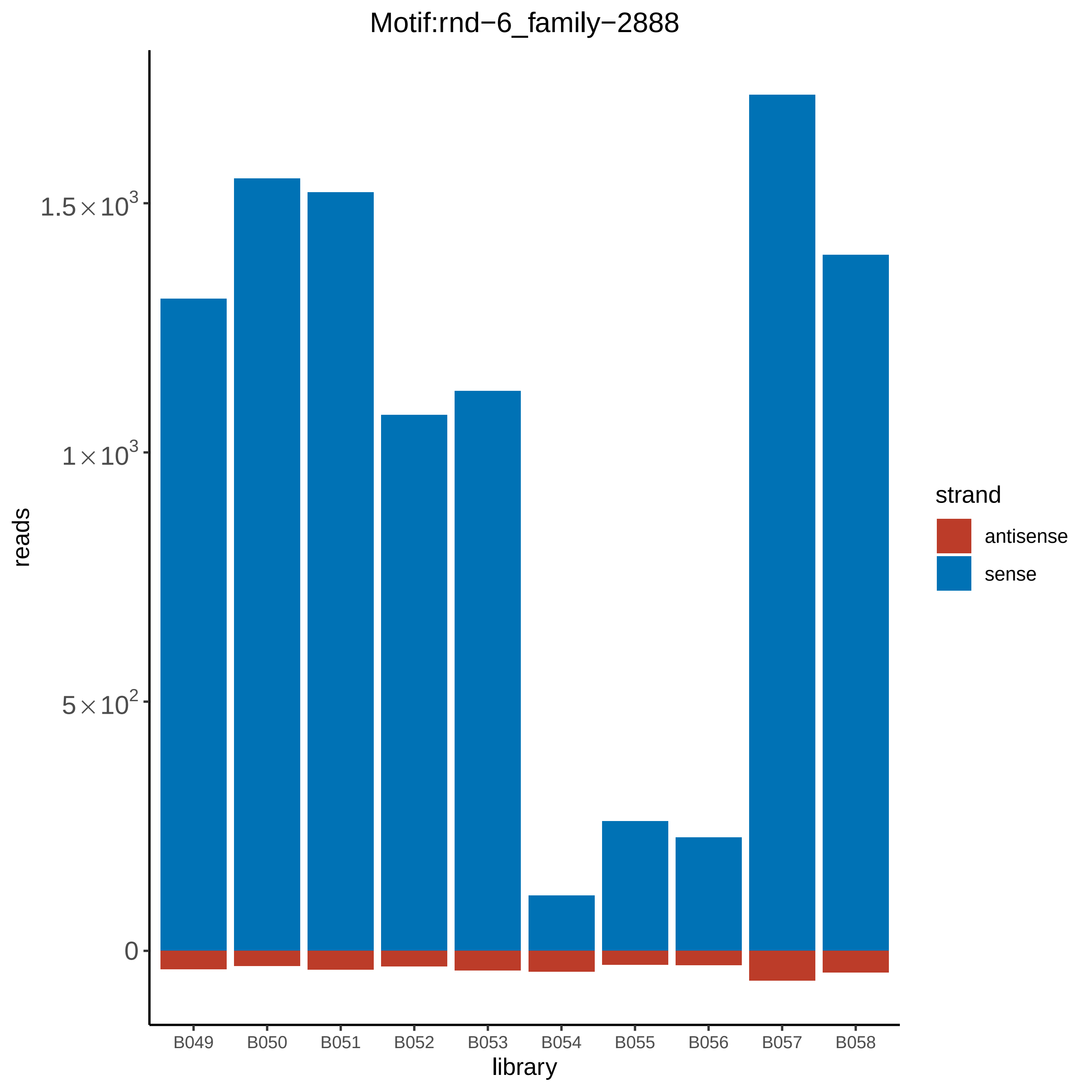

Motif:rnd−6_family−2888
reads
1.5×10^3
1×10^3
5×10^2
0
B049
B050
B051
B052
B053
B054
B055
B056
B057
B058
library
strand
antisense
sense

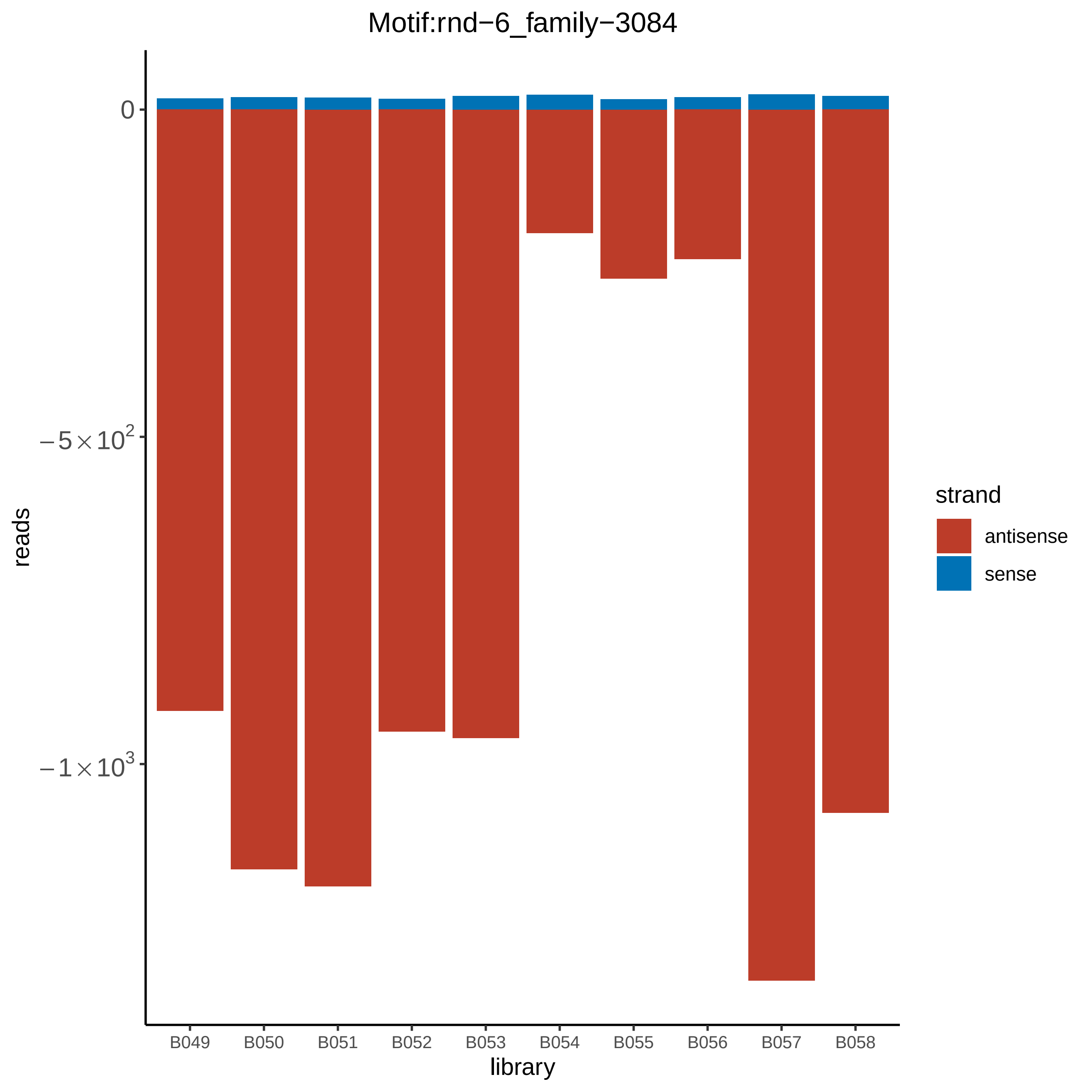
Motif:rnd−6_family−3084
0
$-5\times10^2$
$-1\times10^3$
reads
B049
B050
B051
B052
B053
B054
B055
B056
B057
B058
library
strand
antisense
sense

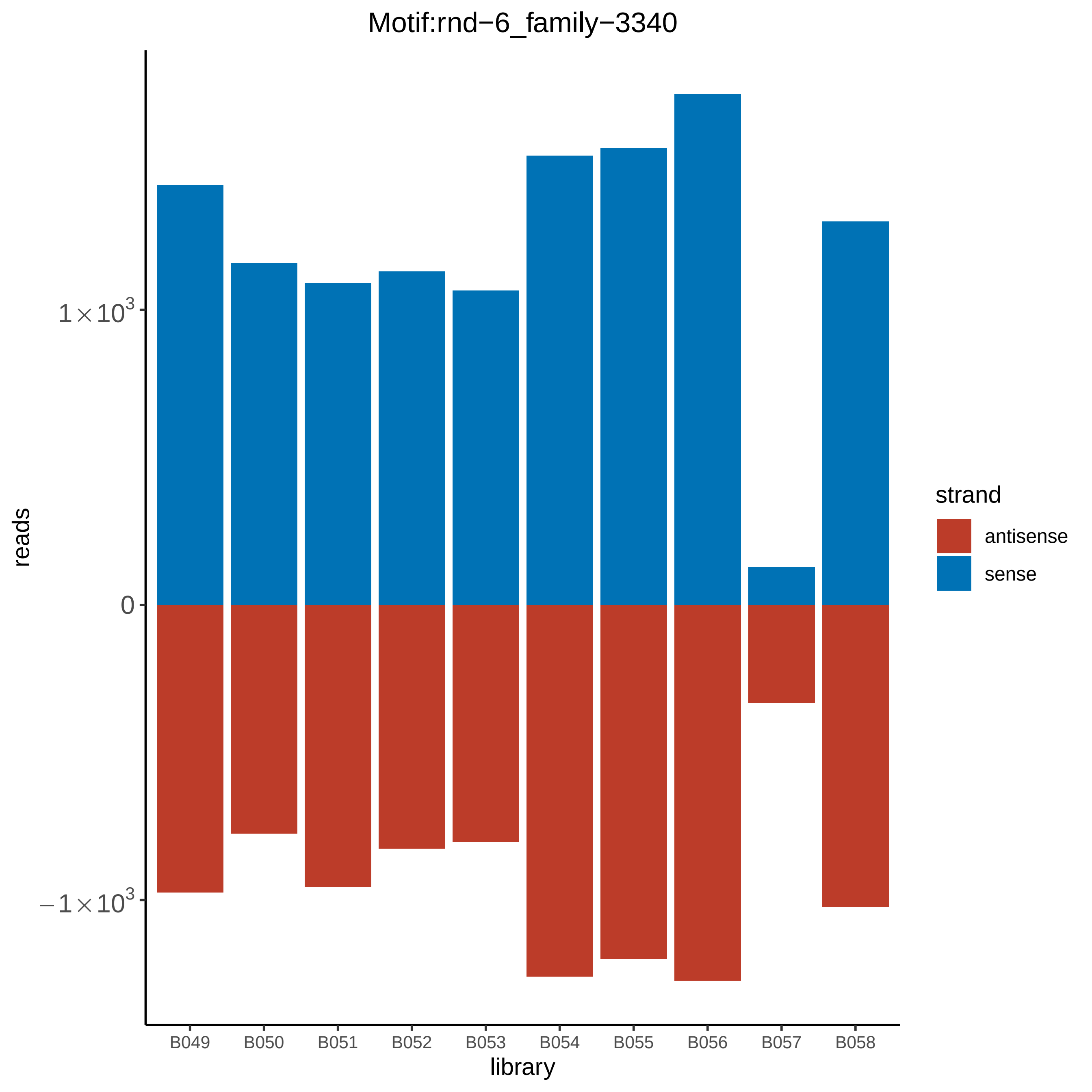

Motif:rnd−6_family−3340
reads
1×10^3
0
−1×10^3
library
B049
B050
B051
B052
B053
B054
B055
B056
B057
B058
strand
antisense
sense

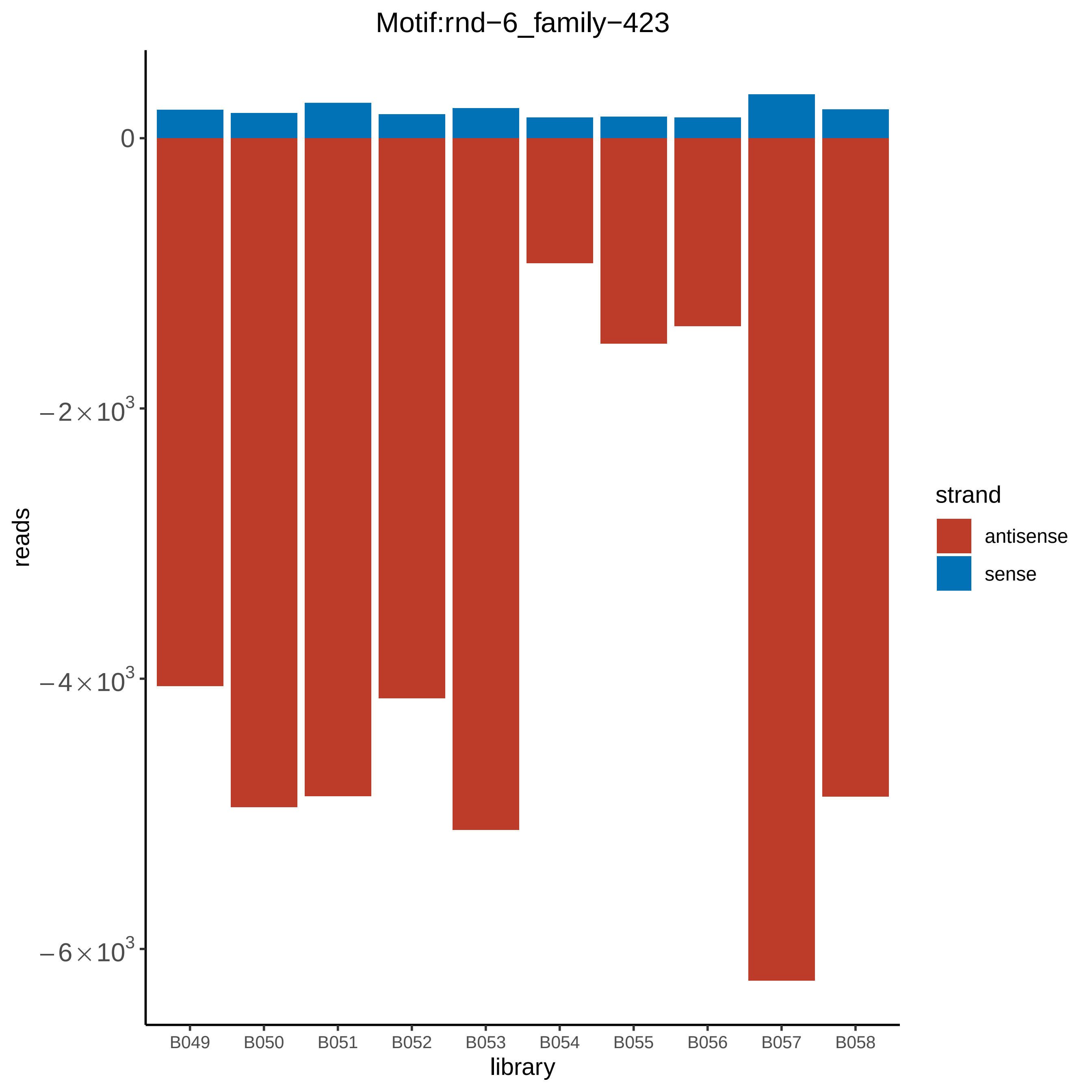

Motif:rnd−6_family−423
0
−2×10^3
−4×10^3
−6×10^3
reads
B049
B050
B051
B052
B053
B054
B055
B056
B057
B058
library
strand
antisense
sense

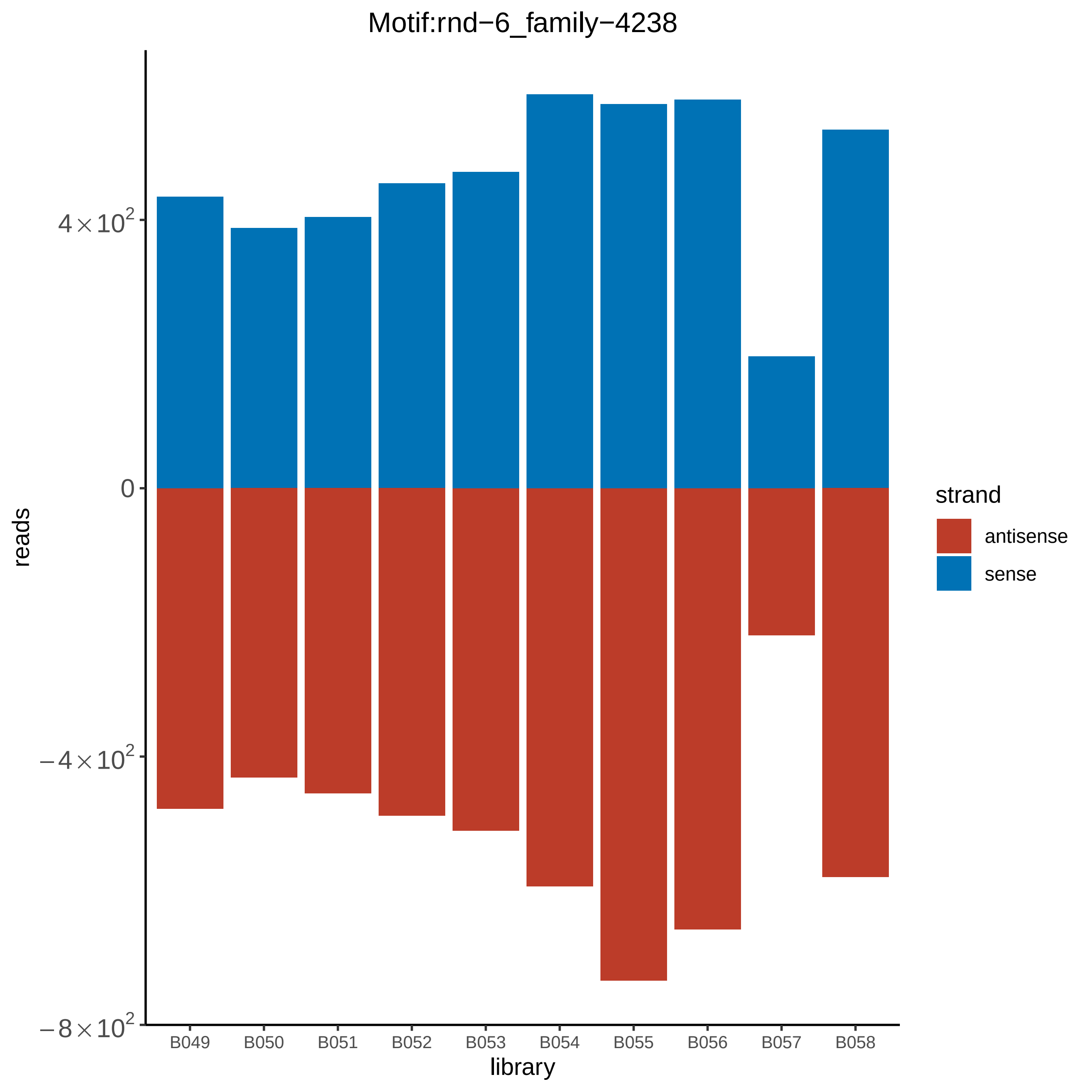
Motif:rnd−6_family−4238
reads
$4\times10^2$
0
$-4\times10^2$
$-8\times10^2$
B049
B050
B051
B052
B053
B054
B055
B056
B057
B058
library
strand
antisense
sense

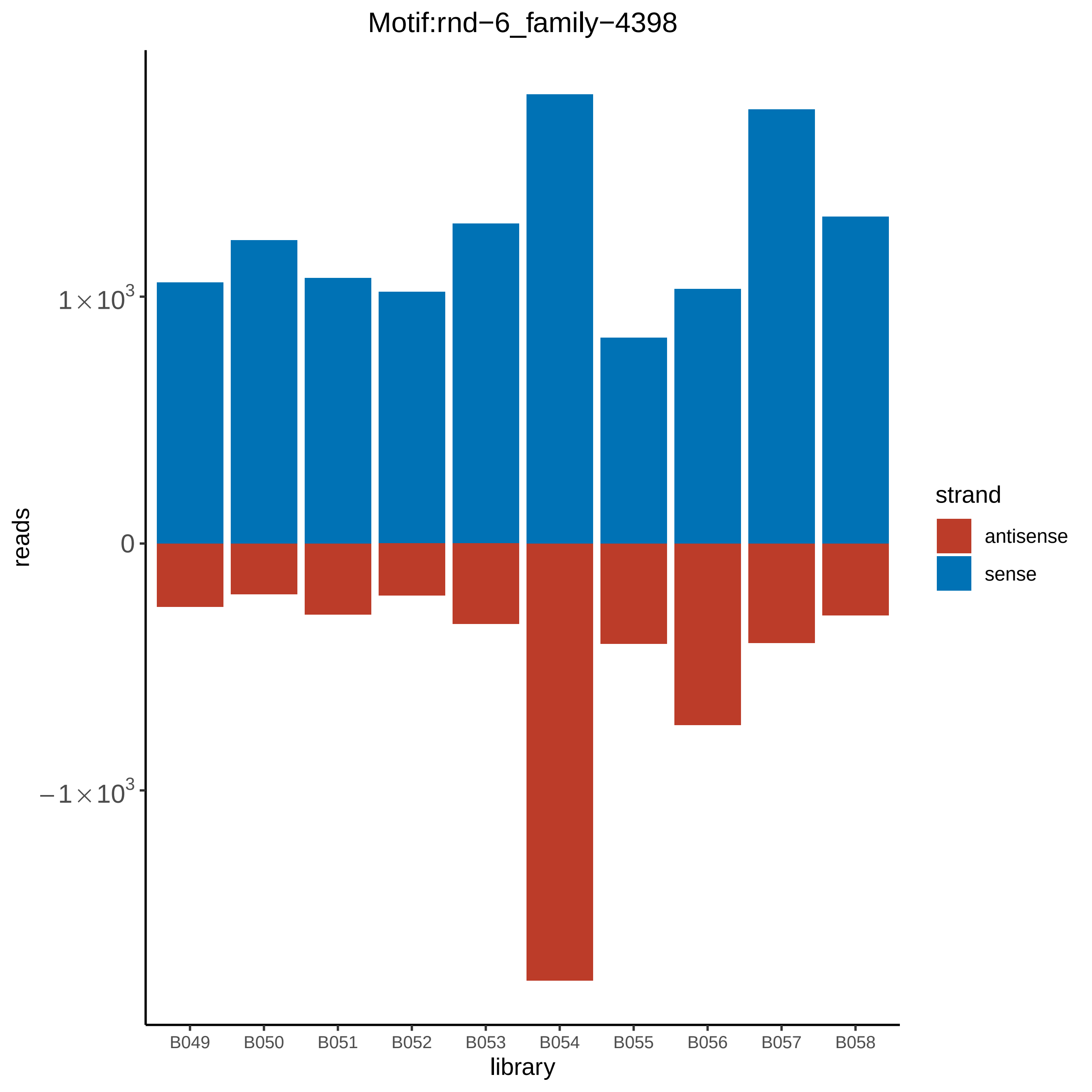

Motif:rnd−6_family−4398
reads
1×10^3
0
−1×10^3
library
B049
B050
B051
B052
B053
B054
B055
B056
B057
B058
strand
antisense
sense

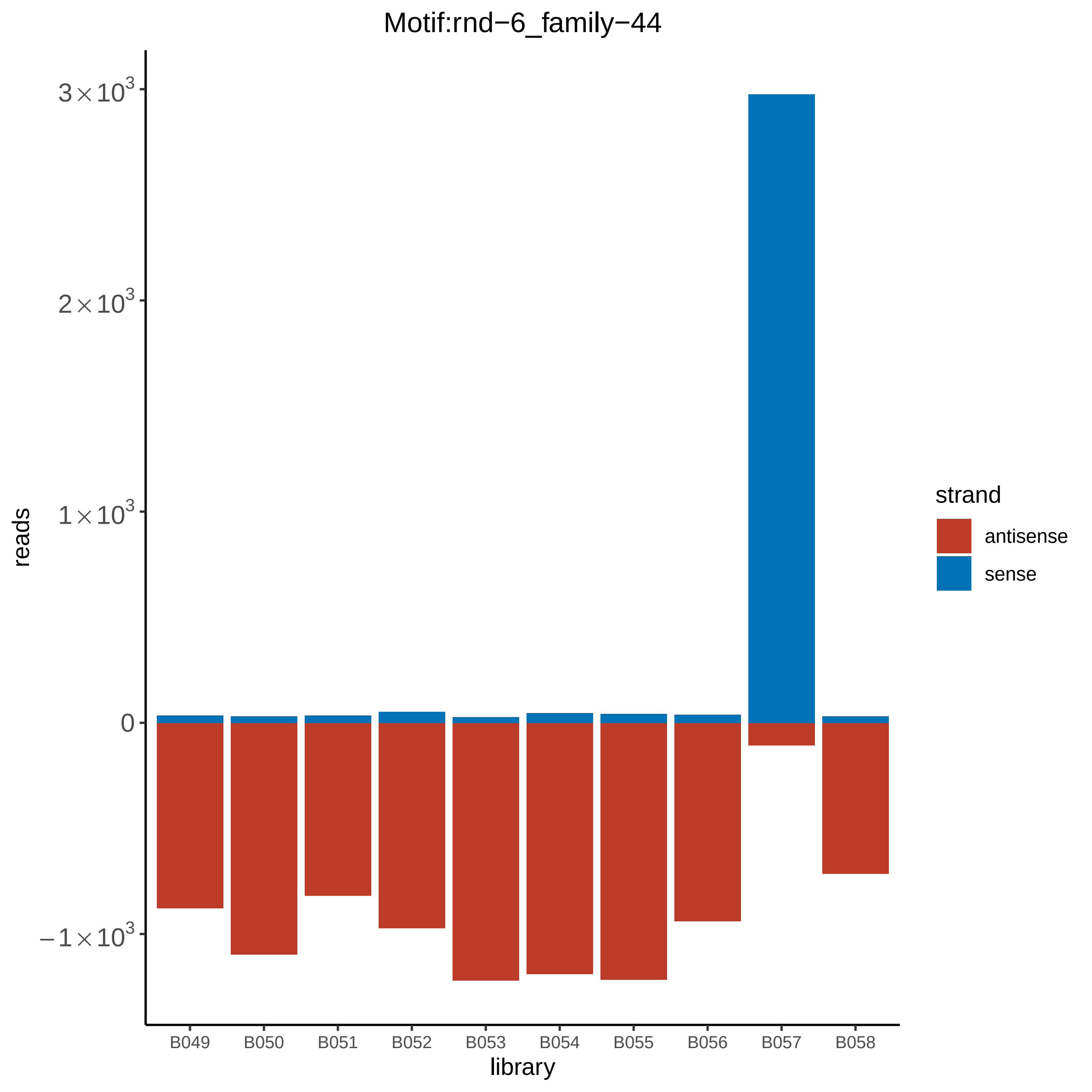

Motif:rnd−6_family−44
reads
$3\times10^3$
$2\times10^3$
$1\times10^3$
0
$-1\times10^3$
library
B049
B050
B051
B052
B053
B054
B055
B056
B057
B058
strand
antisense
sense

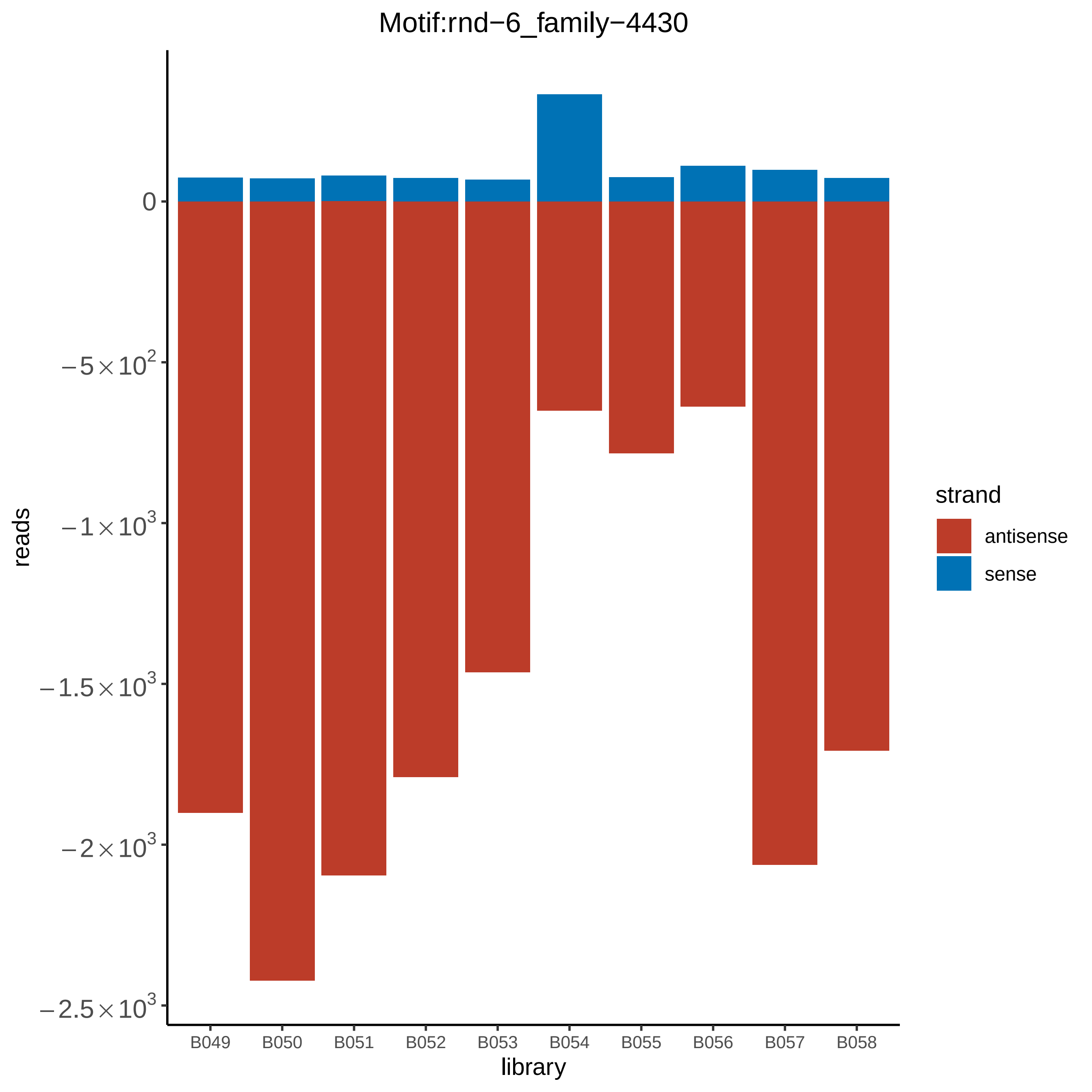
Motif:rnd−6_family−4430
0
−5×10^2
−1×10^3
−1.5×10^3
−2×10^3
−2.5×10^3
reads
B049
B050
B051
B052
B053
B054
B055
B056
B057
B058
library
strand
antisense
sense

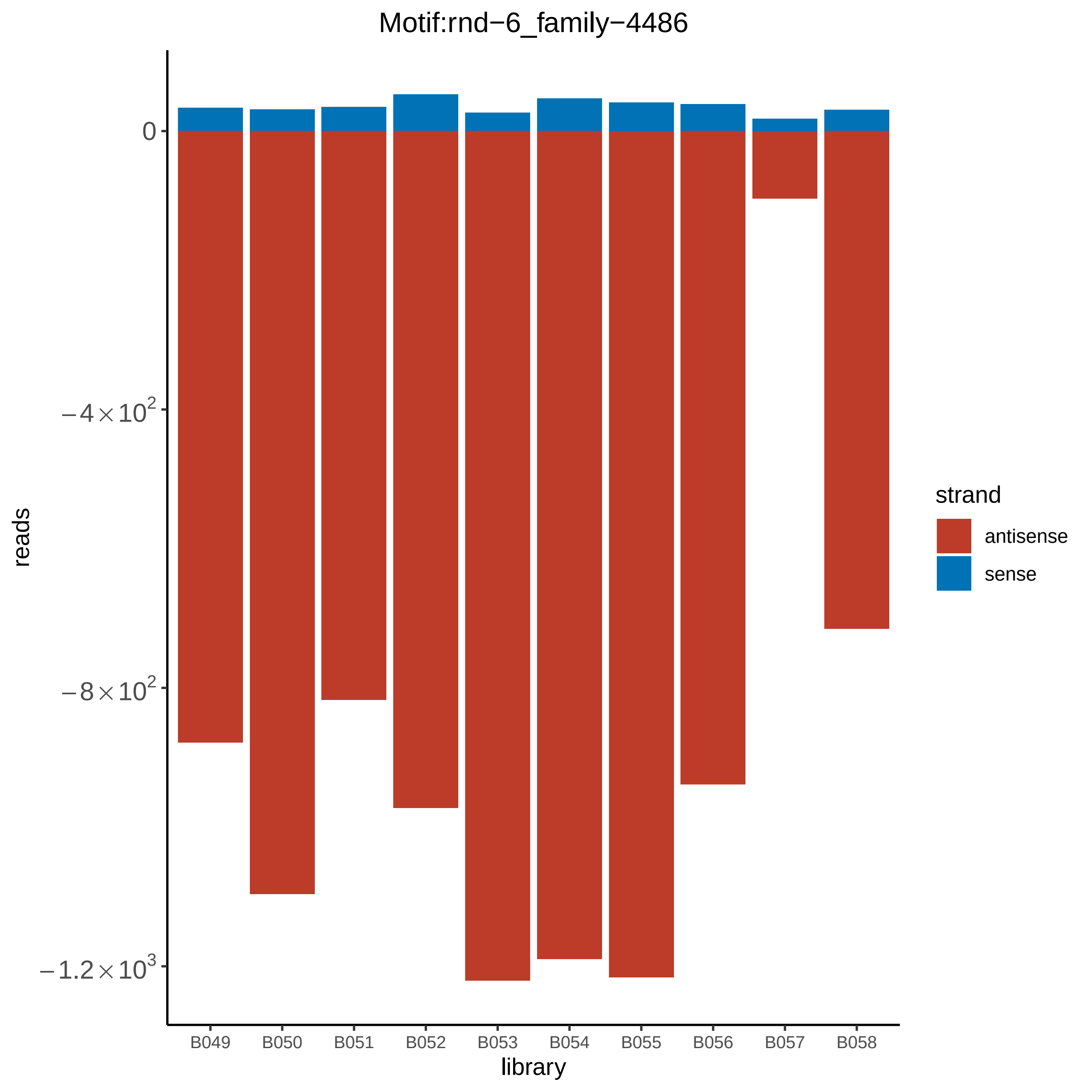

Motif:rnd−6_family−4486
0
−4×10^2
−8×10^2
−1.2×10^3
reads
B049
B050
B051
B052
B053
B054
B055
B056
B057
B058
library
strand
antisense
sense

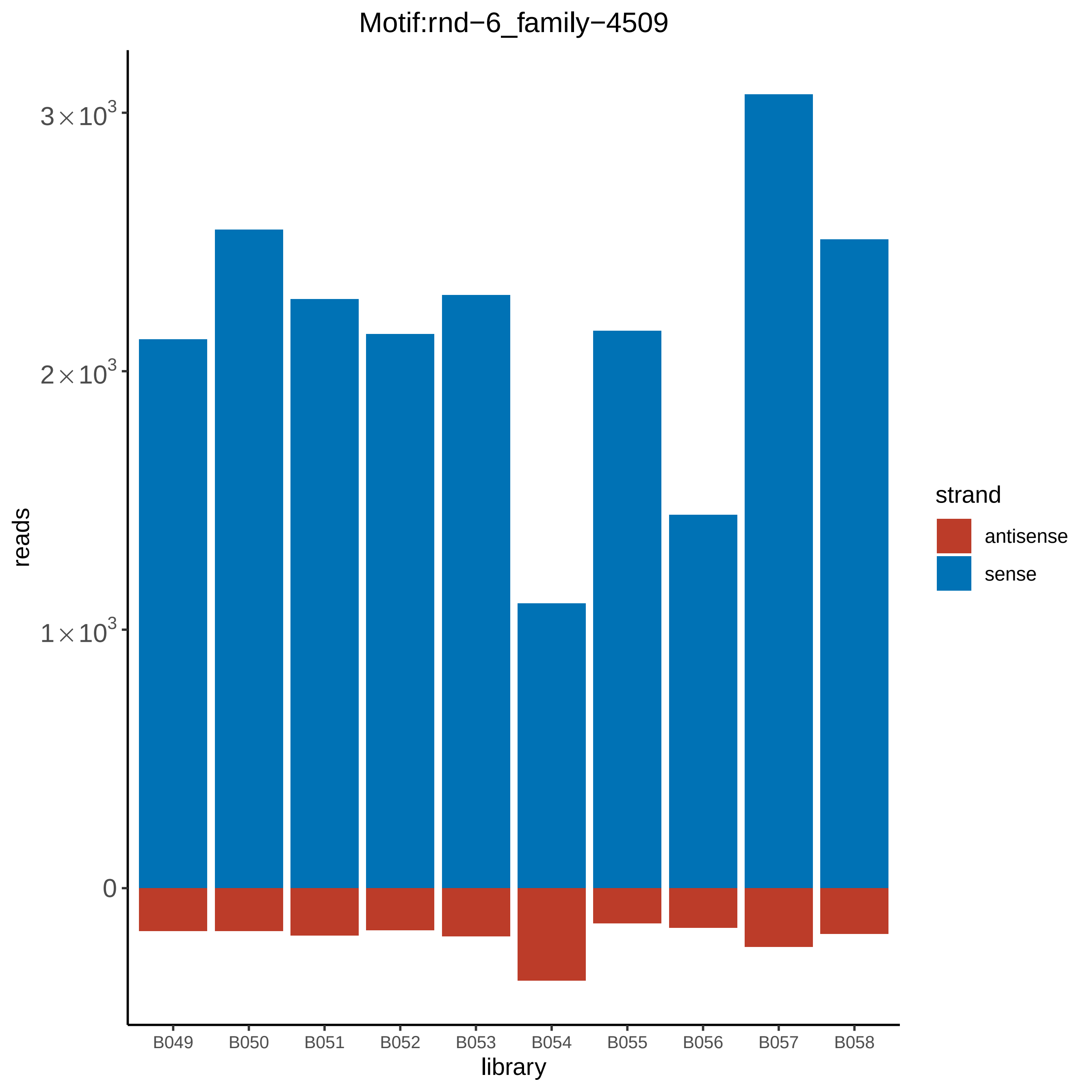

Motif:rnd−6_family−4509
reads
3×10³
2×10³
1×10³
0
B049
B050
B051
B052
B053
B054
B055
B056
B057
B058
library
strand
antisense
sense

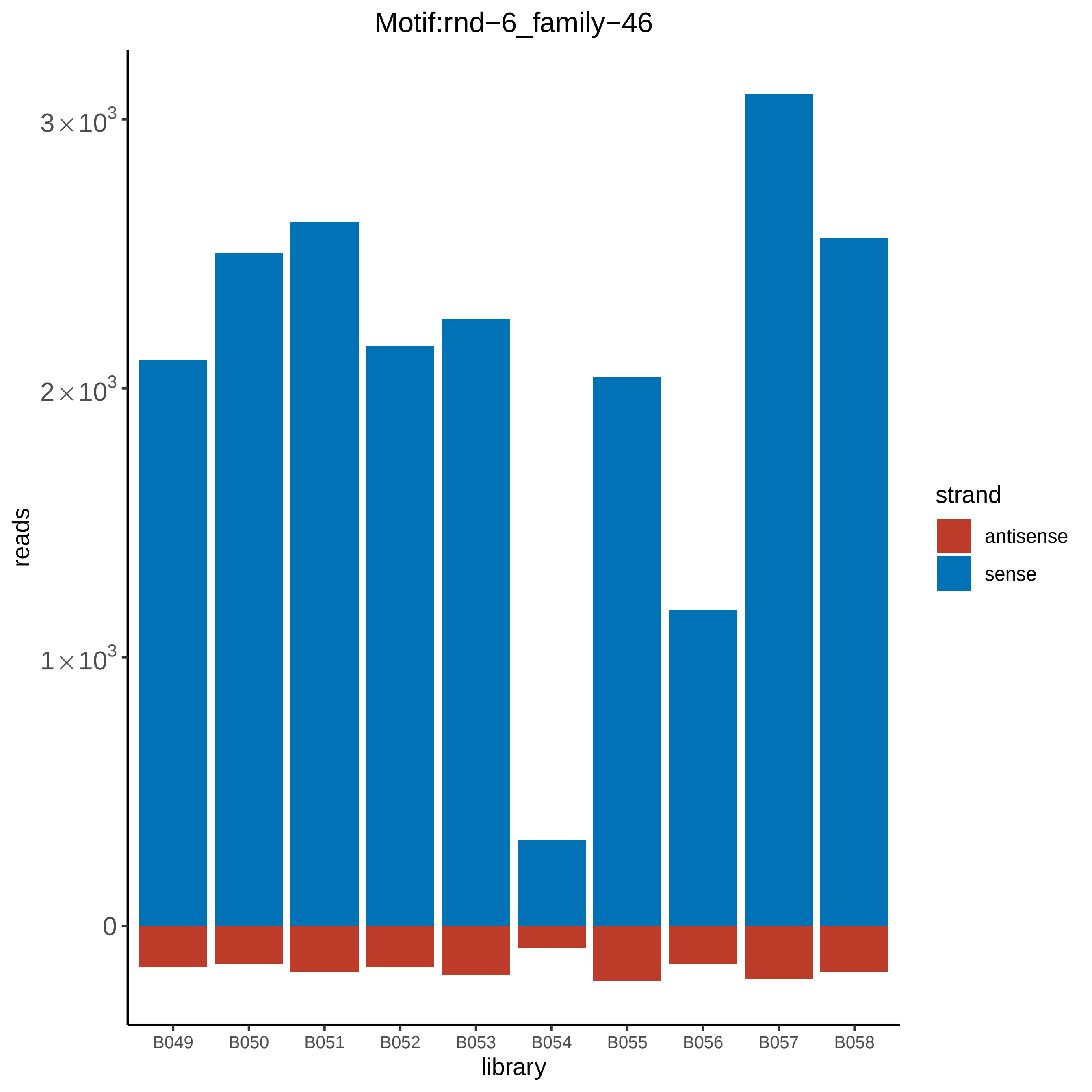

Motif:rnd−6_family−46
$3\times10^3$
$2\times10^3$
$1\times10^3$
0
reads
B049
B050
B051
B052
B053
B054
B055
B056
B057
B058
library
strand
antisense
sense

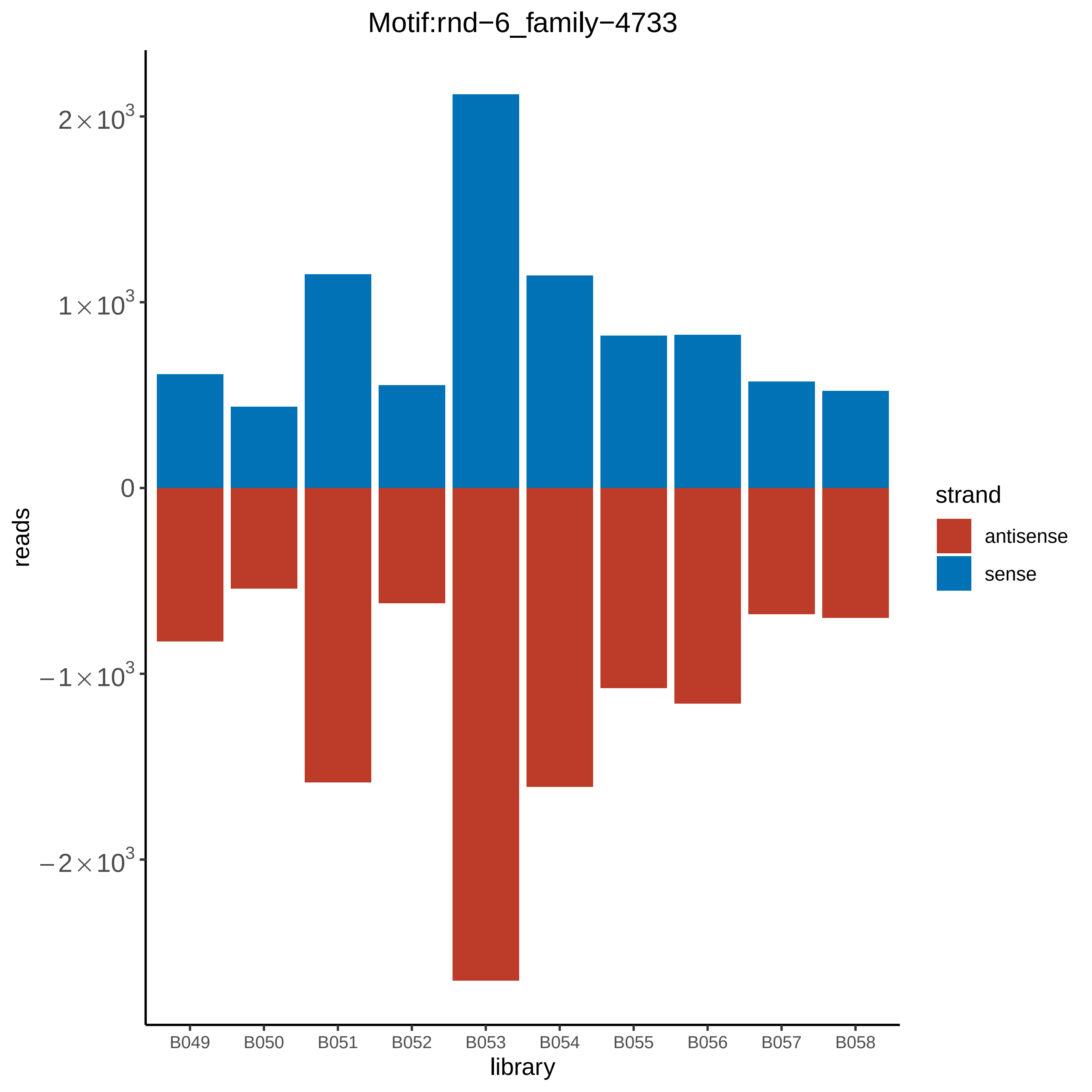

Motif:rnd−6_family−4733
reads
$2\times10^3$
$1\times10^3$
0
$-1\times10^3$
$-2\times10^3$
library
B049
B050
B051
B052
B053
B054
B055
B056
B057
B058
strand
antisense
sense

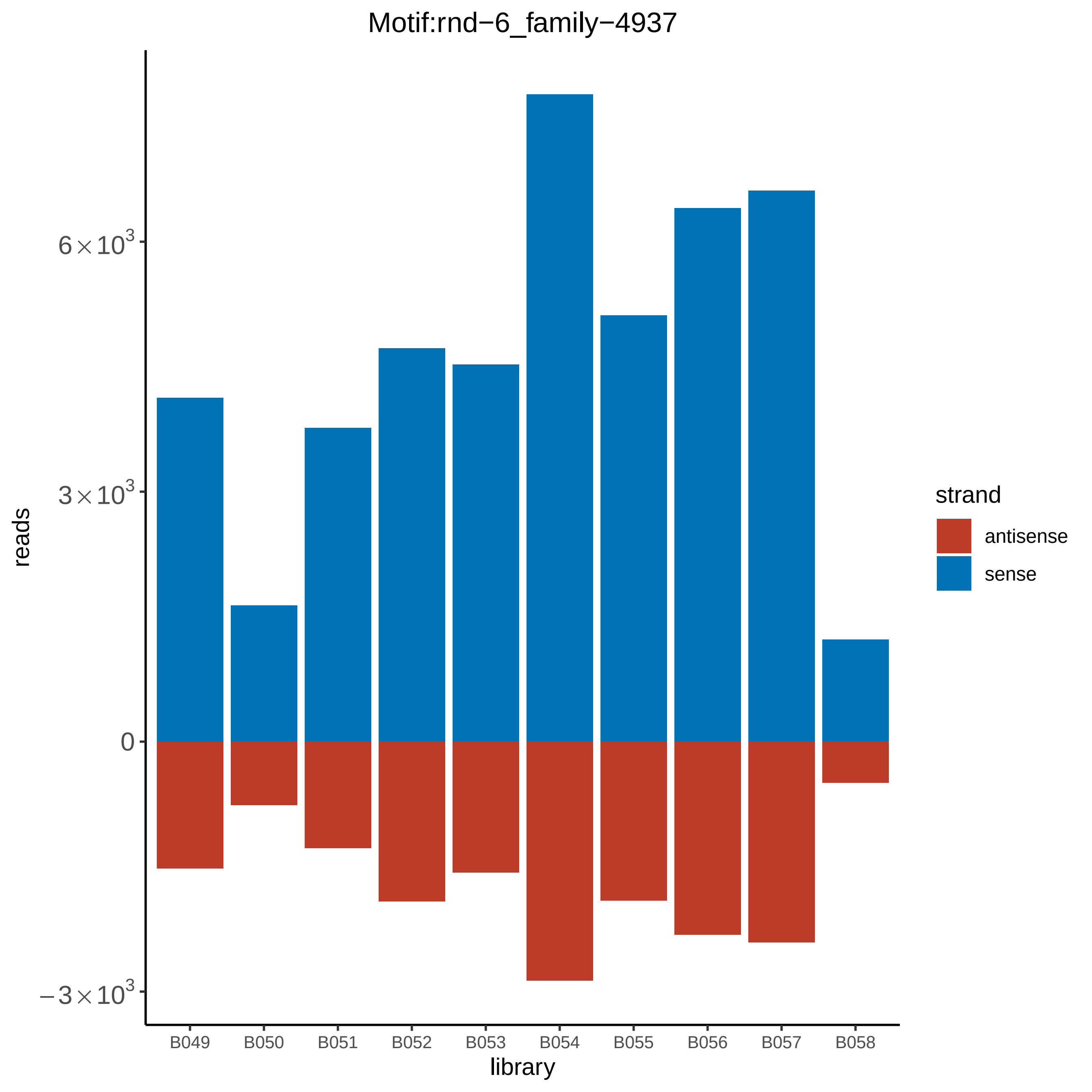
Motif:rnd−6_family−4937
reads
6×10^3
3×10^3
0
−3×10^3
library
B049
B050
B051
B052
B053
B054
B055
B056
B057
B058
strand
antisense
sense

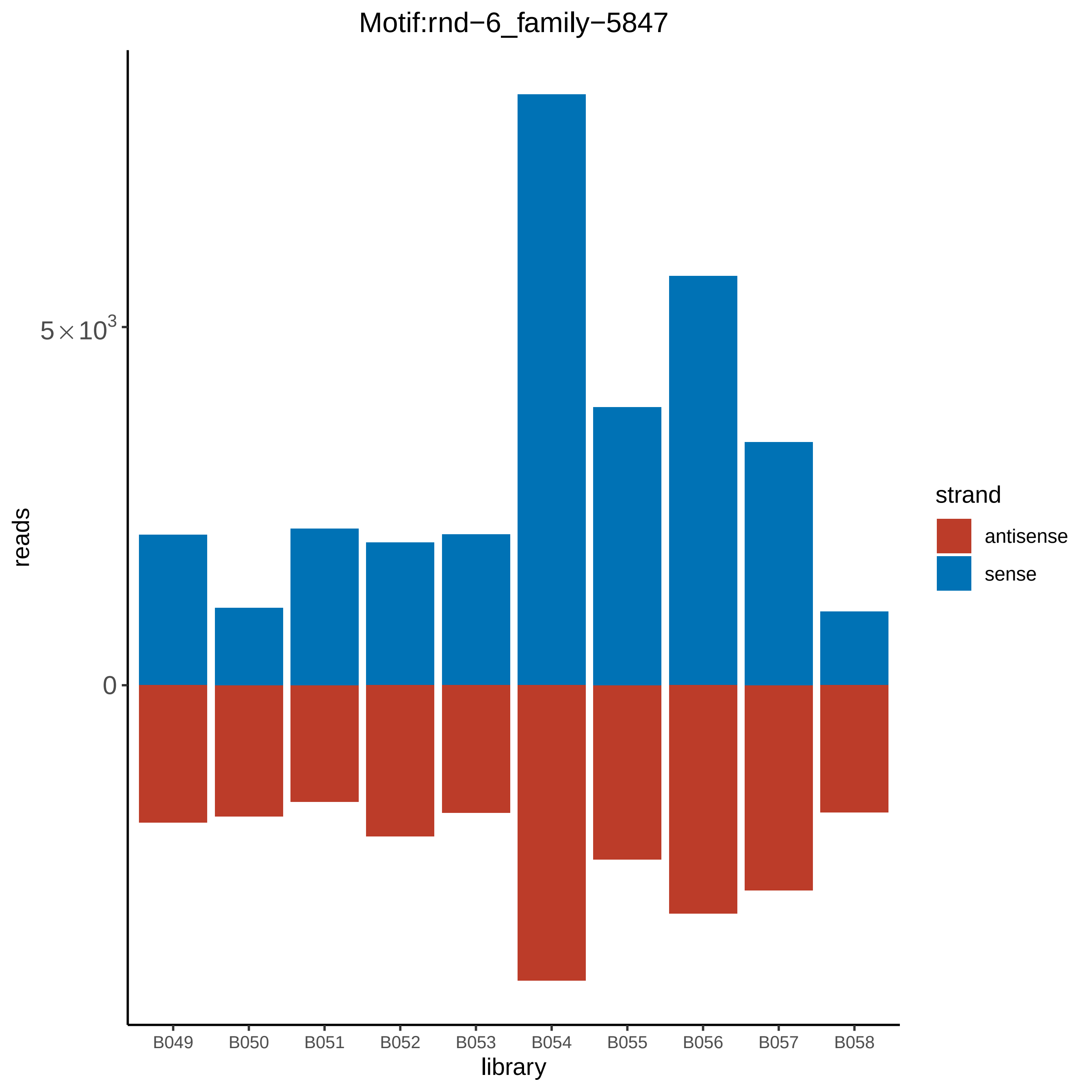

Motif:rnd−6_family−5847
reads
5×10³
0
library
B049
B050
B051
B052
B053
B054
B055
B056
B057
B058
strand
antisense
sense

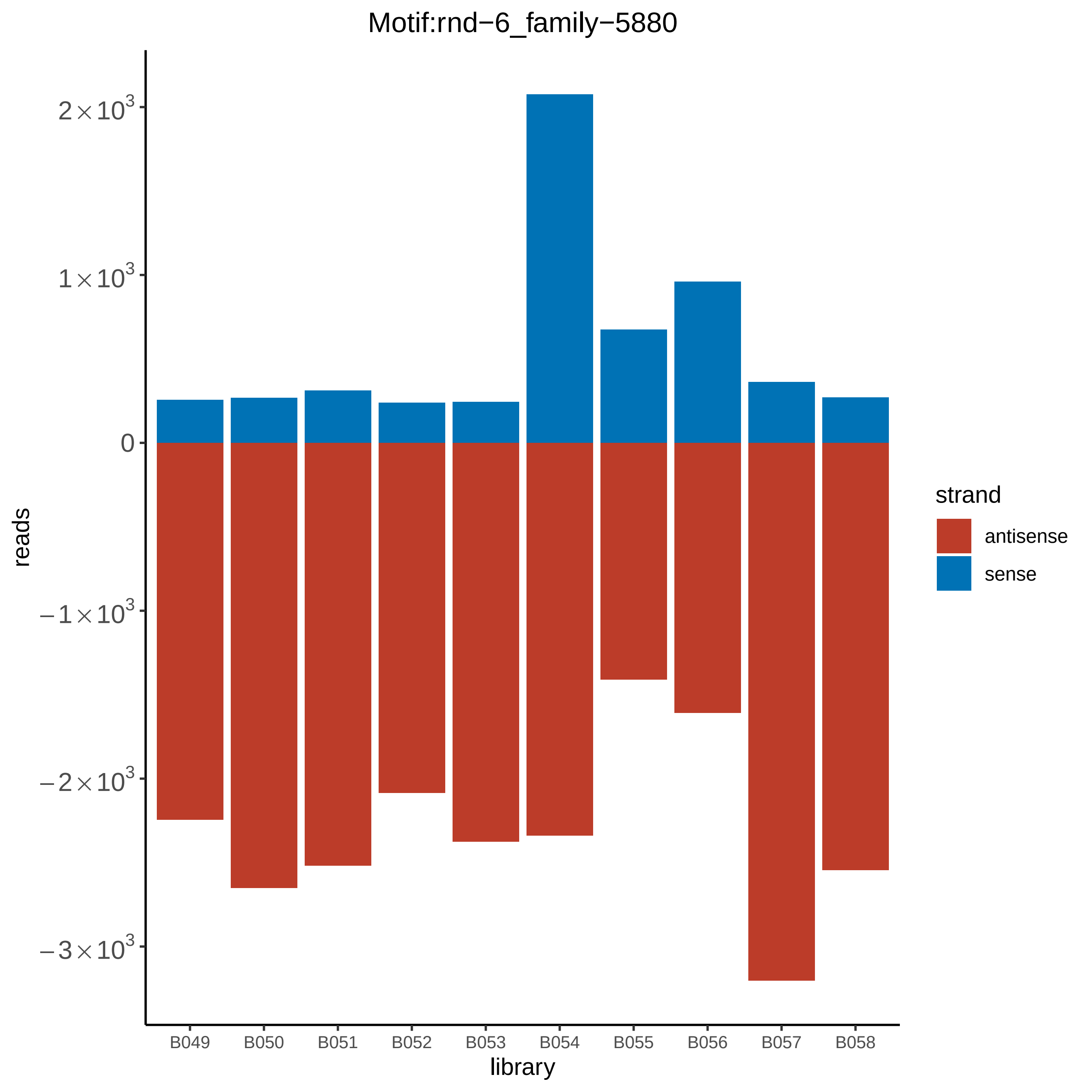
Motif:rnd−6_family−5880
reads
2×10^3
1×10^3
0
−1×10^3
−2×10^3
−3×10^3
library
B049
B050
B051
B052
B053
B054
B055
B056
B057
B058
strand
antisense
sense

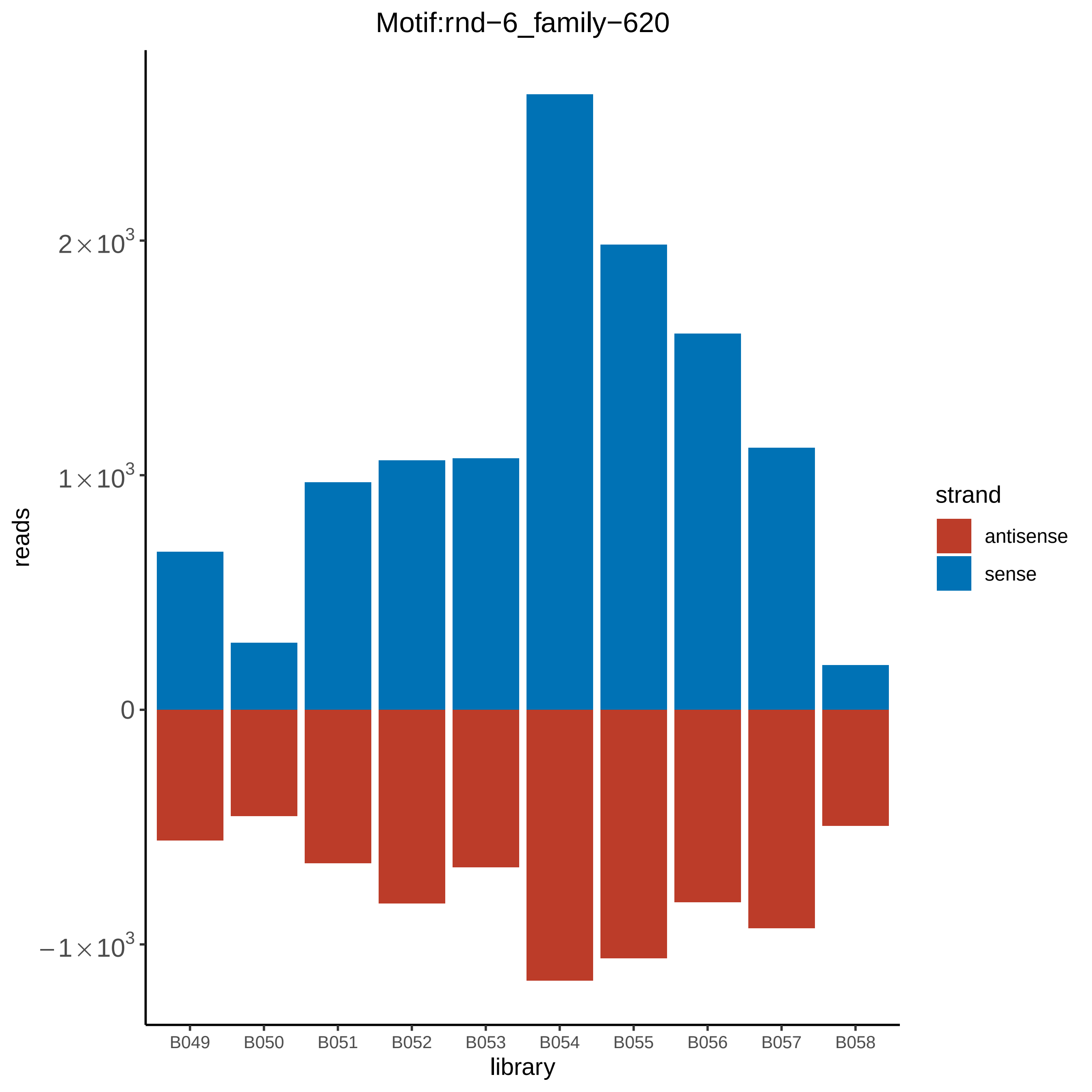

Motif:rnd−6_family−620
reads
$2\times10^3$
$1\times10^3$
0
$-1\times10^3$
B049
B050
B051
B052
B053
B054
B055
B056
B057
B058
library
strand
antisense
sense

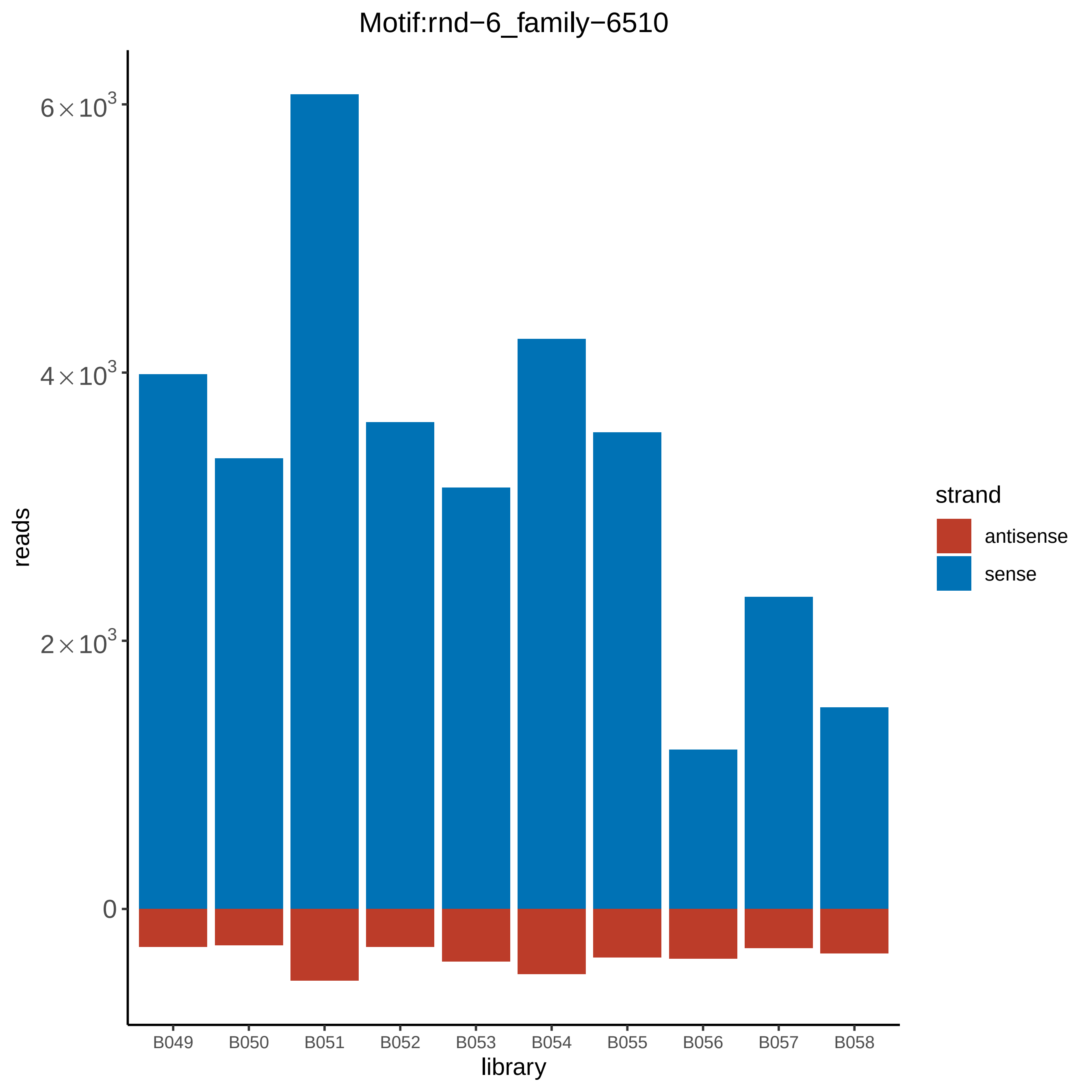

Motif:rnd−6_family−6510
reads
library
6×10^3
4×10^3
2×10^3
0
B049
B050
B051
B052
B053
B054
B055
B056
B057
B058
strand
antisense
sense

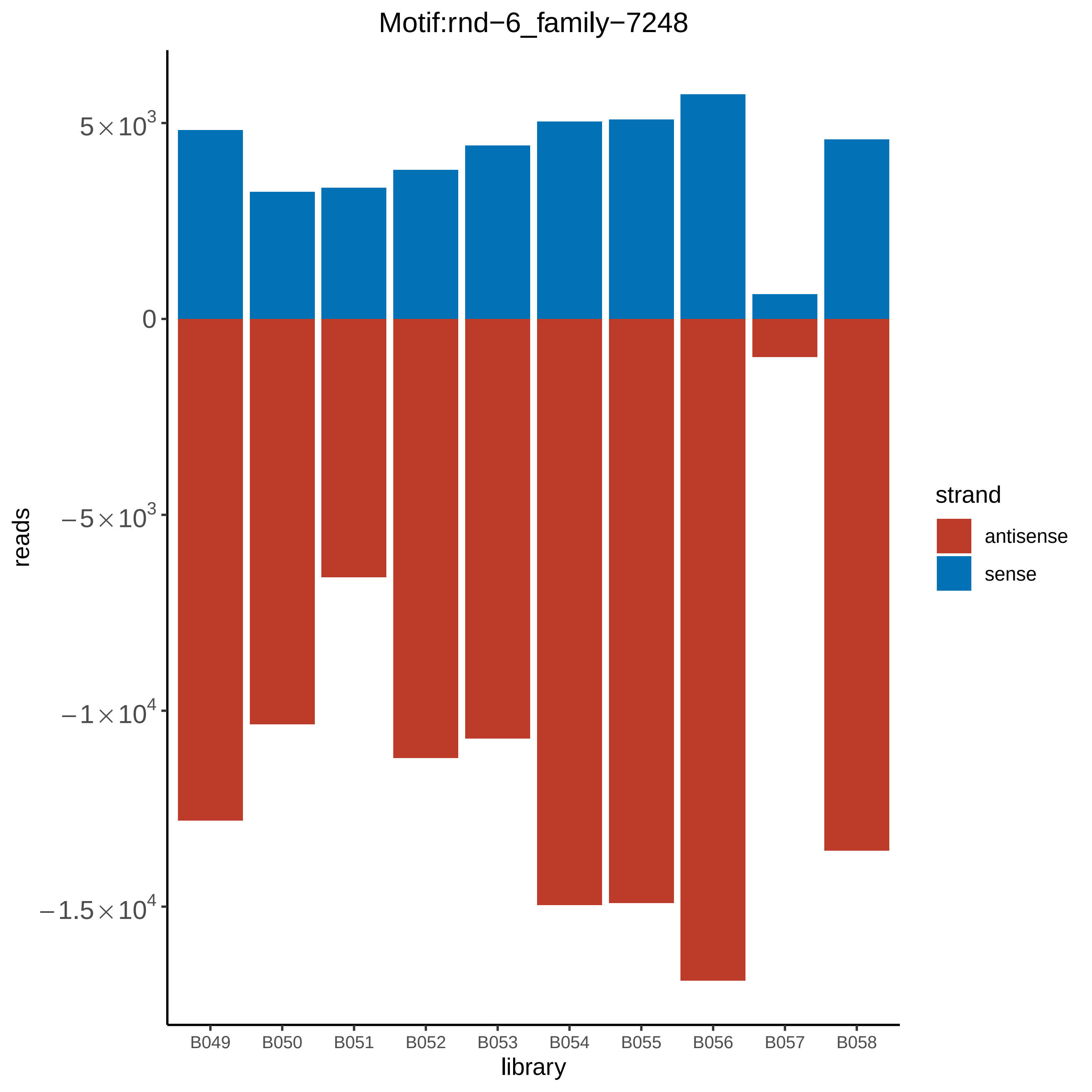
Motif:rnd−6_family−7248
reads
5×10^3
0
−5×10^3
−1×10^4
−1.5×10^4
B049
B050
B051
B052
B053
B054
B055
B056
B057
B058
library
strand
antisense
sense

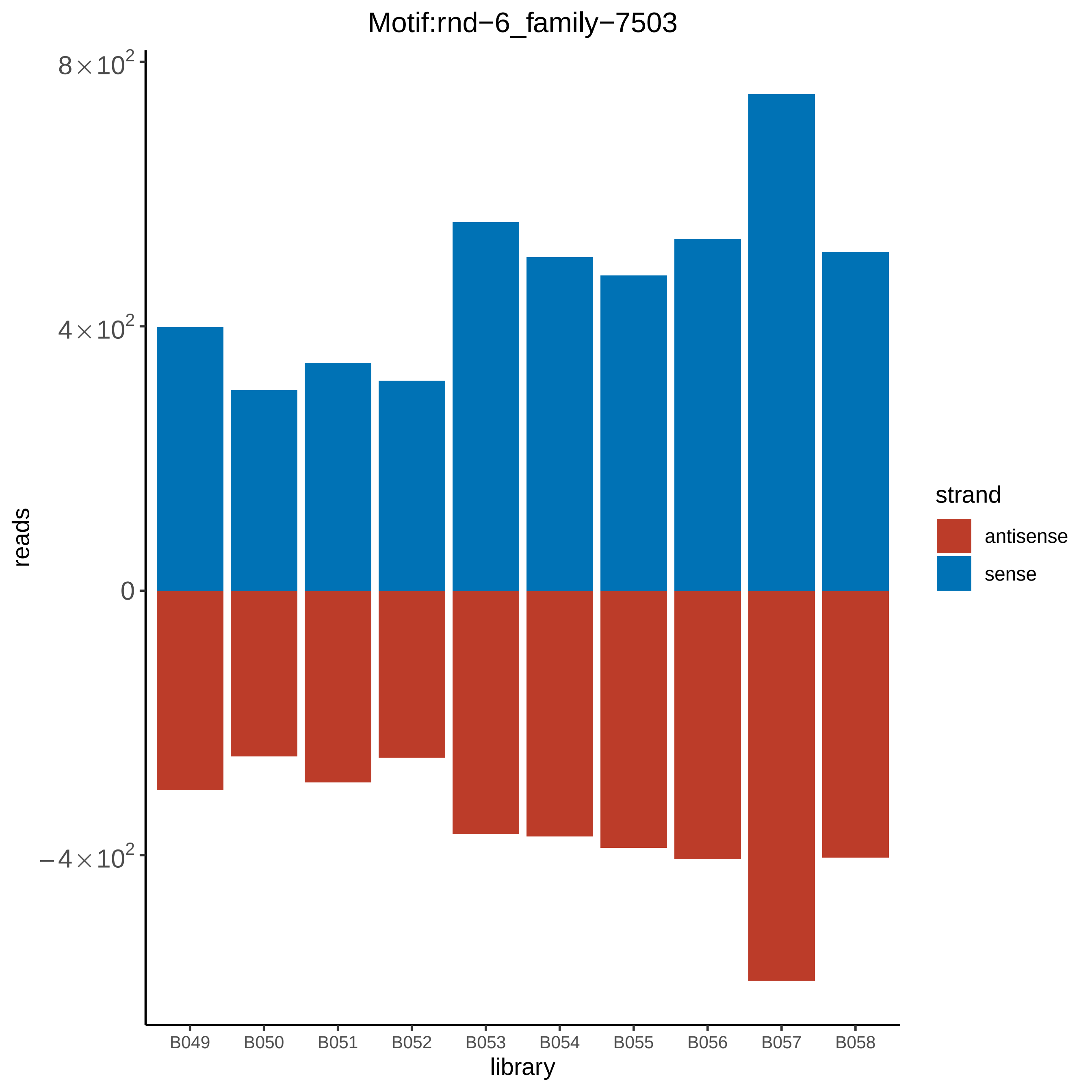

Motif:rnd−6_family−7503
8×10^2
4×10^2
0
−4×10^2
reads
B049
B050
B051
B052
B053
B054
B055
B056
B057
B058
library
strand
antisense
sense

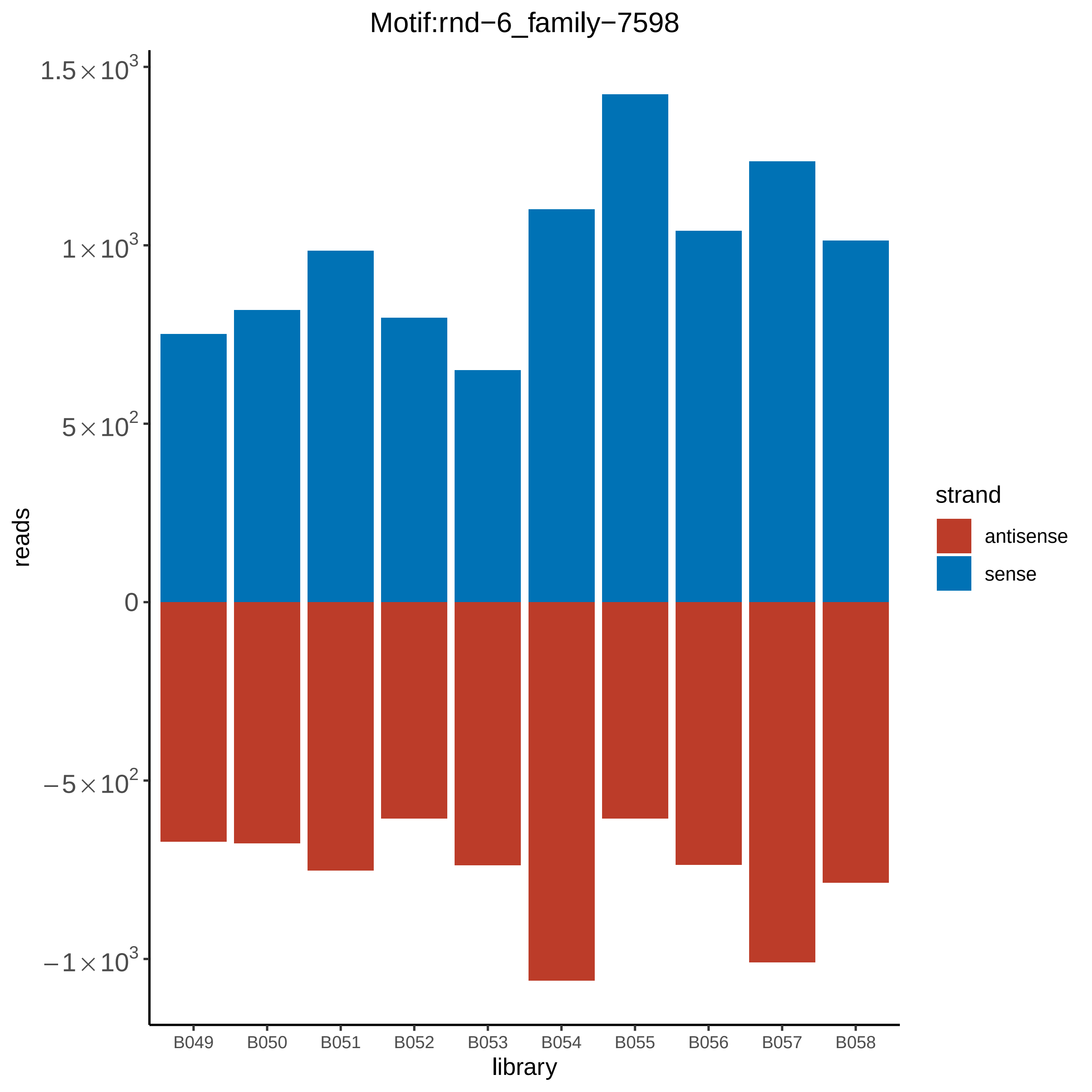

Motif:rnd−6_family−7598
1.5×10^3
1×10^3
5×10^2
0
−5×10^2
−1×10^3
reads
B049
B050
B051
B052
B053
B054
B055
B056
B057
B058
library
strand
antisense
sense

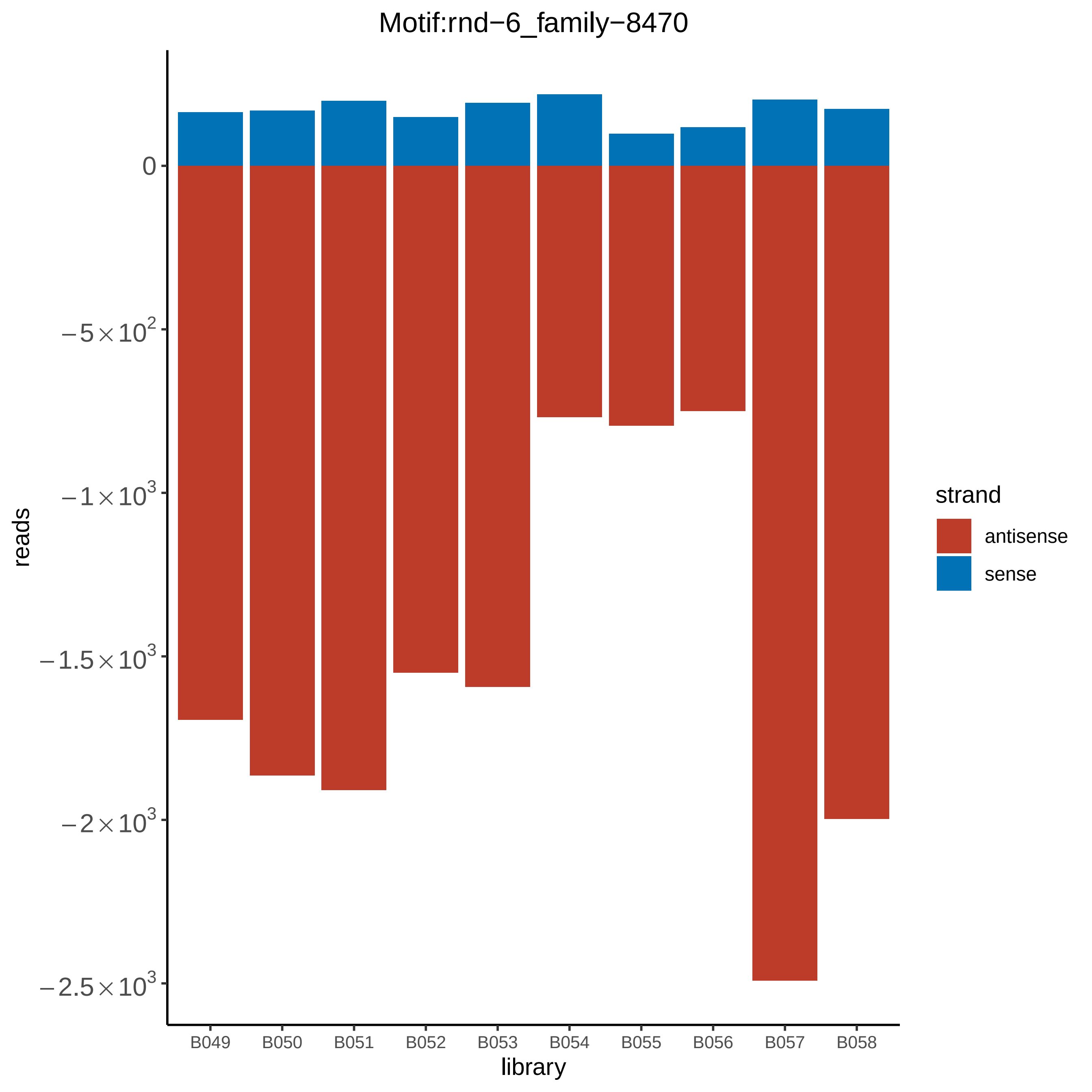
Motif:rnd−6_family−8470
0
−5×10^2
−1×10^3
−1.5×10^3
−2×10^3
−2.5×10^3
reads
B049
B050
B051
B052
B053
B054
B055
B056
B057
B058
library
strand
antisense
sense

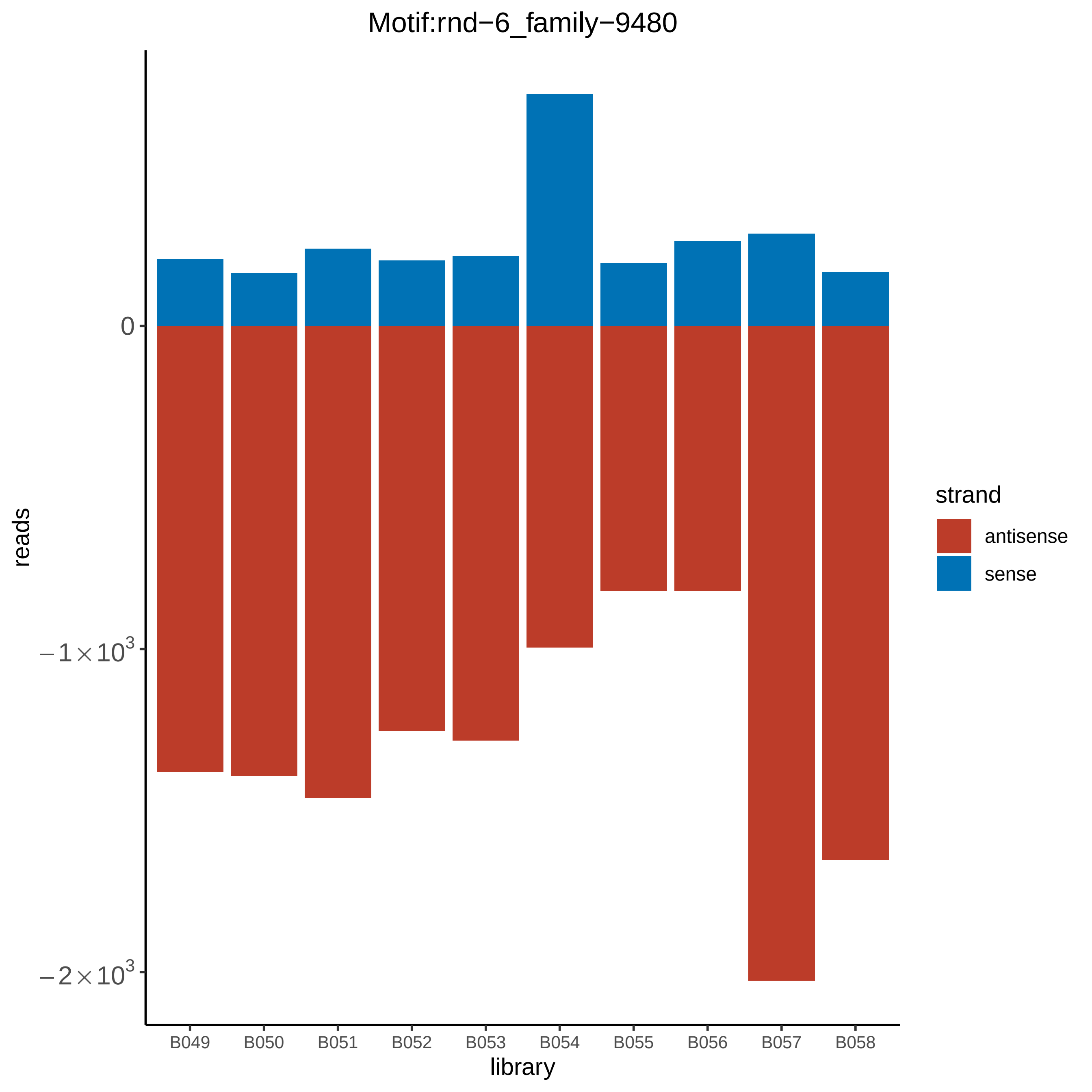

Motif:rnd−6_family−9480
0
−1×10³
−2×10³
reads
B049
B050
B051
B052
B053
B054
B055
B056
B057
B058
library
strand
antisense
sense

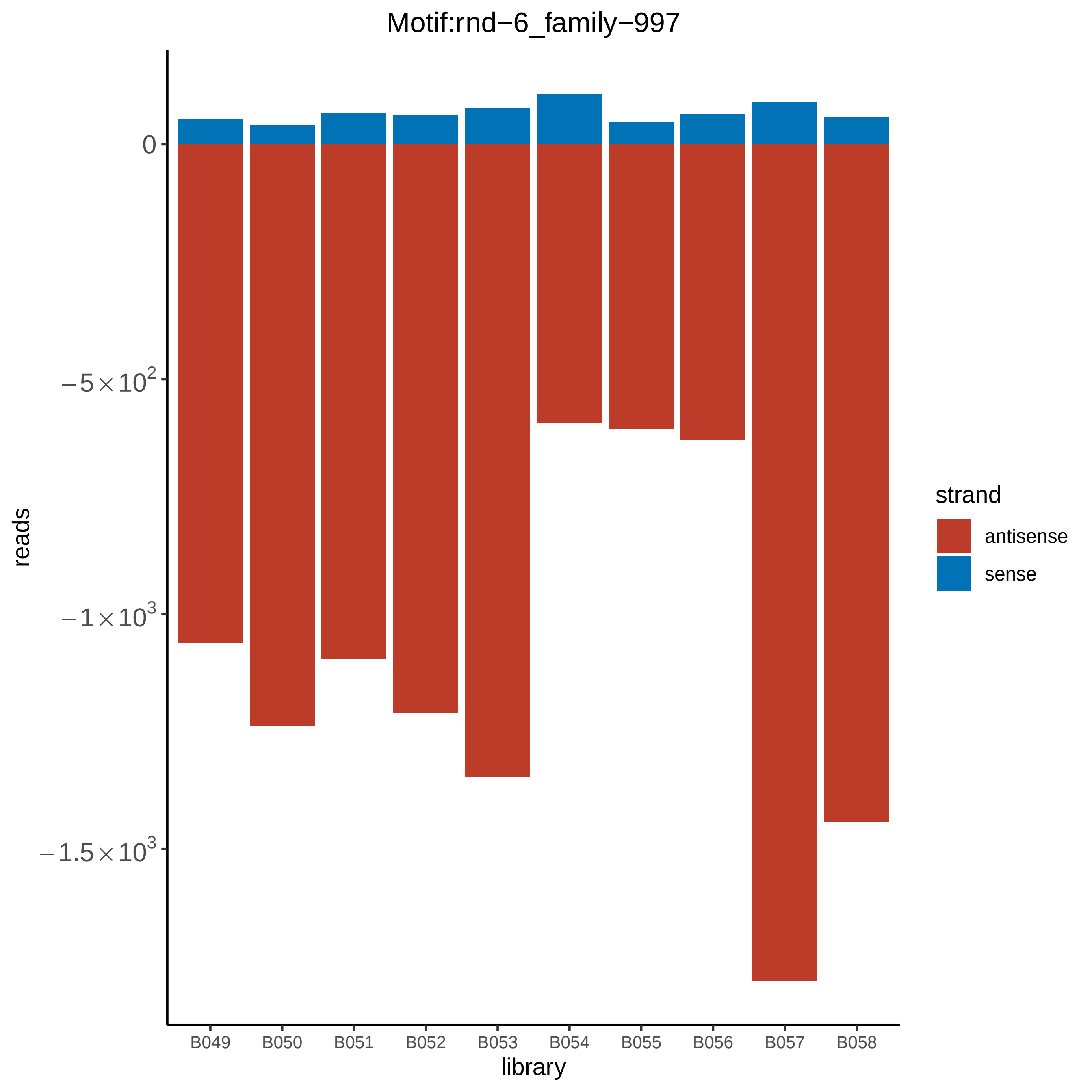
Motif:rnd−6_family−997
reads
0
$-5\times10^2$
$-1\times10^3$
$-1.5\times10^3$
B049
B050
B051
B052
B053
B054
B055
B056
B057
B058
library
strand
antisense
sense
