## Supplementary material for "A novel eukaryotic RdRP-dependent small RNA pathway represses antiviral immunity by controlling an ERK pathway component in the black-legged tick": TE_Relative.html

 


### TE mapping table (RPM)

| ID | GFP | Ago-16 | Ago-30 | Ago-96 | Ago-78 | Aub | AGO3-1 | AGO3-2 | RdRP1 | RdRP3 | SUM\_KD |
| --- | --- | --- | --- | --- | --- | --- | --- | --- | --- | --- | --- |
| Motif:rnd-6\_family-7248 | 1 | 0.77118734 | 0.56433645 | 0.85212062 | 0.85863725 | 1.13484692 | 1.13476291 | 1.28418229 | 0.09097595 | 1.03014828 | 153591.498 |
| Motif:rnd-5\_family-601 | 1 | 0.37053171 | 1.00937787 | 1.38400779 | 1.17568191 | 1.63990999 | 1.32354572 | 1.39968037 | 1.47066642 | 0.23788797 | 153461.943 |
| Motif:rnd-1\_family-272 | 1 | 0.68850832 | 0.78656663 | 0.87195181 | 0.94235489 | 1.18360773 | 1.21784753 | 1.31494006 | 0.10499472 | 1.0567465 | 126463.34 |
| Motif:rnd-1\_family-366 | 1 | 0.69955475 | 0.53898456 | 0.80420013 | 0.82157207 | 1.1113121 | 1.15590533 | 1.32083209 | 0.0955985 | 1.03213335 | 100424.958 |
| Motif:rnd-5\_family-4359 | 1 | 0.84642491 | 0.65920693 | 0.84922198 | 1.00631601 | 1.21608892 | 1.23624197 | 1.23431387 | 0.06398749 | 0.93495658 | 78092.1288 |
| Motif:rnd-1\_family-1035 | 1 | 0.68425899 | 0.5711696 | 0.7434604 | 0.74672116 | 1.12770998 | 1.13079553 | 1.25364629 | 0.08210148 | 0.99908395 | 76048.2511 |
| Motif:rnd-6\_family-4937 | 1 | 0.4247797 | 0.89346711 | 1.17530874 | 1.07978329 | 1.882234 | 1.24334494 | 1.54401137 | 1.5976964 | 0.30452539 | 62972.9124 |
| Motif:rnd-1\_family-1958 | 1 | 0.71064915 | 0.58357853 | 0.80034074 | 0.75139361 | 1.11564464 | 1.13935063 | 1.2701168 | 0.10175325 | 1.00873974 | 60550.1154 |
| Motif:rnd-1\_family-1103 | 1 | 0.73676538 | 0.77441168 | 0.8064254 | 0.93978148 | 1.31222232 | 1.18240509 | 1.2610864 | 0.07154975 | 0.92436842 | 57217.118 |
| Motif:rnd-6\_family-5847 | 1 | 0.72481387 | 0.94843108 | 1.02125089 | 0.96775538 | 3.07855705 | 1.57076899 | 2.21578277 | 1.55760502 | 0.69786753 | 55379.6384 |
| Motif:rnd-1\_family-1744 | 1 | 0.68263738 | 0.46568595 | 0.77582816 | 0.88466378 | 1.11013346 | 1.17852841 | 1.43573918 | 0.09811045 | 1.05862601 | 48674.782 |
| Motif:rnd-1\_family-178 | 1 | 0.53962622 | 1.13965728 | 0.94935234 | 0.90939574 | 1.6287102 | 1.35689002 | 1.61372295 | 1.63530765 | 0.60545668 | 44072.1643 |
| Motif:rnd-5\_family-5812 | 1 | 1.18085774 | 1.13730789 | 0.81971882 | 0.96845669 | 0.65167249 | 0.28307256 | 0.34241582 | 1.05599189 | 0.83347297 | 41979.4074 |
| Motif:rnd-6\_family-423 | 1 | 1.20435487 | 1.20276696 | 1.01378568 | 1.25265717 | 0.25265543 | 0.39364455 | 0.36217543 | 1.53785523 | 1.19304456 | 40138.5092 |
| Motif:rnd-4\_family-2208 | 1 | 1.18032745 | 1.2264092 | 1.04549692 | 1.10072047 | 0.20498453 | 1.14664025 | 0.64464294 | 1.48242445 | 1.25793513 | 38865.0954 |
| Motif:rnd-6\_family-6510 | 1 | 0.85054438 | 1.54716597 | 0.91626287 | 0.82807185 | 1.10921929 | 0.91725356 | 0.36536988 | 0.61381956 | 0.42963958 | 36633.8123 |
| Motif:rnd-6\_family-5880 | 1 | 1.16760284 | 1.13157326 | 0.92857715 | 1.04715286 | 1.76546175 | 0.83367569 | 1.0271807 | 1.42567645 | 1.12550138 | 28635.1917 |
| Motif:rnd-6\_family-1977 | 1 | 1.04988309 | 1.12790827 | 0.9898575 | 1.16219103 | 1.3674364 | 0.63007563 | 0.77106576 | 1.58063226 | 1.20921319 | 25310.7538 |
| Motif:rnd-6\_family-4509 | 1 | 1.18494725 | 1.07496118 | 1.00685644 | 1.08336424 | 0.63735498 | 1.00079187 | 0.69771344 | 1.43989107 | 1.17305665 | 23585.0729 |
| Motif:rnd-5\_family-460 | 1 | 1.14895088 | 1.19544709 | 1.018229 | 1.35481762 | 0.61904037 | 0.54228106 | 0.53185638 | 1.56727121 | 1.17285745 | 23438.1211 |
| Motif:rnd-6\_family-46 | 1 | 1.1700818 | 1.23363153 | 1.02171868 | 1.08023758 | 0.17770832 | 0.99285048 | 0.58302777 | 1.45489291 | 1.20713332 | 22415.9714 |
| Motif:rnd-6\_family-44 | 1 | 1.15843998 | 1.23145756 | 1.02126563 | 1.06059813 | 0.19164538 | 1.10111724 | 0.61619323 | 1.43128035 | 1.20768933 | 21579.2851 |
| Motif:rnd-6\_family-3340 | 1 | 0.8071999 | 0.85396446 | 0.81593934 | 0.78011023 | 1.16132432 | 1.14708415 | 1.25371829 | 0.19108305 | 0.96969693 | 21507.0938 |
| Motif:rnd-1\_family-474 | 1 | 1.14818128 | 1.22309006 | 1.00415744 | 1.06728027 | 0.18297077 | 1.1193614 | 0.64652452 | 1.4537855 | 1.22810306 | 20871.1285 |
| Motif:rnd-6\_family-4733 | 1 | 0.68095142 | 1.9038927 | 0.81674658 | 3.32045791 | 1.91544801 | 1.32079875 | 1.38212175 | 0.87113788 | 0.85061618 | 20200.2007 |
| Motif:rnd-1\_family-423 | 1 | 1.32945687 | 1.08916315 | 0.97529546 | 1.1961366 | 0.43276908 | 0.34407561 | 0.34757915 | 1.44537726 | 1.07568964 | 19906.4641 |
| Motif:rnd-6\_family-2886 | 1 | 1.05279443 | 1.06103543 | 0.80997523 | 0.8494298 | 0.42543823 | 0.85179777 | 0.63932923 | 1.57430376 | 1.21042208 | 19472.5218 |
| Motif:rnd-4\_family-408 | 1 | 0.77297686 | 0.65438022 | 0.89093085 | 0.91770952 | 1.20542538 | 1.24021604 | 1.37312879 | 0.11212536 | 1.07899277 | 19377.6022 |
| Motif:rnd-6\_family-620 | 1 | 0.6007113 | 1.31933973 | 1.53512748 | 1.41784861 | 3.07115992 | 2.47234675 | 1.97011465 | 1.66516187 | 0.55654021 | 19195.9023 |
| Motif:rnd-6\_family-7598 | 1 | 1.05100292 | 1.22175983 | 0.98683865 | 0.97518377 | 1.52050833 | 1.42750606 | 1.24927114 | 1.57865086 | 1.26488128 | 17451.5197 |
| Motif:rnd-6\_family-4398 | 1 | 1.09365053 | 1.03852179 | 0.93772605 | 1.23567092 | 2.73584565 | 0.94537543 | 1.34570952 | 1.64773514 | 1.23096443 | 17332.5085 |
| Motif:rnd-6\_family-8470 | 1 | 1.09442117 | 1.13429919 | 0.91417099 | 0.96168386 | 0.53133894 | 0.48063223 | 0.46701415 | 1.45025048 | 1.16886031 | 17095.9871 |
| Motif:rnd-1\_family-1822 | 1 | 1.0199965 | 0.98434617 | 0.97813743 | 1.08723041 | 0.44344487 | 0.52583346 | 0.61059 | 1.44568075 | 0.86793748 | 16711.3995 |
| Motif:rnd-4\_family-2384 | 1 | 1.30034409 | 1.29013291 | 1.09711116 | 1.40929755 | 0.16663322 | 0.28643214 | 0.24566397 | 1.40090664 | 1.21648203 | 16622.4017 |
| Motif:rnd-6\_family-4430 | 1 | 1.263204 | 1.10225703 | 0.94346389 | 0.77531831 | 0.49802359 | 0.4341356 | 0.37938666 | 1.09413781 | 0.90149974 | 16564.8222 |
| Motif:rnd-1\_family-926 | 1 | 1.12175313 | 1.03757927 | 0.94634004 | 0.84953722 | 0.520073 | 0.45146829 | 0.48222889 | 1.523038 | 1.20291132 | 15943.8976 |
| Motif:rnd-6\_family-9480 | 1 | 0.98103601 | 1.07163232 | 0.91825483 | 0.94501696 | 1.07863492 | 0.63996659 | 0.68290593 | 1.45716657 | 1.14657836 | 15739.927 |
| Motif:rnd-6\_family-1478 | 1 | 1.11224986 | 1.07661945 | 0.93108481 | 1.11522707 | 0.7266859 | 0.41707471 | 0.4345785 | 1.42752118 | 1.1215866 | 14720.3087 |
| Motif:rnd-4\_family-513 | 1 | 0.83735041 | 1.07788617 | 0.72010237 | 0.43299724 | 0.68878533 | 0.81087798 | 0.73388916 | 0.07335264 | 0.62269063 | 14623.2492 |
| Motif:rnd-1\_family-1320 | 1 | 1.24935576 | 1.08110919 | 0.92312713 | 0.7870743 | 1.00987692 | 0.43247083 | 0.53376671 | 1.16175277 | 0.88965811 | 14002.9605 |
| Motif:rnd-4\_family-328 | 1 | 1.07274283 | 1.01527763 | 1.02110213 | 0.96925668 | 0.91240664 | 0.869461 | 0.82236846 | 0.92076703 | 1.0851859 | 13207.2331 |
| Motif:rnd-1\_family-417 | 1 | 1.20197807 | 1.2525623 | 1.04188576 | 1.09063073 | 0.23160144 | 0.95991353 | 0.58095204 | 1.45209301 | 1.20651061 | 12675.2571 |
| Motif:rnd-5\_family-631 | 1 | 1.1408859 | 1.07193823 | 0.9195714 | 0.93085161 | 0.40432951 | 0.45734115 | 0.43992207 | 0.9617947 | 0.7327673 | 12535.4831 |
| Motif:rnd-5\_family-2803 | 1 | 0.84580104 | 0.92287115 | 1.13666029 | 0.7771555 | 2.56600361 | 1.41777554 | 2.42946647 | 1.54874025 | 0.8812025 | 12288.7451 |
| Motif:rnd-6\_family-997 | 1 | 1.14602728 | 1.04199074 | 1.13970322 | 1.27451333 | 0.6270823 | 0.58461746 | 0.62203503 | 1.67512481 | 1.34318346 | 11672.4484 |
| Motif:rnd-6\_family-2652 | 1 | 1.18741355 | 1.16012791 | 1.01862438 | 1.08395168 | 0.2019106 | 0.20256553 | 0.19890974 | 1.39372684 | 1.13176357 | 11612.3979 |
| Motif:rnd-6\_family-1271 | 1 | 1.32301142 | 1.19456949 | 1.06843956 | 1.03257153 | 0.22284496 | 0.54288037 | 0.42264019 | 1.49070959 | 1.24585204 | 11537.0567 |
| Motif:rnd-6\_family-1406 | 1 | 0.99299737 | 1.02358982 | 0.91668786 | 0.89358202 | 1.24362572 | 0.58116152 | 0.77085411 | 1.55804866 | 1.20097503 | 11386.8991 |
| Motif:rnd-6\_family-2888 | 1 | 1.17374424 | 1.15901116 | 0.82223687 | 0.86396726 | 0.11380662 | 0.21432691 | 0.19116322 | 1.32060103 | 1.07002314 | 10674.4716 |
| Motif:rnd-1\_family-528 | 1 | 1.00700258 | 1.06313132 | 0.98487785 | 1.04085649 | 0.74612654 | 0.55600687 | 0.56961038 | 1.42046811 | 1.08527768 | 10190.0693 |
| Motif:rnd-5\_family-37 | 1 | 0.97575144 | 0.84385029 | 1.07032257 | 0.99519324 | 1.40951129 | 1.31266287 | 1.30720288 | 0.25879578 | 1.10830732 | 10008.9093 |
| Motif:rnd-6\_family-4238 | 1 | 0.89859296 | 0.94240577 | 1.03371076 | 1.07705204 | 1.2947083 | 1.43305212 | 1.35667163 | 0.45557094 | 1.22148954 | 9768.60779 |
| Motif:rnd-1\_family-36 | 1 | 1.16384511 | 1.14096622 | 1.04506442 | 1.01626015 | 0.81085779 | 0.58597592 | 0.62726617 | 1.47941247 | 1.17555328 | 9755.13386 |
| Motif:rnd-5\_family-3305 | 1 | 1.23042207 | 0.97175428 | 1.13824145 | 1.17657303 | 0.27417106 | 0.43354966 | 0.46066474 | 1.32913918 | 1.13264076 | 9688.28421 |
| Motif:rnd-5\_family-2342 | 1 | 0.98573285 | 0.99362644 | 0.91016313 | 1.03150043 | 0.42816814 | 0.55002499 | 0.50445239 | 1.44098995 | 1.03986982 | 9673.03897 |
| Motif:rnd-6\_family-4486 | 1 | 1.23606179 | 0.93421379 | 1.12397636 | 1.36721041 | 1.35556811 | 1.37805429 | 1.07187186 | 0.12573634 | 0.8178026 | 9496.07231 |
| Motif:rnd-5\_family-154 | 1 | 1.04040195 | 1.12345948 | 0.94262064 | 0.97584646 | 0.98929136 | 0.49947131 | 0.52436499 | 1.3122379 | 1.07757114 | 9490.57029 |
| Motif:rnd-5\_family-3344 | 1 | 1.19300448 | 0.98110207 | 0.96435535 | 1.0120858 | 0.71456555 | 0.4311968 | 0.4784194 | 1.40049022 | 1.22970733 | 9119.78412 |
| Motif:rnd-1\_family-873 | 1 | 0.88009819 | 1.06248555 | 0.9114809 | 1.0075651 | 1.42715745 | 1.16191671 | 1.35624729 | 1.74807308 | 1.2485925 | 8603.60733 |
| Motif:rnd-6\_family-3084 | 1 | 1.26060436 | 1.28672846 | 1.03251789 | 1.04824512 | 0.22523827 | 0.2920717 | 0.26417239 | 1.44626676 | 1.17003588 | 8449.88057 |
| Motif:rnd-1\_family-412 | 1 | 0.54986469 | 1.05370882 | 0.77501469 | 1.08867179 | 0.67435607 | 0.64222568 | 0.82824576 | 1.14554418 | 0.7221134 | 8404.17454 |
| Motif:rnd-6\_family-7503 | 1 | 0.79152279 | 0.90625092 | 0.81349513 | 1.32110137 | 1.25104832 | 1.23557532 | 1.33869575 | 1.91406432 | 1.30701734 | 8317.90141 |
| Motif:rnd-1\_family-136 | 1 | 0.81011626 | 0.56334901 | 0.87116305 | 0.91941304 | 1.17160897 | 1.14555461 | 1.28420823 | 0.12320911 | 0.99327301 | 8259.5134 |
| Motif:rnd-1\_family-9 | 1 | 1.10816339 | 1.06636876 | 1.03122789 | 0.99014488 | 0.84480684 | 0.7420773 | 0.67628712 | 1.19161925 | 1.13566709 | 8232.88782 |
| Motif:rnd-5\_family-494 | 1 | 0.96259624 | 1.20600328 | 0.94133054 | 1.22450246 | 0.80810611 | 0.79400209 | 0.83441425 | 1.4932508 | 1.02826825 | 8083.26715 |
| Motif:rnd-5\_family-2195 | 1 | 0.95232306 | 1.12960303 | 0.86371896 | 1.04266038 | 0.76446885 | 1.23991297 | 0.89019525 | 1.55997568 | 1.16935736 | 8008.32242 |
| Motif:rnd-1\_family-201 | 1 | 0.82499767 | 0.96743769 | 1.02898941 | 0.91306008 | 1.75568236 | 1.13082203 | 1.28021849 | 0.33294971 | 0.95252382 | 8002.91762 |
| Motif:rnd-6\_family-1239 | 1 | 1.30273914 | 1.23365535 | 0.85531817 | 1.31992241 | 0.41981922 | 0.16647078 | 0.19974387 | 0.47425628 | 0.38934535 | 7781.40252 |
| Motif:rnd-6\_family-8629 | 1 | 1.09819257 | 1.03131769 | 0.93283634 | 1.02543354 | 0.75245543 | 0.6120261 | 0.63865321 | 1.3518384 | 1.09017987 | 7641.49611 |
| Motif:rnd-1\_family-697 | 1 | 1.24419243 | 1.09992572 | 0.98924331 | 1.35483654 | 1.01197141 | 0.5638654 | 0.62379884 | 1.41068846 | 1.14098622 | 7491.13788 |
| Motif:rnd-6\_family-159 | 1 | 1.09959447 | 1.10621658 | 0.9743905 | 1.01746108 | 1.49831396 | 0.70457009 | 0.863241 | 1.47957389 | 1.19201472 | 7444.40865 |
| Motif:rnd-6\_family-2254 | 1 | 1.1577322 | 1.14452025 | 0.93340562 | 1.19436388 | 0.20261368 | 0.42479748 | 0.33352946 | 1.22575243 | 1.05088851 | 7411.8439 |
| Motif:rnd-4\_family-1108 | 1 | 0.84562316 | 0.72703856 | 0.86670565 | 0.95467495 | 0.99686179 | 1.02254173 | 1.15129215 | 0.42300967 | 0.99081516 | 7353.9048 |
| Motif:rnd-1\_family-1829 | 1 | 0.88544882 | 0.99563222 | 1.10147782 | 1.0177507 | 0.59894132 | 0.49289538 | 0.59113355 | 1.56507711 | 0.99331087 | 7330.24797 |
| Motif:rnd-6\_family-2808 | 1 | 1.06424995 | 1.09288194 | 0.94881654 | 1.04724699 | 0.52698913 | 0.61704158 | 0.59362499 | 1.46485359 | 1.16174351 | 7250.06612 |
| Motif:rnd-3\_family-43 | 1 | 0.7221259 | 0.56779765 | 0.70995849 | 0.75012448 | 1.04369079 | 1.11058661 | 1.3138319 | 0.08742687 | 0.98035698 | 7233.29962 |
| Motif:rnd-6\_family-970 | 1 | 0.96625698 | 0.94270171 | 0.96441375 | 0.93339447 | 1.21215309 | 1.07147652 | 1.19700337 | 0.78288168 | 1.17528205 | 7162.60073 |
| Motif:rnd-6\_family-10566 | 1 | 1.1771195 | 1.13615886 | 0.97993627 | 1.20001186 | 0.47422951 | 0.46774581 | 0.48133187 | 1.24990659 | 0.97527467 | 6934.4708 |
| Motif:rnd-6\_family-10254 | 1 | 1.02912696 | 1.07899334 | 0.91119527 | 1.12563203 | 0.5613162 | 0.56000932 | 0.55423434 | 1.31628136 | 1.0219854 | 6774.60508 |
| Motif:rnd-5\_family-557 | 1 | 1.18716512 | 1.07234928 | 1.11404271 | 1.0060065 | 0.35685758 | 0.37594604 | 0.35398159 | 1.43486602 | 1.15109972 | 6733.61198 |
| Motif:rnd-1\_family-81 | 1 | 0.81096033 | 0.97094531 | 0.97563387 | 1.0432237 | 1.33691002 | 1.45402543 | 1.30677222 | 1.79987899 | 1.27411081 | 6651.34287 |
| Motif:rnd-1\_family-311 | 1 | 1.0805068 | 1.29314655 | 1.36203165 | 1.29469565 | 1.75495719 | 1.57251319 | 1.35438171 | 0.2515749 | 1.15670366 | 6589.29971 |
| Motif:rnd-6\_family-4387 | 1 | 0.68725541 | 1.0365592 | 0.97196825 | 1.00929638 | 2.45362853 | 1.35115797 | 1.8837016 | 1.51647535 | 0.78045871 | 6482.6182 |
| Motif:rnd-1\_family-438 | 1 | 1.22714198 | 1.18471806 | 0.99007506 | 1.10480543 | 0.20636748 | 0.8306381 | 0.53615978 | 1.31190121 | 1.0880897 | 6356.93916 |
| Motif:rnd-6\_family-9529 | 1 | 1.07032305 | 1.07890462 | 1.0099174 | 1.16762816 | 0.78683185 | 0.63297253 | 0.78034684 | 1.55750769 | 1.24718784 | 6352.83253 |
| Motif:rnd-1\_family-118 | 1 | 0.78870092 | 0.58665294 | 0.78781672 | 0.98790947 | 1.13883981 | 1.20278868 | 1.41317089 | 0.31117591 | 1.11030205 | 6292.47531 |
| Motif:rnd-5\_family-2258 | 1 | 1.16340363 | 1.2515392 | 1.12825964 | 1.02454804 | 0.65987839 | 0.57172135 | 0.5810964 | 1.6881144 | 1.37059442 | 6254.03011 |
| Motif:rnd-5\_family-1029 | 1 | 0.99716588 | 1.01930142 | 0.8890517 | 0.88699406 | 0.41579314 | 0.45030498 | 0.51293164 | 1.32177776 | 1.0857416 | 6211.39883 |
| Motif:rnd-1\_family-496 | 1 | 0.78110321 | 0.61646431 | 0.88251664 | 1.0162979 | 1.16125888 | 1.15730501 | 1.30363034 | 0.09319601 | 0.96705245 | 6042.82414 |
| Motif:rnd-6\_family-6830 | 1 | 0.84894659 | 0.79139381 | 0.83245344 | 0.90905652 | 0.92761766 | 0.8664778 | 1.03801436 | 0.65544541 | 0.78279599 | 5828.77943 |
| Motif:rnd-6\_family-40 | 1 | 1.09510084 | 1.09240644 | 1.03149885 | 1.0704377 | 0.6301675 | 0.67325626 | 0.62304246 | 1.39644454 | 1.14967243 | 5828.12404 |
| Motif:rnd-6\_family-2044 | 1 | 1.05132129 | 0.9589186 | 0.52315076 | 0.72654138 | 0.01793279 | 0.08720769 | 0.05284683 | 0.16545181 | 0.13992381 | 5758.22673 |
| Motif:rnd-1\_family-117 | 1 | 0.5365006 | 1.18058215 | 1.02140076 | 1.10781862 | 1.79894327 | 1.17837008 | 1.52437927 | 1.10387755 | 0.63268894 | 5735.23461 |
| Motif:rnd-1\_family-463 | 1 | 1.23497152 | 1.13315703 | 0.98848524 | 1.22930212 | 0.78871595 | 0.61977542 | 0.6177222 | 1.52792909 | 1.29149829 | 5734.99627 |
| Motif:rnd-1\_family-940 | 1 | 0.99295098 | 1.07580251 | 0.89832542 | 1.01255927 | 0.71292284 | 0.49565081 | 0.52675142 | 1.42063949 | 1.15883847 | 5691.70792 |
| Motif:rnd-1\_family-96 | 1 | 0.98104825 | 0.76620091 | 1.07433602 | 0.90449823 | 1.35255993 | 1.27653853 | 1.22121605 | 0.31434748 | 1.03437936 | 5619.22038 |
| Motif:rnd-6\_family-8871 | 1 | 1.0737974 | 1.13804101 | 0.95778153 | 0.99673135 | 0.37194157 | 0.4047055 | 0.41015883 | 1.27779317 | 1.04149469 | 5573.87906 |
| Motif:rnd-5\_family-2768 | 1 | 1.25259657 | 1.11748661 | 0.98661243 | 1.07959393 | 0.22349747 | 0.42362734 | 0.36077964 | 1.39529543 | 1.1409496 | 5525.18396 |
| Motif:rnd-6\_family-2915 | 1 | 1.13919053 | 1.04246663 | 1.02688262 | 0.96379465 | 0.63165533 | 0.60415544 | 0.64397884 | 1.58838069 | 1.21682007 | 5400.48539 |
| Motif:rnd-5\_family-558 | 1 | 1.15984315 | 1.17214493 | 1.0379368 | 1.12859789 | 0.49883482 | 0.6085343 | 0.51341264 | 1.51762396 | 1.26871622 | 5234.15712 |
| Motif:rnd-1\_family-1531 | 1 | 1.14421119 | 1.25320258 | 1.11149531 | 0.96714991 | 0.52698763 | 1.40975785 | 0.90175455 | 1.54189492 | 1.30476216 | 5228.15343 |
| Motif:rnd-5\_family-150 | 1 | 1.17286932 | 1.17948372 | 1.20905758 | 1.34933678 | 0.76851844 | 0.91097212 | 0.85277494 | 1.11709026 | 1.22470552 | 5186.99943 |
| Motif:rnd-4\_family-1114 | 1 | 1.21734633 | 1.10602048 | 0.94128124 | 1.12398488 | 0.2768804 | 0.45879159 | 0.41852056 | 1.45200984 | 1.1528618 | 5131.01269 |
| Motif:rnd-1\_family-646 | 1 | 0.79071216 | 0.99097235 | 0.86979569 | 0.95709023 | 1.62743183 | 1.18440625 | 1.17713292 | 0.47870209 | 1.03988129 | 5052.02913 |
| Motif:rnd-6\_family-13609 | 1 | 0.8216799 | 0.79143323 | 0.96593815 | 1.10563446 | 1.56120398 | 1.38240631 | 1.37664294 | 0.08439672 | 1.16205646 | 5039.76151 |
| Motif:rnd-4\_family-200 | 1 | 1.18378374 | 1.19785598 | 0.93657576 | 1.0854543 | 0.33527896 | 0.42169049 | 0.37645617 | 1.46362442 | 1.22969034 | 5015.91255 |
| Motif:rnd-5\_family-1630 | 1 | 1.13851057 | 1.00612489 | 0.94878176 | 1.04545285 | 0.25526516 | 0.44116176 | 0.3820916 | 1.28799926 | 1.05055963 | 4995.28511 |
| Motif:rnd-1\_family-1111 | 1 | 1.26339063 | 1.11318458 | 0.93901479 | 0.80246603 | 0.14384322 | 0.47255756 | 0.4239172 | 1.36697716 | 1.12180352 | 4938.76991 |
| Motif:rnd-1\_family-1207 | 1 | 1.15815352 | 1.1646415 | 0.85508017 | 0.79249864 | 0.14901989 | 0.32978926 | 0.28397294 | 1.39538805 | 1.12178738 | 4924.29581 |
| Motif:rnd-1\_family-862 | 1 | 0.788887 | 0.57378196 | 0.81273747 | 1.09725409 | 1.23628854 | 1.2510095 | 1.51855776 | 0.18810602 | 1.11215307 | 4901.84656 |
| Motif:rnd-1\_family-391 | 1 | 0.81570954 | 0.59002686 | 0.94963952 | 0.95309871 | 1.27758821 | 1.27428137 | 1.4485624 | 0.0998644 | 1.19038612 | 4895.43111 |
| Motif:rnd-1\_family-213 | 1 | 1.10405483 | 0.7354382 | 0.75483784 | 0.95477654 | 0.55328406 | 0.5949826 | 0.59312043 | 1.34044274 | 0.76952157 | 4819.42493 |
| Motif:rnd-1\_family-1792 | 1 | 1.10406335 | 1.01955355 | 0.90618134 | 0.90389596 | 0.47358384 | 0.4921088 | 0.46627858 | 1.54134867 | 1.14390833 | 4768.21945 |
| Motif:rnd-1\_family-425 | 1 | 0.86686232 | 0.86000118 | 0.90358334 | 1.0176048 | 0.92013095 | 0.83940669 | 0.97775824 | 0.85270506 | 1.07783551 | 4758.14619 |
| Motif:rnd-1\_family-839 | 1 | 1.06231305 | 1.09650384 | 0.90618403 | 0.98873897 | 0.59625965 | 0.40978793 | 0.44119018 | 1.46562123 | 1.21934229 | 4498.58543 |
| Motif:rnd-5\_family-930 | 1 | 1.11202408 | 1.09818573 | 0.90270116 | 0.99390421 | 0.26754934 | 0.63767609 | 0.45654832 | 1.39433236 | 1.1627868 | 4470.0336 |
| Motif:rnd-1\_family-1070 | 1 | 1.42485878 | 0.95832078 | 1.01541293 | 0.95947149 | 0.09736001 | 0.1566027 | 0.14256299 | 1.4523319 | 1.27143853 | 4403.97398 |
| Motif:rnd-1\_family-103 | 1 | 1.14891543 | 0.82374958 | 1.11543174 | 0.85742063 | 1.63052867 | 1.39237574 | 1.37833183 | 0.07886737 | 1.06839958 | 4389.60415 |
| Motif:rnd-1\_family-1113 | 1 | 1.05531438 | 1.02023278 | 0.97522202 | 0.96427563 | 1.5149631 | 1.03285362 | 1.03790343 | 1.25806593 | 1.00313554 | 4257.55557 |
| Motif:rnd-6\_family-398 | 1 | 1.04235218 | 1.20213641 | 1.25434942 | 1.32854747 | 1.66348551 | 1.57965022 | 1.4666963 | 0.552434 | 1.20756291 | 4245.08243 |
| Motif:rnd-4\_family-572 | 1 | 1.12764442 | 1.11268024 | 0.92023204 | 1.2886505 | 1.13089784 | 1.17916208 | 0.9725032 | 1.38607026 | 1.12345594 | 4237.80522 |
| Motif:rnd-5\_family-3920 | 1 | 1.09330295 | 1.03347955 | 0.88414989 | 0.94940989 | 0.33274229 | 0.52473091 | 0.48396081 | 1.39051383 | 1.13368571 | 4196.84368 |
| Motif:rnd-1\_family-1650 | 1 | 1.18113465 | 1.23041403 | 1.018997 | 1.11159286 | 0.34497426 | 1.0110639 | 0.55951863 | 1.6397396 | 1.34103277 | 4146.77058 |
| Motif:rnd-5\_family-1583 | 1 | 0.87716837 | 0.98922141 | 0.91287448 | 1.01351151 | 2.09944699 | 0.78114608 | 1.03463832 | 1.49799132 | 1.03445311 | 4087.66519 |
| Motif:rnd-1\_family-61 | 1 | 0.98109562 | 0.95448638 | 0.93383965 | 0.95320058 | 0.5532066 | 0.57486336 | 0.58092946 | 1.4174646 | 1.13286051 | 4013.35593 |
| Motif:rnd-6\_family-3909 | 1 | 1.1463962 | 1.01899176 | 0.95334103 | 0.95431744 | 0.41579489 | 0.3240353 | 0.36303176 | 1.35528407 | 1.08969841 | 3971.94234 |
| Motif:rnd-1\_family-1577 | 1 | 1.05894784 | 1.07646891 | 1.08021326 | 1.11916866 | 0.59905017 | 0.47205022 | 0.5269635 | 1.56480294 | 1.22086478 | 3896.57166 |
| Motif:rnd-5\_family-2666 | 1 | 0.97489602 | 1.02329418 | 0.94943272 | 1.02962766 | 2.77330968 | 0.93009931 | 1.10446036 | 1.42755818 | 1.21121469 | 3847.48559 |
| Motif:rnd-5\_family-1664 | 1 | 1.10395164 | 1.18809187 | 0.8921694 | 0.91826285 | 0.32071354 | 0.40358077 | 0.42883585 | 1.32723908 | 1.11942486 | 3838.87015 |
| Motif:rnd-5\_family-3329 | 1 | 1.07716127 | 1.12271946 | 0.96711559 | 1.00601507 | 0.96931715 | 0.51811904 | 0.62475865 | 1.5897116 | 1.0894165 | 3757.46988 |
| Motif:rnd-5\_family-302 | 1 | 1.57770476 | 1.18224171 | 1.21390196 | 1.33884333 | 0.47642058 | 0.55750341 | 0.48677199 | 1.6907481 | 1.29062226 | 3735.27888 |
| Motif:rnd-6\_family-75 | 1 | 0.87255176 | 0.93923127 | 0.80819895 | 0.85457139 | 0.77477474 | 0.68900978 | 0.70920519 | 1.44011287 | 1.0455536 | 3726.56588 |
| Motif:rnd-4\_family-2199 | 1 | 0.85978204 | 1.11865827 | 1.27775272 | 1.13858772 | 1.78239552 | 1.18906352 | 1.39237112 | 0.17812166 | 1.09886346 | 3722.15448 |
| Motif:rnd-5\_family-283 | 1 | 1.01492046 | 1.05392926 | 0.92668972 | 0.84714311 | 0.60854822 | 0.52694007 | 0.4735514 | 1.4453048 | 1.18729998 | 3703.16153 |
| Motif:rnd-6\_family-427 | 1 | 1.21399687 | 1.02255577 | 0.93584209 | 1.04841767 | 0.2019858 | 0.46706593 | 0.36072027 | 1.57425522 | 1.22553271 | 3702.27512 |
| Motif:rnd-6\_family-10863 | 1 | 1.113454 | 1.09144264 | 0.97815051 | 1.02894734 | 2.0338888 | 0.78756167 | 0.92349297 | 1.47081443 | 1.21733841 | 3644.37278 |
| Motif:rnd-5\_family-610 | 1 | 1.15485785 | 1.05634608 | 0.95242523 | 0.99308233 | 0.63938257 | 0.49243217 | 0.52061984 | 1.41314563 | 1.1304791 | 3634.23883 |
| Motif:rnd-6\_family-5358 | 1 | 1.16784147 | 1.32439731 | 0.97993825 | 1.26295227 | 2.76263393 | 2.18255393 | 2.51482242 | 1.3440836 | 1.12272202 | 3597.82979 |
| Motif:rnd-1\_family-410 | 1 | 1.07803974 | 1.05047631 | 0.921978 | 1.13717213 | 0.61171283 | 0.50394799 | 0.54890945 | 1.45754403 | 1.19996101 | 3510.54779 |
| Motif:rnd-1\_family-1523 | 1 | 1.16262228 | 1.12326264 | 0.98535254 | 1.268404 | 0.22067803 | 0.40536749 | 0.36036892 | 1.82056748 | 1.46343224 | 3509.28148 |
| Motif:rnd-1\_family-400 | 1 | 1.01598716 | 1.11921915 | 0.95268037 | 0.87895072 | 0.51924878 | 0.78911823 | 0.66453056 | 1.60237668 | 1.2612523 | 3458.68221 |
| Motif:rnd-1\_family-1059 | 1 | 0.73868611 | 0.87369733 | 0.89074005 | 0.96878389 | 1.12423843 | 1.17763316 | 1.34938668 | 0.41495463 | 1.01839866 | 3416.14583 |
| Motif:rnd-5\_family-2193 | 1 | 1.10213388 | 1.01571675 | 0.96423616 | 0.98602265 | 0.93319844 | 0.76485844 | 0.71293994 | 1.45285382 | 1.12834713 | 3399.70717 |
| Motif:rnd-6\_family-339 | 1 | 0.54316846 | 0.96661846 | 0.71872832 | 1.75850249 | 1.07065228 | 1.35563724 | 1.28925732 | 2.09275243 | 1.4715078 | 3388.65712 |
| Motif:rnd-1\_family-964 | 1 | 0.79205826 | 0.5314068 | 0.79781603 | 0.91742525 | 1.25708308 | 1.20621395 | 1.49012318 | 0.13197924 | 1.08517531 | 3304.72863 |
| Motif:rnd-5\_family-835 | 1 | 1.04793965 | 1.10410037 | 0.9213481 | 0.86504192 | 0.44924355 | 0.30977567 | 0.35493437 | 1.54543119 | 1.19027269 | 3254.12918 |
| Motif:rnd-1\_family-1055 | 1 | 0.69121517 | 0.82333733 | 0.80244095 | 0.91149018 | 1.08219282 | 1.15293599 | 1.22385741 | 0.16956243 | 0.88371151 | 3245.84144 |
| Motif:rnd-6\_family-1699 | 1 | 1.05683734 | 1.06551788 | 0.90774205 | 0.81497028 | 1.08261937 | 0.52899787 | 0.60623178 | 1.12273093 | 0.90873024 | 3236.88165 |
| Motif:rnd-1\_family-12 | 1 | 0.9928688 | 1.1429005 | 1.38757738 | 1.40566108 | 1.650109 | 1.46434667 | 1.37039652 | 0.34503359 | 1.12506115 | 3200.75238 |
| Motif:rnd-6\_family-4476 | 1 | 1.03460599 | 1.08421337 | 0.93472625 | 0.92632357 | 0.69434388 | 0.49167748 | 0.50862547 | 1.47020221 | 1.22012122 | 3187.52182 |
| Motif:rnd-1\_family-670 | 1 | 0.80997867 | 0.4935774 | 0.73210277 | 0.71644417 | 0.95595717 | 1.00762859 | 1.32316527 | 0.07556117 | 0.90133561 | 3166.49333 |
| Motif:rnd-1\_family-1828 | 1 | 1.07078161 | 1.28353425 | 0.89722727 | 1.05670776 | 0.23009336 | 0.30678705 | 0.21227706 | 0.64996811 | 0.55851305 | 3098.00071 |
| Motif:rnd-6\_family-323 | 1 | 1.04485807 | 1.08285141 | 1.12906731 | 1.1541863 | 0.29513982 | 0.71189814 | 0.63711455 | 1.51531073 | 1.35675597 | 3094.86406 |
| Motif:rnd-6\_family-443 | 1 | 1.17085548 | 1.14825007 | 0.89435216 | 1.05225759 | 5.31497074 | 0.90738479 | 1.56430332 | 1.49524969 | 1.16815621 | 3061.24308 |
| Motif:rnd-6\_family-5719 | 1 | 0.68379682 | 1.10119393 | 1.26609117 | 1.28411708 | 1.41523393 | 1.524057 | 1.37624279 | 1.46043742 | 0.61649501 | 3036.01585 |
| Motif:rnd-6\_family-2158 | 1 | 1.21053253 | 1.24044902 | 1.01969861 | 1.15715125 | 0.4170627 | 0.46435752 | 0.40973996 | 1.49094604 | 1.26546851 | 3011.31798 |
| Motif:rnd-5\_family-8237 | 1 | 0.86569426 | 1.24117674 | 0.94347573 | 0.82545001 | 1.26794819 | 1.38365598 | 1.31008993 | 0.07425332 | 1.01334976 | 2967.48459 |
| Motif:rnd-4\_family-3013 | 1 | 1.15699142 | 1.07661175 | 0.86238952 | 1.1006964 | 0.26317167 | 0.37150231 | 0.38237831 | 1.14221732 | 0.96754148 | 2931.19186 |
| Motif:rnd-6\_family-58 | 1 | 1.04202921 | 0.99071976 | 0.91207742 | 0.78580045 | 0.36151456 | 0.37244083 | 0.37113322 | 1.20124336 | 0.99402414 | 2906.64978 |
| Motif:rnd-6\_family-4001 | 1 | 1.28270175 | 1.26580274 | 0.99556373 | 1.0957263 | 0.18549512 | 0.40825006 | 0.32251255 | 1.57264727 | 1.30491094 | 2899.33536 |
| Motif:rnd-5\_family-317 | 1 | 1.04729549 | 1.13513078 | 0.93250967 | 0.92911066 | 0.93177011 | 0.74474201 | 0.68817979 | 1.40890331 | 1.14164968 | 2867.60646 |
| Motif:rnd-5\_family-1090 | 1 | 0.88160044 | 0.9391635 | 1.02943189 | 0.96868761 | 1.59879558 | 1.43726521 | 1.30450932 | 0.09439518 | 1.2024511 | 2861.62467 |
| Motif:rnd-5\_family-3039 | 1 | 1.14053658 | 1.00130046 | 0.78921799 | 0.98941451 | 0.26261039 | 0.57511559 | 0.48461084 | 1.42979561 | 0.90654139 | 2854.12627 |
| Motif:rnd-5\_family-990 | 1 | 1.25748345 | 1.01960811 | 1.11116345 | 1.12570442 | 0.36798313 | 0.4982655 | 0.46027197 | 1.55237121 | 1.29419542 | 2799.30386 |
| Motif:rnd-1\_family-1639 | 1 | 1.08435769 | 1.15658165 | 0.98907247 | 1.24623111 | 0.28745437 | 0.37794311 | 0.34349274 | 1.47457154 | 1.2502376 | 2788.18847 |
| Motif:rnd-6\_family-5168 | 1 | 0.87684055 | 0.95570623 | 0.84197135 | 0.98600808 | 1.23173119 | 0.83965573 | 1.00030945 | 1.42869182 | 0.97133288 | 2778.89847 |
| Motif:rnd-1\_family-1869 | 1 | 1.05177486 | 1.15175129 | 0.87407286 | 1.02876036 | 1.29360695 | 1.19442469 | 1.46220511 | 1.62329808 | 1.12884742 | 2765.84286 |
| Motif:rnd-1\_family-669 | 1 | 1.11008106 | 1.22810256 | 1.03176677 | 1.02319507 | 0.20748559 | 1.24942993 | 0.70299431 | 1.49371696 | 1.24966968 | 2736.71081 |
| Motif:rnd-6\_family-4780 | 1 | 1.06773251 | 1.04184488 | 1.02401706 | 1.04523828 | 0.40938299 | 0.65868484 | 0.57964693 | 1.56708514 | 1.24926029 | 2732.02705 |
| Motif:rnd-6\_family-9065 | 1 | 1.21073606 | 1.17281867 | 1.08207153 | 1.23887063 | 0.78751658 | 0.84582858 | 0.76543855 | 1.60609961 | 1.27618679 | 2725.59777 |
| Motif:rnd-6\_family-12152 | 1 | 1.17577606 | 0.97222307 | 1.04975612 | 1.01094926 | 0.78667492 | 0.97386292 | 1.00847136 | 1.54864787 | 1.12614379 | 2723.42375 |
| Motif:rnd-6\_family-41 | 1 | 1.15996381 | 1.17407969 | 0.96082587 | 1.07818196 | 0.31852597 | 0.88449801 | 0.57478749 | 1.34910243 | 1.10611957 | 2712.52683 |
| Motif:rnd-6\_family-1065 | 1 | 1.08189524 | 1.08971746 | 0.98418985 | 1.14536099 | 3.57102745 | 1.03108154 | 1.10462983 | 1.47904916 | 1.14536194 | 2700.66244 |
| Motif:rnd-1\_family-82 | 1 | 0.95063416 | 1.08070856 | 1.0231456 | 1.03709851 | 0.80550776 | 0.74640561 | 0.71142128 | 1.5835819 | 1.21684542 | 2687.17717 |
| Motif:rnd-1\_family-93 | 1 | 1.1119877 | 0.8728992 | 1.00295471 | 0.93829392 | 1.05031015 | 0.56555601 | 0.48360629 | 1.45498308 | 1.09373648 | 2679.7876 |
| Motif:rnd-6\_family-7151 | 1 | 1.015982 | 1.10304628 | 0.94741238 | 0.88846724 | 2.38944826 | 0.68897934 | 1.01525486 | 1.42218756 | 1.16295529 | 2629.41322 |
| Motif:rnd-6\_family-5254 | 1 | 1.18582537 | 1.27370854 | 1.08593255 | 0.8914676 | 0.33025269 | 0.48705663 | 0.37224416 | 1.61350131 | 1.39549155 | 2624.28315 |
| Motif:rnd-5\_family-41 | 1 | 0.85998373 | 0.97646656 | 0.822432 | 0.87213229 | 1.17907712 | 1.30858195 | 1.30752405 | 1.72223361 | 1.26629947 | 2608.67271 |
| Motif:rnd-6\_family-862 | 1 | 1.08956658 | 1.04015124 | 0.97259611 | 0.9527165 | 1.32242699 | 0.65892057 | 0.69404793 | 1.37568437 | 1.09091855 | 2598.12912 |
| Motif:rnd-6\_family-337 | 1 | 1.13582347 | 1.16950261 | 0.92417767 | 1.06437419 | 0.28791721 | 0.43248796 | 0.36171066 | 1.55589737 | 1.27071416 | 2583.6359 |
| Motif:rnd-5\_family-2450 | 1 | 0.98199004 | 1.01288361 | 0.9829251 | 0.89893869 | 1.76800331 | 0.68817421 | 0.68029888 | 1.44634716 | 1.21117013 | 2583.03659 |
| Motif:rnd-1\_family-152 | 1 | 0.9723884 | 1.00597557 | 0.92206613 | 0.85966083 | 0.72209178 | 0.99751403 | 0.90315707 | 1.52521515 | 1.22860266 | 2573.35357 |
| Motif:rnd-1\_family-1204 | 1 | 0.98198005 | 0.84946632 | 0.89143284 | 0.85178492 | 2.42243361 | 0.70176206 | 0.78076378 | 1.40120878 | 0.95943837 | 2555.2422 |
| Motif:rnd-5\_family-640 | 1 | 1.22673797 | 1.06284904 | 1.01607128 | 0.98998582 | 0.42511274 | 0.64152057 | 0.48940601 | 1.37931057 | 1.14097964 | 2538.07458 |
| Motif:rnd-1\_family-1549 | 1 | 1.02492686 | 1.04985849 | 0.95012011 | 0.74494799 | 1.18365656 | 1.138677 | 1.09181127 | 1.06457244 | 1.13210967 | 2528.07754 |
| Motif:rnd-6\_family-1183 | 1 | 0.69079137 | 1.00574341 | 0.82876419 | 0.86518935 | 1.59967814 | 0.72425894 | 0.83714951 | 1.38510981 | 0.77495333 | 2503.92922 |
| Motif:rnd-1\_family-385 | 1 | 1.1928954 | 1.10425707 | 0.99878673 | 0.93124262 | 0.15756176 | 0.33577324 | 0.29952953 | 1.28746649 | 1.01957799 | 2480.71606 |
| Motif:rnd-1\_family-488 | 1 | 1.09051189 | 0.76067108 | 0.82654169 | 0.63329654 | 1.28029648 | 1.2545922 | 1.08877458 | 0.09009085 | 0.82327378 | 2475.95856 |
| Motif:rnd-6\_family-2055 | 1 | 1.08921181 | 0.93186762 | 1.13094825 | 0.8513931 | 1.96103221 | 1.21813514 | 1.24332879 | 1.70386395 | 1.2671784 | 2431.60832 |
| Motif:rnd-1\_family-1071 | 1 | 1.22360202 | 1.04367858 | 0.95697972 | 0.99288279 | 0.37236411 | 0.57884269 | 0.46476767 | 1.42660762 | 1.168036 | 2404.53761 |
| Motif:rnd-6\_family-8958 | 1 | 1.22410454 | 1.27352138 | 1.17419857 | 1.44678829 | 0.38896309 | 0.57343688 | 0.58515629 | 2.0895005 | 1.81355733 | 2404.44789 |
| Motif:rnd-1\_family-113 | 1 | 1.00878945 | 1.01649163 | 0.87770132 | 0.87865644 | 0.71589053 | 1.31122788 | 0.81602442 | 1.38519884 | 1.12399082 | 2400.14921 |
| Motif:rnd-5\_family-4686 | 1 | 1.06696167 | 1.11716505 | 1.01765044 | 1.0649407 | 0.86324173 | 0.64747942 | 0.68365758 | 1.45852664 | 1.1226463 | 2395.16195 |
| Motif:rnd-1\_family-243 | 1 | 1.14770881 | 1.09952206 | 1.01656094 | 1.2332544 | 0.55880854 | 0.45320043 | 0.44985843 | 1.64765999 | 1.35964007 | 2323.48404 |
| Motif:rnd-4\_family-294 | 1 | 0.85154855 | 0.94907334 | 0.8799107 | 0.94916527 | 1.30459415 | 1.30373551 | 1.40534851 | 1.8813239 | 1.30968983 | 2314.59142 |
| Motif:rnd-6\_family-10437 | 1 | 1.06414275 | 1.1758936 | 1.01368548 | 1.03073615 | 0.41886021 | 0.61222478 | 0.56040764 | 1.47396675 | 1.17447697 | 2280.24774 |
| Motif:rnd-1\_family-424 | 1 | 0.5596888 | 1.08068454 | 0.80474054 | 1.04926679 | 0.78410833 | 0.74734751 | 0.93112472 | 1.29377396 | 0.81231757 | 2223.34837 |
| Motif:rnd-6\_family-1974 | 1 | 1.05636821 | 1.09139472 | 0.92540774 | 0.93376794 | 2.4126569 | 0.83530096 | 1.20793949 | 1.3512339 | 1.07047861 | 2217.38736 |
| Motif:rnd-6\_family-1496 | 1 | 1.26538026 | 0.940653 | 0.84825474 | 0.95407305 | 0.62978771 | 0.37478303 | 0.4351849 | 1.48564028 | 1.01248334 | 2198.75398 |
| Motif:rnd-6\_family-2368 | 1 | 1.13661733 | 1.009891 | 1.01661632 | 1.21959199 | 0.38081242 | 0.58039071 | 0.54806358 | 1.52356481 | 1.21197363 | 2153.79463 |
| Motif:rnd-6\_family-11285 | 1 | 1.18759316 | 1.02698624 | 0.96403872 | 0.86800683 | 0.99176744 | 0.63391437 | 0.77896588 | 1.46551108 | 1.08084561 | 2131.63973 |
| Motif:rnd-1\_family-636 | 1 | 1.14204607 | 1.17627235 | 0.98746822 | 1.10983014 | 0.1993129 | 1.17678234 | 0.65187221 | 1.4606134 | 1.21005749 | 2128.7236 |
| Motif:rnd-1\_family-435 | 1 | 0.77285142 | 0.56605658 | 0.81614952 | 0.60830322 | 1.01630912 | 1.13062093 | 1.09972693 | 0.08143488 | 0.97945886 | 2117.84809 |
| Motif:rnd-2\_family-70 | 1 | 1.00265468 | 1.19971186 | 1.29859948 | 1.31347179 | 1.34762924 | 1.51952473 | 1.32268444 | 0.58945345 | 1.20696601 | 2080.40801 |
| Motif:rnd-6\_family-850 | 1 | 0.9657294 | 1.03062069 | 0.8387209 | 1.12431011 | 1.44073562 | 0.93893801 | 1.15205117 | 1.38050592 | 1.1041693 | 2056.42652 |
| Motif:rnd-6\_family-1505 | 1 | 1.07083398 | 1.07396914 | 0.92979289 | 1.00613985 | 0.69700242 | 0.83718777 | 0.73740033 | 1.30262952 | 1.12350572 | 2030.20097 |
| Motif:rnd-1\_family-955 | 1 | 1.04403914 | 0.77516062 | 0.97923147 | 1.00285005 | 0.75319033 | 0.65508409 | 0.66314529 | 1.50836532 | 1.06746307 | 1981.92501 |
| Motif:rnd-1\_family-1557 | 1 | 1.05141608 | 1.04835393 | 1.03730379 | 1.17513898 | 1.31369567 | 1.25206506 | 1.60488958 | 1.55432584 | 0.98082156 | 1971.49797 |
| Motif:rnd-6\_family-3651 | 1 | 1.0850492 | 1.03568865 | 0.97835536 | 0.93954076 | 1.16157415 | 0.65070066 | 0.7323051 | 1.4532789 | 1.14723872 | 1966.3436 |
| Motif:rnd-5\_family-1208 | 1 | 1.05410937 | 1.30665757 | 1.01916839 | 1.40187443 | 1.4517408 | 1.29822845 | 1.65181796 | 1.55013282 | 1.01119453 | 1956.19843 |
| Motif:rnd-1\_family-16 | 1 | 0.97508347 | 1.1683616 | 1.23527547 | 1.27327454 | 1.53576425 | 1.37586965 | 1.28052696 | 0.59939911 | 1.09895974 | 1950.92406 |
| Motif:rnd-5\_family-2295 | 1 | 1.06660213 | 1.08890884 | 0.93651782 | 0.97151772 | 0.68601164 | 0.92593057 | 0.82338068 | 1.41381792 | 1.08750083 | 1924.08286 |
| Motif:rnd-5\_family-5166 | 1 | 0.99633361 | 1.07232213 | 0.95657358 | 0.90185385 | 0.25277453 | 0.40449928 | 0.35807378 | 1.1671568 | 1.00084805 | 1921.12982 |
| Motif:rnd-6\_family-9316 | 1 | 1.06576118 | 1.13301777 | 1.19569868 | 1.16840265 | 1.59908273 | 1.1373129 | 1.39996164 | 0.20497794 | 1.08075507 | 1919.72318 |
| Motif:rnd-5\_family-2196 | 1 | 0.89469006 | 1.1632353 | 0.93636577 | 1.16730977 | 1.25463814 | 1.58685276 | 1.20810258 | 1.82371227 | 1.34032437 | 1905.65626 |
| Motif:rnd-3\_family-267 | 1 | 0.90203243 | 1.03662671 | 0.91067888 | 0.88919515 | 1.0863693 | 1.50071121 | 1.40866977 | 1.6714979 | 1.22041134 | 1897.11148 |
| Motif:rnd-1\_family-777 | 1 | 1.05981651 | 1.27061877 | 0.99938528 | 0.98264249 | 0.17023377 | 1.50506748 | 0.74562086 | 1.25507374 | 1.09449508 | 1838.81304 |
| Motif:rnd-5\_family-49 | 1 | 1.08673217 | 1.10231857 | 1.12620487 | 1.13160578 | 1.34468427 | 0.68961356 | 0.853662 | 1.42275622 | 1.11762667 | 1834.23073 |
| Motif:rnd-1\_family-1614 | 1 | 0.97626111 | 1.06242237 | 0.94604186 | 1.07447595 | 1.82195637 | 0.82587334 | 0.99103991 | 1.52506941 | 1.13343766 | 1831.90004 |
| Motif:rnd-1\_family-270 | 1 | 1.21238894 | 1.11350514 | 1.00765743 | 0.9193506 | 0.2536939 | 0.53245148 | 0.38256518 | 1.2233454 | 1.09055373 | 1828.89947 |
| Motif:rnd-5\_family-1662 | 1 | 0.8813409 | 0.86744088 | 0.78456729 | 0.77340493 | 0.57993783 | 0.79902812 | 0.79673862 | 0.6070169 | 0.87767055 | 1809.41965 |
| Motif:rnd-1\_family-264 | 1 | 0.99502183 | 1.00008816 | 0.80767773 | 1.07882722 | 0.31872936 | 0.37039974 | 0.3458661 | 1.09564621 | 0.91182986 | 1803.41937 |
| Motif:rnd-6\_family-3591 | 1 | 1.10531248 | 1.15069709 | 1.05103002 | 1.14163877 | 0.5752024 | 0.94940506 | 0.77438553 | 1.55626369 | 1.24849271 | 1800.36227 |
| Motif:rnd-6\_family-684 | 1 | 1.16300504 | 1.09966704 | 0.96654029 | 1.03114549 | 0.2075483 | 0.28850504 | 0.30323128 | 1.59034348 | 1.36321887 | 1777.20813 |
| Motif:rnd-6\_family-1930 | 1 | 1.23161737 | 1.20961811 | 1.08619709 | 1.22554619 | 0.45375171 | 0.51615874 | 0.41367093 | 1.48452176 | 1.19583366 | 1755.90899 |
| Motif:rnd-5\_family-326 | 1 | 1.16896606 | 0.99062157 | 1.0318407 | 1.07205712 | 0.31968578 | 0.81958022 | 0.58886537 | 1.27454015 | 1.14499596 | 1755.87032 |
| Motif:rnd-6\_family-10883 | 1 | 1.13320986 | 1.1166665 | 0.89883067 | 1.16430175 | 0.47581431 | 0.48645478 | 0.46304018 | 1.37852364 | 1.0953261 | 1751.27026 |
| Motif:rnd-5\_family-1420 | 1 | 0.76049836 | 0.62244413 | 0.84075724 | 0.82124076 | 1.1500951 | 1.17347824 | 1.22497742 | 0.10572374 | 0.95766271 | 1746.29017 |
| Motif:rnd-1\_family-207 | 1 | 0.7923129 | 0.71495049 | 0.87740421 | 0.96459164 | 1.20725771 | 1.18983443 | 1.27728971 | 0.11446243 | 0.94396904 | 1722.88334 |
| Motif:rnd-6\_family-3887 | 1 | 1.01274914 | 1.03678988 | 0.95648653 | 0.98804195 | 0.69649593 | 0.7893248 | 0.72197975 | 1.59504943 | 1.22272498 | 1722.17696 |
| Motif:rnd-1\_family-1154 | 1 | 0.70158484 | 0.53956949 | 0.72371823 | 0.66376534 | 1.16056213 | 1.13598364 | 1.40834532 | 0.09455199 | 0.98301128 | 1720.57523 |
| Motif:rnd-6\_family-1554 | 1 | 1.19049265 | 1.27598415 | 1.11483909 | 1.22608792 | 1.19849303 | 1.20008887 | 1.28691029 | 2.16511439 | 1.63307629 | 1706.98825 |
| Motif:rnd-6\_family-1525 | 1 | 1.24242942 | 1.27243577 | 1.10636536 | 1.21489507 | 0.33411025 | 0.66261362 | 0.49117746 | 1.63385565 | 1.3188997 | 1701.24408 |
| Motif:rnd-6\_family-2386 | 1 | 1.14291763 | 1.07840129 | 1.14235605 | 1.02121764 | 2.67104159 | 1.38062476 | 1.88149856 | 1.56188217 | 1.117071 | 1695.35987 |
| Motif:rnd-1\_family-274 | 1 | 1.08194237 | 1.0969399 | 0.95501726 | 1.10984456 | 0.81571092 | 0.71245846 | 0.69268268 | 1.4539112 | 1.16857705 | 1695.3099 |
| Motif:rnd-6\_family-425 | 1 | 1.12409559 | 1.32294494 | 1.18202851 | 0.96680885 | 0.52692651 | 1.77378998 | 1.10383671 | 1.66085006 | 1.40940301 | 1689.72224 |
| Motif:rnd-1\_family-515 | 1 | 1.22483825 | 1.21770166 | 1.03021635 | 1.07810813 | 0.25095431 | 0.74049937 | 0.51449782 | 1.52091662 | 1.23477611 | 1688.59576 |
| Motif:rnd-6\_family-470 | 1 | 1.03970857 | 1.10033305 | 0.95584366 | 1.08206746 | 0.39865483 | 0.73469917 | 0.65006194 | 1.51616198 | 1.16991905 | 1686.90713 |
| Motif:rnd-1\_family-202 | 1 | 0.71029357 | 0.98941108 | 0.86914293 | 1.06031689 | 1.22023302 | 1.45281249 | 1.50528282 | 1.70097509 | 1.16580061 | 1684.04601 |
| Motif:rnd-1\_family-518 | 1 | 1.07126246 | 1.06289995 | 0.78684704 | 0.74677474 | 0.38487907 | 0.37815958 | 0.43749183 | 0.90386191 | 0.70224805 | 1682.91484 |
| Motif:rnd-6\_family-1887 | 1 | 1.12194312 | 1.14749032 | 1.02083613 | 1.16466771 | 0.67015071 | 1.52283806 | 1.1934444 | 1.53379879 | 1.25143809 | 1652.70074 |
| Motif:rnd-1\_family-911 | 1 | 1.15646525 | 1.12547151 | 0.91367335 | 1.0515241 | 0.21629348 | 0.47707129 | 0.36570106 | 1.45318082 | 1.17602224 | 1651.67083 |
| Motif:rnd-6\_family-12619 | 1 | 1.0785354 | 1.09920088 | 0.89714017 | 0.91772101 | 0.28734556 | 0.402843 | 0.38084734 | 1.28997786 | 0.99280254 | 1648.16518 |
| Motif:rnd-4\_family-469 | 1 | 0.98382862 | 0.93389455 | 0.89334613 | 0.90944835 | 0.77886881 | 1.08116014 | 0.96334583 | 1.54899316 | 1.24584954 | 1643.32371 |
| Motif:rnd-6\_family-5982 | 1 | 1.14191201 | 0.96301527 | 0.98205799 | 1.01547992 | 1.23822431 | 0.68691704 | 0.66658276 | 1.20695486 | 1.00981569 | 1632.08524 |
| Motif:rnd-1\_family-866 | 1 | 0.8619042 | 0.93906044 | 0.84913943 | 0.88167733 | 1.20326776 | 1.14339806 | 1.34695777 | 1.59849812 | 1.255292 | 1626.63462 |
| Motif:rnd-6\_family-1772 | 1 | 1.09029748 | 1.17487035 | 0.92853059 | 1.25012023 | 0.84151757 | 0.5927295 | 0.65790129 | 1.58169578 | 1.23557102 | 1617.36115 |
| Motif:rnd-1\_family-40 | 1 | 0.55750965 | 1.01825519 | 0.96359043 | 1.1926151 | 1.49683084 | 1.19714285 | 1.4569663 | 0.94833546 | 0.81790496 | 1616.6077 |
| Motif:rnd-6\_family-11911 | 1 | 0.84063691 | 1.07784506 | 1.03765382 | 1.37581919 | 1.60568421 | 1.28526146 | 1.46412353 | 1.41309696 | 1.11866784 | 1613.59523 |
| Motif:rnd-1\_family-686 | 1 | 0.81870171 | 0.88357538 | 0.83020668 | 0.79254787 | 1.1954213 | 1.0817323 | 1.12644886 | 1.59017522 | 1.14599451 | 1607.32169 |
| Motif:rnd-5\_family-2058 | 1 | 0.81976791 | 0.89585616 | 0.94614351 | 1.178883 | 1.45195037 | 1.48709857 | 1.30843108 | 0.16968164 | 1.00322315 | 1606.79008 |
| Motif:rnd-1\_family-24 | 1 | 1.07323237 | 1.11318016 | 1.12682317 | 0.96503309 | 0.51137478 | 1.39594296 | 0.78407919 | 1.54103607 | 1.24290945 | 1594.34179 |
| Motif:rnd-6\_family-7331 | 1 | 1.05364331 | 1.03376196 | 0.84680056 | 0.90798193 | 0.21987669 | 0.61943973 | 0.47070905 | 1.46285628 | 1.21634548 | 1590.77087 |
| Motif:rnd-6\_family-5915 | 1 | 1.2412811 | 1.3900689 | 0.95458552 | 1.05218672 | 0.12400214 | 0.49836182 | 0.2754915 | 0.9049756 | 0.74462143 | 1577.74735 |
| Motif:rnd-5\_family-3705 | 1 | 1.10763077 | 1.003142 | 0.94320995 | 0.90330807 | 0.28341623 | 0.90618477 | 0.60474145 | 1.57690717 | 1.31819779 | 1576.39944 |
| Motif:rnd-5\_family-2423 | 1 | 0.95108328 | 1.07857164 | 0.88628752 | 0.92018974 | 0.9119839 | 1.75318031 | 1.34633531 | 1.73148041 | 1.28779403 | 1567.11257 |
| Motif:rnd-1\_family-1913 | 1 | 1.16712813 | 1.13439973 | 0.98039173 | 1.13524435 | 0.43260948 | 0.78936154 | 0.59849695 | 1.21396119 | 0.94620848 | 1554.18592 |
| Motif:rnd-1\_family-477 | 1 | 1.0956095 | 1.18205098 | 0.97645685 | 0.90316128 | 0.12875699 | 2.02853693 | 1.13084195 | 1.47671344 | 1.37055969 | 1550.31082 |
| Motif:rnd-6\_family-2023 | 1 | 1.22811151 | 1.22951435 | 0.99694559 | 1.16996852 | 0.18765461 | 0.26352211 | 0.2342762 | 1.15081994 | 0.96826558 | 1549.6736 |
| Motif:rnd-6\_family-3547 | 1 | 0.92245413 | 1.1017195 | 0.91848595 | 0.88144156 | 0.83928117 | 0.65055085 | 0.56993779 | 1.39796126 | 1.07466314 | 1549.41919 |
| Motif:rnd-1\_family-302 | 1 | 0.88469977 | 0.87421504 | 0.88652664 | 0.92489421 | 0.98337386 | 0.91183193 | 0.84504291 | 0.71452859 | 1.05572008 | 1543.1875 |
| Motif:rnd-6\_family-10182 | 1 | 0.91076654 | 0.97192637 | 0.9827289 | 0.99384662 | 1.12005399 | 0.87413337 | 0.88623508 | 1.51314937 | 1.19324302 | 1540.137 |
| Motif:rnd-1\_family-373 | 1 | 0.8497738 | 0.74337127 | 0.85686304 | 0.74513523 | 1.23328518 | 1.27503334 | 1.54090464 | 2.13253753 | 1.4788358 | 1533.87769 |
| Motif:rnd-6\_family-2079 | 1 | 1.14060313 | 1.04077941 | 0.91798541 | 1.01908738 | 0.76622346 | 0.42366701 | 0.50414894 | 1.38439913 | 1.06292918 | 1515.22356 |
| Motif:rnd-1\_family-726 | 1 | 1.16693686 | 1.31201321 | 1.03458799 | 1.06466499 | 0.21513602 | 1.57524338 | 0.778049 | 1.39471041 | 1.1956482 | 1511.38624 |
| Motif:rnd-6\_family-10536 | 1 | 0.96231856 | 1.06397618 | 0.87497094 | 0.98837757 | 0.31911066 | 0.47340406 | 0.50897258 | 1.31705767 | 0.8924002 | 1489.46668 |
| Motif:rnd-5\_family-866 | 1 | 1.09449926 | 0.99837402 | 0.85141687 | 1.15574141 | 0.4569357 | 0.75947433 | 0.52767706 | 1.63774013 | 1.33805756 | 1485.49814 |
| Motif:rnd-6\_family-4218 | 1 | 1.08969987 | 0.94562502 | 0.94307136 | 0.98848266 | 0.3104697 | 0.4940951 | 0.42914757 | 1.30234597 | 1.10452982 | 1480.78911 |
| Motif:rnd-1\_family-337 | 1 | 1.10364236 | 0.98740819 | 1.05783999 | 1.23607652 | 0.59675566 | 0.9576028 | 0.79055998 | 1.47589918 | 1.26596128 | 1479.25163 |
| Motif:rnd-6\_family-1027 | 1 | 1.39358831 | 1.16676709 | 1.1212912 | 1.25230149 | 0.19239389 | 0.28149811 | 0.2362379 | 1.58586248 | 1.27775241 | 1470.26984 |
| Motif:rnd-4\_family-1511 | 1 | 0.88604454 | 0.88907209 | 0.89103581 | 1.04314519 | 1.09606634 | 0.99890863 | 0.99404652 | 1.48412201 | 1.05436501 | 1448.85759 |
| Motif:rnd-1\_family-1619 | 1 | 0.69238918 | 0.9677718 | 0.80414195 | 0.88437083 | 1.48819929 | 1.30270762 | 1.63239306 | 1.91853058 | 1.29994246 | 1448.04372 |
| Motif:rnd-1\_family-939 | 1 | 1.26786214 | 1.16560515 | 0.87963319 | 1.4698417 | 0.26093093 | 0.66843913 | 0.5494214 | 1.55432458 | 1.19655364 | 1427.31363 |
| Motif:rnd-6\_family-5159 | 1 | 1.04960204 | 1.00107303 | 0.9355959 | 0.85784743 | 0.57789753 | 0.49767106 | 0.51825823 | 1.34253875 | 1.06779566 | 1417.70301 |
| Motif:rnd-1\_family-447 | 1 | 0.83816759 | 0.68722622 | 1.0269163 | 1.14759963 | 1.41050934 | 1.30559348 | 1.36798494 | 0.23773724 | 0.99408219 | 1414.01979 |
| Motif:rnd-1\_family-1107 | 1 | 0.93175219 | 0.94721579 | 0.91278488 | 0.80726071 | 0.83892847 | 0.94501677 | 0.94418075 | 1.61709114 | 1.23938129 | 1413.29207 |
| Motif:rnd-1\_family-84 | 1 | 1.00708169 | 0.83339691 | 0.99782611 | 0.93097434 | 1.24552435 | 1.26826473 | 1.16070086 | 0.20616707 | 0.95373393 | 1408.69353 |
| Motif:rnd-5\_family-139 | 1 | 1.00170379 | 0.97983282 | 0.98992741 | 1.08461982 | 1.27981266 | 1.32319733 | 1.49100617 | 0.22656133 | 1.13054721 | 1405.80346 |
| Motif:rnd-1\_family-1696 | 1 | 0.76812061 | 1.16973181 | 1.06230649 | 1.43477324 | 1.52193546 | 1.16141979 | 1.38732544 | 1.43878422 | 0.61464824 | 1402.9819 |
| Motif:rnd-1\_family-173 | 1 | 1.19119177 | 1.14177731 | 1.01419211 | 1.01802257 | 0.34973556 | 0.61052008 | 0.49379276 | 1.73765499 | 1.31958574 | 1376.21819 |
| Motif:rnd-6\_family-3508 | 1 | 0.71774485 | 1.0468869 | 0.86480141 | 1.45657008 | 1.25785183 | 1.41696687 | 1.47575478 | 2.20086581 | 1.54774668 | 1376.03933 |
| Motif:rnd-5\_family-1764 | 1 | 0.73949919 | 0.87130553 | 0.82983943 | 1.09659131 | 1.36703408 | 1.38255331 | 1.43572237 | 0.61160667 | 1.12157866 | 1370.48576 |
| Motif:rnd-6\_family-2880 | 1 | 1.18793426 | 1.08708172 | 0.98131497 | 1.04904299 | 0.2654288 | 0.38188574 | 0.35543169 | 1.50119832 | 1.28298572 | 1368.73792 |
| Motif:rnd-1\_family-532 | 1 | 0.80775297 | 0.67972958 | 0.83693446 | 0.87004991 | 1.15075907 | 1.22665771 | 1.38481287 | 0.62835214 | 1.03963137 | 1366.9292 |
| Motif:rnd-1\_family-369 | 1 | 0.95749053 | 1.03345217 | 0.88053279 | 1.08358566 | 1.14111197 | 1.14696893 | 1.22280622 | 1.36200688 | 1.24294618 | 1361.42297 |
| Motif:rnd-6\_family-2223 | 1 | 1.22848256 | 1.27884366 | 1.09293439 | 1.20593463 | 0.53800597 | 0.73720524 | 0.58260208 | 1.69824316 | 1.39478641 | 1355.65237 |
| Motif:rnd-1\_family-1749 | 1 | 1.143249 | 1.07192017 | 0.8674734 | 0.97565284 | 0.29532953 | 0.46142635 | 0.40197745 | 1.54703628 | 1.19115253 | 1329.7663 |
| Motif:rnd-6\_family-992 | 1 | 1.09206392 | 0.98450046 | 0.96727124 | 0.86211915 | 0.42482519 | 0.68917057 | 0.7572153 | 1.42832957 | 1.19181027 | 1327.9405 |
| Motif:rnd-5\_family-23 | 1 | 1.27151396 | 1.24525462 | 1.28694996 | 1.22643274 | 0.13368736 | 0.63605963 | 0.4678927 | 2.17180846 | 1.74043901 | 1325.23133 |
| Motif:rnd-1\_family-1262 | 1 | 1.14739904 | 0.95209101 | 0.9851956 | 1.06388789 | 0.37860967 | 0.91963137 | 0.74732368 | 1.67016244 | 1.31513862 | 1323.21308 |
| Motif:rnd-1\_family-1252 | 1 | 1.2649065 | 0.83941072 | 0.88517954 | 0.8893426 | 0.37078669 | 0.66539329 | 0.72324201 | 1.38094185 | 0.90745457 | 1307.9702 |
| Motif:rnd-5\_family-4697 | 1 | 0.7638266 | 1.15271389 | 0.95865452 | 1.05601851 | 0.63522314 | 0.92957174 | 0.8332124 | 1.29407925 | 1.07326364 | 1305.97511 |
| Motif:rnd-1\_family-832 | 1 | 1.22157622 | 1.19561355 | 0.97320638 | 1.22616424 | 0.23899813 | 0.35642609 | 0.2812863 | 1.63403033 | 1.32017922 | 1305.55095 |
| Motif:rnd-6\_family-2943 | 1 | 1.05409527 | 0.95665336 | 1.06354809 | 1.43477525 | 0.53070838 | 1.09941712 | 0.91568083 | 1.81745565 | 1.46522074 | 1303.9829 |
| Motif:rnd-6\_family-7992 | 1 | 1.18222275 | 1.14487464 | 0.93071072 | 1.04343329 | 0.73677455 | 0.75498502 | 0.63006996 | 1.4993999 | 1.2419057 | 1298.14126 |
| Motif:rnd-6\_family-9763 | 1 | 1.18717215 | 0.97080676 | 0.89067112 | 0.92214799 | 0.79145392 | 0.29034331 | 0.38320126 | 1.13527414 | 0.91261038 | 1292.38225 |
| Motif:rnd-6\_family-2729 | 1 | 1.26352167 | 1.18142715 | 0.96833429 | 0.98819309 | 0.14604987 | 0.33142097 | 0.25636431 | 1.16596965 | 1.00106294 | 1284.58336 |
| Motif:rnd-6\_family-16609 | 1 | 1.06239266 | 1.15969564 | 1.05287623 | 1.05951372 | 0.86535147 | 0.77149025 | 0.63790888 | 1.47826077 | 1.12826615 | 1274.86254 |
| Motif:rnd-6\_family-3103 | 1 | 1.00719871 | 0.95290515 | 1.02864676 | 1.03967608 | 0.6379948 | 0.75975286 | 0.80924461 | 1.51690725 | 1.019977 | 1268.21149 |
| Motif:rnd-1\_family-167 | 1 | 0.85152257 | 1.10269512 | 0.89632793 | 0.87708876 | 0.91904211 | 1.72943278 | 1.28615849 | 1.71458065 | 1.23877203 | 1267.21551 |
| Motif:rnd-6\_family-1138 | 1 | 1.30903111 | 1.08268733 | 1.16425079 | 1.43135038 | 0.17568714 | 0.29841623 | 0.31543662 | 1.72967605 | 1.4818533 | 1261.83975 |
| Motif:rnd-6\_family-12116 | 1 | 1.01847618 | 0.99946004 | 1.02376536 | 1.14307313 | 0.46867834 | 0.68130559 | 0.61310839 | 1.46162743 | 1.22553412 | 1257.86292 |
| Motif:rnd-1\_family-1037 | 1 | 1.11337038 | 0.99559157 | 0.99432044 | 1.13767493 | 1.02827618 | 0.81950633 | 0.83420096 | 1.67307808 | 1.24644118 | 1253.14518 |
| Motif:rnd-6\_family-3357 | 1 | 0.75323867 | 0.77687097 | 0.76320026 | 0.70273042 | 1.30541233 | 1.19796533 | 1.48081135 | 1.70375511 | 1.10447437 | 1240.97891 |
| Motif:rnd-6\_family-2490 | 1 | 1.21301836 | 1.18573159 | 0.94888816 | 0.93984901 | 0.20043406 | 0.73559902 | 0.57062189 | 1.32840036 | 1.12713355 | 1232.99223 |
| Motif:rnd-6\_family-2181 | 1 | 0.95321301 | 0.98833858 | 1.06265711 | 1.19067605 | 1.33141985 | 1.39630713 | 1.3170399 | 0.53041763 | 1.07427181 | 1231.57113 |
| Motif:rnd-6\_family-314 | 1 | 1.21771617 | 1.24596217 | 1.03191254 | 1.19320417 | 0.38038126 | 0.76006535 | 0.54445995 | 1.47803607 | 1.2001631 | 1228.02876 |
| Motif:rnd-6\_family-1052 | 1 | 1.25453106 | 1.27067047 | 1.06212173 | 1.08794579 | 0.23650519 | 0.52020863 | 0.44572127 | 1.58371436 | 1.22829194 | 1218.58417 |
| Motif:rnd-5\_family-7700 | 1 | 1.06604569 | 1.09459375 | 1.03578484 | 1.13476948 | 0.26058092 | 0.39467695 | 0.37250838 | 1.44456341 | 1.17316001 | 1215.31047 |
| Motif:rnd-1\_family-731 | 1 | 0.87137085 | 0.94160177 | 0.82778802 | 0.94321669 | 0.78591898 | 0.81261883 | 0.72606639 | 1.42286126 | 1.11365689 | 1214.94916 |
| Motif:rnd-1\_family-131 | 1 | 0.88268091 | 0.98623788 | 0.88220557 | 1.01449306 | 1.3880648 | 1.2443422 | 1.29536493 | 1.63165147 | 1.21981353 | 1213.28965 |
| Motif:rnd-6\_family-10230 | 1 | 0.66825109 | 1.11093499 | 1.04794922 | 0.84407994 | 2.24709173 | 1.37795652 | 1.86669441 | 1.56355931 | 0.41504498 | 1198.25378 |
| Motif:rnd-6\_family-1323 | 1 | 0.92599688 | 1.07754982 | 0.97395786 | 0.93867621 | 0.89245997 | 1.20667411 | 1.40928458 | 1.32288405 | 1.10202792 | 1191.97653 |
| Motif:rnd-5\_family-2038 | 1 | 1.11306825 | 1.05900218 | 1.0091422 | 1.42691339 | 0.35643487 | 0.61866162 | 0.57873795 | 1.49611001 | 1.15379914 | 1185.60293 |
| Motif:rnd-1\_family-98 | 1 | 0.83008196 | 0.91436784 | 0.90657 | 1.02139984 | 1.05951503 | 1.14561827 | 0.93142753 | 0.82172474 | 1.09006368 | 1183.79797 |
| Motif:rnd-1\_family-174 | 1 | 0.88414108 | 0.81259161 | 0.90344497 | 0.82780341 | 0.8916309 | 1.34434376 | 1.18753904 | 0.5545946 | 1.08327442 | 1182.79783 |
| Motif:rnd-6\_family-12996 | 1 | 1.20054561 | 1.04822524 | 0.88585732 | 0.98601038 | 0.25627879 | 0.39974015 | 0.44026683 | 1.42194941 | 1.06751233 | 1178.43658 |
| Motif:rnd-1\_family-1593 | 1 | 0.92827114 | 1.15704122 | 1.17014504 | 1.34727028 | 1.66140344 | 1.54849901 | 1.40328228 | 2.04916164 | 1.42218695 | 1169.5065 |
| Motif:rnd-1\_family-408 | 1 | 0.84496389 | 1.03117743 | 1.04409526 | 0.95071048 | 1.85412796 | 1.31543463 | 1.40478919 | 0.41617878 | 1.04050976 | 1166.60607 |
| Motif:rnd-1\_family-294 | 1 | 0.9774039 | 1.05902199 | 0.87491328 | 0.82824672 | 0.5443429 | 1.45158714 | 0.96729425 | 1.39403225 | 1.21125238 | 1156.4189 |
| Motif:rnd-1\_family-1155 | 1 | 1.08328717 | 1.07032815 | 1.05196634 | 0.96017487 | 0.79150723 | 0.56078131 | 0.66615588 | 1.60886058 | 1.22881699 | 1142.0505 |
| Motif:rnd-3\_family-366 | 1 | 1.10291284 | 1.02917907 | 1.05257146 | 0.85370704 | 0.92986986 | 0.99080781 | 0.91791506 | 1.07246387 | 1.09362296 | 1128.87364 |
| Motif:rnd-5\_family-3272 | 1 | 0.93406076 | 0.72922146 | 1.0127688 | 0.88784511 | 1.42585956 | 1.3693267 | 1.59484326 | 0.11278288 | 1.05202434 | 1128.56589 |
| Motif:rnd-6\_family-1241 | 1 | 0.94133085 | 1.10558196 | 1.25582094 | 0.65515902 | 1.55377443 | 0.76423267 | 0.71936157 | 1.48176502 | 1.1864948 | 1127.89895 |
| Motif:rnd-1\_family-650 | 1 | 1.32987512 | 1.07188828 | 1.00943161 | 1.22955702 | 0.17999645 | 0.45350563 | 0.34482342 | 1.64558848 | 1.59955472 | 1122.40195 |
| Motif:rnd-1\_family-193 | 1 | 0.89204497 | 1.03777959 | 0.90138767 | 0.911877 | 1.01438484 | 2.10342282 | 1.46138022 | 1.67904349 | 1.3025796 | 1116.67179 |
| Motif:rnd-5\_family-2209 | 1 | 0.52587505 | 1.11170115 | 0.76579535 | 1.96401626 | 1.44050292 | 1.2808716 | 1.55999287 | 0.53503853 | 0.95331379 | 1116.16063 |
| Motif:rnd-5\_family-2004 | 1 | 1.12268947 | 0.99199635 | 0.96943194 | 1.09268339 | 0.43477392 | 0.55901362 | 0.50520578 | 1.66284733 | 1.35820929 | 1115.35463 |
| Motif:rnd-5\_family-1311 | 1 | 0.60133759 | 1.36552122 | 1.14885241 | 1.22836313 | 1.8085478 | 1.46162283 | 1.79182936 | 2.01618184 | 0.88343538 | 1109.21223 |
| Motif:rnd-1\_family-3 | 1 | 0.90097638 | 0.96319497 | 0.92758234 | 1.00705383 | 0.95187079 | 0.98178832 | 0.95883368 | 1.46585818 | 1.11530975 | 1107.68934 |
| Motif:rnd-1\_family-1209 | 1 | 1.13877836 | 1.12996164 | 1.03142062 | 0.99205857 | 0.24679964 | 0.59241928 | 0.42561275 | 1.74109241 | 1.44148762 | 1107.5029 |
| Motif:rnd-1\_family-635 | 1 | 1.08613752 | 1.3106775 | 1.00616941 | 0.97237772 | 0.15813894 | 1.72937644 | 0.99069305 | 1.46885441 | 1.33719062 | 1101.20685 |
| Motif:rnd-1\_family-1119 | 1 | 0.85134503 | 1.07598505 | 0.87006062 | 1.05555046 | 0.39140678 | 0.47440313 | 0.41219871 | 1.2831698 | 1.05816156 | 1100.46781 |
| Motif:rnd-5\_family-20 | 1 | 1.30732652 | 0.57346632 | 1.05150344 | 1.00150037 | 0.63151599 | 0.49594862 | 0.54853964 | 1.34847926 | 1.04904291 | 1093.9752 |
| Motif:rnd-5\_family-3317 | 1 | 0.60795242 | 1.28823651 | 0.93869716 | 1.46232079 | 1.00580879 | 0.81912298 | 0.77648341 | 1.42140366 | 0.57337219 | 1081.3996 |
| Motif:rnd-6\_family-3572 | 1 | 0.87623231 | 1.06006434 | 0.98586968 | 1.04558182 | 1.13625978 | 1.02243194 | 1.01772508 | 1.58323668 | 1.19610568 | 1072.52393 |
| Motif:rnd-1\_family-1682 | 1 | 0.88794809 | 1.09836219 | 0.80273532 | 0.50053734 | 0.87913682 | 1.05055612 | 1.01347182 | 1.53151049 | 1.15616453 | 1070.11701 |
| Motif:rnd-6\_family-2764 | 1 | 1.08402723 | 0.69997336 | 1.0324856 | 0.9803891 | 0.64384675 | 0.5360317 | 0.56313723 | 1.24105687 | 1.00580164 | 1069.83735 |
| Motif:rnd-1\_family-603 | 1 | 0.96421787 | 1.02852374 | 0.9166406 | 0.83121986 | 0.38274019 | 0.61687523 | 0.49078676 | 1.29881903 | 1.08086979 | 1067.65262 |
| Motif:rnd-1\_family-1184 | 1 | 1.04683131 | 1.13128653 | 0.98775185 | 0.96239522 | 0.3389701 | 0.66991734 | 0.42354531 | 1.44026811 | 1.15921033 | 1066.38861 |
| Motif:rnd-6\_family-3028 | 1 | 1.11240423 | 1.05744822 | 0.98104366 | 0.95996497 | 0.26204532 | 1.00929692 | 0.70954942 | 1.43297356 | 1.19438488 | 1065.17131 |
| Motif:rnd-5\_family-2753 | 1 | 1.13112295 | 1.29379728 | 1.02899971 | 1.16880511 | 0.3487462 | 0.75163036 | 0.61632735 | 1.24021815 | 1.13359557 | 1063.65533 |
| Motif:rnd-1\_family-1143 | 1 | 1.02480356 | 1.01440856 | 0.94496769 | 1.06195122 | 0.35997377 | 0.7228554 | 0.6458306 | 1.30015609 | 1.17515899 | 1061.83632 |
| Motif:rnd-1\_family-401 | 1 | 1.06220369 | 1.00452573 | 0.95582573 | 1.21228814 | 0.34801929 | 0.44395684 | 0.46718154 | 1.47392447 | 1.09525585 | 1055.23018 |
| Motif:rnd-6\_family-273 | 1 | 1.28159257 | 1.35908511 | 1.02439092 | 1.58076784 | 0.18965731 | 0.2558933 | 0.26984349 | 1.8411336 | 1.58391857 | 1054.70161 |
| Motif:rnd-6\_family-1295 | 1 | 1.14854652 | 1.11423843 | 0.88199817 | 1.1201446 | 0.28089588 | 0.33856432 | 0.3944232 | 1.67140015 | 1.32487933 | 1052.64158 |
| Motif:rnd-1\_family-122 | 1 | 1.03035985 | 0.91473141 | 1.00170158 | 0.99947242 | 1.00560728 | 0.865973 | 0.82910201 | 1.38489885 | 1.0874491 | 1042.54048 |
| Motif:rnd-1\_family-1265 | 1 | 1.06076514 | 0.95627554 | 1.00308864 | 1.09037708 | 0.50376677 | 0.51354014 | 0.59912522 | 1.60311083 | 1.41597528 | 1042.00707 |
| Motif:rnd-6\_family-4563 | 1 | 0.51217352 | 1.30703837 | 0.97249308 | 0.77965827 | 2.44978173 | 1.51371943 | 2.01762077 | 0.76992837 | 0.56064905 | 1034.63693 |
| Motif:rnd-6\_family-2824 | 1 | 0.99757727 | 0.92204813 | 1.01756849 | 1.19039672 | 1.70083392 | 1.50583026 | 1.58037882 | 1.9276375 | 1.35959819 | 1029.34816 |
| Motif:rnd-5\_family-2012 | 1 | 1.11508541 | 1.16275722 | 0.95540634 | 1.2115632 | 0.59808485 | 0.55966442 | 0.60789927 | 1.52151278 | 1.22176383 | 1024.33206 |
| Motif:rnd-6\_family-9696 | 1 | 1.13309414 | 1.19685416 | 1.1248406 | 1.49144585 | 0.34271089 | 0.70023506 | 0.55176755 | 1.77491082 | 1.52214576 | 1020.61303 |
| Motif:rnd-6\_family-5311 | 1 | 1.01174325 | 0.95335734 | 0.83068497 | 1.26217193 | 0.18671124 | 0.65565765 | 0.32425311 | 1.2909235 | 1.0679068 | 1013.25323 |
| Motif:rnd-6\_family-12105 | 1 | 1.10237612 | 1.10257834 | 0.85715282 | 0.89339641 | 0.1605965 | 0.40043515 | 0.28420157 | 1.05553868 | 0.96479807 | 1010.1057 |
| Motif:rnd-6\_family-7506 | 1 | 0.8999543 | 0.99004122 | 0.94778794 | 0.82745609 | 1.90012651 | 1.42215529 | 1.4736127 | 1.86818618 | 1.19088428 | 1009.07555 |
| Motif:rnd-6\_family-6470 | 1 | 0.8976733 | 1.12087256 | 1.10207008 | 0.94882276 | 1.68098219 | 1.24386554 | 1.37440144 | 1.80138062 | 1.26662703 | 1008.52025 |
| Motif:rnd-6\_family-6399 | 1 | 0.89311231 | 1.02100761 | 0.89408598 | 1.06343737 | 0.41963819 | 0.52302825 | 0.56601163 | 1.35519819 | 1.17660369 | 1006.60822 |
| Motif:rnd-6\_family-6882 | 1 | 1.12973603 | 1.05809417 | 1.04341362 | 1.07277669 | 0.62116876 | 0.7454419 | 0.65412259 | 1.36352107 | 1.16419681 | 1002.92047 |
| Motif:rnd-6\_family-5714 | 1 | 0.78961394 | 0.73333199 | 0.89383308 | 1.18287302 | 1.25643666 | 1.23013229 | 1.57079956 | 0.10850659 | 1.11834984 | 1002.25997 |
| Motif:rnd-6\_family-72 | 1 | 1.1062193 | 0.54778988 | 0.80714424 | 1.01980772 | 0.41818634 | 0.47529923 | 0.56432597 | 1.49420853 | 1.02512496 | 998.956031 |
| Motif:rnd-1\_family-164 | 1 | 1.10694127 | 0.99377462 | 0.98153309 | 1.11784404 | 0.35438499 | 0.37135098 | 0.3279615 | 1.28906606 | 1.11319365 | 996.484587 |
| Motif:rnd-4\_family-44 | 1 | 1.12907656 | 1.24141164 | 1.21052521 | 1.03612366 | 0.81884751 | 2.57633239 | 1.39498926 | 1.11716836 | 1.39757233 | 994.580751 |
| Motif:rnd-1\_family-88 | 1 | 1.01001 | 1.04861482 | 0.91681139 | 1.19187659 | 0.85634751 | 0.94445471 | 0.88247478 | 1.5030581 | 1.1384048 | 991.060106 |
| Motif:rnd-1\_family-308 | 1 | 0.85699547 | 0.75181982 | 0.87349596 | 0.95687285 | 1.20153421 | 1.24250836 | 1.17456756 | 0.62399133 | 1.04155255 | 986.210287 |
| Motif:rnd-1\_family-1391 | 1 | 1.30758956 | 1.25403435 | 0.88945263 | 0.97404659 | 0.18147757 | 0.19927954 | 0.21804436 | 1.47975783 | 1.27182449 | 982.722998 |
| Motif:rnd-1\_family-240 | 1 | 0.86195491 | 1.02225369 | 0.8648893 | 0.87714005 | 0.92085145 | 1.36598091 | 1.08045393 | 1.58424352 | 1.21104175 | 978.376164 |
| Motif:rnd-1\_family-1199 | 1 | 0.94527584 | 1.01456668 | 0.82663825 | 0.66196702 | 0.37544813 | 1.24267603 | 0.9462528 | 1.43099382 | 1.05460955 | 973.597193 |
| Motif:rnd-5\_family-291 | 1 | 1.08601402 | 1.07456062 | 0.91201258 | 0.90592528 | 0.69512191 | 0.70691115 | 0.7346394 | 1.59713143 | 1.21722869 | 968.496932 |
| Motif:rnd-6\_family-2543 | 1 | 1.26219804 | 1.17304562 | 1.05691634 | 0.94711648 | 0.19308912 | 1.00988693 | 0.65798472 | 1.64900012 | 1.32525006 | 966.685865 |
| Motif:rnd-6\_family-5252 | 1 | 1.08665856 | 1.08704113 | 0.97246484 | 1.00615962 | 0.92499155 | 0.64336508 | 0.75399167 | 1.31300857 | 1.15345515 | 961.558662 |
| Motif:rnd-1\_family-631 | 1 | 1.04411154 | 1.13836819 | 1.07153726 | 0.87583208 | 0.39821113 | 2.37585645 | 1.10307182 | 1.27423243 | 1.22048396 | 961.243006 |
| Motif:rnd-5\_family-1715 | 1 | 1.0856981 | 0.91728627 | 0.99716508 | 1.28065317 | 0.39596892 | 0.89346266 | 0.67944814 | 1.49936764 | 1.33566474 | 961.156509 |
| Motif:rnd-6\_family-1433 | 1 | 1.06227853 | 1.19302864 | 1.04462652 | 0.94684076 | 0.37900394 | 0.31713638 | 0.31210813 | 1.567825 | 1.2844146 | 958.781453 |
| Motif:rnd-1\_family-480 | 1 | 1.00754514 | 1.08059012 | 0.98387832 | 1.41544523 | 0.2515245 | 0.52892534 | 0.3553541 | 1.52338653 | 1.28637738 | 955.089772 |
| Motif:rnd-1\_family-1612 | 1 | 1.19934892 | 0.96321797 | 0.69347696 | 0.68191052 | 0.38081053 | 0.49725936 | 0.50460823 | 1.56139398 | 0.77753795 | 943.368375 |
| Motif:rnd-1\_family-1176 | 1 | 1.00830694 | 0.92119175 | 0.85849658 | 0.84231988 | 0.50836034 | 0.73822738 | 0.73608974 | 1.54976953 | 1.23811055 | 934.716967 |
| Motif:rnd-5\_family-4863 | 1 | 0.95264615 | 1.00631175 | 1.11337644 | 1.11652243 | 0.95510787 | 1.07838419 | 0.89395878 | 1.69833791 | 1.15417478 | 920.764983 |
| Motif:rnd-6\_family-1122 | 1 | 1.07716777 | 1.0960707 | 0.8770428 | 0.70026323 | 1.245276 | 0.72199163 | 0.70153024 | 1.47608258 | 1.18762611 | 919.251608 |
| Motif:rnd-6\_family-1823 | 1 | 0.96050074 | 0.92356353 | 0.83875258 | 1.07302578 | 0.56287369 | 0.66403414 | 0.59058811 | 1.4150801 | 1.17298164 | 918.980235 |
| Motif:rnd-1\_family-217 | 1 | 0.76319354 | 1.03749109 | 0.84692251 | 1.00145393 | 1.24803714 | 1.39925312 | 1.43549171 | 1.89531849 | 1.30264157 | 918.90665 |
| Motif:rnd-1\_family-1153 | 1 | 1.08356602 | 1.03151869 | 0.9882018 | 1.02963028 | 0.88384762 | 0.71388257 | 0.60758595 | 1.46804942 | 1.18228769 | 918.324572 |
| Motif:rnd-6\_family-1016 | 1 | 0.74620656 | 0.88336015 | 0.73645936 | 0.67479032 | 0.97746378 | 1.43360307 | 1.28518855 | 1.5694799 | 1.11816253 | 917.400993 |
| Motif:rnd-1\_family-970 | 1 | 1.04138871 | 1.03648774 | 0.98891784 | 1.03526778 | 0.64072684 | 1.3683153 | 0.75047758 | 1.3474699 | 1.14110409 | 916.08717 |
| Motif:rnd-6\_family-791 | 1 | 1.12174395 | 1.28930128 | 0.98264382 | 1.25136335 | 0.43884758 | 1.26296362 | 0.78842021 | 1.47402169 | 1.33509087 | 913.935411 |
| Motif:rnd-1\_family-818 | 1 | 1.01483246 | 1.16597191 | 1.01494609 | 1.14911517 | 0.65805913 | 0.61963834 | 0.63794916 | 1.57998624 | 1.32669287 | 907.332244 |
| Motif:rnd-5\_family-720 | 1 | 1.19317885 | 1.17107297 | 1.02931066 | 1.13945836 | 0.37204966 | 0.38594424 | 0.35060422 | 1.28400676 | 1.08819978 | 907.267259 |
| Motif:rnd-6\_family-2474 | 1 | 0.78511339 | 0.98942452 | 0.93838755 | 1.16629037 | 0.58573821 | 0.55325843 | 0.57642774 | 1.67277859 | 1.35018749 | 895.011732 |
| Motif:rnd-1\_family-1719 | 1 | 1.0240262 | 1.16256216 | 0.94597985 | 1.01942859 | 0.39401833 | 0.60101027 | 0.50618226 | 1.37948108 | 1.20470823 | 893.92451 |
| Motif:rnd-1\_family-696 | 1 | 0.82614139 | 1.06308895 | 1.00205425 | 1.11421198 | 1.55379028 | 1.44628296 | 1.42313372 | 1.86242121 | 1.46018766 | 890.370035 |
| Motif:rnd-6\_family-84 | 1 | 0.83350071 | 0.9286221 | 0.86324464 | 0.96352977 | 1.03358619 | 0.89600166 | 0.90428946 | 1.37496931 | 0.97660654 | 889.120748 |
| Motif:rnd-1\_family-63 | 1 | 1.06257944 | 1.1078524 | 0.98712478 | 0.94125054 | 0.43418673 | 0.94206794 | 0.72483479 | 1.59233019 | 1.22023909 | 887.878079 |
| Motif:rnd-6\_family-122 | 1 | 0.92235256 | 0.9328121 | 0.85155213 | 1.05047317 | 1.03819044 | 0.73203074 | 0.70165547 | 1.30304723 | 1.02462512 | 886.649931 |
| Motif:rnd-1\_family-52 | 1 | 0.88146638 | 0.97292034 | 0.89620517 | 0.92896621 | 0.57369837 | 0.84745565 | 0.69964274 | 1.29279355 | 1.19250734 | 880.325506 |
| Motif:rnd-1\_family-525 | 1 | 1.04818294 | 1.33268135 | 0.93020321 | 0.86490089 | 0.26972301 | 0.77410219 | 0.49749538 | 1.0489639 | 0.93847762 | 874.911288 |
| Motif:rnd-5\_family-295 | 1 | 1.0540313 | 1.21225586 | 0.99563083 | 1.11331581 | 0.45513362 | 0.78356254 | 0.5910843 | 1.61699838 | 1.32686682 | 873.619463 |
| Motif:rnd-5\_family-1961 | 1 | 1.25302454 | 1.21180097 | 1.82661517 | 1.26865648 | 2.0255417 | 1.40405163 | 1.49711676 | 2.13674421 | 1.31117845 | 873.566096 |
| Motif:rnd-5\_family-1287 | 1 | 1.14063133 | 1.09264324 | 0.97521496 | 1.04933978 | 0.26382228 | 0.73320426 | 0.45225779 | 1.51632762 | 1.23777649 | 867.026199 |
| Motif:rnd-1\_family-393 | 1 | 0.5502649 | 0.95559856 | 0.6164636 | 0.66064209 | 1.10102175 | 1.11196861 | 1.29211809 | 1.89416083 | 1.17346124 | 866.205429 |
| Motif:rnd-6\_family-1324 | 1 | 1.01135275 | 0.94063454 | 0.90244716 | 0.82263799 | 0.09671869 | 0.94554515 | 0.48365275 | 0.95797492 | 0.75569553 | 864.489261 |
| Motif:rnd-1\_family-877 | 1 | 1.16708927 | 1.32659656 | 1.04497851 | 1.11756409 | 0.24089715 | 0.39896856 | 0.3437304 | 1.46130215 | 1.21179364 | 859.829925 |
| Motif:rnd-6\_family-2423 | 1 | 1.14805912 | 1.11936392 | 1.06700321 | 1.1523883 | 0.19205278 | 0.58269456 | 0.41390256 | 1.34386358 | 1.03215564 | 857.683133 |
| Motif:rnd-1\_family-1665 | 1 | 0.92293487 | 1.06671943 | 0.96167414 | 0.80149217 | 0.44855095 | 2.13481341 | 1.19385137 | 1.35718847 | 1.10206826 | 852.384884 |
| Motif:rnd-1\_family-114 | 1 | 0.77669769 | 0.80166621 | 0.84637815 | 0.66342499 | 1.09415741 | 1.25821319 | 1.18060379 | 0.30964066 | 1.04450672 | 849.169976 |
| Motif:rnd-1\_family-307 | 1 | 0.9349557 | 1.03048852 | 0.89469699 | 1.01930599 | 0.93326508 | 1.30284641 | 1.19338684 | 1.55749583 | 1.25667449 | 844.406783 |
| Motif:rnd-5\_family-366 | 1 | 0.99527581 | 1.15428259 | 0.99259349 | 0.91218331 | 0.34516646 | 2.32758828 | 1.09381696 | 1.32672317 | 1.28838783 | 840.47883 |
| Motif:rnd-1\_family-106 | 1 | 1.0005779 | 1.20251537 | 0.96512523 | 0.84288443 | 1.40481571 | 1.38532307 | 1.48047543 | 1.95127467 | 1.54344127 | 839.954 |
| Motif:rnd-1\_family-1969 | 1 | 1.09542784 | 1.04856175 | 0.88148949 | 0.84383499 | 0.44591873 | 0.38148056 | 0.46567643 | 1.34914588 | 1.05029147 | 833.641215 |
| Motif:rnd-5\_family-2309 | 1 | 1.09546411 | 1.04586298 | 0.98679647 | 1.1203587 | 1.01970146 | 0.73636467 | 0.76261504 | 1.38234567 | 1.15586265 | 826.559698 |
| Motif:rnd-1\_family-1128 | 1 | 1.15940232 | 1.1061872 | 1.02898039 | 1.24016397 | 0.42810958 | 0.77262637 | 0.62454948 | 1.49291957 | 1.3775463 | 821.847886 |
| Motif:rnd-5\_family-3928 | 1 | 0.93137347 | 0.96367984 | 0.87225085 | 0.8909844 | 0.44862031 | 0.37527066 | 0.39323919 | 1.34374343 | 0.9823328 | 820.334882 |
| Motif:rnd-5\_family-1163 | 1 | 0.75770435 | 1.18600748 | 0.99833536 | 0.94977597 | 1.16292985 | 1.2774749 | 1.08385575 | 1.53206398 | 1.2738581 | 819.964343 |
| Motif:rnd-6\_family-15255 | 1 | 0.42883239 | 1.24473247 | 0.9339081 | 1.48645177 | 1.16670609 | 1.18208951 | 1.38934563 | 1.01572227 | 0.37232269 | 819.260165 |
| Motif:rnd-5\_family-4592 | 1 | 1.0892122 | 1.06533295 | 1.01396626 | 1.04334002 | 0.33315571 | 0.8930809 | 0.53819687 | 1.38798957 | 1.19764057 | 816.571416 |
| Motif:rnd-6\_family-10845 | 1 | 1.34696408 | 0.95753223 | 0.90074915 | 1.27090039 | 0.15320377 | 0.24699028 | 0.23926417 | 0.92086871 | 0.81742326 | 807.098321 |
| Motif:rnd-6\_family-8717 | 1 | 1.06384879 | 1.16749717 | 0.75735714 | 1.026024 | 0.134146 | 0.30729106 | 0.28075253 | 1.38153137 | 1.46237605 | 804.58068 |
| Motif:rnd-1\_family-54 | 1 | 0.69248116 | 0.96703061 | 0.65268376 | 0.78052055 | 1.06689229 | 1.36251825 | 1.22533932 | 1.2844526 | 1.10897478 | 799.161934 |
| Motif:rnd-1\_family-685 | 1 | 1.00475412 | 1.11053008 | 0.99792347 | 0.88766327 | 0.43032639 | 2.49658769 | 1.06713762 | 1.35409988 | 1.20821862 | 795.146827 |
| Motif:rnd-1\_family-51 | 1 | 1.07730497 | 1.23979486 | 1.07092933 | 0.9327062 | 0.30254049 | 1.29030645 | 0.71821336 | 1.22845725 | 1.16570771 | 792.949556 |
| Motif:rnd-1\_family-740 | 1 | 1.13428869 | 1.16981511 | 0.97285726 | 1.03227586 | 0.24307595 | 1.11254191 | 0.62653036 | 1.43600074 | 1.14833182 | 790.423355 |
| Motif:rnd-5\_family-739 | 1 | 0.9027247 | 0.942309 | 0.88384002 | 0.86210695 | 1.85473396 | 0.97405514 | 1.08764452 | 1.49132226 | 1.05200289 | 789.728724 |
| Motif:rnd-6\_family-7981 | 1 | 0.68998686 | 1.04241106 | 0.92213113 | 1.01403909 | 1.109374 | 1.15520329 | 1.21014383 | 1.88092416 | 1.25031143 | 782.221293 |
| Motif:rnd-1\_family-162 | 1 | 0.97579945 | 1.10351666 | 0.98218184 | 1.17727776 | 1.07176104 | 2.16221498 | 1.49274489 | 1.41952383 | 1.1316792 | 781.964839 |
| Motif:rnd-1\_family-795 | 1 | 1.15455999 | 1.16686351 | 1.06447319 | 1.21435371 | 0.37015364 | 0.39885871 | 0.34578601 | 1.63609107 | 1.43006777 | 781.613668 |
| Motif:rnd-6\_family-6797 | 1 | 1.24756874 | 1.2664616 | 1.08186021 | 1.23580785 | 0.40568585 | 0.70333916 | 0.59741252 | 1.64638897 | 1.38223592 | 779.996306 |
| Motif:rnd-5\_family-97 | 1 | 0.76171384 | 0.95112154 | 0.84451264 | 0.90871577 | 0.67931987 | 0.67788121 | 0.68955318 | 1.38122118 | 1.03781299 | 778.532819 |
| Motif:rnd-5\_family-208 | 1 | 1.03954281 | 1.16921672 | 1.08839078 | 0.91480043 | 0.30434916 | 2.48297721 | 1.06989225 | 1.87604095 | 1.68864448 | 777.95988 |
| Motif:rnd-1\_family-384 | 1 | 0.75218216 | 0.9080366 | 0.76139209 | 0.97710213 | 1.53378472 | 1.27313522 | 1.63842593 | 1.88492536 | 1.29278439 | 776.566288 |
| Motif:rnd-5\_family-312 | 1 | 1.09450077 | 1.21713257 | 1.04573965 | 1.14710041 | 0.3497992 | 0.94896723 | 0.63054633 | 1.68280824 | 1.2956468 | 773.025079 |
| Motif:rnd-5\_family-478 | 1 | 0.90366791 | 1.10940869 | 0.94663861 | 1.28981929 | 4.19635305 | 1.65533274 | 1.84642906 | 1.40151736 | 1.02863586 | 769.135118 |
| Motif:rnd-1\_family-1837 | 1 | 0.86545279 | 0.67382262 | 0.97069806 | 0.97969025 | 1.38052037 | 1.33321852 | 1.37713776 | 0.1483055 | 1.07374248 | 767.530774 |
| Motif:rnd-6\_family-2714 | 1 | 1.06465997 | 1.00545394 | 1.15132916 | 1.13298446 | 0.40572568 | 0.36622602 | 0.3927183 | 1.56256226 | 1.23169075 | 766.286357 |
| Motif:rnd-1\_family-413 | 1 | 0.81657051 | 1.0317063 | 0.91438268 | 0.94826026 | 2.00962461 | 1.472302 | 1.83131641 | 2.36447077 | 1.51178306 | 762.274207 |
| Motif:rnd-1\_family-1081 | 1 | 1.17910602 | 1.08421083 | 0.94102731 | 1.00769433 | 0.41296317 | 0.58070025 | 0.44192766 | 1.3311139 | 1.15408545 | 762.183005 |
| Motif:rnd-5\_family-2637 | 1 | 1.11293603 | 1.12238644 | 1.13852646 | 1.41939503 | 1.53592882 | 1.29807672 | 1.69040109 | 0.85832781 | 1.16625613 | 759.453475 |
| Motif:rnd-1\_family-443 | 1 | 1.15476858 | 1.0658946 | 1.02778322 | 0.93498564 | 0.45370025 | 0.51935866 | 0.54687562 | 1.5005893 | 1.23516407 | 758.481208 |
| Motif:rnd-6\_family-4859 | 1 | 0.88305003 | 1.0381258 | 0.95513977 | 0.9037611 | 1.27851947 | 1.40840255 | 1.35997987 | 1.87877149 | 1.35048582 | 757.744236 |
| Motif:rnd-1\_family-378 | 1 | 0.63420031 | 1.10485447 | 0.86284685 | 0.83192686 | 1.57592922 | 1.41894478 | 1.59743667 | 2.0747559 | 1.29426902 | 757.413593 |
| Motif:rnd-1\_family-722 | 1 | 1.23964997 | 1.07100931 | 0.91504277 | 1.08431804 | 0.17625135 | 0.400771 | 0.30369043 | 0.95912377 | 0.78327158 | 751.028377 |
| Motif:rnd-1\_family-32 | 1 | 0.90219309 | 0.96426626 | 1.00314026 | 1.2354183 | 1.56160548 | 1.40684057 | 1.56904638 | 1.88056668 | 1.48372602 | 746.963487 |
| Motif:rnd-6\_family-4300 | 1 | 0.90754612 | 1.01830707 | 0.86478265 | 0.8978118 | 1.25948729 | 1.28087899 | 1.17094971 | 1.72732827 | 1.19569805 | 745.774135 |
| Motif:rnd-1\_family-1160 | 1 | 0.6752708 | 0.90869102 | 0.71467685 | 0.85568196 | 1.33475096 | 1.44443079 | 1.52796826 | 1.88035815 | 1.28798446 | 744.545867 |
| Motif:rnd-1\_family-705 | 1 | 0.88744502 | 1.01586114 | 0.90815384 | 1.0642231 | 0.67395224 | 0.75997222 | 0.83223696 | 1.71867676 | 1.30211668 | 729.47755 |
| Motif:rnd-3\_family-857 | 1 | 0.93353444 | 0.92667514 | 1.07920136 | 1.31415198 | 1.50475486 | 1.42018504 | 1.51720639 | 1.61824858 | 1.30455338 | 727.359599 |
| Motif:rnd-5\_family-760 | 1 | 1.25676267 | 1.37199671 | 0.92905843 | 1.53338222 | 0.1081045 | 0.21315695 | 0.1904203 | 0.84959555 | 0.65241863 | 723.340001 |
| Motif:rnd-5\_family-2384 | 1 | 0.66211222 | 1.00702605 | 0.83763532 | 0.84337886 | 0.93797549 | 0.92748525 | 1.14537084 | 1.68594975 | 1.32930263 | 722.677194 |
| Motif:rnd-1\_family-60 | 1 | 0.84583479 | 0.87632312 | 0.86875349 | 1.09270286 | 1.15705099 | 1.19457988 | 1.12652382 | 1.43139761 | 1.16409737 | 721.804201 |
| Motif:rnd-6\_family-5703 | 1 | 1.07301135 | 1.0594825 | 1.1211156 | 1.25106542 | 0.93956026 | 0.93049484 | 0.89107741 | 1.66821952 | 1.21213597 | 721.048108 |
| Motif:rnd-1\_family-19 | 1 | 0.9170793 | 1.01856571 | 0.98842623 | 1.06309375 | 0.8125644 | 0.81543812 | 0.82284299 | 1.59228615 | 1.28358434 | 719.065923 |
| Motif:rnd-6\_family-891 | 1 | 1.1758831 | 1.29387352 | 0.9285793 | 1.16590481 | 0.23126919 | 0.59456525 | 0.34752796 | 1.33710862 | 1.31602681 | 715.630599 |
| Motif:rnd-6\_family-6270 | 1 | 1.00167428 | 1.04811094 | 0.92912192 | 0.79595822 | 0.63104807 | 0.69412966 | 0.76934541 | 1.55517469 | 1.28091968 | 713.796222 |
| Motif:rnd-1\_family-45 | 1 | 1.02211523 | 1.02365286 | 1.1716439 | 1.07218207 | 1.25662225 | 1.3189825 | 1.05942596 | 0.49371323 | 1.00936666 | 708.260787 |
| Motif:rnd-1\_family-868 | 1 | 1.19349667 | 1.13027561 | 0.79088513 | 0.84456804 | 0.54454902 | 0.42539962 | 0.4297275 | 0.88435589 | 0.71812173 | 707.807104 |
| Motif:rnd-1\_family-1060 | 1 | 0.96558267 | 0.83496192 | 0.93346964 | 0.9966441 | 0.5800452 | 0.5394131 | 0.61726407 | 1.61266712 | 1.3074433 | 705.474286 |
| Motif:rnd-1\_family-418 | 1 | 1.01903485 | 1.0493699 | 0.93663496 | 1.04547698 | 0.75353386 | 0.60867324 | 0.69252408 | 1.66620465 | 1.22244924 | 705.062153 |
| Motif:rnd-5\_family-2877 | 1 | 1.09511464 | 0.92503632 | 1.03221616 | 1.08360544 | 1.17529108 | 0.67999062 | 0.71139216 | 1.39344692 | 1.02523594 | 704.290362 |
| Motif:rnd-5\_family-1816 | 1 | 0.98120106 | 1.07121547 | 1.21638223 | 1.14169504 | 1.7859023 | 1.52503495 | 1.59920051 | 1.43738001 | 1.34894186 | 704.161094 |
| Motif:rnd-5\_family-2257 | 1 | 1.11549801 | 1.08197658 | 0.96645048 | 1.07250452 | 0.34247955 | 0.50635439 | 0.47618761 | 1.51321215 | 1.22620227 | 703.405577 |
| Motif:rnd-1\_family-1529 | 1 | 0.99977433 | 1.06433859 | 0.98710404 | 1.02417097 | 0.5144409 | 0.74358743 | 0.56500039 | 1.43739673 | 1.20333069 | 699.6711 |
| Motif:rnd-1\_family-72 | 1 | 0.88925405 | 1.17647781 | 0.84270893 | 1.02680124 | 1.44669996 | 1.63015473 | 1.36931559 | 1.81703067 | 1.32360314 | 699.027434 |
| Motif:rnd-6\_family-12504 | 1 | 1.06610823 | 1.12202176 | 1.01969753 | 0.91684939 | 0.38661166 | 0.94015727 | 0.64038172 | 1.43990126 | 1.20752244 | 697.121356 |
| Motif:rnd-6\_family-902 | 1 | 1.31964153 | 1.08239706 | 0.88551655 | 1.15703494 | 0.79271985 | 0.83470838 | 0.86723125 | 1.34881177 | 1.04856855 | 695.554989 |
| Motif:rnd-5\_family-5765 | 1 | 0.98739223 | 0.97739498 | 1.01569232 | 1.04378714 | 0.43945152 | 0.54774355 | 0.46707346 | 1.53646727 | 1.08852045 | 693.940424 |
| Motif:rnd-1\_family-607 | 1 | 0.98878451 | 0.85457746 | 1.04763975 | 1.20431898 | 1.40478413 | 1.44417594 | 1.49756599 | 0.5617706 | 1.12933553 | 693.920601 |
| Motif:rnd-1\_family-139 | 1 | 0.75540936 | 0.82456605 | 0.77533244 | 0.98313811 | 1.19408201 | 1.00259738 | 1.17403738 | 1.03747359 | 0.83325491 | 692.310224 |
| Motif:rnd-1\_family-553 | 1 | 0.55206617 | 1.05640968 | 0.77981598 | 1.13293164 | 0.67470376 | 0.64582273 | 0.82076681 | 1.16368974 | 0.7272166 | 691.470743 |
| Motif:rnd-6\_family-3172 | 1 | 0.86461019 | 0.8190649 | 0.88335938 | 0.84232474 | 0.57042716 | 0.58621488 | 0.67655061 | 1.39568096 | 1.01140474 | 687.130109 |
| Motif:rnd-6\_family-3044 | 1 | 0.81004271 | 1.22993298 | 0.94664687 | 1.31961723 | 0.96130058 | 0.6267938 | 0.69648515 | 1.96260794 | 1.43592251 | 685.337689 |
| Motif:rnd-6\_family-9354 | 1 | 1.03402447 | 1.19569432 | 0.93888824 | 0.75606662 | 0.3662295 | 1.27565544 | 0.78177978 | 1.49501554 | 1.2661749 | 684.798122 |
| Motif:rnd-1\_family-949 | 1 | 0.90197754 | 1.02044157 | 0.97337674 | 1.13080798 | 1.13115197 | 0.98893963 | 1.00961538 | 1.52655577 | 1.17502445 | 684.516882 |
| Motif:rnd-1\_family-676 | 1 | 0.83908486 | 0.90833758 | 0.8670294 | 1.00915471 | 0.75493596 | 0.60628377 | 0.63811522 | 1.29604977 | 0.96388204 | 683.314846 |
| Motif:rnd-1\_family-637 | 1 | 1.09051707 | 1.0346672 | 1.11868154 | 0.96324667 | 0.65415366 | 0.76073999 | 0.62147482 | 1.55163241 | 1.33585031 | 681.04213 |
| Motif:rnd-1\_family-526 | 1 | 1.38145568 | 1.5119246 | 2.13351552 | 1.53821188 | 2.9893718 | 1.64249407 | 1.98717293 | 2.96282499 | 1.8054264 | 677.198711 |
| Motif:rnd-1\_family-355 | 1 | 1.12507713 | 1.33043782 | 0.95772674 | 0.9553339 | 0.32115485 | 2.27506121 | 1.34202259 | 1.62092642 | 1.35558378 | 676.844471 |
| Motif:rnd-5\_family-3082 | 1 | 1.1961963 | 1.04758678 | 0.96664222 | 0.81952979 | 0.54818742 | 0.62529438 | 0.51350398 | 1.45486307 | 1.23112871 | 676.680962 |
| Motif:rnd-6\_family-2596 | 1 | 0.69999598 | 0.77459165 | 0.7236004 | 0.66387207 | 1.08421727 | 1.04472665 | 1.3044556 | 1.82462302 | 1.15213815 | 672.875584 |
| Motif:rnd-1\_family-220 | 1 | 0.89153429 | 1.05991232 | 0.96101352 | 1.01218289 | 0.9125267 | 0.89561264 | 0.80355109 | 1.61974508 | 1.1748378 | 672.730551 |
| Motif:rnd-1\_family-519 | 1 | 1.03312422 | 1.13350294 | 1.06080424 | 1.1585228 | 0.25782404 | 1.87613475 | 1.04867374 | 1.32145671 | 1.23607612 | 672.173484 |
| Motif:rnd-4\_family-1470 | 1 | 1.01468909 | 1.11419365 | 0.93855357 | 1.72175659 | 0.30377887 | 0.7913954 | 0.52574409 | 1.45807234 | 1.29290639 | 666.629175 |
| Motif:rnd-6\_family-3082 | 1 | 1.12060766 | 1.10935167 | 1.03805979 | 1.21704828 | 0.42407524 | 0.7737337 | 0.64707508 | 1.4743526 | 1.09676922 | 666.295562 |
| Motif:rnd-1\_family-499 | 1 | 0.7711315 | 0.93150006 | 0.80906994 | 0.95343442 | 1.05077836 | 1.17709631 | 1.21263673 | 1.55013198 | 1.11257341 | 665.68642 |
| Motif:rnd-5\_family-1928 | 1 | 1.05613114 | 1.09680013 | 0.98925594 | 0.82055226 | 0.48262299 | 2.65470759 | 1.44123321 | 1.63517004 | 1.18410494 | 664.584167 |
| Motif:rnd-1\_family-237 | 1 | 1.18360609 | 1.04439573 | 1.17014724 | 0.72594324 | 1.52577395 | 1.37082816 | 1.14798471 | 0.16469381 | 0.95131105 | 664.293818 |
| Motif:rnd-1\_family-110 | 1 | 1.06303788 | 0.98073029 | 1.06378597 | 1.26991001 | 1.13043368 | 1.19755265 | 0.93789794 | 1.16779297 | 1.16003026 | 664.012582 |
| Motif:rnd-1\_family-461 | 1 | 1.0599787 | 1.32263327 | 1.04840038 | 0.86480687 | 0.2279261 | 2.38005328 | 1.24601077 | 1.51303182 | 1.2945514 | 663.093557 |
| Motif:rnd-1\_family-169 | 1 | 0.92813501 | 0.86031275 | 0.88126374 | 0.89254635 | 0.89006959 | 1.02457152 | 0.94406321 | 1.27107653 | 1.0914641 | 662.848329 |
| Motif:rnd-1\_family-1863 | 1 | 0.9824104 | 1.0010196 | 0.86966652 | 1.31917605 | 0.14185084 | 0.52220169 | 0.33319398 | 1.35920674 | 1.25417945 | 661.177357 |
| Motif:rnd-1\_family-1926 | 1 | 0.98983695 | 1.0662338 | 0.86412751 | 0.75626341 | 0.14844313 | 1.42987901 | 0.62123275 | 1.18754316 | 1.0764537 | 657.416515 |
| Motif:rnd-6\_family-7307 | 1 | 1.43867196 | 1.42649193 | 1.16142109 | 1.28056769 | 0.3526448 | 0.43430782 | 0.34692642 | 1.4420289 | 1.28662047 | 656.613864 |
| Motif:rnd-6\_family-4269 | 1 | 1.06810529 | 1.21500544 | 0.93490453 | 0.69750081 | 0.15272473 | 1.30938248 | 0.71051485 | 1.69142161 | 1.46696218 | 653.883158 |
| Motif:rnd-1\_family-1300 | 1 | 0.95754515 | 1.0661482 | 0.89294772 | 1.06737769 | 0.56232365 | 0.72531701 | 0.60224171 | 1.40245823 | 1.14714357 | 652.742445 |
| Motif:rnd-5\_family-4365 | 1 | 1.25964332 | 1.09804355 | 0.9680747 | 0.93375346 | 0.34901692 | 0.44735666 | 0.4430233 | 1.46173442 | 1.28079908 | 652.061178 |
| Motif:rnd-4\_family-1676 | 1 | 0.78876489 | 0.81812003 | 0.74090319 | 0.56495743 | 0.7323523 | 0.80293885 | 0.97001436 | 1.60351504 | 1.14552347 | 650.612112 |
| Motif:rnd-1\_family-772 | 1 | 1.06873534 | 1.18181614 | 1.04755669 | 0.89843365 | 0.3153598 | 3.07034472 | 1.24556307 | 1.25526948 | 1.2468407 | 649.776636 |
| Motif:rnd-5\_family-2063 | 1 | 1.00387888 | 1.04199902 | 1.01469339 | 1.12679182 | 0.65046221 | 0.6458099 | 0.5408415 | 1.57699833 | 1.30145778 | 648.75589 |
| Motif:rnd-1\_family-1017 | 1 | 1.06984646 | 1.19199496 | 0.97089599 | 1.13101045 | 0.3036106 | 0.39263106 | 0.2749206 | 1.53021466 | 1.25294564 | 647.367565 |
| Motif:rnd-1\_family-503 | 1 | 0.85078915 | 0.83387548 | 0.84734483 | 0.82822749 | 0.59062205 | 0.6912138 | 0.68977517 | 0.82700902 | 0.978311 | 646.546814 |
| Motif:rnd-6\_family-60 | 1 | 0.99168179 | 0.99024537 | 0.94597046 | 1.04742692 | 0.68371507 | 0.99907196 | 0.78485044 | 1.36772786 | 1.14220542 | 645.88636 |
| Motif:rnd-6\_family-4493 | 1 | 0.92614836 | 1.04342941 | 1.01044383 | 0.8637768 | 3.64966752 | 1.00936557 | 1.46714345 | 1.43910131 | 1.11088452 | 645.673914 |
| Motif:rnd-6\_family-7183 | 1 | 0.96903161 | 1.06931845 | 0.89865153 | 0.89459833 | 5.11126887 | 2.44279443 | 2.792768 | 1.65549262 | 1.34537037 | 642.48651 |
| Motif:rnd-6\_family-906 | 1 | 0.75123032 | 1.20958269 | 1.09921777 | 1.15950551 | 1.94566011 | 1.33548558 | 1.52048402 | 0.59602039 | 0.80718264 | 640.99364 |
| Motif:rnd-1\_family-628 | 1 | 0.76685605 | 1.03040777 | 0.99606251 | 0.88748348 | 1.36435782 | 1.5999697 | 2.01366314 | 2.77826154 | 2.00873655 | 639.630691 |
| Motif:rnd-4\_family-2329 | 1 | 0.95881968 | 1.09800787 | 1.03672019 | 0.81341292 | 0.4184271 | 0.4747226 | 0.42148937 | 1.41713363 | 1.15158959 | 633.905126 |
| Motif:rnd-5\_family-195 | 1 | 0.92967075 | 1.14520612 | 1.22137985 | 1.25621475 | 1.55322087 | 1.43112413 | 1.35809807 | 0.72749313 | 1.10258913 | 633.126004 |
| Motif:rnd-1\_family-150 | 1 | 0.85786922 | 0.91009737 | 1.00025632 | 0.88407229 | 1.07746712 | 1.05770668 | 1.09727866 | 1.03279037 | 1.13004467 | 633.106142 |
| Motif:rnd-6\_family-9012 | 1 | 1.21536174 | 1.12169781 | 1.03673395 | 0.78721565 | 0.13768369 | 0.53820104 | 0.3707057 | 1.33040159 | 1.04691863 | 632.714351 |
| Motif:rnd-1\_family-744 | 1 | 0.71194313 | 1.0439221 | 0.90580692 | 0.98846302 | 1.5064762 | 1.25547106 | 1.49306471 | 1.88165186 | 1.23329098 | 630.459929 |
| Motif:rnd-6\_family-5458 | 1 | 0.92242141 | 0.95884136 | 0.95509245 | 1.01072887 | 1.10397134 | 1.25772191 | 1.09406911 | 1.60347047 | 1.13867329 | 630.05567 |
| Motif:rnd-1\_family-252 | 1 | 1.05167092 | 1.07539272 | 0.91146181 | 0.81176558 | 0.40521113 | 0.64289108 | 0.4956304 | 1.40497055 | 1.22262708 | 628.954665 |
| Motif:rnd-1\_family-377 | 1 | 0.75142918 | 0.73929663 | 0.86045772 | 0.99853755 | 1.18463703 | 1.23156826 | 1.34202545 | 0.29555138 | 1.04435764 | 628.025813 |
| Motif:rnd-1\_family-814 | 1 | 0.90579577 | 1.00712594 | 0.9124942 | 0.91217852 | 0.80104936 | 1.20840212 | 0.9260891 | 1.3694812 | 1.19462569 | 622.832227 |
| Motif:rnd-1\_family-97 | 1 | 0.98899814 | 1.195843 | 1.1293013 | 1.18765697 | 1.49093525 | 1.46127038 | 1.31586599 | 0.46762537 | 1.22145376 | 620.874882 |
| Motif:rnd-6\_family-5078 | 1 | 1.05682531 | 1.08487634 | 1.0159675 | 1.1081975 | 0.88689007 | 0.47578824 | 0.58211821 | 1.38825911 | 1.0413709 | 618.421809 |
| Motif:rnd-1\_family-206 | 1 | 1.01853317 | 0.93773984 | 0.89084711 | 0.92745981 | 1.66724082 | 1.05385345 | 1.26726894 | 1.49338177 | 1.0953505 | 617.616861 |
| Motif:rnd-6\_family-6331 | 1 | 1.0084977 | 0.92860431 | 0.84378106 | 1.00031592 | 0.33784514 | 0.76526129 | 0.53119454 | 1.35713022 | 1.10578778 | 616.951955 |
| Motif:rnd-1\_family-708 | 1 | 0.95716073 | 1.03204527 | 0.93145082 | 0.97101071 | 0.84806352 | 1.75974398 | 1.27442916 | 1.36350423 | 1.19168055 | 616.130772 |
| Motif:rnd-1\_family-361 | 1 | 1.06276988 | 1.05164097 | 0.97502219 | 1.0323078 | 0.81663335 | 0.76802199 | 0.6975074 | 1.66708125 | 1.20710241 | 615.470063 |
| Motif:rnd-1\_family-134 | 1 | 1.08705554 | 1.15164548 | 1.04990393 | 1.09316883 | 0.51643438 | 1.87452969 | 1.09861561 | 1.53257596 | 1.28983928 | 613.535491 |
| Motif:rnd-5\_family-1513 | 1 | 1.22431347 | 1.26469169 | 0.99766627 | 1.22683248 | 0.41839121 | 0.63512941 | 0.44500947 | 1.49121889 | 1.21458911 | 613.533059 |
| Motif:rnd-1\_family-157 | 1 | 1.01862135 | 1.16399382 | 1.01322157 | 1.10624901 | 0.46243114 | 0.93741566 | 0.59899643 | 1.4539547 | 1.21556022 | 608.013255 |
| Motif:rnd-5\_family-6090 | 1 | 0.67383876 | 1.26501335 | 0.86545156 | 1.11374799 | 2.29566782 | 1.25705259 | 1.49300223 | 1.60926861 | 0.95323795 | 607.983038 |
| Motif:rnd-1\_family-534 | 1 | 1.04251582 | 1.24182225 | 1.08230266 | 1.09283304 | 0.73183108 | 1.44661924 | 0.91544359 | 1.52513323 | 1.24329861 | 607.197882 |
| Motif:rnd-6\_family-9788 | 1 | 1.01123506 | 1.22597452 | 1.32084677 | 1.83213625 | 1.24480211 | 1.13564155 | 1.37482546 | 0.67754466 | 1.08849831 | 606.548976 |
| Motif:rnd-1\_family-473 | 1 | 0.70962783 | 0.45076502 | 0.78321777 | 0.76616921 | 1.13875976 | 1.12282084 | 1.29736515 | 0.51372847 | 1.0835434 | 605.828862 |
| Motif:rnd-1\_family-1979 | 1 | 0.995807 | 1.061928 | 0.97225732 | 0.82153494 | 0.22522398 | 0.2956561 | 0.24365401 | 1.25859385 | 0.94200643 | 605.083554 |
| Motif:rnd-1\_family-1751 | 1 | 1.18503303 | 1.26252636 | 1.07524366 | 1.08006709 | 0.69519644 | 0.81747865 | 0.70539641 | 1.55808744 | 1.22578842 | 604.190004 |
| Motif:rnd-6\_family-1731 | 1 | 0.66165576 | 1.02670846 | 0.73546289 | 0.9434302 | 1.22013107 | 1.19795171 | 1.2360315 | 1.59939788 | 1.20342458 | 602.95753 |
| Motif:rnd-6\_family-7508 | 1 | 1.1370011 | 1.27370225 | 0.99704764 | 0.99088589 | 0.33074755 | 0.92429814 | 0.48872169 | 1.63272964 | 1.33972437 | 600.6895 |
| Motif:rnd-6\_family-3794 | 1 | 0.83493905 | 1.05742167 | 1.0110632 | 1.12423013 | 1.10827705 | 1.47125496 | 1.31048225 | 1.69079931 | 1.2698327 | 600.45471 |
| Motif:rnd-6\_family-3678 | 1 | 1.00723147 | 1.07255353 | 0.54406252 | 1.20298287 | 0.30275069 | 0.42569773 | 0.33953847 | 0.44296308 | 0.32776567 | 598.747688 |
| Motif:rnd-6\_family-5796 | 1 | 1.14840505 | 1.22936547 | 1.03184475 | 0.98806074 | 0.95171902 | 0.95970786 | 0.76859471 | 1.55874938 | 1.29281011 | 597.958206 |
| Motif:rnd-6\_family-71 | 1 | 0.88341218 | 0.98976787 | 0.89835924 | 1.17465558 | 1.50029141 | 1.50936033 | 1.63102197 | 1.91127978 | 1.30254483 | 597.396315 |
| Motif:rnd-4\_family-490 | 1 | 0.59084553 | 1.21933881 | 0.91850478 | 0.8490462 | 1.13787945 | 1.11806303 | 1.33719217 | 1.73343492 | 1.38056519 | 593.619379 |
| Motif:rnd-6\_family-3067 | 1 | 1.0373992 | 1.1078286 | 0.99836033 | 1.27586453 | 0.86516668 | 1.33127562 | 1.03237514 | 1.35597148 | 1.12212317 | 592.339705 |
| Motif:rnd-1\_family-338 | 1 | 0.91654285 | 1.13824885 | 0.91529646 | 0.92713659 | 0.55840193 | 1.69686311 | 1.12304357 | 1.5052718 | 1.18618063 | 592.205343 |
| Motif:rnd-6\_family-1081 | 1 | 0.88734642 | 0.82865358 | 0.78338416 | 0.76820404 | 1.20448002 | 1.29126268 | 1.45448287 | 2.10074024 | 1.33599496 | 591.741368 |
| Motif:rnd-6\_family-5454 | 1 | 1.31570264 | 1.0877412 | 1.06431009 | 1.00272685 | 0.24060953 | 0.70235139 | 0.48297233 | 1.52107124 | 1.23312811 | 591.140415 |
| Motif:rnd-6\_family-5274 | 1 | 1.15435729 | 1.03425254 | 0.98244391 | 0.9715339 | 0.45443697 | 0.85095105 | 0.61876689 | 1.36097079 | 1.19028916 | 590.447826 |
| Motif:rnd-5\_family-36 | 1 | 0.79433285 | 0.81986854 | 0.80253465 | 0.7819638 | 1.24751921 | 1.38659285 | 1.51483938 | 1.89363088 | 1.43146119 | 587.828904 |
| Motif:rnd-4\_family-927 | 1 | 1.09484172 | 1.152317 | 1.04870919 | 0.95662262 | 0.86843779 | 2.20975009 | 1.52089247 | 1.75433275 | 1.2431934 | 587.381694 |
| Motif:rnd-5\_family-164 | 1 | 1.38785085 | 1.40680395 | 1.22135255 | 1.13089107 | 0.40090826 | 0.53255717 | 0.53797395 | 1.67731366 | 1.35740166 | 587.109378 |
| Motif:rnd-1\_family-246 | 1 | 0.99391899 | 1.17000808 | 0.861593 | 0.76244458 | 0.48095702 | 1.5620709 | 1.03076124 | 1.42899248 | 1.12478444 | 583.98878 |
| Motif:rnd-5\_family-4690 | 1 | 1.05808272 | 1.09561106 | 0.90192755 | 1.30609079 | 0.60439417 | 0.81387135 | 0.72841664 | 1.60204392 | 1.30255707 | 583.199447 |
| Motif:rnd-1\_family-429 | 1 | 0.99713864 | 1.13343119 | 0.9402059 | 0.82835432 | 0.33388717 | 1.23897543 | 0.74873688 | 1.15996005 | 1.02096215 | 576.852627 |
| Motif:rnd-5\_family-1400 | 1 | 0.95411503 | 1.05165882 | 0.88282731 | 1.13334253 | 1.96533776 | 1.54112999 | 1.6993499 | 1.82941963 | 1.31068495 | 576.834192 |
| Motif:rnd-1\_family-305 | 1 | 1.06000483 | 1.07174055 | 0.88930418 | 1.31164418 | 0.22005294 | 0.2620118 | 0.27036955 | 1.55749445 | 1.2837169 | 575.585072 |
| Motif:rnd-1\_family-142 | 1 | 0.85515922 | 0.96727342 | 0.87343825 | 0.94210755 | 0.77255122 | 0.75697502 | 0.76537418 | 1.42467053 | 0.89154406 | 575.028157 |
| Motif:rnd-1\_family-10 | 1 | 0.79392851 | 1.05380431 | 0.92084291 | 1.05348102 | 1.53712555 | 1.30144693 | 1.47571242 | 1.74755388 | 1.2535722 | 572.195375 |
| Motif:rnd-6\_family-1900 | 1 | 1.28108251 | 1.10486435 | 0.84798203 | 0.98858981 | 0.35164239 | 0.36384196 | 0.36213733 | 1.15972298 | 0.84627973 | 571.635286 |
| Motif:rnd-1\_family-1741 | 1 | 1.13888615 | 1.53944842 | 1.01591252 | 1.09345534 | 0.63574213 | 0.76563375 | 0.78144467 | 1.67559091 | 1.3454592 | 568.582719 |
| Motif:rnd-1\_family-1373 | 1 | 1.26706471 | 1.14862952 | 0.93517995 | 0.90225287 | 0.41644845 | 0.51658646 | 0.42540244 | 1.40793616 | 1.26379014 | 568.253036 |
| Motif:rnd-1\_family-905 | 1 | 1.00666033 | 1.00136562 | 0.86656482 | 0.87278637 | 0.3236705 | 0.29309111 | 0.28927004 | 1.43163048 | 1.15505493 | 567.216766 |
| Motif:rnd-1\_family-138 | 1 | 0.98685838 | 1.1145028 | 0.9025001 | 1.20010987 | 0.78450269 | 1.1239635 | 0.96998085 | 1.38443468 | 1.1492741 | 565.255082 |
| Motif:rnd-6\_family-1544 | 1 | 1.11800147 | 1.12338632 | 0.94270736 | 0.94311235 | 0.37066656 | 0.74233728 | 0.5436501 | 1.54893007 | 1.15364626 | 564.937034 |
| Motif:rnd-4\_family-88 | 1 | 0.88729085 | 0.91090371 | 1.02443693 | 1.12741403 | 1.34842837 | 1.54686182 | 1.51026524 | 0.96942152 | 1.1895058 | 558.918569 |
| Motif:rnd-5\_family-5961 | 1 | 0.94836893 | 1.02225522 | 0.99404855 | 0.92050847 | 0.52161753 | 0.83587037 | 0.73593132 | 1.47271817 | 1.21570535 | 558.329097 |
| Motif:rnd-1\_family-56 | 1 | 0.85987802 | 0.9239219 | 0.99234781 | 1.00990623 | 1.24412391 | 2.19001188 | 1.60005662 | 1.17579363 | 1.0329608 | 554.766555 |
| Motif:rnd-1\_family-109 | 1 | 0.9450971 | 1.09118512 | 0.98663821 | 0.95355092 | 0.4948915 | 0.53612572 | 0.51444574 | 1.52544176 | 1.25538199 | 547.381073 |
| Motif:rnd-4\_family-236 | 1 | 0.5785868 | 0.97755694 | 0.87117661 | 1.55368552 | 1.54928622 | 1.28468225 | 1.290539 | 1.83951699 | 1.3161342 | 546.816414 |
| Motif:rnd-1\_family-855 | 1 | 1.25354968 | 1.15028725 | 0.94644214 | 0.908957 | 0.22309644 | 0.94818259 | 0.71440106 | 1.43722598 | 1.15042567 | 541.256835 |
| Motif:rnd-1\_family-6 | 1 | 0.93310618 | 1.08495055 | 1.0752048 | 1.13343893 | 1.06804917 | 1.11507705 | 1.05425034 | 1.06564775 | 1.03795843 | 540.679219 |
| Motif:rnd-6\_family-5246 | 1 | 0.98273245 | 1.05175105 | 1.10114719 | 1.00253334 | 1.55234307 | 1.3286934 | 1.38333583 | 1.80846575 | 1.12932731 | 538.836681 |
| Motif:rnd-4\_family-281 | 1 | 0.77331655 | 0.96379994 | 0.87347313 | 0.85629356 | 1.00253605 | 0.82284949 | 1.03854349 | 1.46931229 | 1.02407902 | 533.509262 |
| Motif:rnd-5\_family-233 | 1 | 0.92217223 | 0.92905945 | 0.99570594 | 1.20349289 | 1.40657152 | 1.40417806 | 1.44725917 | 1.43014465 | 1.20428028 | 533.320742 |
| Motif:rnd-6\_family-30 | 1 | 0.76674735 | 0.87622729 | 0.82770979 | 0.80082008 | 1.23490006 | 1.09715618 | 1.29393706 | 1.60613813 | 1.11036702 | 530.885539 |
| Motif:rnd-1\_family-184 | 1 | 1.34009229 | 0.9672354 | 0.98425589 | 1.02250547 | 0.31923455 | 0.53454843 | 0.47996624 | 1.38952166 | 1.04383332 | 528.11992 |
| Motif:rnd-5\_family-1416 | 1 | 0.90235145 | 1.18709721 | 0.98108779 | 0.99043404 | 1.73290469 | 1.45501269 | 1.40164176 | 0.75555815 | 1.07850999 | 527.02433 |
| Motif:rnd-6\_family-4724 | 1 | 1.25685788 | 1.20405286 | 0.84225027 | 1.55669914 | 0.25845987 | 0.61599156 | 0.54906545 | 1.38783229 | 1.19005994 | 525.834859 |
| Motif:rnd-5\_family-1129 | 1 | 0.75341862 | 1.3734175 | 1.07273537 | 1.41807548 | 1.76583323 | 1.50031394 | 1.62084024 | 2.21920812 | 1.70317597 | 525.0644 |
| Motif:rnd-6\_family-6087 | 1 | 0.99361661 | 0.89360727 | 1.03438003 | 1.15947219 | 0.55179994 | 0.92839997 | 0.70777694 | 1.34648152 | 1.13045704 | 522.557686 |
| Motif:rnd-6\_family-7467 | 1 | 1.15766854 | 1.00310338 | 0.91154626 | 0.89834138 | 0.25350231 | 0.36653848 | 0.32254999 | 1.31381685 | 1.07374115 | 522.399411 |
| Motif:rnd-4\_family-2163 | 1 | 0.74237604 | 0.85422019 | 0.80626651 | 0.790447 | 1.22169229 | 1.27161847 | 1.1707268 | 0.22398121 | 0.9187386 | 518.272188 |
| Motif:rnd-1\_family-614 | 1 | 0.78296472 | 0.66217739 | 0.80254908 | 0.85456933 | 0.84243907 | 1.12574093 | 1.09808425 | 0.39854398 | 0.99280469 | 517.961674 |
| Motif:rnd-1\_family-238 | 1 | 0.95939693 | 1.07117978 | 0.8659631 | 0.79769935 | 0.84830715 | 1.92722564 | 1.48987633 | 1.34651184 | 1.16685787 | 517.714532 |
| Motif:rnd-1\_family-1879 | 1 | 1.18940043 | 1.17362865 | 0.95537411 | 0.74665402 | 0.31395806 | 1.35424875 | 0.81678854 | 1.616927 | 1.2762954 | 516.957778 |
| Motif:rnd-5\_family-3626 | 1 | 1.28657132 | 1.20962451 | 1.08483735 | 0.99136972 | 0.48068012 | 1.42763545 | 1.0497923 | 1.67990561 | 1.42775504 | 515.857083 |
| Motif:rnd-1\_family-20 | 1 | 1.10506938 | 1.12623045 | 1.018628 | 1.0312771 | 0.19118085 | 3.08457037 | 1.3180584 | 1.59971866 | 1.27887583 | 515.245979 |
| Motif:rnd-1\_family-800 | 1 | 1.27414169 | 1.19185506 | 1.02649032 | 1.01718927 | 0.22548533 | 0.64041218 | 0.39360241 | 1.47740276 | 1.29540977 | 510.574058 |
| Motif:rnd-1\_family-809 | 1 | 1.17086231 | 1.17404999 | 1.10378871 | 1.4401336 | 0.64878032 | 1.51500071 | 1.18687824 | 1.66049784 | 1.33982386 | 508.85243 |
| Motif:rnd-1\_family-68 | 1 | 0.67267339 | 1.09102162 | 0.71284492 | 0.79942355 | 1.29678036 | 1.23447623 | 1.3380982 | 1.64086477 | 1.25242352 | 507.955258 |
| Motif:rnd-6\_family-7630 | 1 | 0.8926912 | 1.21302422 | 0.90528458 | 0.94247903 | 0.40384377 | 4.31744591 | 1.73362088 | 1.31063199 | 1.07867224 | 506.693946 |
| Motif:rnd-6\_family-4336 | 1 | 1.20214073 | 1.10394903 | 0.90705462 | 1.20999945 | 1.82618747 | 1.41314629 | 1.86744455 | 1.64670895 | 1.17230674 | 503.253197 |
| Motif:rnd-5\_family-5461 | 1 | 0.95960823 | 1.16537511 | 0.80079491 | 1.10198309 | 0.44296663 | 0.38525962 | 0.41988181 | 1.41903177 | 1.11582621 | 503.166304 |
| Motif:rnd-6\_family-11892 | 1 | 0.98311239 | 1.05022774 | 0.99909001 | 1.06787974 | 0.41554295 | 0.46461483 | 0.51931248 | 1.46293648 | 1.2874864 | 502.235497 |
