## Supplementary material for "A novel eukaryotic RdRP-dependent small RNA pathway represses antiviral immunity by controlling an ERK pathway component in the black-legged tick": TE_RPM.html

 


### TE mapping table (RPM)

| ID | GFP | Ago-16 | Ago-30 | Ago-96 | Ago-78 | Aub | AGO3-1 | AGO3-2 | RdRP1 | RdRP3 | SUM\_KD |
| --- | --- | --- | --- | --- | --- | --- | --- | --- | --- | --- | --- |
| Motif:rnd-6\_family-7248 | 17611.2843 | 13581.5995 | 9938.68959 | 15006.9386 | 15121.7048 | 19986.1119 | 19984.6323 | 22616.0995 | 1602.20315 | 18142.2342 | 153591.498 |
| Motif:rnd-5\_family-601 | 13936.7818 | 5164.01949 | 14067.4792 | 19288.6146 | 16385.2223 | 22855.0677 | 18445.968 | 19507.0399 | 20496.357 | 3315.3927 | 153461.943 |
| Motif:rnd-1\_family-272 | 13794.7193 | 9497.77892 | 10850.4659 | 12028.3305 | 12999.5211 | 16327.5364 | 16799.8649 | 18139.2291 | 1448.37264 | 14577.5213 | 126463.34 |
| Motif:rnd-1\_family-366 | 11704.4138 | 8187.87824 | 6308.49831 | 9412.69108 | 9616.01951 | 13007.2567 | 13529.1944 | 15459.5654 | 1118.92427 | 12080.5159 | 100424.958 |
| Motif:rnd-5\_family-4359 | 8632.05618 | 7306.38734 | 5690.31121 | 7330.53186 | 8686.57634 | 10497.3479 | 10671.3101 | 10654.6667 | 552.343556 | 8070.59768 | 78092.1288 |
| Motif:rnd-1\_family-1035 | 9119.64638 | 6240.19996 | 5208.86472 | 6780.0959 | 6809.83294 | 10284.3163 | 10312.4553 | 11432.8108 | 748.736413 | 9111.29232 | 76048.2511 |
| Motif:rnd-6\_family-4937 | 5650.25209 | 2400.11236 | 5048.31437 | 6640.79066 | 6101.04778 | 10635.0967 | 7025.21238 | 8724.05352 | 9027.38745 | 1720.64513 | 62972.9124 |
| Motif:rnd-1\_family-1958 | 7139.02454 | 5073.3417 | 4166.18138 | 5713.65216 | 5364.21736 | 7964.61448 | 8133.85214 | 9067.39505 | 726.418828 | 7201.41776 | 60550.1154 |
| Motif:rnd-1\_family-1103 | 6351.09524 | 4679.26706 | 4918.3623 | 5121.68452 | 5968.64165 | 8334.04898 | 7509.56735 | 8009.27987 | 454.419191 | 5870.75188 | 57217.118 |
| Motif:rnd-6\_family-5847 | 4018.01572 | 2912.31351 | 3810.81098 | 4103.40213 | 3888.45631 | 12369.6908 | 6311.37455 | 8903.05013 | 6258.48153 | 2804.04266 | 55379.6384 |
| Motif:rnd-1\_family-1744 | 5601.27118 | 3823.63705 | 2608.43323 | 4345.62391 | 4955.24174 | 6218.15856 | 6601.25724 | 8041.96455 | 549.543163 | 5929.65137 | 48674.782 |
| Motif:rnd-1\_family-178 | 3873.41386 | 2090.19564 | 4414.3643 | 3677.2345 | 3522.46606 | 6308.66873 | 5255.79665 | 6250.61689 | 6334.2234 | 2345.18426 | 44072.1643 |
| Motif:rnd-5\_family-5812 | 5074.28691 | 5992.01101 | 5771.02655 | 4159.48848 | 4914.22712 | 3306.77316 | 1436.39132 | 1737.51604 | 5358.40583 | 4229.28098 | 41979.4074 |
| Motif:rnd-6\_family-423 | 4264.18417 | 5135.59099 | 5128.81984 | 4322.96884 | 5341.56092 | 1077.3692 | 1678.57281 | 1544.38267 | 6557.69799 | 5087.36175 | 40138.5092 |
| Motif:rnd-4\_family-2208 | 3777.13087 | 4458.25126 | 4632.30806 | 3948.97871 | 4157.56529 | 774.253315 | 4331.01031 | 2434.90071 | 5599.31122 | 4751.38565 | 38865.0954 |
| Motif:rnd-6\_family-6510 | 4270.99577 | 3632.67145 | 6607.93936 | 3913.35482 | 3536.69135 | 4737.47088 | 3917.58605 | 1560.49314 | 2621.62069 | 1834.98877 | 36633.8123 |
| Motif:rnd-6\_family-5880 | 2500.36555 | 2919.43393 | 2829.3468 | 2321.7823 | 2618.26494 | 4414.29981 | 2084.49395 | 2568.32723 | 3564.71233 | 2814.16489 | 28635.1917 |
| Motif:rnd-6\_family-1977 | 2324.59057 | 2440.54832 | 2621.92494 | 2301.01339 | 2701.61832 | 3178.7298 | 1464.66783 | 1792.41216 | 3674.32291 | 2810.9256 | 25310.7538 |
| Motif:rnd-6\_family-4509 | 2290.04921 | 2713.58753 | 2461.71402 | 2305.7508 | 2480.95743 | 1459.57424 | 2291.86263 | 1597.79809 | 3297.42147 | 2686.35748 | 23585.0729 |
| Motif:rnd-5\_family-460 | 2309.00364 | 2652.93177 | 2760.2917 | 2351.09446 | 3128.27885 | 1429.36642 | 1252.12888 | 1228.05827 | 3618.83498 | 2708.13212 | 23438.1211 |
| Motif:rnd-6\_family-46 | 2259.38246 | 2643.66232 | 2787.24546 | 2308.45327 | 2440.66984 | 401.510978 | 2243.22896 | 1317.28268 | 3287.15956 | 2727.37587 | 22415.9714 |
| Motif:rnd-6\_family-44 | 2153.68858 | 2494.91896 | 2652.17611 | 2199.48812 | 2284.19808 | 412.744392 | 2371.46364 | 1327.08829 | 3082.53219 | 2600.98674 | 21579.2851 |
| Motif:rnd-6\_family-3340 | 2394.96713 | 1933.21722 | 2045.21679 | 1954.14788 | 1868.33834 | 2781.3336 | 2747.22884 | 3002.61413 | 457.637543 | 2322.39228 | 21507.0938 |
| Motif:rnd-1\_family-474 | 2071.89389 | 2378.9098 | 2534.11285 | 2080.50768 | 2211.29148 | 379.095937 | 2319.19807 | 1339.53018 | 3012.08935 | 2544.49926 | 20871.1285 |
| Motif:rnd-6\_family-4733 | 1436.49228 | 978.181422 | 2734.92725 | 1173.25014 | 4769.81237 | 2751.52636 | 1897.31724 | 1985.40727 | 1251.38282 | 1221.90355 | 20200.2007 |
| Motif:rnd-1\_family-423 | 2155.41896 | 2865.53659 | 2347.60291 | 2102.17032 | 2578.17554 | 932.79863 | 741.627033 | 749.178636 | 3115.39359 | 2318.56186 | 19906.4641 |
| Motif:rnd-6\_family-2886 | 2055.25024 | 2163.75601 | 2180.69334 | 1664.70177 | 1745.79079 | 874.38197 | 1750.65757 | 1313.98152 | 3235.58825 | 2487.72029 | 19472.5218 |
| Motif:rnd-4\_family-408 | 2095.8081 | 1620.01113 | 1371.45533 | 1867.22008 | 1923.34303 | 2526.34029 | 2599.25485 | 2877.81447 | 234.993155 | 2261.36178 | 19377.6022 |
| Motif:rnd-6\_family-620 | 1229.8482 | 738.783665 | 1622.58762 | 1887.97381 | 1743.7386 | 3777.0607 | 3040.61134 | 2422.94205 | 2047.89639 | 684.459924 | 19195.9023 |
| Motif:rnd-6\_family-7598 | 1421.64256 | 1494.15049 | 1736.90579 | 1402.93182 | 1386.36275 | 2161.61941 | 2029.4034 | 1776.01704 | 2244.2773 | 1798.20909 | 17451.5197 |
| Motif:rnd-6\_family-4398 | 1311.95568 | 1434.82103 | 1362.49457 | 1230.25501 | 1621.14551 | 3589.30842 | 1240.29065 | 1765.51128 | 2161.75555 | 1614.9708 | 17332.5085 |
| Motif:rnd-6\_family-8470 | 1857.72007 | 2033.12819 | 2107.21039 | 1698.27379 | 1786.5394 | 987.078973 | 892.88009 | 867.581497 | 2694.15946 | 2171.41527 | 17095.9871 |
| Motif:rnd-1\_family-1822 | 1864.4463 | 1901.72871 | 1835.26058 | 1823.6847 | 2027.08273 | 826.779098 | 980.388196 | 1138.41223 | 2695.39416 | 1618.22282 | 16711.3995 |
| Motif:rnd-4\_family-2384 | 1765.89771 | 2296.27468 | 2278.24279 | 1937.38609 | 2488.67536 | 294.257135 | 505.809787 | 433.817368 | 2473.85788 | 2148.18286 | 16622.4017 |
| Motif:rnd-6\_family-4430 | 1974.01742 | 2493.58673 | 2175.8746 | 1862.41415 | 1530.49183 | 983.107201 | 856.99119 | 748.915816 | 2159.84711 | 1779.57619 | 16564.8222 |
| Motif:rnd-1\_family-926 | 1745.37726 | 1957.88242 | 1810.96728 | 1651.72039 | 1482.76293 | 907.723543 | 787.982433 | 841.671283 | 2658.27595 | 2099.53409 | 15943.8976 |
| Motif:rnd-6\_family-9480 | 1586.49548 | 1556.40918 | 1700.13984 | 1456.80713 | 1499.26512 | 1711.24943 | 1015.30407 | 1083.42714 | 2311.78822 | 1819.0414 | 15739.927 |
| Motif:rnd-6\_family-1478 | 1572.24111 | 1748.72496 | 1692.70536 | 1463.88981 | 1753.40586 | 1142.52542 | 655.741946 | 683.262121 | 2244.40753 | 1763.40457 | 14720.3087 |
| Motif:rnd-4\_family-513 | 2089.65301 | 1749.77178 | 2252.40808 | 1504.76406 | 904.813921 | 1439.32231 | 1694.45358 | 1533.57367 | 153.281467 | 1301.2073 | 14623.2492 |
| Motif:rnd-1\_family-1320 | 1544.18444 | 1929.23575 | 1669.43199 | 1425.47854 | 1215.38786 | 1559.43622 | 667.814671 | 824.234201 | 1793.96057 | 1373.7962 | 14002.9605 |
| Motif:rnd-4\_family-328 | 1363.17697 | 1462.33833 | 1384.00308 | 1391.94291 | 1321.26838 | 1243.77171 | 1185.22919 | 1121.03372 | 1255.1684 | 1479.30043 | 13207.2331 |
| Motif:rnd-1\_family-417 | 1265.23216 | 1520.78133 | 1584.78212 | 1318.22738 | 1379.90108 | 293.029508 | 1214.51347 | 735.039159 | 1837.23482 | 1526.51604 | 12675.2571 |
| Motif:rnd-5\_family-631 | 1555.3863 | 1774.51832 | 1667.27804 | 1430.28875 | 1447.83383 | 628.888519 | 711.342101 | 684.248711 | 1495.9623 | 1139.7362 | 12535.4831 |
| Motif:rnd-5\_family-2803 | 908.549372 | 768.451986 | 838.473998 | 1032.71201 | 706.084118 | 2331.34112 | 1288.11912 | 2207.29038 | 1407.10704 | 800.615965 | 12288.7451 |
| Motif:rnd-6\_family-997 | 1116.52366 | 1279.5666 | 1163.40732 | 1272.50562 | 1423.02432 | 700.152189 | 652.73919 | 694.516795 | 1870.31655 | 1499.69615 | 11672.4484 |
| Motif:rnd-6\_family-2652 | 1353.58508 | 1607.26528 | 1570.33185 | 1378.79477 | 1467.22084 | 273.303091 | 274.1896 | 269.241171 | 1886.5279 | 1531.9383 | 11612.3979 |
| Motif:rnd-6\_family-1271 | 1208.88915 | 1599.37417 | 1444.10212 | 1291.625 | 1248.26452 | 269.394772 | 656.282142 | 510.925085 | 1802.1027 | 1506.09703 | 11537.0567 |
| Motif:rnd-6\_family-1406 | 1118.38868 | 1110.55702 | 1144.77127 | 1025.21332 | 999.372007 | 1390.85695 | 649.964421 | 862.114493 | 1742.50404 | 1343.1569 | 11386.8991 |
| Motif:rnd-6\_family-2888 | 1346.2773 | 1580.18525 | 1560.35043 | 1106.95881 | 1163.1395 | 153.215182 | 288.543375 | 257.358621 | 1777.89522 | 1440.54787 | 10674.4716 |
| Motif:rnd-1\_family-528 | 1075.65549 | 1083.18786 | 1143.56305 | 1059.38926 | 1119.60301 | 802.575081 | 598.071802 | 612.704485 | 1527.93437 | 1167.38491 | 10190.0693 |
| Motif:rnd-5\_family-37 | 973.478015 | 949.872574 | 821.469687 | 1041.9355 | 968.798738 | 1372.12829 | 1277.84848 | 1272.53329 | 251.931932 | 1078.91282 | 10008.9093 |
| Motif:rnd-6\_family-4238 | 911.824518 | 819.359085 | 859.308678 | 942.562814 | 982.082468 | 1180.5468 | 1306.6921 | 1237.04649 | 415.400701 | 1113.78413 | 9768.60779 |
| Motif:rnd-1\_family-36 | 971.123758 | 1130.23765 | 1108.01942 | 1014.8869 | 986.91438 | 787.443242 | 569.055095 | 609.153043 | 1436.69264 | 1141.60773 | 9755.13386 |
| Motif:rnd-5\_family-3305 | 1059.15807 | 1303.21148 | 1029.24138 | 1205.57762 | 1246.17683 | 290.390419 | 459.197558 | 487.916725 | 1407.76852 | 1199.64561 | 9688.28421 |
| Motif:rnd-5\_family-2342 | 1088.75102 | 1073.21765 | 1081.8118 | 990.941029 | 1123.04715 | 466.168445 | 598.84022 | 549.223005 | 1568.87932 | 1132.15933 | 9673.03897 |
| Motif:rnd-6\_family-4486 | 912.163328 | 1127.49026 | 852.155554 | 1025.25003 | 1247.11923 | 1236.49955 | 1257.01063 | 977.722208 | 114.691987 | 745.969524 | 9496.07231 |
| Motif:rnd-5\_family-154 | 1000.5593 | 1040.98385 | 1124.08784 | 943.147839 | 976.392248 | 989.844673 | 499.750612 | 524.658219 | 1312.97187 | 1078.17384 | 9490.57029 |
| Motif:rnd-5\_family-3344 | 969.681549 | 1156.83445 | 951.356572 | 935.117584 | 981.400928 | 692.901003 | 418.12352 | 463.914413 | 1358.02957 | 1192.42453 | 9119.78412 |
| Motif:rnd-1\_family-873 | 728.895839 | 641.499898 | 774.441301 | 664.374624 | 734.41001 | 1040.24917 | 846.916275 | 988.563045 | 1274.16327 | 910.093906 | 8603.60733 |
| Motif:rnd-6\_family-3084 | 936.183497 | 1180.15702 | 1204.61398 | 966.62621 | 981.349783 | 210.864271 | 273.43263 | 247.313759 | 1353.97112 | 1095.3683 | 8449.88057 |
| Motif:rnd-1\_family-412 | 991.088132 | 544.964322 | 1044.31831 | 768.107836 | 1078.9697 | 668.346263 | 636.502213 | 820.864522 | 1135.33525 | 715.677994 | 8404.17454 |
| Motif:rnd-6\_family-7503 | 700.232461 | 554.24993 | 634.586299 | 569.635675 | 925.078095 | 876.024667 | 865.189971 | 937.398254 | 1340.29006 | 915.216 | 8317.90141 |
| Motif:rnd-1\_family-136 | 929.926918 | 753.348894 | 523.873368 | 810.117958 | 854.986928 | 1089.51073 | 1065.28208 | 1194.21983 | 114.575385 | 923.671306 | 8259.5134 |
| Motif:rnd-1\_family-9 | 841.261279 | 932.254961 | 897.094756 | 867.532095 | 832.970548 | 710.703264 | 624.280873 | 568.934136 | 1002.46315 | 955.392759 | 8232.88782 |
| Motif:rnd-5\_family-494 | 785.357058 | 755.981745 | 947.14321 | 739.280576 | 961.671674 | 634.651821 | 623.575126 | 655.3131 | 1172.73511 | 807.557731 | 8083.26715 |
| Motif:rnd-5\_family-2195 | 754.632465 | 718.653895 | 852.435128 | 651.790357 | 786.825376 | 576.892986 | 935.678602 | 671.770222 | 1177.20835 | 882.435042 | 8008.32242 |
| Motif:rnd-1\_family-201 | 785.625602 | 648.139277 | 760.043818 | 808.400429 | 717.323366 | 1379.30908 | 888.402749 | 1005.77245 | 261.573749 | 748.327098 | 8002.91762 |
| Motif:rnd-6\_family-1239 | 1057.07333 | 1377.09083 | 1304.06419 | 904.13401 | 1395.2548 | 443.779645 | 175.971737 | 211.143837 | 501.323616 | 411.566526 | 7781.40252 |
| Motif:rnd-6\_family-8629 | 801.589189 | 880.299299 | 826.693115 | 747.75152 | 821.976442 | 603.160115 | 490.593466 | 511.937473 | 1083.61908 | 873.876409 | 7641.49611 |
| Motif:rnd-1\_family-697 | 717.575733 | 892.802316 | 789.280015 | 709.856992 | 972.197858 | 726.166128 | 404.616085 | 447.62287 | 1012.27584 | 818.744038 | 7491.13788 |
| Motif:rnd-6\_family-159 | 680.763822 | 748.564145 | 753.072235 | 663.329799 | 692.650695 | 1019.99799 | 479.645795 | 587.663231 | 1007.24042 | 811.480517 | 7444.40865 |
| Motif:rnd-6\_family-2254 | 855.120339 | 990.000368 | 978.702558 | 798.174119 | 1021.32487 | 173.258995 | 363.252907 | 285.207758 | 1048.16585 | 898.636139 | 7411.8439 |
| Motif:rnd-4\_family-1108 | 819.05145 | 692.60886 | 595.481961 | 709.876507 | 781.927898 | 816.481091 | 837.514285 | 942.967523 | 346.466625 | 811.528595 | 7353.9048 |
| Motif:rnd-1\_family-1829 | 793.173722 | 702.314727 | 789.709311 | 873.663269 | 807.253112 | 475.064475 | 390.951615 | 468.87156 | 1241.3781 | 787.868083 | 7330.24797 |
| Motif:rnd-6\_family-2808 | 761.765759 | 810.709176 | 832.520047 | 722.77595 | 797.756905 | 401.442224 | 470.041107 | 452.203153 | 1115.87535 | 884.976444 | 7250.06612 |
| Motif:rnd-3\_family-43 | 872.964927 | 630.390555 | 495.667387 | 619.768836 | 654.83234 | 911.105457 | 969.503173 | 1146.9292 | 76.3204986 | 855.817254 | 7233.29962 |
| Motif:rnd-6\_family-970 | 699.092892 | 675.503382 | 659.03606 | 674.214795 | 652.529431 | 847.407629 | 749.061623 | 836.816568 | 547.306999 | 821.631347 | 7162.60073 |
| Motif:rnd-6\_family-10566 | 758.552518 | 892.906981 | 861.836176 | 743.333121 | 910.272037 | 359.727935 | 354.809707 | 365.115449 | 948.119816 | 739.797056 | 6934.4708 |
| Motif:rnd-6\_family-10254 | 739.684701 | 761.22947 | 798.11487 | 673.997189 | 832.612803 | 415.19696 | 414.230281 | 409.958618 | 973.633217 | 755.946965 | 6774.60508 |
| Motif:rnd-5\_family-557 | 743.85529 | 883.079074 | 797.672696 | 828.686575 | 748.32326 | 265.450331 | 279.64939 | 263.311014 | 1067.33272 | 856.25163 | 6733.61198 |
| Motif:rnd-1\_family-81 | 555.55352 | 450.531847 | 539.412082 | 542.016827 | 579.566604 | 742.725103 | 807.788991 | 725.981936 | 999.929189 | 707.836772 | 6651.34287 |
| Motif:rnd-1\_family-311 | 543.648642 | 587.416065 | 703.017394 | 740.466692 | 703.859558 | 954.080166 | 854.894715 | 736.307812 | 136.768278 | 628.840387 | 6589.29971 |
| Motif:rnd-6\_family-4387 | 510.824413 | 351.06681 | 529.499751 | 496.50511 | 515.573233 | 1253.3735 | 690.204511 | 962.240855 | 774.65268 | 398.677339 | 6482.6182 |
| Motif:rnd-1\_family-438 | 670.570508 | 822.885246 | 794.437009 | 663.915133 | 740.84995 | 138.383866 | 557.001395 | 359.532891 | 879.722291 | 729.640873 | 6356.93916 |
| Motif:rnd-6\_family-9529 | 614.892196 | 658.133298 | 663.410041 | 620.99033 | 717.96546 | 483.816741 | 389.20983 | 479.829159 | 957.699376 | 766.886093 | 6352.83253 |
| Motif:rnd-1\_family-118 | 674.62574 | 532.077919 | 395.771131 | 531.481417 | 666.469156 | 768.290664 | 811.432222 | 953.361501 | 209.92721 | 749.038355 | 6292.47531 |
| Motif:rnd-5\_family-2258 | 599.093489 | 696.987555 | 749.789013 | 675.933017 | 613.800064 | 395.328814 | 342.514495 | 348.131026 | 1011.33841 | 821.114228 | 6254.03011 |
| Motif:rnd-5\_family-1029 | 724.018411 | 721.966457 | 737.992996 | 643.689792 | 642.200019 | 301.041829 | 326.029041 | 371.371899 | 956.991468 | 786.096915 | 6211.39883 |
| Motif:rnd-1\_family-496 | 673.008372 | 525.688979 | 414.885604 | 593.941075 | 683.977 | 781.536966 | 778.875977 | 877.354163 | 62.7216072 | 650.834395 | 6042.82414 |
| Motif:rnd-6\_family-6830 | 673.675886 | 571.914831 | 533.142903 | 560.803795 | 612.409447 | 624.913644 | 583.725186 | 699.285247 | 441.557731 | 527.350762 | 5828.77943 |
| Motif:rnd-6\_family-40 | 597.019866 | 653.796965 | 652.188357 | 615.825309 | 639.072577 | 376.22248 | 401.94733 | 371.968689 | 833.705175 | 686.377297 | 5828.12404 |
| Motif:rnd-6\_family-2044 | 1219.11235 | 1281.67878 | 1169.02951 | 637.779508 | 885.735548 | 21.8619933 | 106.315878 | 64.4261275 | 201.704267 | 170.582764 | 5758.22673 |
| Motif:rnd-1\_family-117 | 517.407444 | 277.58936 | 610.842012 | 528.480357 | 573.19361 | 930.786719 | 609.697467 | 788.725236 | 571.154474 | 327.357929 | 5735.23461 |
| Motif:rnd-1\_family-463 | 549.773759 | 678.954956 | 622.980015 | 543.443247 | 675.838069 | 433.615309 | 340.736225 | 339.607416 | 840.015373 | 710.0319 | 5734.99627 |
| Motif:rnd-1\_family-940 | 612.377641 | 608.060977 | 658.797413 | 550.114394 | 620.06866 | 436.577976 | 303.525424 | 322.570744 | 869.967902 | 709.646786 | 5691.70792 |
| Motif:rnd-1\_family-96 | 566.161183 | 555.431435 | 433.79319 | 608.247362 | 512.091781 | 765.766968 | 722.726593 | 691.405145 | 177.971275 | 585.625443 | 5619.22038 |
| Motif:rnd-6\_family-8871 | 642.711387 | 690.141822 | 731.431931 | 615.577093 | 640.610589 | 239.051019 | 260.108773 | 263.61369 | 821.252251 | 669.380503 | 5573.87906 |
| Motif:rnd-5\_family-2768 | 615.246542 | 770.655735 | 687.529785 | 607.009886 | 664.216441 | 137.505966 | 260.635201 | 221.968361 | 858.450727 | 701.965311 | 5525.18396 |
| Motif:rnd-6\_family-2915 | 547.865218 | 624.122885 | 571.131214 | 562.593276 | 528.029565 | 346.061948 | 330.99571 | 352.813571 | 870.218591 | 666.653414 | 5400.48539 |
| Motif:rnd-5\_family-558 | 528.40146 | 612.862829 | 619.363107 | 548.447323 | 596.352788 | 263.584998 | 321.550372 | 271.287942 | 801.914769 | 670.391528 | 5234.15712 |
| Motif:rnd-1\_family-1531 | 468.421475 | 535.973107 | 587.027027 | 520.648284 | 453.033782 | 246.852273 | 660.360895 | 422.401187 | 722.256746 | 611.178647 | 5228.15343 |
| Motif:rnd-5\_family-150 | 480.95423 | 564.096478 | 567.277702 | 581.501379 | 648.969266 | 369.622169 | 438.135886 | 410.145702 | 537.269297 | 589.027323 | 5186.99943 |
| Motif:rnd-4\_family-1114 | 560.907594 | 682.818822 | 620.375297 | 527.971788 | 630.451669 | 155.304246 | 257.339632 | 234.751304 | 814.443388 | 646.648952 | 5131.01269 |
| Motif:rnd-1\_family-646 | 499.403597 | 394.884475 | 494.895154 | 434.379085 | 477.974298 | 812.745372 | 591.49676 | 587.864431 | 239.065492 | 519.320463 | 5052.02913 |
| Motif:rnd-6\_family-13609 | 491.617277 | 403.952016 | 389.082227 | 474.871879 | 543.549012 | 767.514905 | 679.614862 | 676.781494 | 41.4907943 | 571.287047 | 5039.76151 |
| Motif:rnd-4\_family-200 | 543.411672 | 643.281921 | 650.928939 | 508.946196 | 589.848544 | 182.194434 | 229.151477 | 204.570613 | 795.350641 | 668.228108 | 5015.91255 |
| Motif:rnd-5\_family-1630 | 583.837765 | 664.705482 | 587.413707 | 553.934617 | 610.374858 | 149.033366 | 257.56684 | 223.079443 | 751.98264 | 613.35639 | 4995.28511 |
| Motif:rnd-1\_family-1111 | 571.143945 | 721.577933 | 635.788646 | 536.312603 | 458.323596 | 82.1550974 | 269.898338 | 242.117687 | 780.740764 | 640.711299 | 4938.76991 |
| Motif:rnd-1\_family-1207 | 596.860389 | 691.255975 | 695.128396 | 510.363471 | 473.011027 | 88.9439854 | 196.838082 | 169.49213 | 832.851893 | 669.550465 | 4924.29581 |
| Motif:rnd-1\_family-862 | 511.740425 | 403.705346 | 293.627381 | 415.910601 | 561.509286 | 632.658848 | 640.192158 | 777.107445 | 96.2613739 | 569.133695 | 4901.84656 |
| Motif:rnd-1\_family-391 | 509.98552 | 416.000035 | 300.905115 | 484.302402 | 486.066539 | 651.551517 | 649.865075 | 738.745892 | 50.9293055 | 607.079705 | 4895.43111 |
| Motif:rnd-1\_family-213 | 573.709746 | 633.407027 | 421.928035 | 433.057802 | 547.764602 | 317.424415 | 341.347276 | 340.278933 | 769.025099 | 441.481999 | 4819.42493 |
| Motif:rnd-1\_family-1792 | 526.821392 | 581.644203 | 537.122621 | 477.395706 | 476.191718 | 249.494044 | 259.253393 | 245.645475 | 812.015509 | 602.635394 | 4768.21945 |
| Motif:rnd-1\_family-425 | 510.756055 | 442.755168 | 439.250797 | 461.510652 | 519.747817 | 469.962445 | 428.732035 | 499.395941 | 435.524258 | 550.511021 | 4758.14619 |
| Motif:rnd-1\_family-839 | 489.725052 | 520.241319 | 536.985409 | 443.78101 | 484.210244 | 292.003248 | 200.683356 | 216.061827 | 717.751477 | 597.142489 | 4498.58543 |
| Motif:rnd-5\_family-930 | 495.255736 | 550.736317 | 543.882793 | 447.067916 | 492.236761 | 132.505274 | 315.812704 | 226.10812 | 690.551136 | 575.876848 | 4470.0336 |
| Motif:rnd-1\_family-1070 | 519.437023 | 740.124447 | 497.787286 | 527.443071 | 498.38501 | 50.5723028 | 81.3451553 | 74.0524114 | 754.395003 | 660.432272 | 4403.97398 |
| Motif:rnd-1\_family-103 | 418.29574 | 480.586443 | 344.570923 | 466.580357 | 358.655382 | 682.04326 | 582.42488 | 576.550372 | 32.9897916 | 446.907 | 4389.60415 |
| Motif:rnd-1\_family-1113 | 391.969125 | 413.650659 | 399.899753 | 382.256921 | 377.966272 | 593.818812 | 404.846732 | 406.826103 | 493.123029 | 393.198159 | 4257.55557 |
| Motif:rnd-6\_family-398 | 345.206813 | 359.827077 | 414.985698 | 433.009991 | 458.62367 | 574.246599 | 545.306078 | 506.313604 | 190.703937 | 416.858965 | 4245.08243 |
| Motif:rnd-4\_family-572 | 376.985438 | 425.105538 | 419.464259 | 346.914073 | 485.802501 | 426.332032 | 444.526953 | 366.619543 | 522.528343 | 423.526543 | 4237.80522 |
| Motif:rnd-5\_family-3920 | 475.510456 | 519.876996 | 491.430337 | 420.422508 | 451.454325 | 158.222372 | 249.514988 | 230.128373 | 661.203904 | 539.079425 | 4196.84368 |
| Motif:rnd-1\_family-1650 | 397.258549 | 469.215854 | 488.792514 | 404.805274 | 441.589779 | 137.043909 | 401.653779 | 222.273517 | 651.400637 | 532.736768 | 4146.77058 |
| Motif:rnd-5\_family-1583 | 363.656658 | 318.988106 | 359.736953 | 331.972873 | 368.57021 | 763.477988 | 284.068952 | 376.253118 | 544.754568 | 376.185765 | 4087.66519 |
| Motif:rnd-1\_family-61 | 441.904818 | 433.550879 | 421.792126 | 412.668233 | 421.223922 | 244.464615 | 254.034845 | 256.715483 | 626.384479 | 500.616529 | 4013.35593 |
| Motif:rnd-6\_family-3909 | 460.734574 | 528.18438 | 469.484737 | 439.237169 | 439.687037 | 191.571021 | 149.294199 | 167.261218 | 624.426262 | 502.06174 | 3971.94234 |
| Motif:rnd-1\_family-1577 | 400.942485 | 424.577186 | 431.602129 | 433.103398 | 448.722277 | 240.184623 | 189.264937 | 211.282009 | 627.396036 | 489.496579 | 3896.57166 |
| Motif:rnd-5\_family-2666 | 309.684365 | 301.910051 | 316.89821 | 294.024464 | 318.85959 | 858.850824 | 288.037205 | 342.034115 | 442.092491 | 375.094274 | 3847.48559 |
| Motif:rnd-5\_family-1664 | 441.134364 | 486.991013 | 524.108171 | 393.56657 | 405.077288 | 141.477696 | 178.033285 | 189.174172 | 585.490801 | 493.816787 | 3838.87015 |
| Motif:rnd-5\_family-3329 | 377.091911 | 406.188811 | 423.368441 | 364.691461 | 379.360146 | 365.521655 | 195.378451 | 235.591396 | 599.467446 | 410.81016 | 3757.46988 |
| Motif:rnd-5\_family-302 | 345.38718 | 544.919055 | 408.33115 | 419.266197 | 462.419356 | 164.549507 | 192.554485 | 168.124754 | 583.962789 | 445.764411 | 3735.27888 |
| Motif:rnd-6\_family-75 | 408.023702 | 356.021787 | 383.228615 | 329.764307 | 348.685366 | 316.126436 | 281.132292 | 289.372496 | 587.600227 | 426.610656 | 3726.56588 |
| Motif:rnd-4\_family-2199 | 337.286212 | 289.992612 | 377.308023 | 430.968403 | 384.029952 | 601.177511 | 401.054749 | 469.62762 | 60.0778975 | 370.631502 | 3722.15448 |
| Motif:rnd-5\_family-283 | 407.642924 | 413.725145 | 429.62681 | 377.758499 | 345.33188 | 248.070337 | 214.803344 | 193.039823 | 589.16832 | 483.994454 | 3703.16153 |
| Motif:rnd-6\_family-427 | 409.074355 | 496.615008 | 418.301345 | 382.828992 | 428.880787 | 82.6271296 | 191.064642 | 147.561346 | 643.987495 | 501.334023 | 3702.27512 |
| Motif:rnd-6\_family-10863 | 312.95356 | 348.459405 | 341.57087 | 306.115683 | 322.012737 | 636.512844 | 246.470207 | 289.010404 | 460.296657 | 380.970409 | 3644.37278 |
| Motif:rnd-5\_family-610 | 388.573501 | 448.747174 | 410.468099 | 370.087201 | 385.885479 | 248.447087 | 191.346041 | 202.299026 | 549.110986 | 439.274233 | 3634.23883 |
| Motif:rnd-6\_family-5358 | 229.717904 | 268.274112 | 304.237807 | 225.109359 | 290.122774 | 634.626651 | 501.371831 | 577.699886 | 308.760101 | 257.909362 | 3597.82979 |
| Motif:rnd-1\_family-410 | 369.152816 | 397.961413 | 387.786291 | 340.350766 | 419.790306 | 225.815474 | 186.03377 | 202.631425 | 538.056528 | 442.969005 | 3510.54779 |
| Motif:rnd-1\_family-1523 | 357.722894 | 415.896624 | 401.816776 | 352.483162 | 453.737177 | 78.9415064 | 145.009172 | 128.91215 | 651.258751 | 523.503264 | 3509.28148 |
| Motif:rnd-1\_family-400 | 352.805654 | 358.446016 | 394.866855 | 336.111016 | 310.098772 | 183.193857 | 278.405353 | 234.450104 | 565.327612 | 444.976969 | 3458.68221 |
| Motif:rnd-1\_family-1059 | 357.467598 | 264.056325 | 312.318472 | 318.410697 | 346.308846 | 401.878825 | 420.965713 | 482.362051 | 148.332775 | 364.044524 | 3416.14583 |
| Motif:rnd-5\_family-2193 | 337.932738 | 372.447129 | 343.243942 | 325.846964 | 333.209333 | 315.358296 | 258.470684 | 240.925719 | 490.966916 | 381.30545 | 3399.70717 |
| Motif:rnd-6\_family-339 | 276.245642 | 150.047874 | 267.024135 | 198.545539 | 485.778727 | 295.763035 | 374.488915 | 356.151744 | 578.113848 | 406.497665 | 3388.65712 |
| Motif:rnd-1\_family-964 | 358.847631 | 284.228209 | 190.694024 | 286.294371 | 329.21587 | 451.101312 | 432.84704 | 534.727224 | 47.3603498 | 389.412597 | 3304.72863 |
| Motif:rnd-5\_family-835 | 370.288678 | 388.040191 | 408.835876 | 341.164762 | 320.315214 | 166.349745 | 114.706355 | 131.428112 | 572.255728 | 440.744521 | 3254.12918 |
| Motif:rnd-1\_family-1055 | 371.346151 | 256.680061 | 305.743131 | 297.983338 | 338.478359 | 401.868147 | 428.138358 | 454.47476 | 62.9662735 | 328.162858 | 3245.84144 |
| Motif:rnd-6\_family-1699 | 355.921189 | 376.15081 | 379.240399 | 323.084621 | 290.065172 | 385.327184 | 188.281503 | 215.770695 | 399.603738 | 323.436338 | 3236.88165 |
| Motif:rnd-1\_family-12 | 269.333928 | 267.413252 | 307.821896 | 373.721706 | 378.592259 | 444.430402 | 394.398288 | 369.094314 | 92.9291862 | 303.01715 | 3200.75238 |
| Motif:rnd-6\_family-4476 | 340.371227 | 352.150113 | 369.035044 | 318.153914 | 315.293882 | 236.334649 | 167.352817 | 173.121425 | 500.414575 | 415.294177 | 3187.52182 |
| Motif:rnd-1\_family-670 | 395.033927 | 319.969034 | 194.979768 | 289.205407 | 283.019725 | 377.635511 | 398.047481 | 522.695204 | 29.8491322 | 356.058136 | 3166.49333 |
| Motif:rnd-1\_family-1828 | 426.376011 | 456.5556 | 547.268242 | 382.556173 | 450.554843 | 98.1062112 | 130.80657 | 90.509768 | 277.130775 | 238.136521 | 3098.00071 |
| Motif:rnd-6\_family-323 | 311.756546 | 325.741347 | 337.586024 | 351.994138 | 359.82515 | 92.0116997 | 221.938878 | 198.624594 | 472.408092 | 422.977592 | 3094.86406 |
| Motif:rnd-6\_family-443 | 194.787818 | 228.068401 | 223.665141 | 174.208896 | 204.966965 | 1035.29199 | 176.747494 | 304.707286 | 291.256475 | 227.542615 | 3061.24308 |
| Motif:rnd-6\_family-5719 | 258.876395 | 177.018824 | 285.073124 | 327.761144 | 332.427629 | 366.3707 | 394.542435 | 356.27681 | 378.072821 | 159.595968 | 3036.01585 |
| Motif:rnd-6\_family-2158 | 311.234274 | 376.759234 | 386.070273 | 317.365159 | 360.145147 | 129.804149 | 144.523923 | 127.525061 | 464.033556 | 393.857198 | 3011.31798 |
| Motif:rnd-5\_family-8237 | 298.988062 | 258.832234 | 371.097052 | 282.087974 | 246.799681 | 379.101397 | 413.696659 | 391.70128 | 22.2007648 | 302.979483 | 2967.48459 |
| Motif:rnd-4\_family-3013 | 352.158583 | 407.444473 | 379.138076 | 303.697856 | 387.619695 | 92.6780895 | 130.827666 | 134.657742 | 402.241647 | 340.728032 | 2931.19186 |
| Motif:rnd-6\_family-58 | 361.929542 | 377.14116 | 358.570749 | 330.107753 | 284.404374 | 130.842734 | 134.797277 | 134.324012 | 434.765479 | 359.766703 | 2906.64978 |
| Motif:rnd-6\_family-4001 | 307.341014 | 394.226886 | 389.033126 | 305.977567 | 336.761643 | 57.0101766 | 125.471929 | 99.1212665 | 483.339065 | 401.052684 | 2899.33536 |
| Motif:rnd-5\_family-317 | 287.932778 | 301.550704 | 326.841373 | 268.500092 | 267.521405 | 268.287148 | 214.435611 | 198.149486 | 405.669484 | 328.718377 | 2867.60646 |
| Motif:rnd-5\_family-1090 | 273.674691 | 241.271716 | 257.025273 | 281.729457 | 265.10528 | 437.549945 | 393.343156 | 357.011216 | 25.8334819 | 329.080454 | 2861.62467 |
| Motif:rnd-5\_family-3039 | 332.681982 | 379.435984 | 333.114622 | 262.558583 | 329.160378 | 87.3656722 | 191.330552 | 161.221245 | 475.66728 | 301.589976 | 2854.12627 |
| Motif:rnd-5\_family-990 | 288.973925 | 363.379954 | 294.640158 | 321.097273 | 325.299238 | 106.337466 | 143.985688 | 133.006543 | 448.594857 | 373.98876 | 2799.30386 |
| Motif:rnd-1\_family-1639 | 302.736811 | 328.274998 | 350.139855 | 299.428643 | 377.280057 | 87.0229476 | 114.41723 | 103.987832 | 446.407132 | 378.492968 | 2788.18847 |
| Motif:rnd-6\_family-5168 | 274.262795 | 240.484727 | 262.114659 | 230.921399 | 270.42533 | 337.818061 | 230.286313 | 274.347665 | 391.837054 | 266.400467 | 2778.89847 |
| Motif:rnd-1\_family-1869 | 234.219934 | 246.346643 | 269.763127 | 204.725275 | 240.956187 | 302.988563 | 279.758091 | 342.47763 | 380.20883 | 264.398582 | 2765.84286 |
| Motif:rnd-1\_family-669 | 265.791892 | 295.050557 | 326.419725 | 274.235245 | 271.956956 | 55.1479087 | 332.088369 | 186.850157 | 397.017905 | 332.152094 | 2736.71081 |
| Motif:rnd-6\_family-4780 | 283.320276 | 302.510276 | 295.175783 | 290.124797 | 296.137204 | 115.986444 | 186.618737 | 164.225687 | 443.98705 | 353.940794 | 2732.02705 |
| Motif:rnd-6\_family-9065 | 248.107146 | 300.392289 | 290.98471 | 268.469687 | 307.37268 | 195.388469 | 209.8561 | 189.910751 | 398.48485 | 316.631091 | 2725.59777 |
| Motif:rnd-6\_family-12152 | 255.660391 | 300.599384 | 248.558927 | 268.381064 | 258.459684 | 201.121596 | 248.978172 | 257.826183 | 395.927974 | 287.910374 | 2723.42375 |
| Motif:rnd-6\_family-41 | 282.375903 | 327.545845 | 331.531829 | 271.314068 | 304.452612 | 89.9439899 | 249.760913 | 162.306093 | 380.954051 | 312.341523 | 2712.52683 |
| Motif:rnd-6\_family-1065 | 198.107393 | 214.331453 | 215.881093 | 194.975284 | 226.904494 | 707.447195 | 204.264879 | 218.835346 | 293.010622 | 226.904683 | 2700.66244 |
| Motif:rnd-1\_family-82 | 264.607077 | 251.54452 | 285.963141 | 270.731568 | 274.423609 | 213.143034 | 197.504181 | 188.247077 | 419.027036 | 321.985931 | 2687.17717 |
| Motif:rnd-1\_family-93 | 279.89304 | 311.23763 | 244.318399 | 280.720043 | 262.621932 | 293.974507 | 158.295146 | 135.357983 | 407.239683 | 306.129238 | 2679.7876 |
| Motif:rnd-6\_family-7151 | 226.016276 | 229.628471 | 249.306423 | 214.130614 | 200.808045 | 540.054337 | 155.720514 | 229.464123 | 321.437578 | 262.84684 | 2629.41322 |
| Motif:rnd-6\_family-5254 | 272.356237 | 322.966953 | 346.902493 | 295.760511 | 242.79675 | 89.9463136 | 132.652861 | 101.382957 | 439.447207 | 380.070868 | 2624.28315 |
| Motif:rnd-5\_family-41 | 230.555426 | 198.273902 | 225.129661 | 189.616142 | 201.07482 | 271.842645 | 301.700701 | 301.456795 | 397.070377 | 291.952242 | 2608.67271 |
| Motif:rnd-6\_family-862 | 254.792758 | 277.613683 | 265.023006 | 247.810443 | 242.745258 | 336.944851 | 167.888155 | 176.838355 | 350.514452 | 277.958155 | 2598.12912 |
| Motif:rnd-6\_family-337 | 280.750493 | 318.883014 | 328.338452 | 259.46333 | 298.823586 | 80.8328271 | 121.421152 | 101.550383 | 436.81901 | 356.753655 | 2583.6359 |
| Motif:rnd-5\_family-2450 | 242.067436 | 237.707808 | 245.186139 | 237.934158 | 217.603773 | 427.976104 | 166.584535 | 164.678174 | 350.113594 | 293.184868 | 2583.03659 |
| Motif:rnd-1\_family-152 | 253.865731 | 246.85609 | 255.382723 | 234.080985 | 218.23841 | 183.314331 | 253.23463 | 229.280621 | 387.199911 | 311.900137 | 2573.35357 |
| Motif:rnd-1\_family-1204 | 235.717555 | 231.469934 | 200.234108 | 210.126358 | 200.780643 | 571.01027 | 165.417608 | 184.039707 | 330.289549 | 226.156463 | 2555.2422 |
| Motif:rnd-5\_family-640 | 270.815384 | 332.219538 | 287.835879 | 275.167737 | 268.10339 | 115.127013 | 173.733605 | 132.538625 | 373.538561 | 308.994854 | 2538.07458 |
| Motif:rnd-1\_family-1549 | 243.536782 | 249.607392 | 255.679164 | 231.389189 | 181.422209 | 288.263927 | 277.309746 | 265.896213 | 259.262553 | 275.71036 | 2528.07754 |
| Motif:rnd-6\_family-1183 | 257.827694 | 178.105116 | 259.308504 | 213.678344 | 223.069761 | 412.441387 | 186.733986 | 215.840312 | 357.119708 | 199.804406 | 2503.92922 |
| Motif:rnd-1\_family-385 | 297.909111 | 355.374427 | 328.968252 | 297.547668 | 277.425654 | 46.9389985 | 100.029841 | 89.2325072 | 383.548027 | 303.741575 | 2480.71606 |
| Motif:rnd-1\_family-488 | 279.831029 | 305.159072 | 212.859346 | 231.291994 | 177.215985 | 358.26671 | 351.073851 | 304.672919 | 25.2101235 | 230.377532 | 2475.95856 |
| Motif:rnd-6\_family-2055 | 196.145525 | 213.64403 | 182.781657 | 221.830451 | 166.996931 | 384.647787 | 238.931777 | 243.873402 | 334.205359 | 248.551399 | 2431.60832 |
| Motif:rnd-1\_family-1071 | 260.576497 | 318.841951 | 271.958113 | 249.366419 | 258.721918 | 97.0292737 | 150.832758 | 121.107477 | 371.740459 | 304.362745 | 2404.53761 |
| Motif:rnd-6\_family-8958 | 207.831323 | 254.407288 | 264.677661 | 244.03526 | 300.687969 | 80.8386533 | 119.178103 | 121.613764 | 434.263763 | 376.9141 | 2404.44789 |
| Motif:rnd-1\_family-113 | 236.841917 | 238.923628 | 240.747829 | 207.87645 | 208.102664 | 169.552857 | 310.553757 | 193.268771 | 328.073187 | 266.208154 | 2400.14921 |
| Motif:rnd-5\_family-4686 | 238.508032 | 254.478934 | 266.452849 | 242.717805 | 253.996918 | 205.890073 | 154.429007 | 163.057792 | 347.870364 | 267.760172 | 2395.16195 |
| Motif:rnd-1\_family-243 | 233.136085 | 267.572354 | 256.33828 | 236.997039 | 287.516126 | 130.278392 | 105.657319 | 104.878179 | 384.129066 | 316.981201 | 2323.48404 |
| Motif:rnd-4\_family-294 | 195.581799 | 166.547383 | 185.621466 | 172.094505 | 185.639446 | 255.154902 | 254.986967 | 274.86063 | 367.952801 | 256.151524 | 2314.59142 |
| Motif:rnd-6\_family-10437 | 239.411317 | 254.767823 | 281.522252 | 242.687777 | 246.769901 | 100.279816 | 146.573503 | 134.167887 | 352.884367 | 281.183095 | 2280.24774 |
| Motif:rnd-1\_family-424 | 245.32004 | 137.302835 | 265.113583 | 197.418962 | 257.406176 | 192.357466 | 183.339295 | 228.423545 | 317.388709 | 199.277759 | 2223.34837 |
| Motif:rnd-6\_family-1974 | 186.577318 | 197.094354 | 203.629508 | 172.660087 | 174.21991 | 450.147194 | 155.848197 | 225.374131 | 252.109631 | 199.727035 | 2217.38736 |
| Motif:rnd-6\_family-1496 | 245.774091 | 310.99771 | 231.18813 | 208.479023 | 234.486433 | 154.785466 | 92.1118947 | 106.957118 | 365.131938 | 248.842175 | 2198.75398 |
| Motif:rnd-6\_family-2368 | 223.712261 | 254.275246 | 225.925 | 227.429538 | 272.837704 | 85.1923453 | 129.840477 | 122.608496 | 340.840182 | 271.133382 | 2153.79463 |
| Motif:rnd-6\_family-11285 | 213.214519 | 253.212124 | 218.96838 | 205.547048 | 185.071645 | 211.459217 | 135.159711 | 166.086814 | 312.468287 | 230.451986 | 2131.63973 |
| Motif:rnd-1\_family-636 | 210.467659 | 240.363776 | 247.567305 | 207.830122 | 233.583362 | 41.9488394 | 247.674641 | 137.197983 | 307.411929 | 254.677987 | 2128.7236 |
| Motif:rnd-1\_family-435 | 262.405094 | 202.800126 | 148.536087 | 214.161774 | 159.621824 | 266.684692 | 296.680706 | 288.573958 | 21.3688363 | 257.014991 | 2117.84809 |
| Motif:rnd-2\_family-70 | 176.295355 | 176.763364 | 211.503649 | 228.937086 | 231.559007 | 237.58081 | 267.885204 | 233.183154 | 103.917864 | 212.782521 | 2080.40801 |
| Motif:rnd-6\_family-850 | 187.360371 | 180.939416 | 193.097477 | 157.143043 | 210.651171 | 269.936804 | 175.919768 | 215.848749 | 258.65214 | 206.877581 | 2056.42652 |
| Motif:rnd-6\_family-1505 | 207.619671 | 222.326205 | 222.977128 | 193.043286 | 208.894424 | 144.711383 | 173.816634 | 153.098789 | 270.451544 | 233.261901 | 2030.20097 |
| Motif:rnd-1\_family-955 | 209.760163 | 218.997824 | 162.597796 | 205.40375 | 210.357991 | 157.989302 | 137.410511 | 139.101431 | 316.395005 | 223.911234 | 1981.92501 |
| Motif:rnd-1\_family-1557 | 164.045269 | 172.479839 | 171.977507 | 170.164783 | 192.776009 | 215.505591 | 205.395375 | 263.274603 | 254.979856 | 160.899136 | 1971.49797 |
| Motif:rnd-6\_family-3651 | 193.086746 | 209.508627 | 199.977755 | 188.907451 | 181.412862 | 224.284588 | 125.641638 | 141.398382 | 280.608939 | 221.516606 | 1966.3436 |
| Motif:rnd-5\_family-1208 | 153.488409 | 161.793575 | 200.556822 | 156.430536 | 215.171516 | 222.825431 | 199.26305 | 253.534976 | 237.927475 | 155.206641 | 1956.19843 |
| Motif:rnd-1\_family-16 | 169.020698 | 164.809286 | 197.47731 | 208.787146 | 215.209778 | 259.575999 | 232.550485 | 216.435588 | 101.310816 | 185.746952 | 1950.92406 |
| Motif:rnd-5\_family-2295 | 192.404666 | 205.219233 | 209.51115 | 180.190393 | 186.924539 | 131.99181 | 178.153354 | 158.422268 | 272.025206 | 209.240243 | 1924.08286 |
| Motif:rnd-5\_family-5166 | 236.871372 | 236.002908 | 254.002421 | 226.584892 | 213.623349 | 59.8749753 | 95.8142396 | 84.8173639 | 276.466049 | 237.072251 | 1921.12982 |
| Motif:rnd-6\_family-9316 | 174.759056 | 186.251424 | 198.00513 | 208.959192 | 204.18896 | 279.454248 | 198.755743 | 244.656015 | 35.821671 | 188.871743 | 1919.72318 |
| Motif:rnd-5\_family-2196 | 153.989531 | 137.772892 | 179.126076 | 144.190519 | 179.753501 | 193.201165 | 244.358771 | 186.035172 | 280.83268 | 206.395955 | 1905.65626 |
| Motif:rnd-3\_family-267 | 163.17563 | 147.1897 | 169.15222 | 148.600591 | 145.094967 | 177.269003 | 244.879547 | 229.860618 | 272.74779 | 199.141411 | 1897.11148 |
| Motif:rnd-1\_family-777 | 182.368484 | 193.277137 | 231.720846 | 182.256377 | 179.203018 | 31.0451918 | 274.476925 | 135.977721 | 228.88592 | 199.601418 | 1838.81304 |
| Motif:rnd-5\_family-49 | 168.661721 | 183.290126 | 185.918958 | 189.947665 | 190.858591 | 226.796797 | 116.311379 | 143.980087 | 239.964555 | 188.50085 | 1834.23073 |
| Motif:rnd-1\_family-1614 | 161.307386 | 157.478125 | 171.376581 | 152.603534 | 173.320914 | 293.895101 | 133.219451 | 159.862056 | 246.005012 | 182.831879 | 1831.90004 |
| Motif:rnd-1\_family-270 | 209.363759 | 253.830327 | 233.127634 | 210.966949 | 192.47869 | 53.1142348 | 111.475997 | 80.0952236 | 256.124214 | 228.322437 | 1828.89947 |
| Motif:rnd-5\_family-1662 | 227.110166 | 200.161467 | 197.00463 | 178.183185 | 175.648101 | 131.709736 | 181.467388 | 180.94742 | 137.85967 | 199.327892 | 1809.41965 |
| Motif:rnd-1\_family-264 | 227.587071 | 226.454104 | 227.607134 | 183.81699 | 245.527135 | 72.5386145 | 84.2981292 | 78.7145866 | 249.354921 | 207.520679 | 1803.41937 |
| Motif:rnd-6\_family-3591 | 170.611186 | 188.578683 | 196.32181 | 179.317484 | 194.776359 | 98.1359219 | 161.979118 | 132.118811 | 265.516049 | 213.006847 | 1800.36227 |
| Motif:rnd-6\_family-684 | 197.17828 | 229.31935 | 216.830465 | 190.580749 | 203.319496 | 40.9239372 | 56.8868571 | 59.7905525 | 313.58125 | 268.797188 | 1777.20813 |
| Motif:rnd-6\_family-1930 | 178.865653 | 220.294067 | 216.359153 | 194.28336 | 219.208142 | 81.160541 | 92.3230218 | 73.9914615 | 265.530002 | 213.893588 | 1755.90899 |
| Motif:rnd-5\_family-326 | 186.573355 | 218.097937 | 184.823589 | 192.513984 | 200.017301 | 59.6447808 | 152.911813 | 109.866547 | 237.795261 | 213.625753 | 1755.87032 |
| Motif:rnd-6\_family-10883 | 190.104043 | 215.427788 | 212.282829 | 170.871335 | 221.338486 | 90.4541714 | 92.4769681 | 88.0257567 | 262.062955 | 208.225929 | 1751.27026 |
| Motif:rnd-5\_family-1420 | 201.72288 | 153.409895 | 125.561185 | 169.599957 | 165.663033 | 232.000511 | 236.717428 | 247.105995 | 21.3268074 | 193.182477 | 1746.29017 |
| Motif:rnd-1\_family-207 | 189.701572 | 150.302981 | 135.627203 | 166.444946 | 182.984547 | 229.018705 | 225.71348 | 242.303892 | 21.7136141 | 179.072404 | 1722.88334 |
| Motif:rnd-6\_family-3887 | 171.880082 | 174.071406 | 178.203533 | 164.400978 | 169.824731 | 119.713747 | 135.66919 | 124.093911 | 274.157286 | 210.162093 | 1722.17696 |
| Motif:rnd-1\_family-1154 | 204.56028 | 143.516362 | 110.374441 | 148.043977 | 135.779991 | 237.404931 | 232.377144 | 288.091555 | 19.3414917 | 201.085061 | 1720.57523 |
| Motif:rnd-6\_family-1554 | 128.431028 | 152.896213 | 163.875983 | 143.179941 | 157.467754 | 153.923712 | 154.128667 | 165.27924 | 278.067983 | 209.737729 | 1706.98825 |
| Motif:rnd-6\_family-1525 | 165.542482 | 205.674875 | 210.642202 | 183.150479 | 201.116767 | 55.3093737 | 109.690669 | 81.3106854 | 270.472584 | 218.333963 | 1701.24408 |
| Motif:rnd-6\_family-2386 | 121.122968 | 138.433589 | 130.619172 | 138.365569 | 123.692913 | 323.524652 | 167.225406 | 227.892778 | 189.17986 | 135.302966 | 1695.35987 |
| Motif:rnd-1\_family-274 | 168.067384 | 181.839231 | 184.359829 | 160.507248 | 186.528684 | 137.094382 | 119.741001 | 116.417335 | 244.355097 | 196.399706 | 1695.3099 |
| Motif:rnd-6\_family-425 | 139.985605 | 157.357215 | 185.193281 | 165.466995 | 135.339319 | 73.7620785 | 248.305141 | 154.521261 | 232.495167 | 197.296175 | 1689.72224 |
| Motif:rnd-1\_family-515 | 172.086042 | 210.777589 | 209.549482 | 177.285858 | 185.527368 | 43.1856589 | 127.42958 | 88.5378454 | 261.728575 | 212.487757 | 1688.59576 |
| Motif:rnd-6\_family-470 | 174.855243 | 181.798499 | 192.399013 | 167.134271 | 189.205176 | 69.7068271 | 128.465976 | 113.666704 | 265.108923 | 204.566496 | 1686.90713 |
| Motif:rnd-1\_family-202 | 144.252794 | 102.461803 | 142.725312 | 125.376283 | 152.95368 | 176.022045 | 209.572306 | 217.141302 | 245.370479 | 168.170011 | 1684.04601 |
| Motif:rnd-1\_family-518 | 225.156473 | 241.201685 | 239.318811 | 177.163684 | 168.141142 | 86.6579521 | 85.1450143 | 98.5040608 | 203.51035 | 158.115664 | 1682.91484 |
| Motif:rnd-6\_family-1887 | 142.148137 | 159.482137 | 163.113627 | 145.109956 | 165.555363 | 95.260642 | 216.468645 | 169.645918 | 218.026695 | 177.889619 | 1652.70074 |
| Motif:rnd-1\_family-911 | 184.84571 | 213.767657 | 208.038594 | 168.888591 | 194.369725 | 39.9808436 | 88.1845292 | 67.5982083 | 268.614286 | 217.382685 | 1651.67083 |
| Motif:rnd-6\_family-12619 | 197.469883 | 212.978267 | 217.059078 | 177.158155 | 181.222251 | 56.7420234 | 79.5492994 | 75.2058174 | 254.731807 | 196.0486 | 1648.16518 |
| Motif:rnd-4\_family-469 | 158.948233 | 156.377819 | 148.440882 | 141.995778 | 144.555199 | 123.7998 | 171.848503 | 153.122113 | 246.20978 | 198.025607 | 1643.32371 |
| Motif:rnd-6\_family-5982 | 164.674791 | 188.044137 | 158.584335 | 161.720193 | 167.223945 | 203.904354 | 113.117888 | 109.769343 | 198.75506 | 166.291189 | 1632.08524 |
| Motif:rnd-1\_family-866 | 146.818834 | 126.543756 | 137.871753 | 124.669647 | 129.446826 | 176.66239 | 167.872384 | 197.758805 | 234.68969 | 184.300535 | 1626.63462 |
| Motif:rnd-6\_family-1772 | 156.217964 | 170.324062 | 183.535872 | 145.053151 | 195.291263 | 131.460146 | 92.5949557 | 102.775966 | 247.089354 | 193.018413 | 1617.36115 |
| Motif:rnd-1\_family-40 | 151.806249 | 84.6334051 | 154.577502 | 146.279046 | 181.046444 | 227.228324 | 181.733785 | 221.176634 | 143.963244 | 124.163065 | 1616.6077 |
| Motif:rnd-6\_family-11911 | 132.058505 | 111.013238 | 142.338615 | 137.031016 | 181.688663 | 212.044318 | 169.729736 | 193.350011 | 186.611513 | 147.729615 | 1613.59523 |
| Motif:rnd-1\_family-686 | 153.593099 | 125.746915 | 135.711068 | 127.514 | 121.729862 | 183.608481 | 166.146624 | 173.014783 | 244.239999 | 176.016862 | 1607.32169 |
| Motif:rnd-5\_family-2058 | 156.591415 | 128.3686 | 140.283374 | 148.157946 | 184.602975 | 227.363008 | 232.866917 | 204.889106 | 26.5706043 | 157.096132 | 1606.79008 |
| Motif:rnd-1\_family-24 | 148.261052 | 159.118568 | 165.041274 | 167.064002 | 143.076818 | 75.8169135 | 206.964011 | 116.248384 | 228.475683 | 184.275088 | 1594.34179 |
| Motif:rnd-6\_family-7331 | 180.1264 | 189.788983 | 186.207824 | 152.531121 | 163.551508 | 39.6055182 | 111.577411 | 84.7870732 | 263.499083 | 219.095954 | 1590.77087 |
| Motif:rnd-6\_family-5915 | 192.747313 | 239.25362 | 267.932084 | 183.993789 | 202.806168 | 23.9009919 | 96.0578517 | 53.1001734 | 174.431605 | 143.523753 | 1577.74735 |
| Motif:rnd-5\_family-3705 | 163.412694 | 181.000939 | 163.926136 | 154.132474 | 147.611995 | 46.313738 | 148.082085 | 98.8223899 | 257.686707 | 215.410283 | 1576.39944 |
| Motif:rnd-5\_family-2423 | 132.057367 | 125.597549 | 142.433339 | 117.040785 | 121.517827 | 120.434184 | 231.520452 | 177.793531 | 228.654818 | 170.062718 | 1567.11257 |
| Motif:rnd-1\_family-1913 | 165.377612 | 193.016879 | 187.604331 | 162.134842 | 187.744013 | 71.5438655 | 130.542706 | 98.9779564 | 200.762024 | 156.481694 | 1554.18592 |
| Motif:rnd-1\_family-477 | 137.284475 | 150.410185 | 162.277267 | 134.052363 | 123.990012 | 17.676249 | 278.48673 | 155.247057 | 202.729877 | 188.156605 | 1550.31082 |
| Motif:rnd-6\_family-2023 | 183.84854 | 225.78653 | 226.044441 | 183.28699 | 215.097022 | 34.4999441 | 48.4480806 | 43.0712611 | 211.576581 | 178.01421 | 1549.6736 |
| Motif:rnd-6\_family-3547 | 165.598249 | 152.756781 | 182.44283 | 152.099658 | 145.965168 | 138.983477 | 107.730048 | 94.3806566 | 231.499976 | 177.962342 | 1549.41919 |
| Motif:rnd-1\_family-302 | 169.93899 | 150.344974 | 148.563208 | 150.65543 | 157.17558 | 167.113559 | 154.955788 | 143.605723 | 121.426238 | 179.40801 | 1543.1875 |
| Motif:rnd-6\_family-10182 | 147.436788 | 134.280485 | 143.297699 | 144.89039 | 146.529552 | 165.137175 | 128.879403 | 130.663643 | 223.093934 | 175.927937 | 1540.137 |
| Motif:rnd-1\_family-373 | 129.378473 | 109.942421 | 96.1762134 | 110.859616 | 96.4044325 | 159.560576 | 164.961893 | 199.359942 | 275.904562 | 191.329565 | 1533.87769 |
| Motif:rnd-6\_family-2079 | 163.634189 | 186.641683 | 170.307098 | 150.21379 | 166.75754 | 125.380331 | 69.3263498 | 82.495954 | 226.535068 | 173.931561 | 1515.22356 |
| Motif:rnd-1\_family-726 | 140.764419 | 164.263206 | 184.684809 | 145.633181 | 149.866955 | 30.283419 | 221.738277 | 109.521594 | 196.325641 | 168.304744 | 1511.38624 |
| Motif:rnd-6\_family-10536 | 177.30506 | 170.623947 | 188.648366 | 155.136763 | 175.244343 | 56.5798672 | 83.9368831 | 90.2433646 | 233.521021 | 158.227061 | 1489.46668 |
| Motif:rnd-5\_family-866 | 151.274013 | 165.569306 | 151.028044 | 128.797231 | 174.833657 | 69.1224426 | 114.888705 | 79.8237798 | 247.747586 | 202.413371 | 1485.49814 |
| Motif:rnd-6\_family-4218 | 172.035425 | 187.466988 | 162.680997 | 162.241676 | 170.054033 | 53.4117178 | 85.00181 | 73.8285272 | 224.049672 | 190.018267 | 1480.78911 |
| Motif:rnd-1\_family-337 | 141.261217 | 155.901874 | 139.482482 | 149.43177 | 174.609697 | 84.2983898 | 135.272133 | 111.675444 | 208.487362 | 178.831257 | 1479.25163 |
| Motif:rnd-6\_family-1027 | 154.640028 | 215.504574 | 180.428913 | 173.396514 | 193.655962 | 29.7517163 | 43.5308035 | 36.5317588 | 245.237877 | 197.591696 | 1470.26984 |
| Motif:rnd-4\_family-1511 | 140.164915 | 124.192346 | 124.616703 | 124.891949 | 146.212361 | 153.630055 | 140.011943 | 139.330445 | 208.021884 | 147.784988 | 1448.85759 |
| Motif:rnd-1\_family-1619 | 120.766436 | 83.6173425 | 116.874347 | 97.1133379 | 106.802301 | 179.724573 | 157.323387 | 197.138355 | 231.694191 | 156.989447 | 1448.04372 |
| Motif:rnd-1\_family-939 | 142.551579 | 180.735777 | 166.158871 | 125.393088 | 209.528302 | 37.1960423 | 95.2870205 | 78.3208431 | 221.571478 | 170.570631 | 1427.31363 |
| Motif:rnd-6\_family-5159 | 160.22359 | 168.171011 | 160.395514 | 149.904528 | 137.44738 | 92.5927749 | 79.7385927 | 83.0371455 | 215.106412 | 171.08606 | 1417.70301 |
| Motif:rnd-1\_family-447 | 141.178677 | 118.331376 | 97.0216571 | 144.978687 | 162.016613 | 199.133884 | 184.321992 | 193.130341 | 33.5633524 | 140.343208 | 1414.01979 |
| Motif:rnd-1\_family-1107 | 138.781019 | 129.309511 | 131.455567 | 126.677207 | 112.032445 | 116.427332 | 131.150385 | 131.034362 | 224.421618 | 172.002622 | 1413.29207 |
| Motif:rnd-1\_family-84 | 146.682838 | 147.721601 | 122.245007 | 146.363966 | 136.557951 | 182.69707 | 186.032697 | 170.254912 | 30.2410908 | 139.896394 | 1408.69353 |
| Motif:rnd-5\_family-139 | 133.794186 | 134.022143 | 131.095933 | 132.446531 | 145.115834 | 171.231521 | 177.036142 | 199.488006 | 30.3125111 | 151.260657 | 1405.80346 |
| Motif:rnd-1\_family-1696 | 121.375227 | 93.23079 | 141.976481 | 128.937697 | 174.145971 | 184.725313 | 140.967607 | 168.386979 | 174.632805 | 74.6030307 | 1402.9819 |
| Motif:rnd-1\_family-173 | 139.343086 | 165.984355 | 159.098787 | 141.32066 | 141.854409 | 48.7331667 | 85.0717135 | 68.806556 | 242.130281 | 183.875181 | 1376.21819 |
| Motif:rnd-6\_family-3508 | 105.969887 | 76.0593128 | 110.938491 | 91.6428943 | 154.352613 | 133.294442 | 150.155861 | 156.385615 | 233.225621 | 164.014596 | 1376.03933 |
| Motif:rnd-5\_family-1764 | 131.075079 | 96.9298885 | 114.206429 | 108.771252 | 143.735803 | 179.184138 | 181.218324 | 188.187467 | 80.1663546 | 147.011025 | 1370.48576 |
| Motif:rnd-6\_family-2880 | 150.538079 | 178.829361 | 163.647203 | 147.725268 | 157.920921 | 39.9570687 | 57.4882834 | 53.5059394 | 225.987561 | 193.138234 | 1368.73792 |
| Motif:rnd-1\_family-532 | 142.023343 | 114.719758 | 96.5374353 | 118.864213 | 123.567383 | 163.434665 | 174.214051 | 196.675792 | 89.240635 | 147.651927 | 1366.9292 |
| Motif:rnd-1\_family-369 | 122.973083 | 117.745559 | 127.086803 | 108.281819 | 133.251877 | 140.326071 | 141.04632 | 150.372274 | 167.490221 | 152.848948 | 1361.42297 |
| Motif:rnd-6\_family-2223 | 126.024681 | 154.819145 | 161.165892 | 137.736716 | 151.977547 | 67.8019849 | 92.906029 | 73.4221994 | 214.020622 | 175.777551 | 1355.65237 |
| Motif:rnd-1\_family-1749 | 148.490687 | 169.761844 | 159.17017 | 128.811707 | 144.875358 | 43.8536149 | 68.5174615 | 59.6898473 | 229.720534 | 176.875078 | 1329.7663 |
| Motif:rnd-6\_family-992 | 141.310776 | 154.32041 | 139.120523 | 136.685847 | 121.826712 | 60.0323206 | 97.3871974 | 107.002658 | 201.838403 | 168.415654 | 1327.9405 |
| Motif:rnd-5\_family-23 | 118.535479 | 150.719544 | 147.606879 | 152.549259 | 145.375816 | 15.8466085 | 75.3955967 | 55.4618319 | 257.436474 | 206.303847 | 1325.23133 |
| Motif:rnd-1\_family-1262 | 129.988796 | 149.149035 | 123.761159 | 128.064388 | 138.293513 | 49.2149537 | 119.541767 | 97.1436799 | 217.102472 | 170.953317 | 1323.21308 |
| Motif:rnd-1\_family-1252 | 146.524079 | 185.339287 | 122.993866 | 129.700106 | 130.310095 | 54.3291158 | 97.4961056 | 105.972342 | 202.341271 | 132.963935 | 1307.9702 |
| Motif:rnd-5\_family-4697 | 134.684325 | 102.875446 | 155.252507 | 129.115734 | 142.229145 | 85.5545641 | 125.198736 | 112.220633 | 174.292219 | 144.551796 | 1305.97511 |
| Motif:rnd-1\_family-832 | 138.19039 | 168.810116 | 165.222322 | 134.487766 | 169.444136 | 33.0271685 | 49.2545957 | 38.8709919 | 225.807352 | 182.436113 | 1305.55095 |
| Motif:rnd-6\_family-2943 | 115.014463 | 121.236207 | 110.028968 | 122.323419 | 165.019948 | 61.0390927 | 126.448879 | 105.31653 | 209.033767 | 168.521623 | 1303.9829 |
| Motif:rnd-6\_family-7992 | 127.714793 | 150.987352 | 146.217442 | 118.86552 | 133.261871 | 94.0969832 | 96.4227317 | 80.4692172 | 191.495598 | 158.609754 | 1298.14126 |
| Motif:rnd-6\_family-9763 | 152.337458 | 180.850807 | 147.890231 | 135.682563 | 140.477673 | 120.568057 | 44.2300909 | 58.3758437 | 172.94479 | 139.024736 | 1292.38225 |
| Motif:rnd-6\_family-2729 | 154.725405 | 195.498929 | 182.796812 | 149.825912 | 152.898575 | 22.5975397 | 51.2791764 | 39.6659981 | 180.405144 | 154.889869 | 1284.58336 |
| Motif:rnd-6\_family-16609 | 124.793755 | 132.579976 | 144.722789 | 131.392384 | 132.220702 | 107.990445 | 96.277142 | 79.6070086 | 184.47776 | 140.800582 | 1274.86254 |
| Motif:rnd-6\_family-3103 | 129.776111 | 130.710332 | 123.66432 | 133.493779 | 134.925122 | 82.7964475 | 98.5977479 | 105.020599 | 196.858376 | 132.368651 | 1268.21149 |
| Motif:rnd-1\_family-167 | 109.095795 | 92.8975174 | 120.299411 | 97.7855979 | 95.6866831 | 100.263621 | 188.673918 | 140.314512 | 187.053611 | 135.144844 | 1267.21551 |
| Motif:rnd-6\_family-1138 | 126.330658 | 165.370792 | 136.776612 | 147.080585 | 180.823479 | 22.194589 | 37.699049 | 39.8492469 | 218.511187 | 187.203551 | 1261.83975 |
| Motif:rnd-6\_family-12116 | 130.551035 | 132.963122 | 130.480543 | 133.65363 | 149.229395 | 61.1863899 | 88.9451187 | 80.0418963 | 190.81702 | 159.99477 | 1257.86292 |
| Motif:rnd-1\_family-1037 | 115.577562 | 128.680646 | 115.068046 | 114.921132 | 131.489708 | 118.845657 | 94.7165257 | 96.4148965 | 193.370353 | 144.060658 | 1253.14518 |
| Motif:rnd-6\_family-3357 | 115.02837 | 86.6437916 | 89.3621791 | 87.7896582 | 80.8339046 | 150.159483 | 137.800019 | 170.335364 | 195.980244 | 127.045896 | 1240.97891 |
| Motif:rnd-6\_family-2490 | 133.301135 | 161.696746 | 158.059385 | 126.487863 | 125.282934 | 26.7180083 | 98.0561571 | 76.0645025 | 177.077308 | 150.248194 | 1232.99223 |
| Motif:rnd-6\_family-2181 | 113.568084 | 108.254571 | 112.243718 | 120.683938 | 135.222817 | 151.206835 | 158.575966 | 149.57373 | 60.2384669 | 122.002999 | 1231.57113 |
| Motif:rnd-6\_family-314 | 122.16881 | 148.766956 | 152.21774 | 126.06753 | 145.772353 | 46.4706632 | 92.856255 | 66.5159784 | 180.569955 | 146.622517 | 1228.02876 |
| Motif:rnd-6\_family-1052 | 125.760641 | 157.770656 | 159.80036 | 133.573115 | 136.820769 | 29.7429676 | 65.4217229 | 56.0541375 | 199.168992 | 154.470805 | 1218.58417 |
| Motif:rnd-5\_family-7700 | 135.385255 | 144.326874 | 148.191863 | 140.229998 | 153.631069 | 35.2787406 | 53.4333784 | 50.4320791 | 195.57263 | 158.828584 | 1215.31047 |
| Motif:rnd-1\_family-731 | 128.632757 | 112.086822 | 121.120826 | 106.480638 | 121.328558 | 101.094905 | 104.529383 | 93.3958941 | 183.02661 | 143.252768 | 1214.94916 |
| Motif:rnd-1\_family-131 | 105.09353 | 92.7640409 | 103.647218 | 92.7140853 | 106.616658 | 145.876668 | 130.772338 | 136.134503 | 171.476075 | 128.194531 | 1213.28965 |
| Motif:rnd-6\_family-10230 | 98.69023 | 65.9498208 | 109.638441 | 103.422355 | 83.3024279 | 221.766125 | 135.990884 | 184.224587 | 154.308084 | 40.9608261 | 1198.25378 |
| Motif:rnd-6\_family-1323 | 109.864528 | 101.734202 | 118.38451 | 107.003418 | 103.127212 | 98.0496818 | 132.570702 | 154.830425 | 145.338063 | 121.073787 | 1191.97653 |
| Motif:rnd-5\_family-2038 | 120.833541 | 134.495989 | 127.962989 | 121.938226 | 172.41904 | 43.069223 | 74.7550362 | 69.9309142 | 180.78032 | 139.417651 | 1185.60293 |
| Motif:rnd-1\_family-98 | 121.780284 | 101.0876 | 111.351967 | 110.402342 | 124.386365 | 129.028047 | 139.513733 | 113.429502 | 100.069855 | 132.748273 | 1183.79797 |
| Motif:rnd-1\_family-174 | 124.644593 | 110.203393 | 101.285132 | 112.60952 | 103.181202 | 111.13696 | 167.565215 | 148.020338 | 69.1271729 | 135.024308 | 1182.79783 |
| Motif:rnd-6\_family-12996 | 135.353142 | 162.497641 | 141.880585 | 119.903561 | 133.459602 | 34.6880654 | 54.1060251 | 59.5914427 | 192.465363 | 144.491156 | 1178.43658 |
| Motif:rnd-1\_family-1593 | 85.4448622 | 79.3159927 | 98.8632429 | 99.982899 | 115.117358 | 141.958454 | 132.311339 | 119.903301 | 175.090439 | 121.51861 | 1169.5065 |
| Motif:rnd-1\_family-408 | 107.008561 | 90.4183545 | 110.344815 | 111.727136 | 101.734155 | 198.407649 | 140.762798 | 150.32451 | 44.5346336 | 111.343456 | 1166.60607 |
| Motif:rnd-1\_family-294 | 112.18551 | 109.650552 | 118.806928 | 98.1525801 | 92.9172631 | 61.0673396 | 162.847088 | 108.516396 | 156.390258 | 135.884986 | 1156.4189 |
| Motif:rnd-1\_family-1155 | 113.955732 | 123.44679 | 121.970035 | 119.877599 | 109.417426 | 90.1967648 | 63.9042007 | 75.9122478 | 183.338945 | 140.030762 | 1142.0505 |
| Motif:rnd-3\_family-366 | 112.403467 | 123.971238 | 115.683299 | 118.312687 | 95.9596168 | 104.520589 | 111.370233 | 103.176827 | 120.548664 | 122.927022 | 1128.87364 |
| Motif:rnd-5\_family-3272 | 111.532332 | 104.177969 | 81.3317433 | 112.956467 | 99.0234247 | 159.029485 | 152.724238 | 177.876648 | 12.5788494 | 117.334733 | 1128.56589 |
| Motif:rnd-6\_family-1241 | 105.771711 | 99.5661688 | 116.939306 | 132.830355 | 69.2972562 | 164.345435 | 80.8341734 | 76.0880755 | 156.728869 | 125.497604 | 1127.89895 |
| Motif:rnd-1\_family-650 | 113.785163 | 151.32009 | 121.96499 | 114.858342 | 139.905369 | 20.4808436 | 51.6021569 | 39.2357234 | 187.243618 | 182.005654 | 1122.40195 |
| Motif:rnd-1\_family-193 | 90.7575271 | 80.9597846 | 94.1863129 | 81.8077057 | 82.7596927 | 92.0630614 | 190.901564 | 132.631301 | 152.385903 | 118.218934 | 1116.67179 |
| Motif:rnd-5\_family-2209 | 100.219964 | 52.7031315 | 111.41466 | 76.7479591 | 196.833735 | 144.367194 | 128.368933 | 156.342485 | 53.6214956 | 95.5410687 | 1116.16063 |
| Motif:rnd-5\_family-2004 | 115.022356 | 129.1344 | 114.101756 | 111.506343 | 125.683027 | 50.0086638 | 64.299019 | 58.1099096 | 191.264684 | 156.224468 | 1115.35463 |
| Motif:rnd-5\_family-1311 | 83.3637162 | 50.1296967 | 113.83496 | 95.7726208 | 102.400938 | 150.767347 | 121.846357 | 149.373633 | 168.076513 | 73.6464445 | 1109.21223 |
| Motif:rnd-1\_family-3 | 107.830882 | 97.1530676 | 103.862159 | 100.022015 | 108.591504 | 102.641062 | 105.867098 | 103.391877 | 158.064827 | 120.264845 | 1107.68934 |
| Motif:rnd-1\_family-1209 | 113.710975 | 129.491611 | 128.489053 | 117.283848 | 112.807946 | 28.0637528 | 67.3645327 | 48.3967836 | 197.981389 | 163.913006 | 1107.5029 |
| Motif:rnd-1\_family-635 | 99.5700743 | 108.146802 | 130.504287 | 100.184363 | 96.8197193 | 15.745822 | 172.194214 | 98.6433793 | 146.253989 | 133.144203 | 1101.20685 |
| Motif:rnd-1\_family-1119 | 129.890397 | 110.581528 | 139.760133 | 113.012506 | 137.105874 | 50.8399204 | 61.6203586 | 53.5405955 | 166.671463 | 137.44503 | 1100.46781 |
| Motif:rnd-5\_family-20 | 121.453987 | 158.780048 | 69.6497277 | 127.70929 | 121.636213 | 76.7000983 | 60.234886 | 66.622281 | 163.778217 | 127.410449 | 1093.9752 |
| Motif:rnd-5\_family-3317 | 109.305178 | 66.452308 | 140.81095 | 102.604453 | 159.83928 | 109.940109 | 89.5343646 | 84.8736345 | 155.366822 | 62.6725065 | 1081.3996 |
| Motif:rnd-6\_family-3572 | 98.1849338 | 86.0327985 | 104.082353 | 96.7975476 | 102.660386 | 111.563605 | 100.387415 | 99.925271 | 155.450047 | 117.439577 | 1072.52393 |
| Motif:rnd-1\_family-1682 | 107.870103 | 95.7830405 | 118.480452 | 86.5911211 | 53.9929647 | 94.8325673 | 113.323601 | 109.323311 | 165.204246 | 124.715602 | 1070.11701 |
| Motif:rnd-6\_family-2764 | 121.755773 | 131.986581 | 85.2257676 | 125.711085 | 119.36803 | 78.3920229 | 65.2649074 | 68.5651644 | 151.105862 | 122.462157 | 1069.83735 |
| Motif:rnd-1\_family-603 | 123.991501 | 119.554817 | 127.528205 | 113.655636 | 103.064181 | 47.4564693 | 76.4872474 | 60.8533365 | 161.042552 | 134.018676 | 1067.65262 |
| Motif:rnd-1\_family-1184 | 116.415742 | 121.867648 | 131.699575 | 114.989863 | 112.037949 | 39.4613898 | 77.9888917 | 49.3072841 | 167.669925 | 134.950346 | 1066.38861 |
| Motif:rnd-6\_family-3028 | 109.59555 | 121.914565 | 115.891625 | 107.518018 | 105.207885 | 28.7189272 | 110.614453 | 77.76343 | 157.04757 | 130.899287 | 1065.17131 |
| Motif:rnd-5\_family-2753 | 109.505691 | 123.864414 | 141.678195 | 112.681328 | 127.990829 | 38.189629 | 82.307777 | 67.4913141 | 135.81097 | 124.13518 | 1063.65533 |
| Motif:rnd-1\_family-1143 | 114.79181 | 117.639058 | 116.445796 | 108.474547 | 121.903309 | 41.3219769 | 82.977852 | 74.1360282 | 149.247301 | 134.898645 | 1061.83632 |
| Motif:rnd-1\_family-401 | 116.43045 | 123.67286 | 116.957383 | 111.287216 | 141.147275 | 40.5199778 | 51.6900393 | 54.394104 | 171.609737 | 127.521141 | 1055.23018 |
| Motif:rnd-6\_family-273 | 101.547551 | 130.142614 | 138.0118 | 104.024391 | 160.523161 | 19.2591539 | 25.9852638 | 27.4018723 | 186.962692 | 160.84311 | 1054.70161 |
| Motif:rnd-6\_family-1295 | 113.491253 | 130.349998 | 126.456326 | 100.099065 | 127.126625 | 31.8791536 | 38.4240228 | 44.7635223 | 189.689364 | 150.362247 | 1052.64158 |
| Motif:rnd-1\_family-122 | 103.025004 | 106.152831 | 94.240199 | 103.20031 | 102.970651 | 103.602695 | 89.2168591 | 85.4182216 | 142.679249 | 112.034457 | 1042.54048 |
| Motif:rnd-1\_family-1265 | 106.916116 | 113.412894 | 102.241262 | 107.246341 | 116.578891 | 53.8607371 | 54.9056683 | 64.0561007 | 171.398443 | 151.390618 | 1042.00707 |
| Motif:rnd-6\_family-4563 | 87.0681888 | 44.5939723 | 113.801495 | 84.6732086 | 67.8834113 | 213.298203 | 131.79686 | 175.670688 | 67.036246 | 48.8146532 | 1034.63693 |
| Motif:rnd-6\_family-2824 | 77.9698551 | 77.7809551 | 71.8919515 | 79.3396697 | 92.8150786 | 132.613844 | 117.409418 | 123.221966 | 150.297709 | 106.00771 | 1029.34816 |
| Motif:rnd-5\_family-2012 | 102.909293 | 114.752662 | 119.65854 | 98.3201867 | 124.681133 | 61.5484487 | 57.5946255 | 62.5584449 | 156.577856 | 125.730873 | 1024.33206 |
| Motif:rnd-6\_family-9696 | 94.1698181 | 106.703283 | 112.707558 | 105.926047 | 140.449234 | 32.2729563 | 65.9409782 | 51.9598051 | 167.143107 | 143.340242 | 1020.61303 |
| Motif:rnd-6\_family-5311 | 118.047885 | 119.434152 | 112.541813 | 98.0605869 | 148.996754 | 22.0407852 | 77.3989642 | 38.2773261 | 152.390819 | 126.064146 | 1013.25323 |
| Motif:rnd-6\_family-12105 | 129.151823 | 142.373895 | 142.400012 | 110.702835 | 115.383764 | 20.7412472 | 51.7168694 | 36.7050797 | 136.324751 | 124.605426 | 1010.1057 |
| Motif:rnd-6\_family-7506 | 80.5957518 | 72.5324832 | 79.7931156 | 76.387676 | 66.689428 | 153.142214 | 114.619717 | 118.766971 | 150.567956 | 95.9802327 | 1009.07555 |
| Motif:rnd-6\_family-6470 | 81.0922804 | 72.7943651 | 90.8941243 | 89.3693863 | 76.9421964 | 136.314747 | 100.867918 | 111.453384 | 146.078142 | 102.713701 | 1008.52025 |
| Motif:rnd-6\_family-6399 | 112.948206 | 100.875422 | 115.32098 | 100.985397 | 120.11335 | 47.3973224 | 59.0750545 | 63.929955 | 153.06724 | 132.895293 | 1006.60822 |
| Motif:rnd-6\_family-6882 | 101.793796 | 115.000131 | 107.707427 | 106.213037 | 109.202019 | 63.2310882 | 75.881335 | 66.5855867 | 138.798021 | 118.508029 | 1002.92047 |
| Motif:rnd-6\_family-5714 | 101.403526 | 80.0696175 | 74.3624233 | 90.6378159 | 119.947514 | 127.407134 | 124.739775 | 159.284672 | 11.0028619 | 113.404629 | 1002.25997 |
| Motif:rnd-6\_family-72 | 118.106366 | 130.651552 | 64.6974275 | 95.328854 | 120.445786 | 49.3904109 | 56.1358124 | 66.6504456 | 176.475589 | 121.073787 | 998.956031 |
| Motif:rnd-1\_family-164 | 115.120026 | 127.431118 | 114.403359 | 112.994113 | 128.686246 | 40.7967448 | 42.749871 | 37.7548694 | 148.397347 | 128.150893 | 996.484587 |
| Motif:rnd-4\_family-44 | 76.9677177 | 86.9024585 | 95.548645 | 93.1713837 | 79.7480771 | 63.0248055 | 198.294582 | 107.369179 | 85.9859105 | 107.567992 | 994.580751 |
| Motif:rnd-1\_family-88 | 94.458166 | 95.4036935 | 99.0502372 | 86.6003145 | 112.582496 | 80.8890005 | 89.2114544 | 83.3569378 | 141.976162 | 107.531644 | 991.060106 |
| Motif:rnd-1\_family-308 | 101.427134 | 86.9225802 | 76.2549042 | 88.5961785 | 97.052866 | 121.868191 | 126.024086 | 119.133039 | 63.2896146 | 105.641694 | 986.210287 |
| Motif:rnd-1\_family-1391 | 111.984769 | 146.430146 | 140.432773 | 99.6051371 | 109.07838 | 20.3226419 | 22.3161938 | 24.4175695 | 165.710387 | 142.425 | 982.722998 |
| Motif:rnd-1\_family-240 | 90.6843413 | 78.1657993 | 92.7024047 | 78.4319029 | 79.5428557 | 83.5067996 | 123.873116 | 97.980261 | 143.666139 | 109.822545 | 978.376164 |
| Motif:rnd-1\_family-1199 | 102.50088 | 96.8916005 | 103.993979 | 84.7311315 | 67.852168 | 38.483701 | 127.375412 | 96.9917392 | 146.678169 | 108.098413 | 973.597193 |
| Motif:rnd-5\_family-291 | 97.5368893 | 105.926438 | 104.809307 | 88.9548616 | 88.3611246 | 67.7999983 | 68.9498857 | 71.6544151 | 155.779291 | 118.724722 | 968.496932 |
| Motif:rnd-6\_family-2543 | 94.0860402 | 118.755242 | 110.367234 | 99.441079 | 89.1104341 | 18.1669102 | 95.0162633 | 61.9071429 | 155.147956 | 124.687563 | 966.685865 |
| Motif:rnd-6\_family-5252 | 96.7252286 | 105.107306 | 105.144311 | 94.0618808 | 97.3210195 | 89.4700119 | 62.2295984 | 72.9299919 | 127.001086 | 111.568229 | 961.558662 |
| Motif:rnd-1\_family-631 | 83.5739455 | 87.2605253 | 95.1379353 | 89.5526035 | 73.1967299 | 33.2800155 | 198.559835 | 92.1880748 | 106.492659 | 102.000682 | 961.243006 |
| Motif:rnd-5\_family-1715 | 95.308249 | 103.475993 | 87.4249402 | 95.0380571 | 122.056839 | 37.7390436 | 85.1543508 | 64.75698 | 142.902155 | 127.299901 | 961.156509 |
| Motif:rnd-6\_family-1433 | 105.27659 | 111.833068 | 125.598007 | 109.974723 | 99.6801614 | 39.9001802 | 33.3869689 | 32.8576107 | 165.055327 | 135.218818 | 958.781453 |
| Motif:rnd-1\_family-480 | 101.249563 | 102.013507 | 109.409285 | 99.6172486 | 143.313253 | 25.4666708 | 53.5534121 | 35.9793828 | 154.242273 | 130.245177 | 955.089772 |
| Motif:rnd-1\_family-1612 | 114.21529 | 136.984006 | 110.014217 | 79.2056415 | 77.8845764 | 43.4943233 | 56.7945722 | 57.6339256 | 178.335123 | 88.8067009 | 943.368375 |
| Motif:rnd-1\_family-1176 | 99.4287507 | 100.2547 | 91.5929369 | 85.359228 | 83.7507979 | 50.5455844 | 73.4010001 | 73.1884569 | 154.091704 | 123.103809 | 934.716967 |
| Motif:rnd-5\_family-4863 | 83.9438392 | 79.9687702 | 84.4736726 | 93.4611038 | 93.7251907 | 80.1754167 | 90.5237168 | 75.0423213 | 142.565074 | 96.8858773 | 920.764983 |
| Motif:rnd-6\_family-1122 | 91.168 | 98.2032393 | 99.9265828 | 79.9582257 | 63.8415682 | 113.529347 | 65.8225052 | 63.9570789 | 134.571545 | 108.273516 | 919.251608 |
| Motif:rnd-6\_family-1823 | 99.8739631 | 95.9290116 | 92.2399423 | 83.7695281 | 107.167344 | 56.2163824 | 66.3196873 | 58.9843337 | 141.329699 | 117.150343 | 918.980235 |
| Motif:rnd-1\_family-217 | 77.026121 | 58.7858146 | 79.9139183 | 65.2351405 | 77.1381115 | 96.1314844 | 107.77908 | 110.570402 | 145.989121 | 100.337457 | 918.90665 |
| Motif:rnd-1\_family-1153 | 91.9375416 | 99.620404 | 94.8352958 | 90.8528425 | 94.6616801 | 81.2587657 | 65.6325797 | 55.8599191 | 134.968902 | 108.696642 | 918.324572 |
| Motif:rnd-6\_family-1016 | 88.0025036 | 65.6680201 | 77.7378934 | 64.810241 | 59.3832048 | 86.0192574 | 126.160703 | 113.099838 | 138.118218 | 98.4011135 | 917.400993 |
| Motif:rnd-1\_family-970 | 88.509502 | 92.1728007 | 91.7390175 | 87.5286246 | 91.6310394 | 56.7103775 | 121.108943 | 66.4243717 | 119.263925 | 100.998569 | 916.08717 |
| Motif:rnd-6\_family-791 | 83.507147 | 93.6736495 | 107.665901 | 82.05778 | 104.497808 | 36.6468533 | 105.466515 | 65.8387012 | 123.091393 | 111.489663 | 913.935411 |
| Motif:rnd-1\_family-818 | 89.2411871 | 90.5648549 | 104.052734 | 90.574995 | 102.548417 | 58.7259439 | 55.2972234 | 56.9313038 | 140.999906 | 118.395679 | 907.332244 |
| Motif:rnd-5\_family-720 | 100.652865 | 120.096889 | 117.871866 | 103.603069 | 114.689762 | 37.4478016 | 38.8463321 | 35.2892537 | 129.238987 | 109.530434 | 907.267259 |
| Motif:rnd-6\_family-2474 | 93.0597222 | 73.0624124 | 92.0755698 | 87.3260788 | 108.534675 | 54.5085938 | 51.486031 | 53.642163 | 155.668379 | 125.648108 | 895.011732 |
| Motif:rnd-1\_family-1719 | 96.7723472 | 99.097421 | 112.503885 | 91.5446849 | 98.6524989 | 38.1300185 | 58.1611345 | 48.9843959 | 133.49566 | 116.582464 | 893.92451 |
| Motif:rnd-1\_family-696 | 69.825735 | 57.6859122 | 74.2309733 | 69.9691746 | 77.8006817 | 108.494604 | 100.987815 | 99.3714003 | 130.045016 | 101.958723 | 890.370035 |
| Motif:rnd-6\_family-84 | 90.9646919 | 75.819119 | 84.4718161 | 78.5247688 | 87.6471849 | 94.0198527 | 81.5045049 | 82.2584028 | 125.073697 | 88.8367104 | 889.120748 |
| Motif:rnd-1\_family-63 | 88.6772635 | 94.2266434 | 98.24133 | 87.5355232 | 83.4675163 | 38.5024348 | 83.5400009 | 64.2763383 | 141.203543 | 108.207485 | 887.878079 |
| Motif:rnd-6\_family-122 | 92.7774608 | 85.5735211 | 86.5439318 | 79.0048292 | 97.4602388 | 96.320677 | 67.9159265 | 65.0977829 | 120.893443 | 95.0621195 | 886.649931 |
| Motif:rnd-1\_family-52 | 94.8048906 | 83.5673119 | 92.2376035 | 84.9646227 | 88.0705332 | 54.3893689 | 80.3429246 | 66.3295232 | 122.563181 | 113.055547 | 880.325506 |
| Motif:rnd-1\_family-525 | 100.50988 | 105.352746 | 133.947676 | 93.4946063 | 86.9310716 | 27.1097542 | 77.8048961 | 50.0031508 | 105.431241 | 94.326267 | 874.911288 |
| Motif:rnd-5\_family-295 | 86.0803847 | 90.7314248 | 104.351472 | 85.7042841 | 95.8346646 | 39.1780223 | 67.4493432 | 50.8807229 | 139.191905 | 114.217239 | 873.619463 |
| Motif:rnd-5\_family-1961 | 58.4922264 | 73.2922206 | 70.8809577 | 106.842871 | 74.2065691 | 118.478546 | 82.126146 | 87.5697421 | 124.98304 | 76.6937778 | 873.566096 |
| Motif:rnd-5\_family-1287 | 91.6400273 | 104.527501 | 100.129865 | 89.368723 | 96.1615312 | 24.176607 | 67.1908317 | 41.4448617 | 138.956356 | 113.429895 | 867.026199 |
| Motif:rnd-1\_family-393 | 83.6452794 | 46.0270166 | 79.9313043 | 51.5642321 | 55.2595581 | 92.0952821 | 93.0109367 | 108.079608 | 158.437701 | 98.1545107 | 866.205429 |
| Motif:rnd-6\_family-1324 | 109.198769 | 110.438476 | 102.716127 | 98.5461094 | 89.8310384 | 10.5614715 | 103.252361 | 52.8142328 | 104.609678 | 82.5209976 | 864.489261 |
| Motif:rnd-1\_family-877 | 92.3265704 | 107.753366 | 122.480143 | 96.4792869 | 103.180871 | 22.2411321 | 36.8353387 | 31.7353836 | 134.917062 | 111.880772 | 859.829925 |
| Motif:rnd-6\_family-2423 | 94.7560929 | 108.785611 | 106.066564 | 101.105062 | 109.195828 | 18.1980903 | 55.2138177 | 39.2197306 | 127.339297 | 97.8030392 | 857.683133 |
| Motif:rnd-1\_family-1665 | 77.5650245 | 71.5874584 | 82.7401253 | 74.5922742 | 62.1677399 | 34.79181 | 165.586968 | 92.60113 | 105.270392 | 85.4819614 | 852.384884 |
| Motif:rnd-1\_family-114 | 94.6120061 | 73.4849042 | 75.8472285 | 80.077519 | 62.7679351 | 103.520437 | 119.0421 | 111.699311 | 29.2956547 | 98.8228804 | 849.169976 |
| Motif:rnd-1\_family-307 | 75.9145802 | 70.9767627 | 78.2291066 | 67.9205361 | 77.3801885 | 70.8484203 | 98.9050685 | 90.59548 | 118.236698 | 95.3999423 | 844.406783 |
| Motif:rnd-5\_family-366 | 73.4939989 | 73.1467985 | 84.8328587 | 72.9496639 | 67.0399907 | 25.367598 | 171.063903 | 80.388992 | 97.5062237 | 94.6888028 | 840.47883 |
| Motif:rnd-1\_family-106 | 65.7424272 | 65.78042 | 79.0562995 | 63.4496718 | 55.4132522 | 92.3560353 | 91.0745397 | 97.330096 | 128.281628 | 101.46963 | 839.954 |
| Motif:rnd-1\_family-1969 | 97.3672262 | 106.65878 | 102.095554 | 85.8281742 | 82.1618568 | 43.417814 | 37.1436417 | 45.341569 | 131.362627 | 102.263972 | 833.641215 |
| Motif:rnd-5\_family-2309 | 80.2066813 | 87.8635499 | 83.8852029 | 79.1476688 | 89.8602655 | 81.7868723 | 59.06134 | 61.1667978 | 110.873397 | 92.7079228 | 826.559698 |
| Motif:rnd-1\_family-1128 | 80.3332246 | 93.1385428 | 88.863595 | 82.6613158 | 99.6263947 | 34.3913655 | 62.0675447 | 50.1720364 | 119.931092 | 110.662774 | 821.847886 |
| Motif:rnd-5\_family-3928 | 100.022626 | 93.1584135 | 96.3897843 | 87.2448073 | 89.1185886 | 44.8721264 | 37.5354949 | 39.3327558 | 134.404781 | 98.2555044 | 820.334882 |
| Motif:rnd-5\_family-1163 | 73.0675282 | 55.3635596 | 86.6586538 | 72.9458967 | 69.3977776 | 84.9724256 | 93.341961 | 79.1946687 | 111.944181 | 93.0776902 | 819.964343 |
| Motif:rnd-6\_family-15255 | 80.1615705 | 34.3758208 | 99.7797343 | 74.8635338 | 119.156357 | 93.5250094 | 94.7581697 | 111.372166 | 81.4218941 | 29.8459087 | 819.260165 |
| Motif:rnd-5\_family-4592 | 85.3983179 | 93.0168983 | 90.9776483 | 86.5910144 | 89.0994871 | 28.4508705 | 76.2675956 | 45.9610608 | 118.532013 | 102.27651 | 816.571416 |
| Motif:rnd-6\_family-10845 | 102.764097 | 138.419582 | 98.3999308 | 92.5646629 | 130.602959 | 15.7437628 | 25.3816576 | 24.5876902 | 94.6322335 | 84.0017446 | 807.098321 |
| Motif:rnd-6\_family-8717 | 93.7649824 | 99.7517693 | 109.470369 | 71.0135551 | 96.2051245 | 12.578111 | 28.8130719 | 26.3246838 | 129.539302 | 137.119711 | 804.58068 |
| Motif:rnd-1\_family-54 | 78.80587 | 54.5715493 | 76.2076855 | 51.4352771 | 61.5095787 | 84.0773817 | 107.374472 | 96.5639536 | 101.222433 | 87.3937329 | 799.161934 |
| Motif:rnd-1\_family-685 | 68.8007338 | 69.1278213 | 76.4052955 | 68.657867 | 61.0718732 | 29.6067142 | 171.767215 | 73.4198581 | 93.163101 | 83.1263484 | 795.146827 |
| Motif:rnd-1\_family-51 | 79.0896338 | 85.203663 | 98.0549452 | 84.6994157 | 73.767385 | 23.9277469 | 102.049894 | 56.8032031 | 97.1582572 | 92.1954122 | 792.949556 |
| Motif:rnd-1\_family-740 | 80.0370556 | 90.7851407 | 93.6285736 | 77.864628 | 82.6203234 | 19.4550078 | 89.0445901 | 50.1456082 | 114.933315 | 91.9091129 | 790.423355 |
| Motif:rnd-5\_family-739 | 71.4638711 | 64.5121919 | 67.3410432 | 63.1626176 | 61.6094858 | 132.546554 | 69.6097487 | 77.7272966 | 106.575711 | 75.1802039 | 789.728724 |
| Motif:rnd-6\_family-7981 | 69.3795239 | 47.8709287 | 72.3219869 | 63.9770108 | 70.3535508 | 76.9678508 | 80.14747 | 83.9592237 | 130.497711 | 86.7460368 | 782.221293 |
| Motif:rnd-1\_family-162 | 62.4737039 | 60.9618037 | 68.9407835 | 61.3605356 | 73.5489199 | 66.9568888 | 135.081694 | 93.2573513 | 88.6829531 | 70.7002045 | 781.964839 |
| Motif:rnd-1\_family-795 | 79.9097332 | 92.2605961 | 93.2437684 | 85.0617751 | 97.0387023 | 29.578816 | 31.8726327 | 27.6316024 | 130.739664 | 114.276377 | 781.613668 |
| Motif:rnd-6\_family-6797 | 73.816022 | 92.0905861 | 93.485184 | 79.8586253 | 91.2224427 | 29.9460562 | 51.9176695 | 44.0985754 | 121.529949 | 102.031195 | 779.996306 |
| Motif:rnd-5\_family-97 | 87.1636597 | 66.3937422 | 82.903229 | 73.6107966 | 79.2069826 | 59.2119739 | 59.0865746 | 60.1039477 | 120.392331 | 90.4595823 | 778.532819 |
| Motif:rnd-5\_family-208 | 61.5773751 | 64.0123214 | 71.9973137 | 67.0202564 | 56.3310008 | 18.740953 | 152.895368 | 65.8811633 | 115.521765 | 103.982363 | 777.95988 |
| Motif:rnd-1\_family-384 | 64.5966589 | 48.5884298 | 58.6561211 | 49.1833613 | 63.1175307 | 99.0774216 | 82.240309 | 105.836905 | 121.759969 | 83.5095816 | 776.566288 |
| Motif:rnd-5\_family-312 | 74.2419449 | 81.2578751 | 90.3623108 | 77.6377501 | 85.16298 | 25.9697083 | 70.4531681 | 46.8129487 | 124.935025 | 96.1913681 | 773.025079 |
| Motif:rnd-5\_family-478 | 50.0158964 | 45.1977511 | 55.4880809 | 47.3469732 | 64.511497 | 209.884679 | 82.7930162 | 92.3508893 | 70.0981874 | 51.4481476 | 769.135118 |
| Motif:rnd-1\_family-1837 | 78.2987887 | 67.7638916 | 52.7594622 | 76.0044791 | 76.7085579 | 108.093111 | 104.389428 | 107.828257 | 11.6120556 | 84.0727428 | 767.530774 |
| Motif:rnd-6\_family-2714 | 82.2782759 | 87.5983935 | 82.7270172 | 94.7293934 | 93.2200211 | 33.3823503 | 30.1323821 | 32.3121238 | 128.564985 | 101.341415 | 766.286357 |
| Motif:rnd-1\_family-413 | 54.8381993 | 44.7792381 | 56.5769191 | 50.1430912 | 52.0008801 | 110.204296 | 80.7384378 | 100.426178 | 129.663456 | 82.903512 | 762.274207 |
| Motif:rnd-1\_family-1081 | 83.4553125 | 98.4026792 | 90.4831624 | 78.5337227 | 84.0974457 | 34.4639113 | 48.4624792 | 36.8811552 | 111.08856 | 96.314577 | 762.183005 |
| Motif:rnd-5\_family-2637 | 61.5328804 | 68.4821711 | 69.0636827 | 70.0568262 | 87.3395063 | 94.5101778 | 79.8744293 | 104.015317 | 52.8153682 | 71.7631159 | 759.453475 |
| Motif:rnd-1\_family-443 | 80.3550829 | 92.7915401 | 85.6500552 | 82.5876084 | 75.1308417 | 36.4570664 | 41.7330599 | 43.9441908 | 120.580028 | 99.2517346 | 758.481208 |
| Motif:rnd-6\_family-4859 | 62.8507966 | 55.5003863 | 65.2470372 | 60.0312907 | 56.8020957 | 80.355995 | 88.5192628 | 85.4758544 | 118.082373 | 84.8791444 | 757.744236 |
| Motif:rnd-1\_family-378 | 61.1055527 | 38.7531241 | 67.5127535 | 52.7247202 | 50.8353339 | 96.2980837 | 86.7054467 | 97.6123106 | 126.779214 | 79.0870534 | 757.413593 |
| Motif:rnd-1\_family-722 | 94.6699163 | 117.357583 | 101.392369 | 86.6270136 | 102.652306 | 16.6856182 | 37.9408974 | 28.7502776 | 90.8001629 | 74.1522328 | 751.028377 |
| Motif:rnd-1\_family-32 | 57.4286536 | 51.8117249 | 55.3765092 | 57.6089949 | 70.9484329 | 89.6809565 | 80.7930002 | 90.1082779 | 107.998501 | 85.208436 | 746.963487 |
| Motif:rnd-6\_family-4300 | 65.8648625 | 59.7753913 | 67.0706571 | 56.9587766 | 59.1342403 | 82.9559831 | 84.3649469 | 77.1244591 | 113.770312 | 78.754507 | 745.774135 |
| Motif:rnd-1\_family-1160 | 64.0204351 | 43.2310977 | 58.1747854 | 45.7538944 | 54.781117 | 85.4513708 | 92.4731322 | 97.8212455 | 120.381435 | 82.4573545 | 744.545867 |
| Motif:rnd-1\_family-705 | 71.7803328 | 63.7010875 | 72.918852 | 65.1875755 | 76.3902946 | 48.3764837 | 54.5510348 | 59.7382288 | 123.367262 | 93.4663986 | 729.47755 |
| Motif:rnd-3\_family-857 | 57.642247 | 53.8110162 | 53.4156301 | 62.2075991 | 75.7507041 | 86.7375018 | 81.862699 | 87.455237 | 93.2795463 | 75.1974184 | 727.359599 |
| Motif:rnd-5\_family-760 | 89.2473136 | 112.162718 | 122.447057 | 82.9159616 | 136.850297 | 9.64794712 | 19.0236066 | 16.9944188 | 75.8241054 | 58.2265752 | 723.340001 |
| Motif:rnd-5\_family-2384 | 69.6473296 | 46.1143142 | 70.1366758 | 58.3390471 | 58.7390697 | 65.3274819 | 64.5968637 | 79.7720347 | 117.421966 | 92.5824113 | 722.677194 |
| Motif:rnd-1\_family-60 | 67.0992298 | 56.7548479 | 58.8005944 | 58.2926771 | 73.3195299 | 77.6372458 | 80.1554096 | 75.5888933 | 96.0457204 | 78.1100533 | 721.804201 |
| Motif:rnd-6\_family-5703 | 64.6902439 | 69.4133735 | 68.5381873 | 72.5252536 | 80.9317519 | 60.7803763 | 60.1939309 | 57.6440041 | 107.917594 | 78.4133926 | 721.048108 |
| Motif:rnd-1\_family-19 | 69.7182653 | 63.93717 | 71.0126362 | 68.9113606 | 74.1170583 | 56.6505619 | 56.8509129 | 57.367168 | 111.011488 | 89.4893021 | 719.065923 |
| Motif:rnd-6\_family-891 | 76.206004 | 89.60937 | 98.6009596 | 70.7633105 | 88.848963 | 17.6240242 | 45.3094016 | 26.4836522 | 101.895738 | 100.289176 | 715.630599 |
| Motif:rnd-6\_family-6270 | 73.5456711 | 73.6688076 | 77.0840274 | 68.3328879 | 58.5392612 | 46.4108171 | 51.050201 | 56.5820011 | 114.376422 | 94.2061256 | 713.796222 |
| Motif:rnd-1\_family-45 | 67.921059 | 69.4231513 | 69.527589 | 79.5793117 | 72.8237492 | 85.3511398 | 89.5867203 | 71.957339 | 33.533475 | 68.5572533 | 708.260787 |
| Motif:rnd-1\_family-868 | 88.9051118 | 106.107974 | 100.487292 | 70.3137104 | 75.0864007 | 48.4131463 | 37.8201431 | 38.204914 | 78.6237475 | 63.8446642 | 707.807104 |
| Motif:rnd-1\_family-1060 | 75.1504677 | 72.5639855 | 62.747762 | 70.1506736 | 74.8982696 | 43.5906264 | 40.5371004 | 46.3876455 | 121.192749 | 98.255006 | 705.474286 |
| Motif:rnd-1\_family-418 | 70.5492381 | 71.8921338 | 74.0322517 | 66.0788766 | 73.7576087 | 53.1612152 | 42.9413939 | 48.8570158 | 117.549535 | 86.2428847 | 705.062153 |
| Motif:rnd-5\_family-2877 | 69.584768 | 76.2033074 | 64.3684302 | 71.8265252 | 75.4024416 | 81.782375 | 47.3169574 | 49.5020297 | 96.9627202 | 71.3408075 | 704.290362 |
| Motif:rnd-5\_family-1816 | 53.724215 | 52.7142548 | 57.5502171 | 65.3492019 | 61.3366841 | 95.946278 | 81.9313579 | 85.9158519 | 77.2221564 | 72.4708773 | 704.161094 |
| Motif:rnd-5\_family-2257 | 75.627977 | 84.3628697 | 81.8277084 | 73.0906915 | 81.1113543 | 25.9009697 | 38.2945087 | 36.0130533 | 114.441225 | 92.73522 | 703.405577 |
| Motif:rnd-1\_family-1529 | 73.3473718 | 73.3308196 | 78.0664444 | 72.4014857 | 75.1202512 | 37.7328394 | 54.5401583 | 41.4412503 | 105.429316 | 88.2611637 | 699.6711 |
| Motif:rnd-1\_family-72 | 55.8237188 | 49.6414572 | 65.675384 | 47.0431304 | 57.3198666 | 80.7602166 | 91.001362 | 76.4403253 | 101.433491 | 73.8884817 | 699.027434 |
| Motif:rnd-6\_family-12504 | 71.5785396 | 76.3104766 | 80.3126909 | 72.9884619 | 65.6267324 | 27.6730365 | 67.2950786 | 45.8375525 | 103.066073 | 86.4327138 | 697.121356 |
| Motif:rnd-6\_family-902 | 67.2903029 | 88.7991103 | 72.8348343 | 59.5866654 | 77.8572475 | 53.3423379 | 56.1677628 | 58.3562401 | 90.7619876 | 70.5584999 | 695.554989 |
| Motif:rnd-5\_family-5765 | 76.2276889 | 75.2666262 | 74.5045585 | 77.4238798 | 79.565486 | 33.498318 | 41.7531795 | 35.6038768 | 117.121403 | 82.9754074 | 693.940424 |
| Motif:rnd-1\_family-607 | 62.3303174 | 61.6312511 | 53.2660699 | 65.299723 | 75.065605 | 87.560681 | 90.0159892 | 93.3438133 | 35.0152962 | 70.391855 | 693.920601 |
| Motif:rnd-1\_family-139 | 72.2670278 | 54.5911645 | 59.5889202 | 56.0309484 | 71.0484678 | 86.2927775 | 72.4547331 | 84.8442093 | 74.9751366 | 60.2168393 | 692.310224 |
| Motif:rnd-1\_family-553 | 80.841422 | 44.6297695 | 85.4016667 | 63.041411 | 91.5878179 | 54.5439785 | 52.2091926 | 66.351938 | 94.07435 | 58.789197 | 691.470743 |
| Motif:rnd-6\_family-3172 | 79.4403509 | 68.6849236 | 65.0667847 | 70.1743675 | 66.9145569 | 45.3148908 | 46.5690745 | 53.7453854 | 110.873425 | 80.346349 | 687.130109 |
| Motif:rnd-6\_family-3044 | 62.363798 | 50.5173207 | 76.7033149 | 59.036489 | 82.2963744 | 59.9503513 | 39.0892049 | 43.4354287 | 122.395782 | 89.5496251 | 685.337689 |
| Motif:rnd-6\_family-9354 | 67.737886 | 70.0426352 | 80.9938253 | 63.5982986 | 51.2143303 | 24.8075485 | 86.4102304 | 52.9560881 | 101.269242 | 85.7680379 | 684.798122 |
| Motif:rnd-1\_family-949 | 63.0432554 | 56.8635905 | 64.3319605 | 61.3648356 | 71.2898296 | 71.3115156 | 62.3459723 | 63.6494412 | 96.2390977 | 74.0773839 | 684.516882 |
| Motif:rnd-1\_family-676 | 76.9249921 | 64.54658 | 69.8738518 | 66.6962166 | 77.6292187 | 58.0734183 | 46.6383349 | 49.0869719 | 99.6986476 | 74.1466144 | 683.314846 |
| Motif:rnd-1\_family-637 | 67.2238239 | 73.3087363 | 69.5542892 | 75.2020625 | 64.7531206 | 43.9746755 | 51.1398274 | 41.7778763 | 104.306719 | 89.8009996 | 681.04213 |
| Motif:rnd-1\_family-526 | 35.7315112 | 49.3615371 | 54.0234019 | 76.2338471 | 54.9626889 | 106.814971 | 58.6888595 | 71.0047904 | 105.866411 | 64.510694 | 677.198711 |
| Motif:rnd-1\_family-355 | 55.102692 | 61.9947909 | 73.3107384 | 52.7733173 | 52.6414652 | 17.6964286 | 125.362125 | 73.9490918 | 89.3174711 | 74.6963511 | 676.844471 |
| Motif:rnd-5\_family-3082 | 71.9648908 | 86.0841555 | 75.3894731 | 69.5642988 | 58.9773541 | 39.4502024 | 44.9992047 | 36.9542092 | 104.699107 | 88.5980661 | 676.680962 |
| Motif:rnd-6\_family-2596 | 65.50439 | 45.8527796 | 50.7391313 | 47.3989751 | 43.4865011 | 71.0209994 | 68.4341864 | 85.4475987 | 119.520901 | 75.470122 | 672.875584 |
| Motif:rnd-1\_family-220 | 65.1181847 | 58.0550839 | 69.0195719 | 62.5794523 | 65.9115138 | 59.4220737 | 58.3206589 | 52.3257683 | 105.474921 | 76.5033225 | 672.730551 |
| Motif:rnd-1\_family-519 | 60.4139986 | 62.4151686 | 68.4794584 | 64.0874318 | 69.9910106 | 15.576107 | 113.34489 | 63.3545785 | 79.8345162 | 74.6763245 | 672.173484 |
| Motif:rnd-4\_family-1470 | 65.6060678 | 66.5697627 | 73.0978755 | 61.5748031 | 112.957752 | 19.9296673 | 51.9203193 | 34.4919553 | 95.6584386 | 84.8225335 | 666.629175 |
| Motif:rnd-6\_family-3082 | 67.2952877 | 75.4116267 | 74.6541504 | 69.8565357 | 81.9016359 | 28.5382078 | 52.0686091 | 43.5450685 | 99.21703 | 73.8074098 | 666.295562 |
| Motif:rnd-1\_family-499 | 62.9886588 | 48.5725163 | 58.6739326 | 50.9622115 | 60.055551 | 66.1871245 | 74.1437354 | 76.3823825 | 97.6407895 | 70.079518 | 665.68642 |
| Motif:rnd-5\_family-1928 | 53.7664111 | 56.7843867 | 58.9710166 | 53.1887403 | 44.1181322 | 25.9488543 | 142.734265 | 77.4899813 | 87.917288 | 63.6650915 | 664.584167 |
| Motif:rnd-1\_family-237 | 64.5905884 | 76.4498323 | 67.458139 | 75.580516 | 46.8890738 | 98.5506894 | 88.5426346 | 74.1490227 | 10.6375865 | 61.4457357 | 664.293818 |
| Motif:rnd-1\_family-110 | 60.5233886 | 64.3386607 | 59.3571186 | 64.3839383 | 76.8592842 | 68.4176897 | 72.4799642 | 56.7647553 | 70.6788043 | 70.2089783 | 664.012582 |
| Motif:rnd-1\_family-461 | 55.4546784 | 58.7807842 | 73.3462348 | 58.1387105 | 47.9575736 | 12.6394914 | 131.985227 | 69.0971514 | 83.9047446 | 71.7889611 | 663.093557 |
| Motif:rnd-1\_family-169 | 67.751639 | 62.8826611 | 58.2875851 | 59.7070508 | 60.4714675 | 60.3036628 | 69.4164024 | 63.9618243 | 86.1175453 | 73.9484907 | 662.848329 |
| Motif:rnd-1\_family-1863 | 75.2800421 | 73.9558947 | 75.3567975 | 65.4685191 | 99.3076604 | 10.6784513 | 39.3113175 | 25.08279 | 102.321177 | 94.4147076 | 661.177357 |
| Motif:rnd-1\_family-1926 | 71.9273126 | 71.1963104 | 76.6913384 | 62.1543558 | 54.3959702 | 10.67703 | 102.847397 | 44.6835642 | 85.416807 | 77.4264293 | 657.416515 |
| Motif:rnd-6\_family-7307 | 64.5658249 | 92.8890855 | 92.1026707 | 74.9881269 | 82.680937 | 22.7687376 | 28.0413863 | 22.3995254 | 93.1058296 | 83.0717404 | 656.613864 |
| Motif:rnd-6\_family-4269 | 63.8151305 | 68.1612855 | 77.5357521 | 59.661048 | 44.5110747 | 9.74606378 | 83.5584448 | 45.341569 | 107.93836 | 93.6144295 | 653.883158 |
| Motif:rnd-1\_family-1300 | 69.2675015 | 66.3267562 | 73.8494284 | 61.8522465 | 73.9345924 | 38.9507108 | 50.2408695 | 41.7157387 | 97.1448177 | 79.4597835 | 652.742445 |
| Motif:rnd-5\_family-4365 | 70.5583623 | 88.8783959 | 77.4761643 | 68.3057624 | 65.8841086 | 24.6259973 | 31.5646982 | 31.2589426 | 103.137633 | 90.3711136 | 652.061178 |
| Motif:rnd-4\_family-1676 | 70.9726016 | 55.9806751 | 58.0640889 | 52.583801 | 40.0964548 | 51.9769216 | 56.9866393 | 68.8444398 | 113.805694 | 81.3007958 | 650.612112 |
| Motif:rnd-1\_family-772 | 52.6991599 | 56.3214612 | 62.2807359 | 55.2053621 | 47.3466882 | 16.6191279 | 161.804795 | 65.6401521 | 66.1516727 | 65.707482 | 649.776636 |
| Motif:rnd-5\_family-2063 | 65.5114915 | 65.7656032 | 68.2629139 | 66.4740789 | 73.8178255 | 42.6127149 | 42.3079346 | 35.4312872 | 103.31157 | 85.2604704 | 648.75589 |
| Motif:rnd-1\_family-1017 | 70.9983169 | 75.9573049 | 84.6296548 | 68.9319784 | 80.2998513 | 21.5557717 | 27.8760839 | 19.5188275 | 108.642718 | 88.9570569 | 647.367565 |
| Motif:rnd-1\_family-503 | 79.4560222 | 67.6003067 | 66.2564118 | 67.3266347 | 65.8076445 | 46.9284377 | 54.9210679 | 54.8067599 | 65.7108301 | 77.7326986 | 646.546814 |
| Motif:rnd-6\_family-60 | 64.8943192 | 64.3545137 | 64.2612984 | 61.3881039 | 67.9720618 | 44.3691926 | 64.8340945 | 50.9323133 | 88.7578052 | 74.1226572 | 645.88636 |
| Motif:rnd-6\_family-4493 | 47.7570588 | 44.2301145 | 49.8311239 | 48.2558265 | 41.251426 | 174.297651 | 48.2043319 | 70.0665029 | 68.7272899 | 53.0525883 | 645.673914 |
| Motif:rnd-6\_family-7183 | 35.3416191 | 34.2471431 | 37.7914524 | 31.75979 | 31.6165429 | 180.640929 | 86.3324545 | 98.7011222 | 58.5078552 | 47.5476017 | 642.48651 |
| Motif:rnd-6\_family-906 | 56.1075624 | 42.1496774 | 67.866757 | 61.6744394 | 65.0570437 | 109.16634 | 74.9308744 | 85.3107039 | 33.4412107 | 45.289031 | 640.99364 |
| Motif:rnd-1\_family-628 | 44.2779431 | 33.9547853 | 45.6243398 | 44.1035988 | 39.2959317 | 60.4109942 | 70.8434272 | 89.1609635 | 123.015884 | 88.9428236 | 639.630691 |
| Motif:rnd-4\_family-2329 | 72.1139884 | 69.1443068 | 79.1817366 | 74.7620311 | 58.6584311 | 30.1743889 | 34.2340876 | 30.3952214 | 102.1952 | 83.0457336 | 633.905126 |
| Motif:rnd-5\_family-195 | 53.9979535 | 50.2003109 | 61.8388015 | 65.9520344 | 67.8330511 | 83.8708037 | 77.2778171 | 73.3345522 | 39.2831131 | 59.5375667 | 633.126004 |
| Motif:rnd-1\_family-150 | 63.0107914 | 54.0550042 | 57.3459464 | 63.0269425 | 55.7060828 | 67.8920635 | 66.6469406 | 69.1404063 | 65.0769421 | 71.2050222 | 633.106142 |
| Motif:rnd-6\_family-9012 | 73.7006877 | 89.5730173 | 82.6699121 | 76.4080088 | 58.0183135 | 10.1472962 | 39.6657409 | 27.3212018 | 98.051545 | 77.1586278 | 632.714351 |
| Motif:rnd-1\_family-744 | 52.4504997 | 37.3417442 | 54.7542404 | 47.5100159 | 51.8453782 | 79.0154803 | 65.8501102 | 78.3120396 | 98.6936686 | 64.6867517 | 630.459929 |
| Motif:rnd-6\_family-5458 | 57.0444657 | 52.619029 | 54.6965889 | 54.4827343 | 57.6564894 | 62.9754659 | 71.7461002 | 62.4105971 | 91.4691765 | 64.9550232 | 630.05567 |
| Motif:rnd-1\_family-252 | 69.7163785 | 73.318693 | 74.9724933 | 63.5438079 | 56.5933373 | 28.2497932 | 44.820002 | 34.553506 | 97.9494992 | 85.2371547 | 628.954665 |
| Motif:rnd-1\_family-377 | 66.4728111 | 49.949585 | 49.1430989 | 57.1970294 | 66.3755977 | 78.7461721 | 81.8658275 | 89.2082381 | 19.6460605 | 69.4213928 | 628.025813 |
| Motif:rnd-1\_family-814 | 60.8398443 | 55.1084641 | 61.2733859 | 55.5159962 | 55.4967903 | 48.7356987 | 73.5190175 | 56.3431094 | 83.3190598 | 72.6808607 | 622.832227 |
| Motif:rnd-1\_family-97 | 54.1825148 | 53.5864053 | 64.7938005 | 61.1883974 | 64.3502602 | 80.7826702 | 79.1753501 | 71.2969602 | 25.337065 | 66.1814587 | 620.874882 |
| Motif:rnd-6\_family-5078 | 64.1496927 | 67.7950246 | 69.5944922 | 65.1740046 | 71.0905396 | 56.893714 | 30.5216169 | 37.3426628 | 89.0564339 | 66.8036275 | 618.421809 |
| Motif:rnd-1\_family-206 | 54.4075393 | 55.4158853 | 51.0201109 | 48.468788 | 50.4607987 | 90.7105372 | 57.3375782 | 68.9490115 | 81.2512767 | 59.5953351 | 617.616861 |
| Motif:rnd-6\_family-6331 | 69.4889643 | 70.0794614 | 64.5277446 | 58.633456 | 69.510917 | 23.4764429 | 53.1771907 | 36.9121118 | 94.3056088 | 76.8400578 | 616.951955 |
| Motif:rnd-1\_family-708 | 54.3848355 | 52.0550244 | 56.1276155 | 50.6567929 | 52.8082547 | 46.1217796 | 95.7034626 | 69.3096479 | 74.1539894 | 64.8093698 | 616.130772 |
| Motif:rnd-1\_family-361 | 59.8817679 | 63.6405458 | 62.9741258 | 58.3860501 | 61.8164194 | 48.9014306 | 45.9904912 | 41.7679461 | 99.8278394 | 72.2834469 | 615.470063 |
| Motif:rnd-1\_family-134 | 52.4668596 | 57.0343992 | 60.4232371 | 55.0851672 | 57.3551449 | 27.0956416 | 98.3507738 | 57.640921 | 80.4095008 | 67.6738452 | 613.535491 |
| Motif:rnd-5\_family-1513 | 61.8615488 | 75.7379501 | 78.235813 | 61.7171806 | 75.89378 | 25.88227 | 39.2900528 | 27.5289196 | 92.2491595 | 75.1363848 | 613.533059 |
| Motif:rnd-1\_family-157 | 60.9815636 | 62.1171245 | 70.9821795 | 61.7878369 | 67.4608049 | 28.1997204 | 57.1650661 | 36.5276988 | 88.6644764 | 74.1267842 | 608.013255 |
| Motif:rnd-5\_family-6090 | 48.5365763 | 32.7057936 | 61.3994433 | 42.0060424 | 54.0575255 | 111.423986 | 61.0130546 | 72.465266 | 78.1084494 | 46.2669017 | 607.983038 |
| Motif:rnd-1\_family-534 | 53.6308516 | 55.9110155 | 66.6000093 | 58.0448214 | 58.6095758 | 39.2486971 | 77.5834667 | 49.096011 | 81.7942464 | 66.6791874 | 607.197882 |
| Motif:rnd-6\_family-9788 | 50.9212564 | 51.493361 | 62.4281857 | 67.2592092 | 93.2947631 | 63.3869118 | 57.8283079 | 70.0078773 | 34.5013934 | 55.4277102 | 606.548976 |
| Motif:rnd-1\_family-473 | 68.3317336 | 48.4900711 | 30.8015002 | 53.5186062 | 52.353647 | 77.8134421 | 76.724307 | 88.6512394 | 35.103908 | 74.0404072 | 605.828862 |
| Motif:rnd-1\_family-1979 | 77.4094878 | 77.0849093 | 82.203309 | 75.2619381 | 63.5945809 | 17.4343953 | 22.8865169 | 18.8610561 | 97.4271313 | 72.9202294 | 605.083554 |
| Motif:rnd-1\_family-1751 | 56.973158 | 67.5150924 | 71.93014 | 61.2600346 | 61.5348411 | 39.6075063 | 46.5743222 | 40.1886319 | 88.7692174 | 69.8370599 | 604.190004 |
| Motif:rnd-6\_family-1731 | 55.7046045 | 36.8572387 | 57.1923911 | 40.9686429 | 52.5534006 | 67.9669409 | 66.7314457 | 68.8526693 | 89.0938862 | 67.0363104 | 602.95753 |
| Motif:rnd-6\_family-7508 | 59.3868419 | 67.5229183 | 75.6411813 | 59.2115101 | 58.8455826 | 19.6419854 | 54.8911401 | 29.0235868 | 96.9627202 | 79.562033 | 600.6895 |
| Motif:rnd-6\_family-3794 | 50.5505423 | 42.2066053 | 53.4532444 | 51.1097943 | 56.830455 | 56.0240167 | 74.3727831 | 66.2456196 | 85.4708912 | 64.1907585 | 600.45471 |
| Motif:rnd-6\_family-3678 | 89.8273028 | 90.4768865 | 96.3445983 | 48.8716231 | 108.060727 | 27.1952081 | 38.2392216 | 30.4997591 | 39.7901227 | 29.4422385 | 598.747688 |
| Motif:rnd-6\_family-5796 | 54.7116888 | 62.8311948 | 67.2606837 | 56.4539721 | 54.0584708 | 52.07015 | 52.507234 | 42.0510915 | 85.2818666 | 70.7318537 | 597.958206 |
| Motif:rnd-6\_family-71 | 46.6690378 | 41.2279847 | 46.191513 | 41.9255512 | 54.8200633 | 70.0172064 | 70.4404454 | 76.1182893 | 89.1976796 | 60.7885441 | 597.396315 |
| Motif:rnd-4\_family-490 | 52.6031089 | 31.0802708 | 64.1410342 | 48.3161988 | 44.6624547 | 59.8560107 | 58.8136029 | 70.3404992 | 91.1841394 | 72.622059 | 593.619379 |
| Motif:rnd-6\_family-3067 | 53.2374774 | 55.2285201 | 58.9780108 | 53.1501852 | 67.923837 | 46.0592782 | 70.873789 | 54.9610513 | 72.1885365 | 59.7390193 | 592.339705 |
| Motif:rnd-1\_family-338 | 53.9989071 | 49.492304 | 61.464208 | 49.4249999 | 50.0643551 | 30.1530496 | 91.6288229 | 60.6431378 | 81.2830827 | 64.0524761 | 592.205343 |
| Motif:rnd-6\_family-1081 | 50.7734108 | 45.053593 | 42.0735513 | 39.7750642 | 39.0043163 | 61.1555795 | 65.5618395 | 73.8491018 | 106.661857 | 67.8330543 | 591.741368 |
| Motif:rnd-6\_family-5454 | 61.2541843 | 80.5923233 | 66.6287084 | 65.1934528 | 61.4212158 | 14.7382646 | 43.0219316 | 29.5840244 | 93.17203 | 75.5342797 | 591.140415 |
| Motif:rnd-6\_family-5274 | 61.3898638 | 70.8658524 | 63.4926261 | 60.3120959 | 59.6423308 | 27.8977693 | 52.2397544 | 37.985977 | 83.5498475 | 73.0717086 | 590.447826 |
| Motif:rnd-5\_family-36 | 50.3590904 | 40.0018591 | 41.2878157 | 40.4148951 | 39.3789639 | 62.8239575 | 69.8275932 | 76.285985 | 95.3616181 | 72.0871264 | 587.828904 |
| Motif:rnd-4\_family-927 | 45.7138278 | 50.0494155 | 52.6768362 | 47.9405163 | 43.7308774 | 39.6996023 | 101.016256 | 69.5258686 | 80.1973408 | 56.8311534 | 587.381694 |
| Motif:rnd-5\_family-164 | 55.1118357 | 76.4870465 | 77.5315887 | 67.3110032 | 62.3254961 | 22.0947303 | 29.3501565 | 29.6486857 | 92.4399027 | 74.8089332 | 587.109378 |
| Motif:rnd-1\_family-246 | 56.0690332 | 55.728076 | 65.6012386 | 48.3086728 | 42.7495068 | 26.9667432 | 87.5838614 | 57.7937894 | 80.1222698 | 63.0655887 | 583.98878 |
| Motif:rnd-5\_family-4690 | 56.0068828 | 59.2599207 | 61.36177 | 50.5141406 | 73.1501044 | 33.8501941 | 45.5823786 | 40.796318 | 89.7255463 | 72.9521916 | 583.199447 |
| Motif:rnd-1\_family-429 | 61.3565258 | 61.1809626 | 69.5434136 | 57.6877616 | 50.8249259 | 20.48609 | 76.0192519 | 45.9398683 | 71.1711349 | 62.6426927 | 576.852627 |
| Motif:rnd-5\_family-1400 | 43.1507811 | 41.1708042 | 45.3799048 | 38.0946761 | 48.9046286 | 84.805956 | 66.5010168 | 73.3283455 | 78.9409687 | 56.5571103 | 576.834192 |
| Motif:rnd-1\_family-305 | 64.4816599 | 68.3508772 | 69.107617 | 57.3437986 | 84.5770248 | 14.1893008 | 16.8948818 | 17.4338043 | 100.429883 | 82.7762251 | 575.585072 |
| Motif:rnd-1\_family-142 | 62.1713075 | 53.1663524 | 60.1366502 | 54.3027852 | 58.5720523 | 48.0304965 | 47.0621022 | 47.5842899 | 88.5736721 | 55.4284491 | 575.028157 |
| Motif:rnd-1\_family-10 | 47.1428781 | 37.4280542 | 49.6793734 | 43.411177 | 49.6641326 | 72.4645763 | 61.353984 | 69.5693781 | 82.3847944 | 59.0970267 | 572.195375 |
| Motif:rnd-6\_family-1900 | 68.8208052 | 88.1651578 | 76.0376646 | 58.3587907 | 68.0355453 | 24.2002475 | 25.0398329 | 24.9225191 | 79.8130854 | 58.2416371 | 571.635286 |
| Motif:rnd-1\_family-1741 | 51.7289578 | 58.9134073 | 79.6341163 | 52.5520974 | 56.5633147 | 32.8862412 | 39.6054127 | 40.4232963 | 86.6766392 | 69.5992366 | 568.582719 |
| Motif:rnd-1\_family-1373 | 61.2124651 | 77.5601808 | 70.3104594 | 57.2446636 | 55.2291126 | 25.4917776 | 31.6214824 | 26.0398747 | 86.1832838 | 77.3597361 | 568.253036 |
| Motif:rnd-1\_family-905 | 68.8362206 | 69.2946934 | 68.9302247 | 59.6510337 | 60.0793027 | 22.2801864 | 20.1752137 | 19.9121853 | 98.5480745 | 79.5096316 | 567.216766 |
| Motif:rnd-1\_family-138 | 53.2449378 | 52.5452119 | 59.3416439 | 48.0535519 | 63.8997952 | 41.7707754 | 59.8453792 | 51.646567 | 73.714177 | 61.1930428 | 565.255082 |
| Motif:rnd-6\_family-1544 | 59.5520784 | 66.5793232 | 66.9000024 | 56.1401767 | 56.1642951 | 22.0739011 | 44.2077019 | 32.3754475 | 92.2420598 | 68.702048 | 564.937034 |
| Motif:rnd-4\_family-88 | 48.5402793 | 43.0693346 | 44.2155115 | 49.7264572 | 54.7250046 | 65.4531245 | 75.0851593 | 73.3087477 | 47.0559884 | 57.7389626 | 558.918569 |
| Motif:rnd-5\_family-5961 | 57.7560515 | 54.7740395 | 59.0414273 | 57.4123185 | 53.1649266 | 30.126521 | 48.276556 | 42.5044606 | 85.0584338 | 70.2143625 | 558.329097 |
| Motif:rnd-1\_family-56 | 46.1190719 | 39.6567622 | 42.6104127 | 45.7661593 | 46.5759388 | 57.3778642 | 101.001434 | 73.7931863 | 54.2265285 | 47.6391966 | 554.766555 |
| Motif:rnd-1\_family-109 | 58.8407373 | 55.6102045 | 64.2061459 | 58.0545183 | 56.1076348 | 29.1197303 | 31.5459865 | 30.270318 | 89.7581702 | 73.8676273 | 547.381073 |
| Motif:rnd-4\_family-236 | 44.5974104 | 25.8034309 | 43.596506 | 38.8522082 | 69.2904061 | 69.0942086 | 57.2935303 | 57.5547265 | 82.037778 | 58.6962086 | 546.816414 |
| Motif:rnd-1\_family-855 | 55.6129556 | 69.713628 | 63.970889 | 52.6344394 | 50.5497764 | 12.4069745 | 52.7312313 | 39.7299257 | 79.9284283 | 63.9785866 | 541.256835 |
| Motif:rnd-1\_family-6 | 51.163453 | 47.7409278 | 55.5098249 | 55.0111978 | 57.9906626 | 54.6450904 | 57.0512038 | 53.939093 | 54.5222249 | 53.1055409 | 540.679219 |
| Motif:rnd-6\_family-5246 | 43.6646729 | 42.9106892 | 45.9243708 | 48.0812421 | 43.7752906 | 67.7826077 | 58.0169957 | 60.402945 | 78.9661461 | 49.3117207 | 538.836681 |
| Motif:rnd-4\_family-281 | 54.3056013 | 41.9953974 | 52.3397316 | 47.4344708 | 46.5015224 | 54.4433235 | 44.6853187 | 56.3987327 | 79.7919344 | 55.6132294 | 533.509262 |
| Motif:rnd-5\_family-233 | 44.6560004 | 41.1805154 | 41.4880722 | 44.4642444 | 53.7431992 | 62.8118989 | 62.7050165 | 64.628851 | 63.8645829 | 53.7783611 | 533.320742 |
| Motif:rnd-6\_family-30 | 50.0174608 | 38.3507324 | 43.826652 | 41.3999246 | 40.054967 | 61.7665891 | 54.876976 | 64.7194755 | 80.3350116 | 55.5377499 | 530.885539 |
| Motif:rnd-1\_family-184 | 58.1553544 | 77.9335759 | 56.2499143 | 57.2397485 | 59.4641703 | 18.5651305 | 31.086807 | 27.9125549 | 80.8081635 | 60.7045009 | 528.11992 |
| Motif:rnd-5\_family-1416 | 45.8896508 | 41.4085833 | 54.4754953 | 45.0217742 | 45.4506712 | 79.5224643 | 66.7700697 | 64.3208911 | 34.6722754 | 49.4924548 | 527.02433 |
| Motif:rnd-6\_family-4724 | 53.3232446 | 67.0197658 | 64.2040255 | 44.9115011 | 83.0083046 | 13.7818449 | 32.8466301 | 29.2779061 | 74.0037597 | 63.4578765 | 525.834859 |
| Motif:rnd-5\_family-1129 | 36.3944884 | 27.4202605 | 49.9848648 | 39.0416621 | 51.6101736 | 64.2666736 | 54.6032083 | 58.9897134 | 80.7670663 | 61.9862886 | 525.0644 |
| Motif:rnd-6\_family-6087 | 53.6177065 | 53.2754431 | 47.9131614 | 55.4610881 | 62.1682555 | 29.5862025 | 49.7786699 | 37.9493469 | 72.1952853 | 60.6125266 | 522.557686 |
| Motif:rnd-6\_family-7467 | 62.9335794 | 72.8562405 | 63.1288867 | 57.3668601 | 56.5358283 | 15.953733 | 23.0675154 | 20.2991579 | 82.6832287 | 67.5743813 | 522.399411 |
| Motif:rnd-4\_family-2163 | 57.5853804 | 42.749981 | 49.1905801 | 46.4291445 | 45.5181703 | 70.3516372 | 73.2266603 | 67.4167654 | 12.8979655 | 52.9059035 | 518.272188 |
| Motif:rnd-1\_family-614 | 60.5104528 | 47.3775283 | 40.0686198 | 48.5625886 | 51.7103625 | 50.9763535 | 68.1191062 | 66.445585 | 24.1160165 | 60.0750603 | 517.961674 |
| Motif:rnd-1\_family-238 | 45.1245161 | 43.292318 | 48.3364764 | 39.0761522 | 35.9957768 | 38.2794343 | 86.965217 | 67.2299973 | 60.7607298 | 52.6539136 | 517.714532 |
| Motif:rnd-1\_family-1879 | 49.501496 | 58.8771195 | 58.0963911 | 47.2924434 | 36.9604656 | 15.541325 | 67.0373743 | 40.4322363 | 80.0403672 | 63.1785594 | 516.957778 |
| Motif:rnd-5\_family-3626 | 44.3245679 | 57.0267464 | 53.6161046 | 48.0849552 | 43.9420334 | 21.3058865 | 63.2793672 | 46.5315951 | 74.4611583 | 63.2846681 | 515.857083 |
| Motif:rnd-1\_family-20 | 40.3999921 | 44.6448048 | 45.4997137 | 41.1525649 | 41.6635897 | 7.72362383 | 124.616827 | 53.2495806 | 64.6286813 | 51.6666013 | 515.245979 |
| Motif:rnd-1\_family-800 | 53.5081433 | 68.1769836 | 63.7739704 | 54.9255939 | 54.4279109 | 12.0652242 | 34.2672306 | 21.0608734 | 79.0531265 | 69.3150009 | 510.574058 |
| Motif:rnd-1\_family-809 | 41.573519 | 48.6768836 | 48.8094069 | 45.8883913 | 59.8714655 | 26.972046 | 62.9839623 | 49.3427236 | 69.0328045 | 55.7012268 | 508.85243 |
| Motif:rnd-1\_family-68 | 46.0162387 | 30.9538664 | 50.2047206 | 32.8024132 | 36.786445 | 59.6729842 | 56.8059761 | 61.5742799 | 75.5064891 | 57.631845 | 507.955258 |
| Motif:rnd-6\_family-7630 | 36.7230621 | 32.7823435 | 44.545985 | 33.2448123 | 34.61071 | 14.8303202 | 158.550166 | 63.6639406 | 48.130451 | 39.6121556 | 506.693946 |
| Motif:rnd-6\_family-4336 | 37.6998431 | 45.3205372 | 41.6187157 | 34.1958077 | 45.6168105 | 68.8470638 | 53.2754347 | 70.4024535 | 62.0807337 | 44.1957974 | 503.253197 |
| Motif:rnd-5\_family-5461 | 57.1083863 | 54.8016736 | 66.5527085 | 45.7320853 | 62.932486 | 25.297054 | 22.0014935 | 23.9787144 | 81.0386564 | 63.7230458 | 503.166304 |
| Motif:rnd-6\_family-11892 | 54.2945459 | 53.3776393 | 57.0216431 | 54.2451385 | 57.9800523 | 22.5616574 | 25.2259974 | 28.1957874 | 79.4295179 | 69.9035179 | 502.235497 |
