## Supplementary material for "A novel eukaryotic RdRP-dependent small RNA pathway represses antiviral immunity by controlling an ERK pathway component in the black-legged tick": TE_cDNA_links.html

ISE6 sRNA counts


### Links for sRNA count tables

```
        TE Normalized sRNA level
        TE Relative sRNA level
        
        CDS Normalized sRNA level
        CDS Relative sRNA level
```

##### Note that TEs with less than 50RPM or Coding Genes with less than 3.5RPM on average in KD libraries are not shown.

##### See Supplementary table for the full information
