## Supplementary figures and images for "A novel eukaryotic RdRP-dependent small RNA pathway represses antiviral immunity by controlling an ERK pathway component in the black-legged tick"

### ISCI002834.RACDS_lengthdistribution.pdf

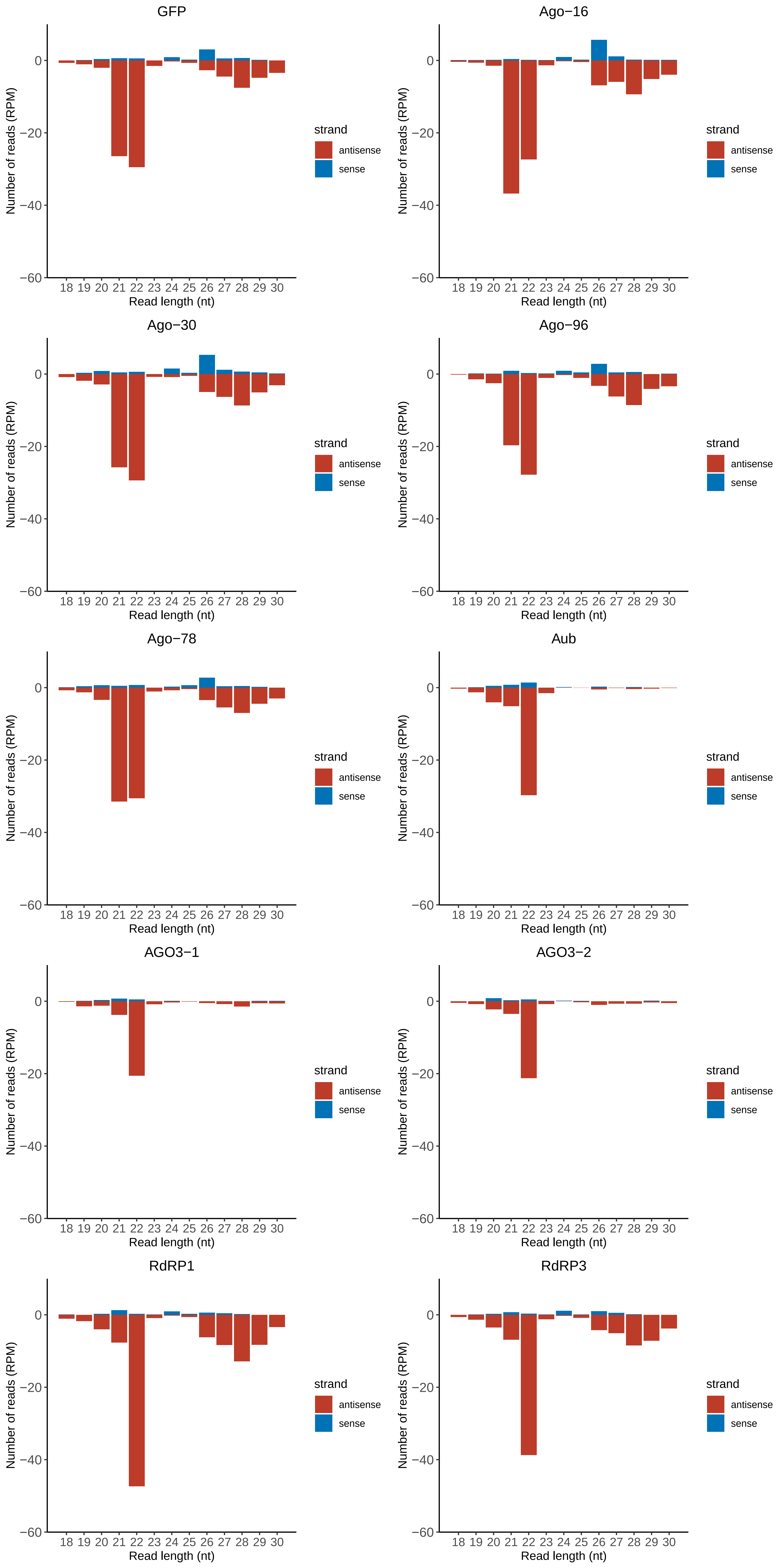

### ISCI007053.RACDS_lengthdistribution.pdf

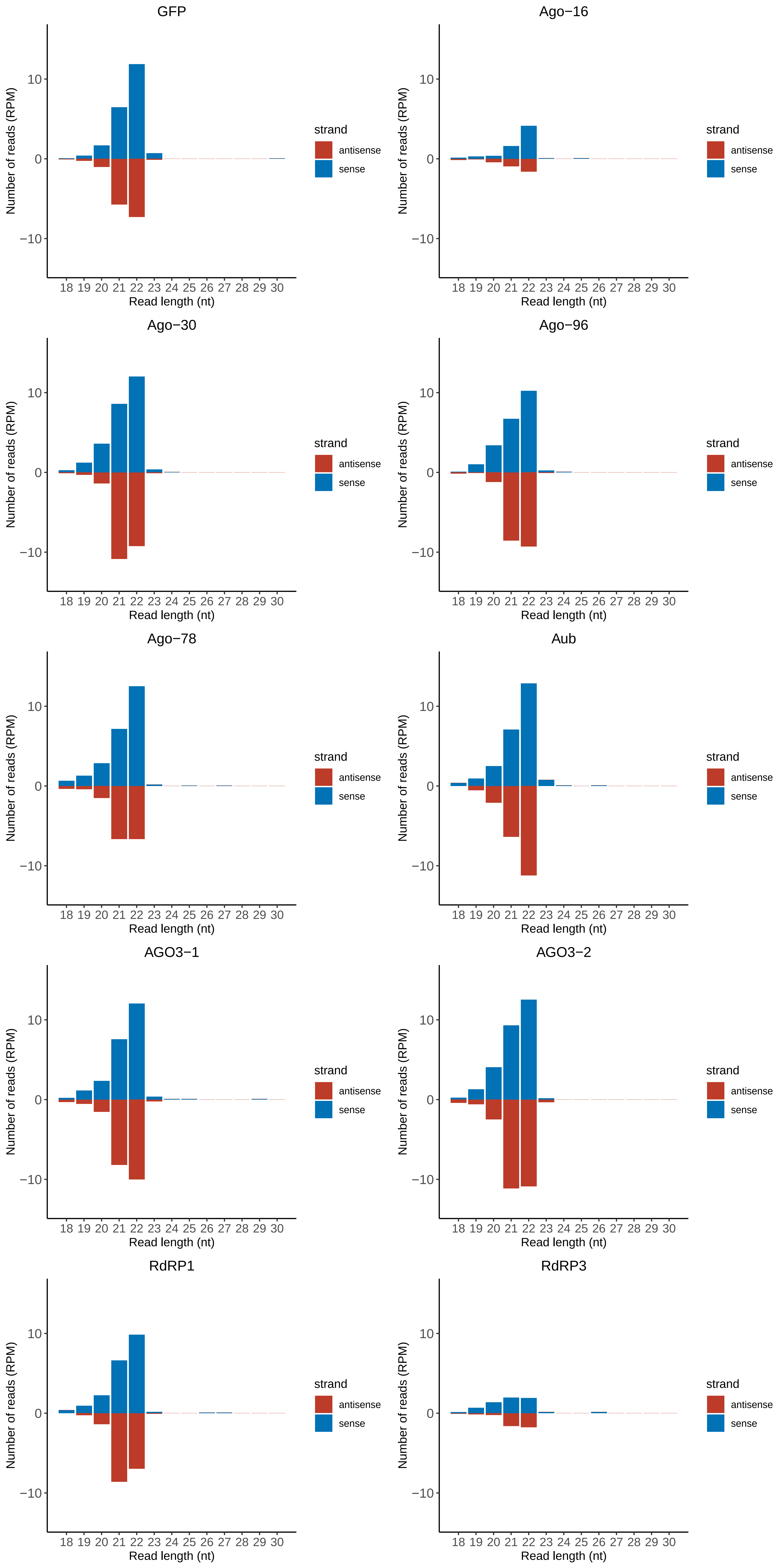

### ISCI007738.RACDS_lengthdistribution.pdf

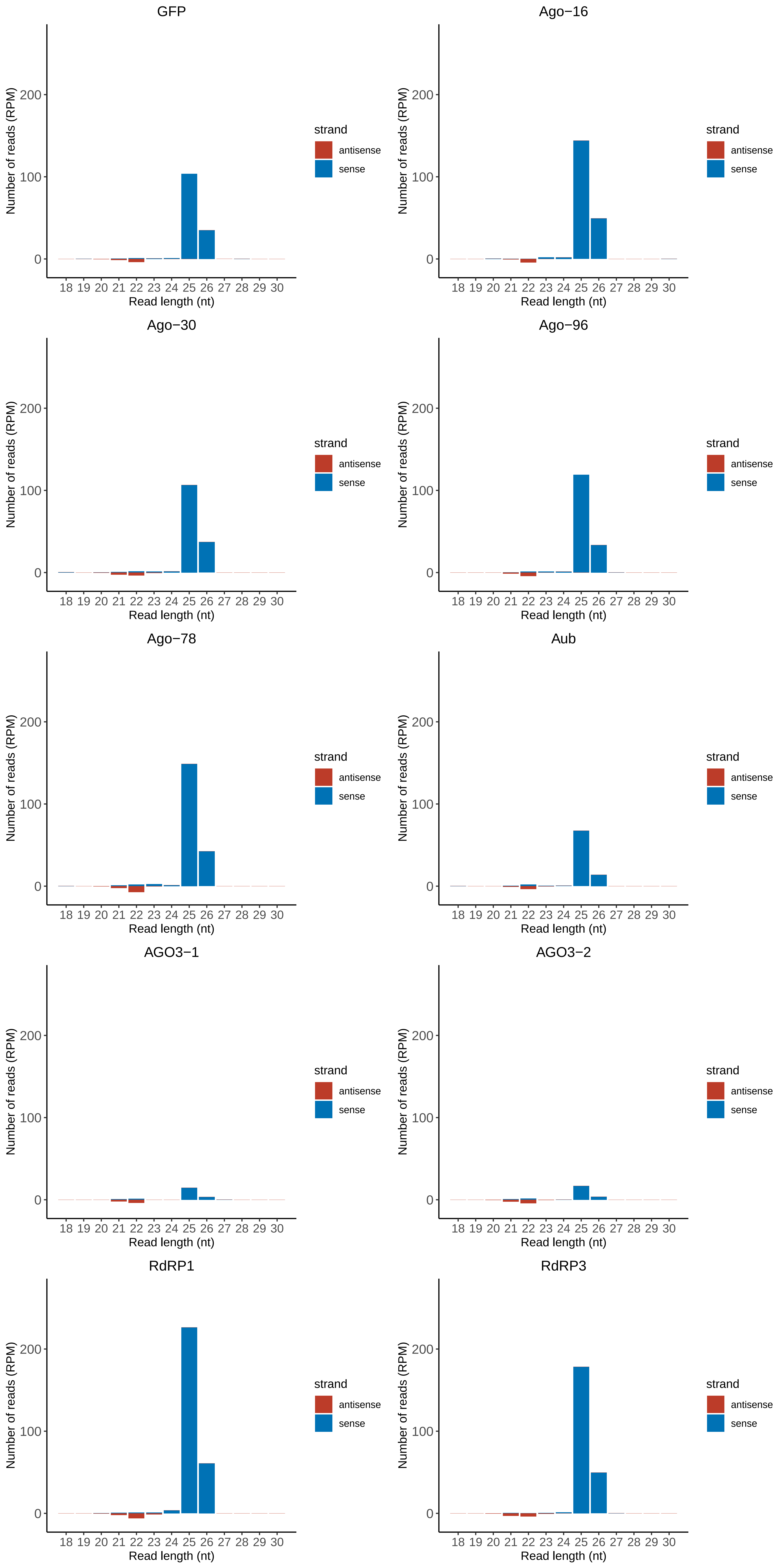

### ISCI011355.RACDS_lengthdistribution.pdf

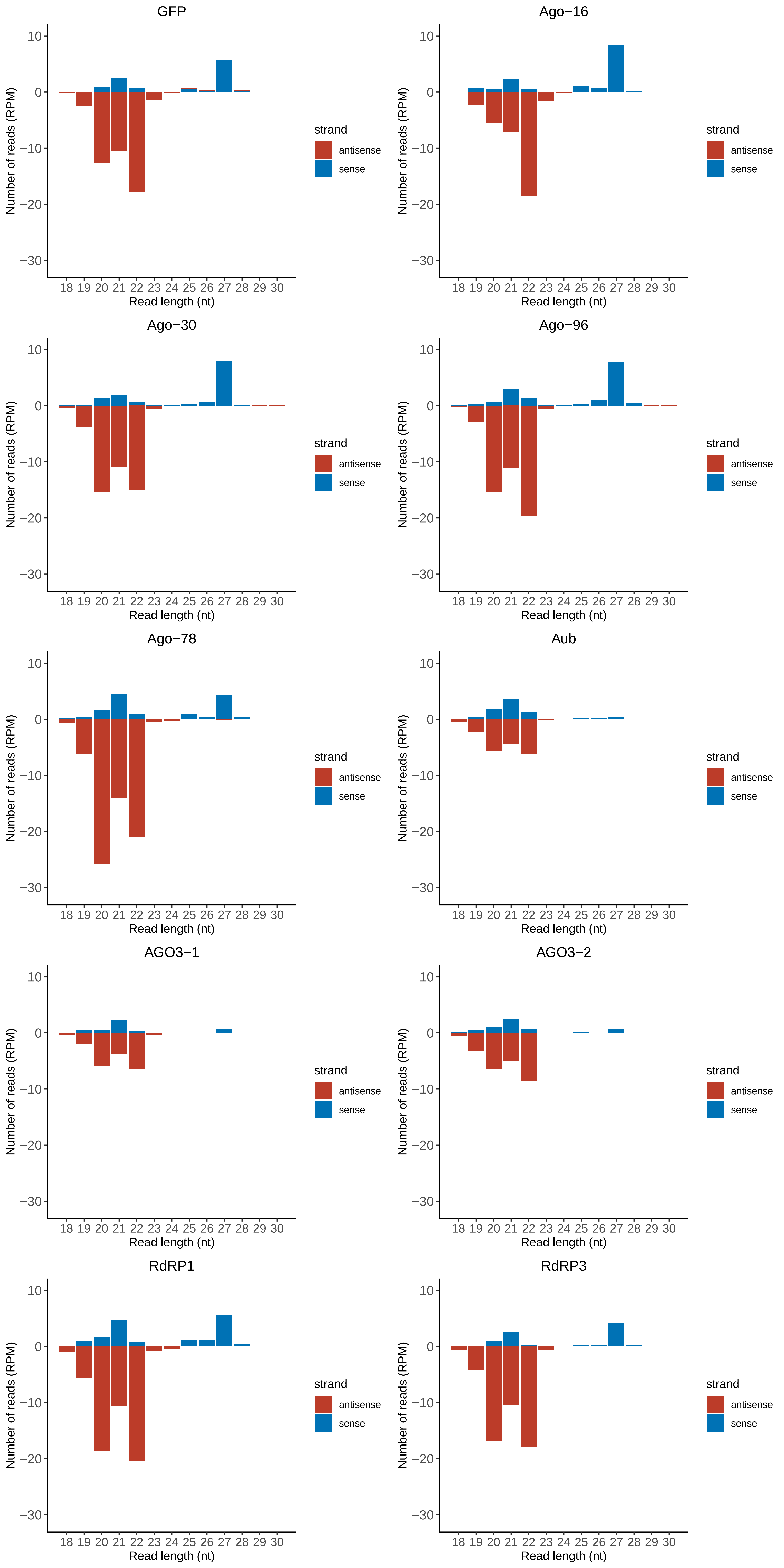

### ISCI015077.RACDS_lengthdistribution.pdf

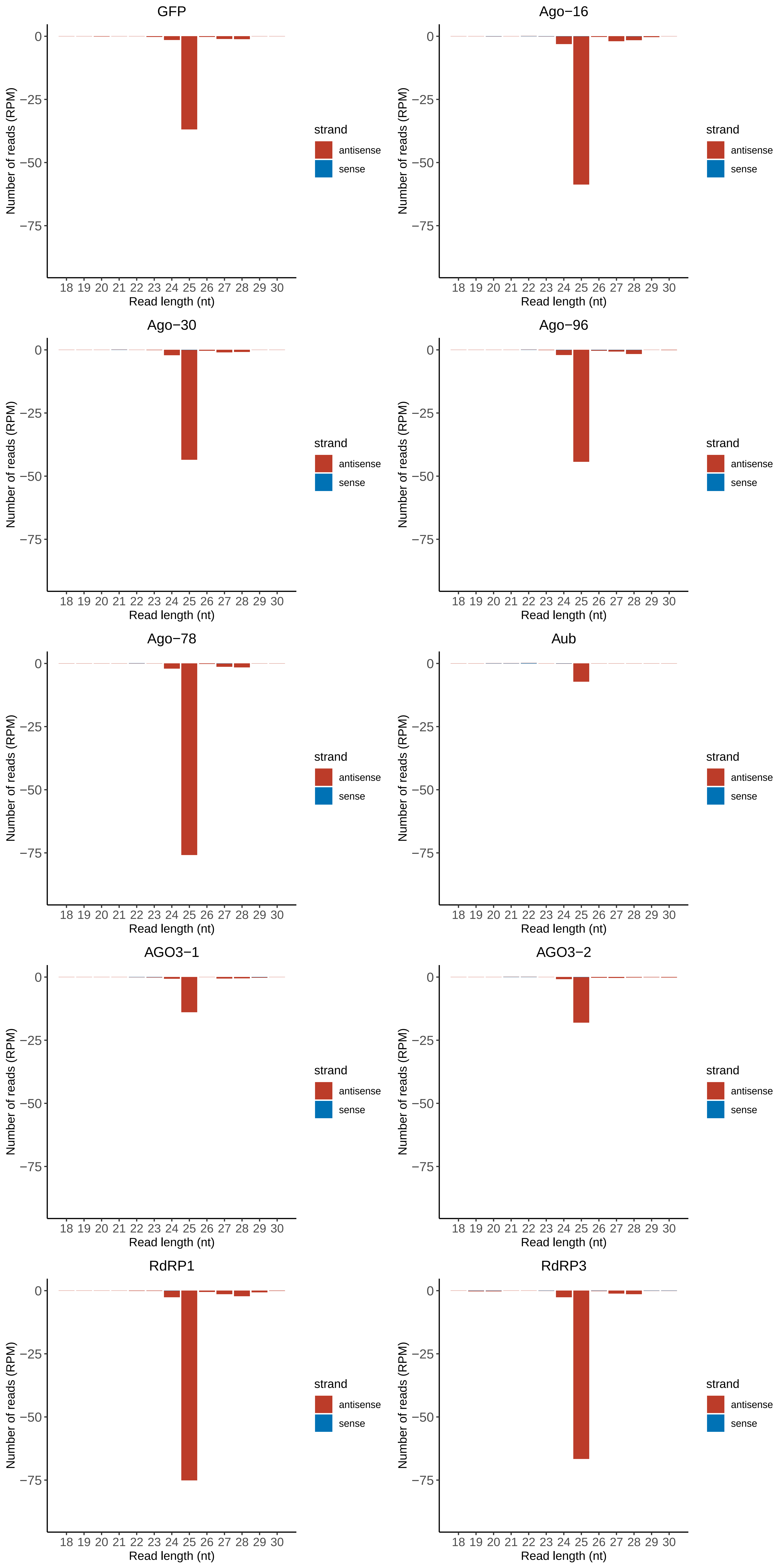

### ISCI017059.RACDS_lengthdistribution.pdf

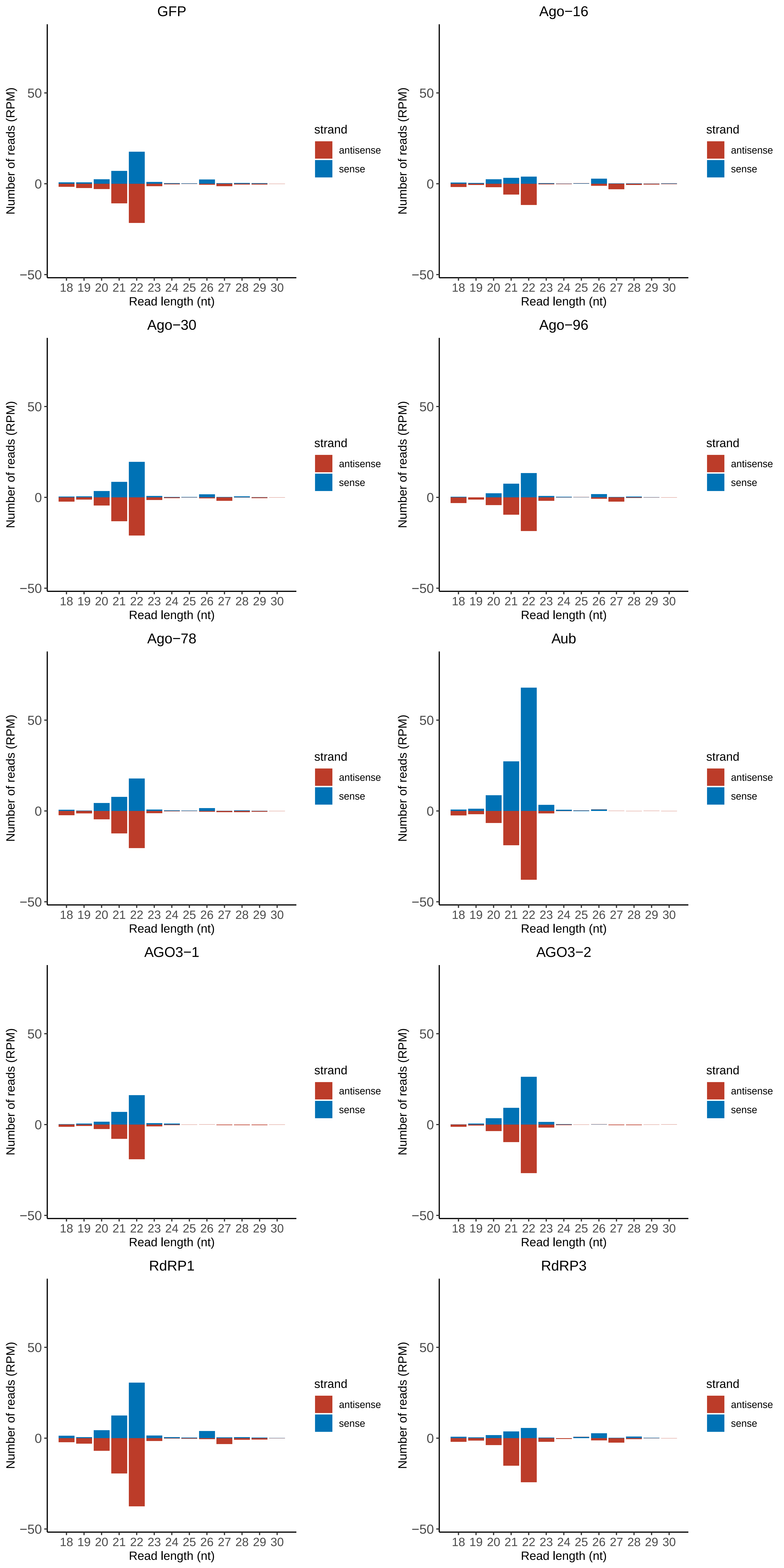

### Motif:rnd-1_family-54TE_lengthdistribution.pdf

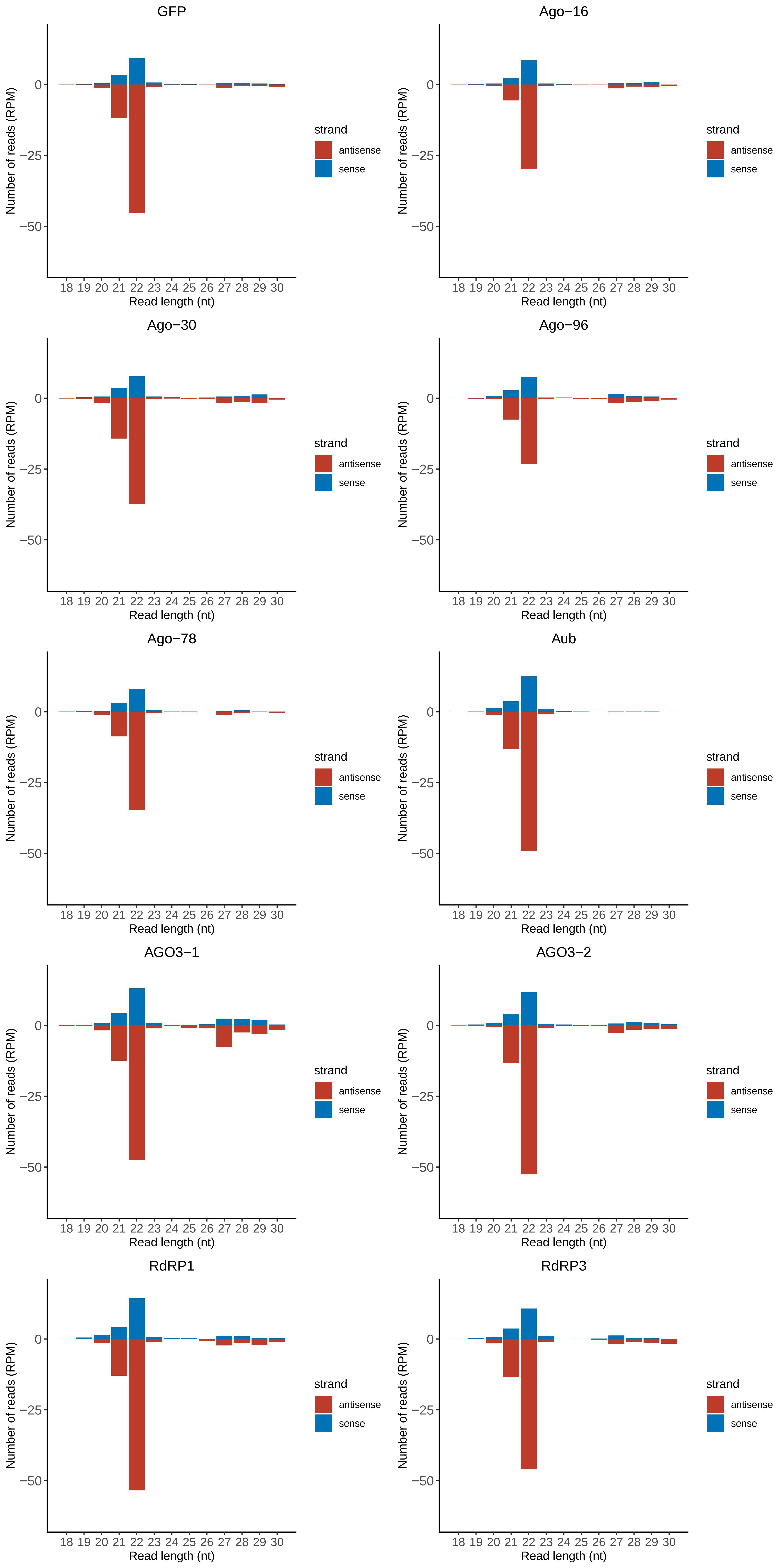

### Motif:rnd-1_family-63TE_lengthdistribution.pdf

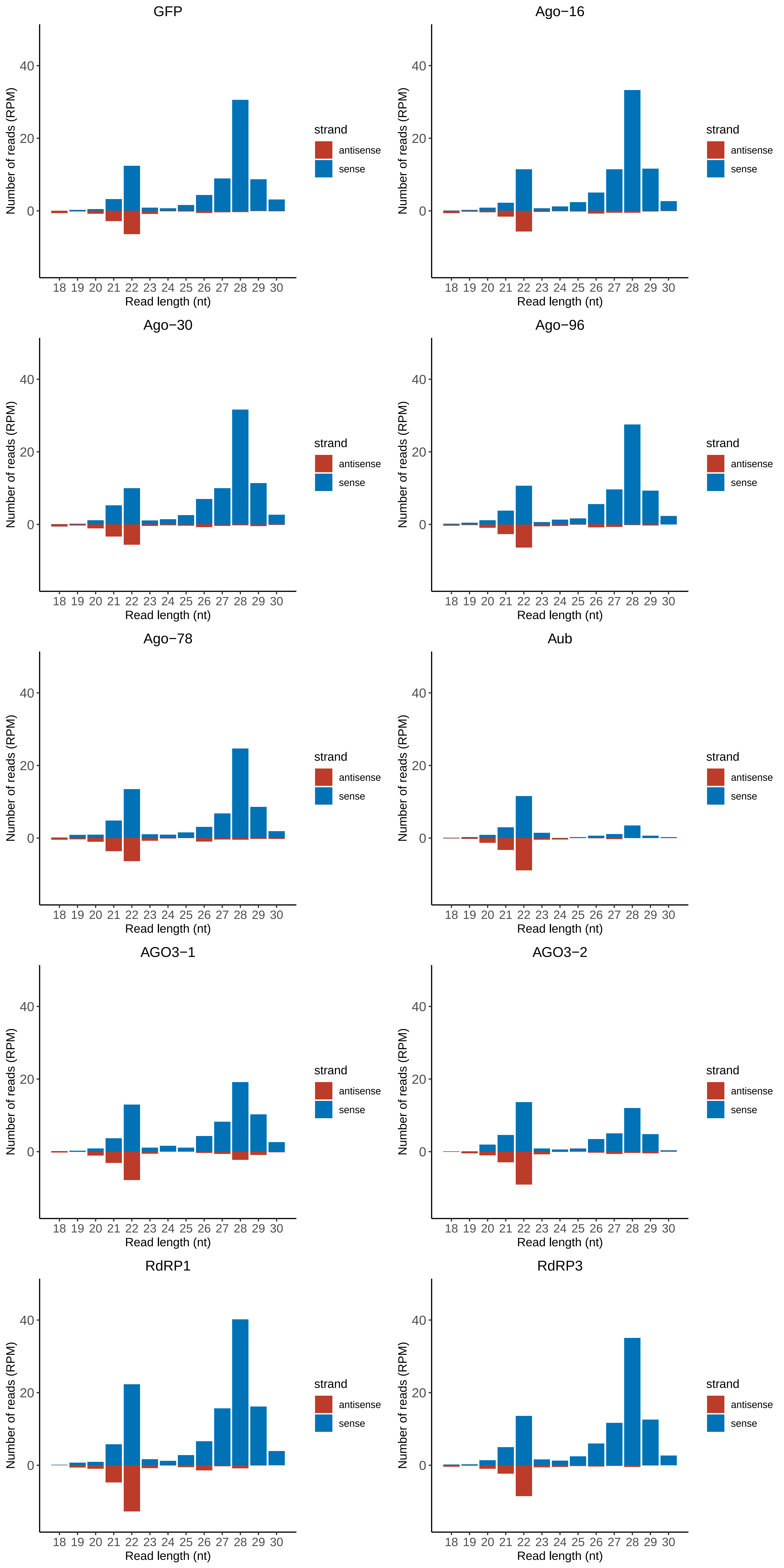

### Motif:rnd-1_family-84TE_lengthdistribution.pdf

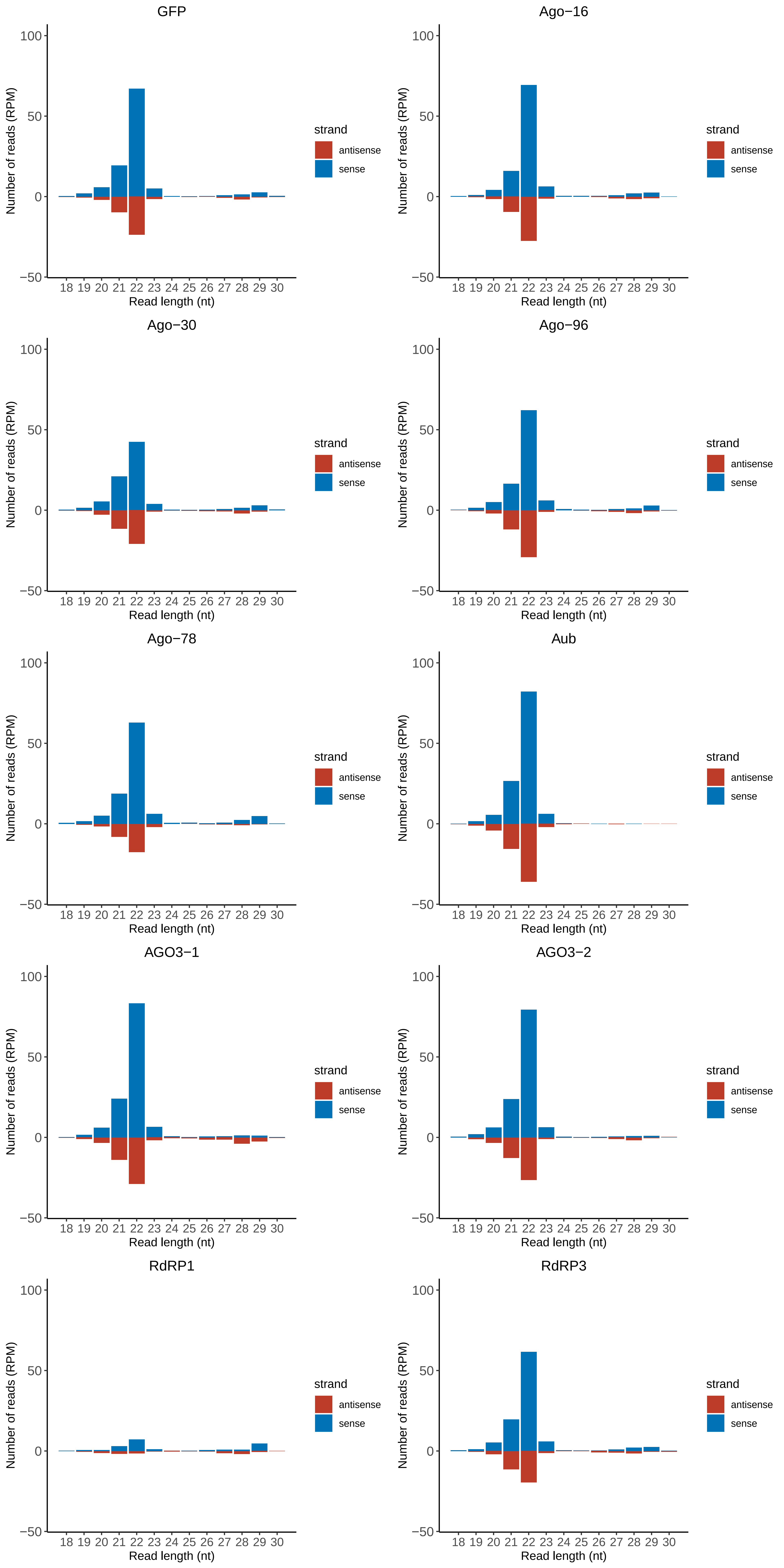

### Motif:rnd-1_family-88TE_lengthdistribution.pdf

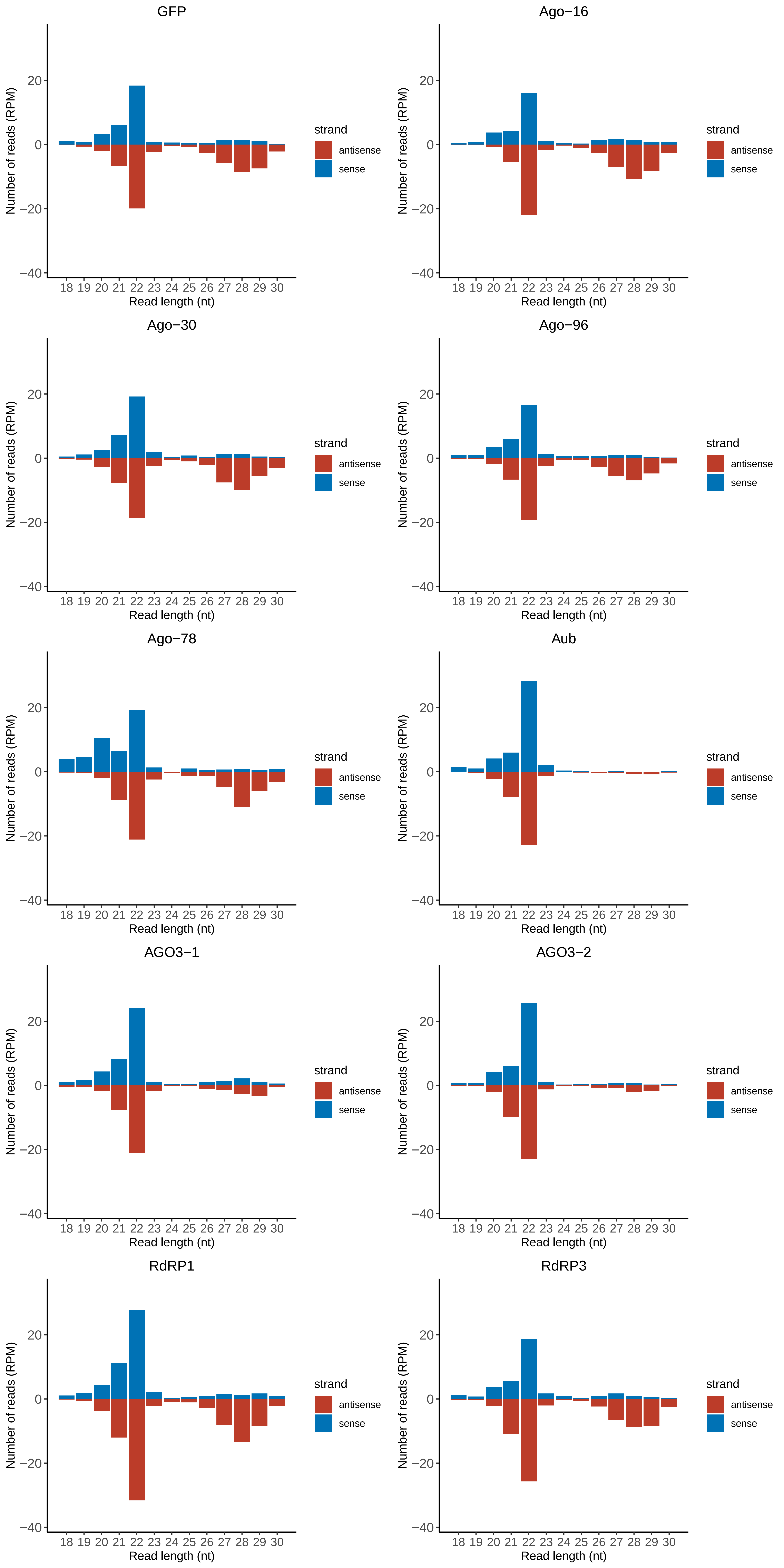

### Motif:rnd-1_family-103TE_lengthdistribution.pdf

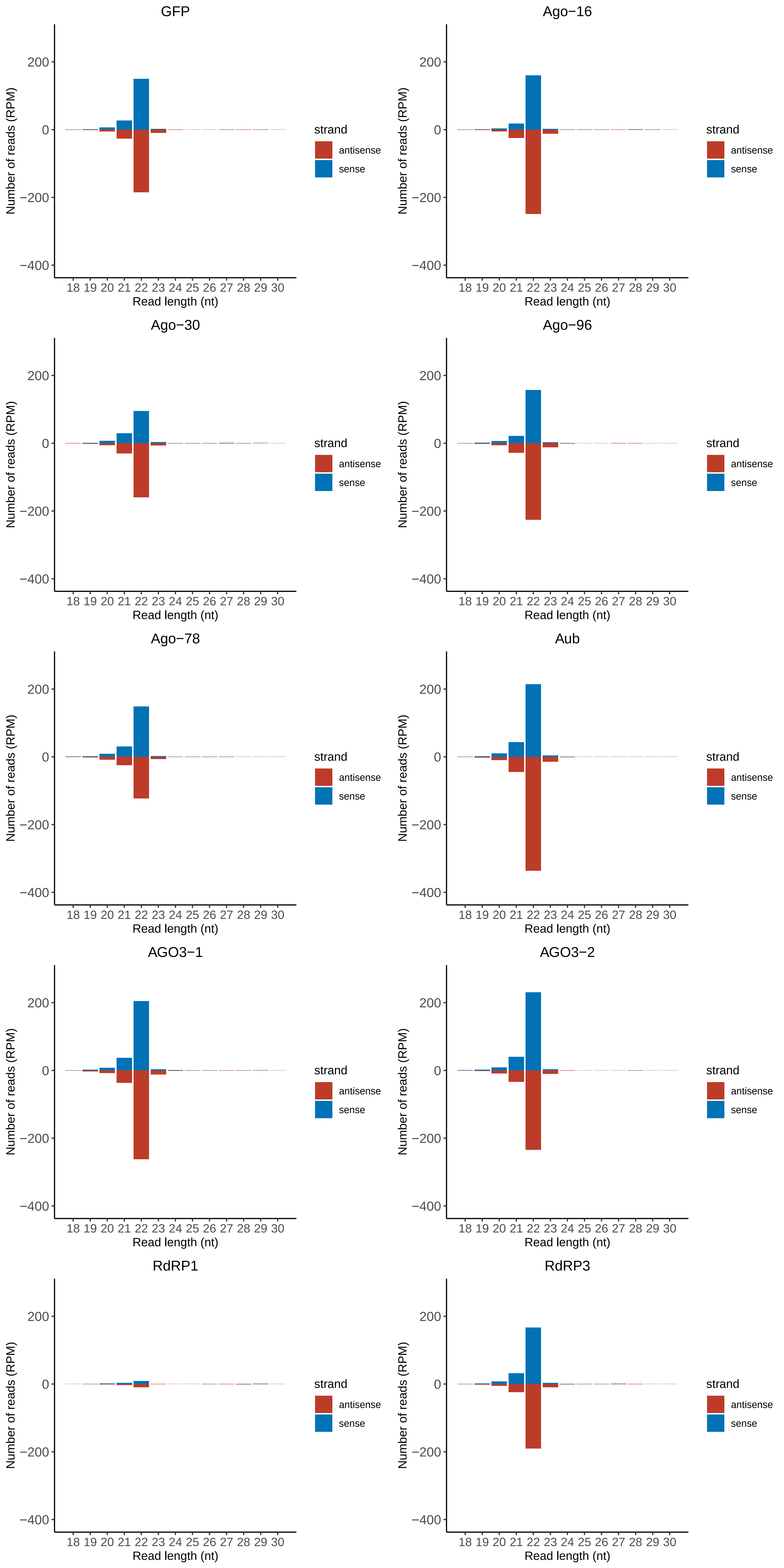

### Motif:rnd-1_family-134TE_lengthdistribution.pdf

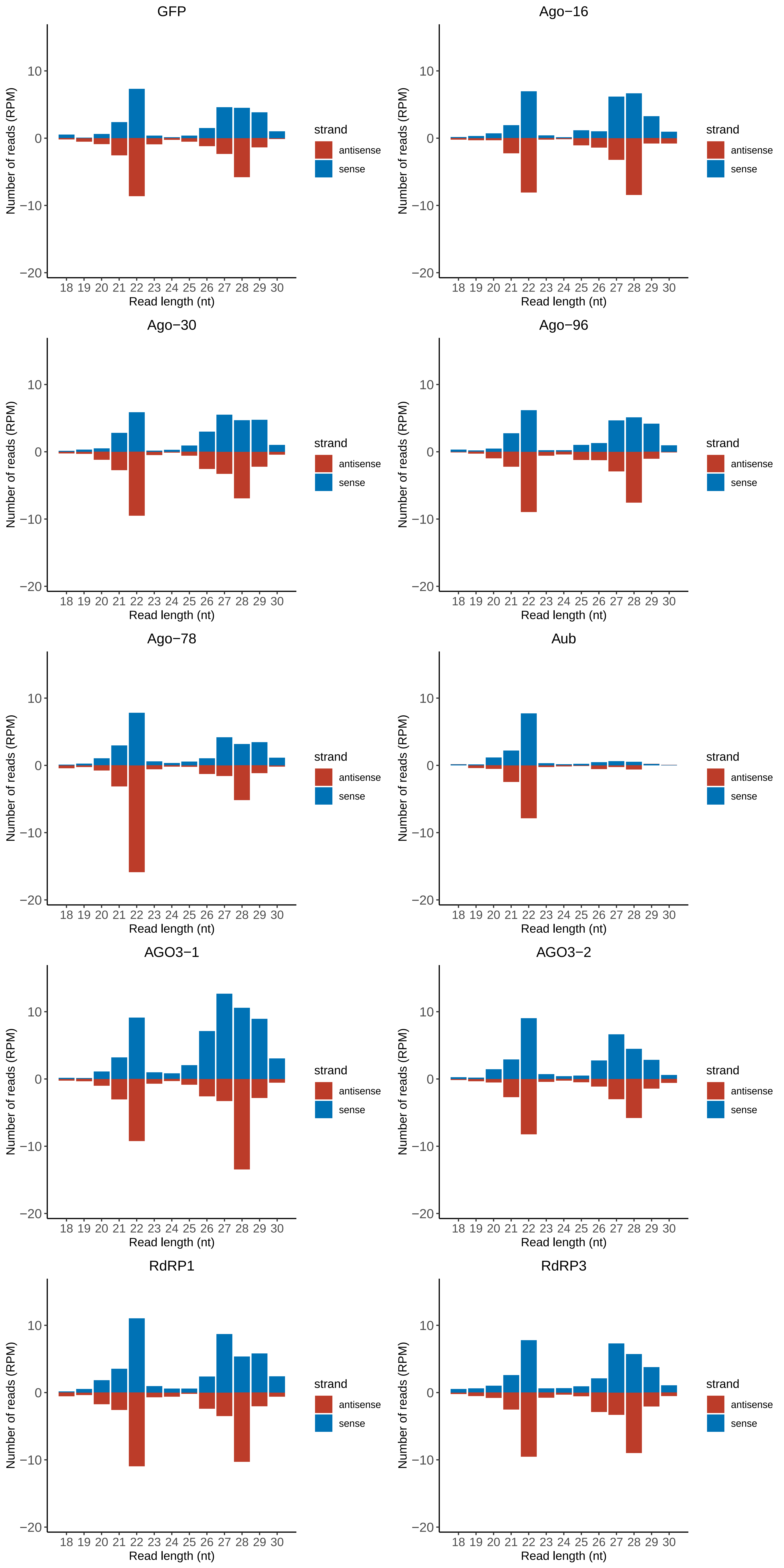

### Motif:rnd-1_family-139TE_lengthdistribution.pdf

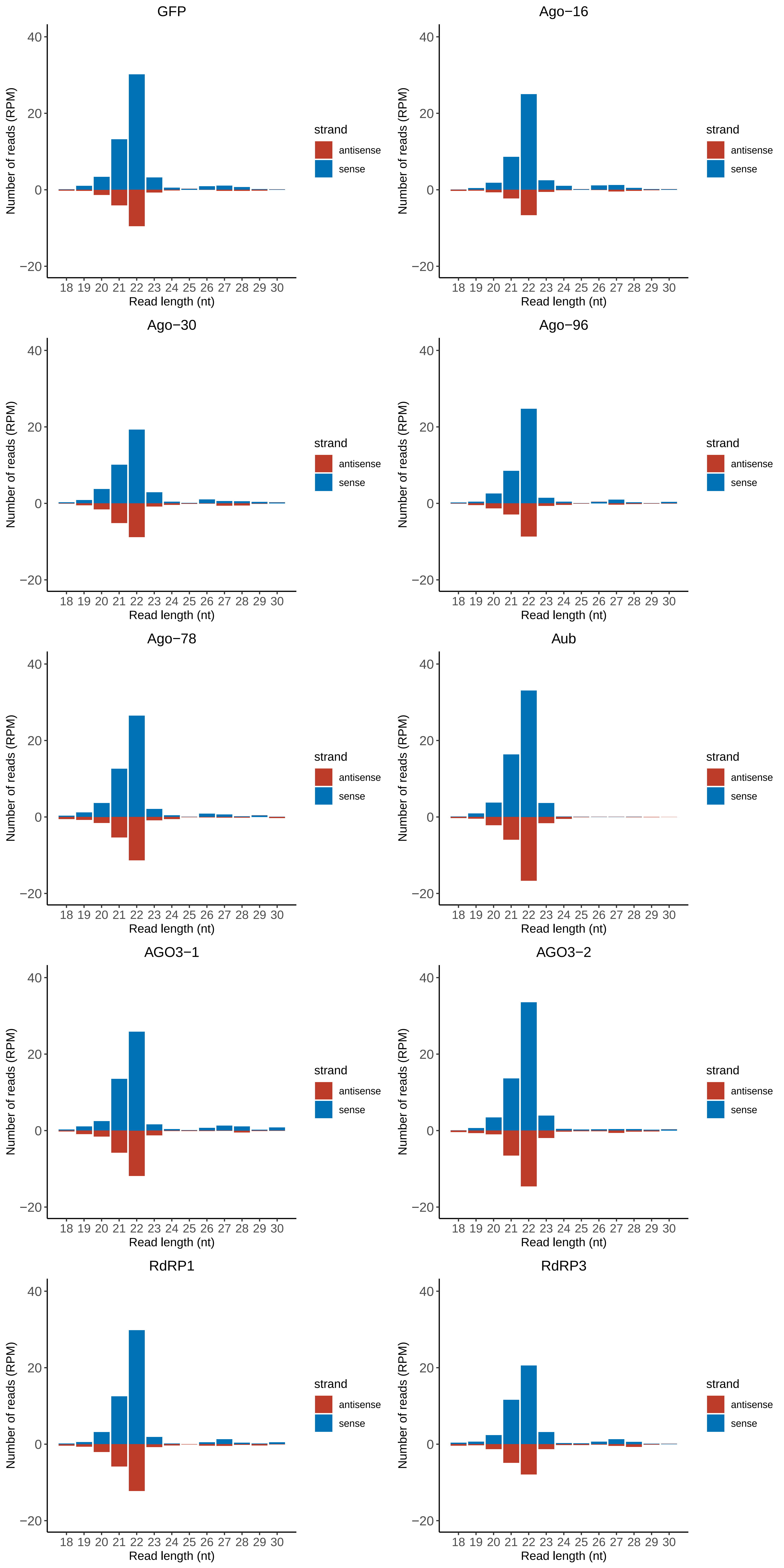

### Motif:rnd-1_family-152TE_lengthdistribution.pdf

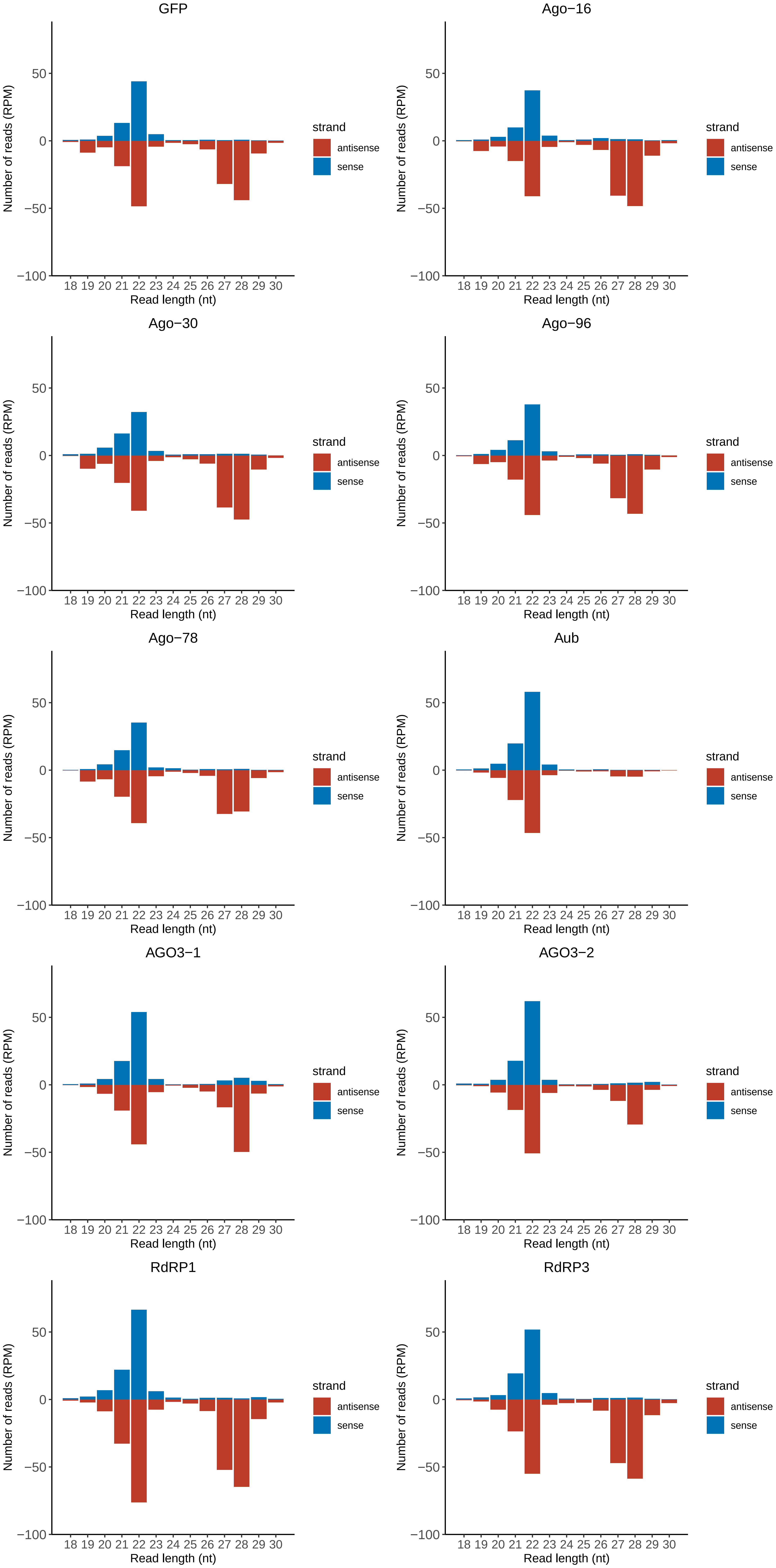

### Motif:rnd-1_family-157TE_lengthdistribution.pdf

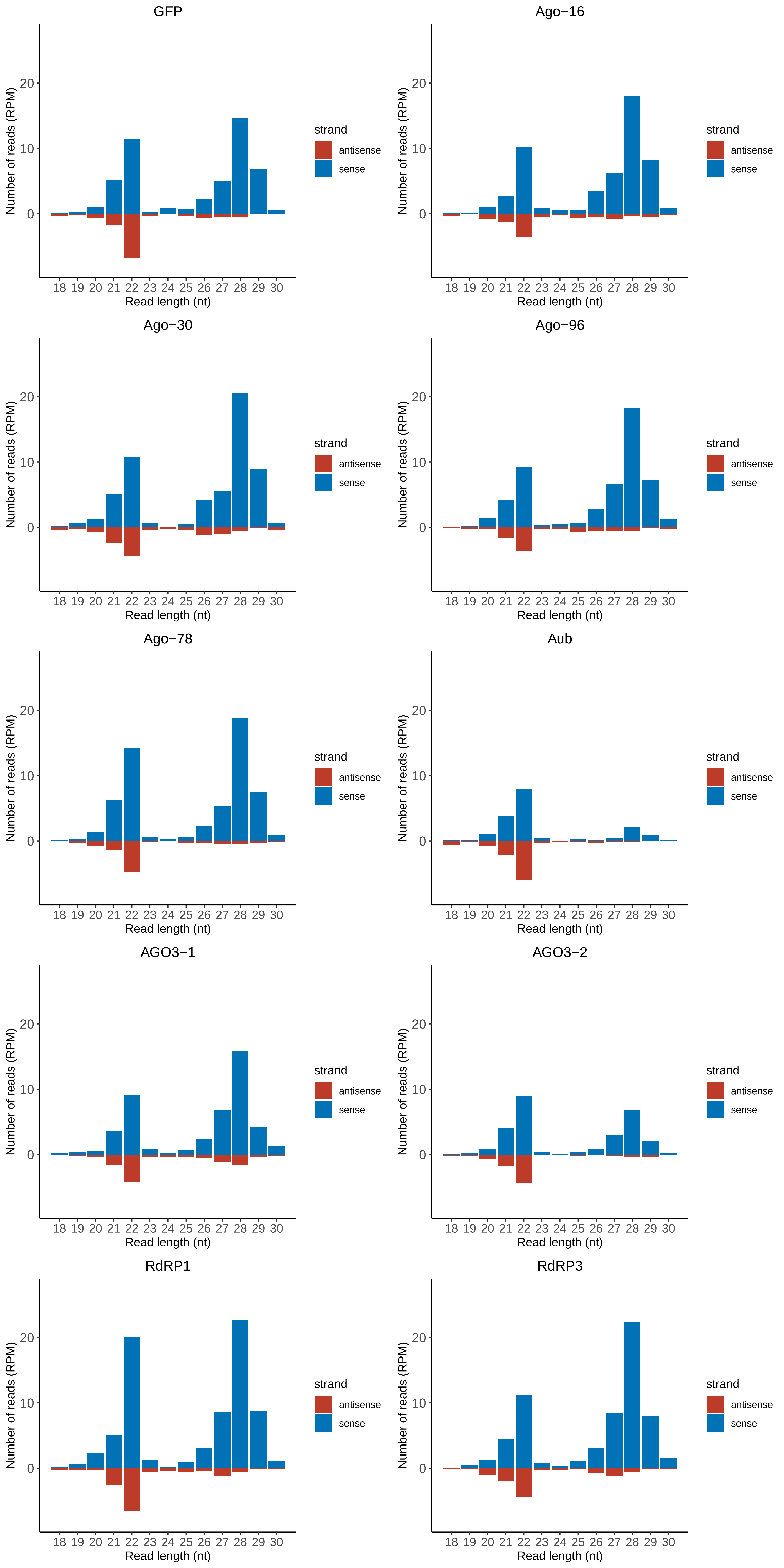

### Motif:rnd-1_family-164TE_lengthdistribution.pdf

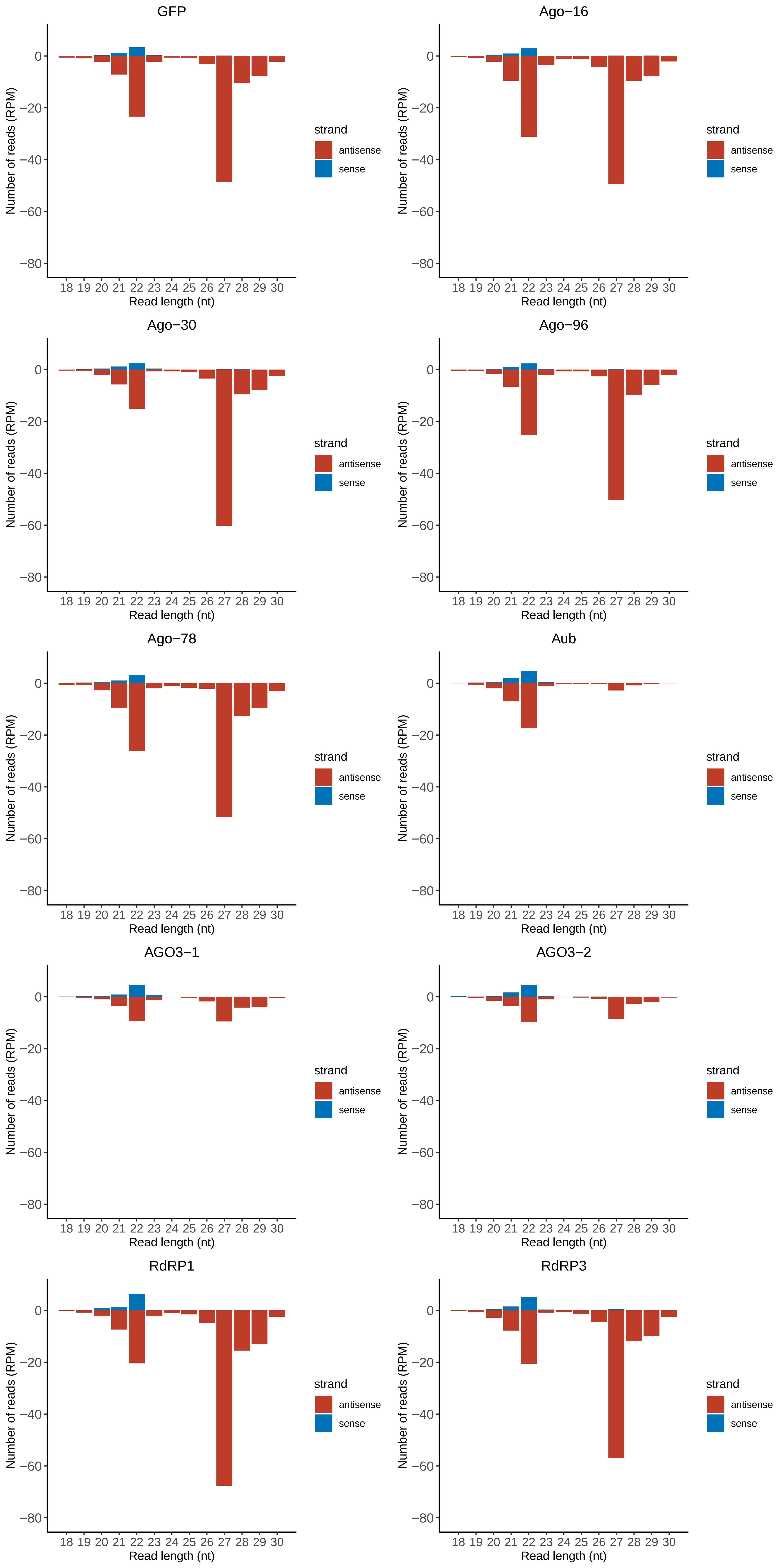

### Motif:rnd-1_family-169TE_lengthdistribution.pdf

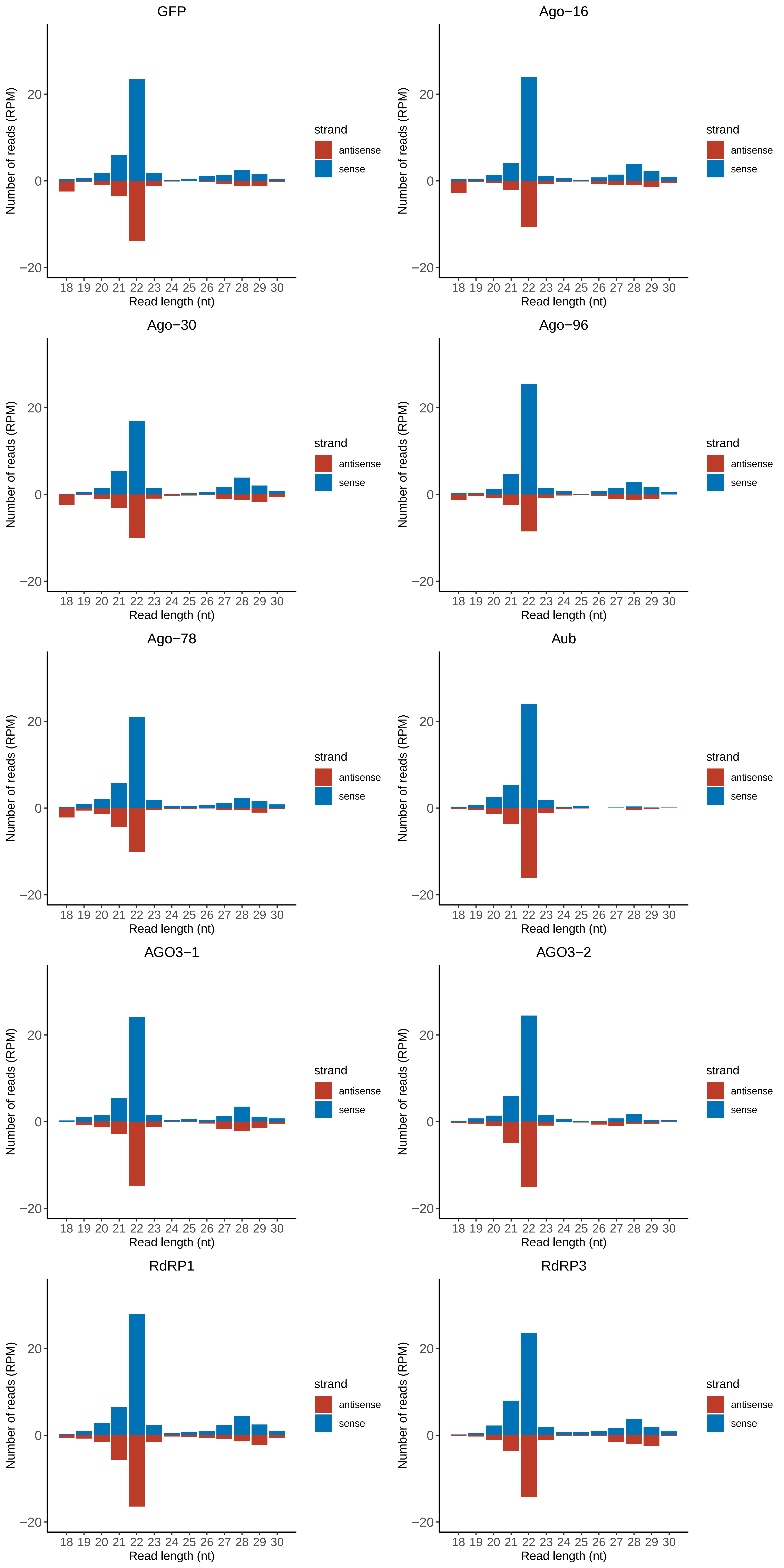

### Motif:rnd-1_family-201TE_lengthdistribution.pdf

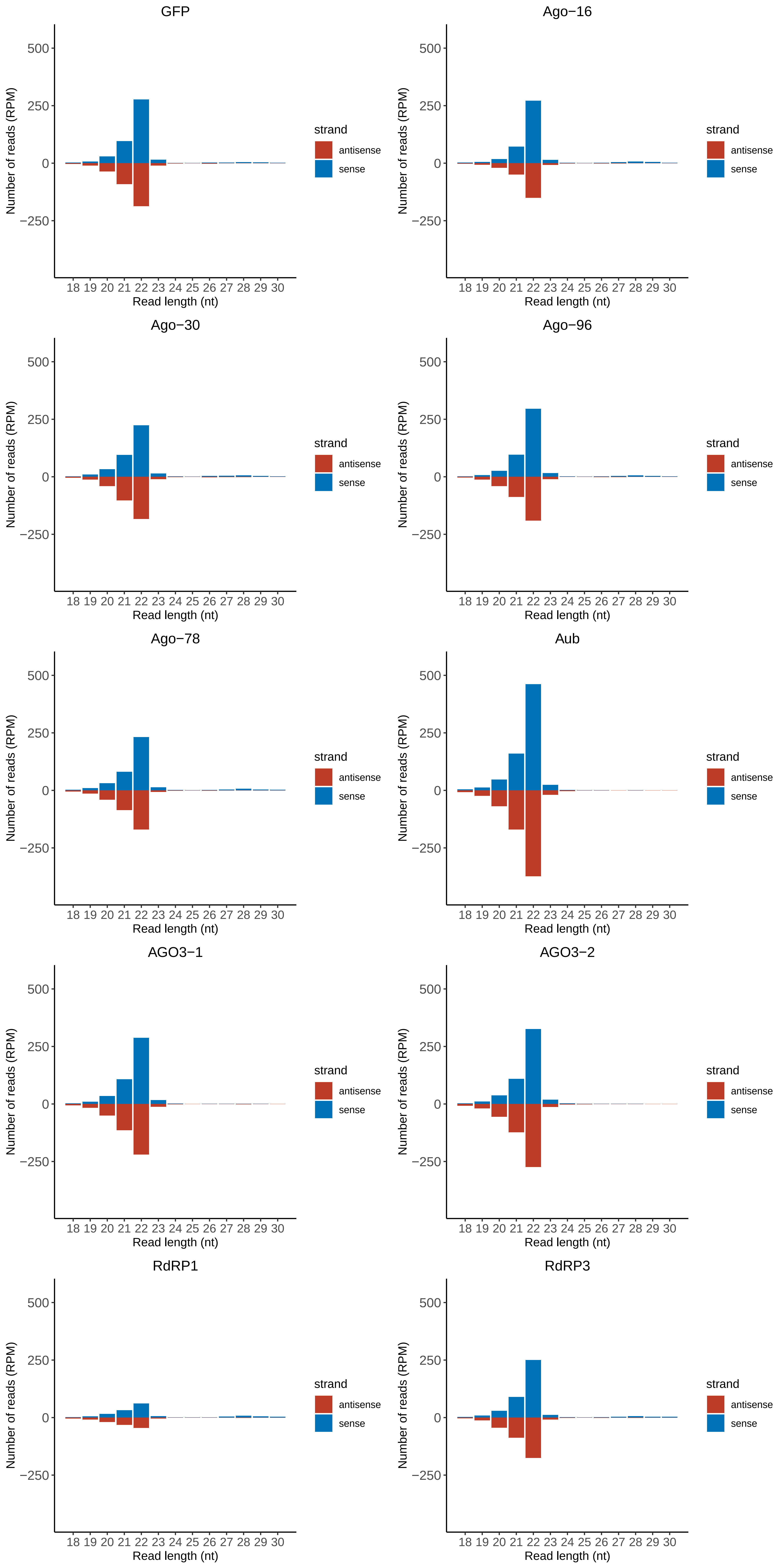

### Motif:rnd-1_family-337TE_lengthdistribution.pdf

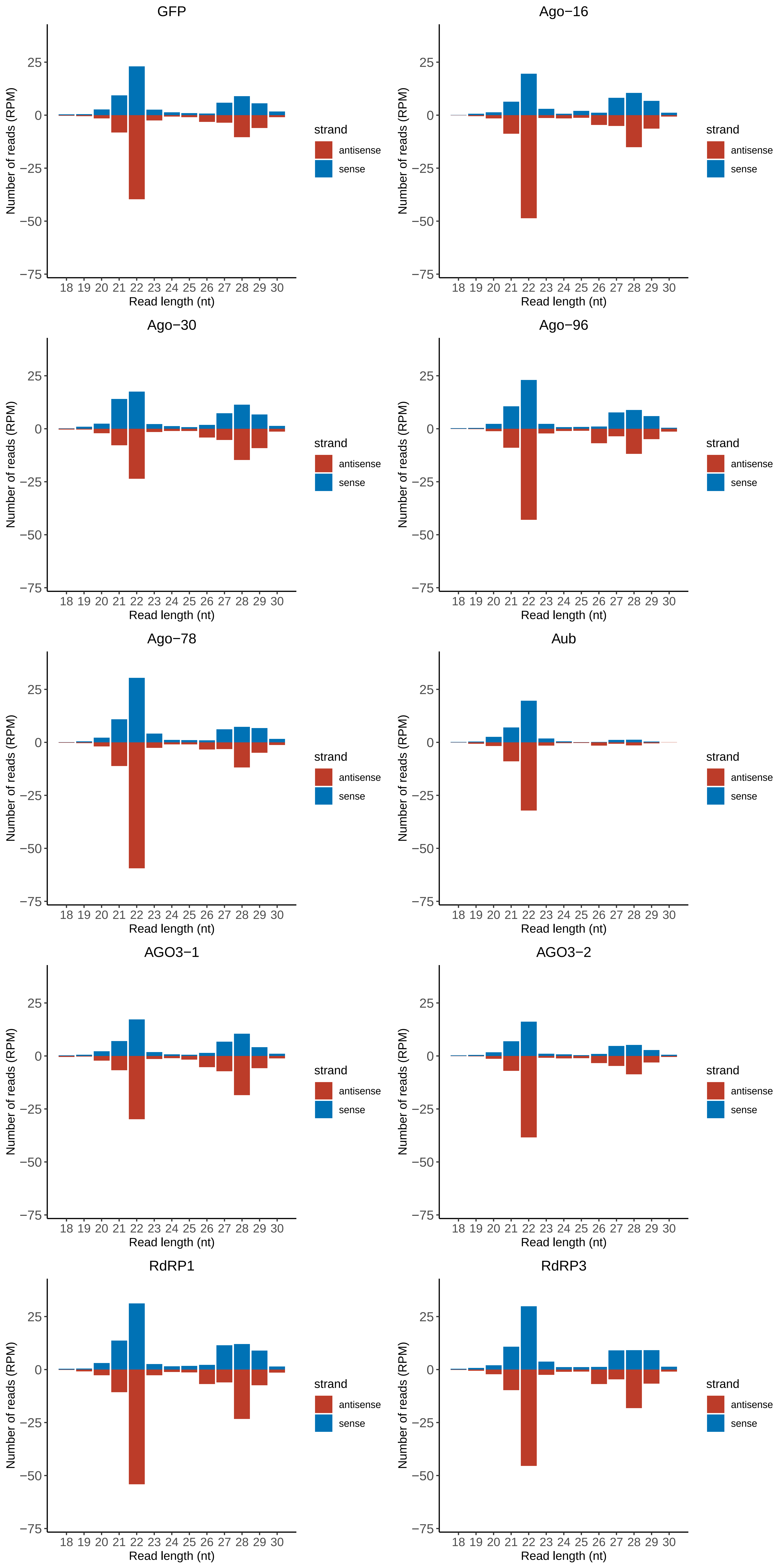

### Motif:rnd-1_family-384TE_lengthdistribution.pdf

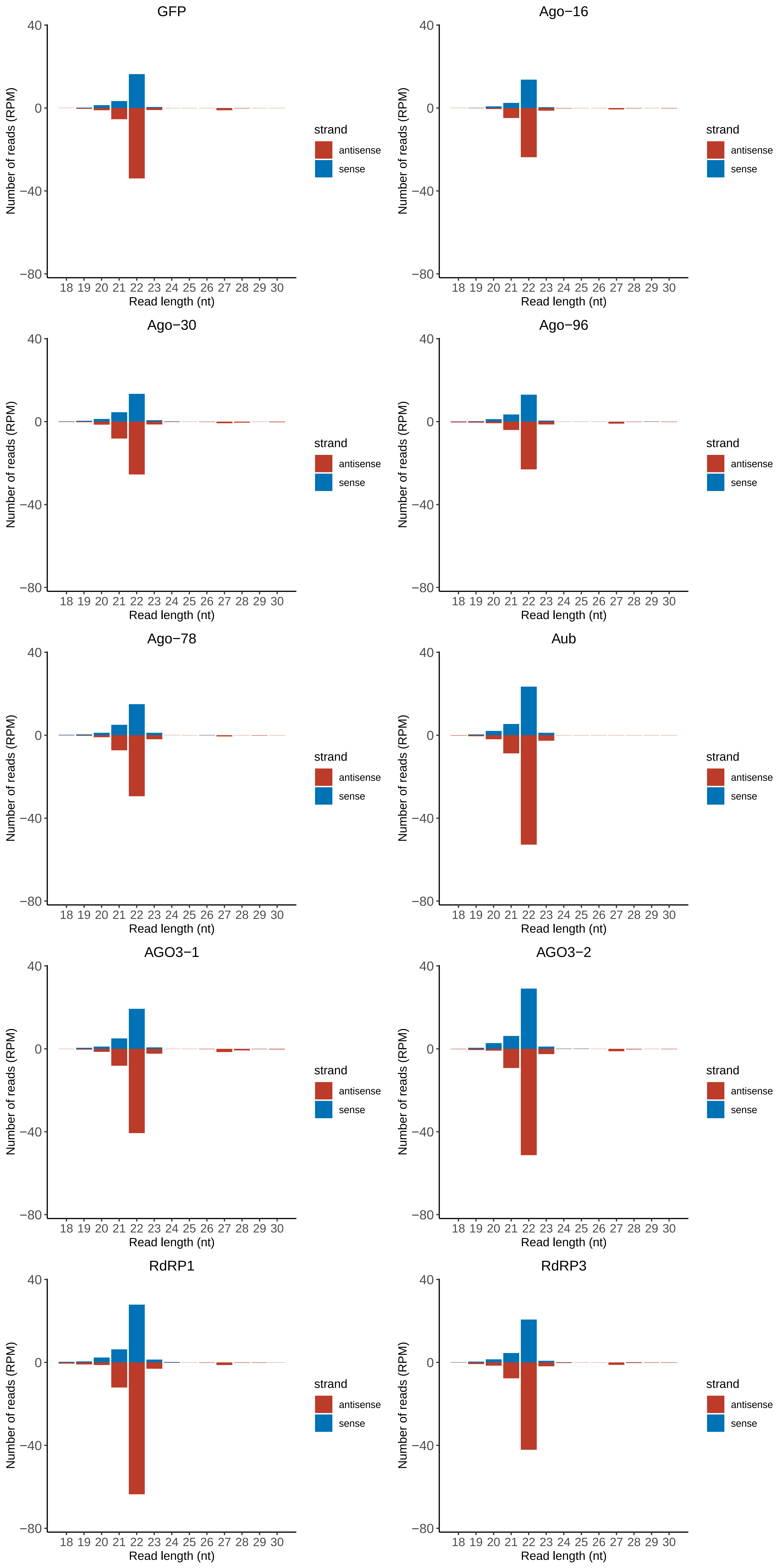

### Motif:rnd-1_family-385TE_lengthdistribution.pdf

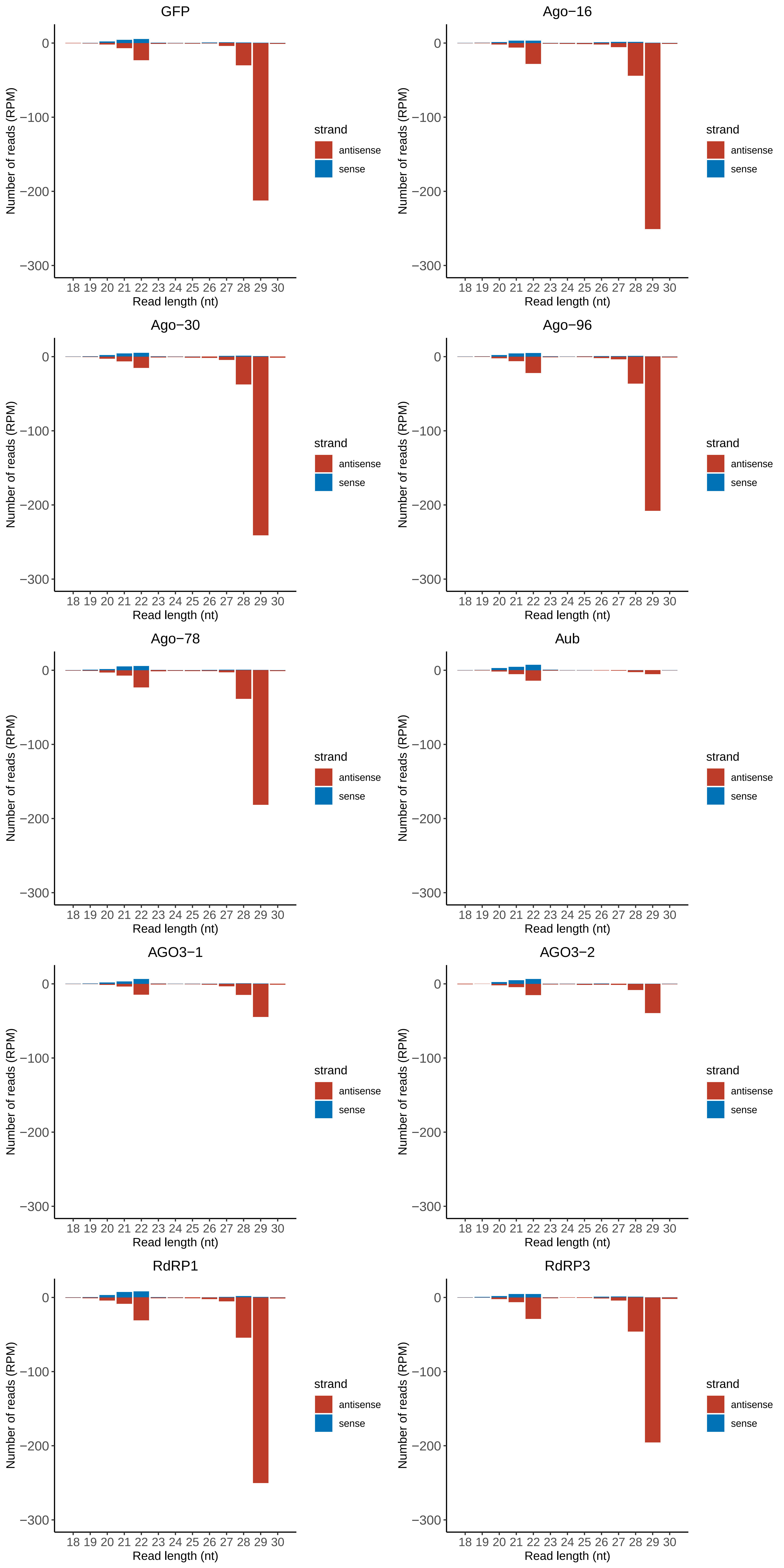

### Motif:rnd-1_family-413TE_lengthdistribution.pdf

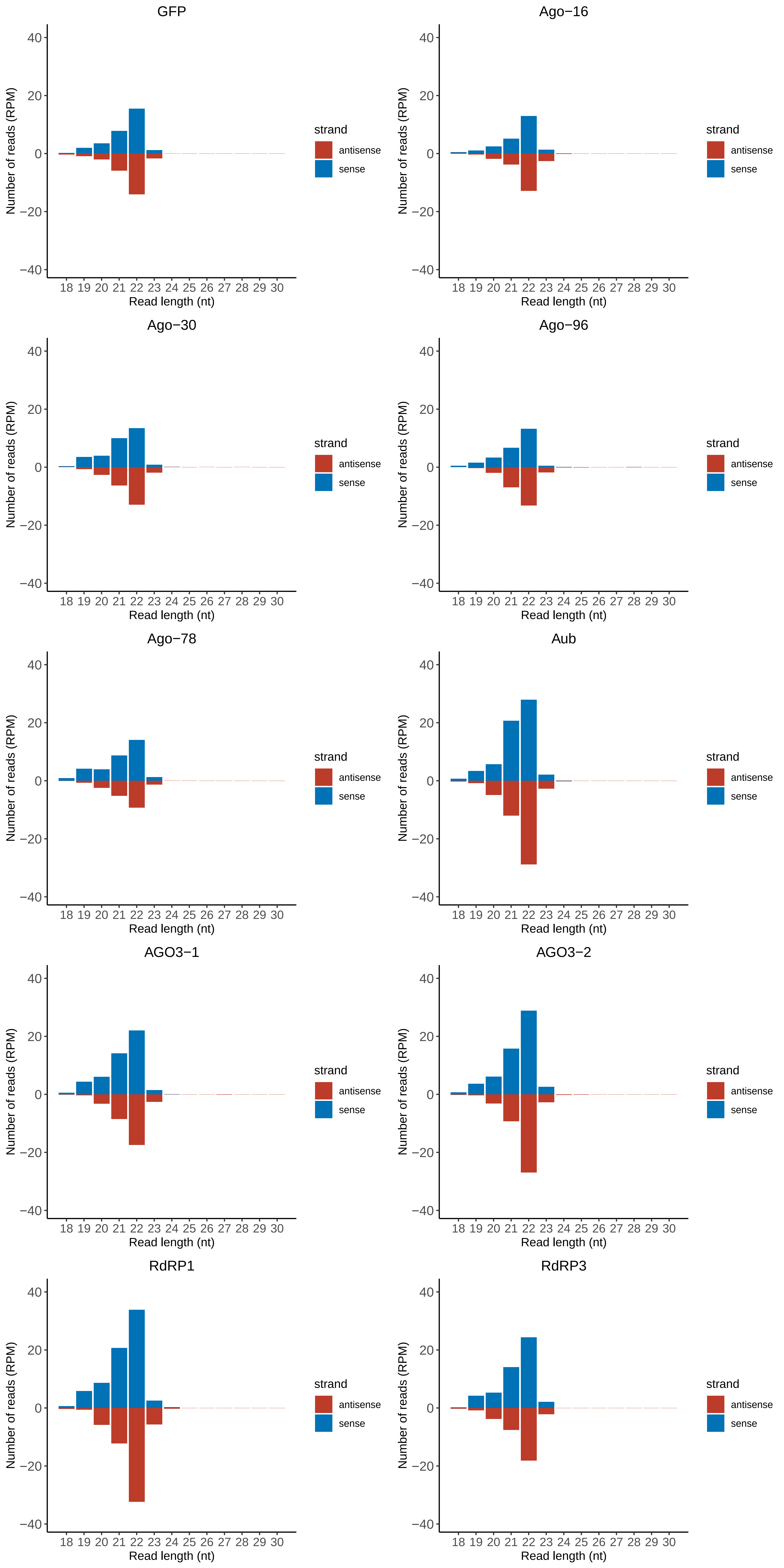

### Motif:rnd-1_family-417TE_lengthdistribution.pdf

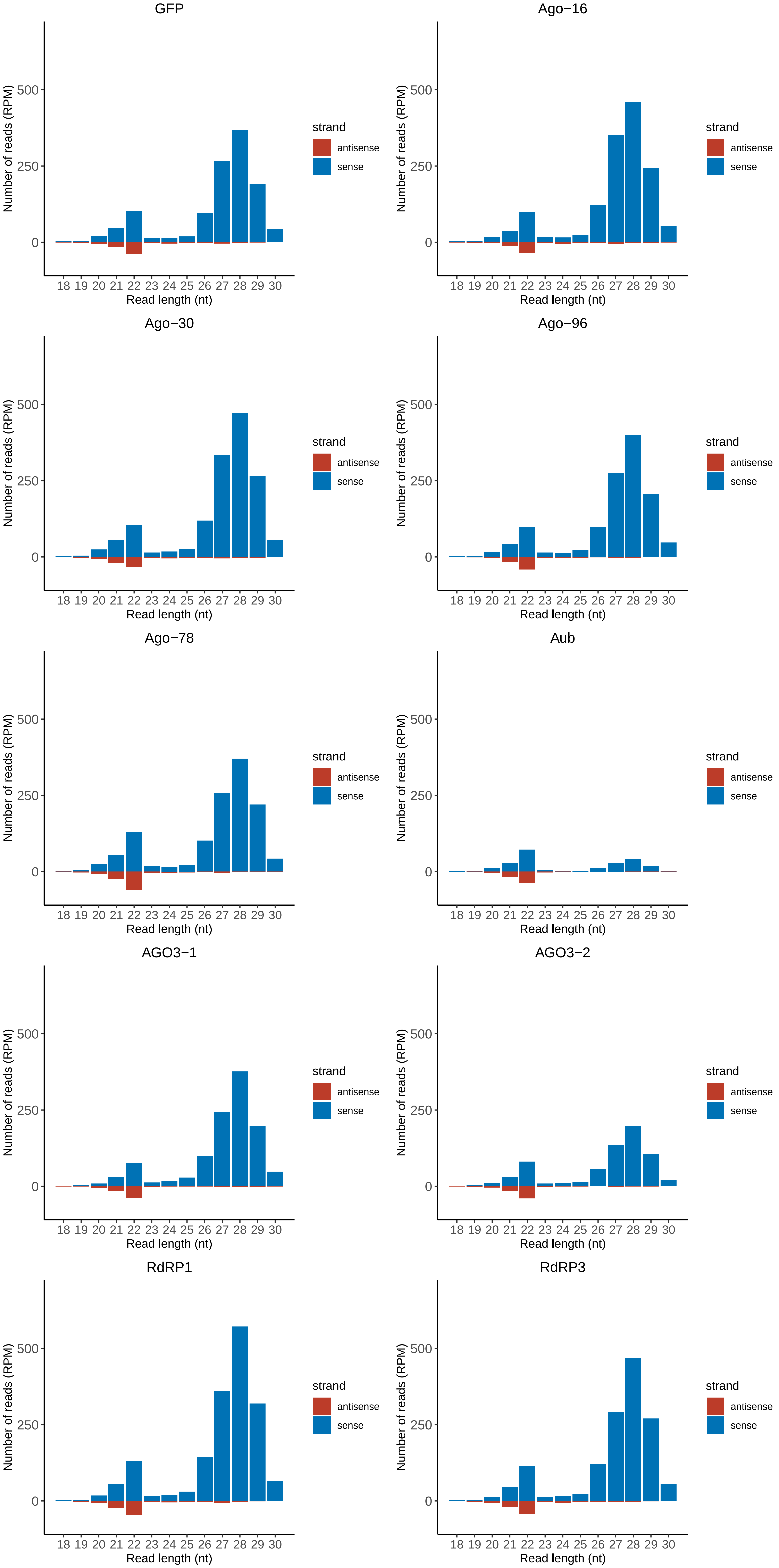

### Motif:rnd-1_family-424TE_lengthdistribution.pdf

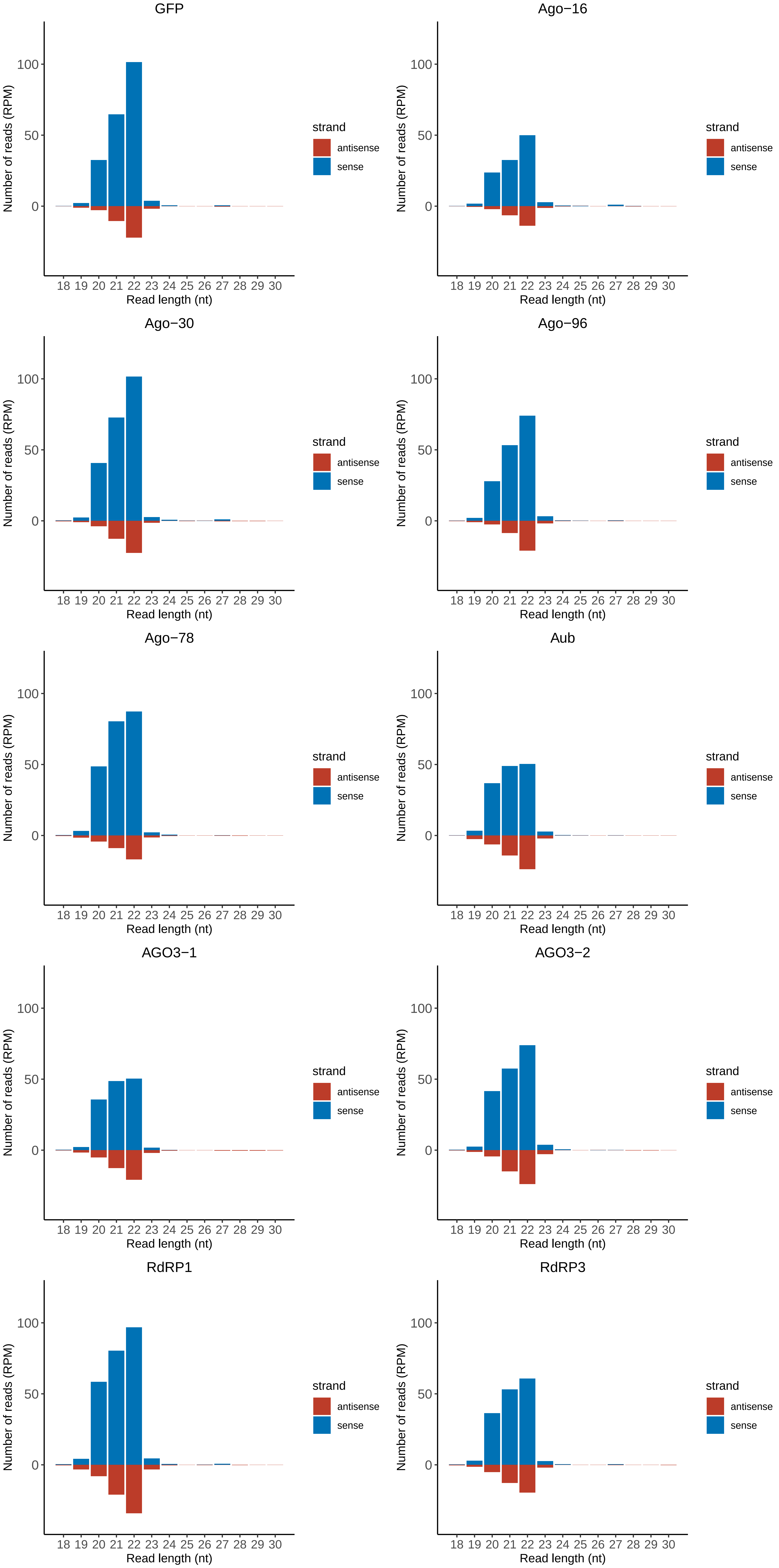

### Motif:rnd-1_family-429TE_lengthdistribution.pdf

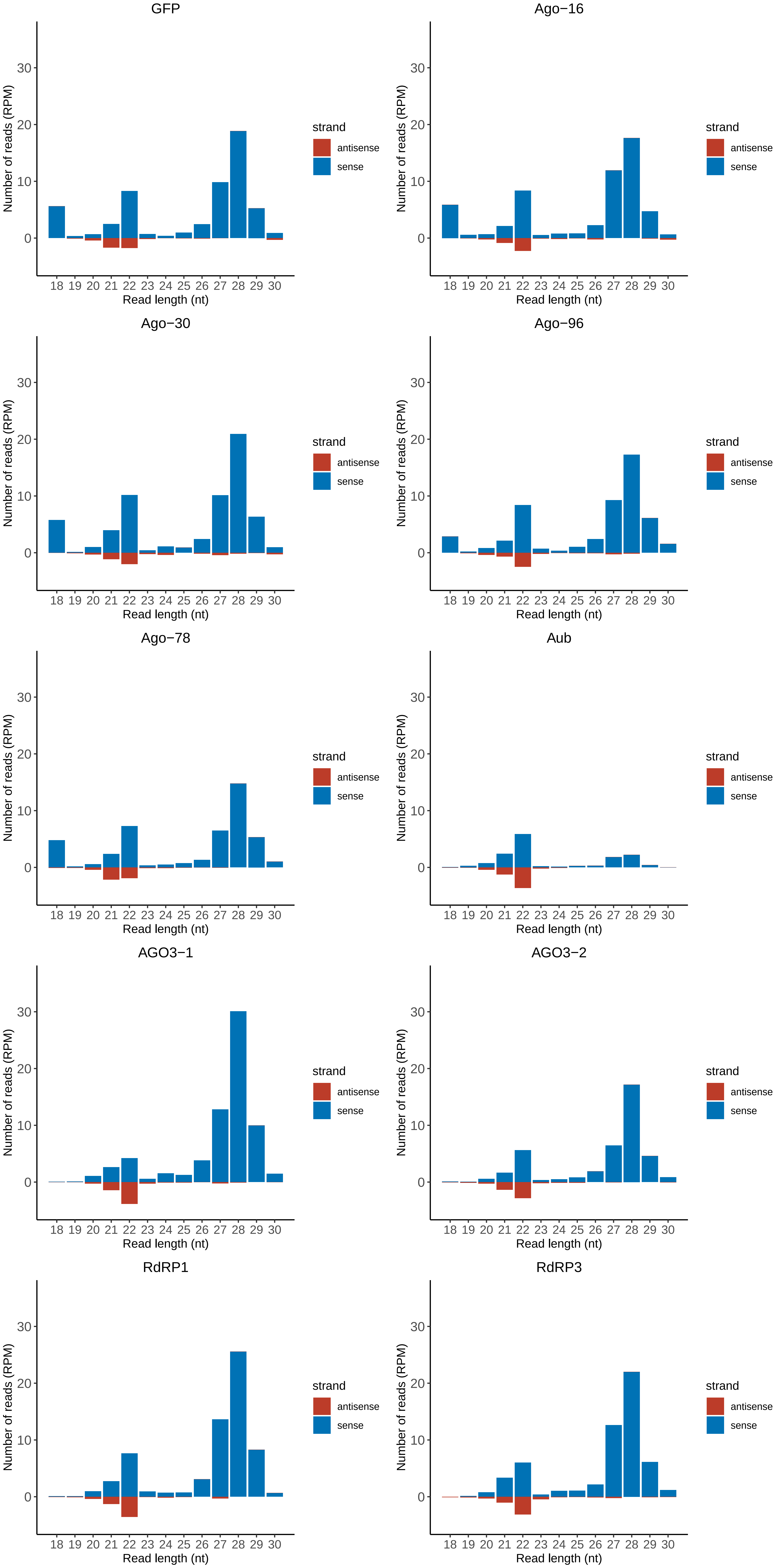

### Motif:rnd-1_family-443TE_lengthdistribution.pdf

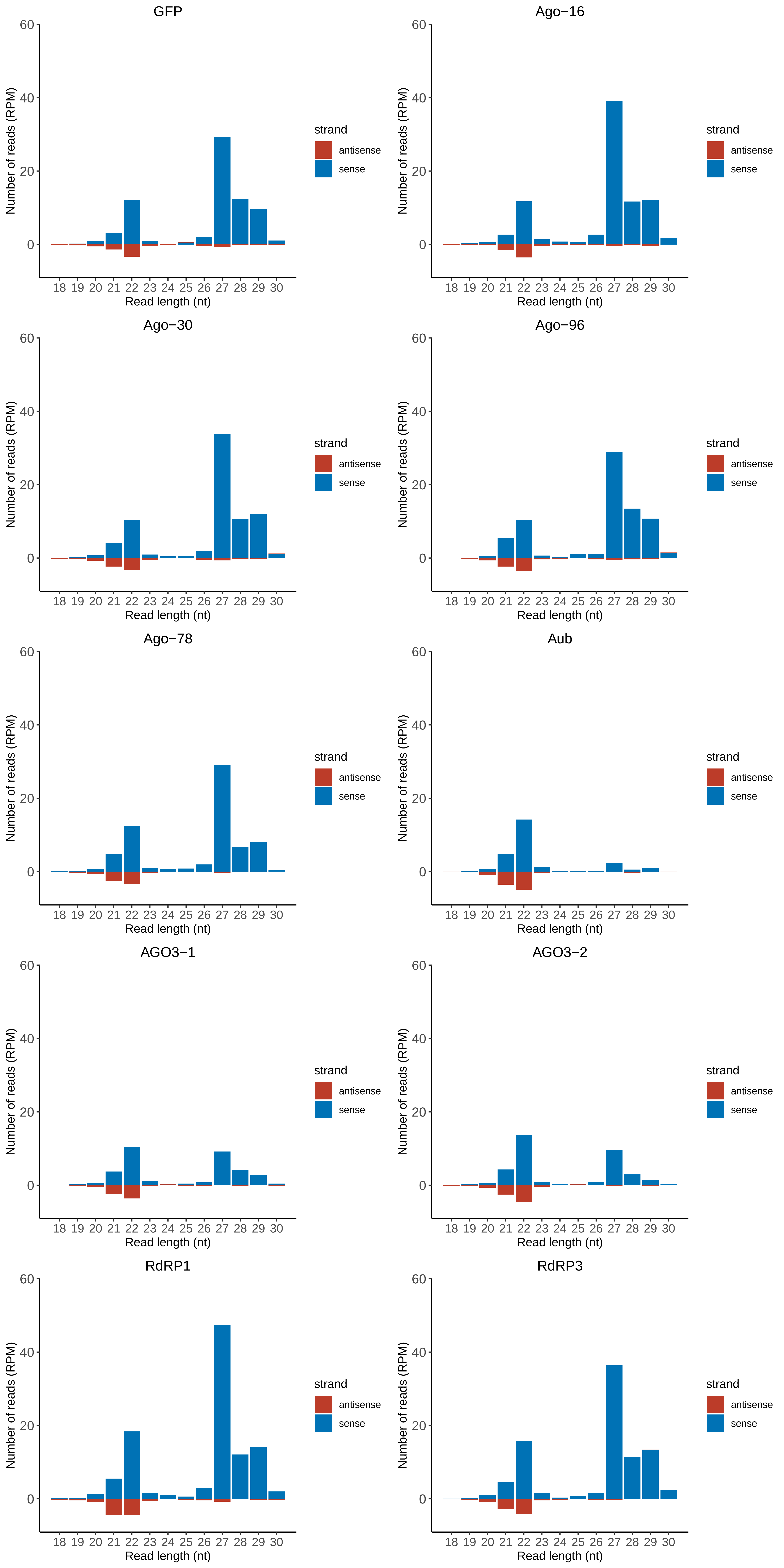

### Motif:rnd-1_family-447TE_lengthdistribution.pdf

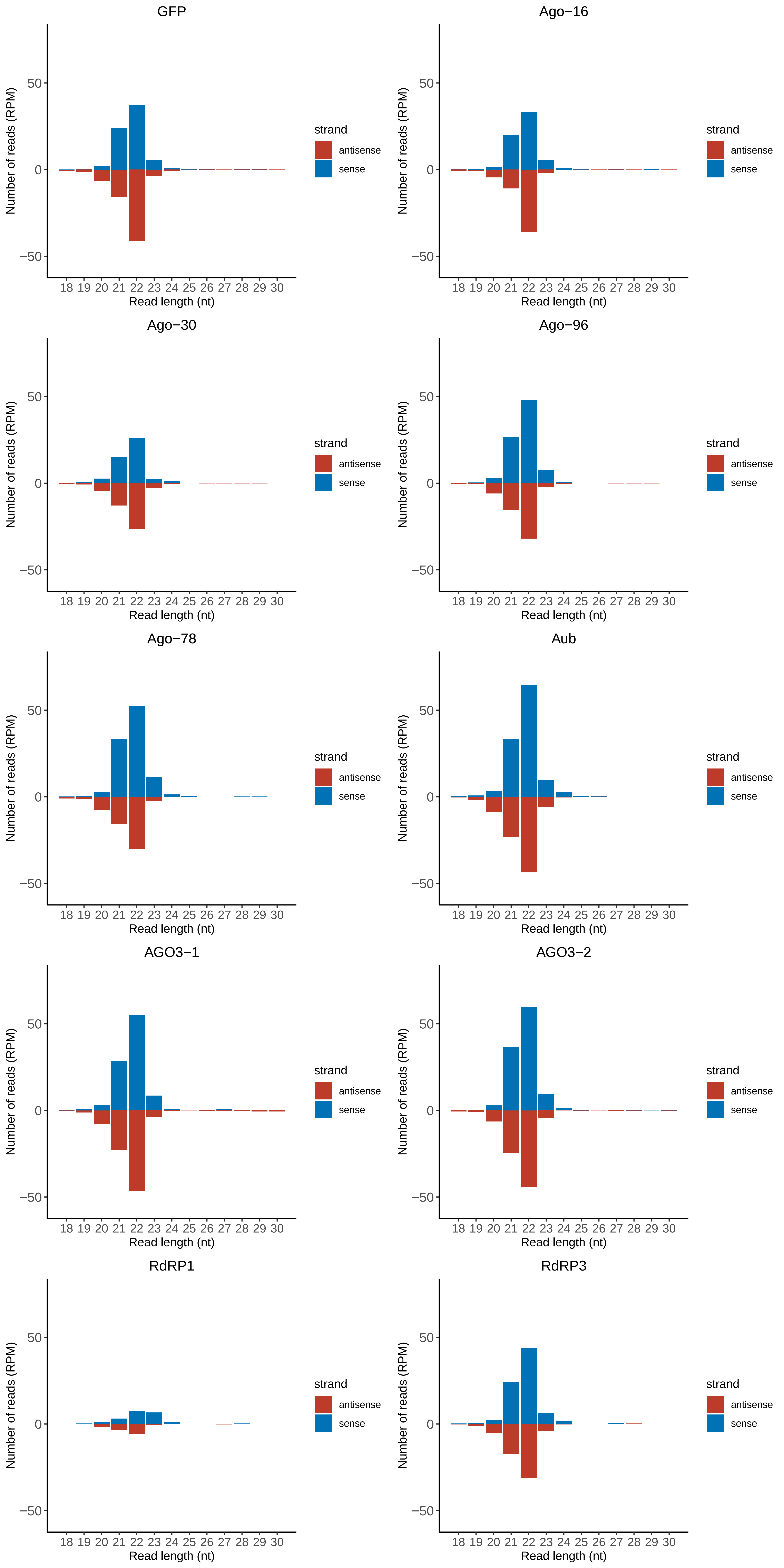

### Motif:rnd-1_family-474TE_lengthdistribution.pdf

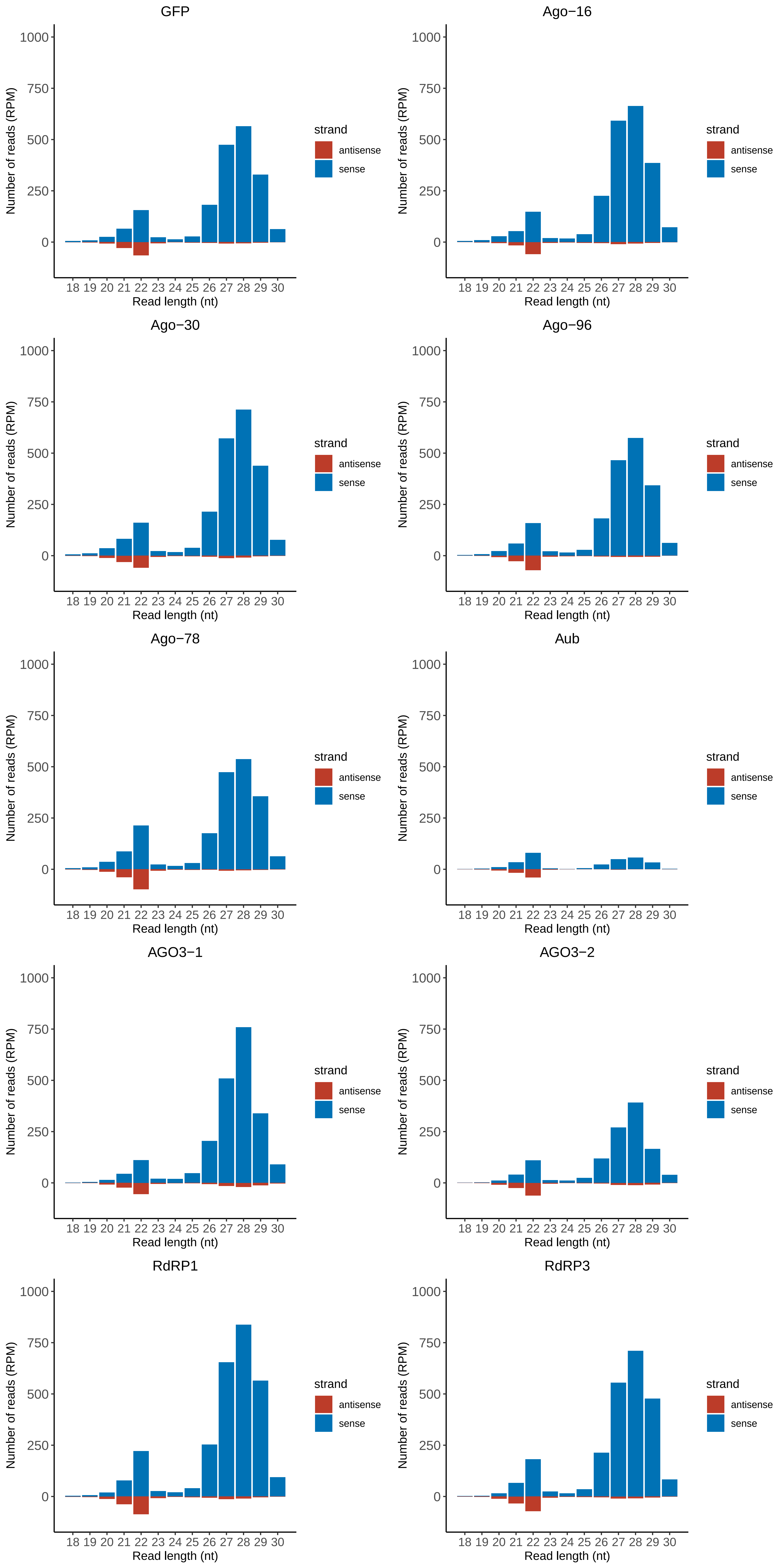

### Motif:rnd-1_family-525TE_lengthdistribution.pdf

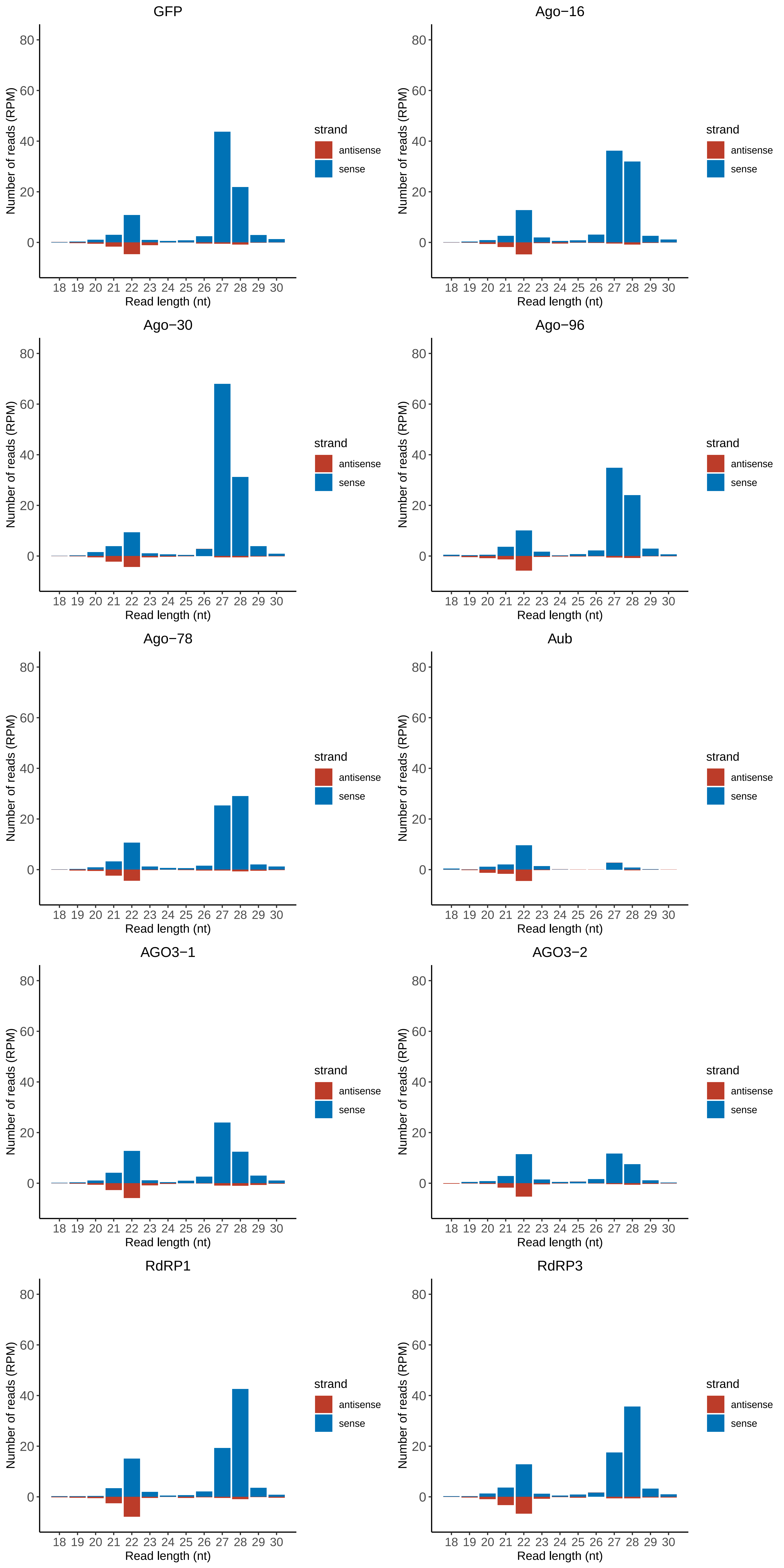

### Motif:rnd-1_family-614TE_lengthdistribution.pdf

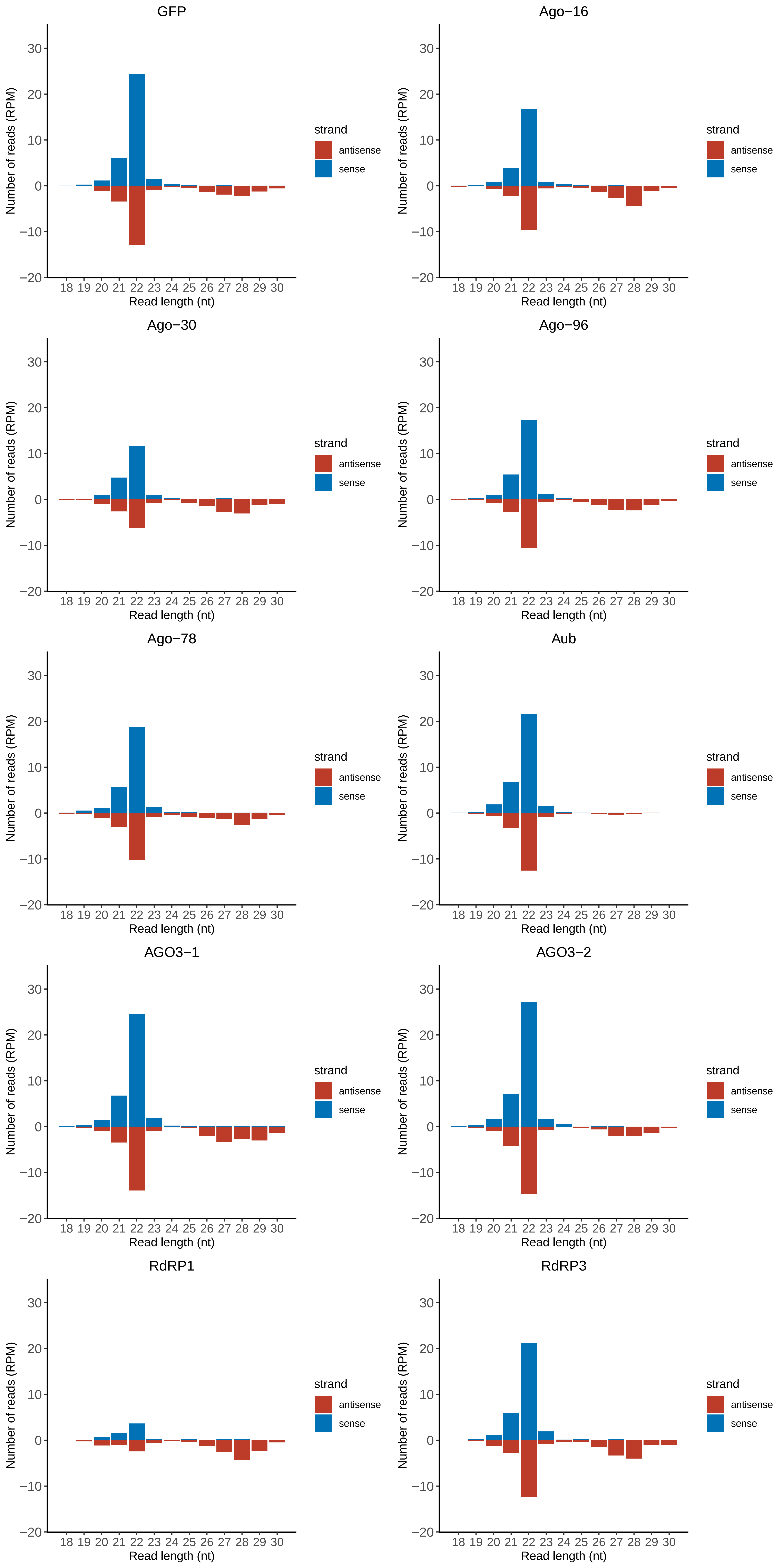

### Motif:rnd-1_family-722TE_lengthdistribution.pdf

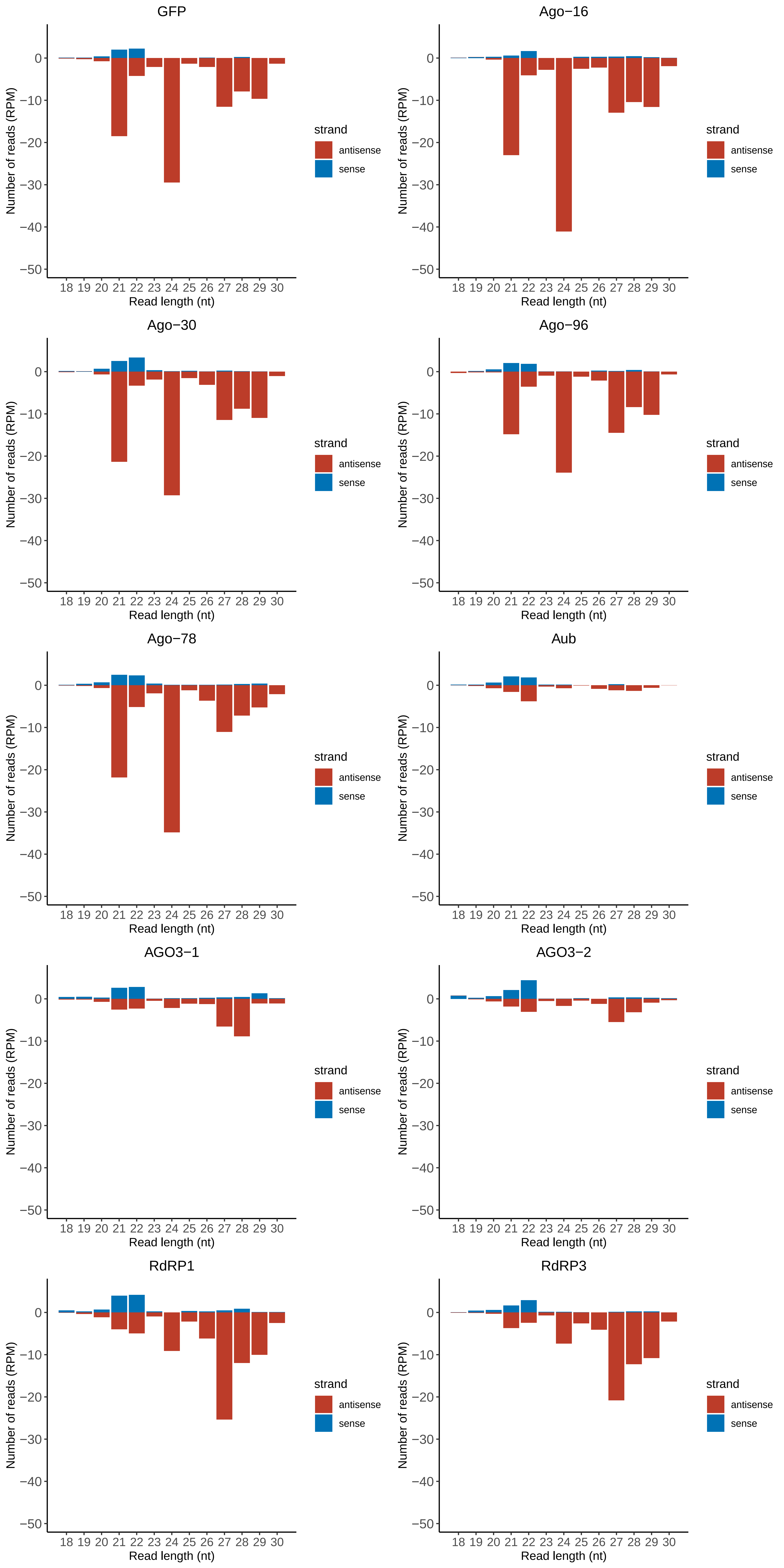

### Motif:rnd-1_family-726TE_lengthdistribution.pdf

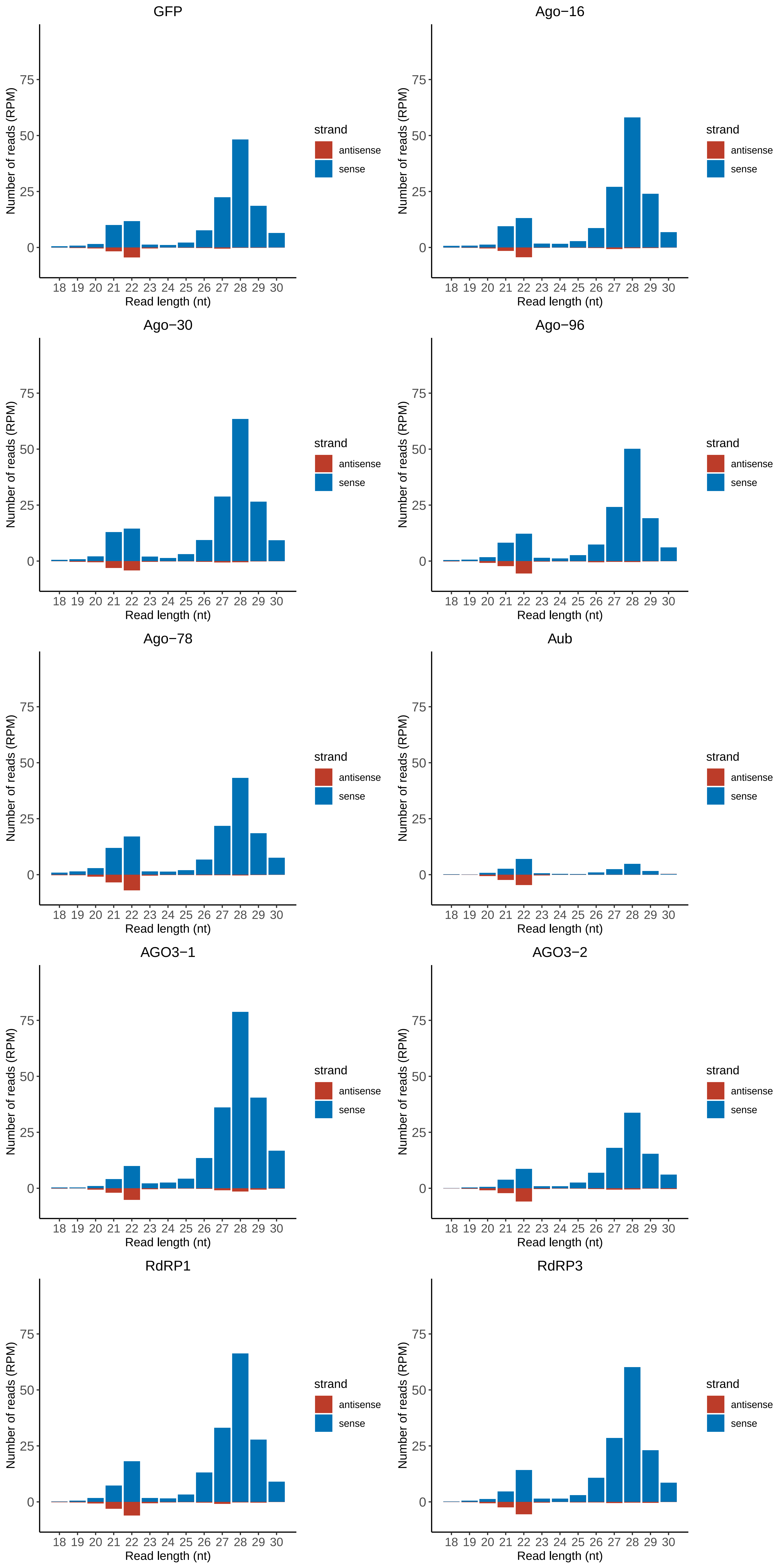

### Motif:rnd-1_family-772TE_lengthdistribution.pdf

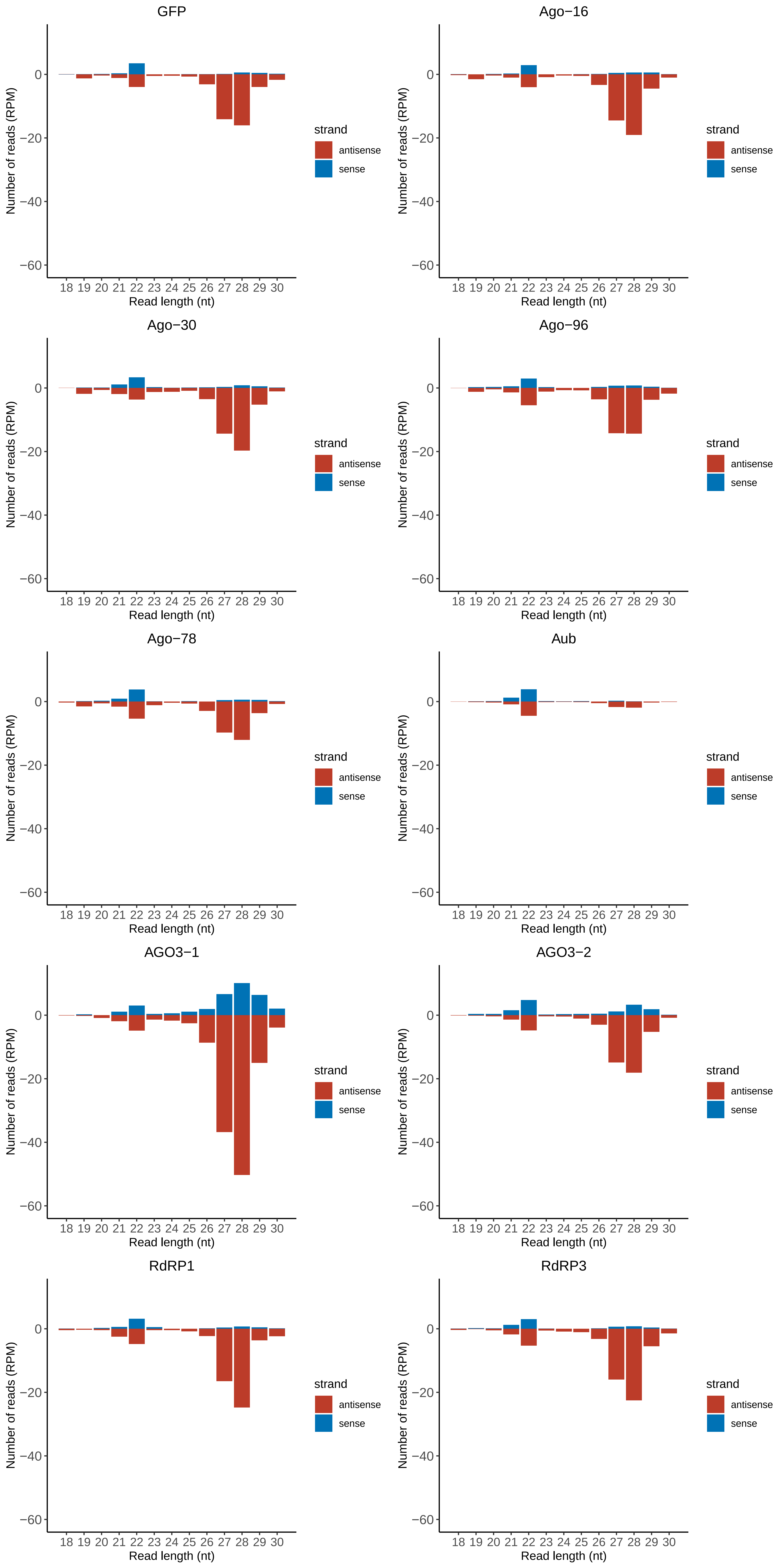

### Motif:rnd-1_family-795TE_lengthdistribution.pdf

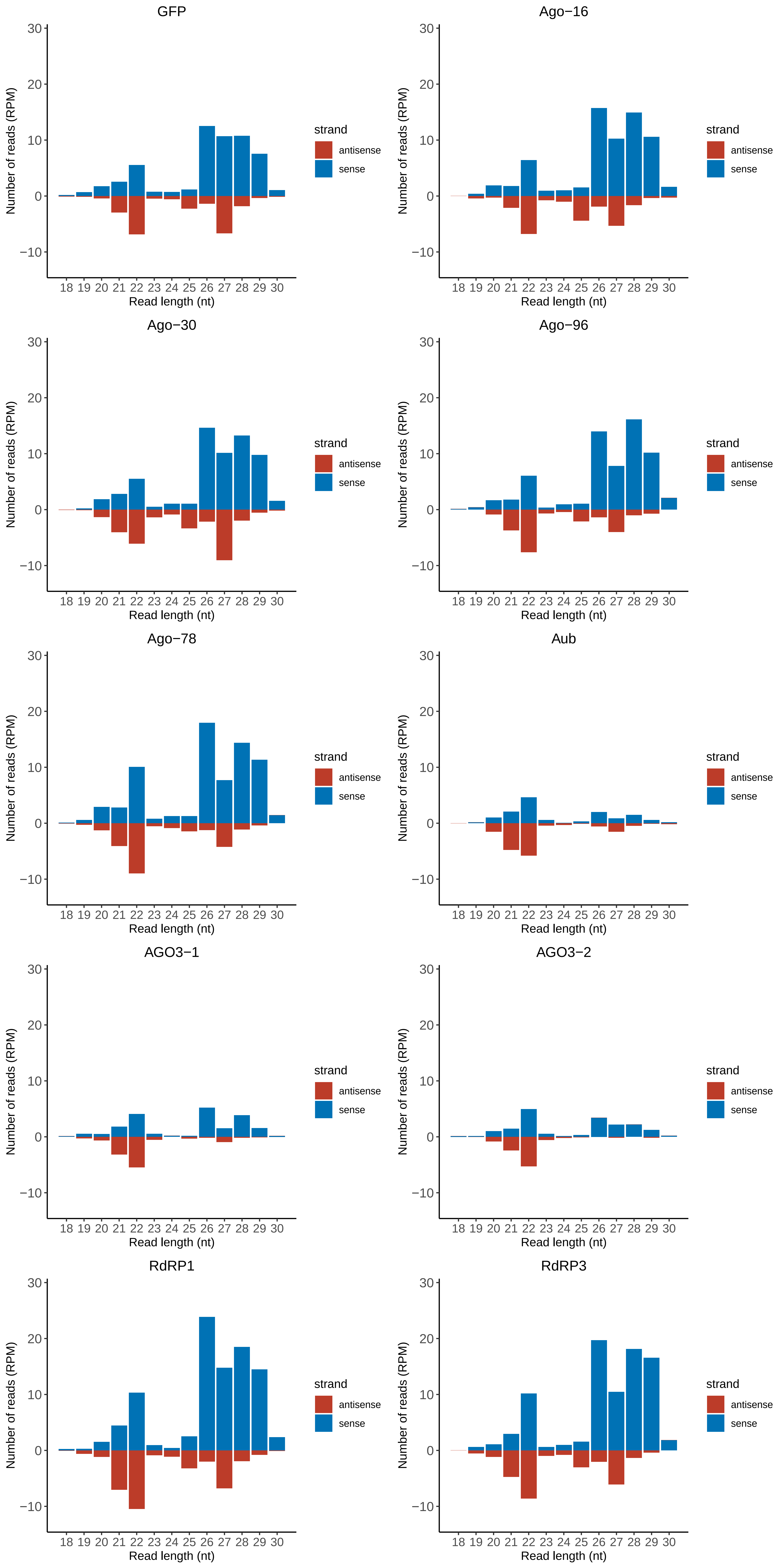

### Motif:rnd-1_family-1037TE_lengthdistribution.pdf

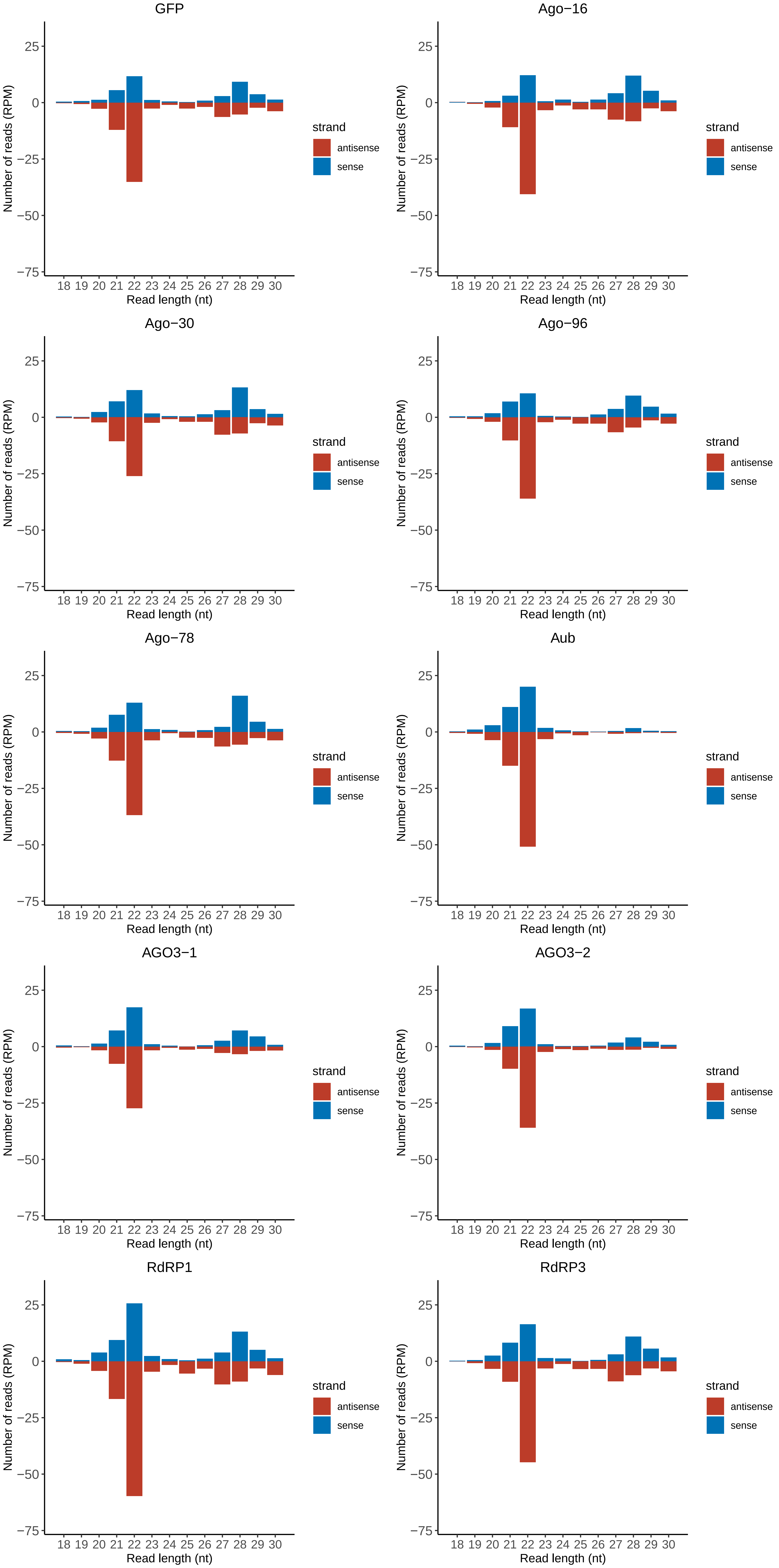

### Motif:rnd-1_family-1059TE_lengthdistribution.pdf

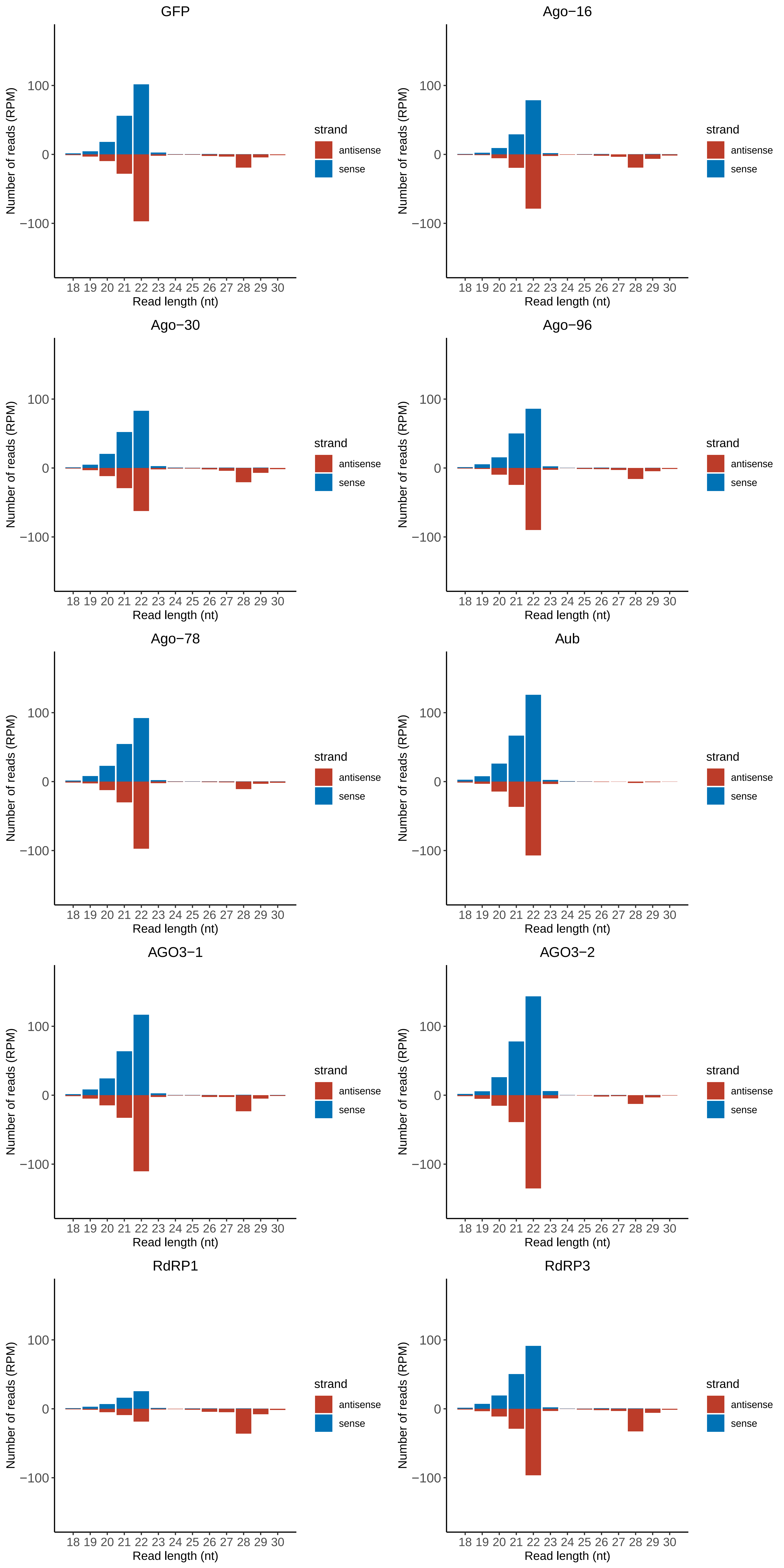

### Motif:rnd-1_family-1154TE_lengthdistribution.pdf

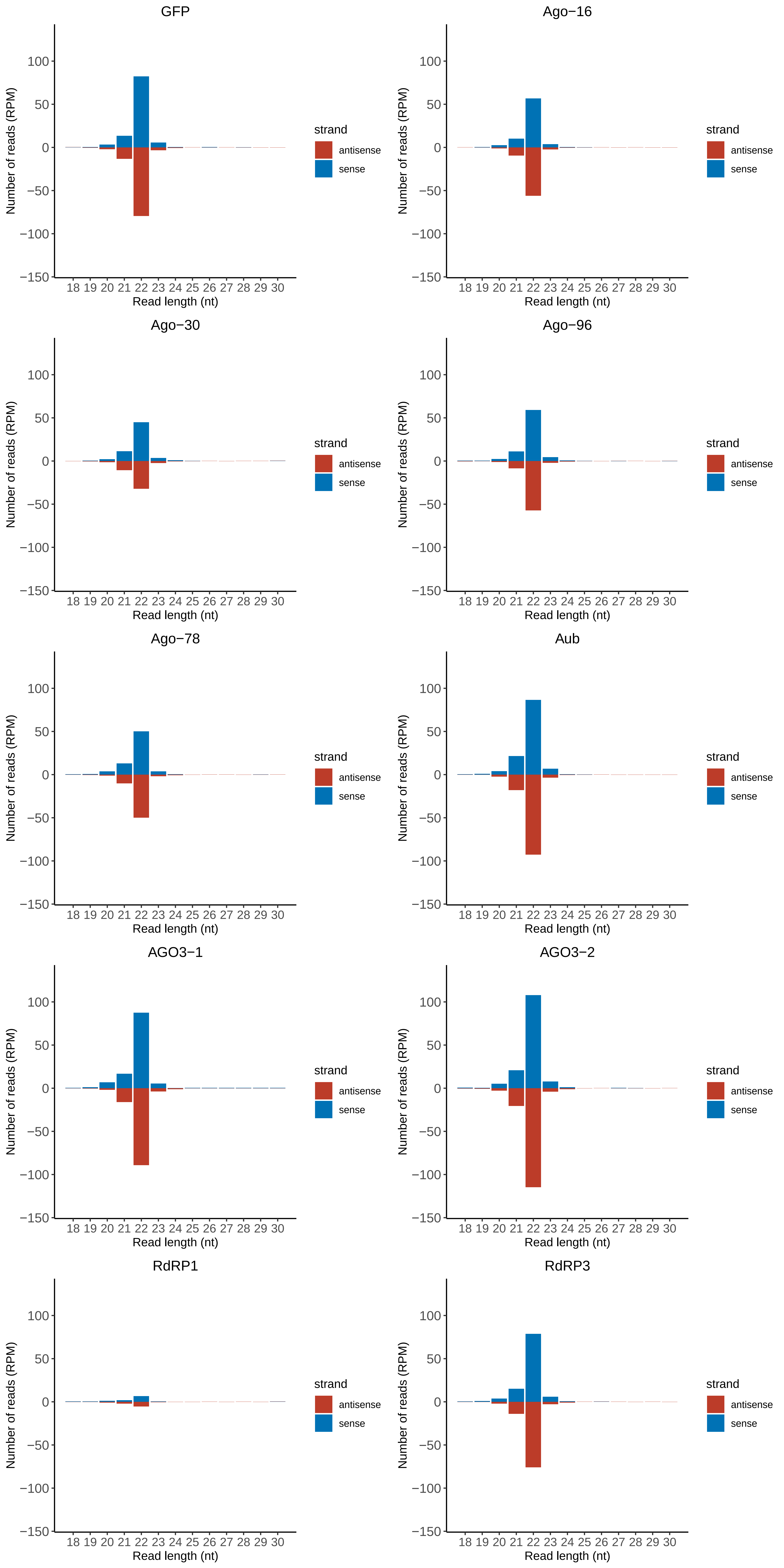

### Motif:rnd-1_family-1155TE_lengthdistribution.pdf

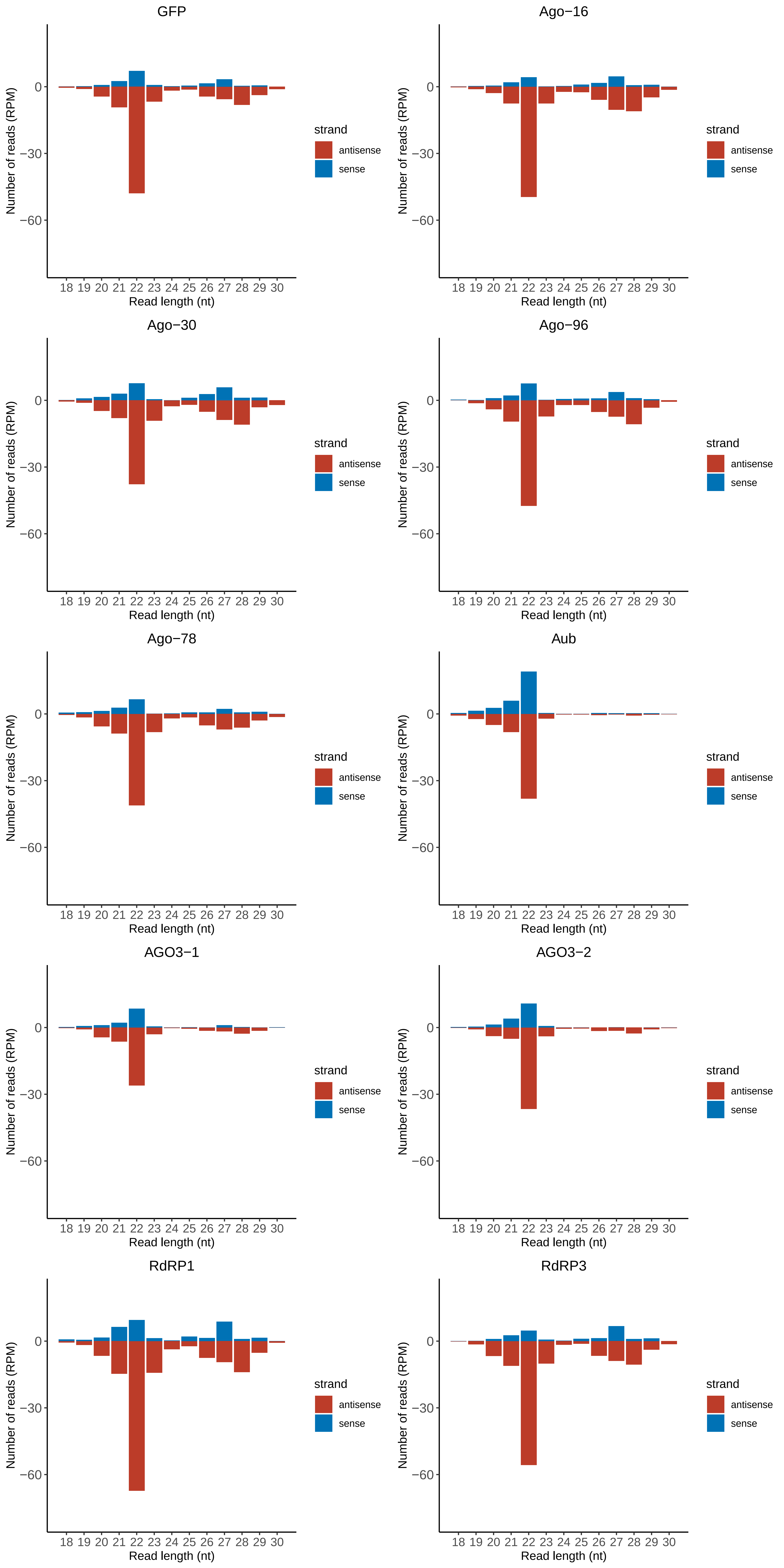

### Motif:rnd-1_family-1207TE_lengthdistribution.pdf

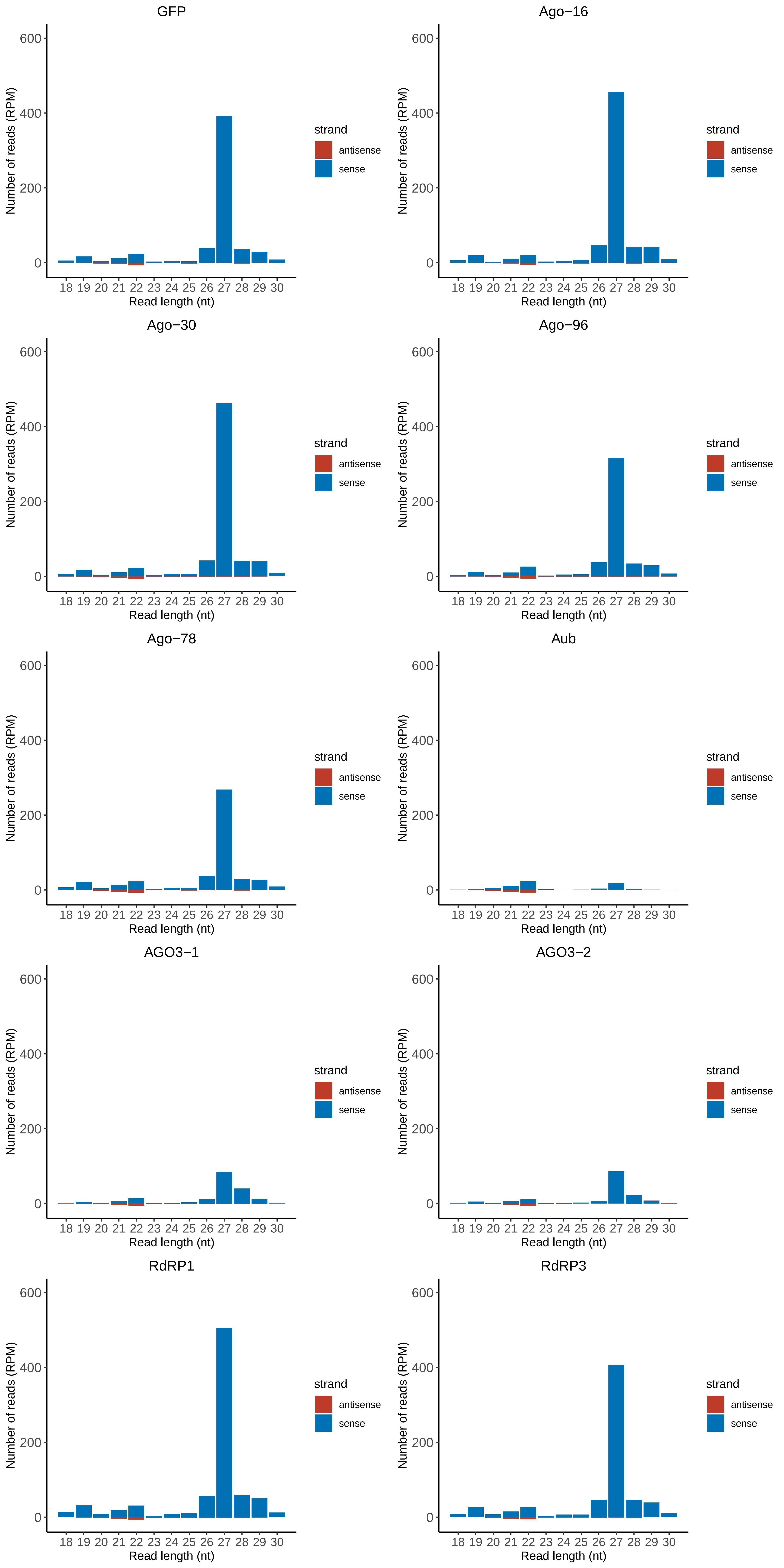

### Motif:rnd-1_family-1523TE_lengthdistribution.pdf

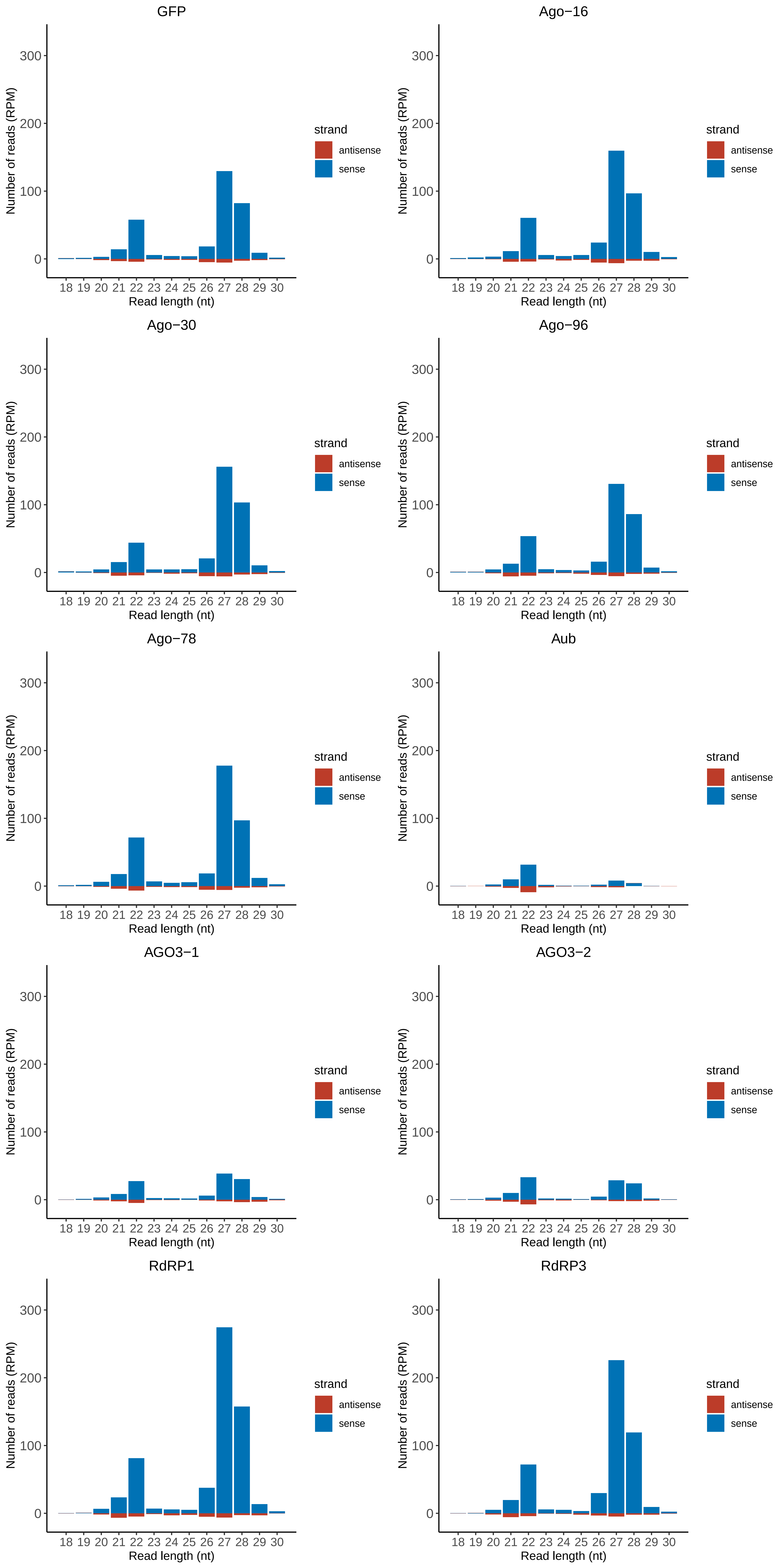

### Motif:rnd-1_family-1549TE_lengthdistribution.pdf

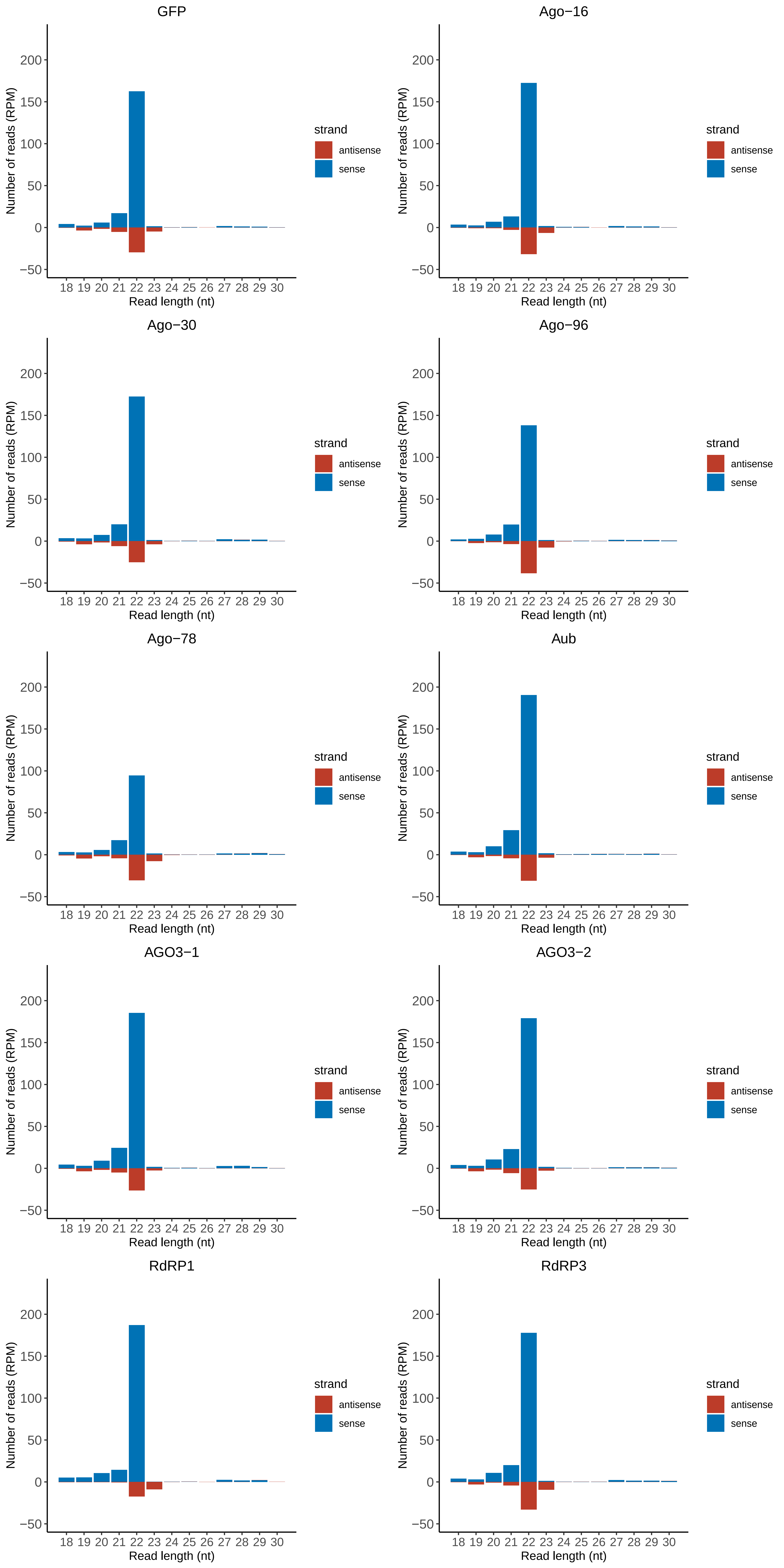

### Motif:rnd-1_family-1577TE_lengthdistribution.pdf

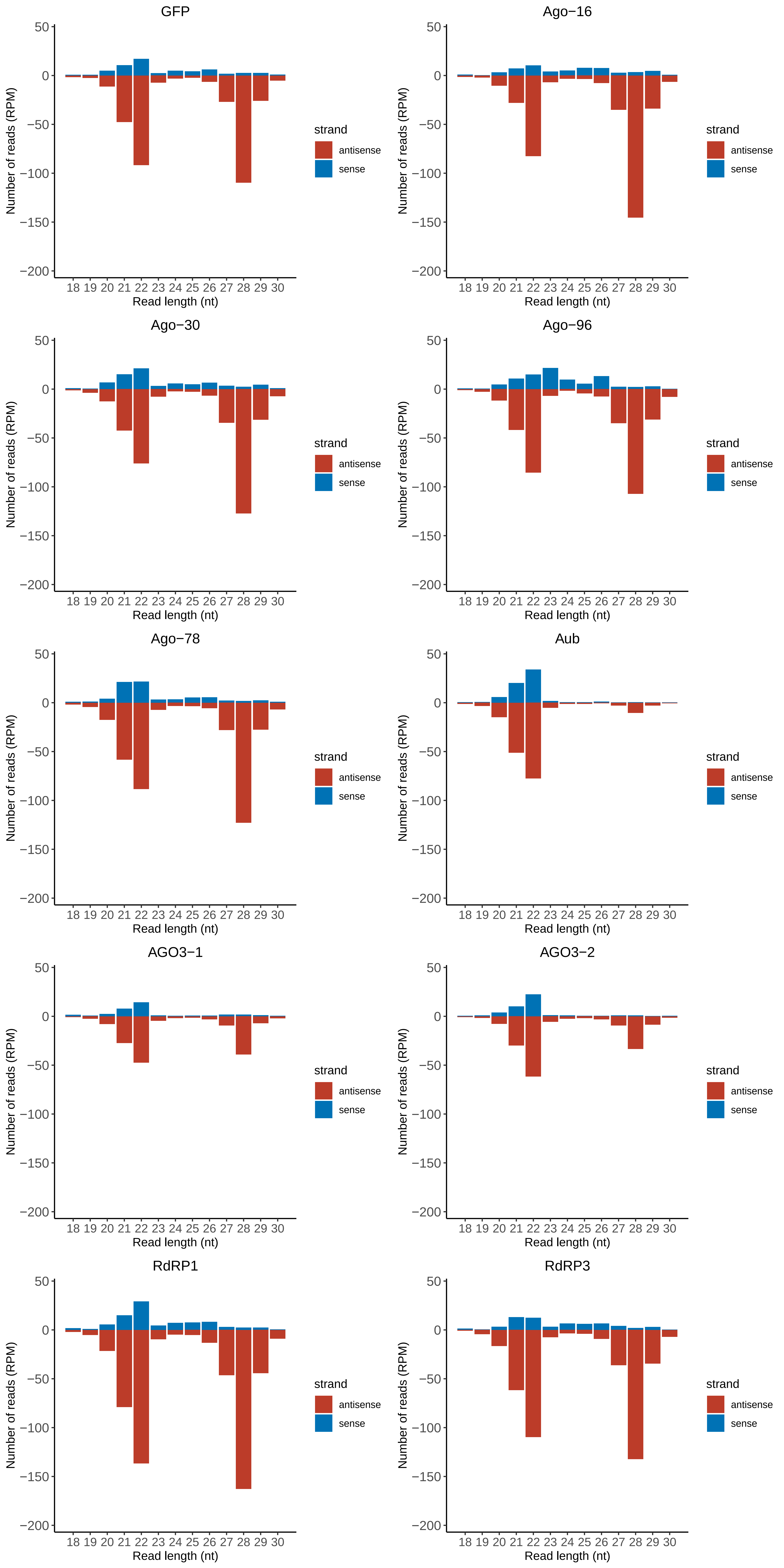

### Motif:rnd-1_family-1612TE_lengthdistribution.pdf

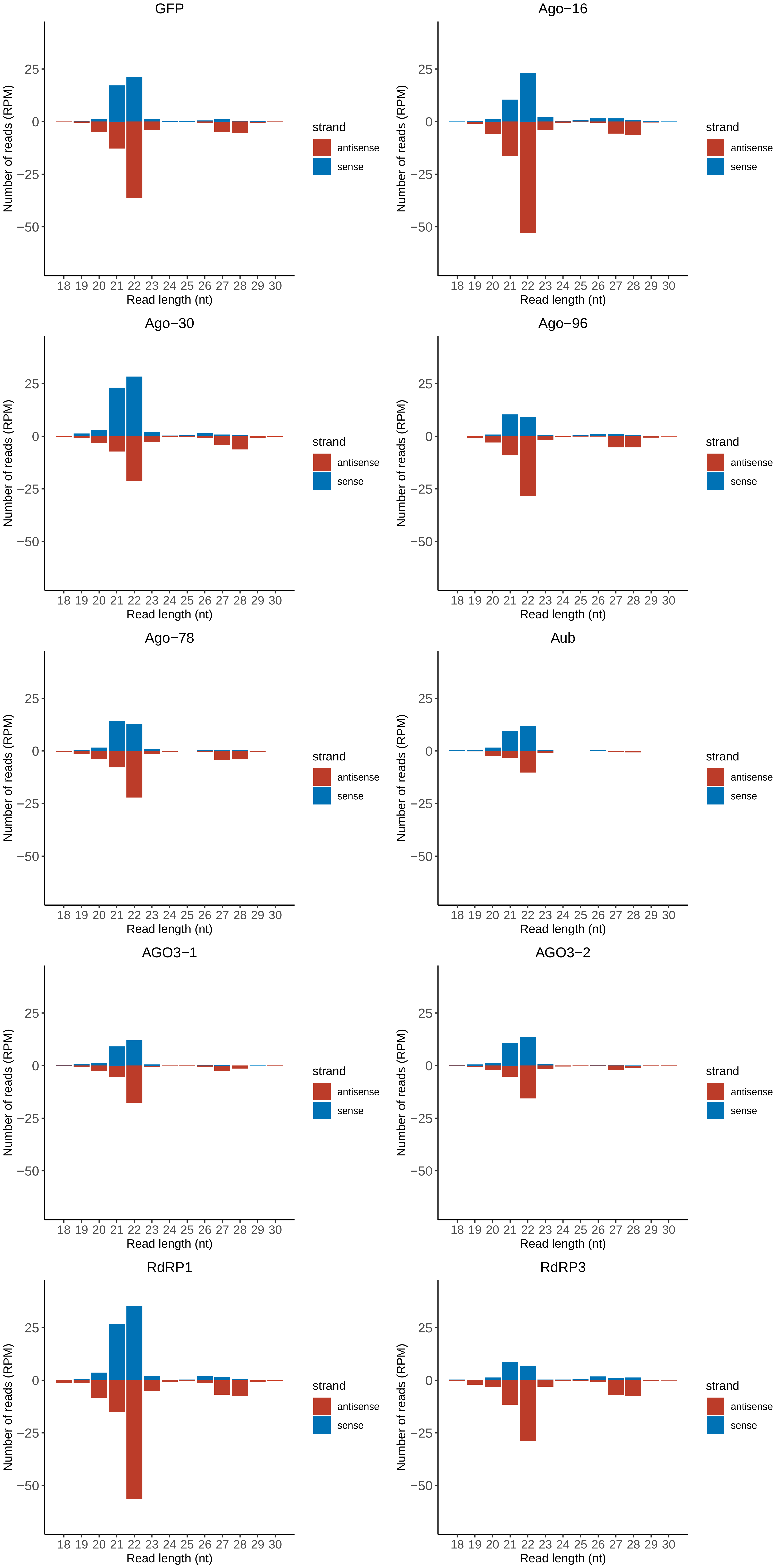

### Motif:rnd-1_family-1696TE_lengthdistribution.pdf

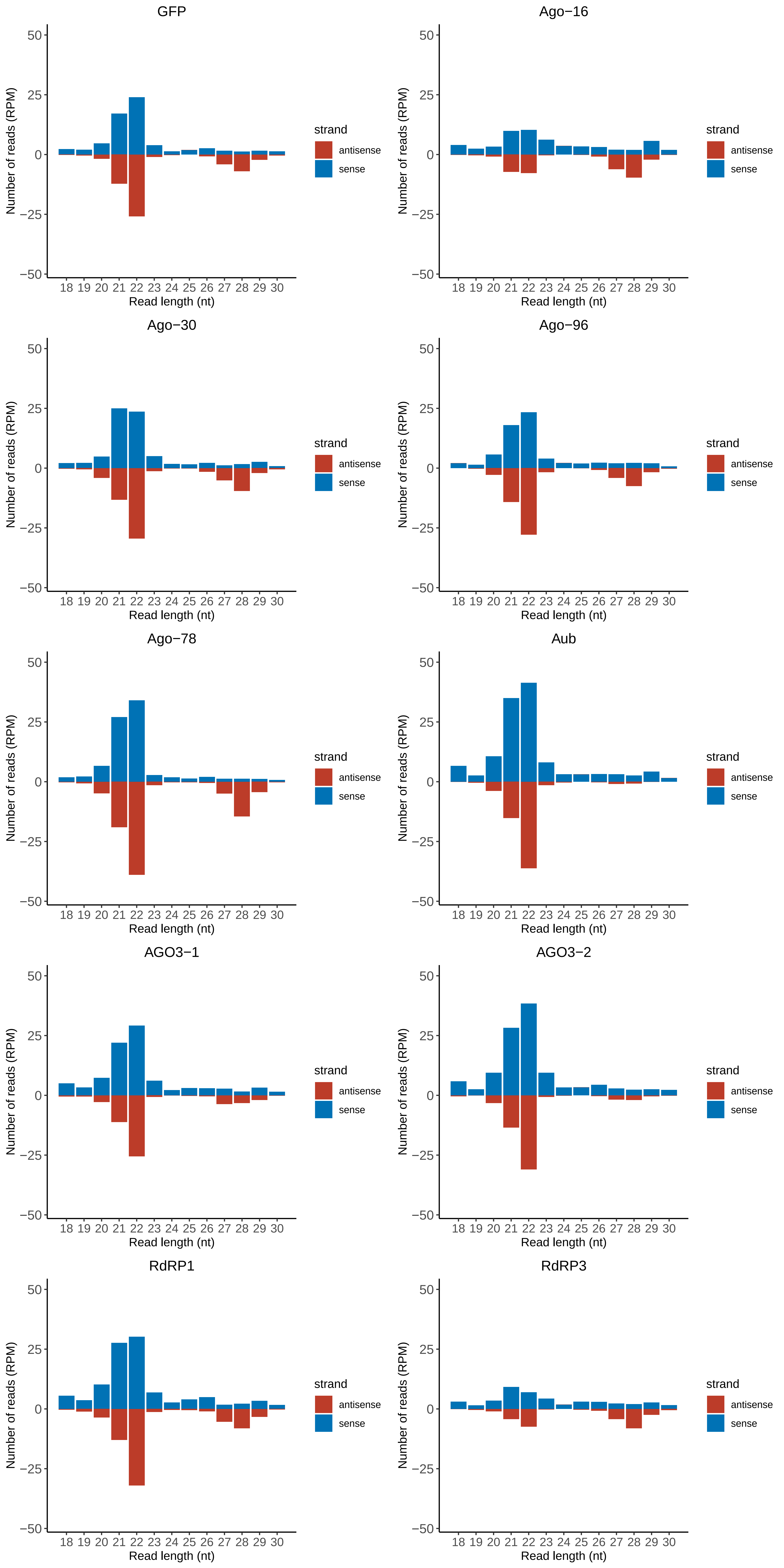

### Motif:rnd-1_family-1822TE_lengthdistribution.pdf

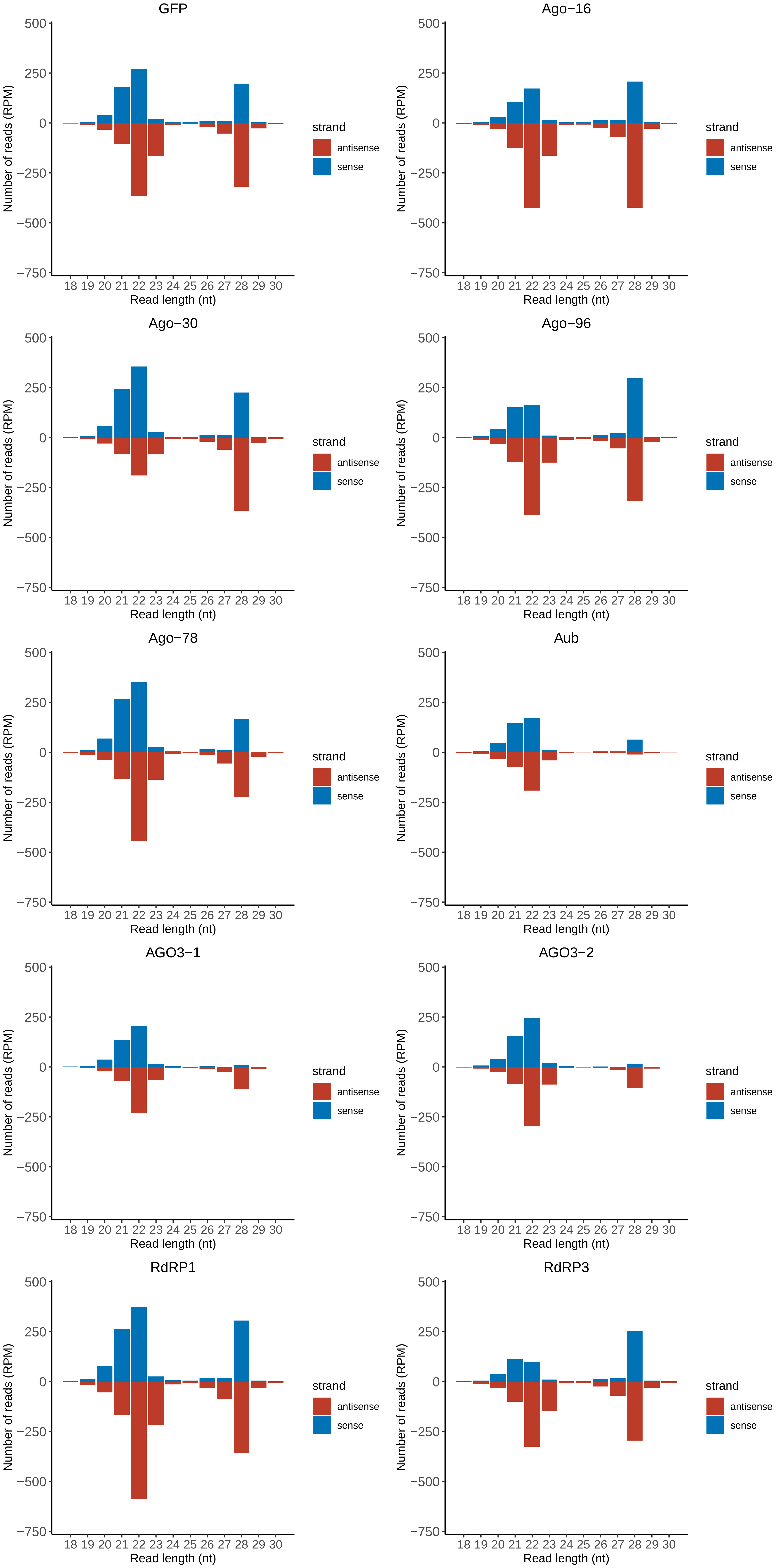

### Motif:rnd-4_family-44TE_lengthdistribution.pdf

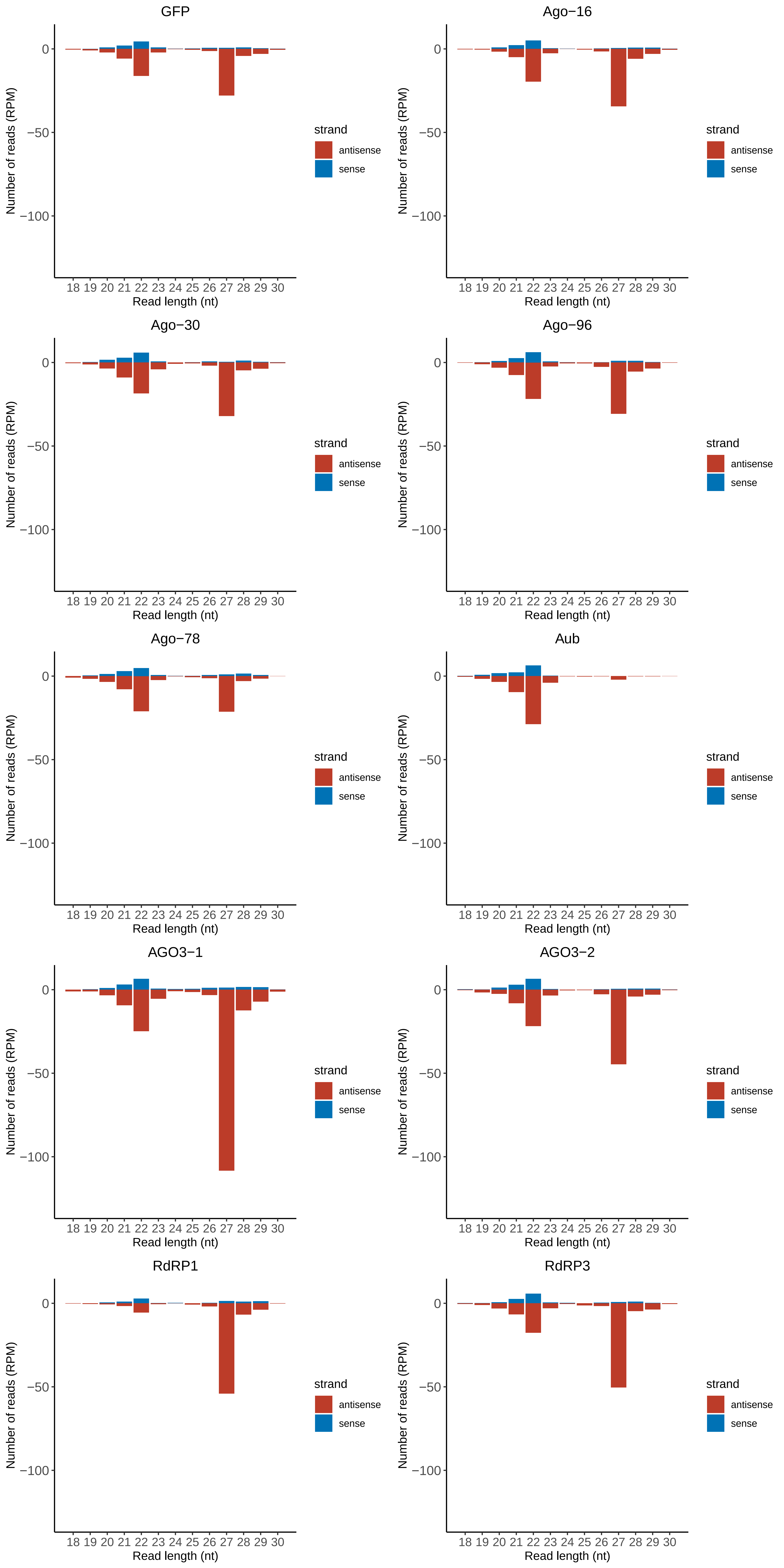

### Motif:rnd-4_family-200TE_lengthdistribution.pdf

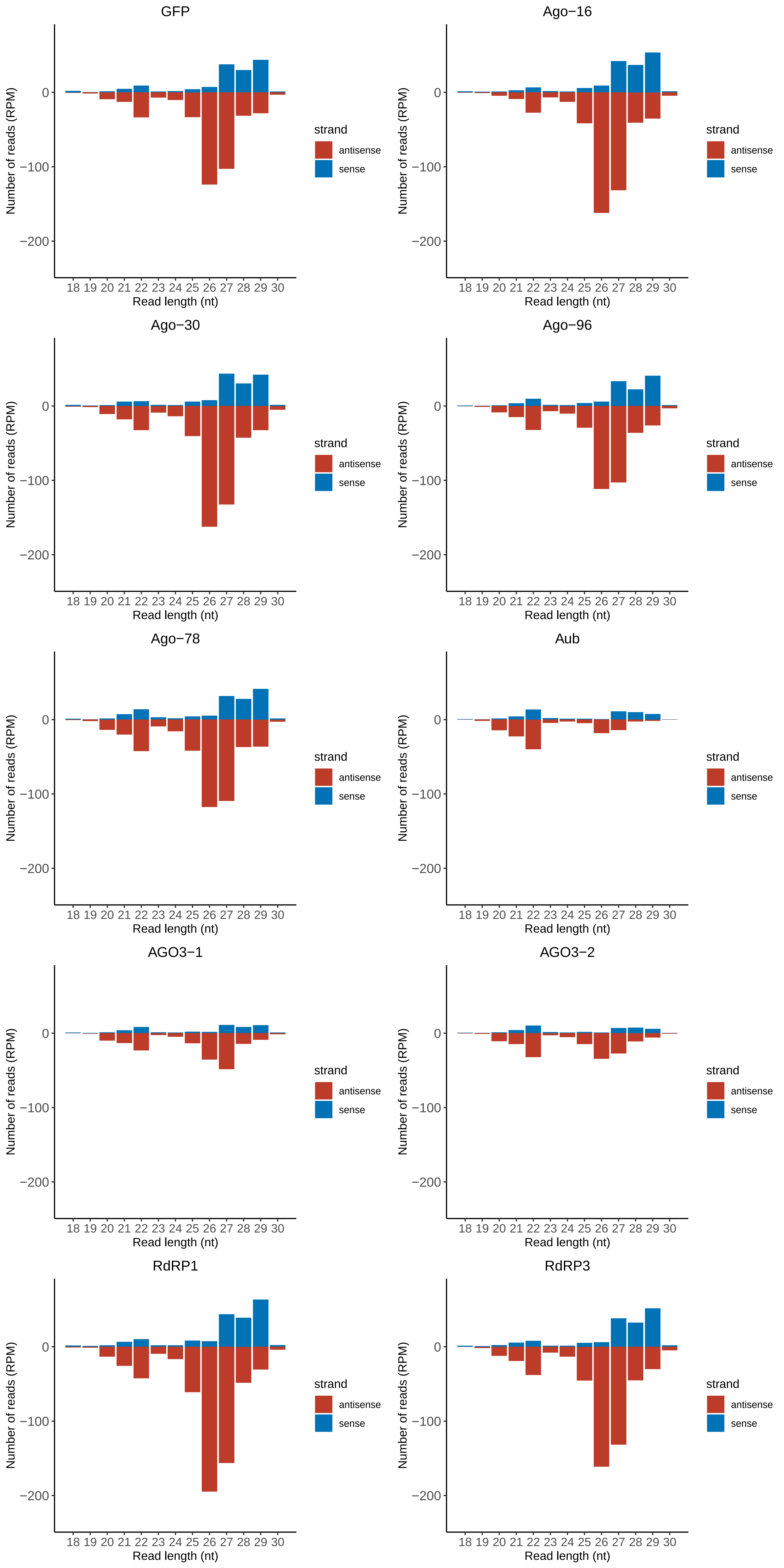

### Motif:rnd-4_family-236TE_lengthdistribution.pdf

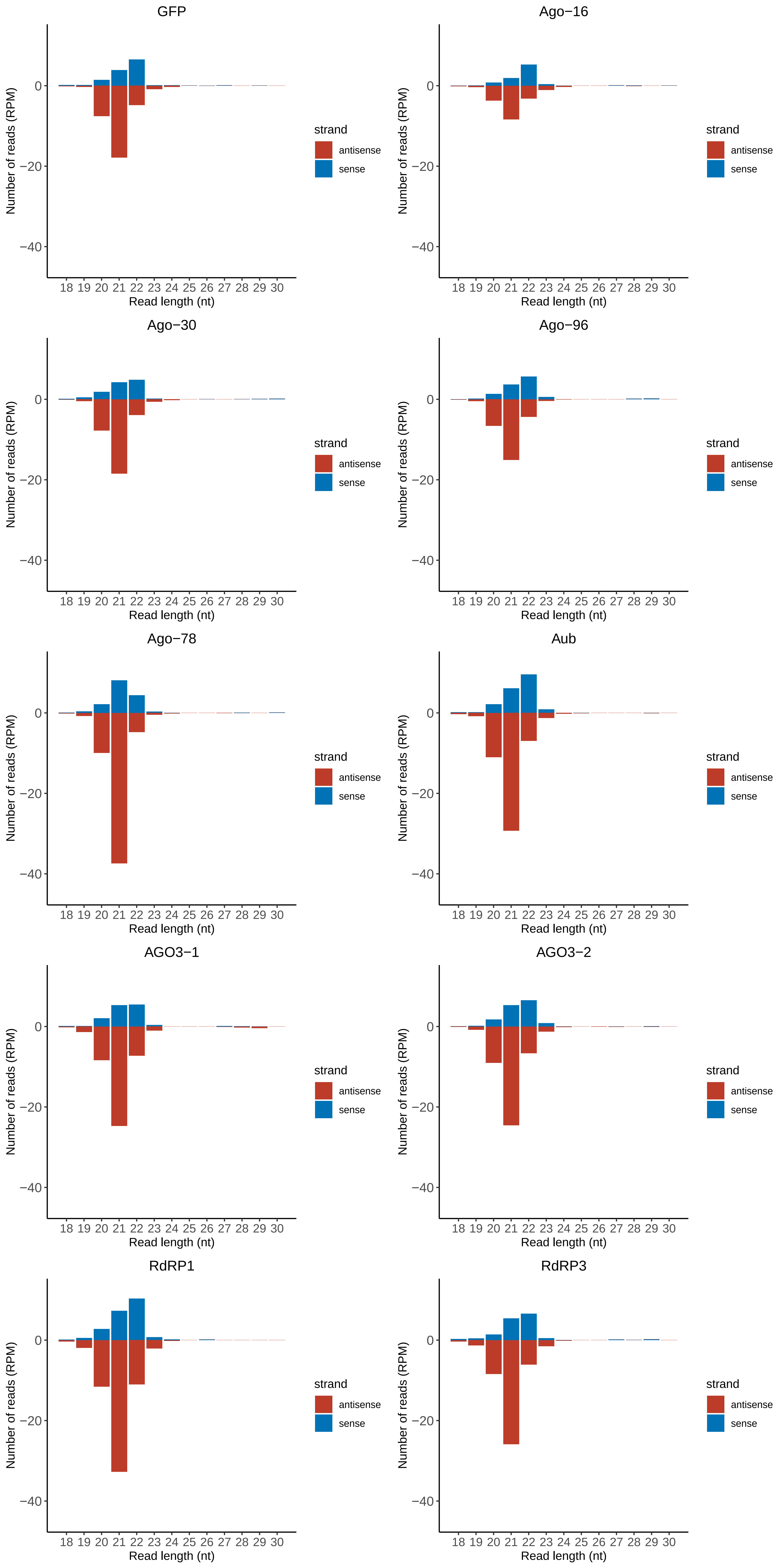

### Motif:rnd-4_family-513TE_lengthdistribution.pdf

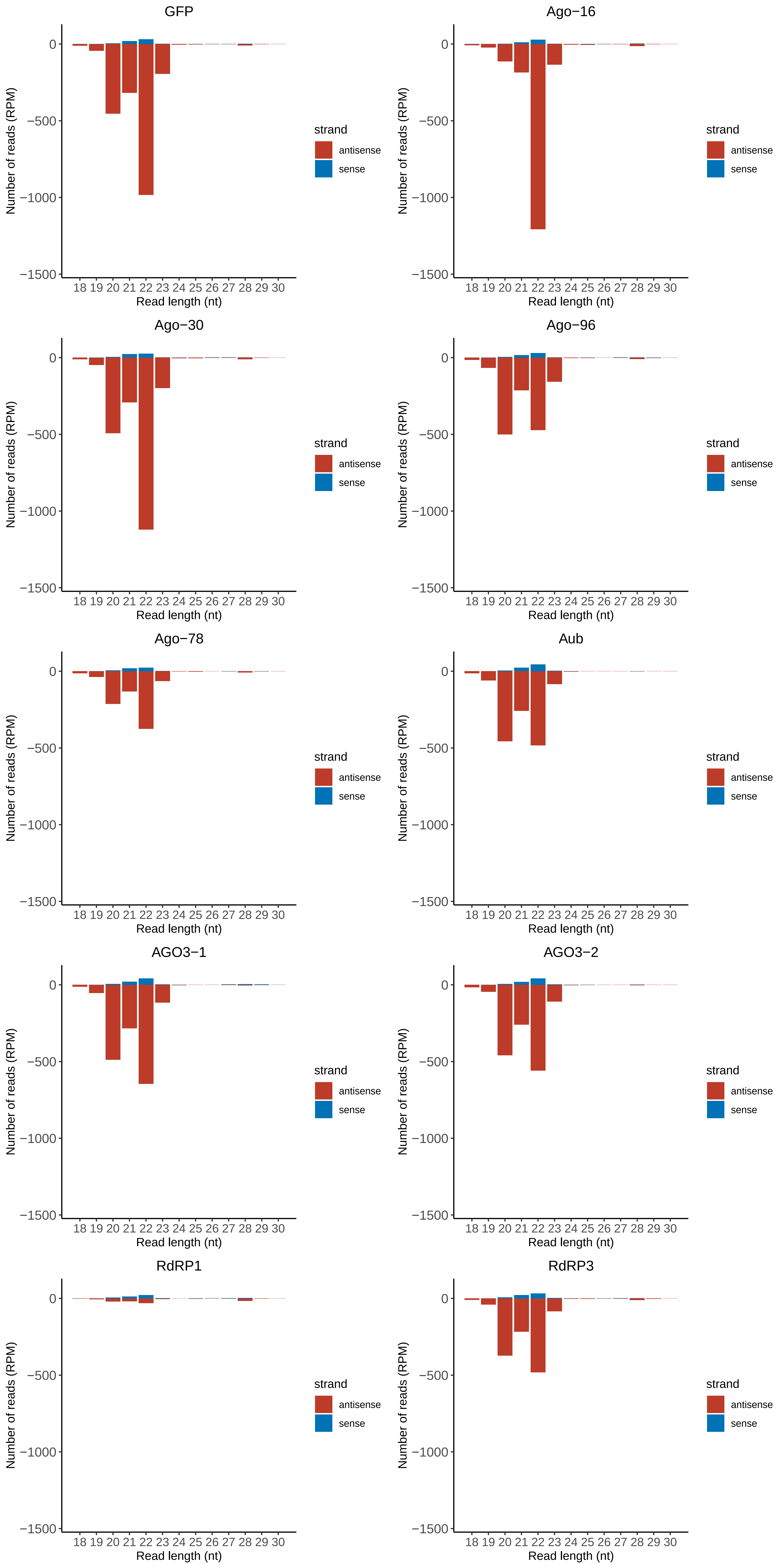

### Motif:rnd-4_family-2163TE_lengthdistribution.pdf

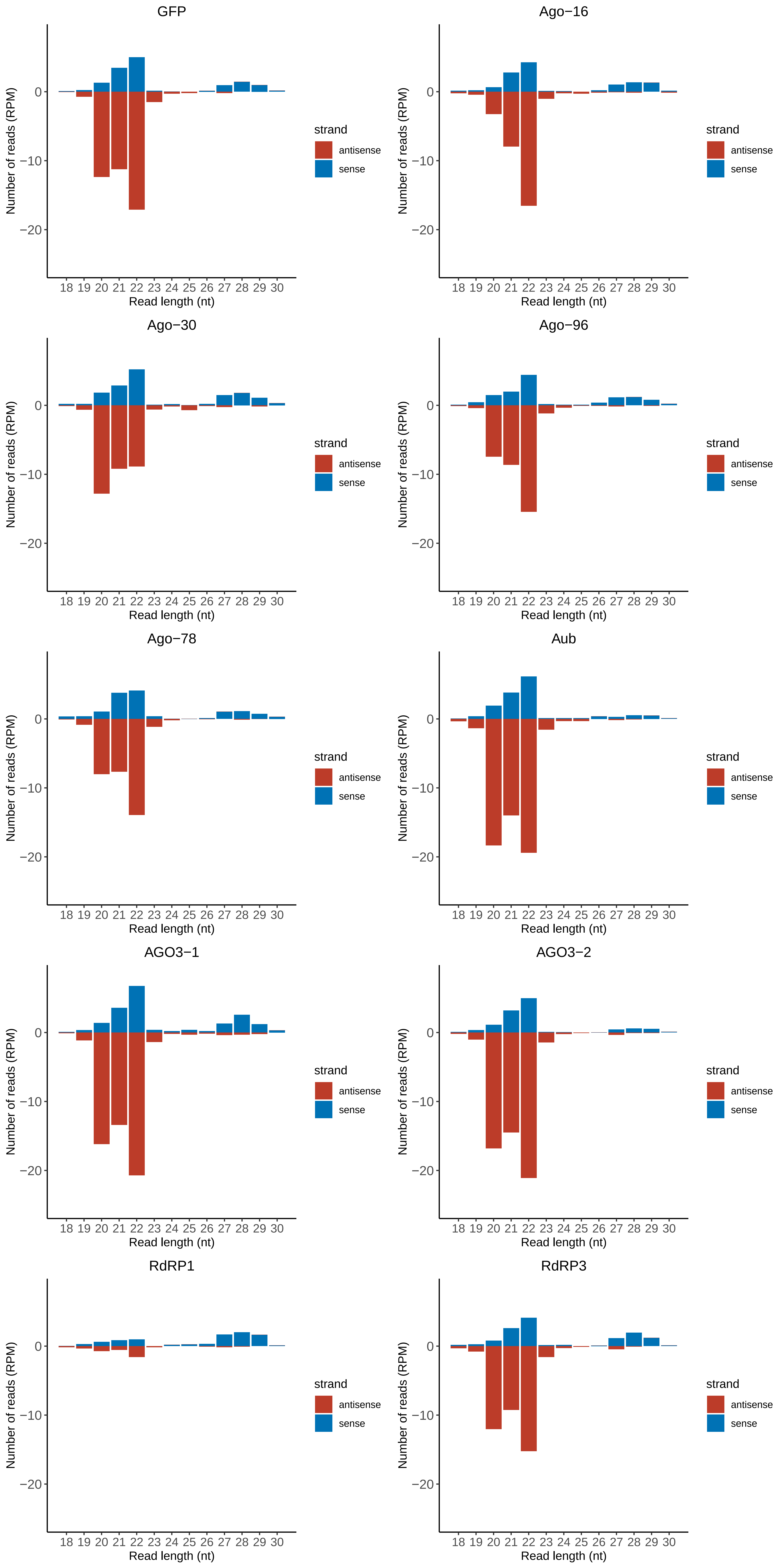

### Motif:rnd-4_family-3013TE_lengthdistribution.pdf

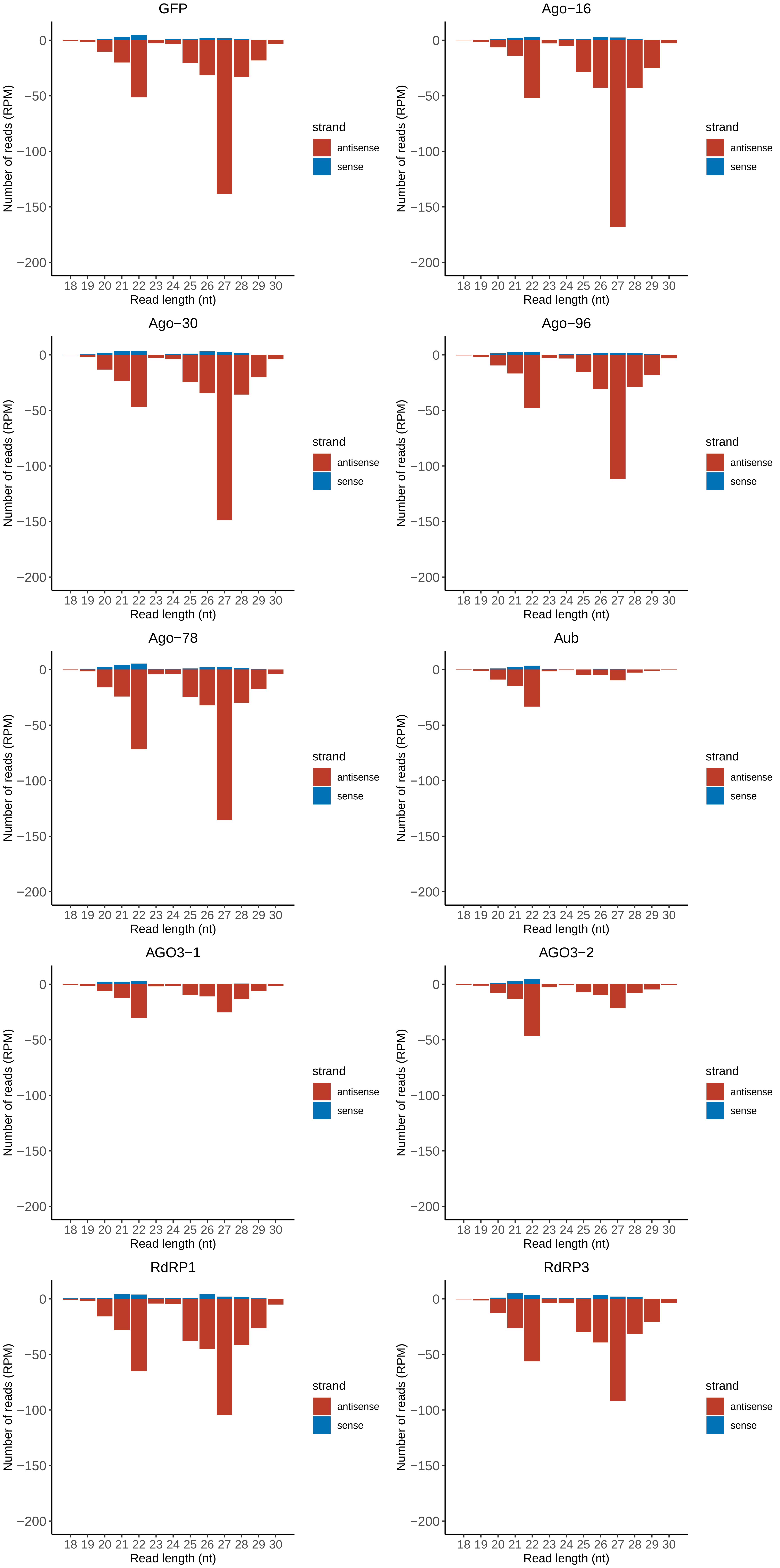

### Motif:rnd-5_family-326TE_lengthdistribution.pdf

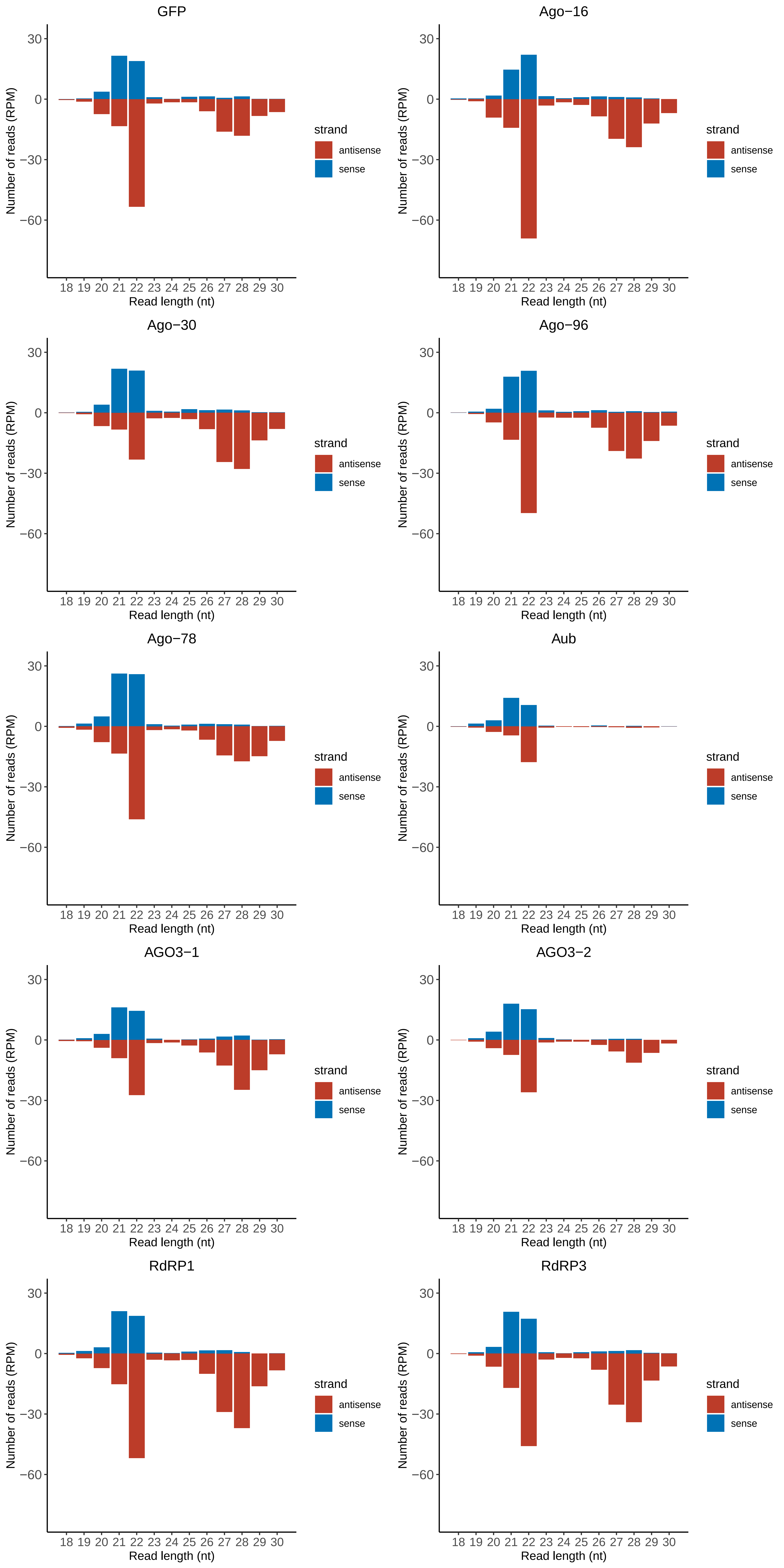

### Motif:rnd-5_family-557TE_lengthdistribution.pdf

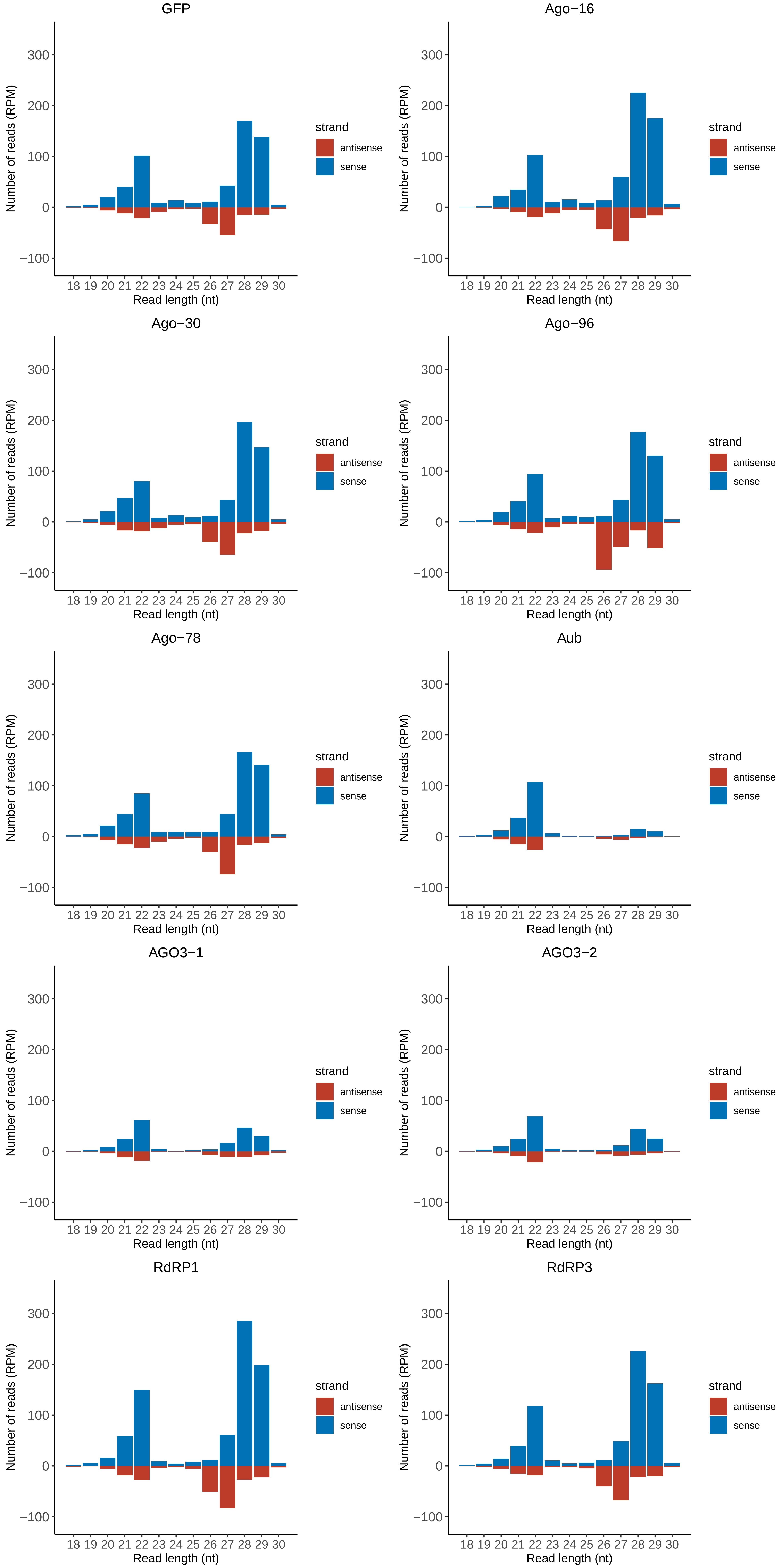

### Motif:rnd-5_family-601TE_lengthdistribution.pdf

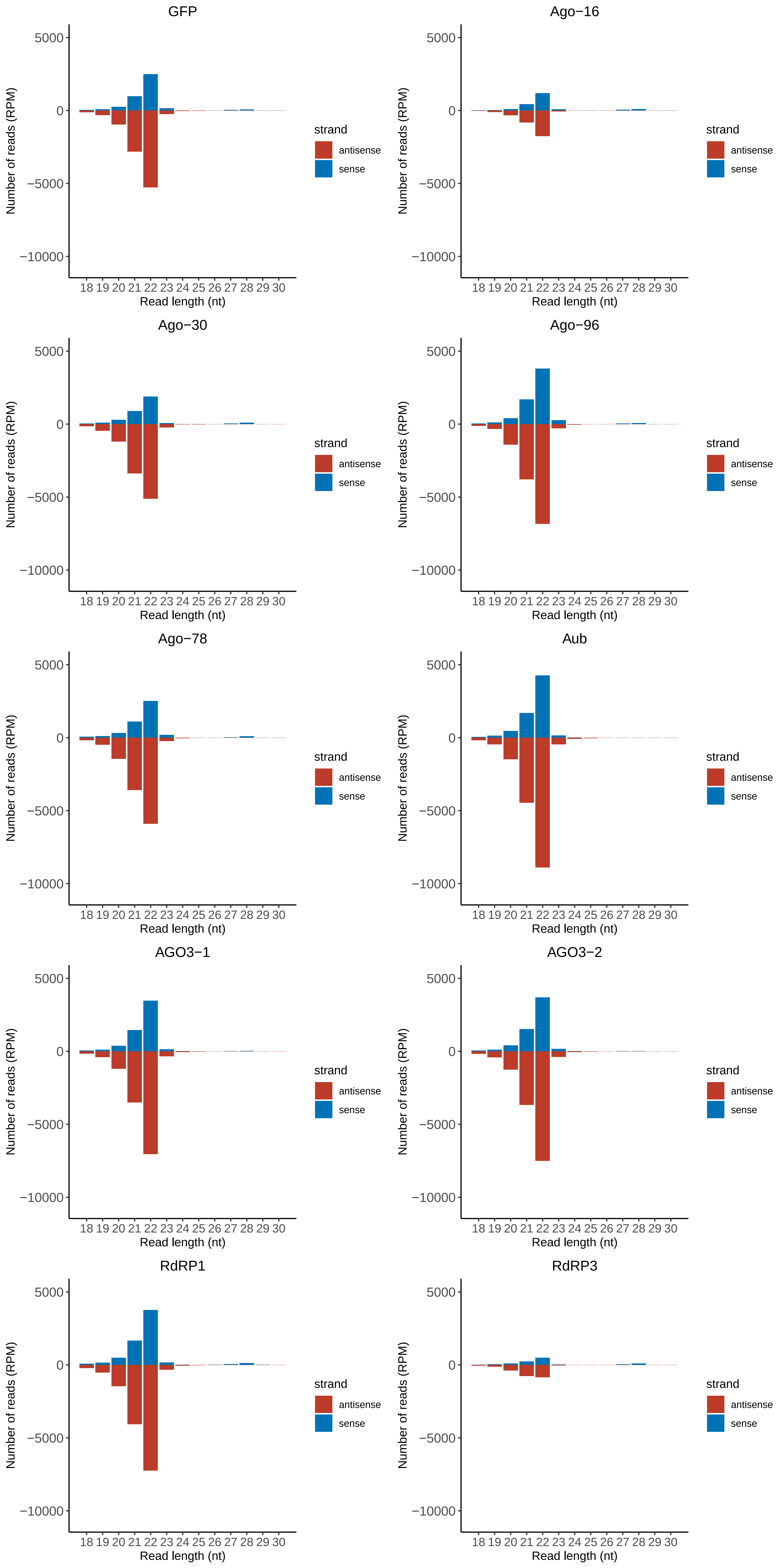

### Motif:rnd-5_family-835TE_lengthdistribution.pdf

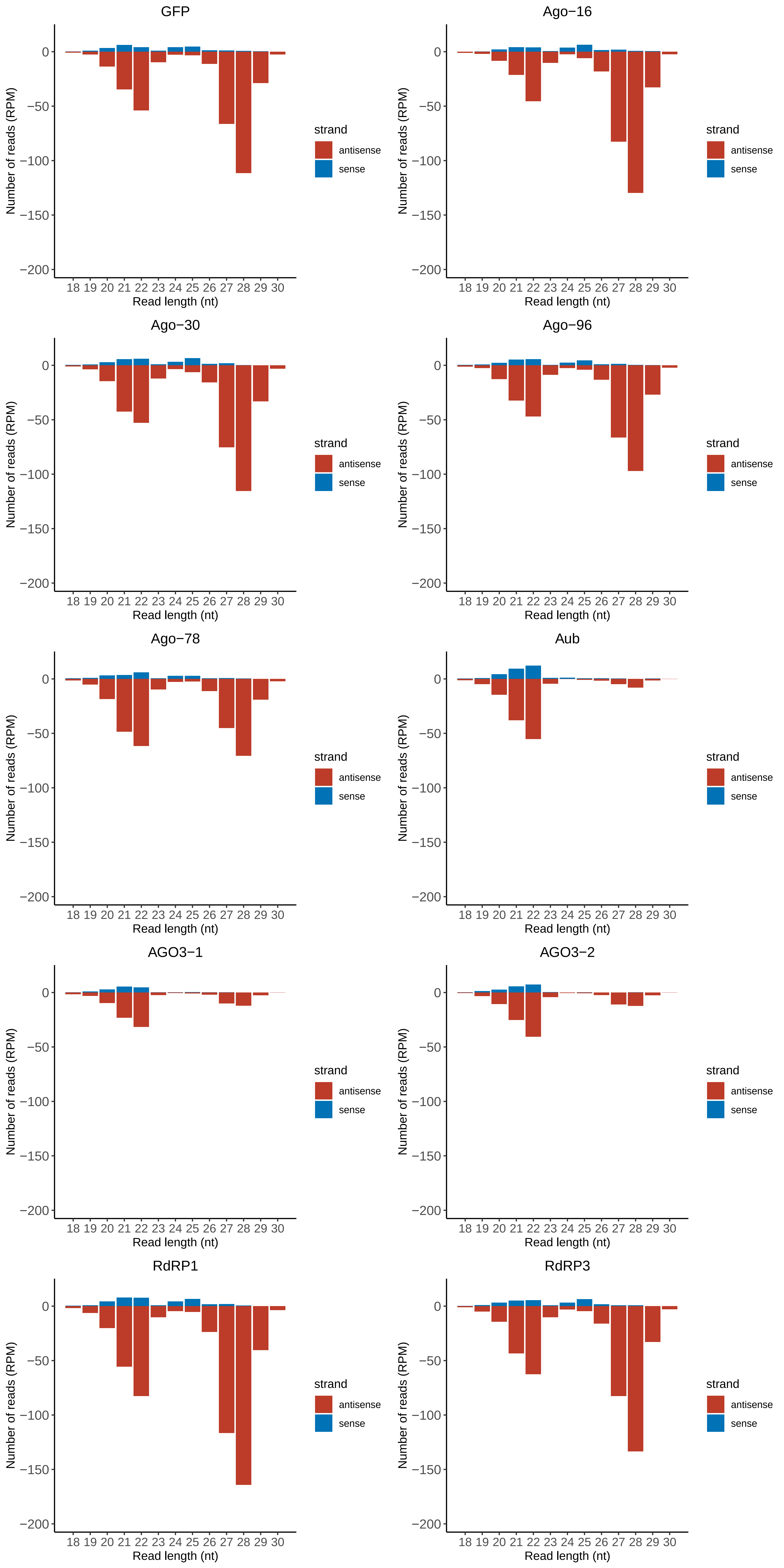

### Motif:rnd-5_family-1208TE_lengthdistribution.pdf

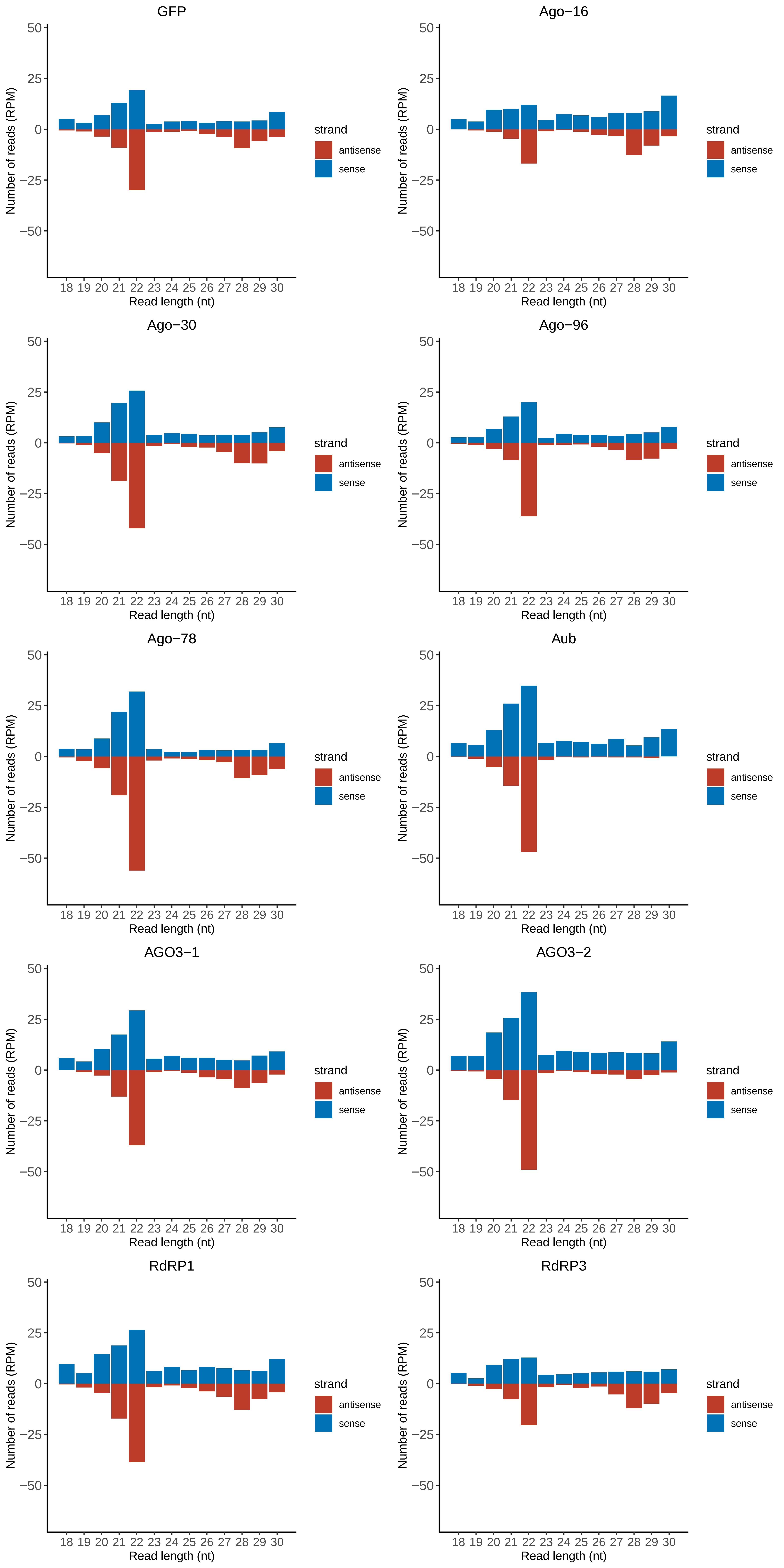

### Motif:rnd-5_family-1416TE_lengthdistribution.pdf

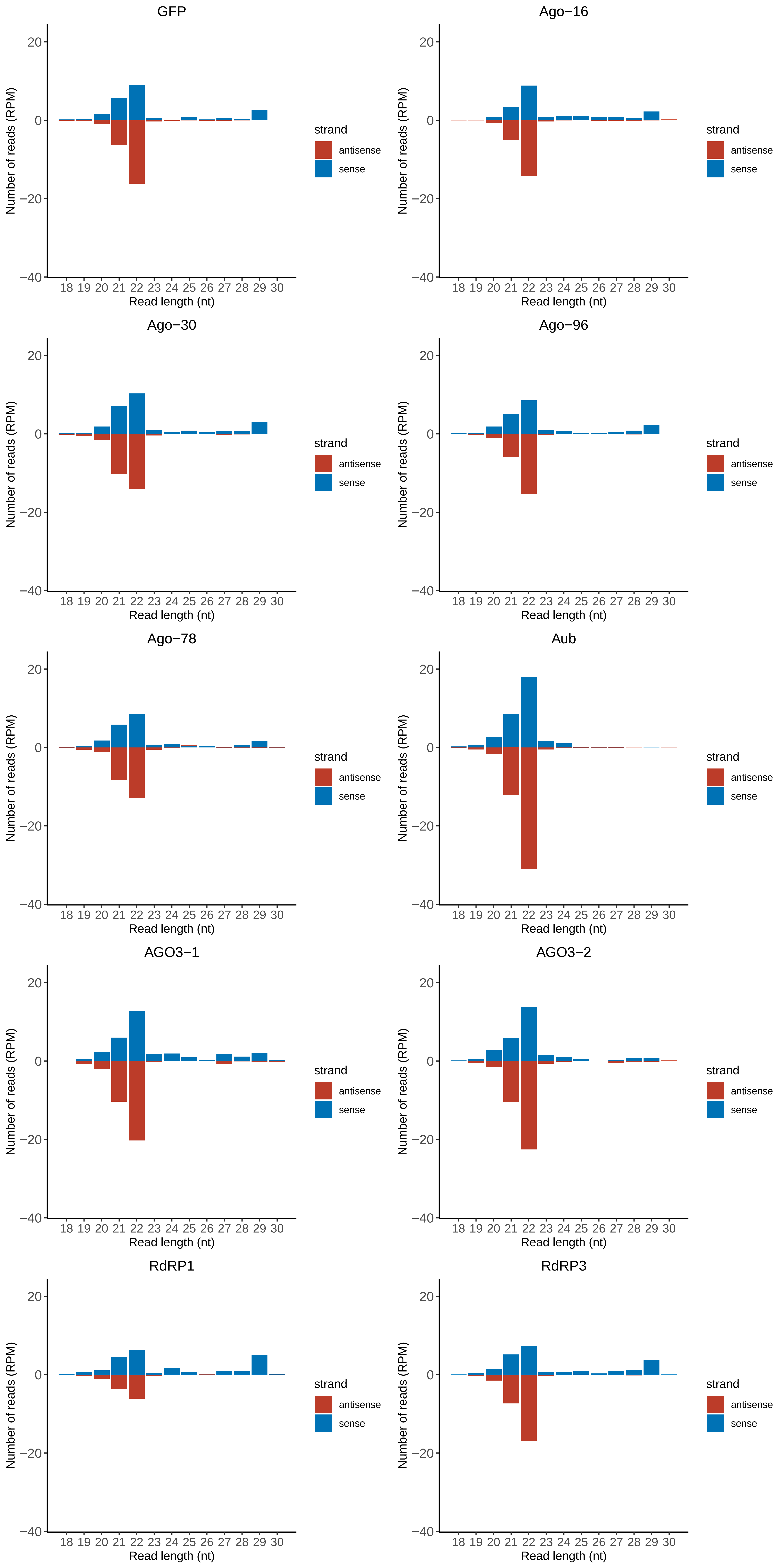

### Motif:rnd-5_family-2196TE_lengthdistribution.pdf

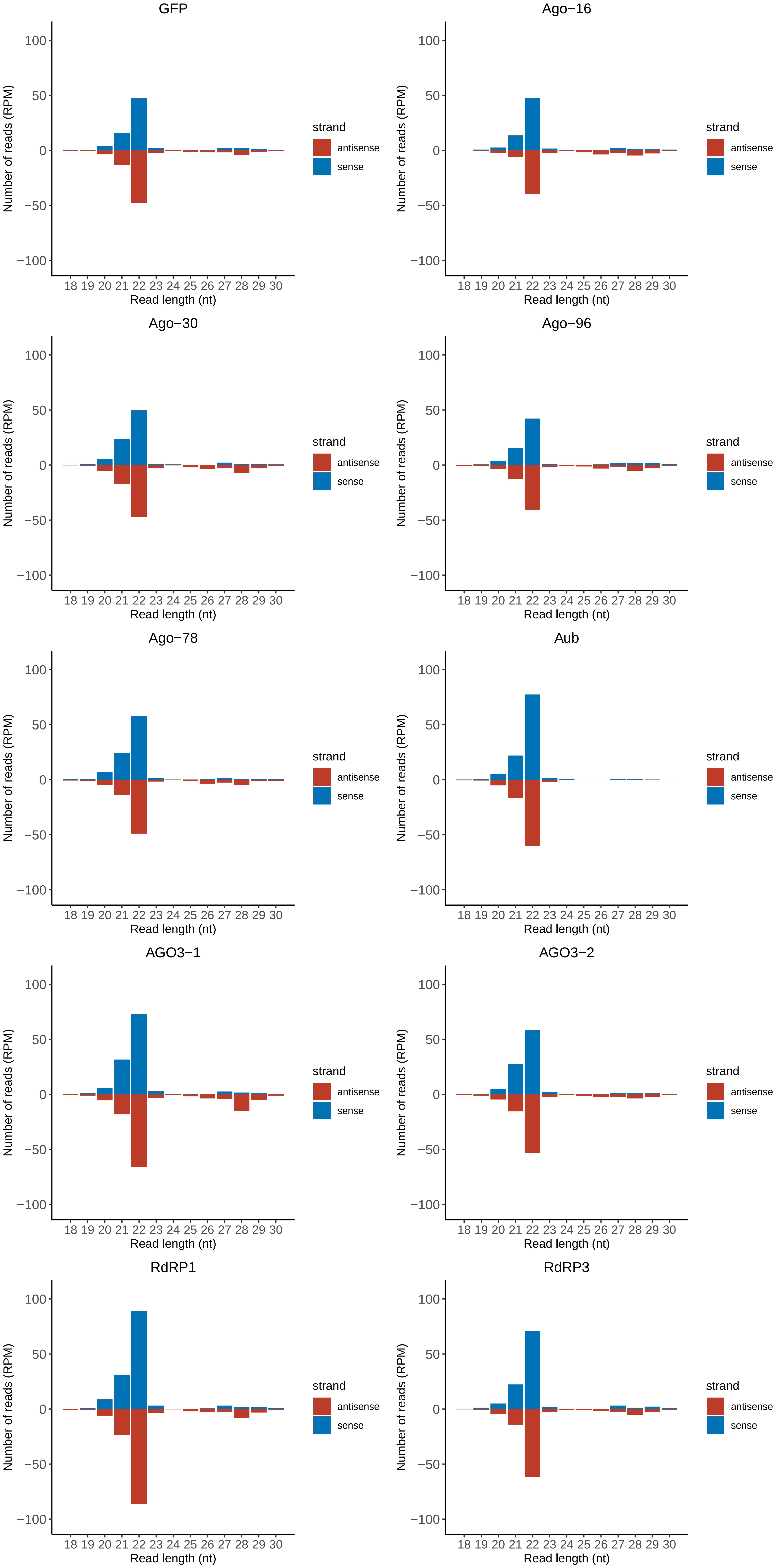

### Motif:rnd-5_family-2666TE_lengthdistribution.pdf

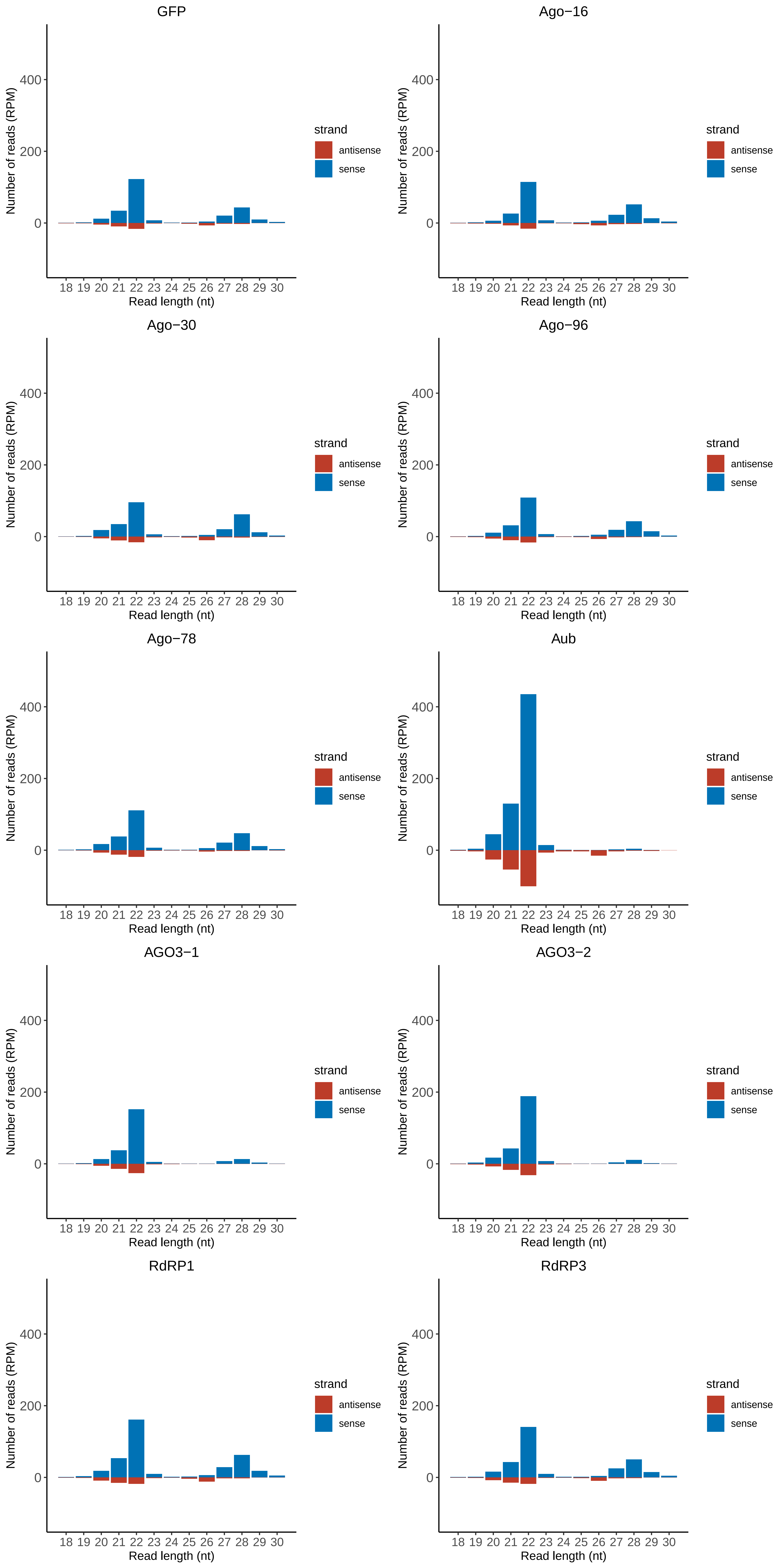

### Motif:rnd-5_family-4359TE_lengthdistribution.pdf

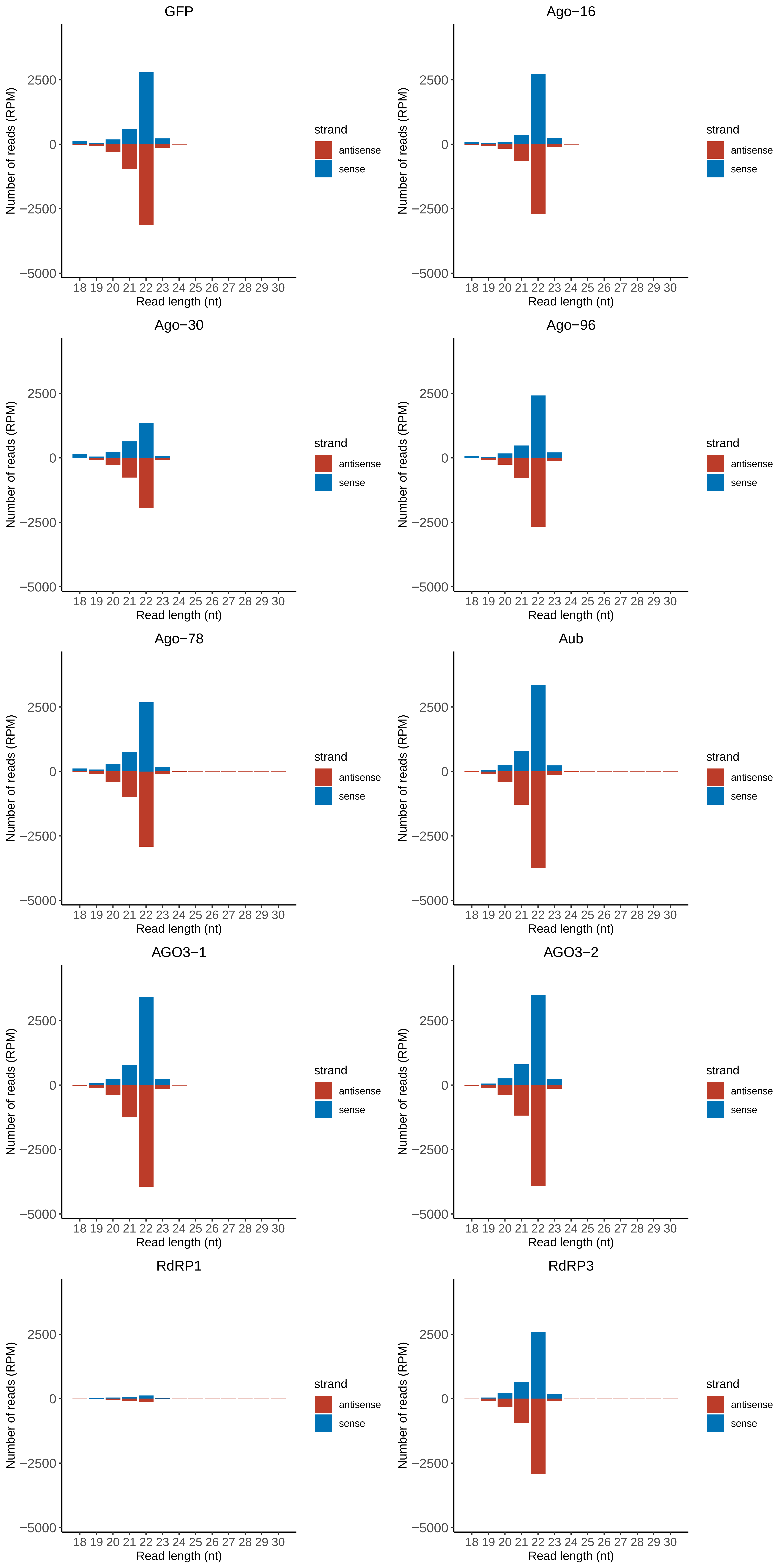

### Motif:rnd-5_family-4592TE_lengthdistribution.pdf

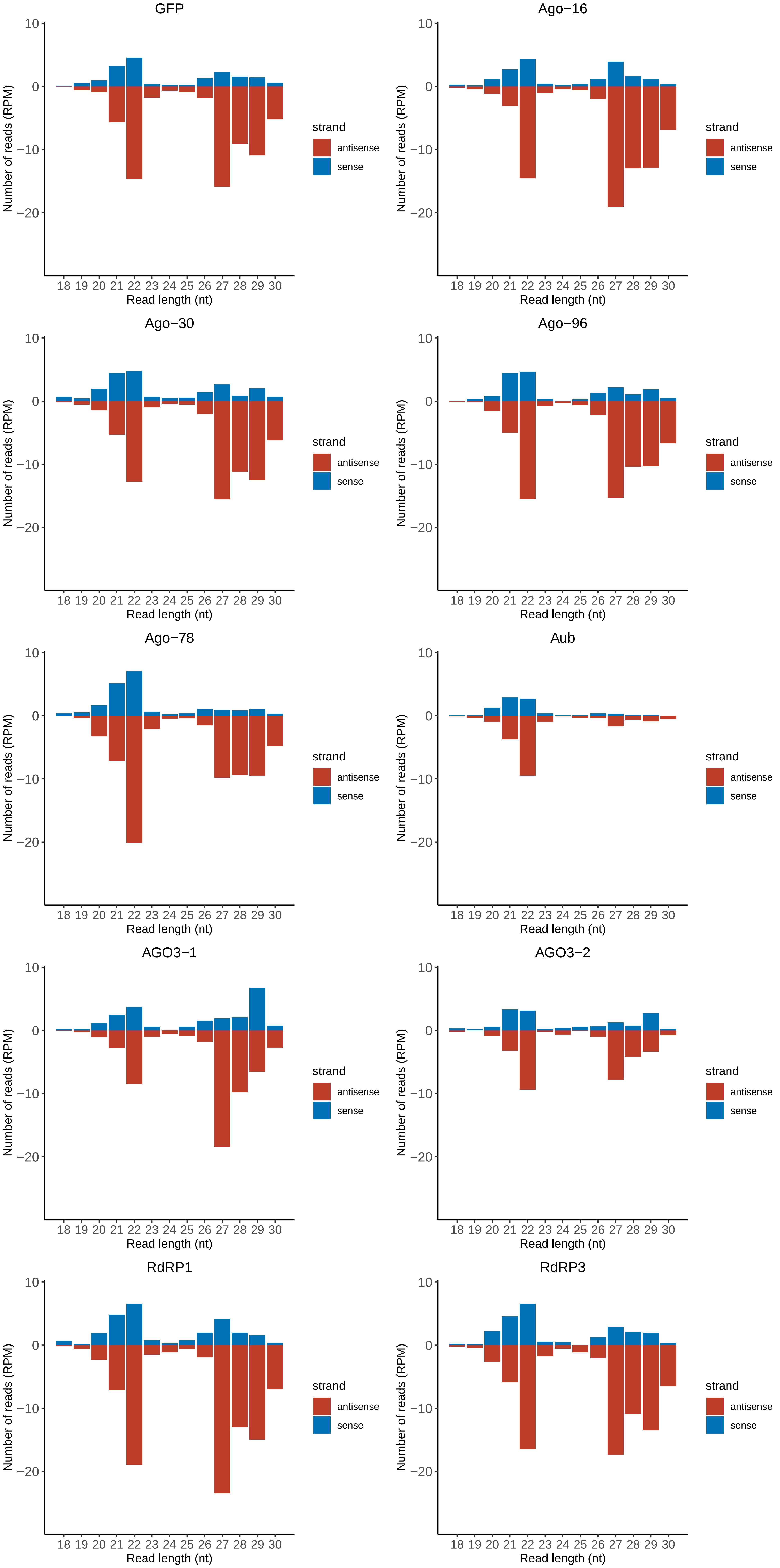

### Motif:rnd-5_family-4690TE_lengthdistribution.pdf

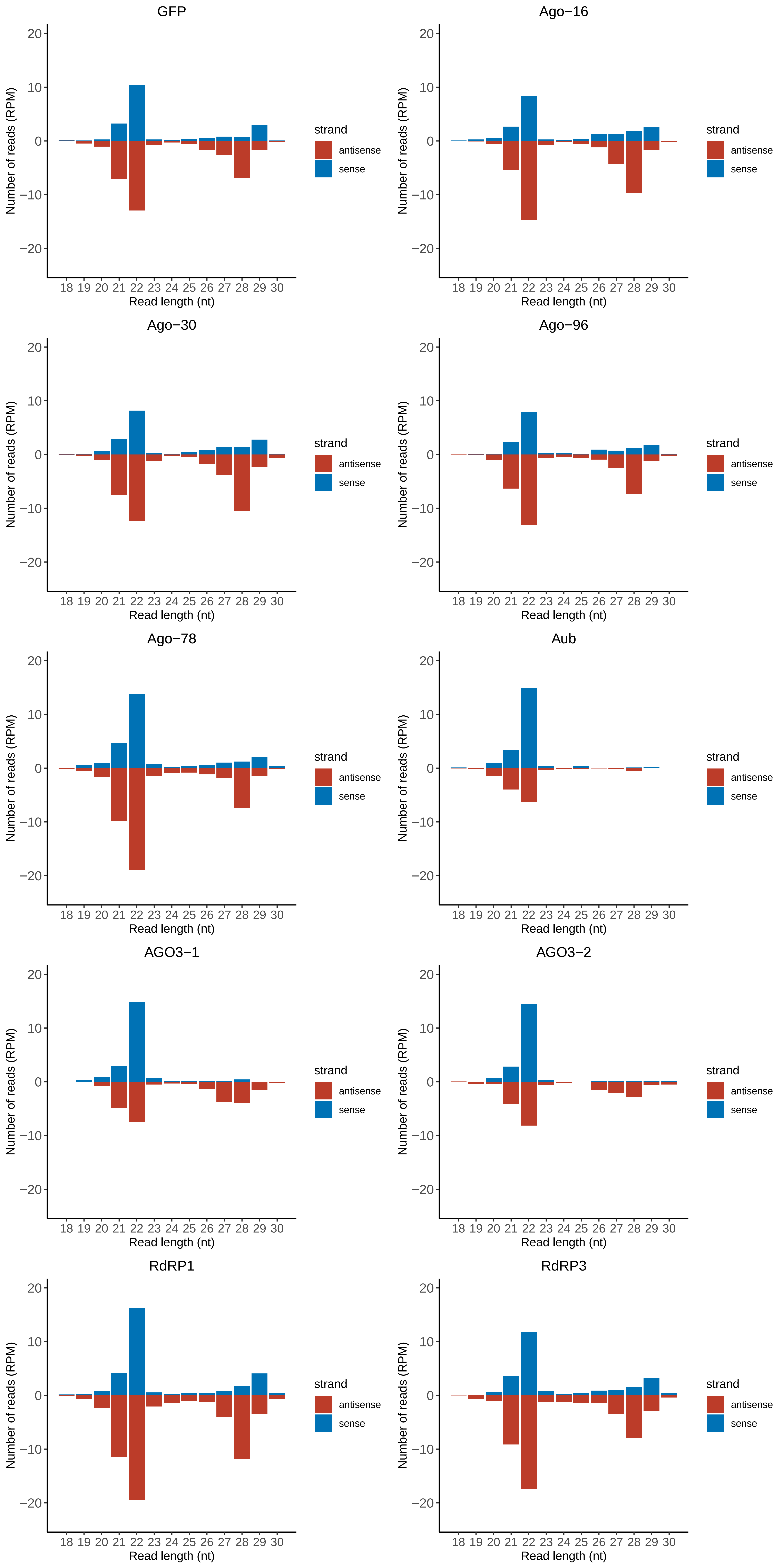

### Motif:rnd-6_family-84TE_lengthdistribution.pdf

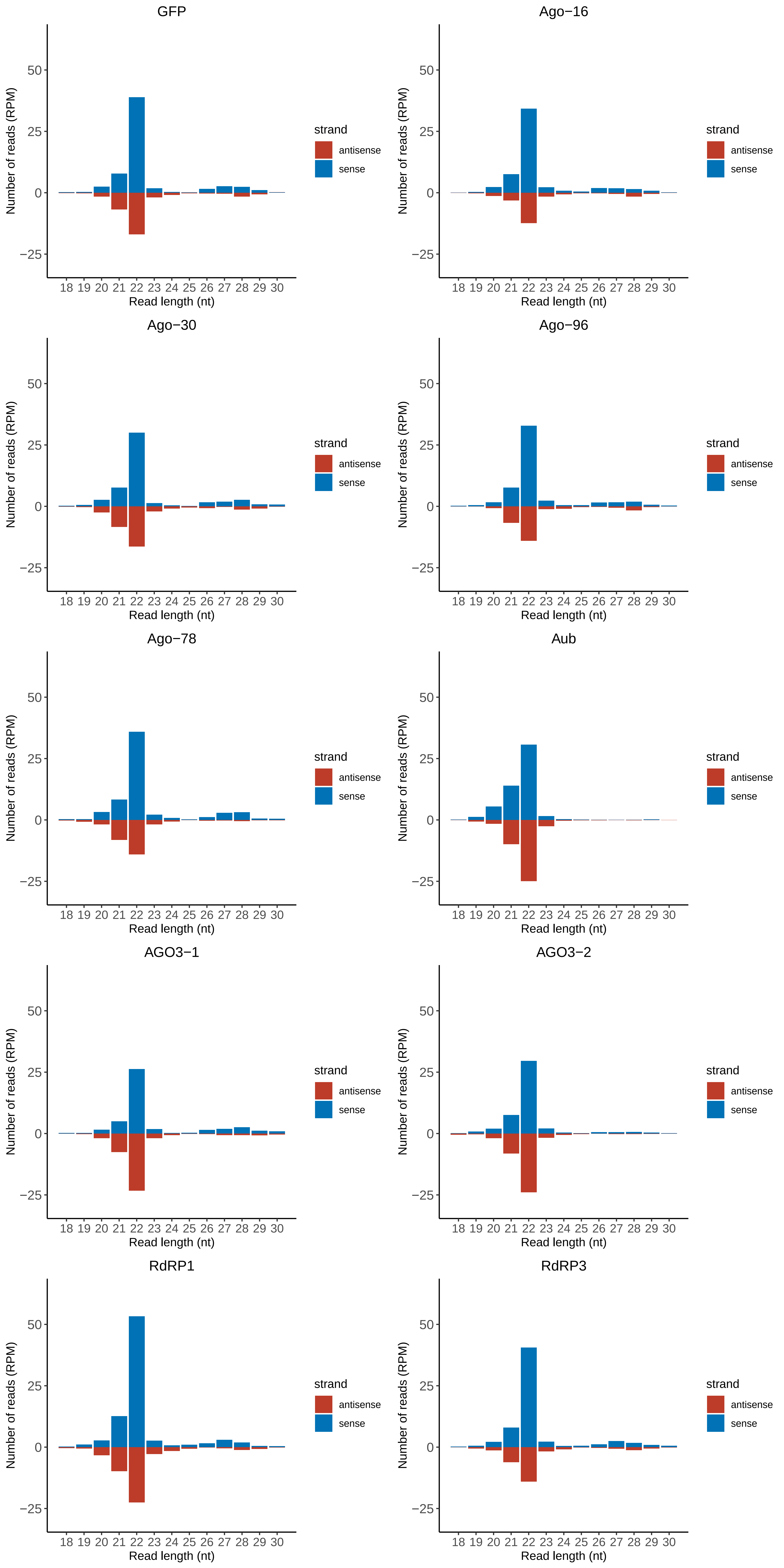

### Motif:rnd-6_family-159TE_lengthdistribution.pdf

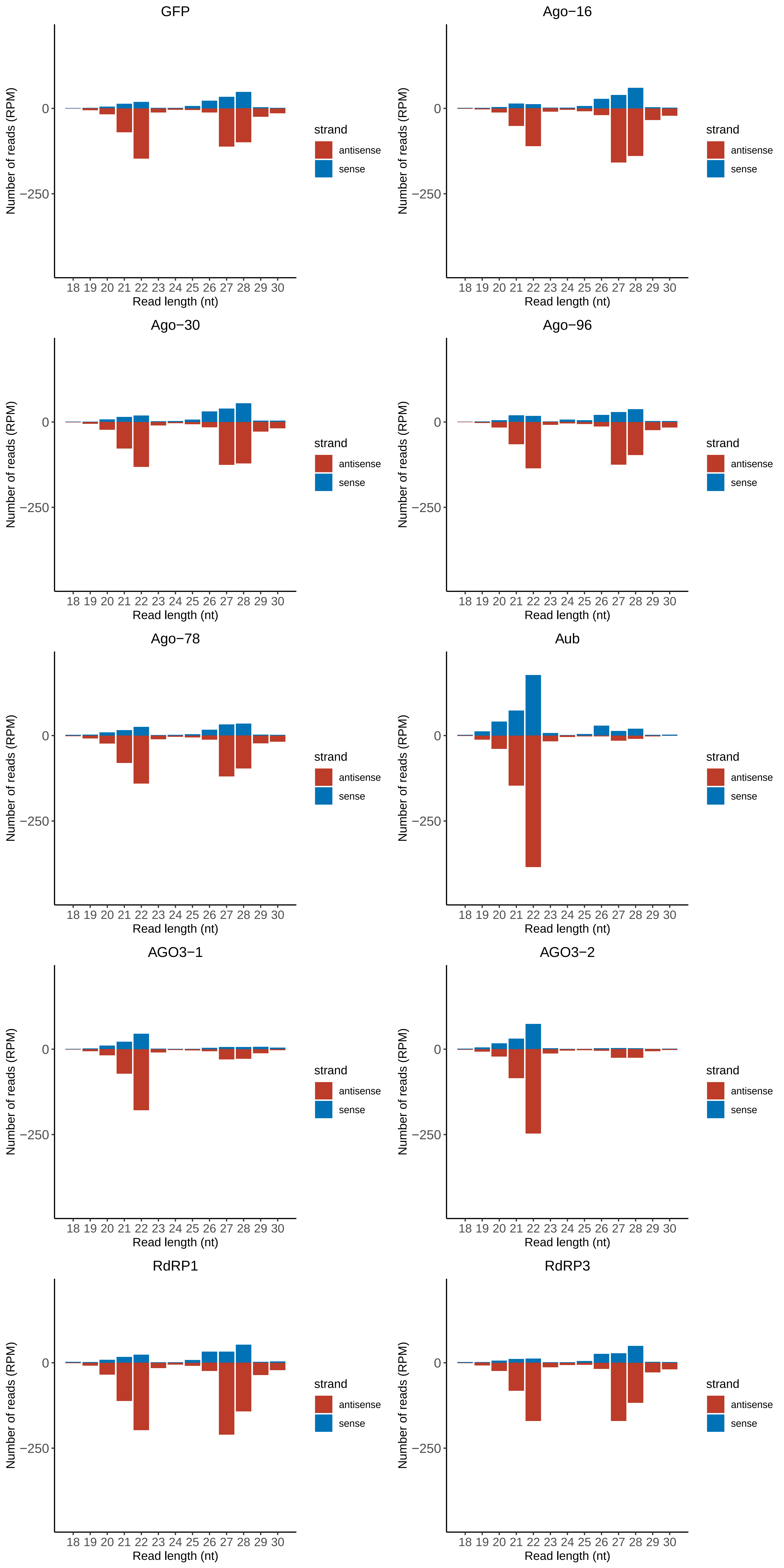

### Motif:rnd-6_family-339TE_lengthdistribution.pdf

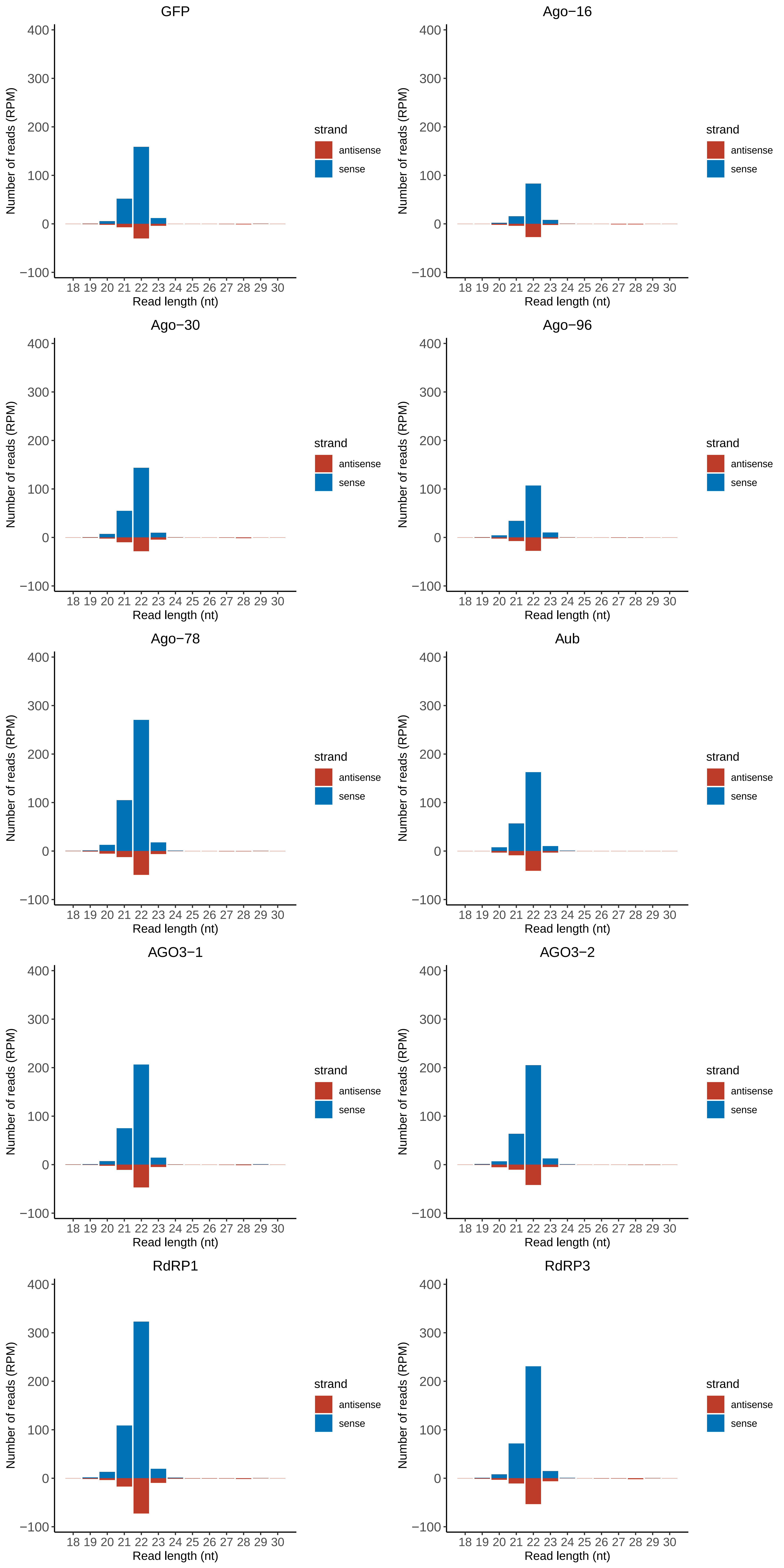

### Motif:rnd-6_family-423TE_lengthdistribution.pdf

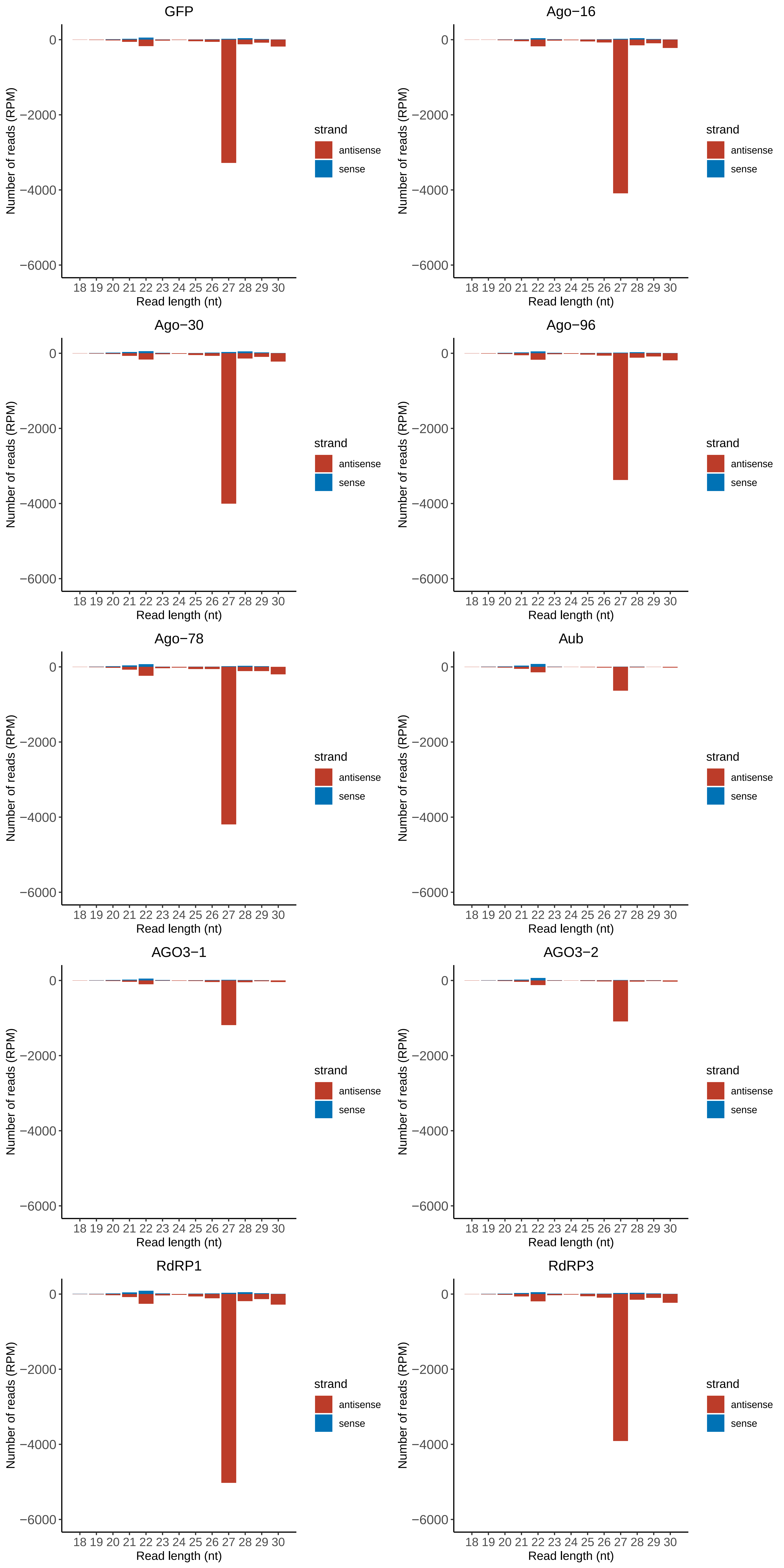

### Motif:rnd-6_family-427TE_lengthdistribution.pdf

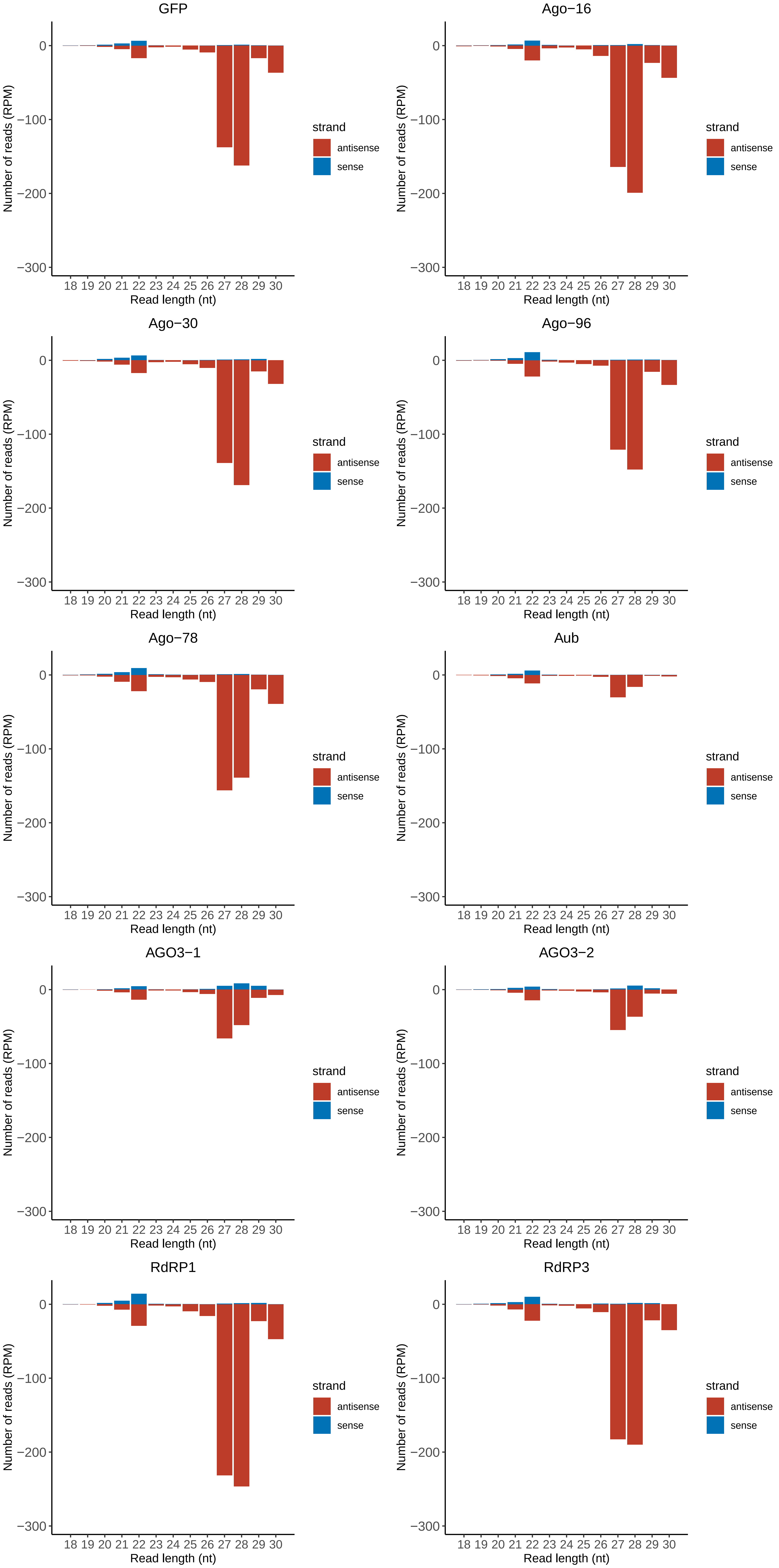

### Motif:rnd-6_family-620TE_lengthdistribution.pdf

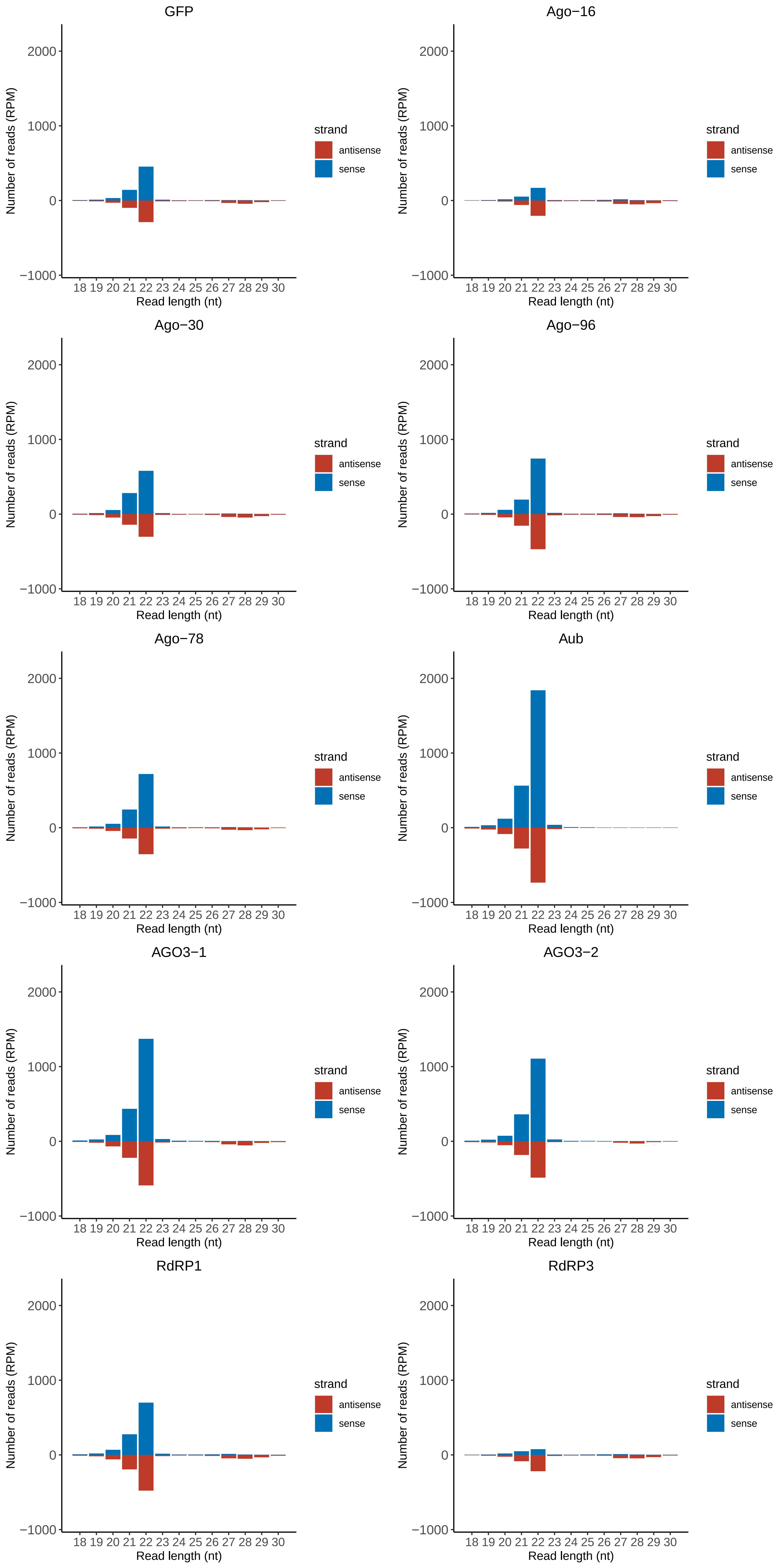

### Motif:rnd-6_family-891TE_lengthdistribution.pdf

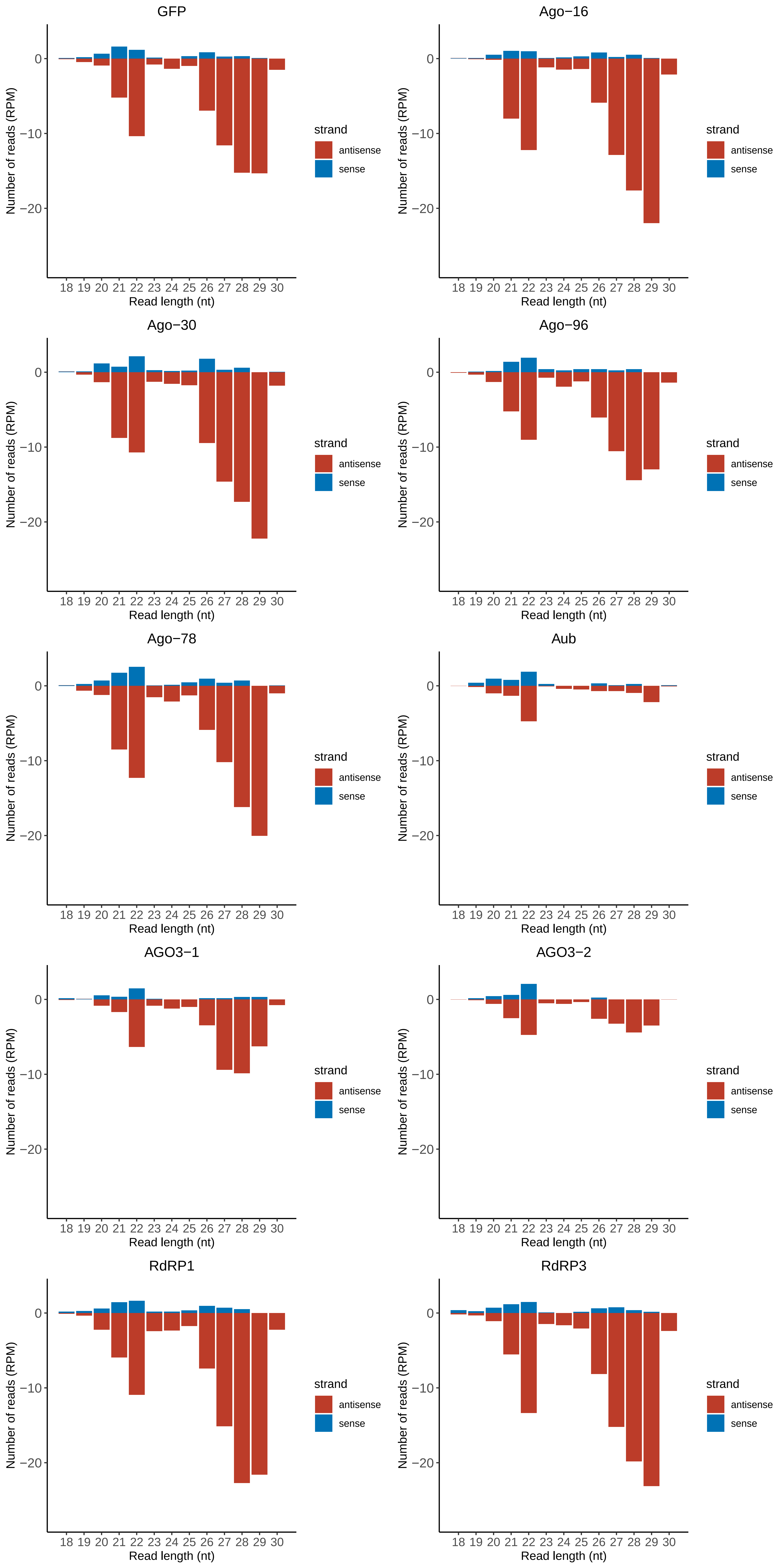

### Motif:rnd-6_family-1239TE_lengthdistribution.pdf

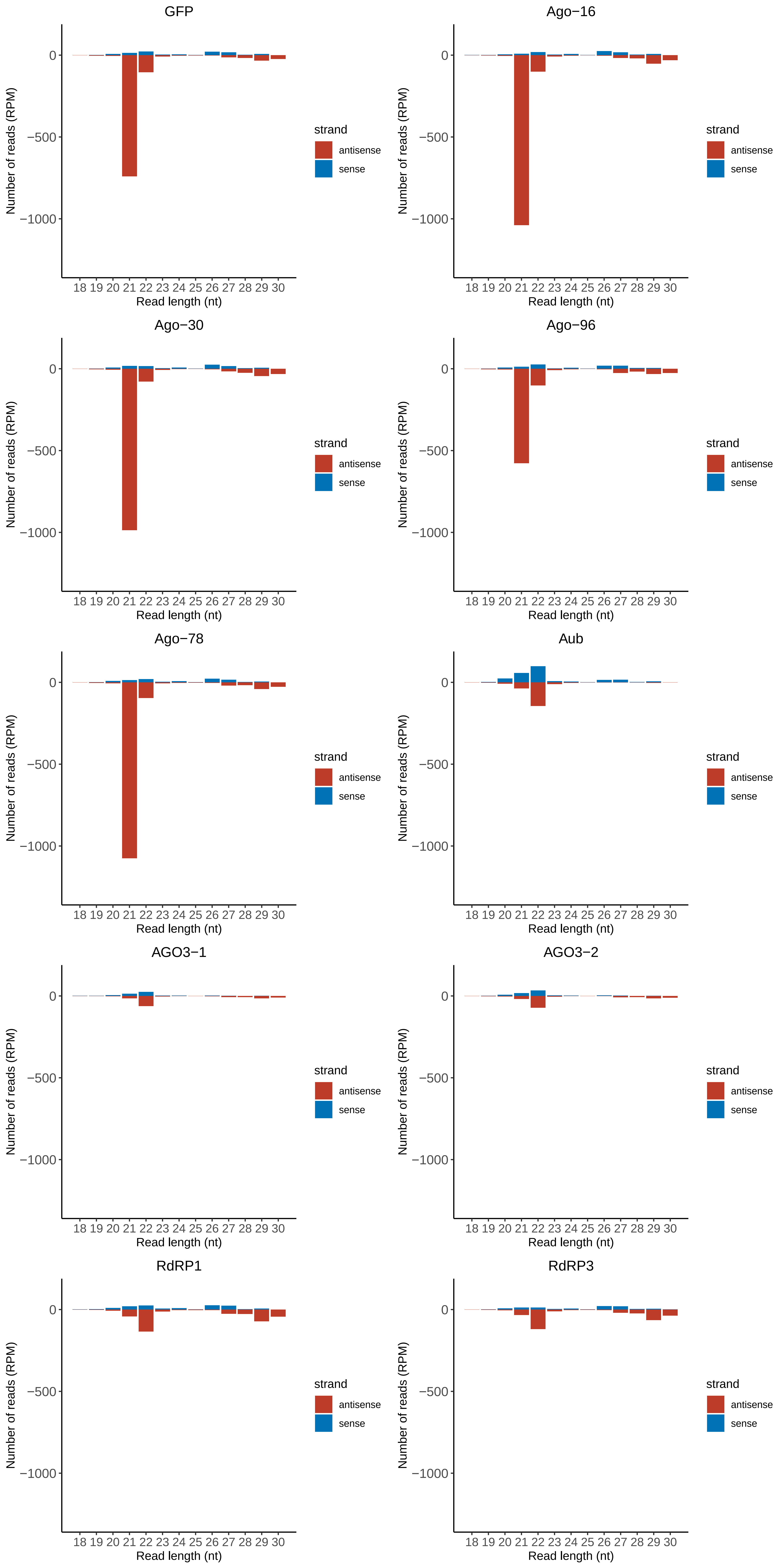

### Motif:rnd-6_family-1544TE_lengthdistribution.pdf

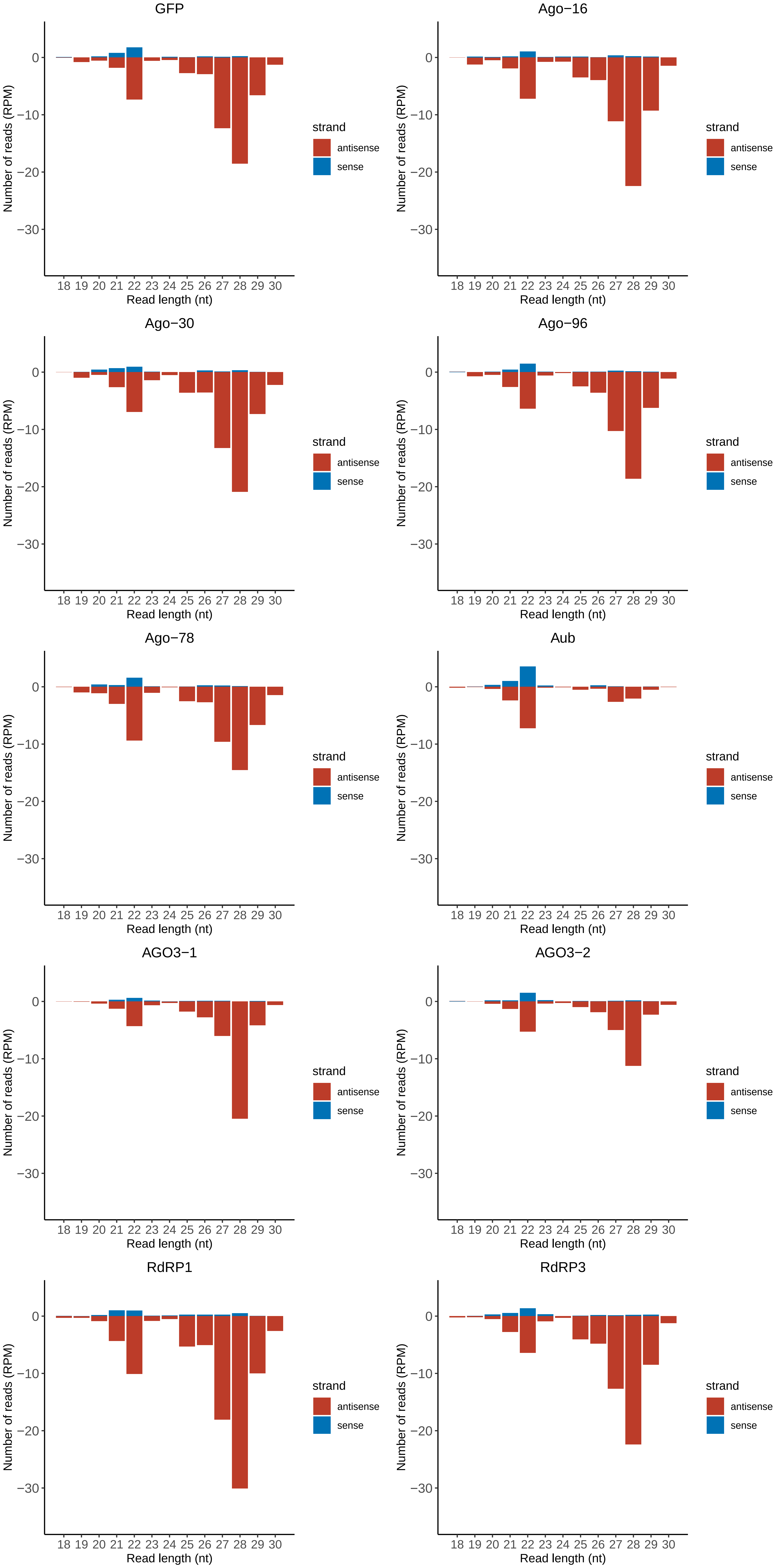

### Motif:rnd-6_family-1977TE_lengthdistribution.pdf

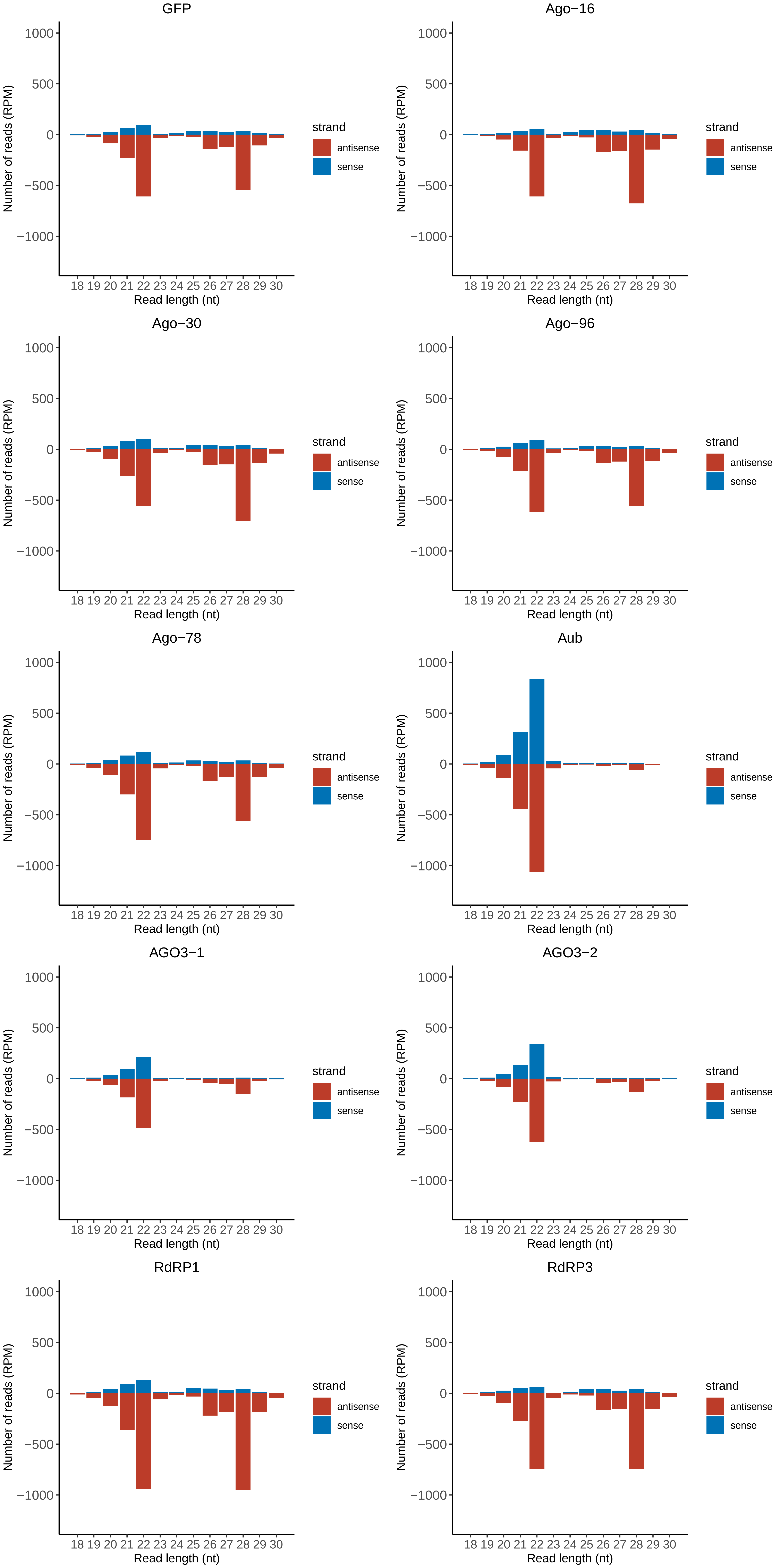

### Motif:rnd-6_family-2023TE_lengthdistribution.pdf

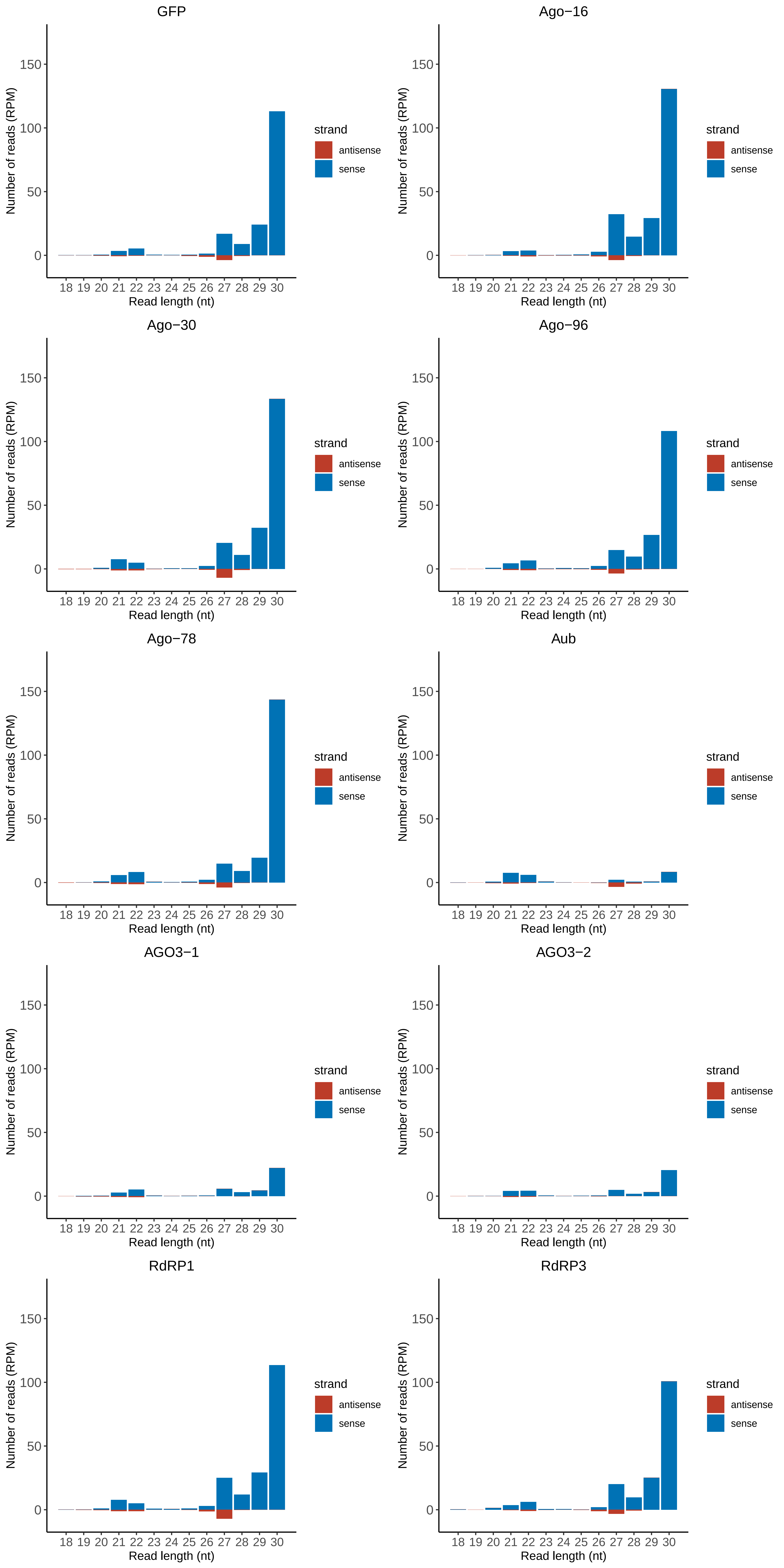

### Motif:rnd-6_family-2044TE_lengthdistribution.pdf

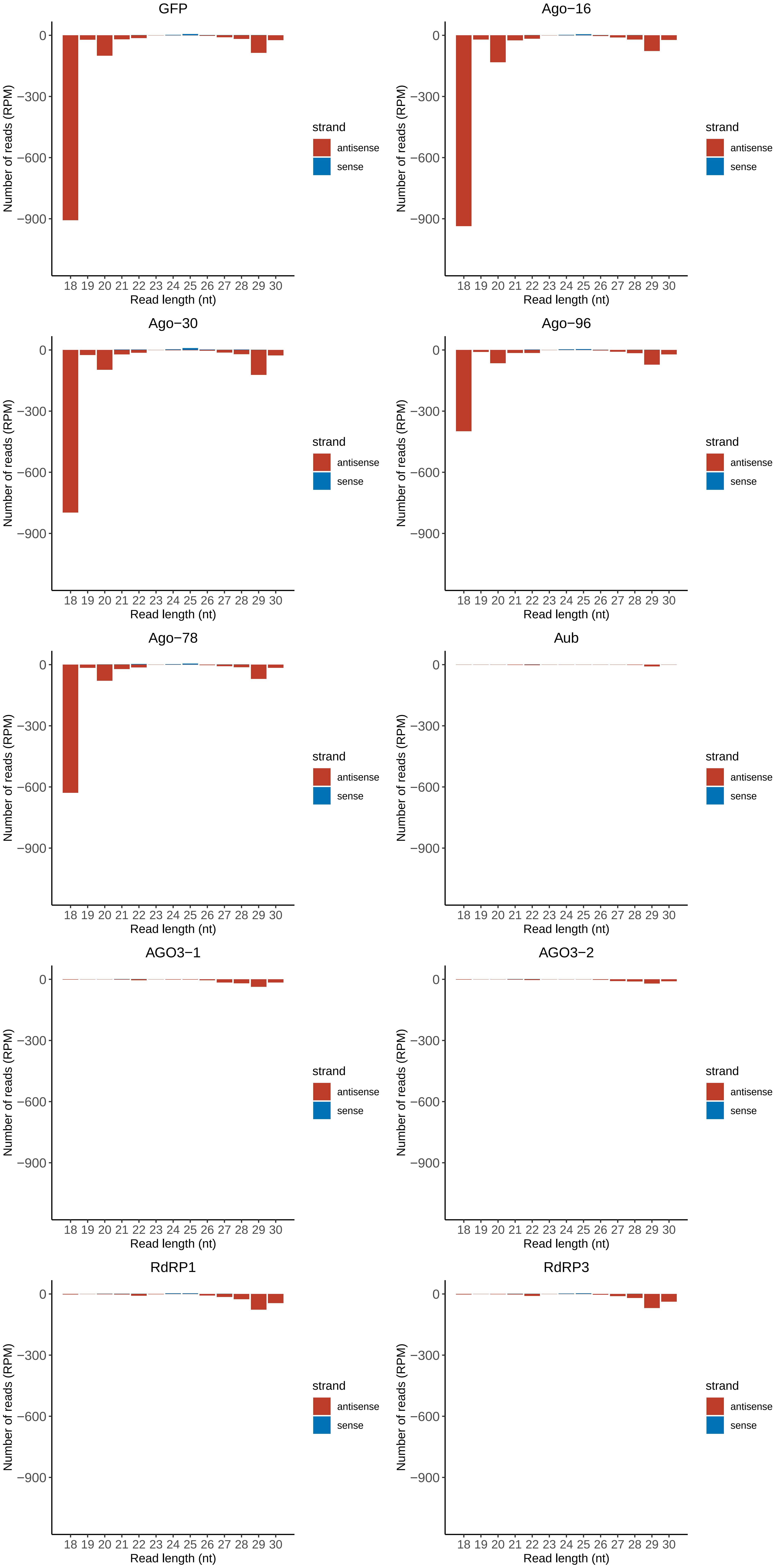

### Motif:rnd-6_family-2652TE_lengthdistribution.pdf

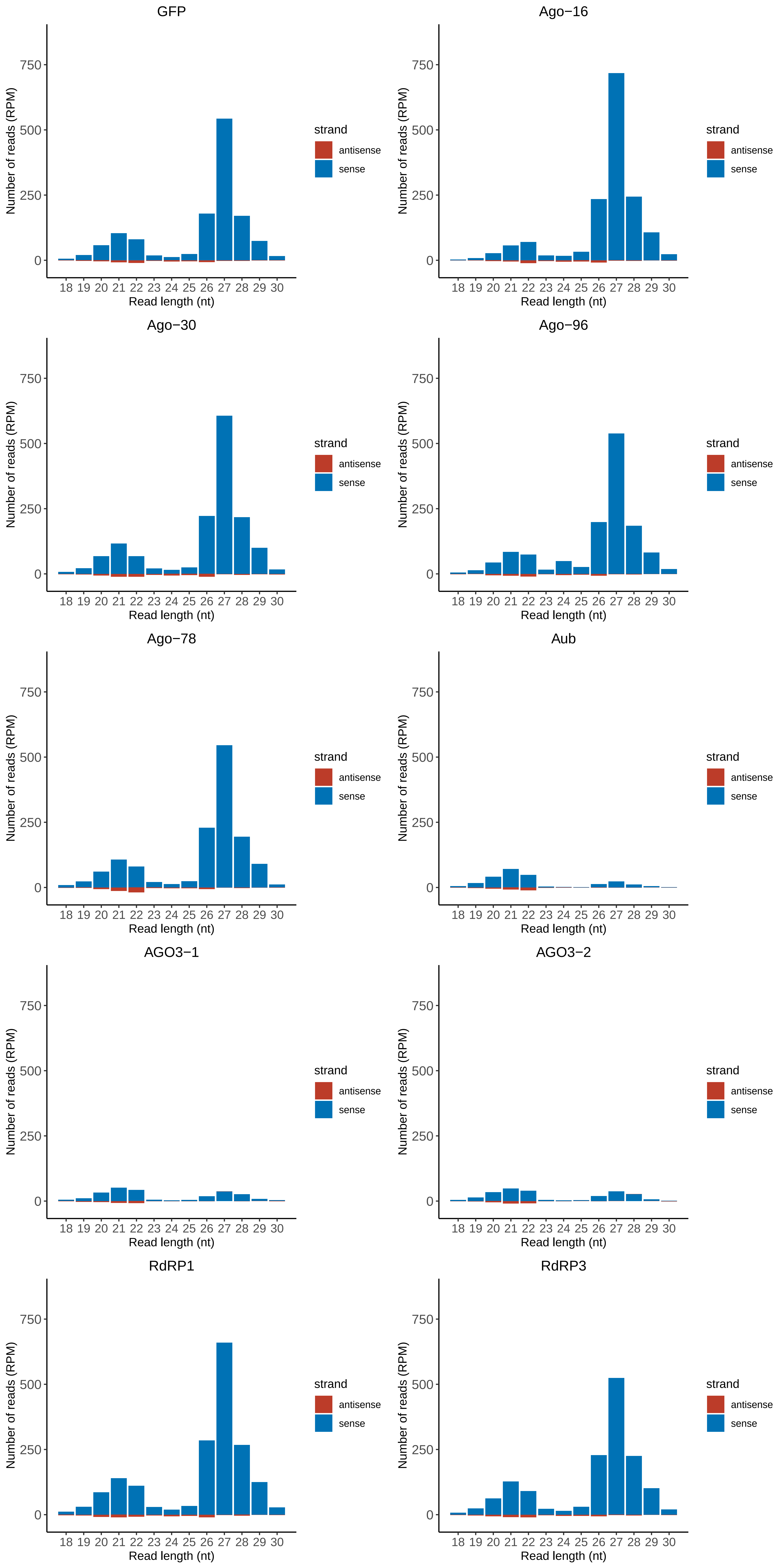

### Motif:rnd-6_family-2764TE_lengthdistribution.pdf

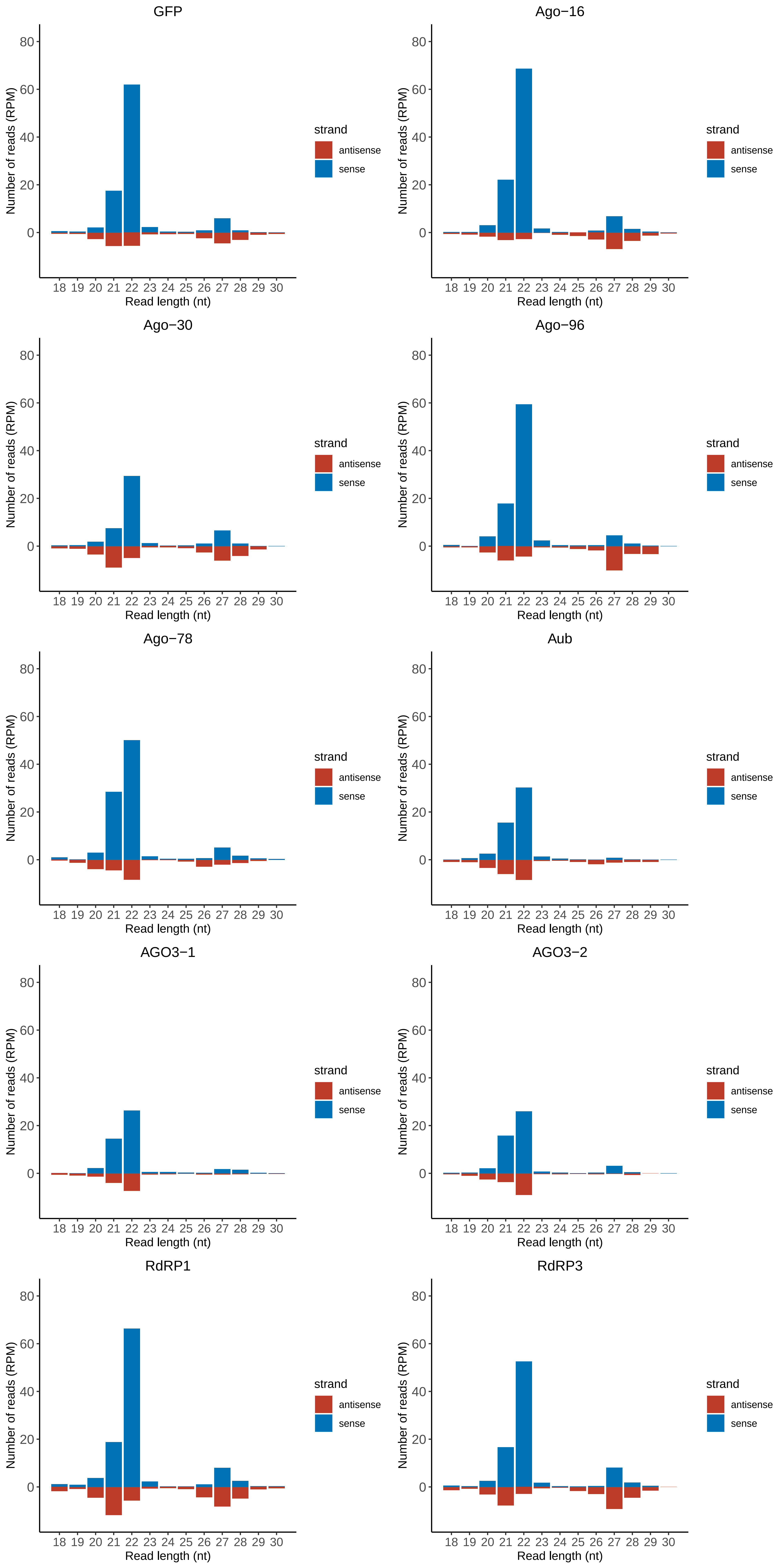

### Motif:rnd-6_family-2808TE_lengthdistribution.pdf

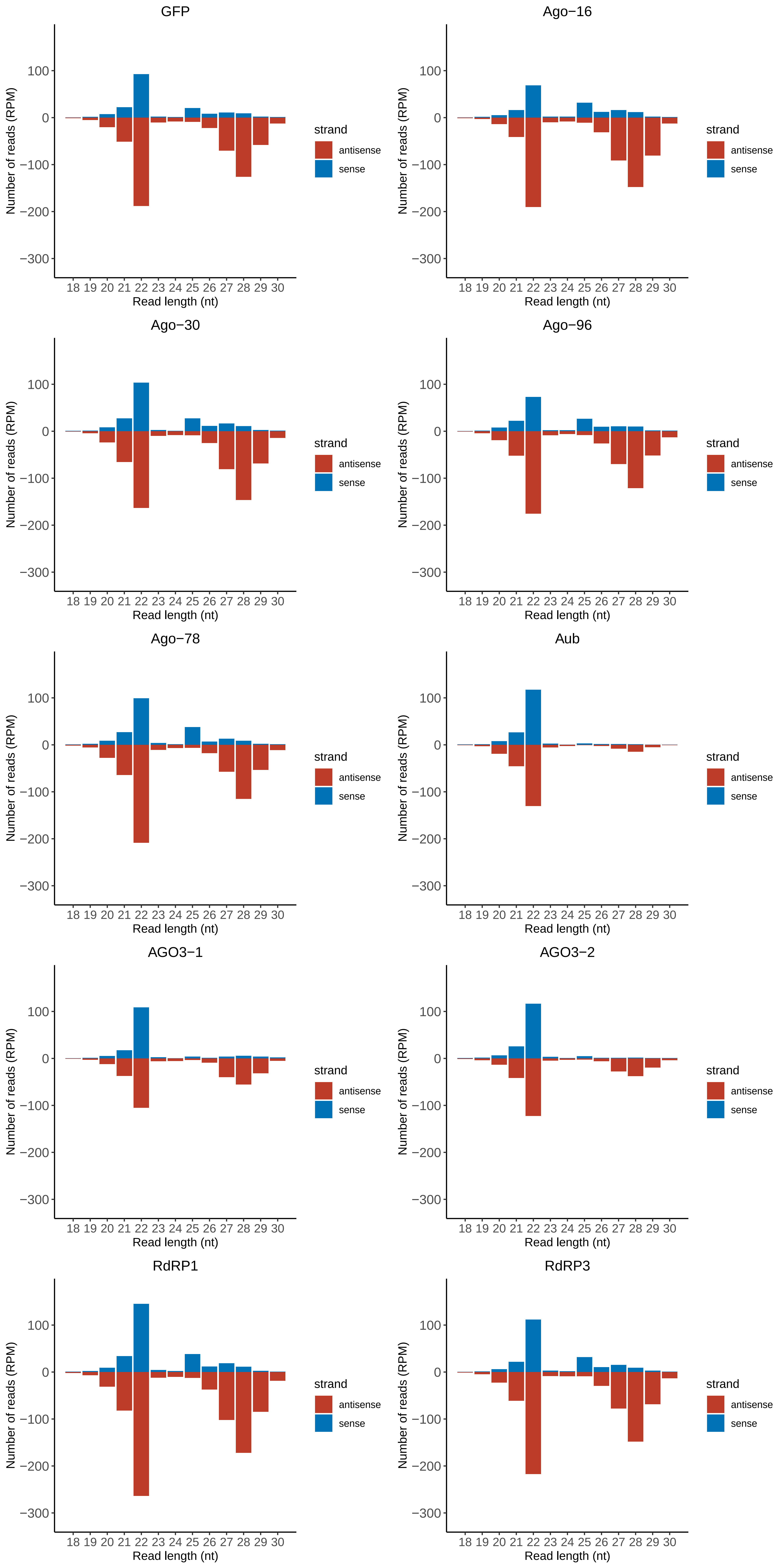

### Motif:rnd-6_family-2888TE_lengthdistribution.pdf

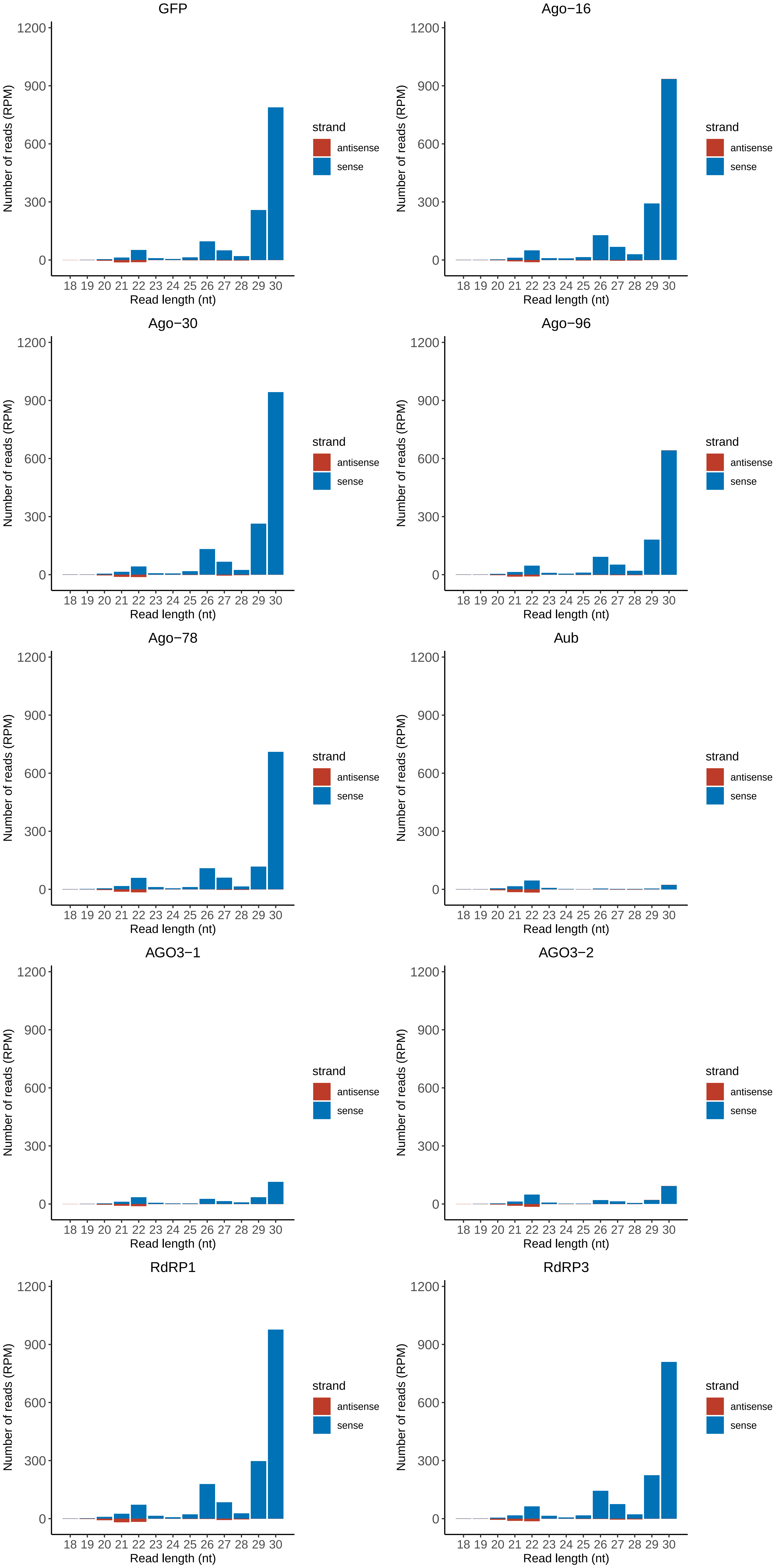

### Motif:rnd-6_family-3082TE_lengthdistribution.pdf

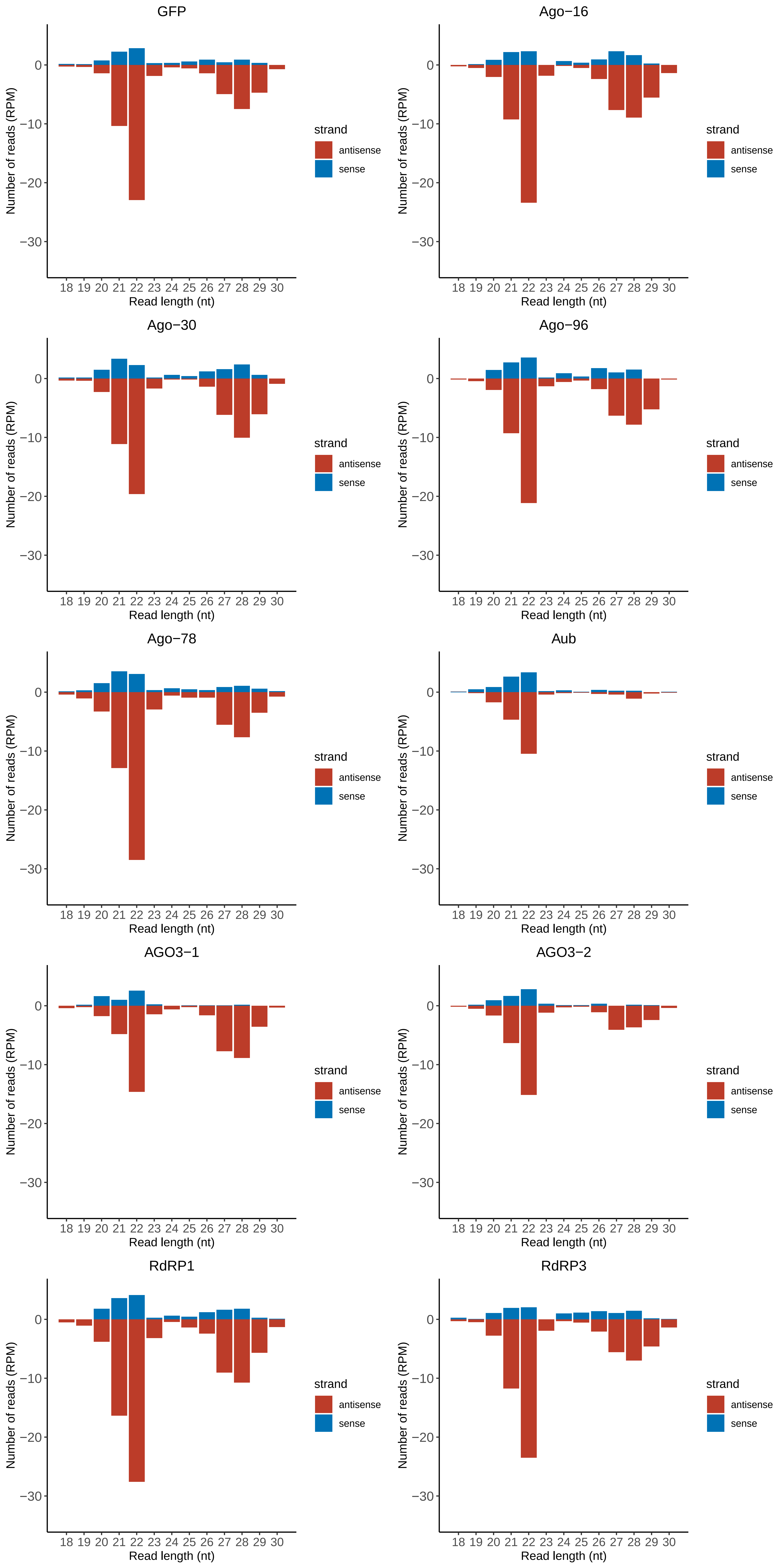

### Motif:rnd-6_family-3103TE_lengthdistribution.pdf

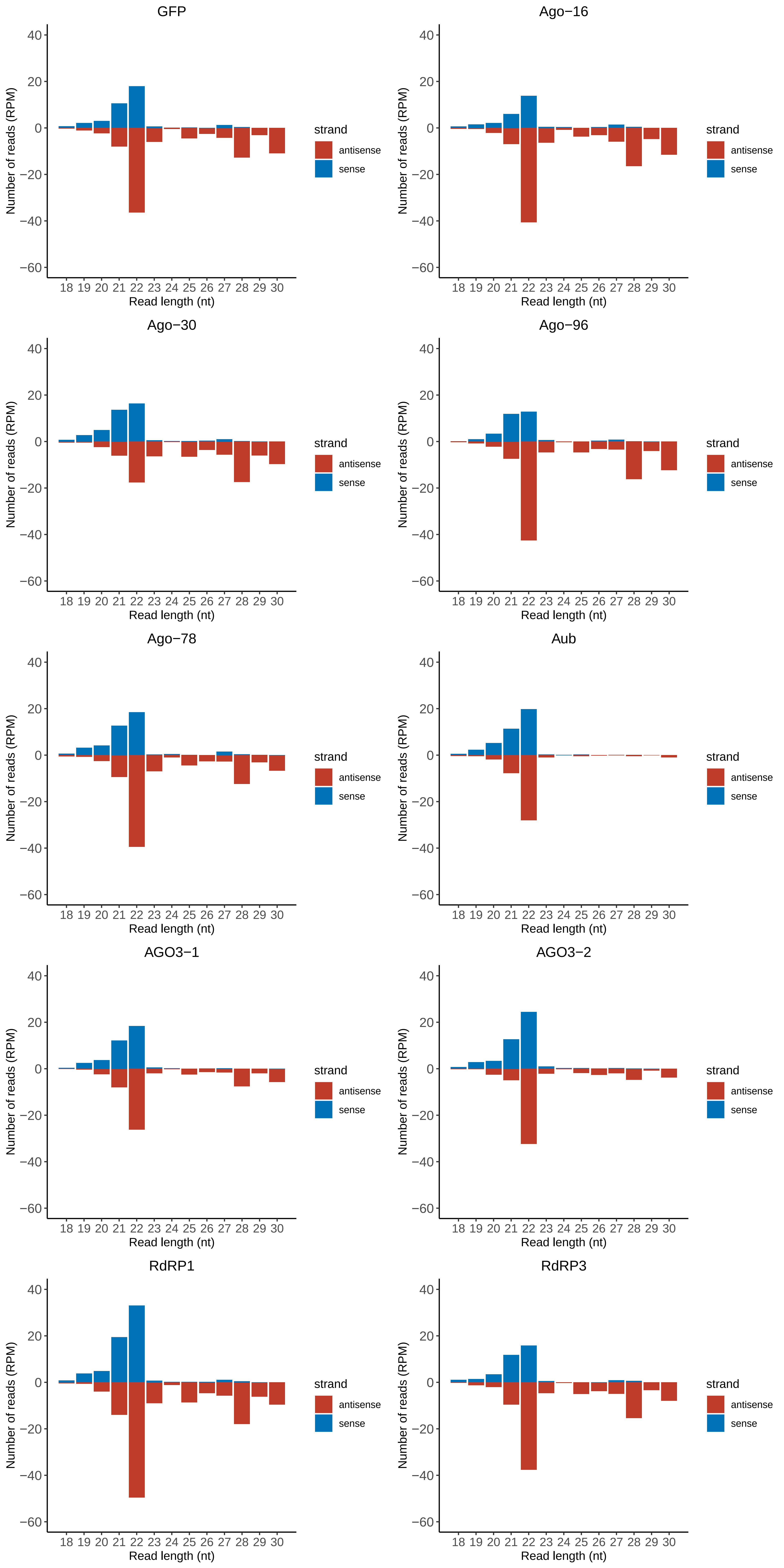

### Motif:rnd-6_family-4001TE_lengthdistribution.pdf

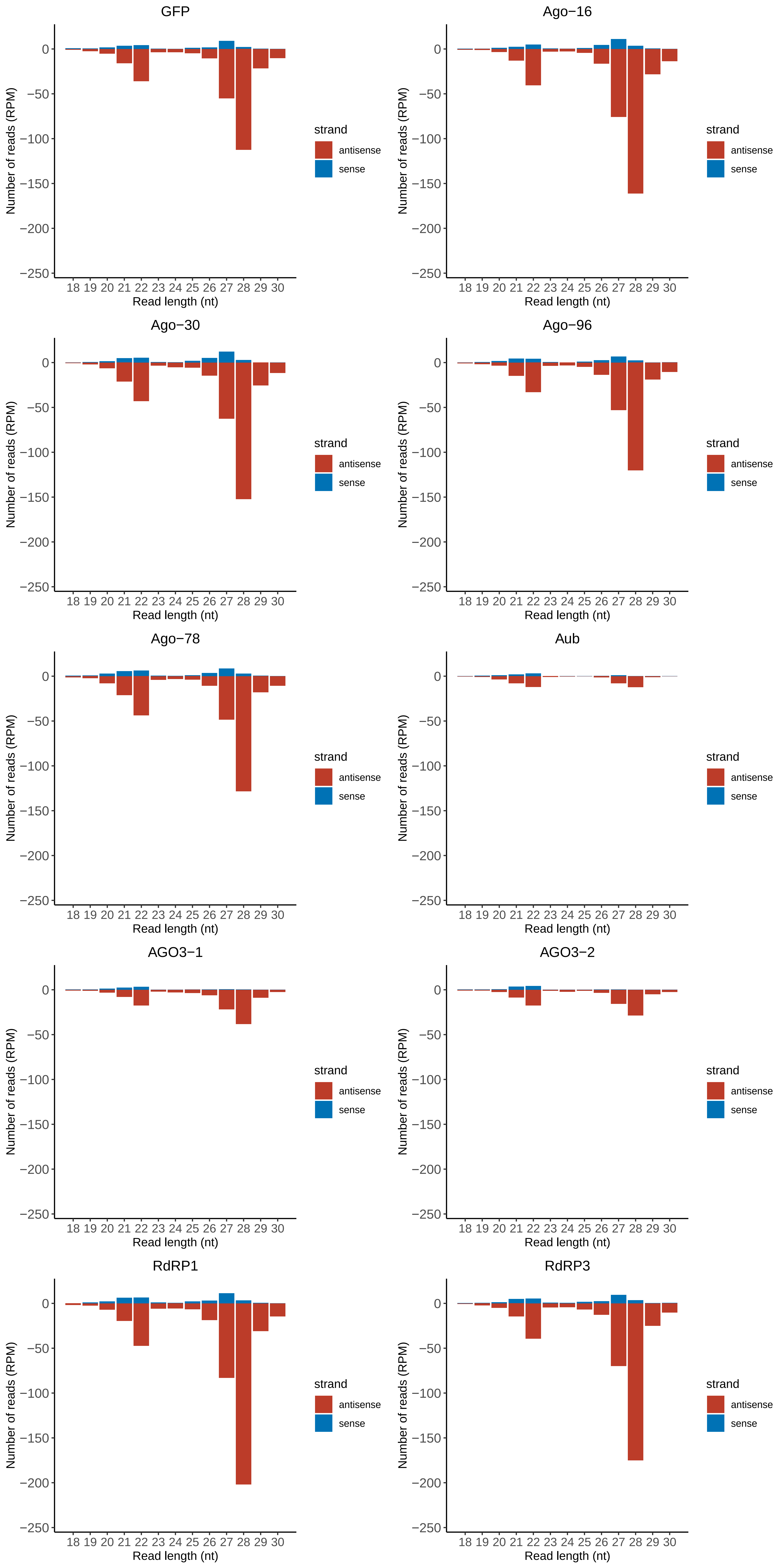

### Motif:rnd-6_family-4238TE_lengthdistribution.pdf

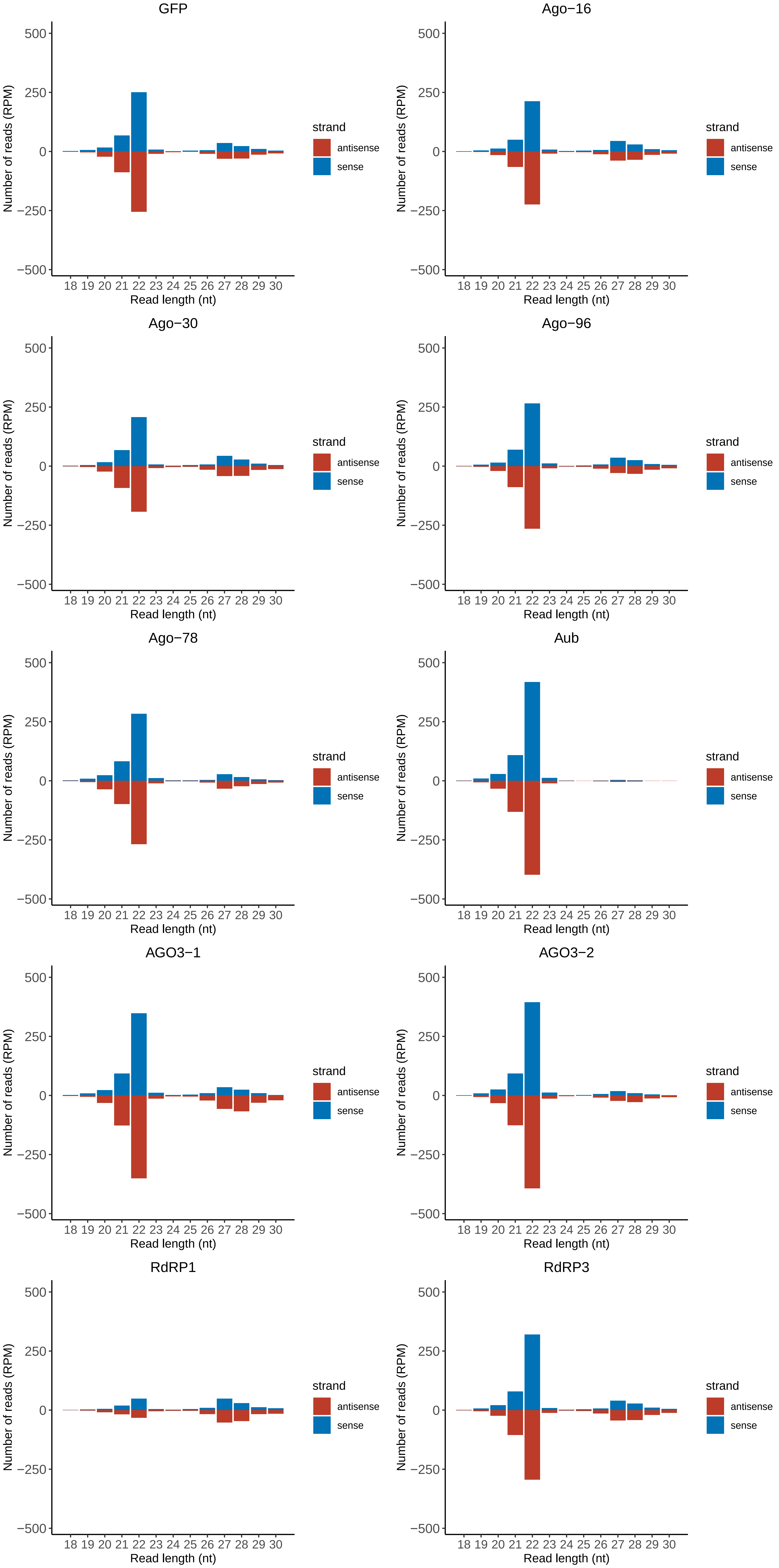

### Motif:rnd-6_family-4387TE_lengthdistribution.pdf

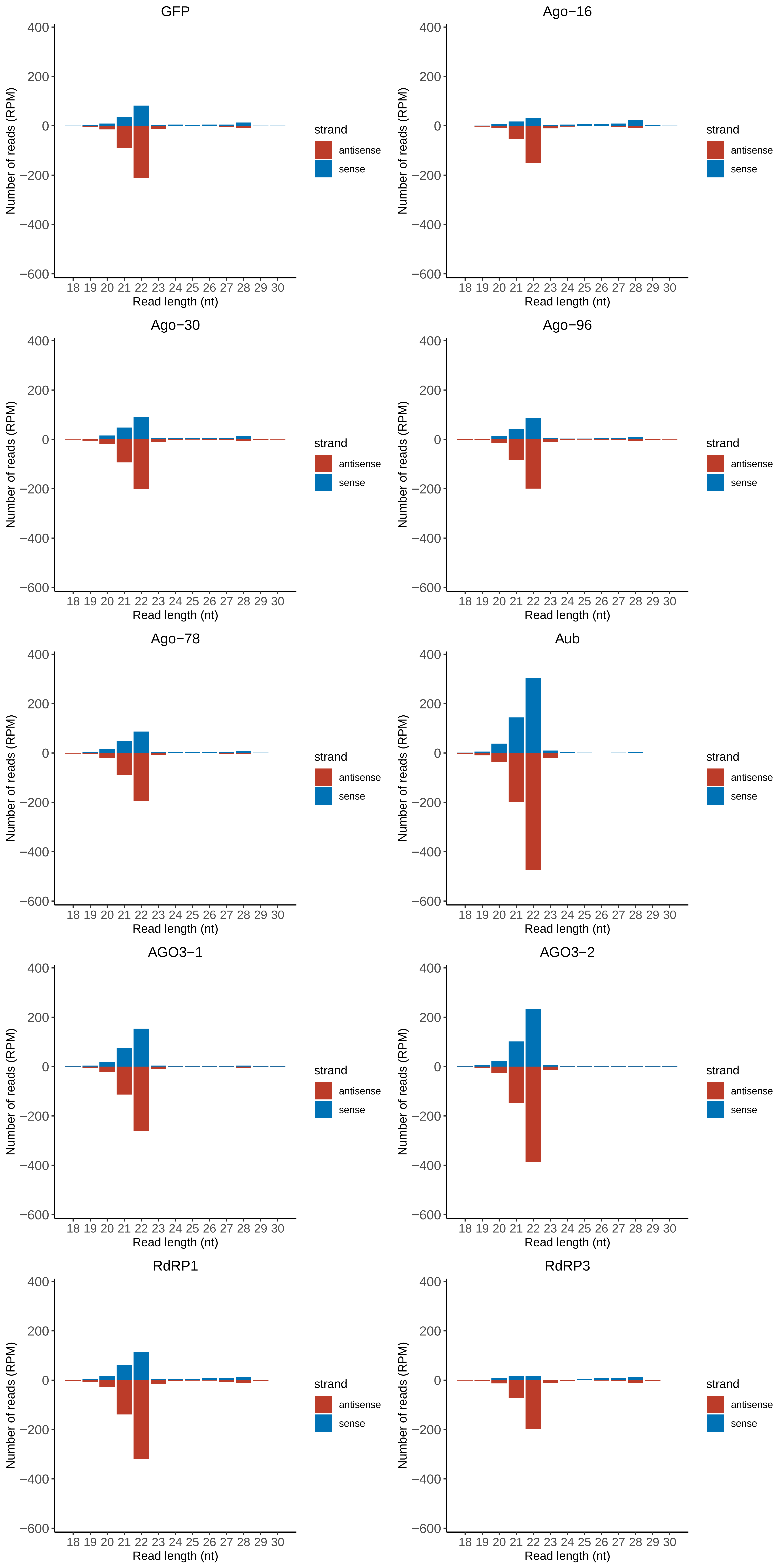

### Motif:rnd-6_family-5311TE_lengthdistribution.pdf

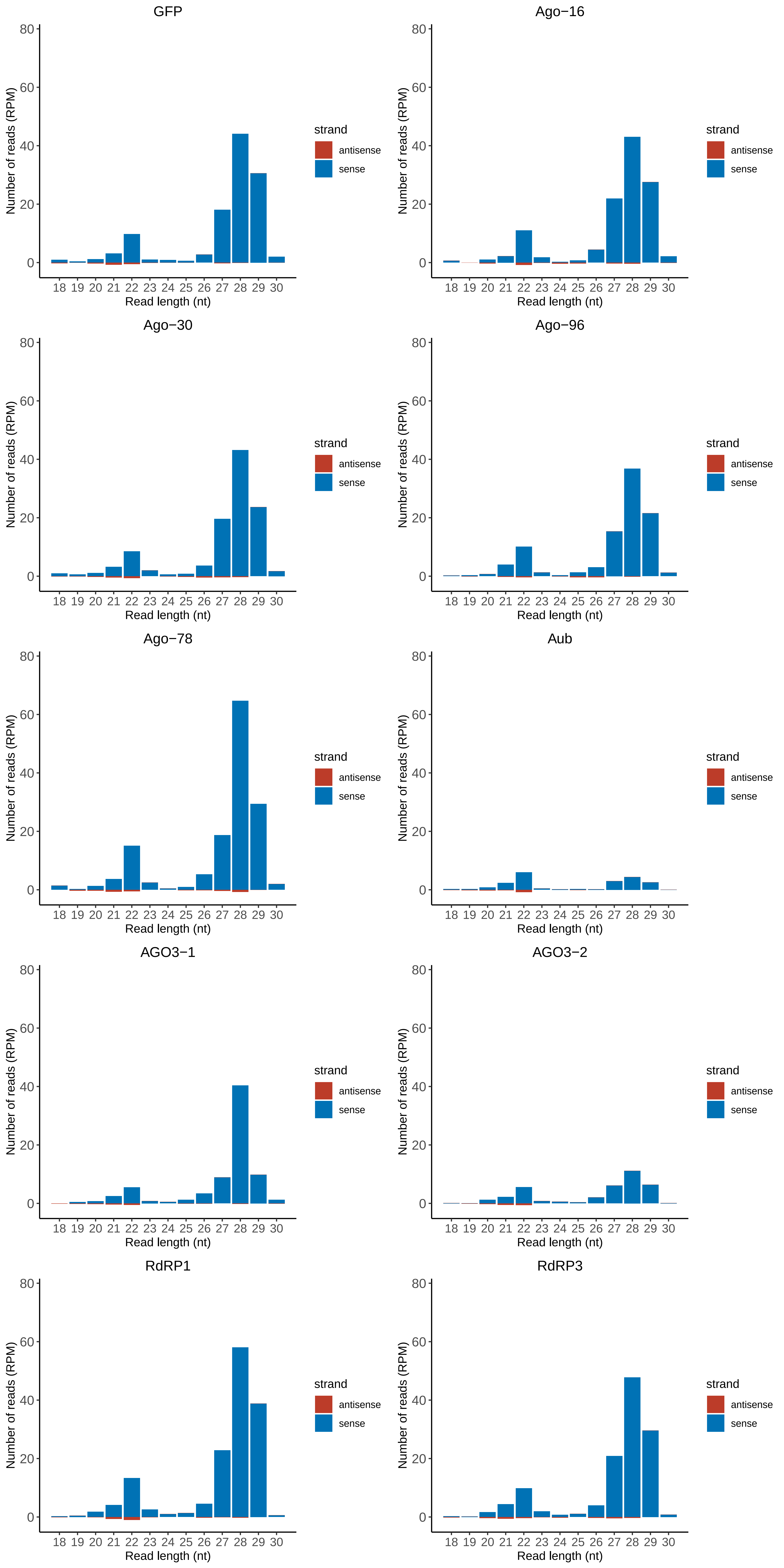

### Motif:rnd-6_family-6087TE_lengthdistribution.pdf

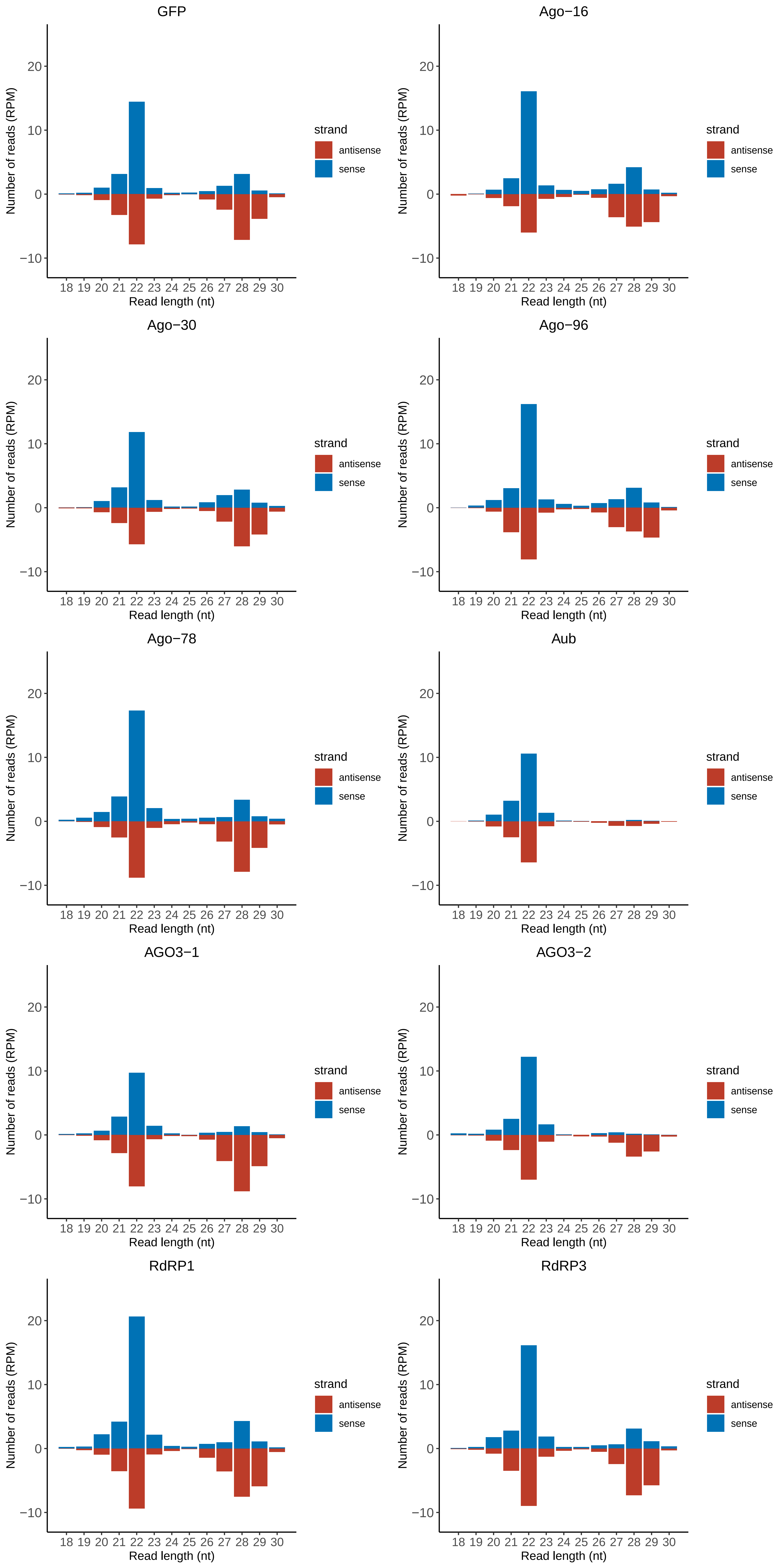

### Motif:rnd-6_family-7467TE_lengthdistribution.pdf

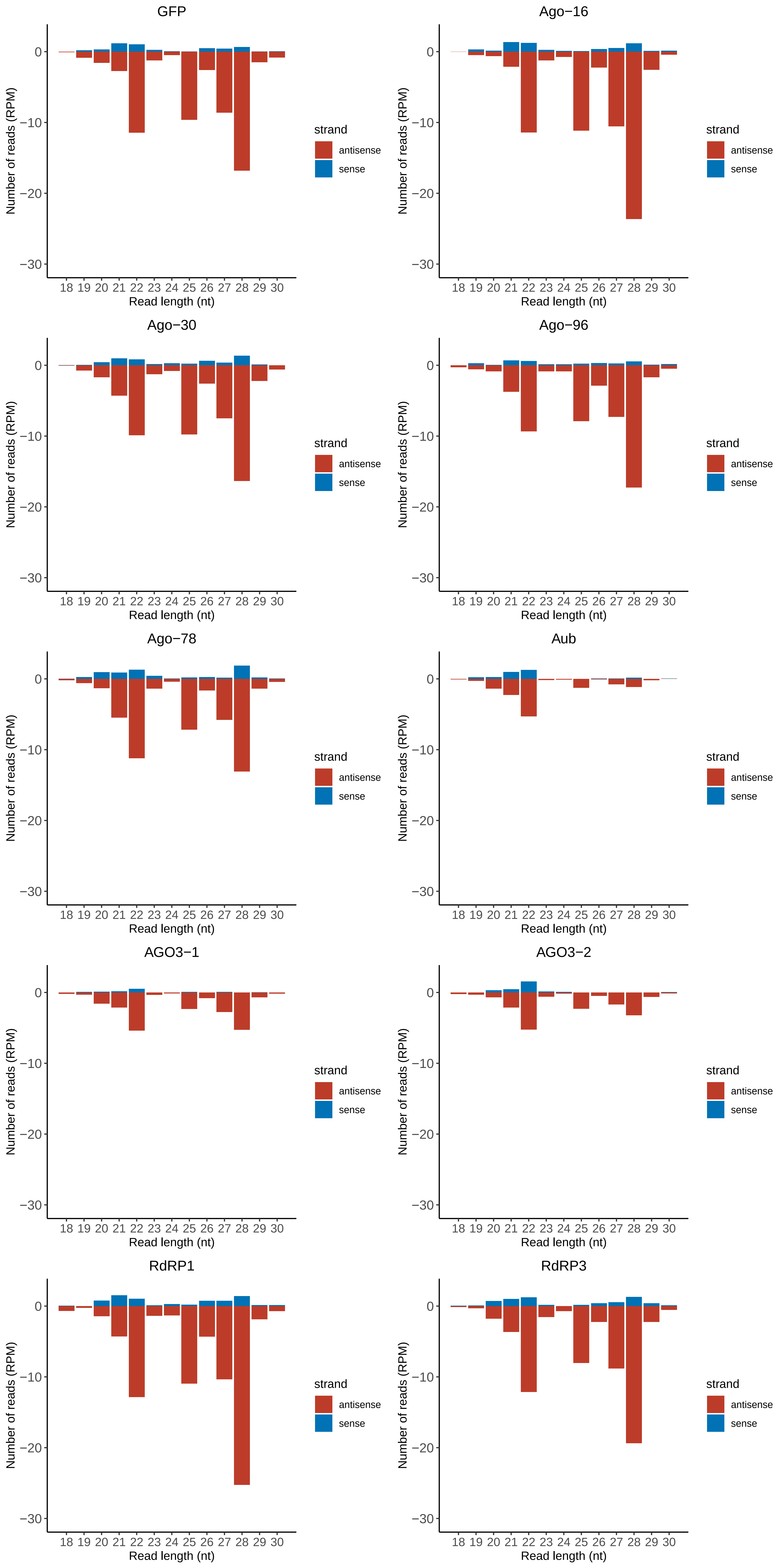

### Motif:rnd-6_family-7508TE_lengthdistribution.pdf

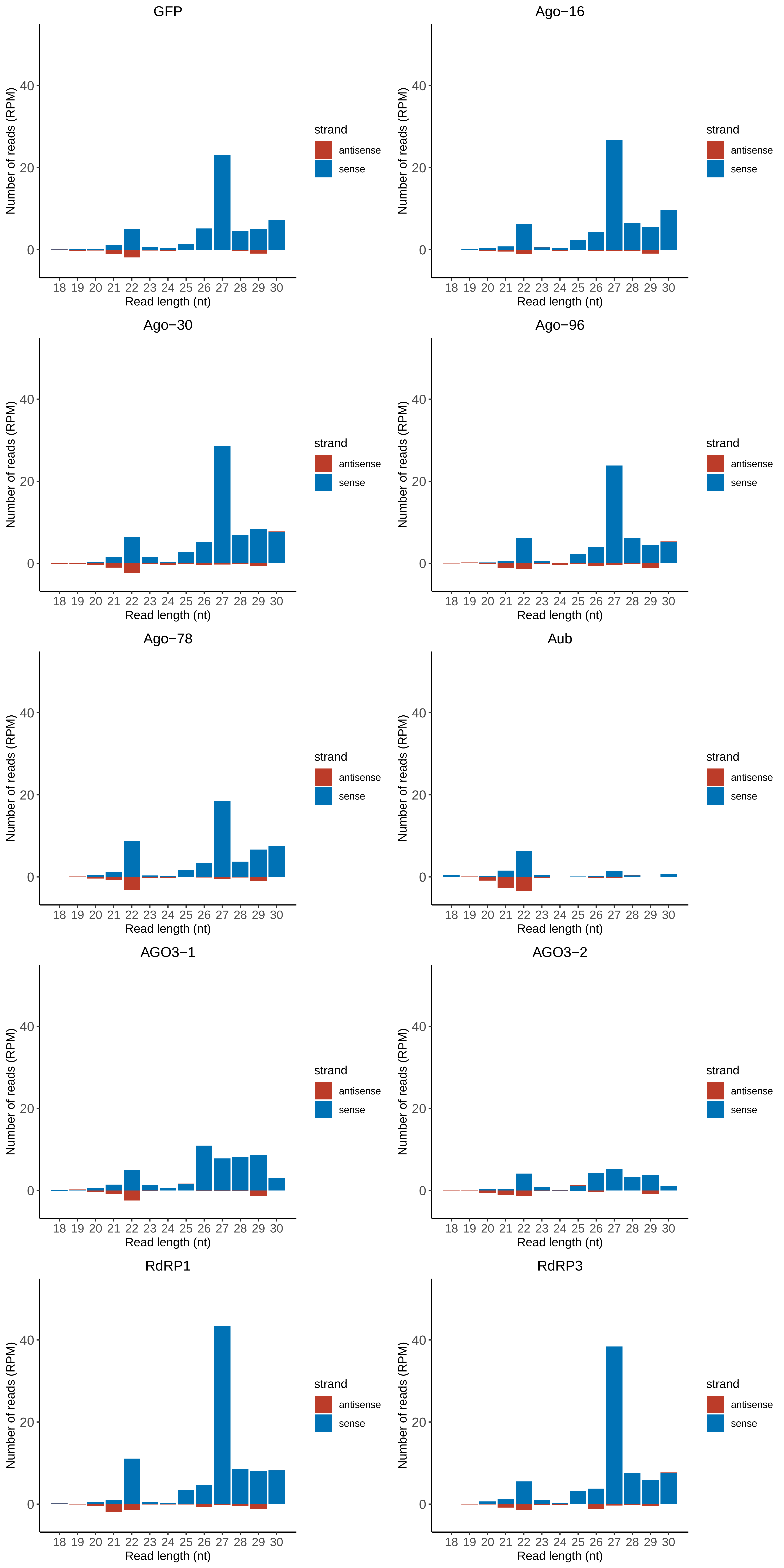

### Motif:rnd-6_family-7630TE_lengthdistribution.pdf

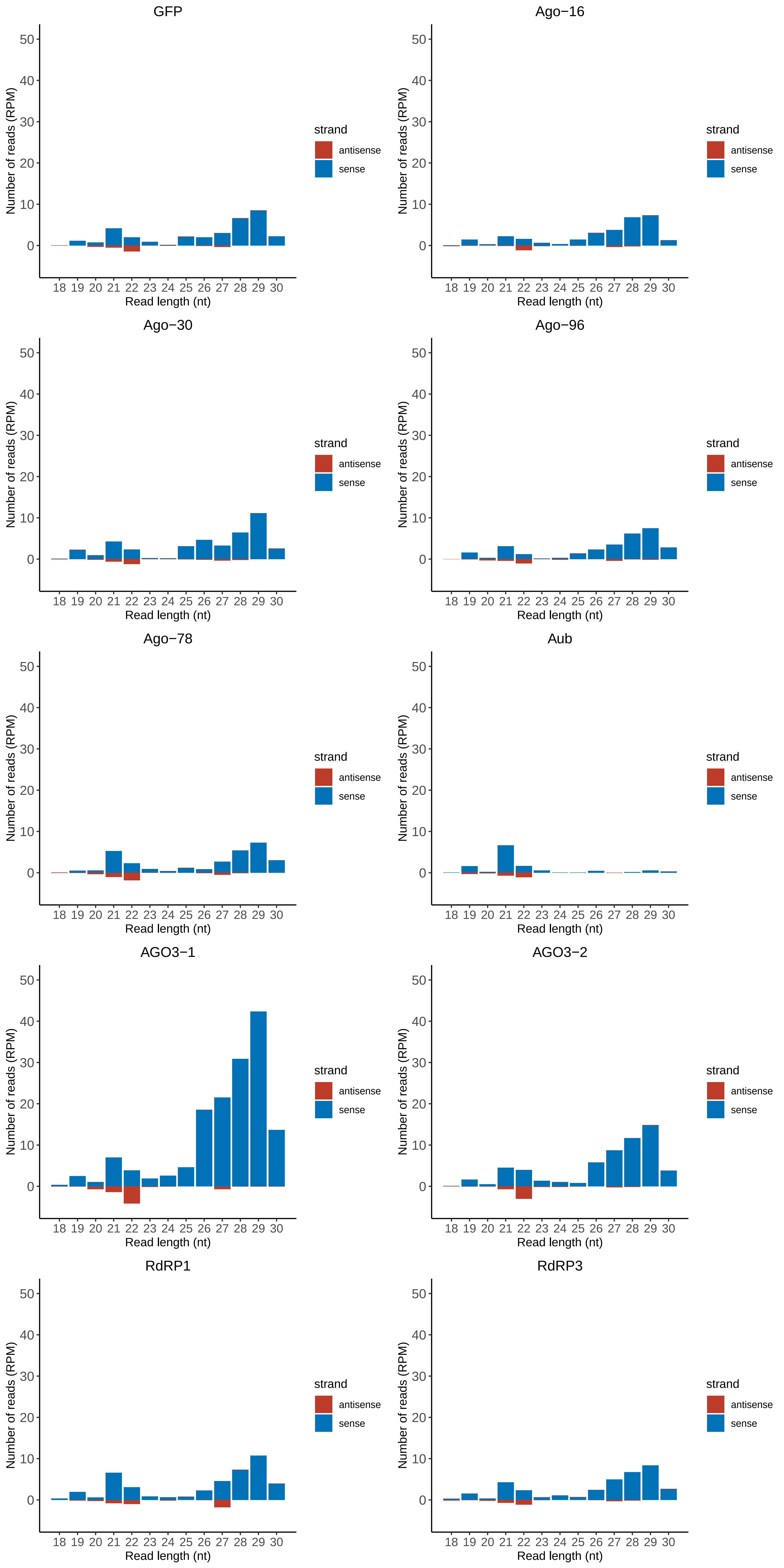

### Motif:rnd-6_family-7981TE_lengthdistribution.pdf

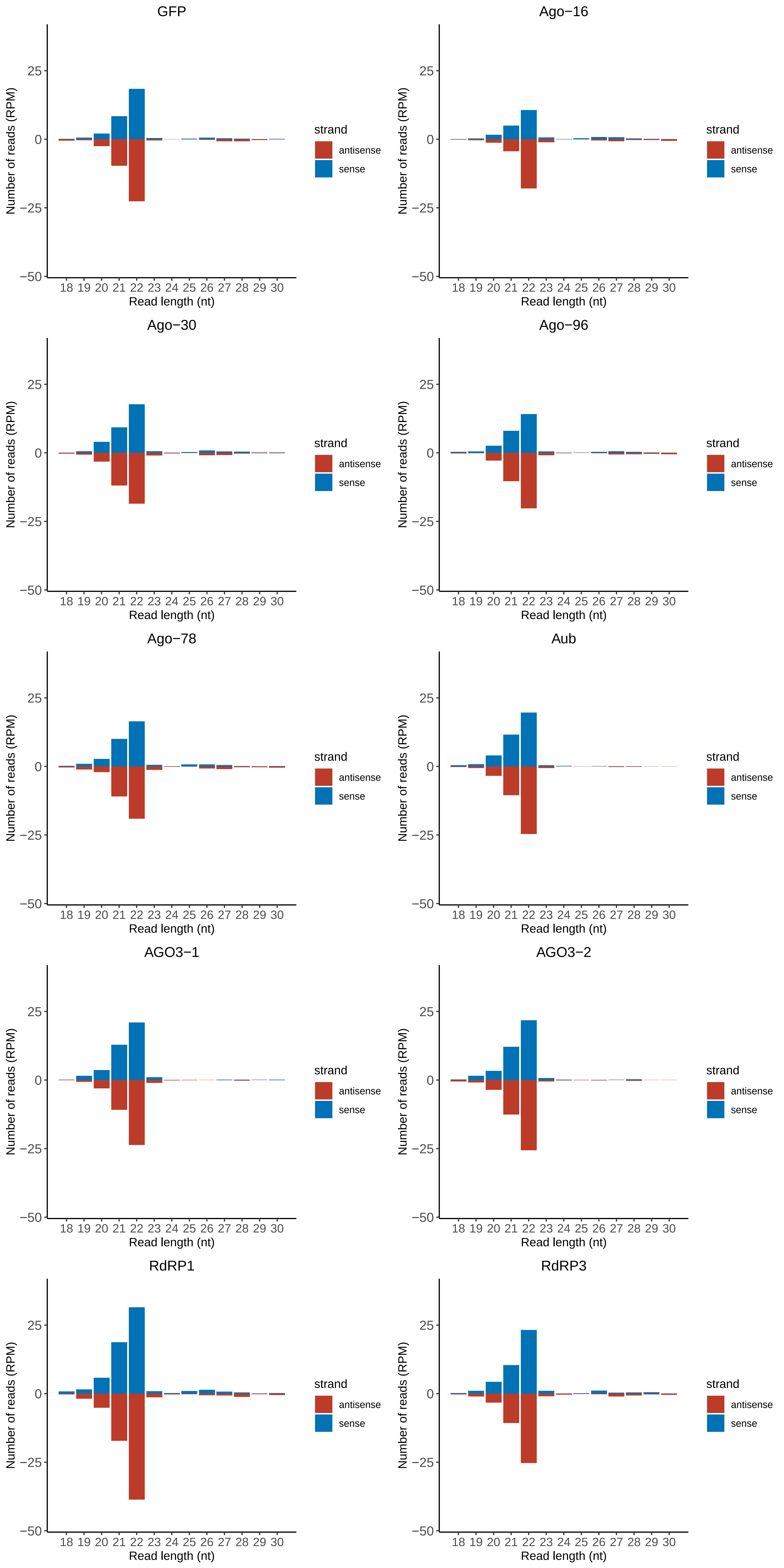

### Motif:rnd-6_family-9316TE_lengthdistribution.pdf

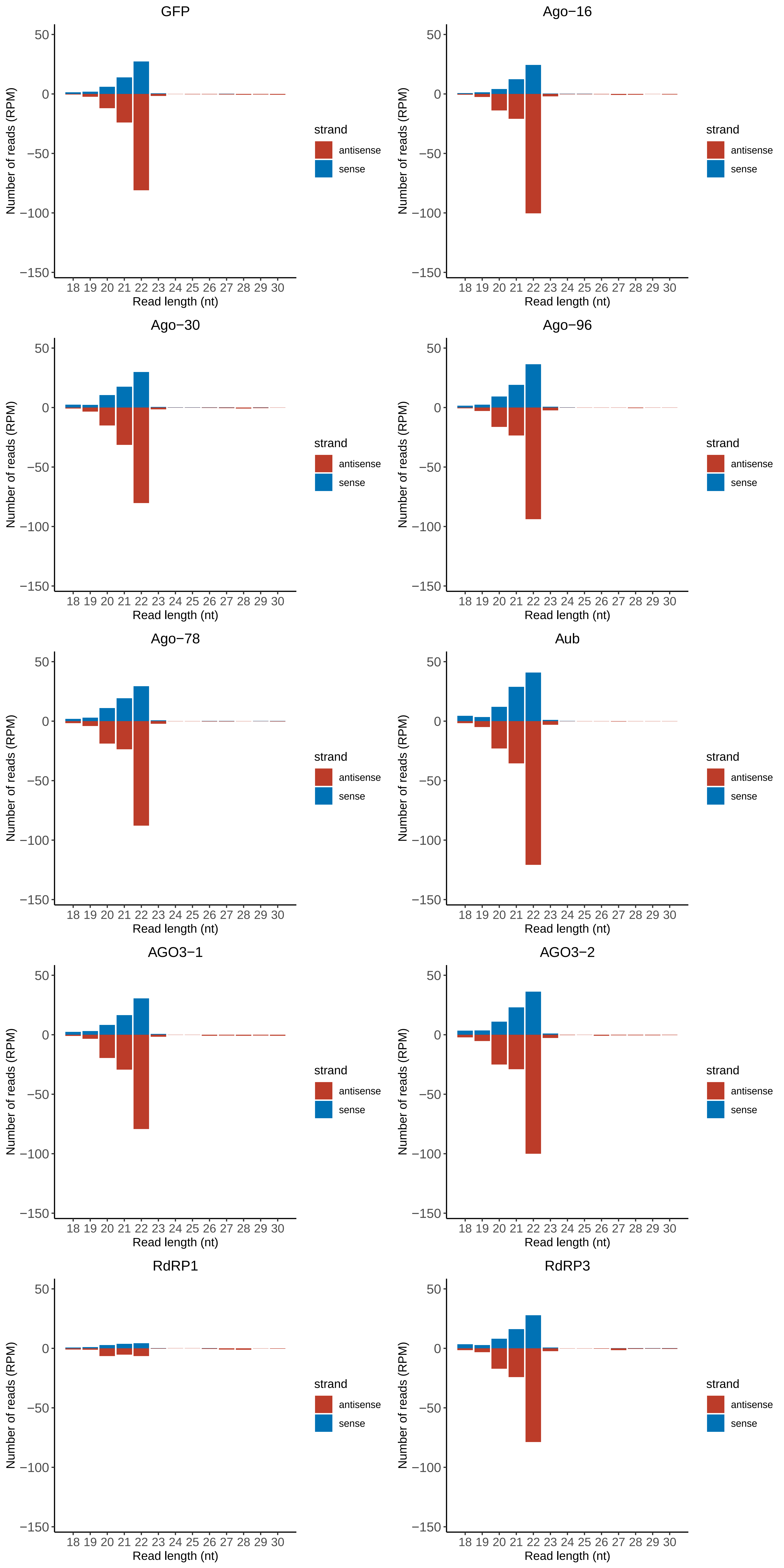

### Motif:rnd-6_family-10883TE_lengthdistribution.pdf

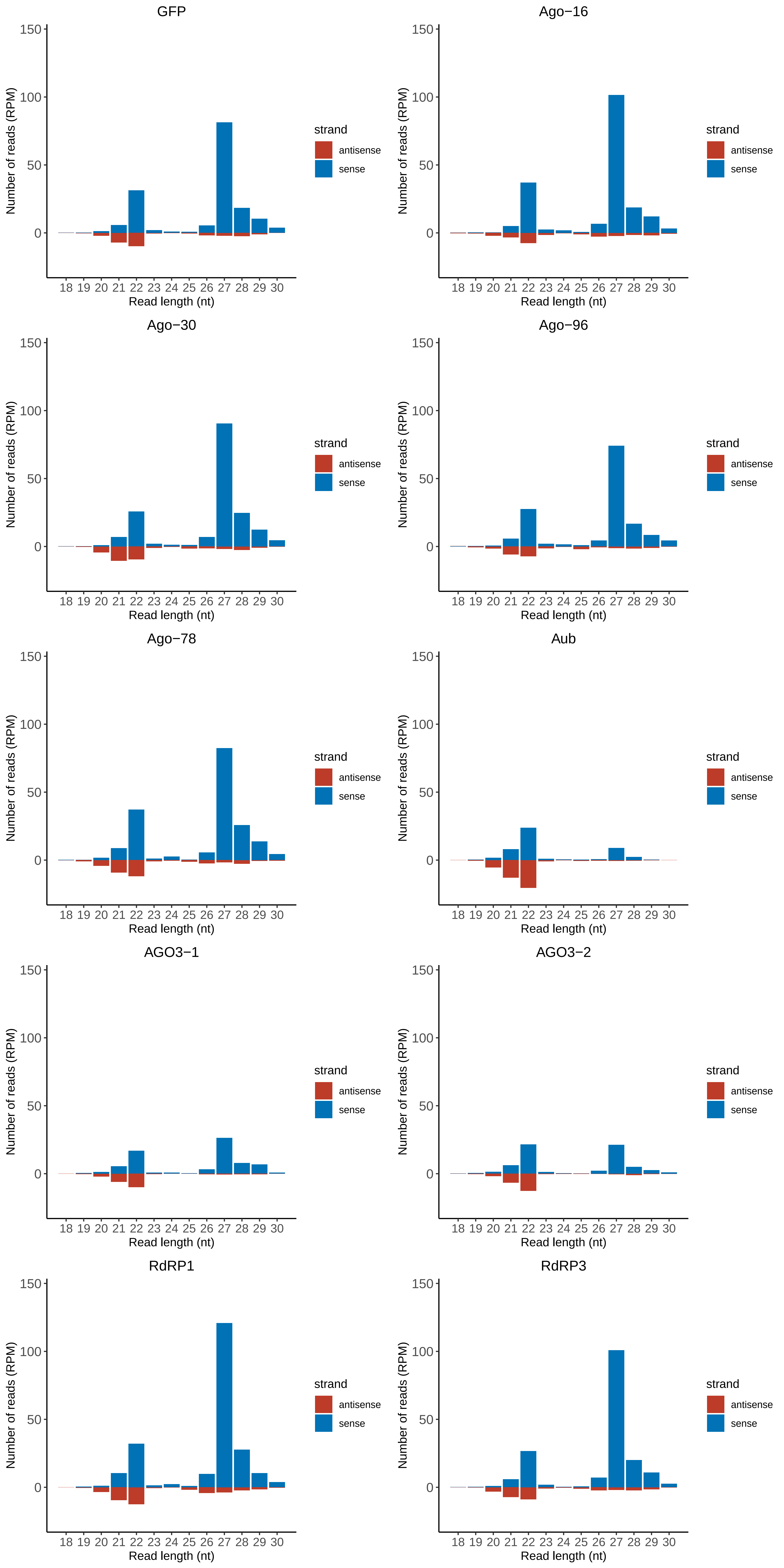

### Motif:rnd-6_family-12504TE_lengthdistribution.pdf

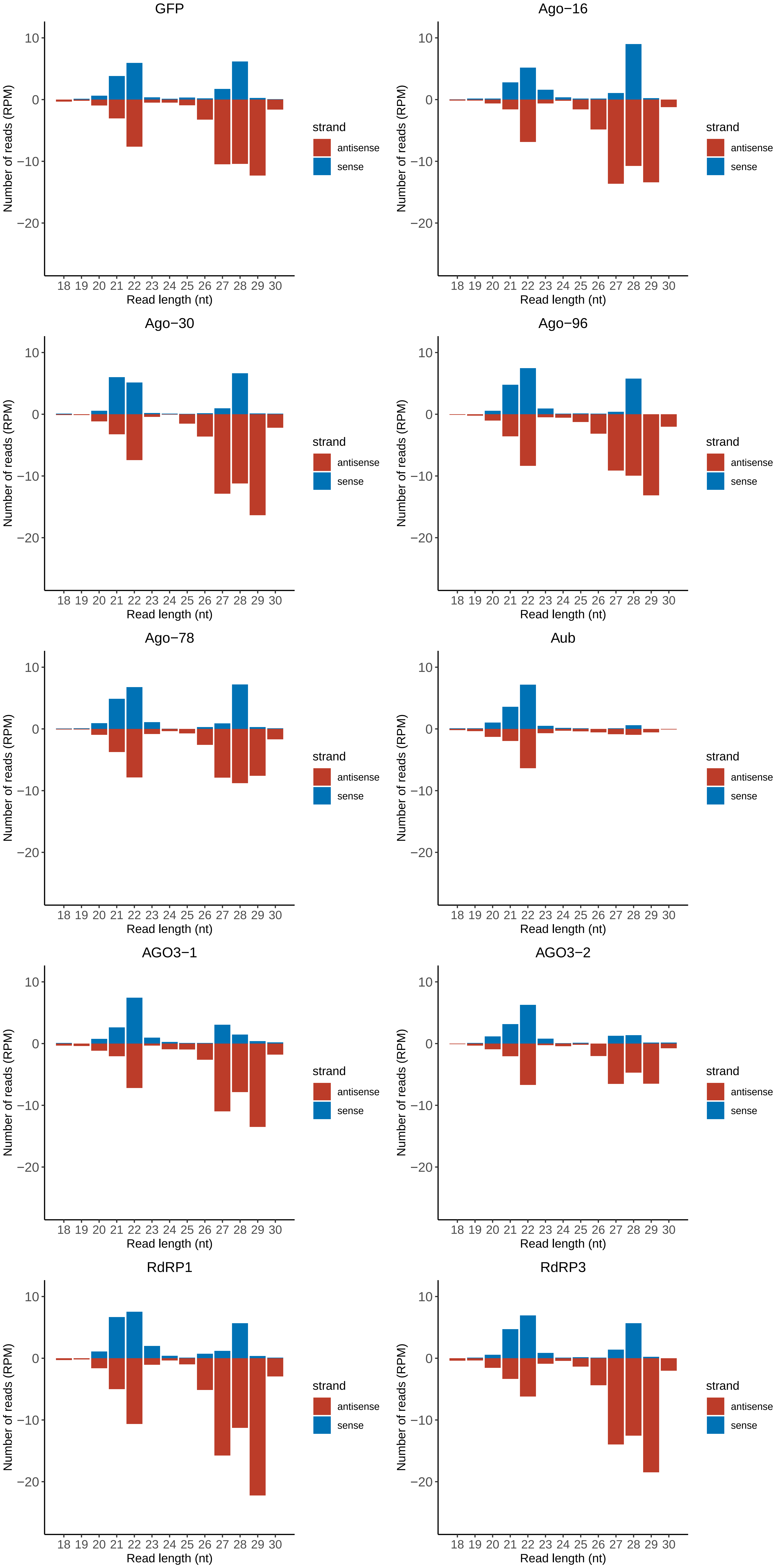
