## Supplementary figures and images for "A novel eukaryotic RdRP-dependent small RNA pathway represses antiviral immunity by controlling an ERK pathway component in the black-legged tick"

### ISCI003367.RACDS_lengthdistribution.pdf

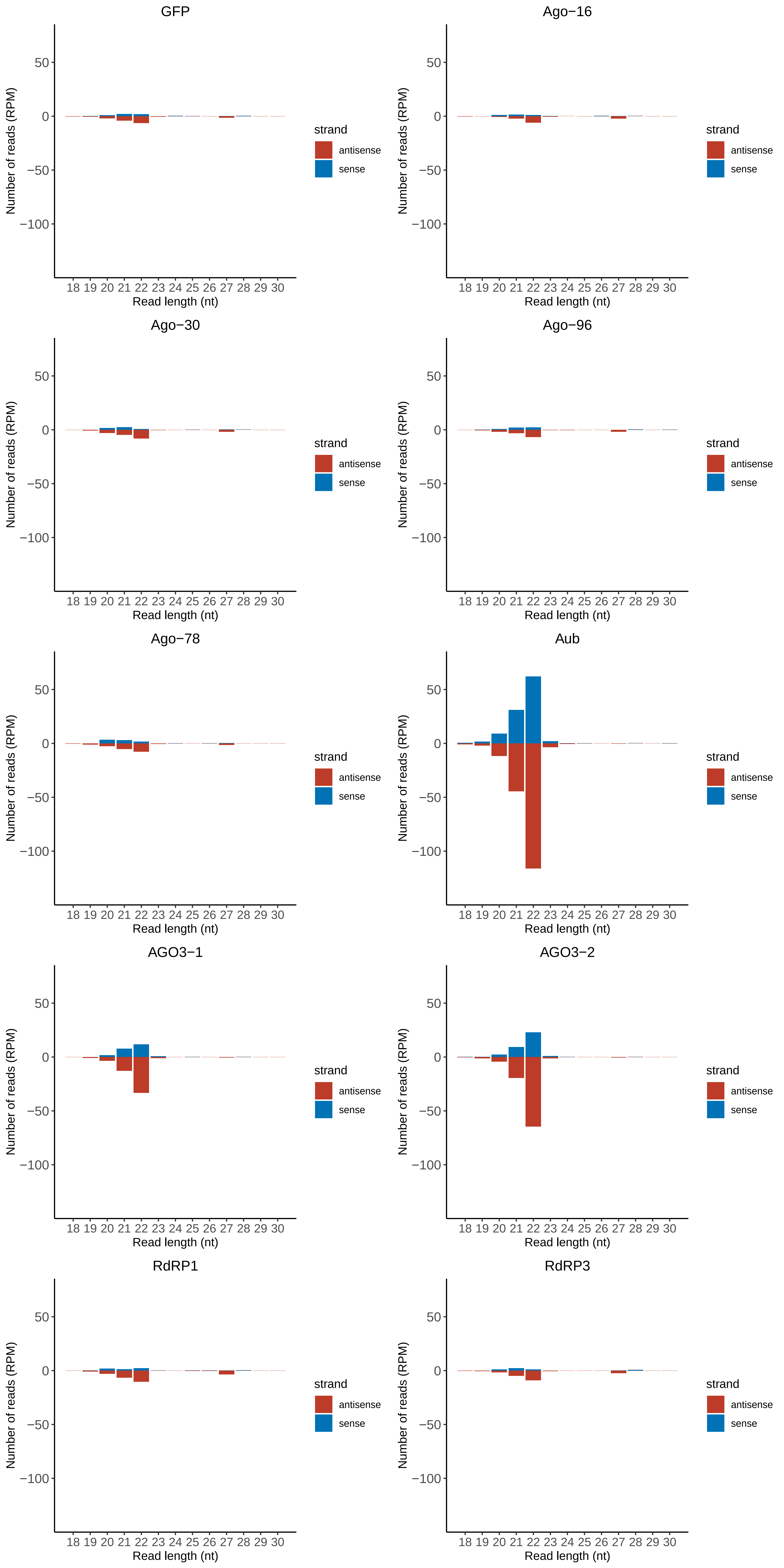

### ISCI004588.RACDS_lengthdistribution.pdf

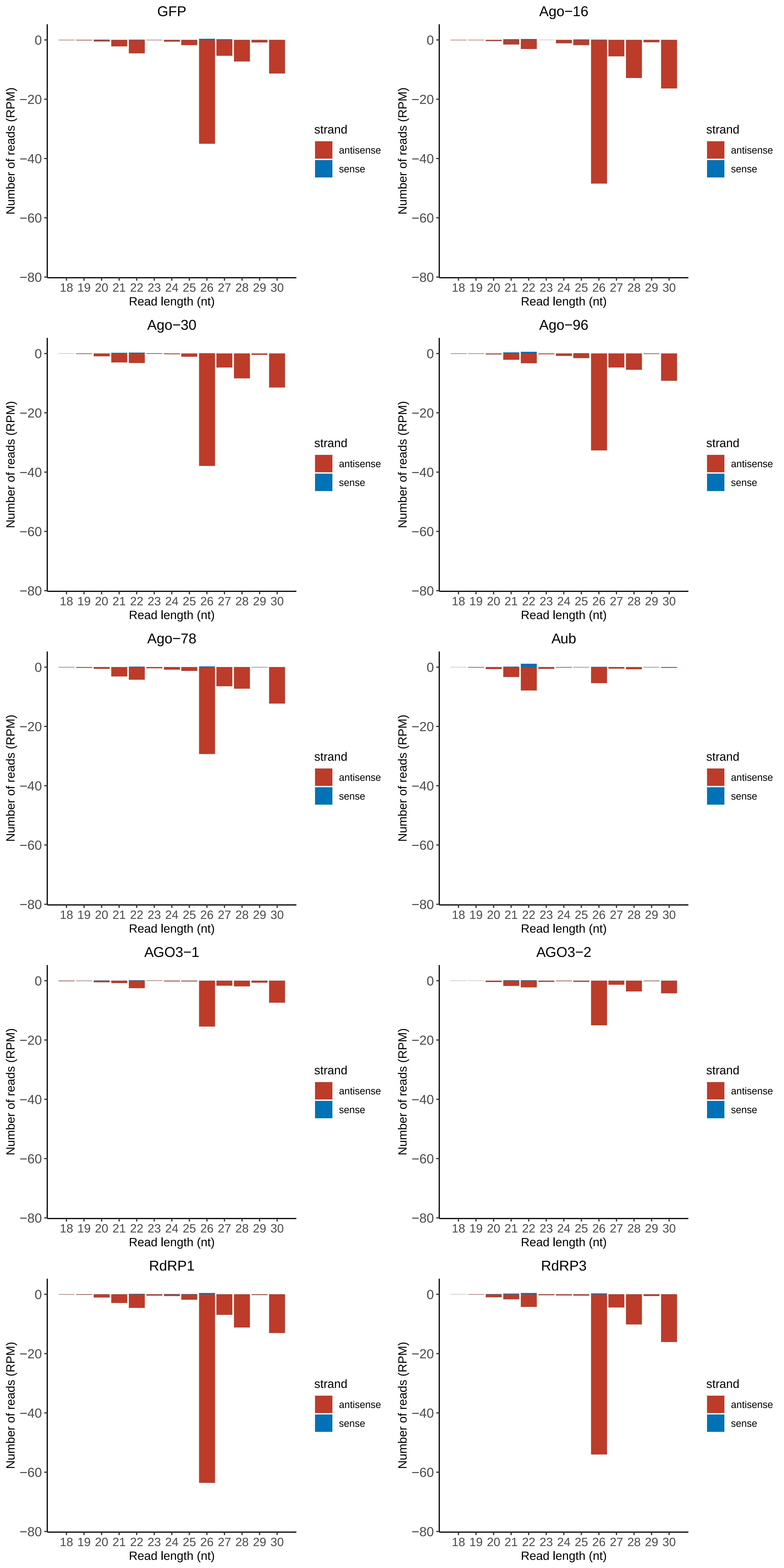

### ISCI005507.RACDS_lengthdistribution.pdf

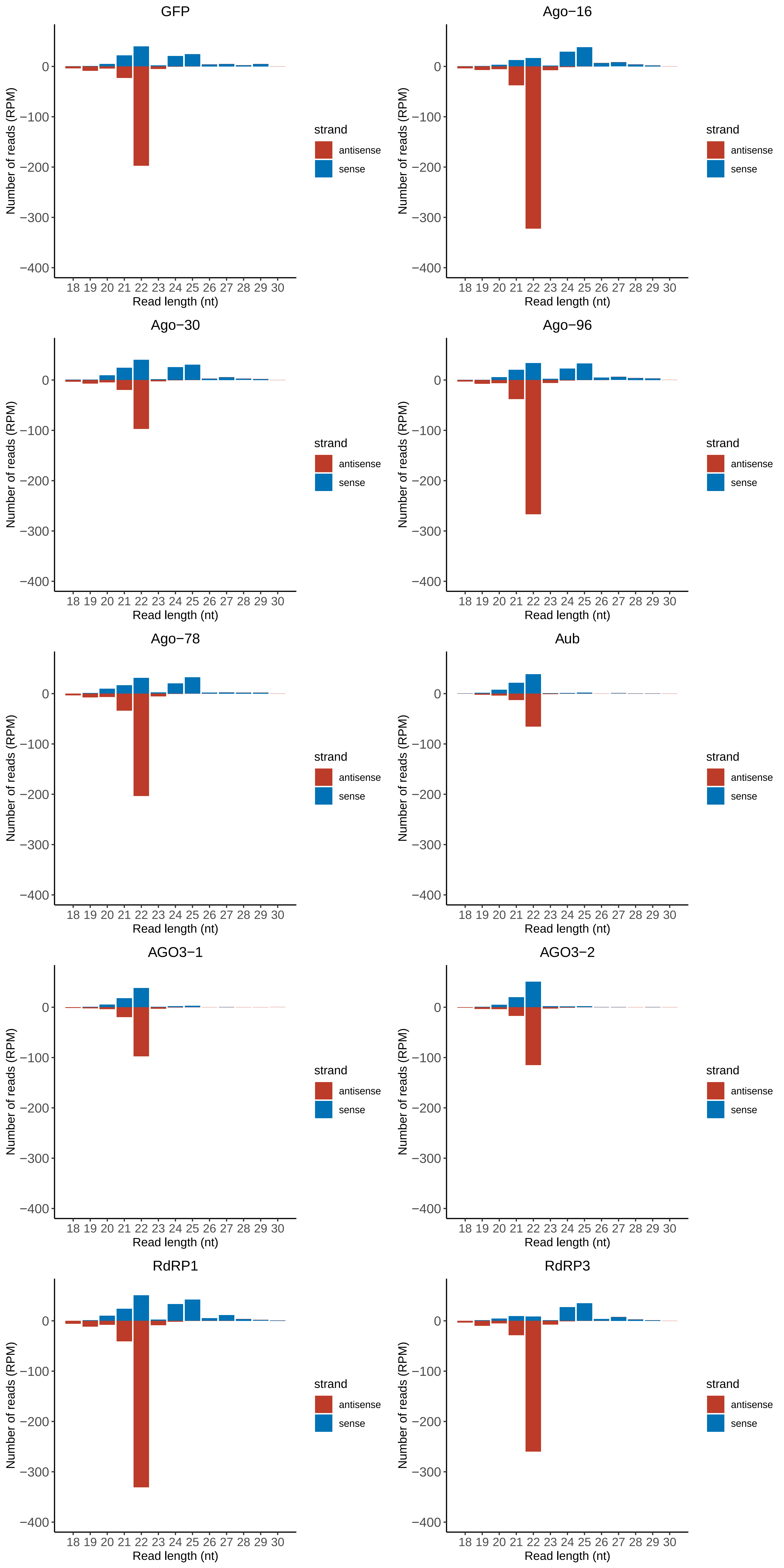

### ISCI006668.RACDS_lengthdistribution.pdf

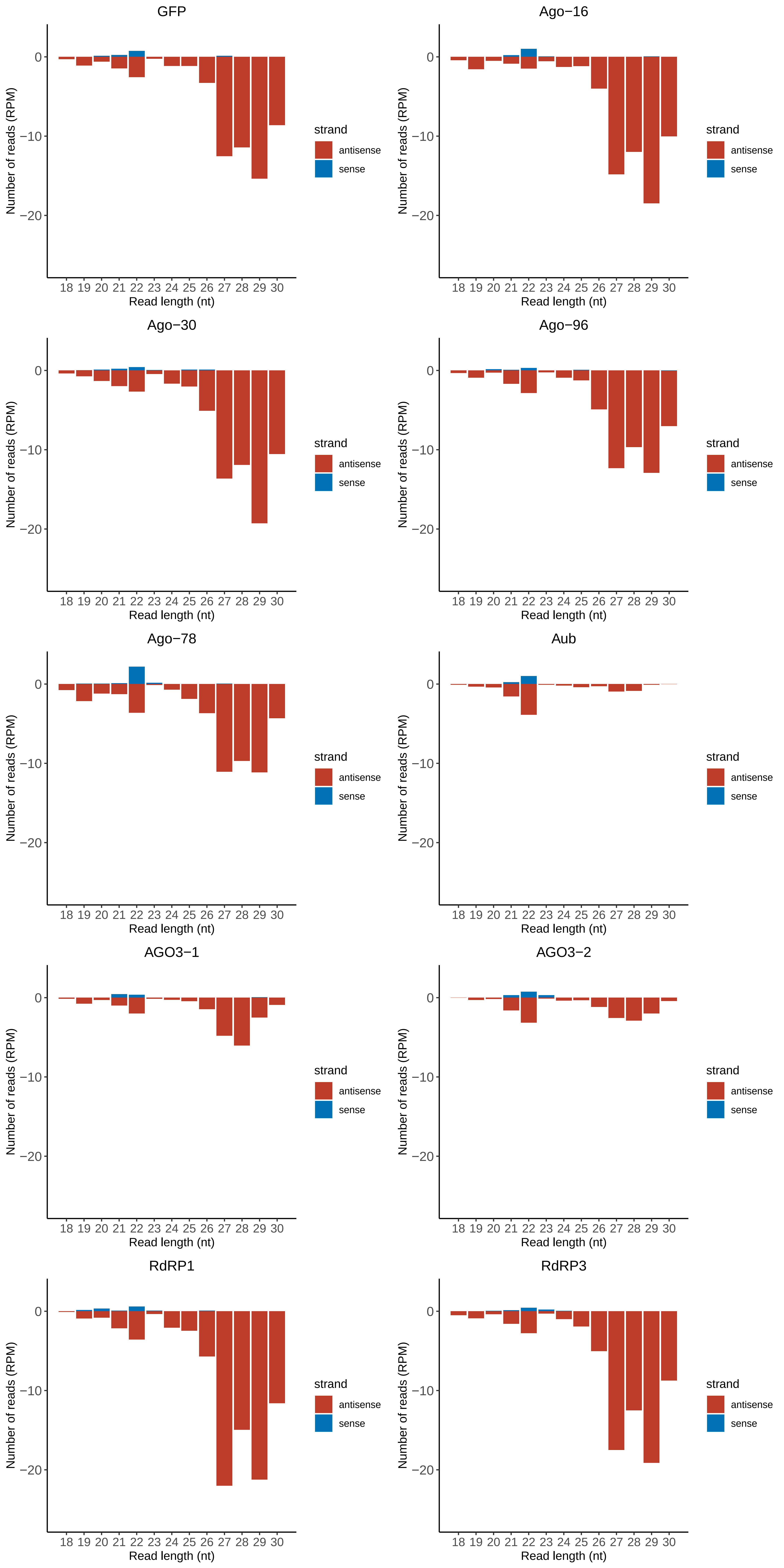

### ISCI007332.RACDS_lengthdistribution.pdf

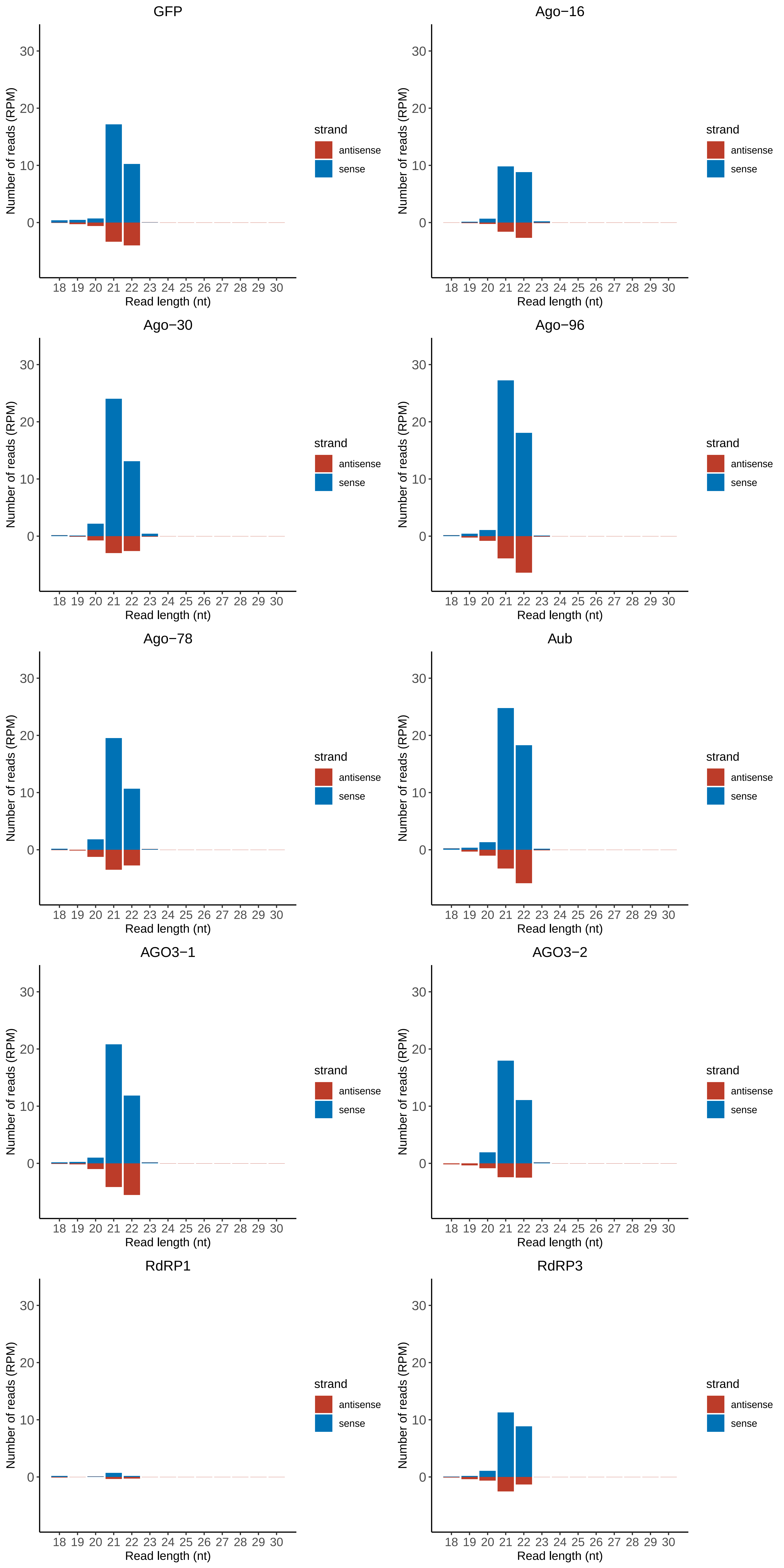

### ISCI010315.RACDS_lengthdistribution.pdf

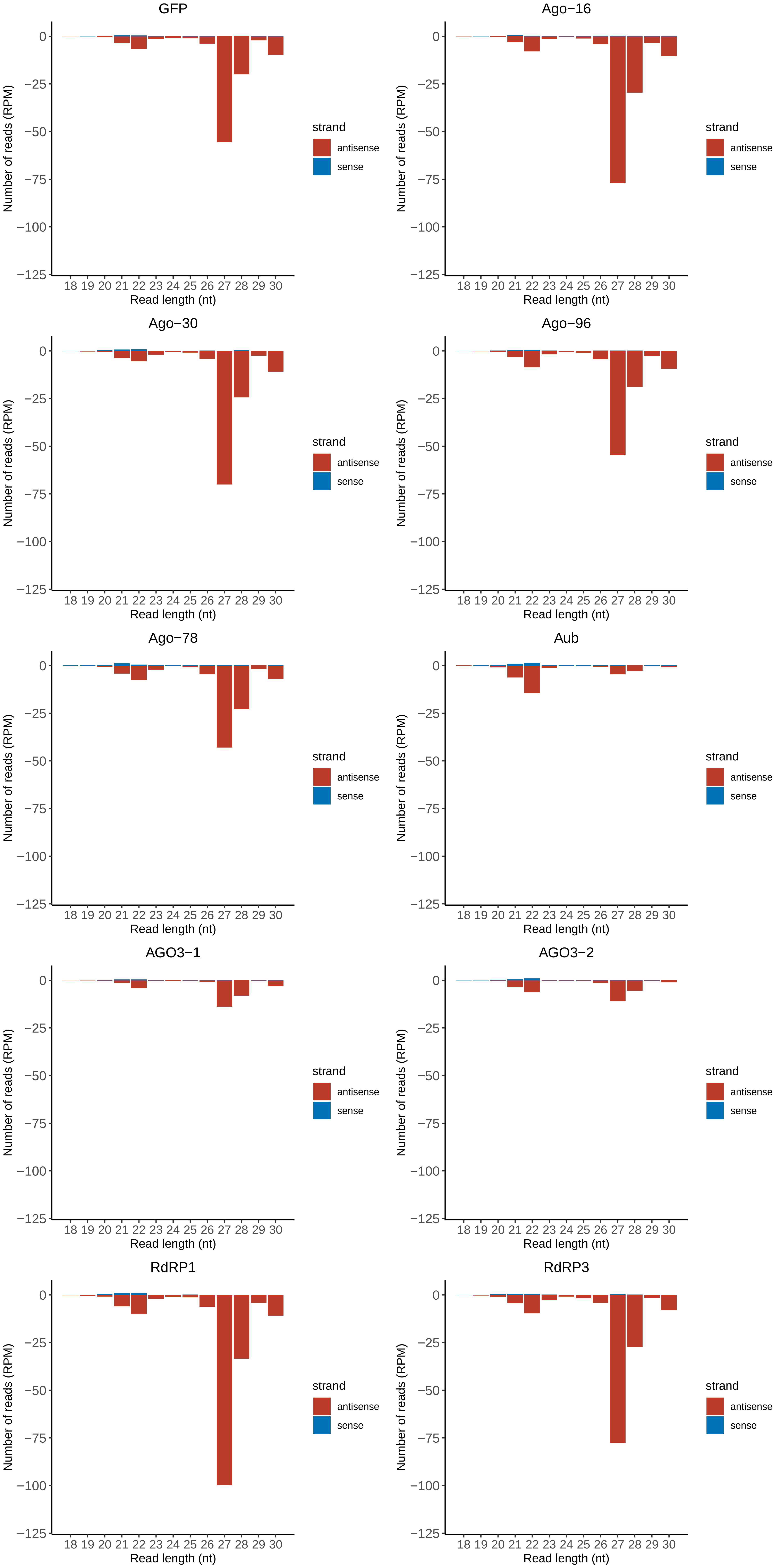

### ISCI018019.RACDS_lengthdistribution.pdf

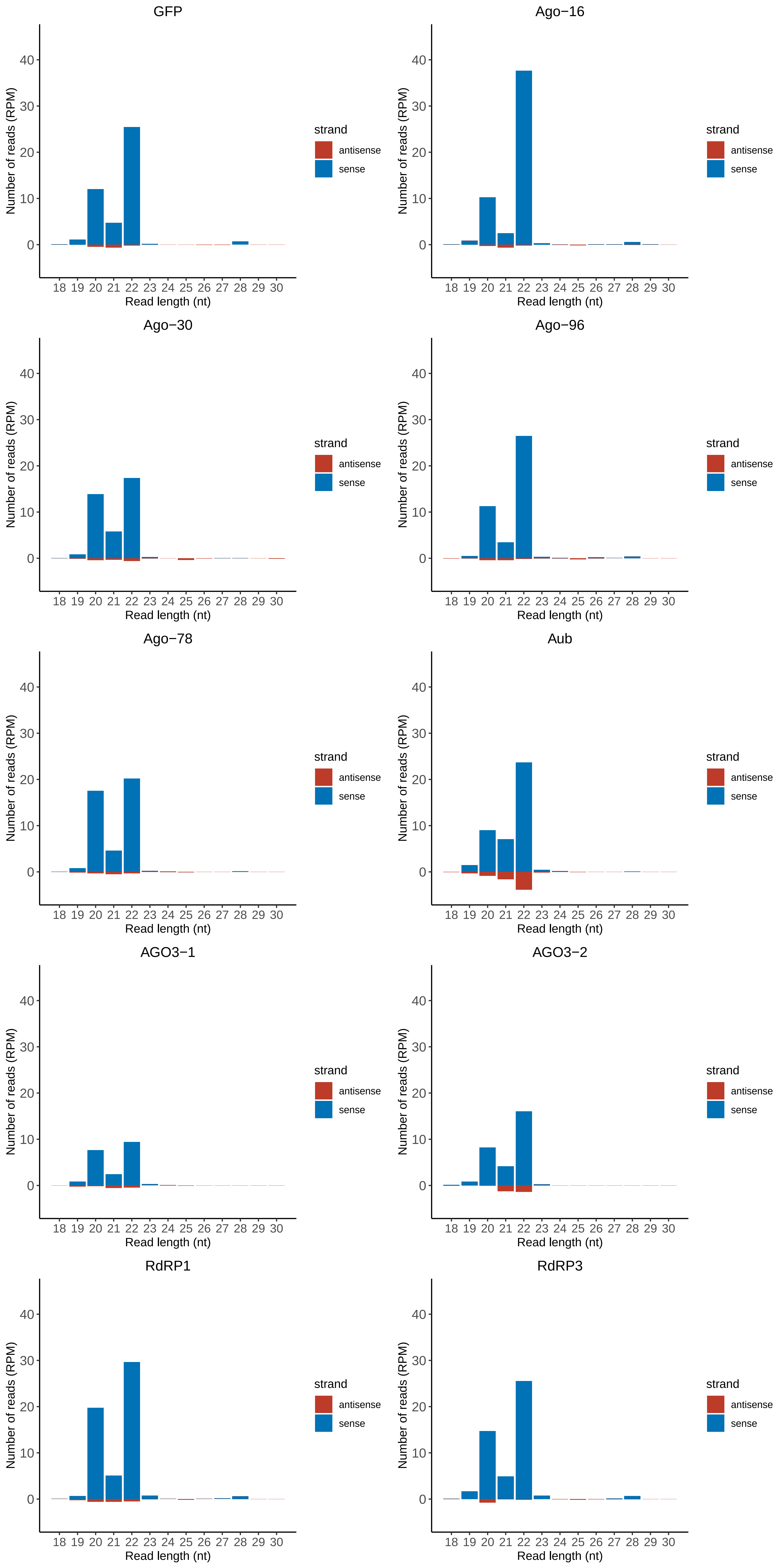

### ISCI024649.RACDS_lengthdistribution.pdf

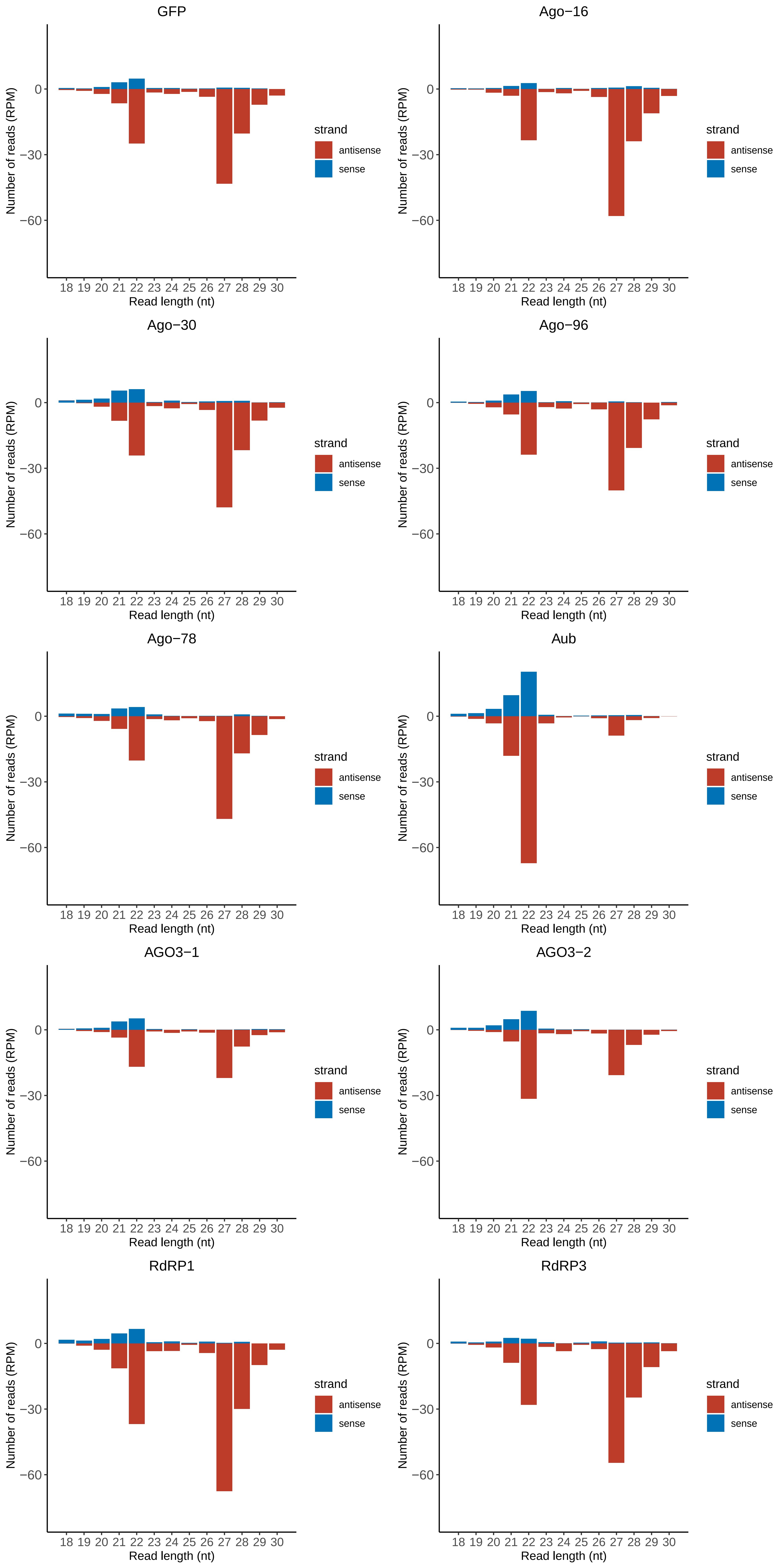

### Motif:rnd-1_family-9TE_lengthdistribution.pdf

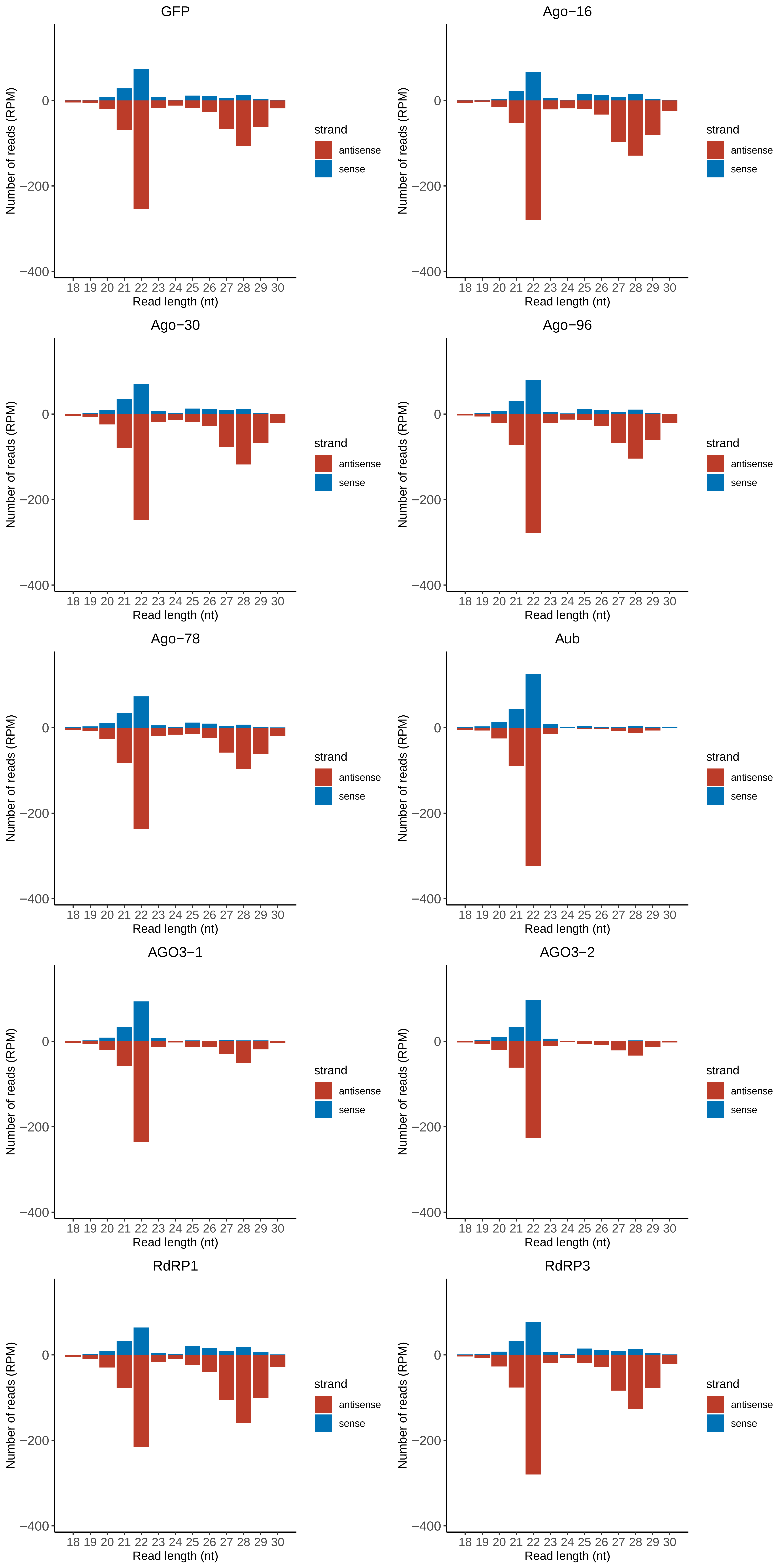

### Motif:rnd-1_family-12TE_lengthdistribution.pdf

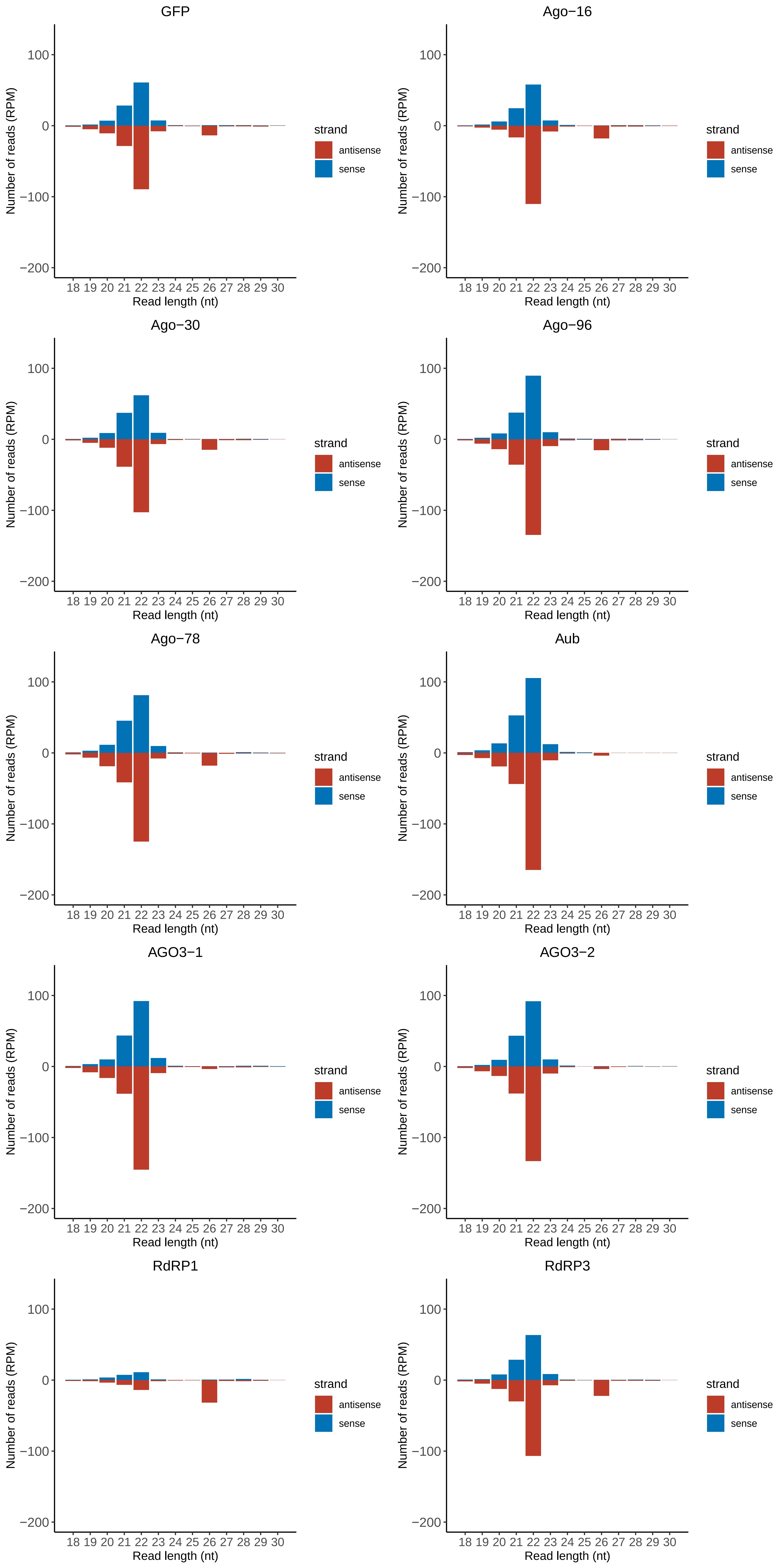

### Motif:rnd-1_family-16TE_lengthdistribution.pdf

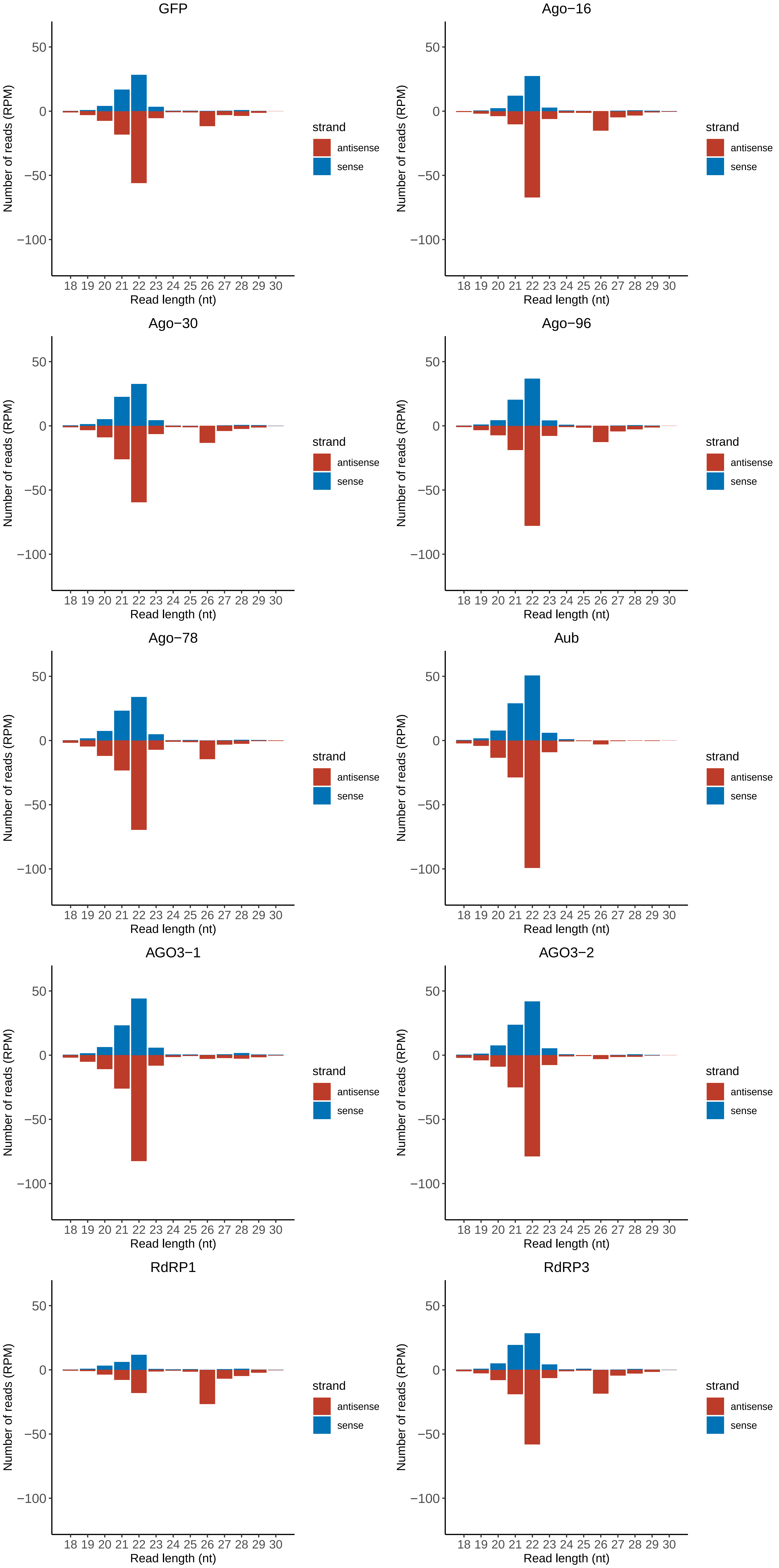

### Motif:rnd-1_family-51TE_lengthdistribution.pdf

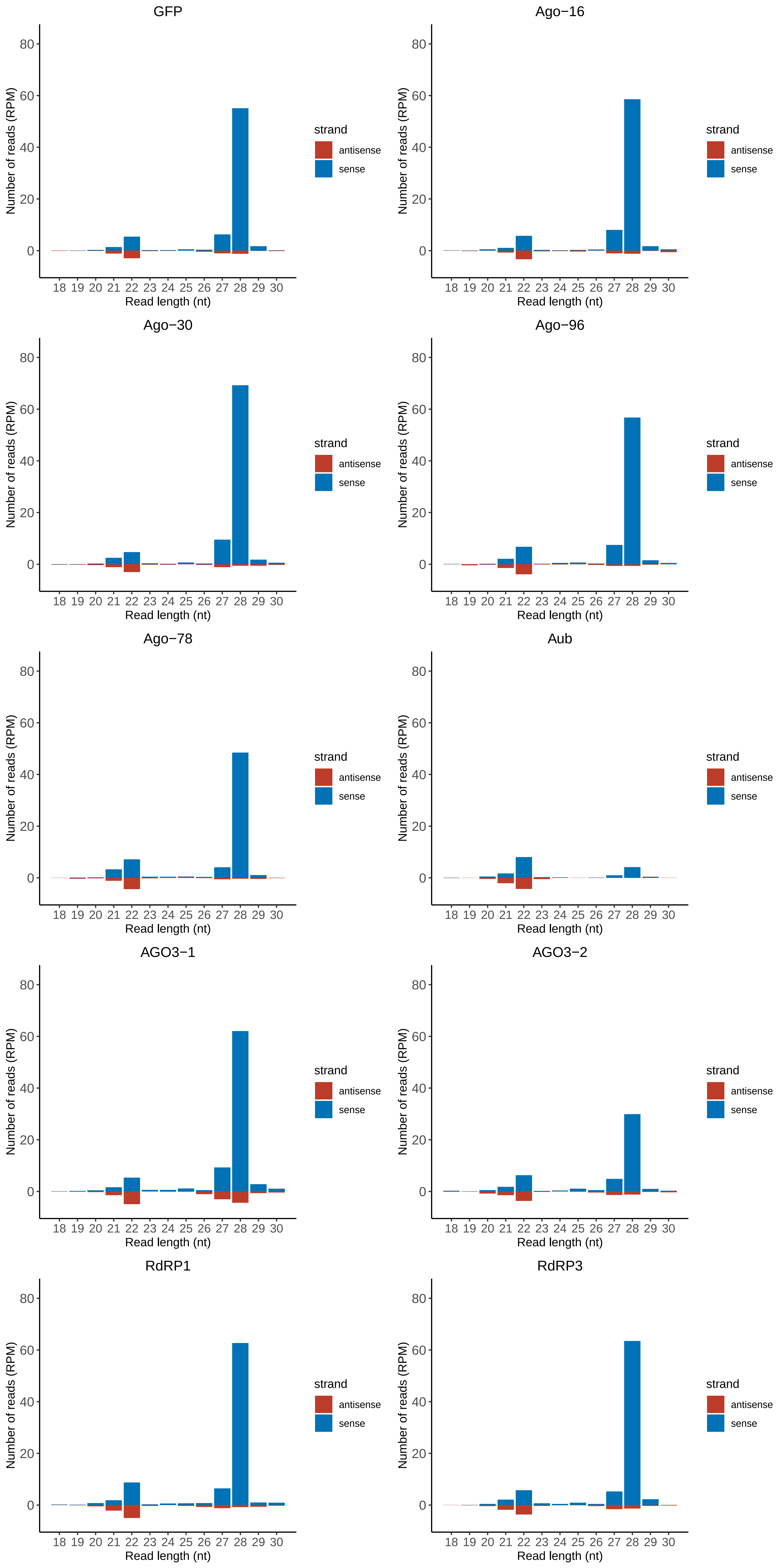

### Motif:rnd-1_family-96TE_lengthdistribution.pdf

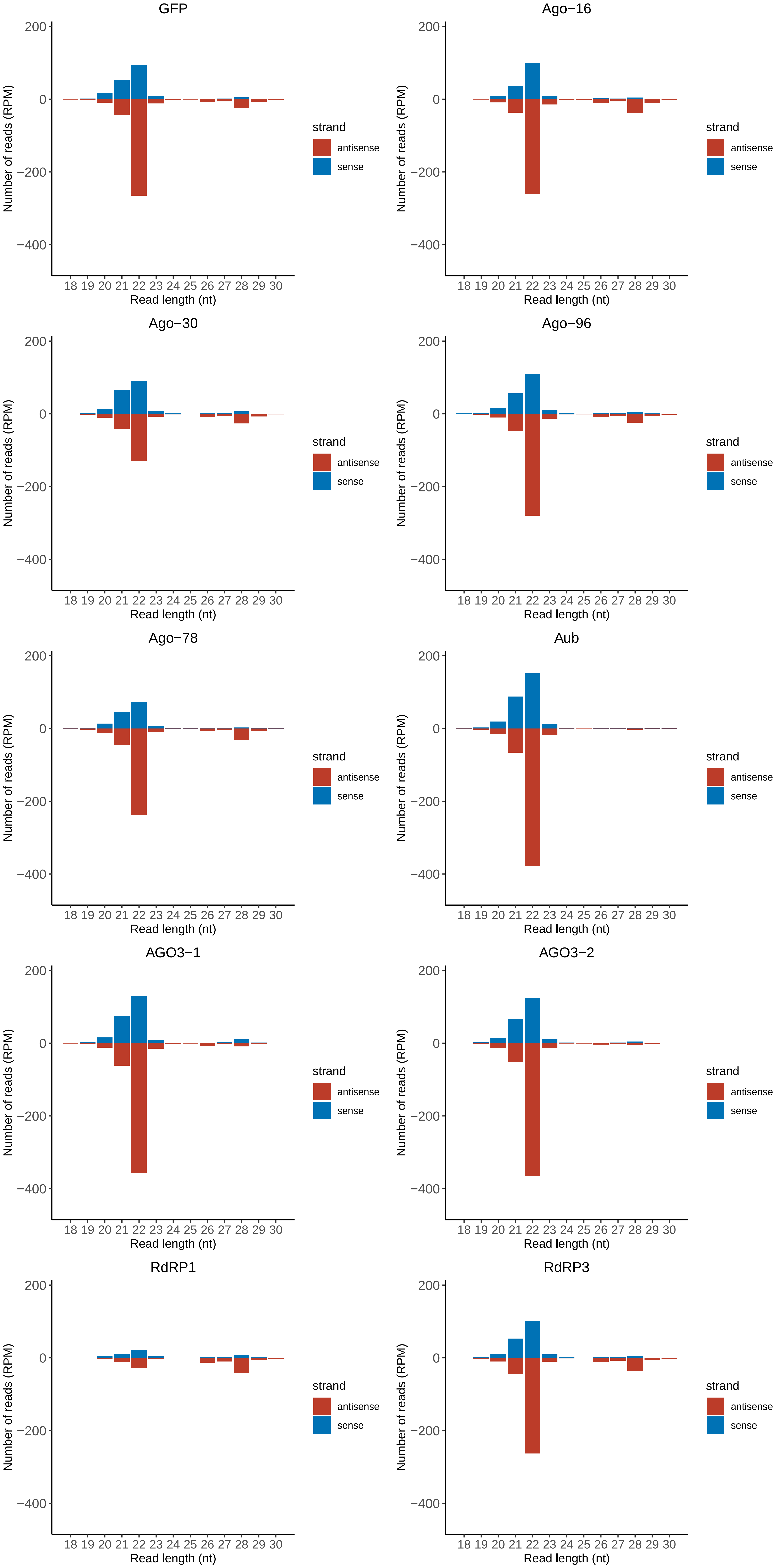

### Motif:rnd-1_family-122TE_lengthdistribution.pdf

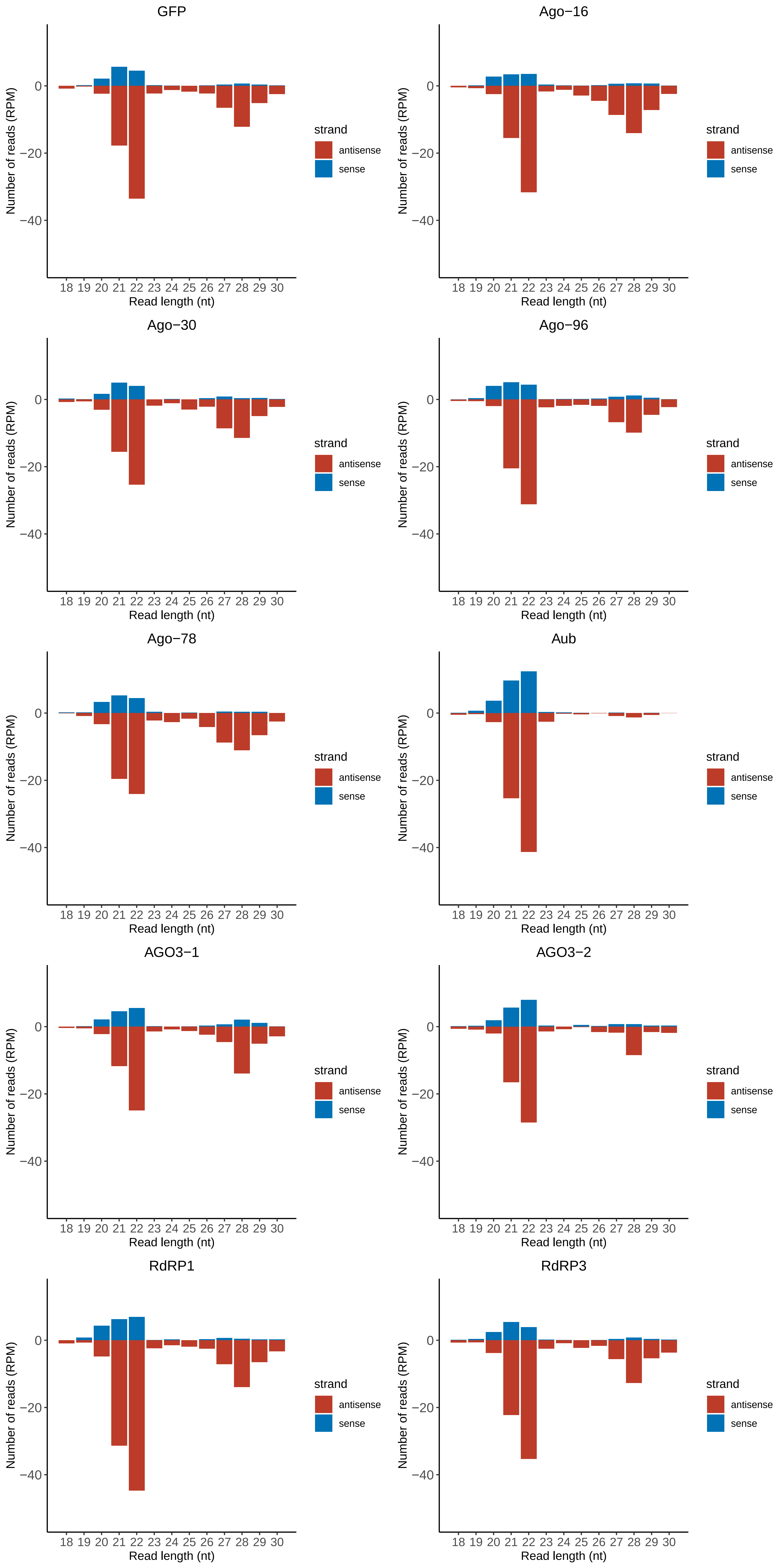

### Motif:rnd-1_family-138TE_lengthdistribution.pdf

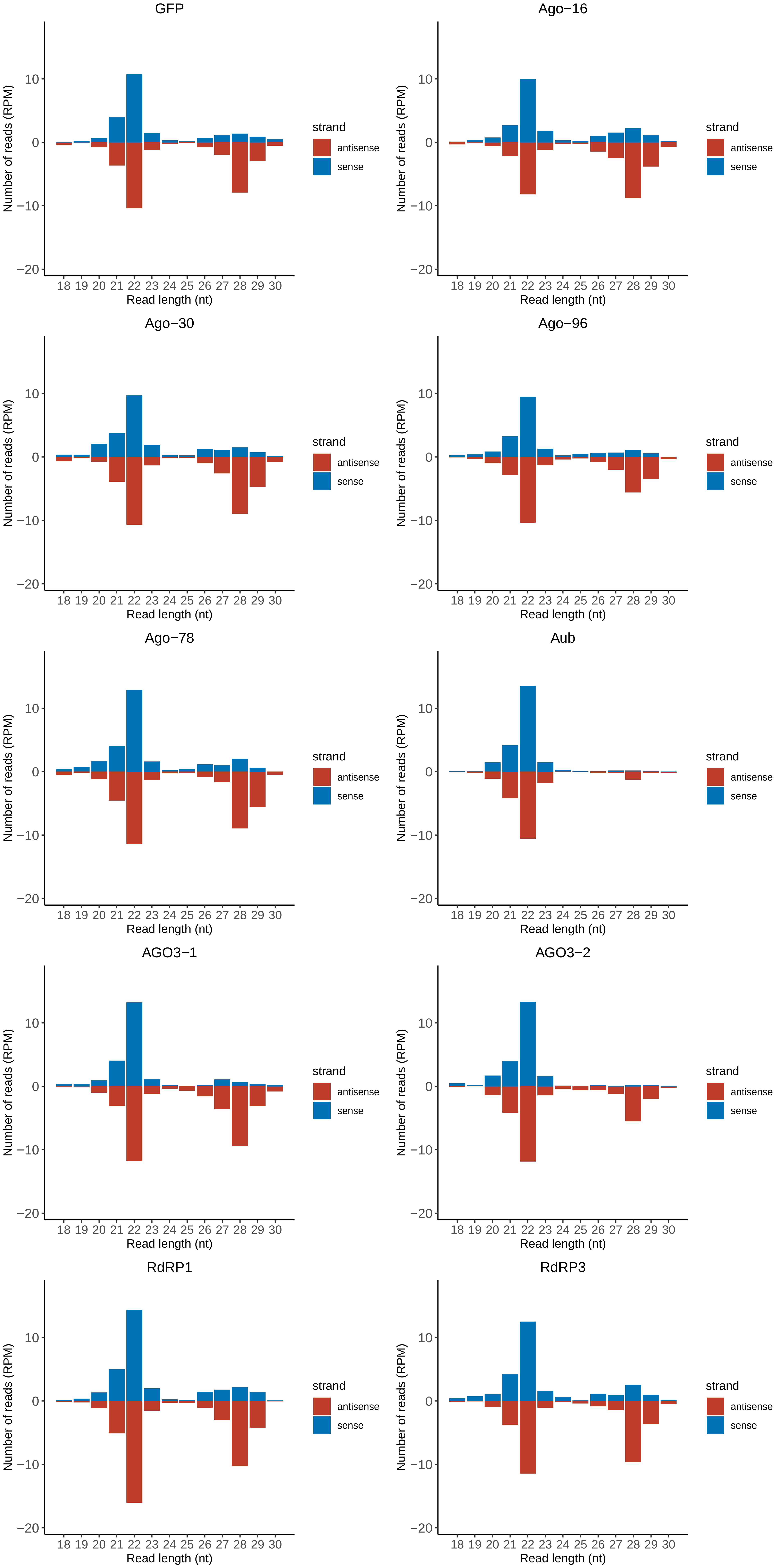

### Motif:rnd-1_family-173TE_lengthdistribution.pdf

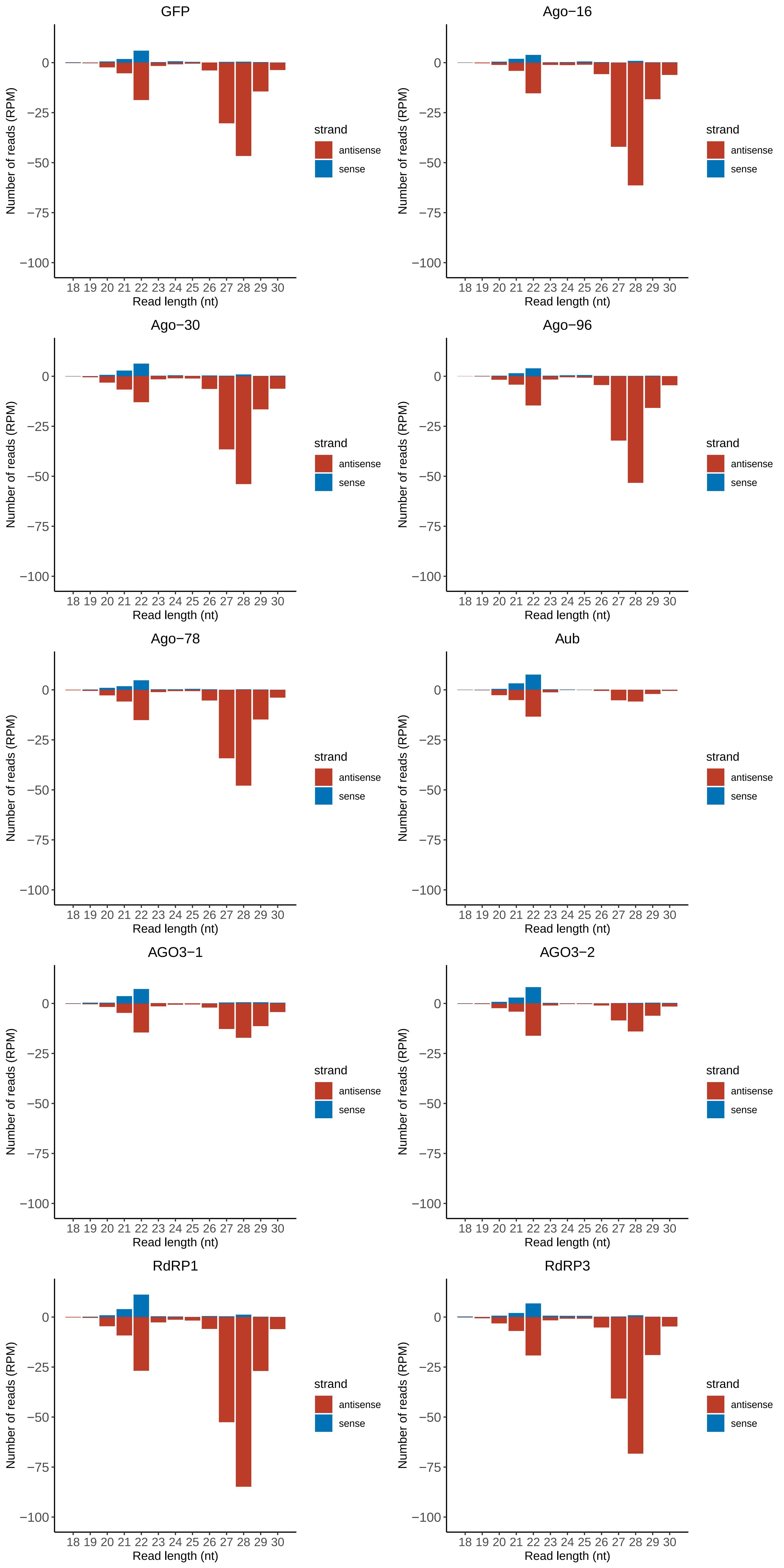

### Motif:rnd-1_family-237TE_lengthdistribution.pdf

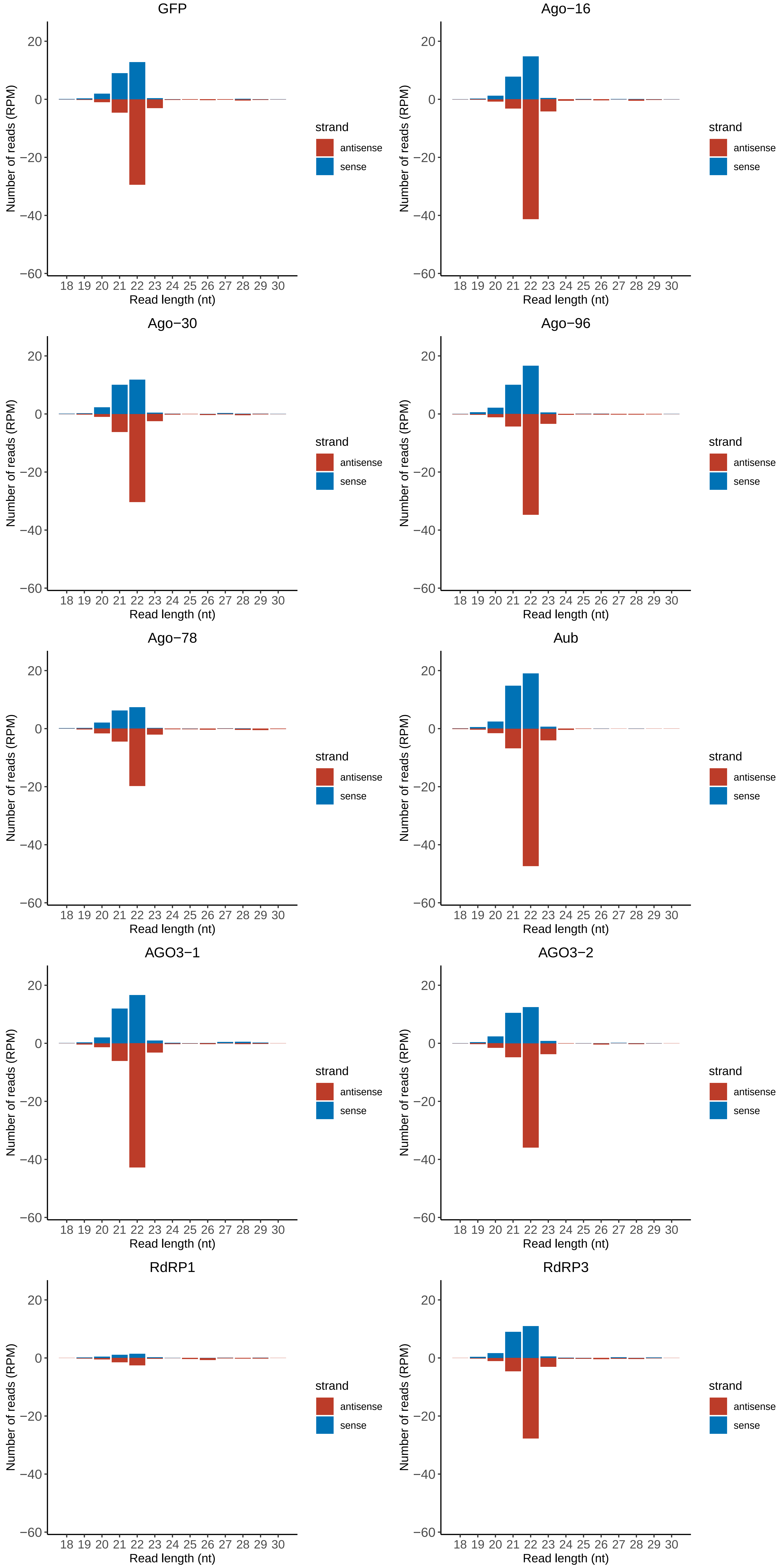

### Motif:rnd-1_family-243TE_lengthdistribution.pdf

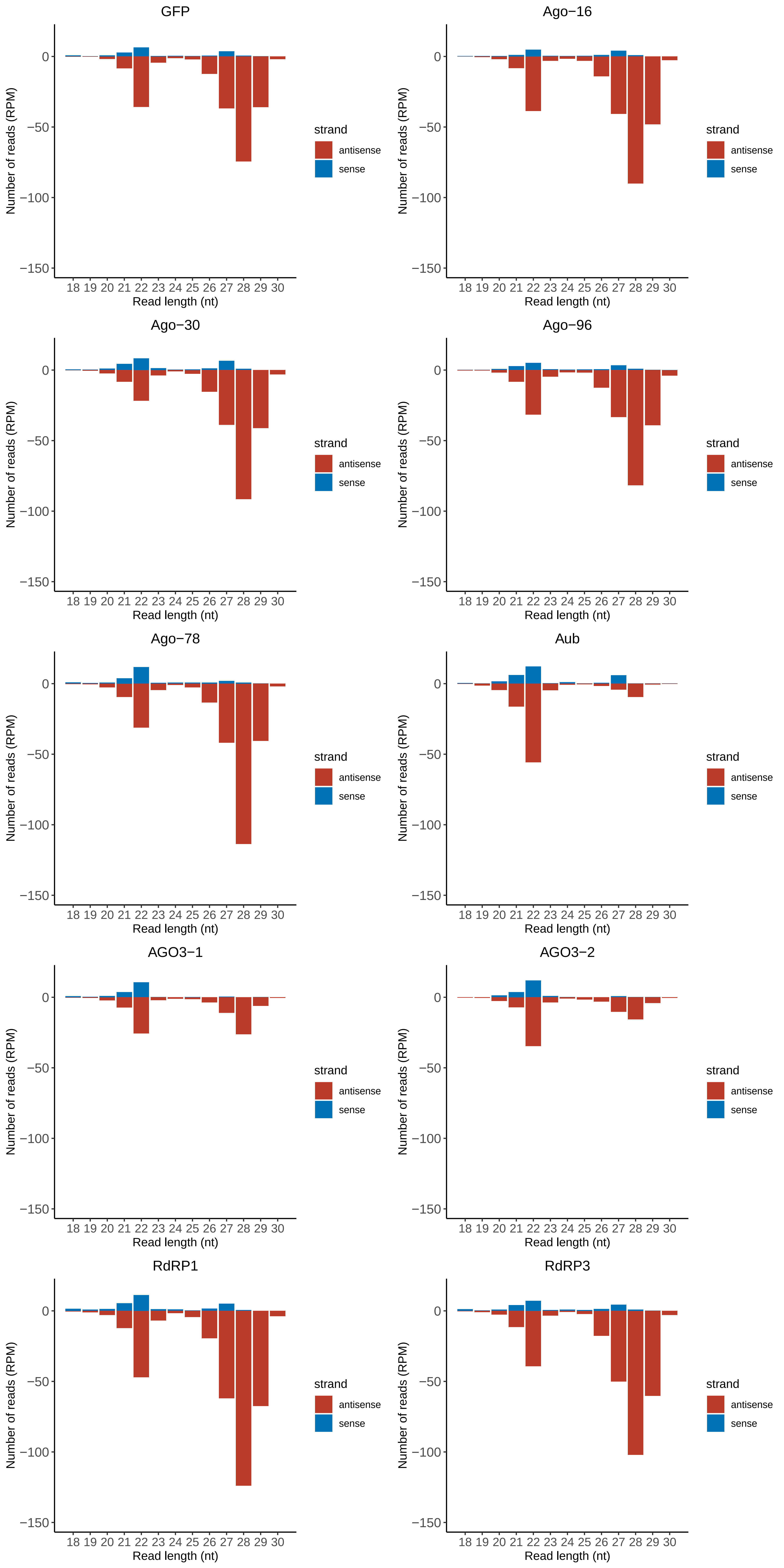

### Motif:rnd-1_family-270TE_lengthdistribution.pdf

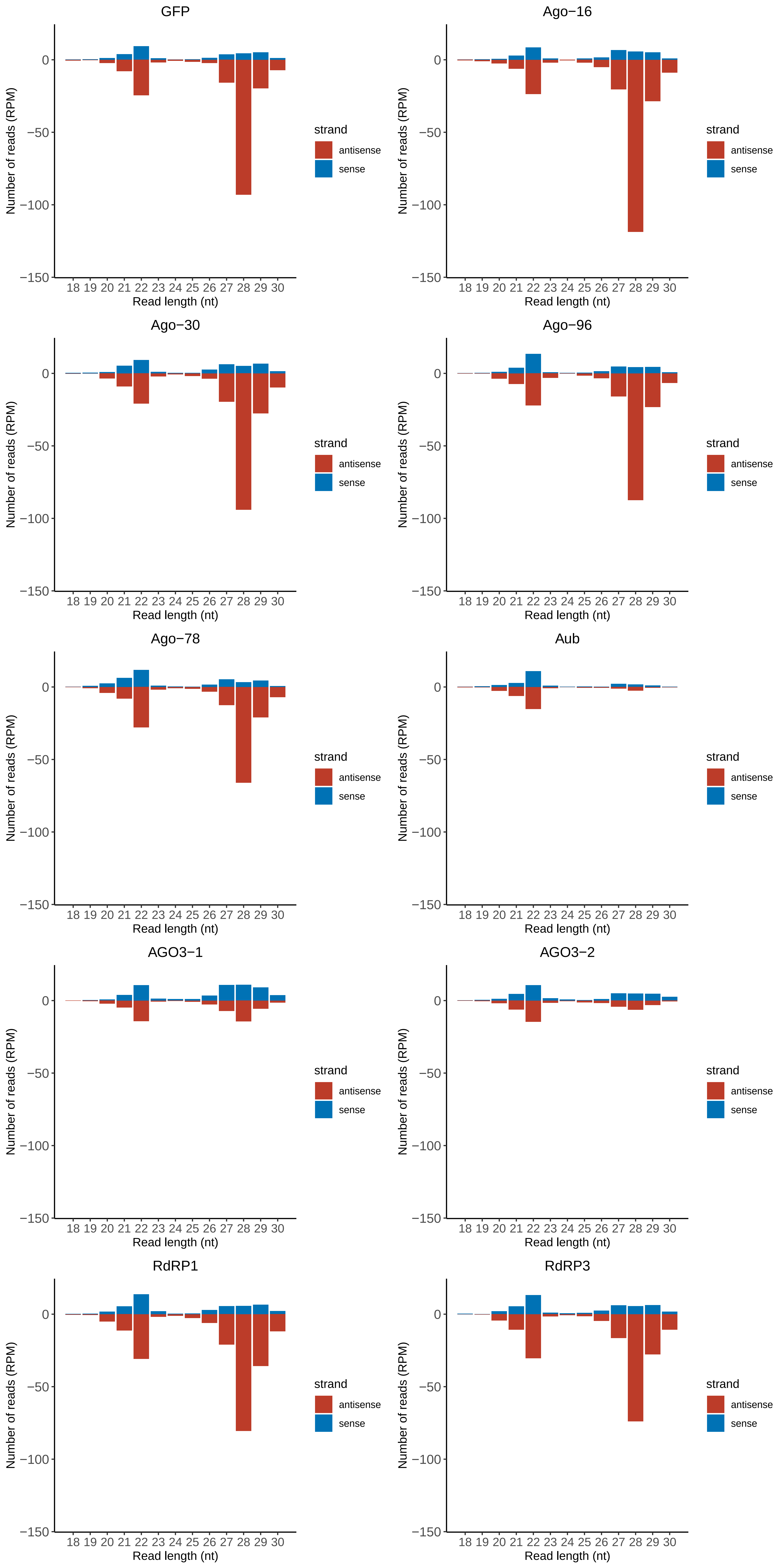

### Motif:rnd-1_family-274TE_lengthdistribution.pdf

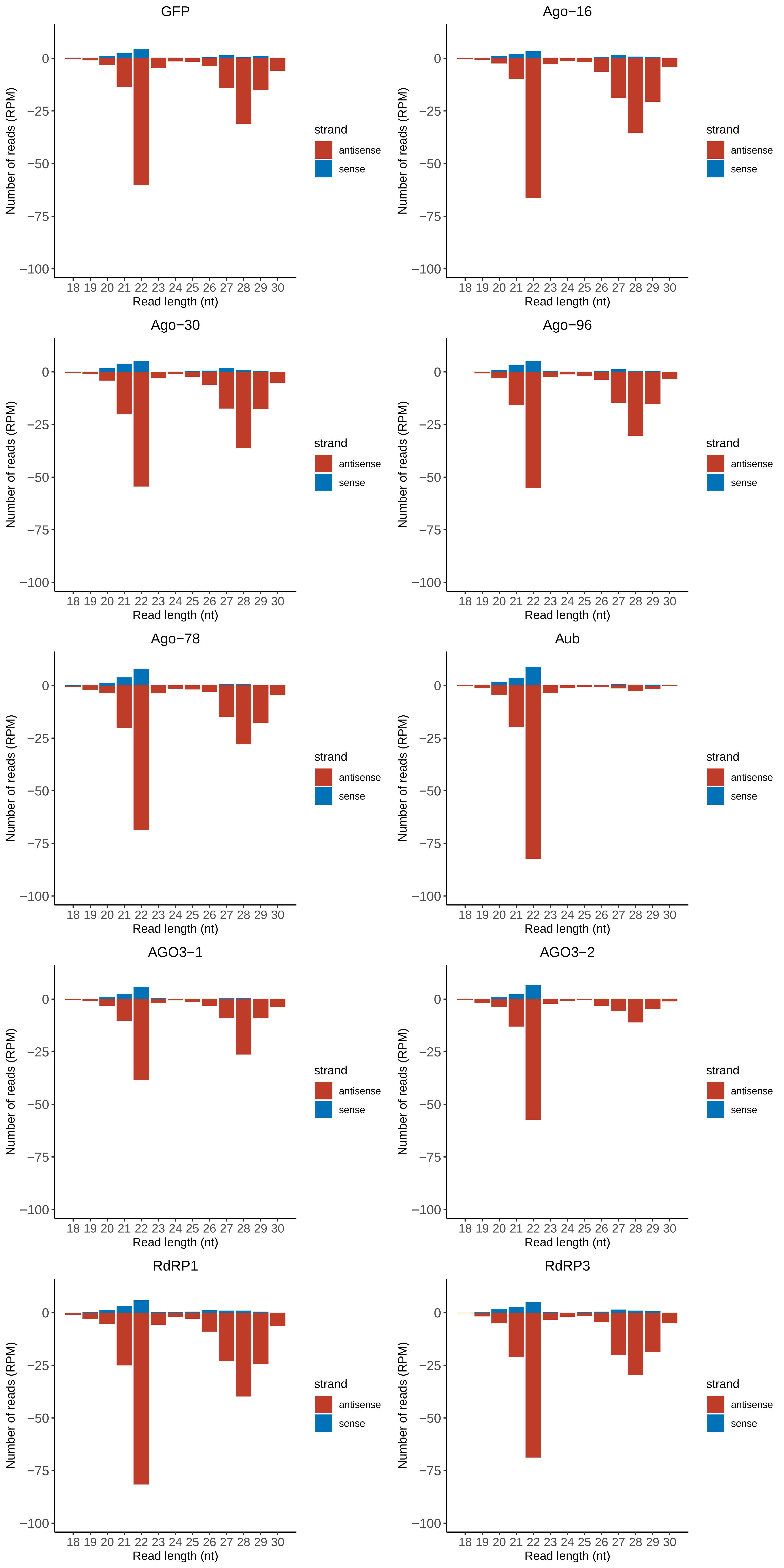

### Motif:rnd-1_family-305TE_lengthdistribution.pdf

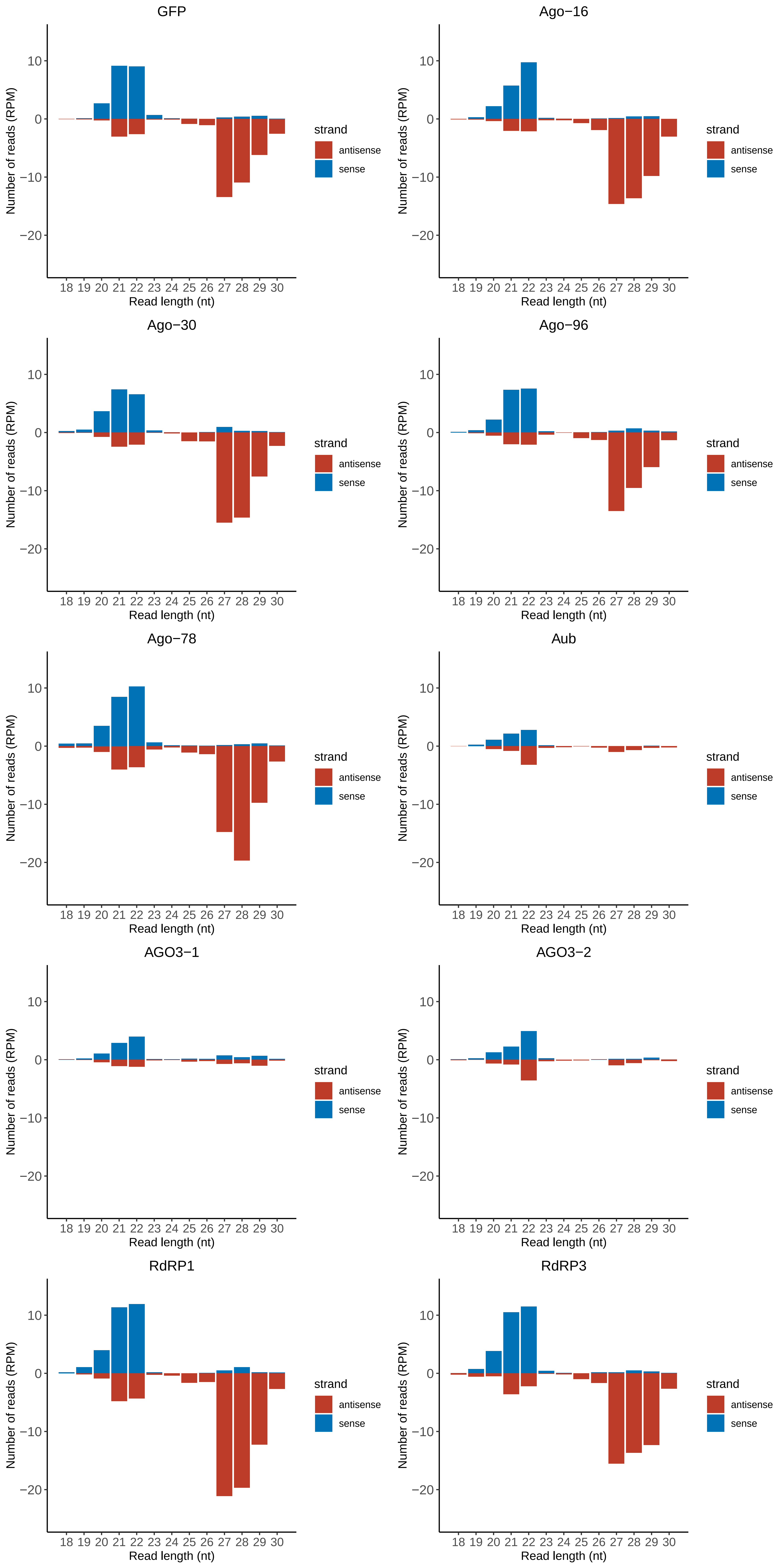

### Motif:rnd-1_family-393TE_lengthdistribution.pdf

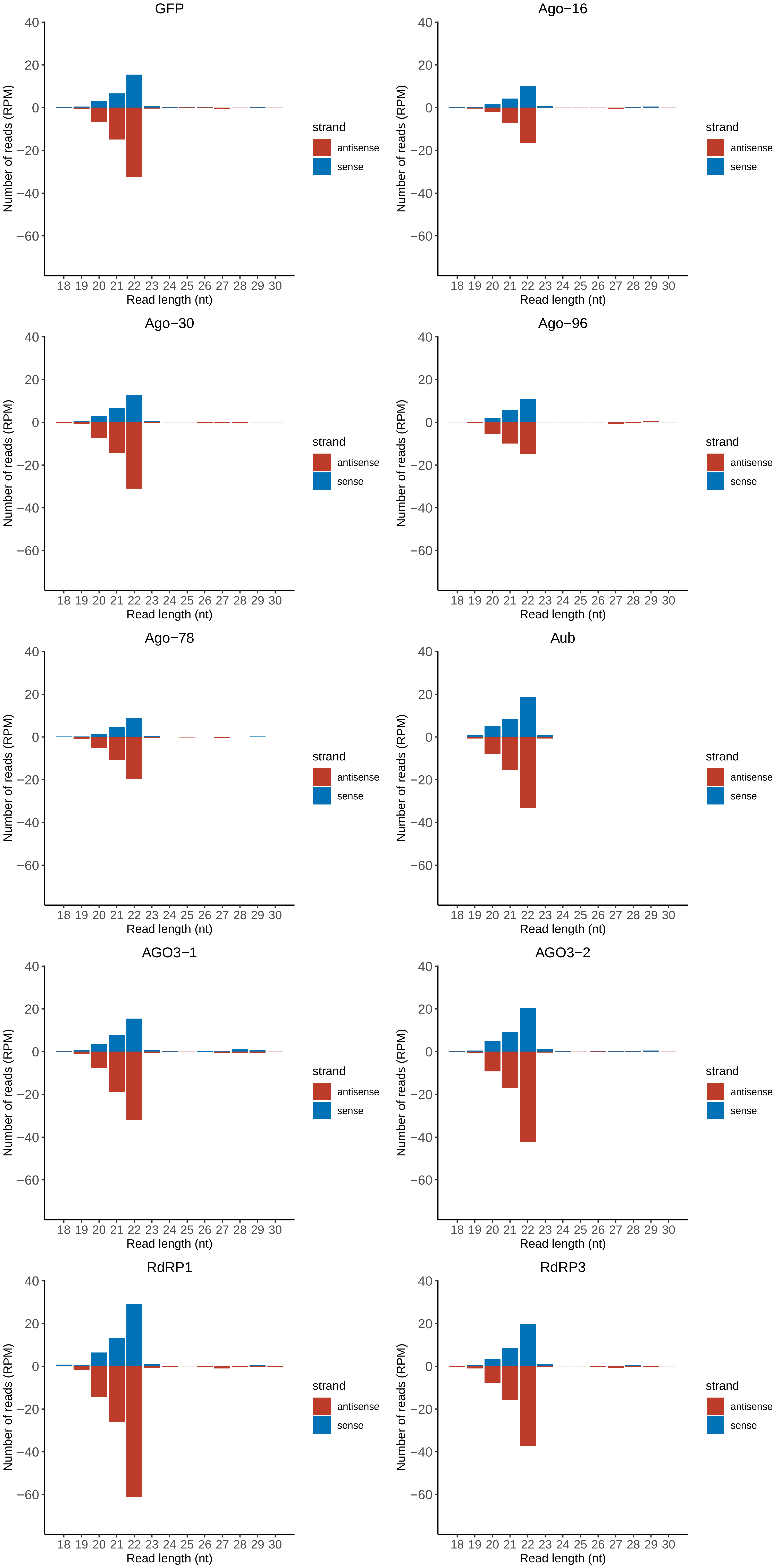

### Motif:rnd-1_family-401TE_lengthdistribution.pdf

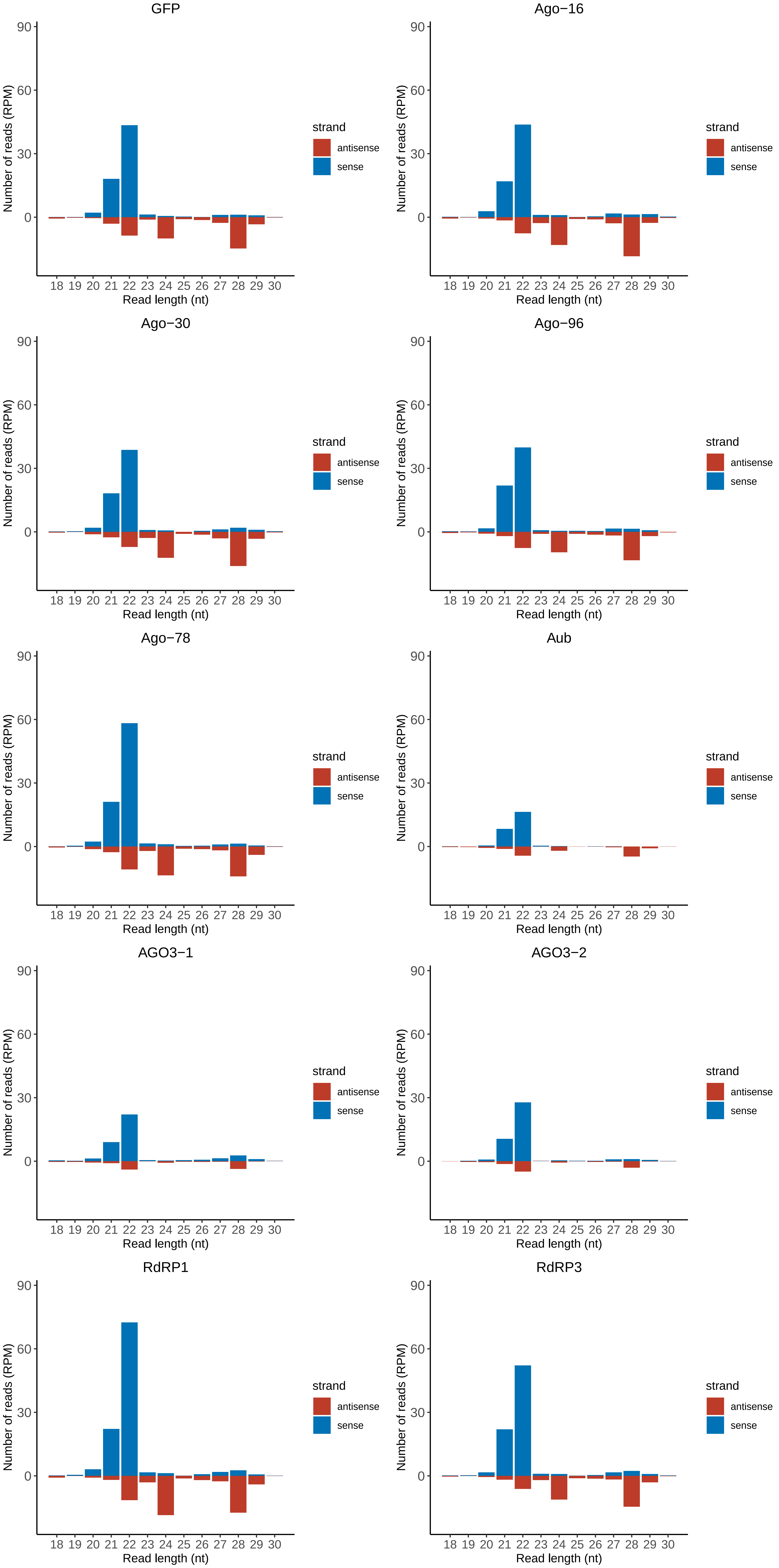

### Motif:rnd-1_family-463TE_lengthdistribution.pdf

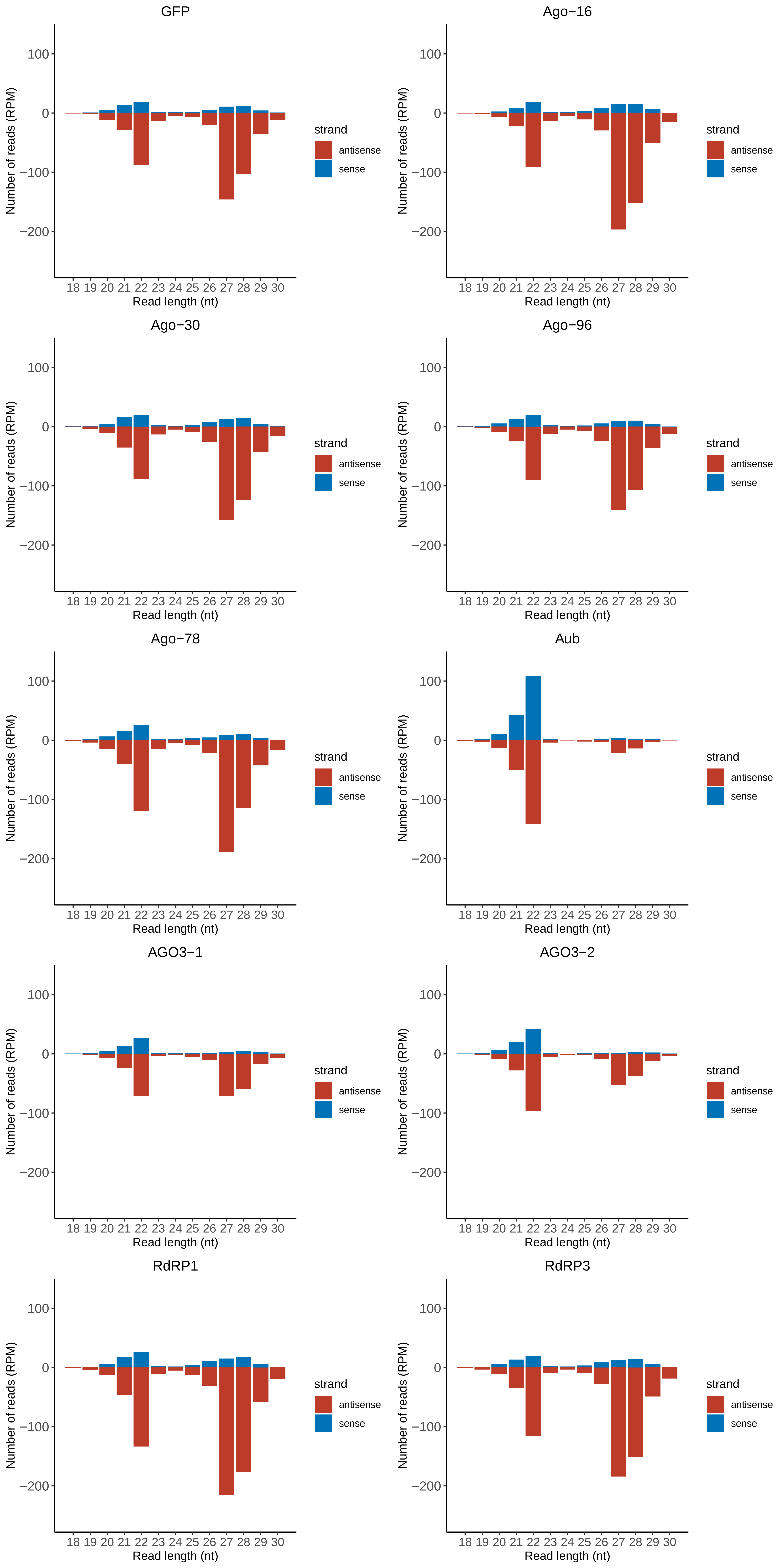

### Motif:rnd-1_family-603TE_lengthdistribution.pdf

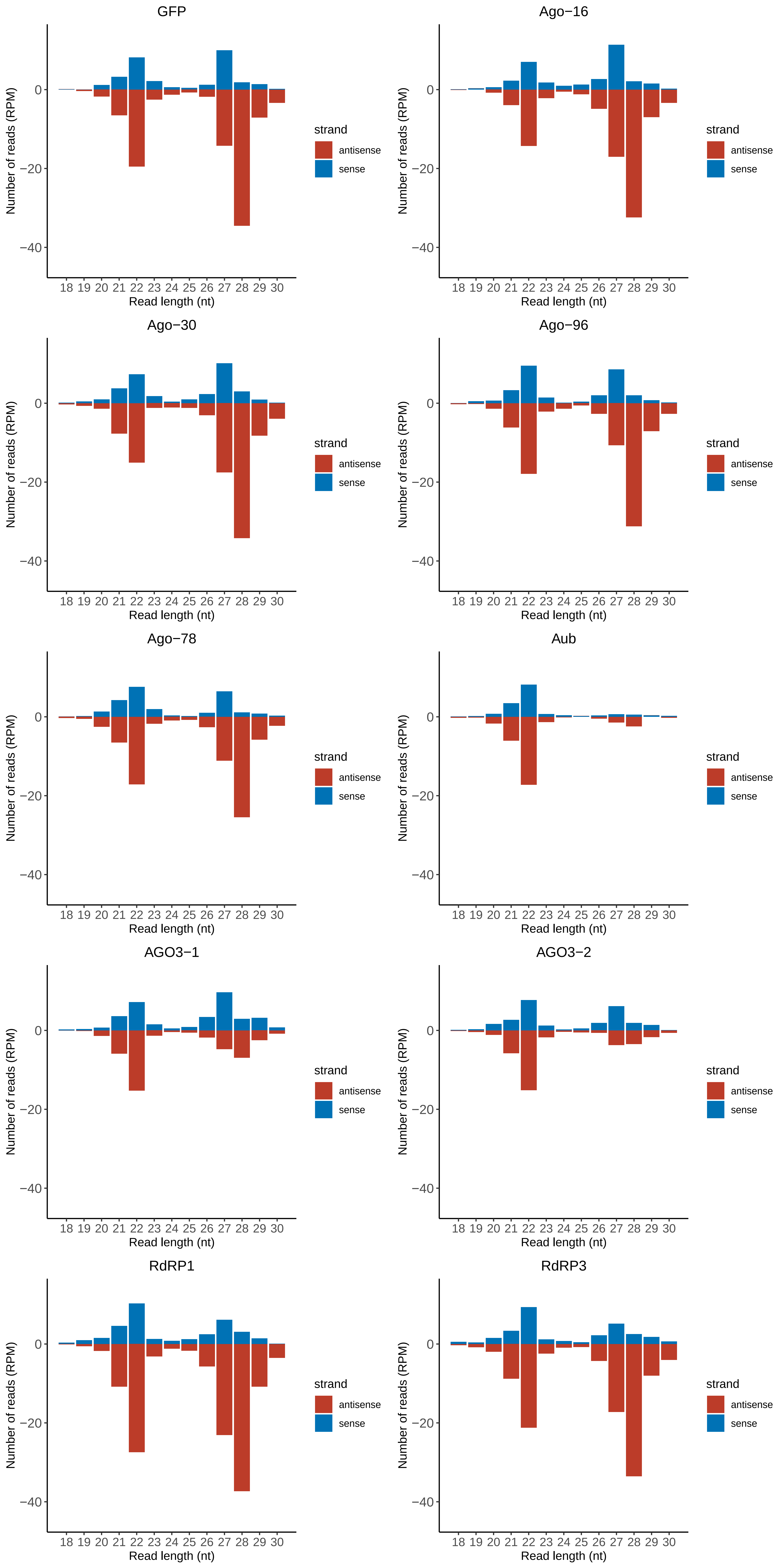

### Motif:rnd-1_family-607TE_lengthdistribution.pdf

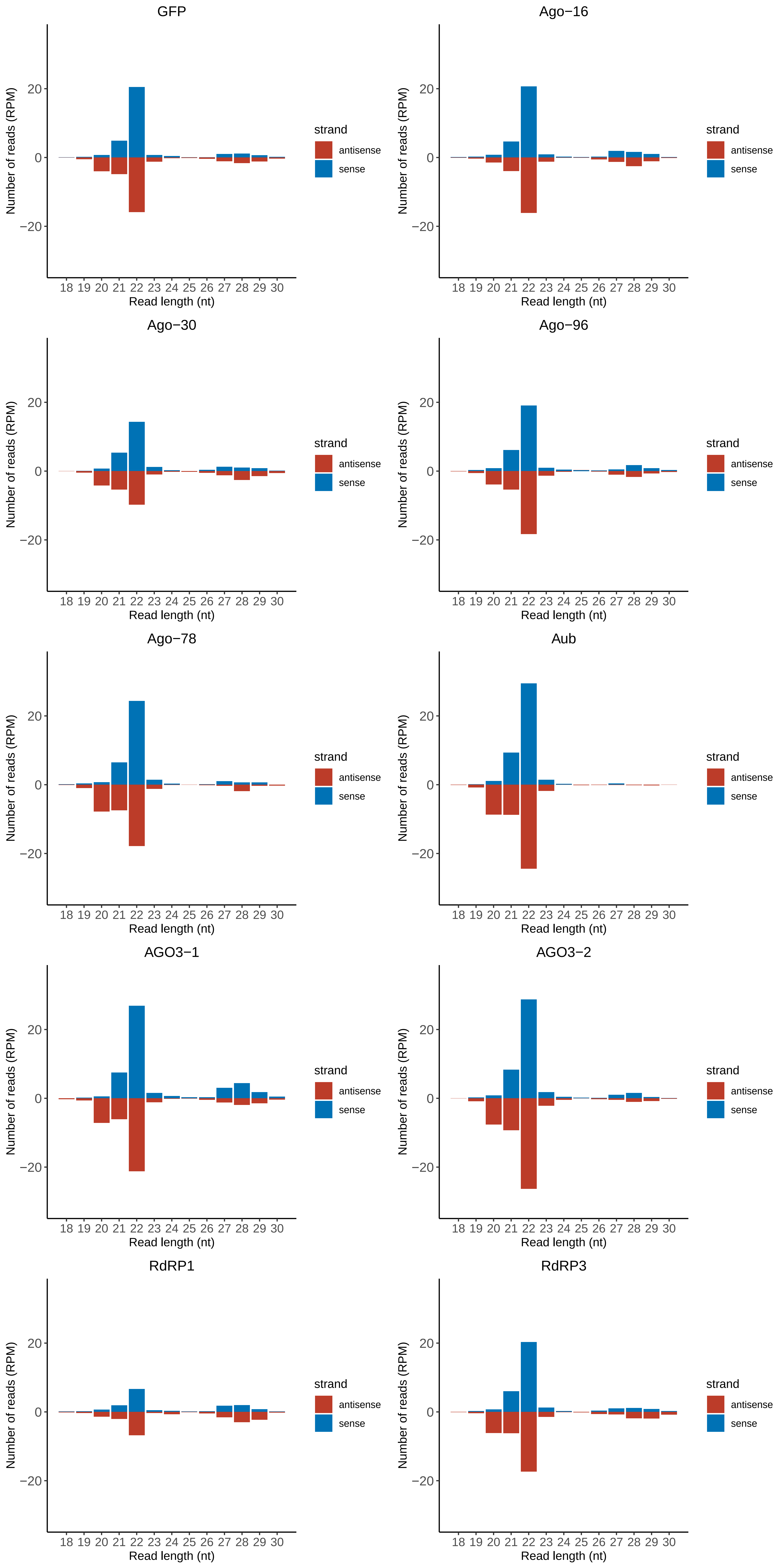

### Motif:rnd-1_family-669TE_lengthdistribution.pdf

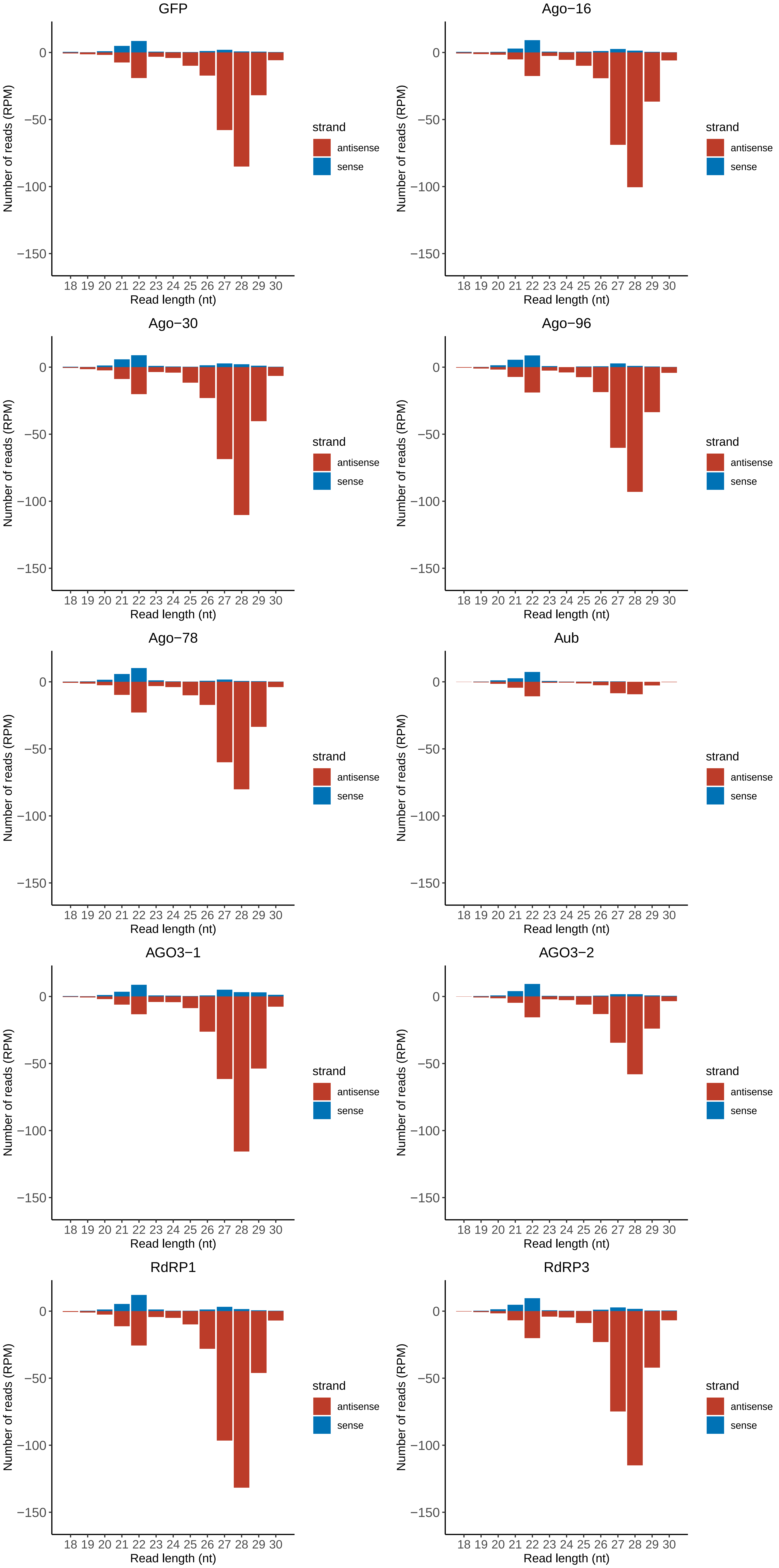

### Motif:rnd-1_family-731TE_lengthdistribution.pdf

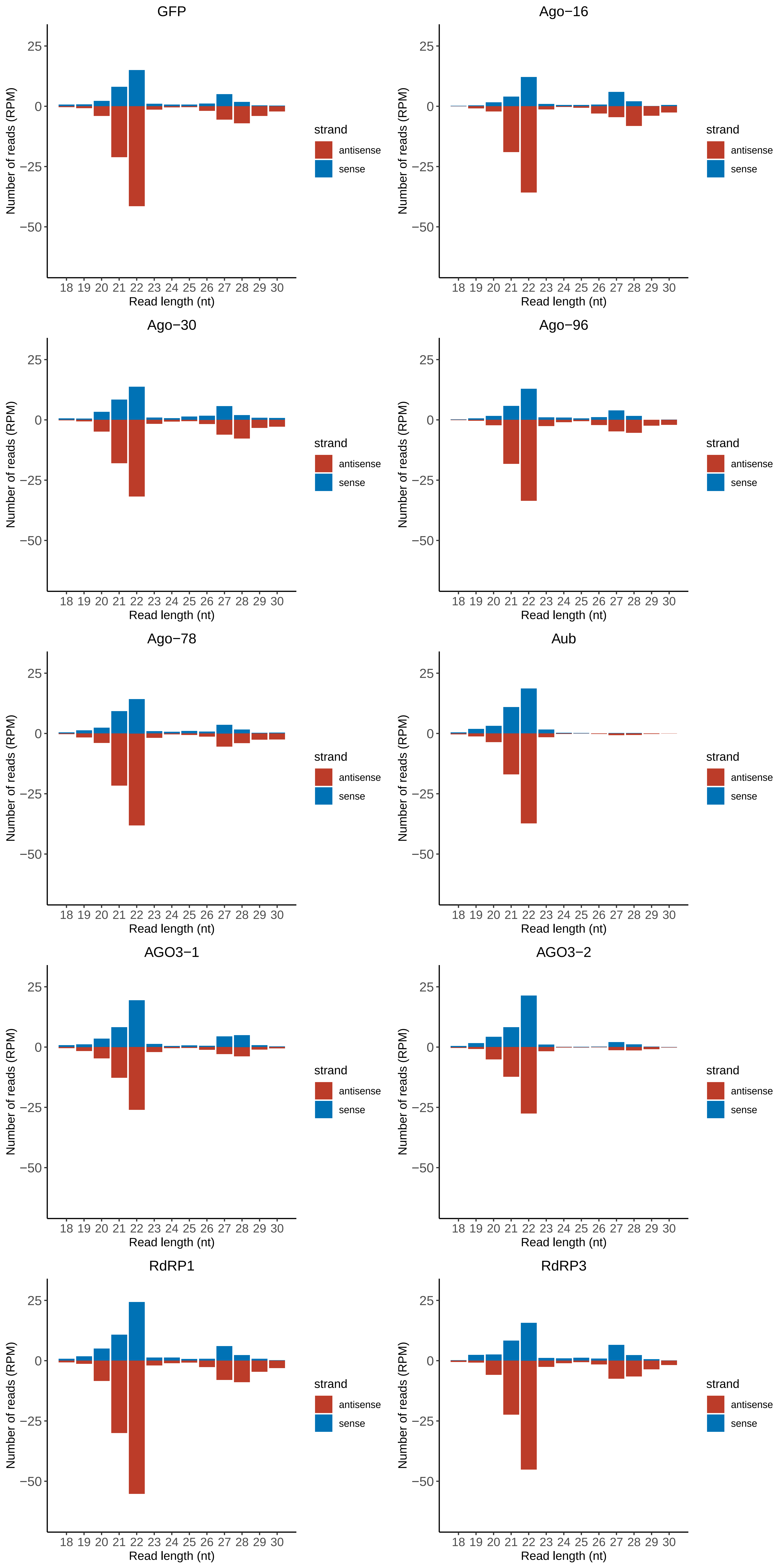

### Motif:rnd-1_family-809TE_lengthdistribution.pdf

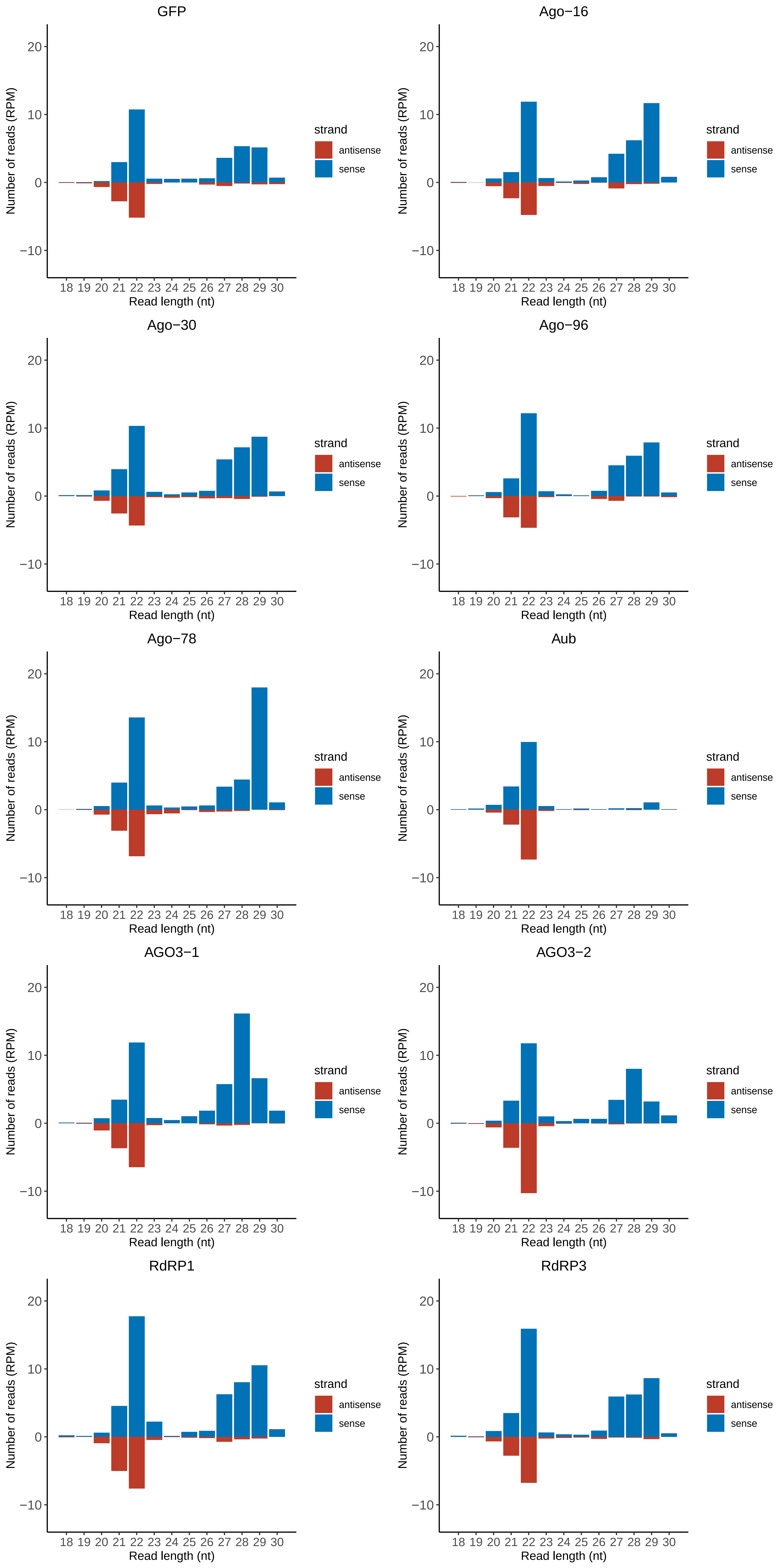

### Motif:rnd-1_family-832TE_lengthdistribution.pdf

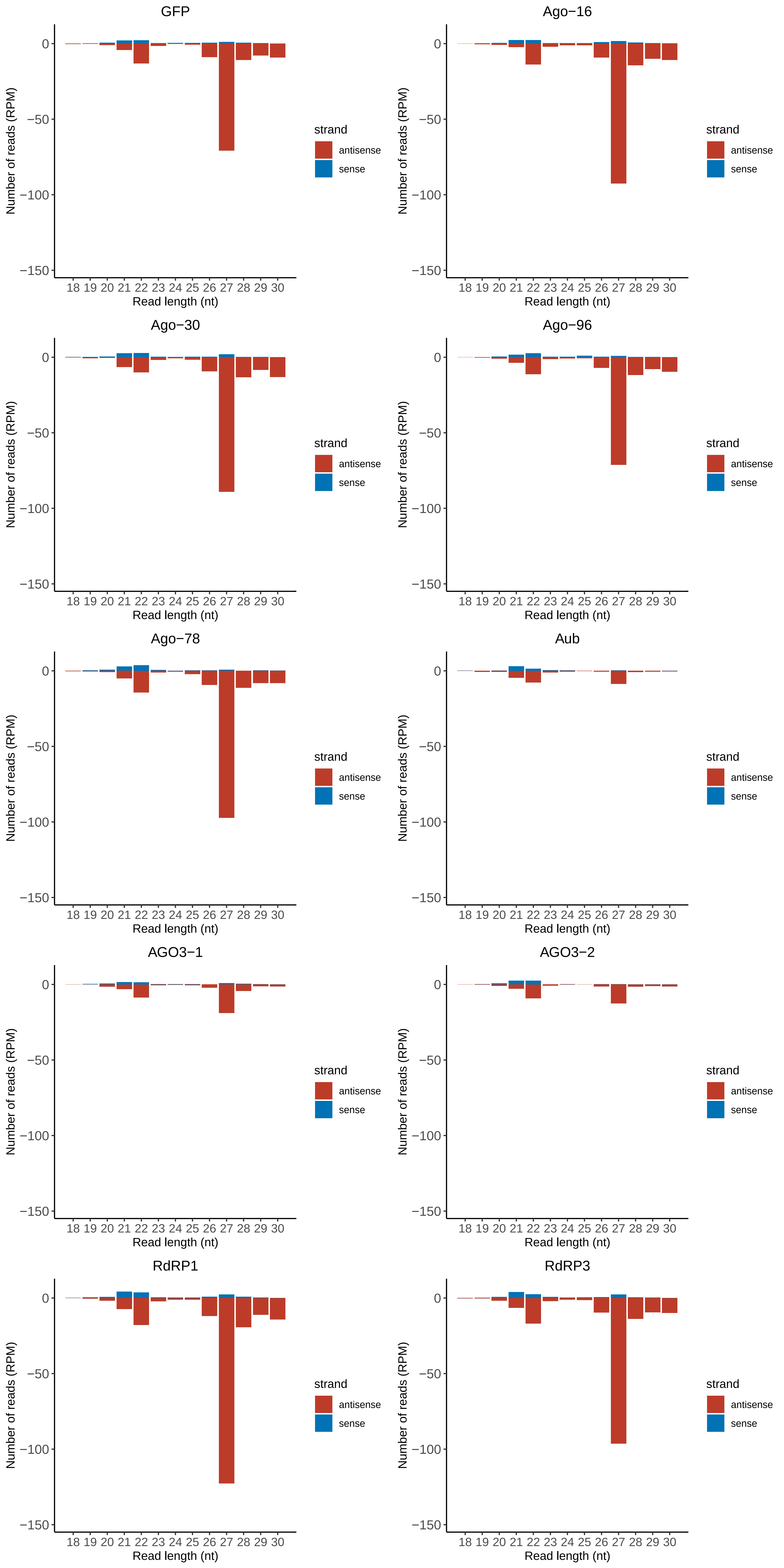

### Motif:rnd-1_family-955TE_lengthdistribution.pdf

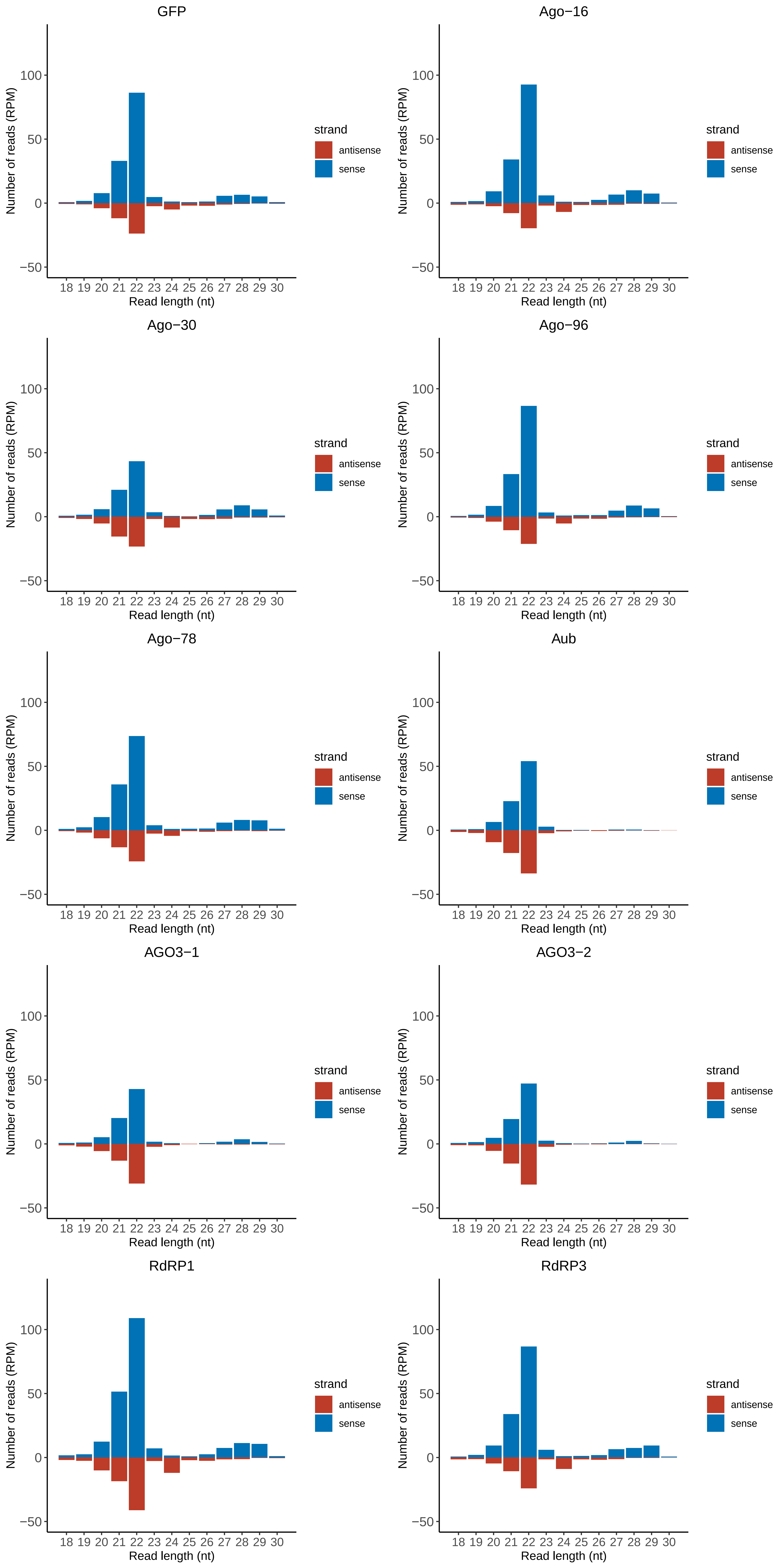

### Motif:rnd-1_family-970TE_lengthdistribution.pdf

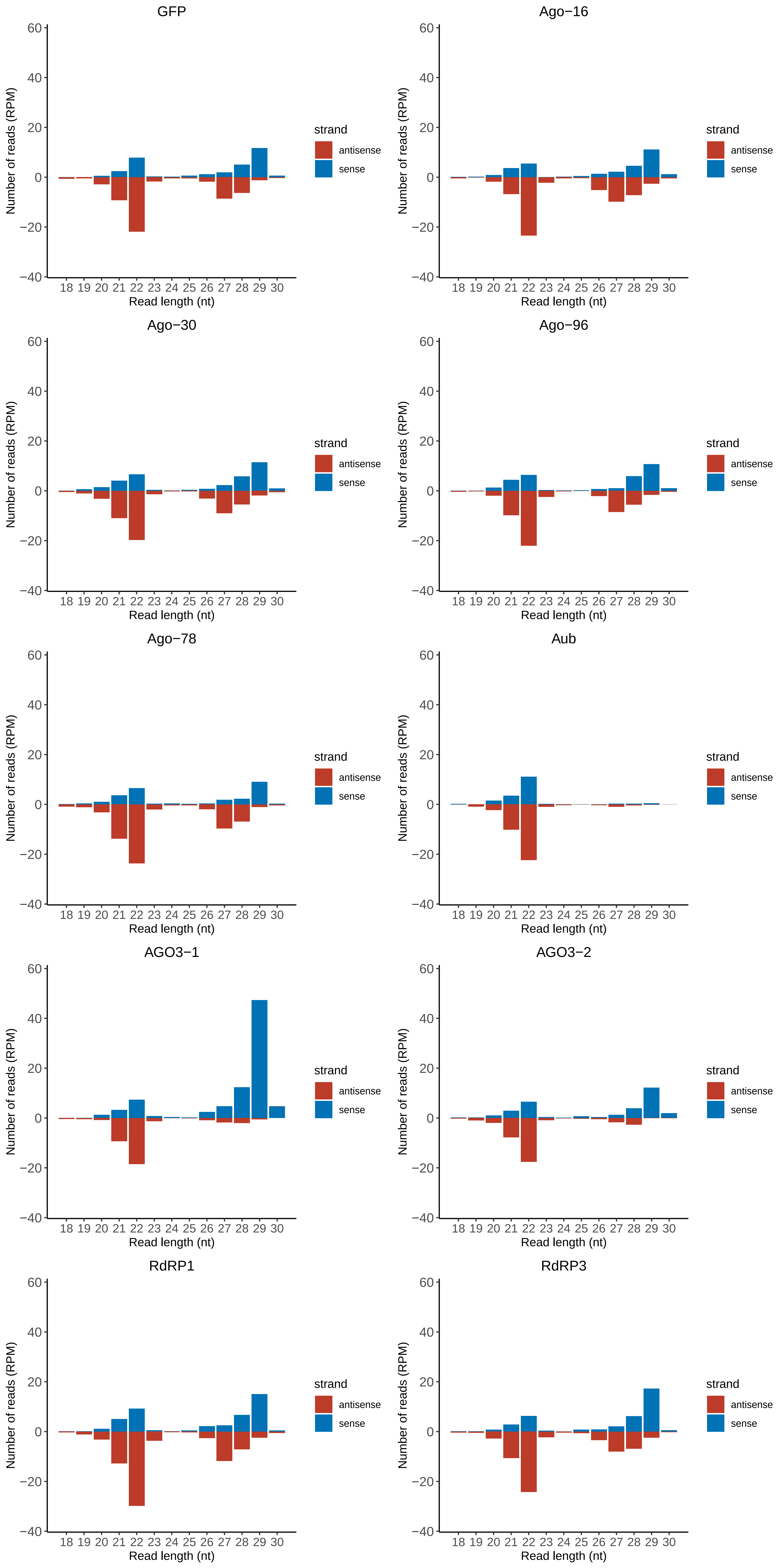

### Motif:rnd-1_family-1055TE_lengthdistribution.pdf

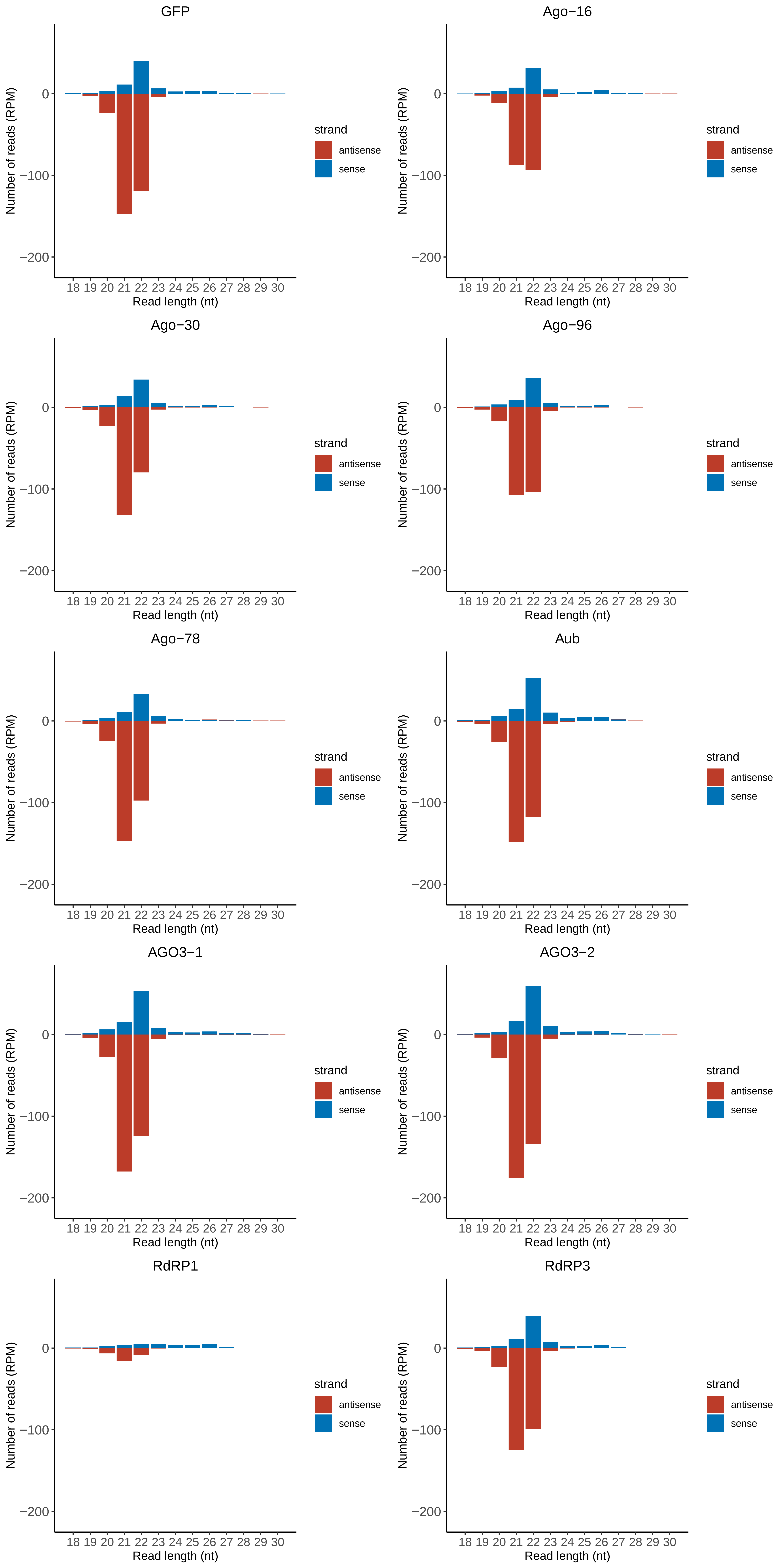

### Motif:rnd-1_family-1070TE_lengthdistribution.pdf

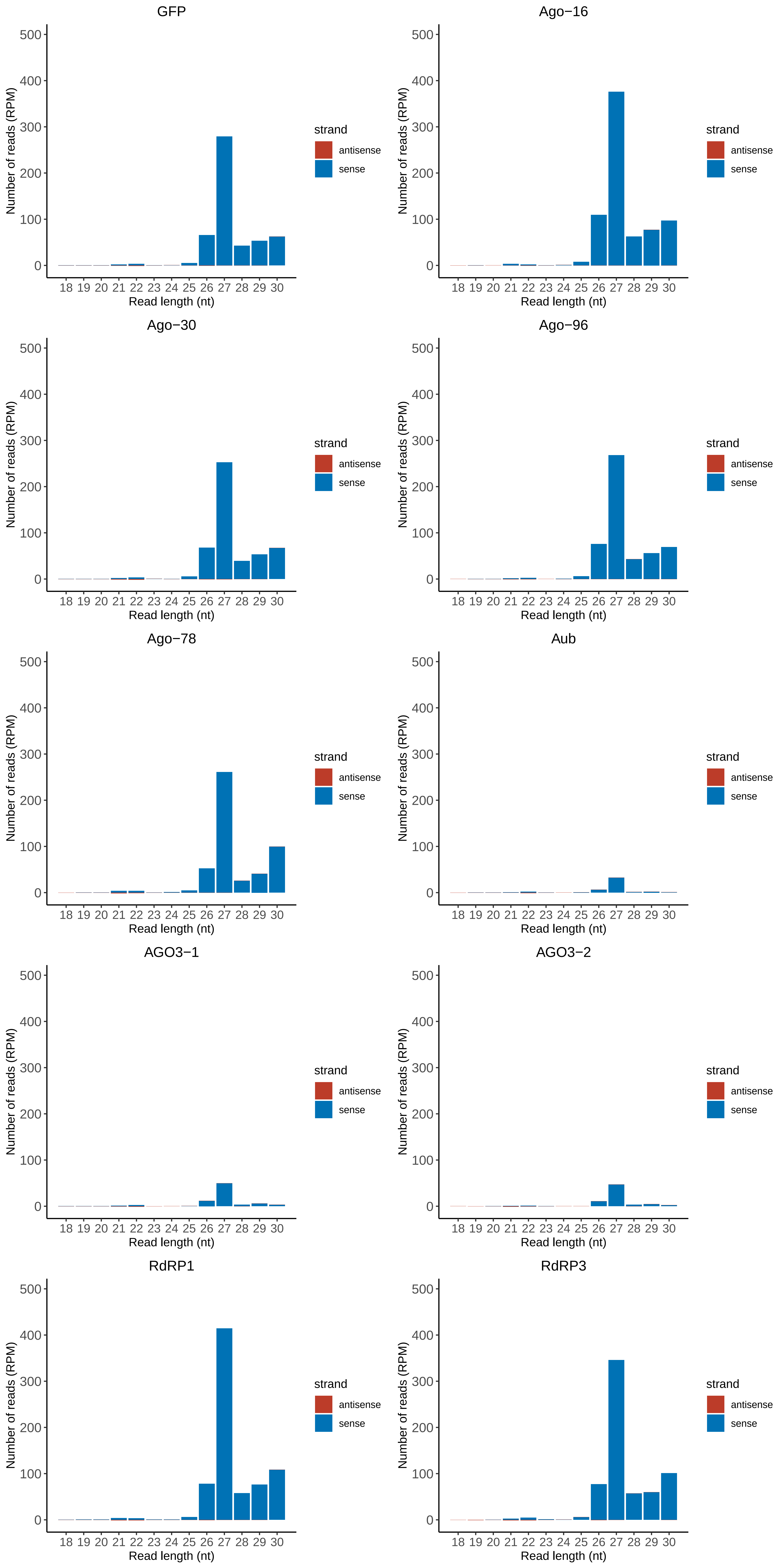

### Motif:rnd-1_family-1113TE_lengthdistribution.pdf

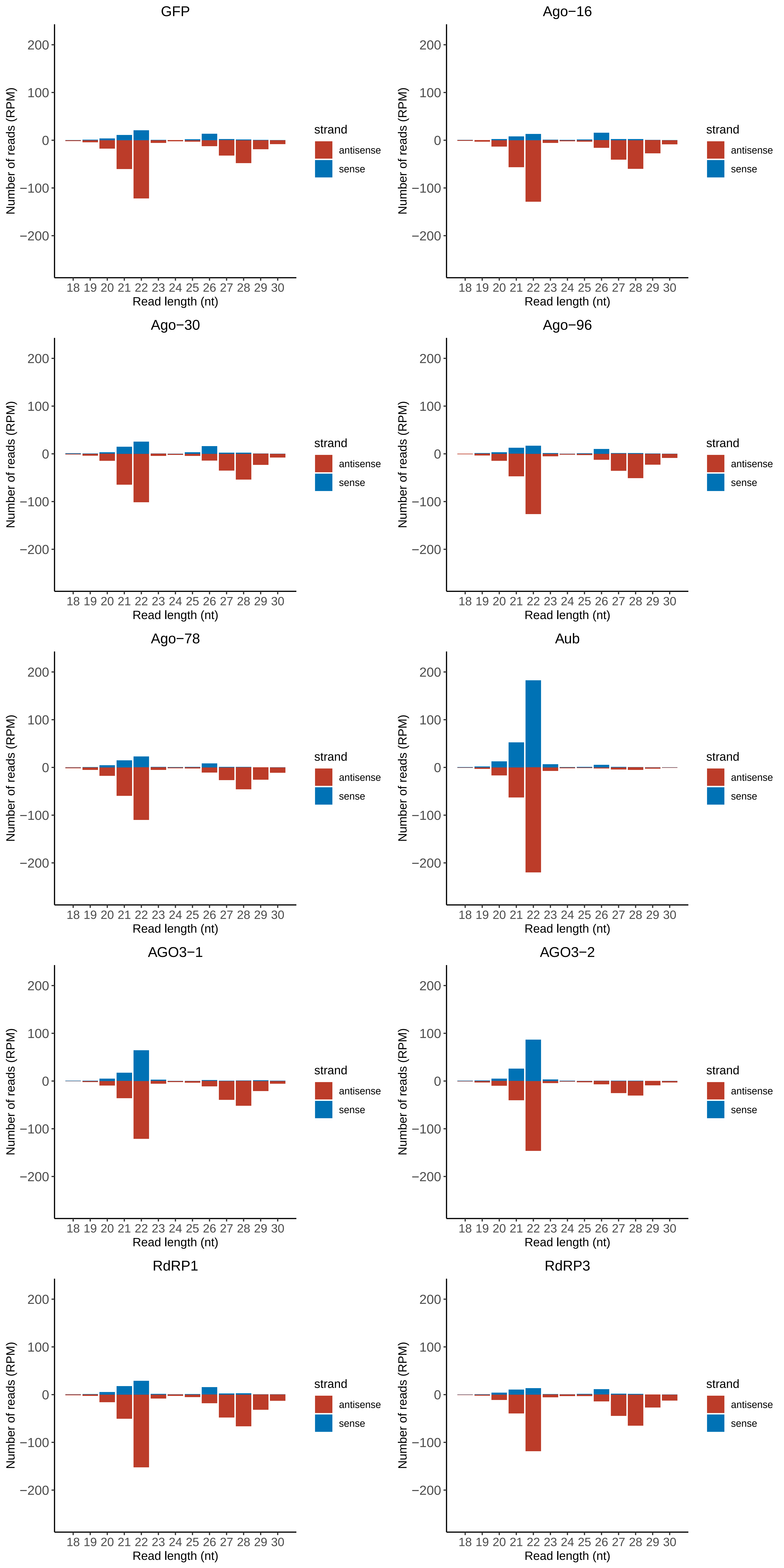

### Motif:rnd-1_family-1128TE_lengthdistribution.pdf

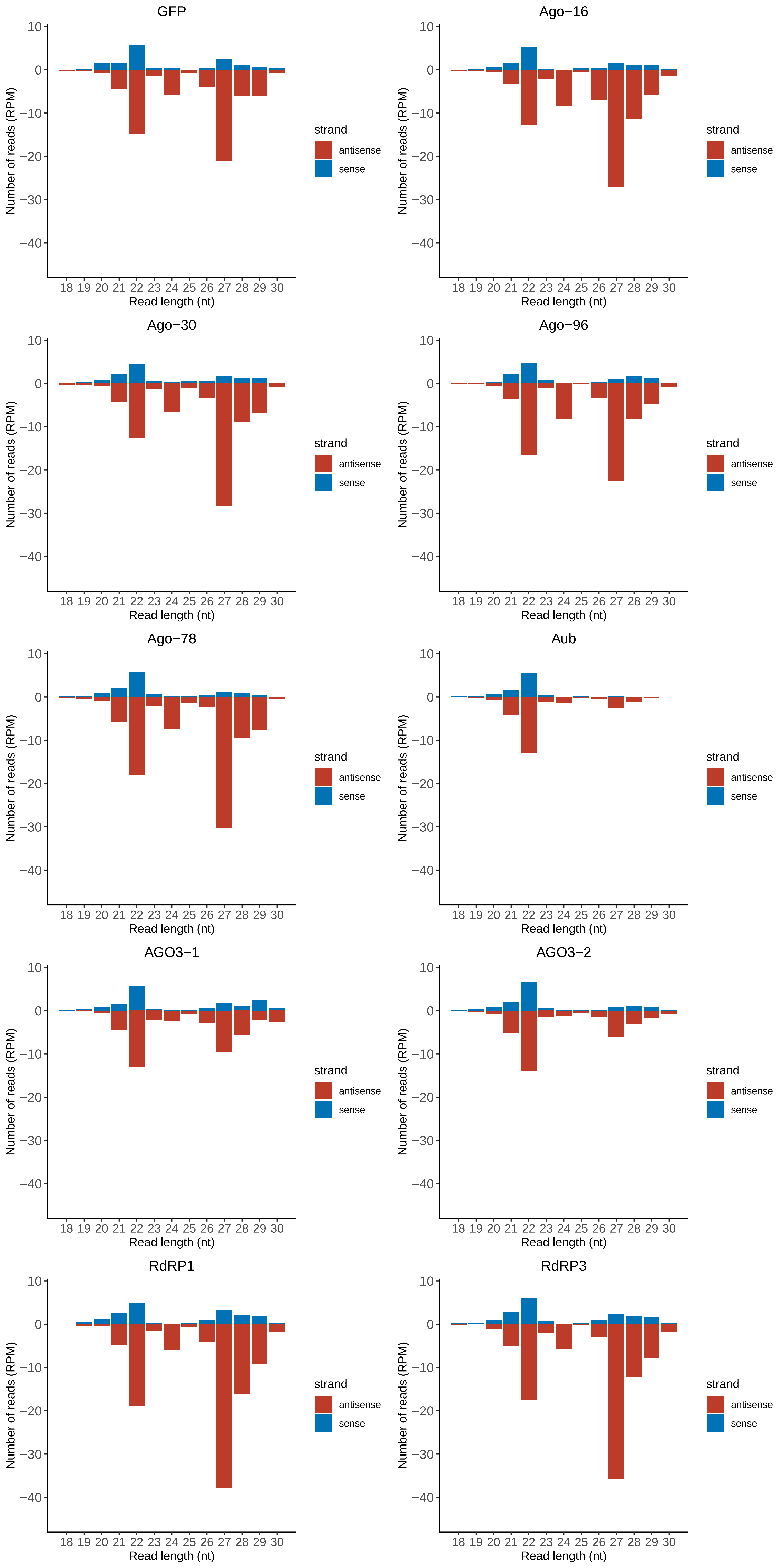

### Motif:rnd-1_family-1373TE_lengthdistribution.pdf

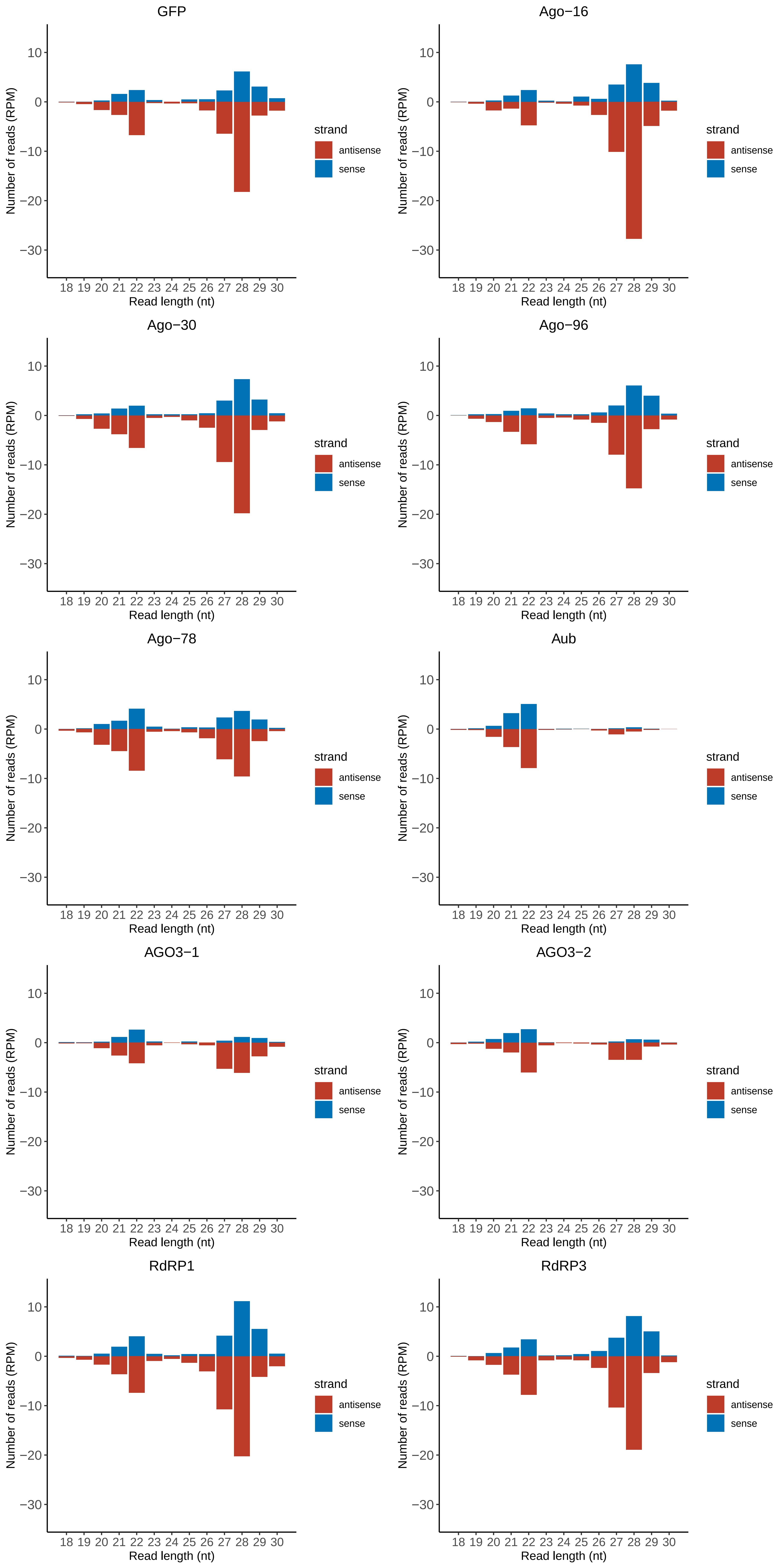

### Motif:rnd-1_family-1391TE_lengthdistribution.pdf

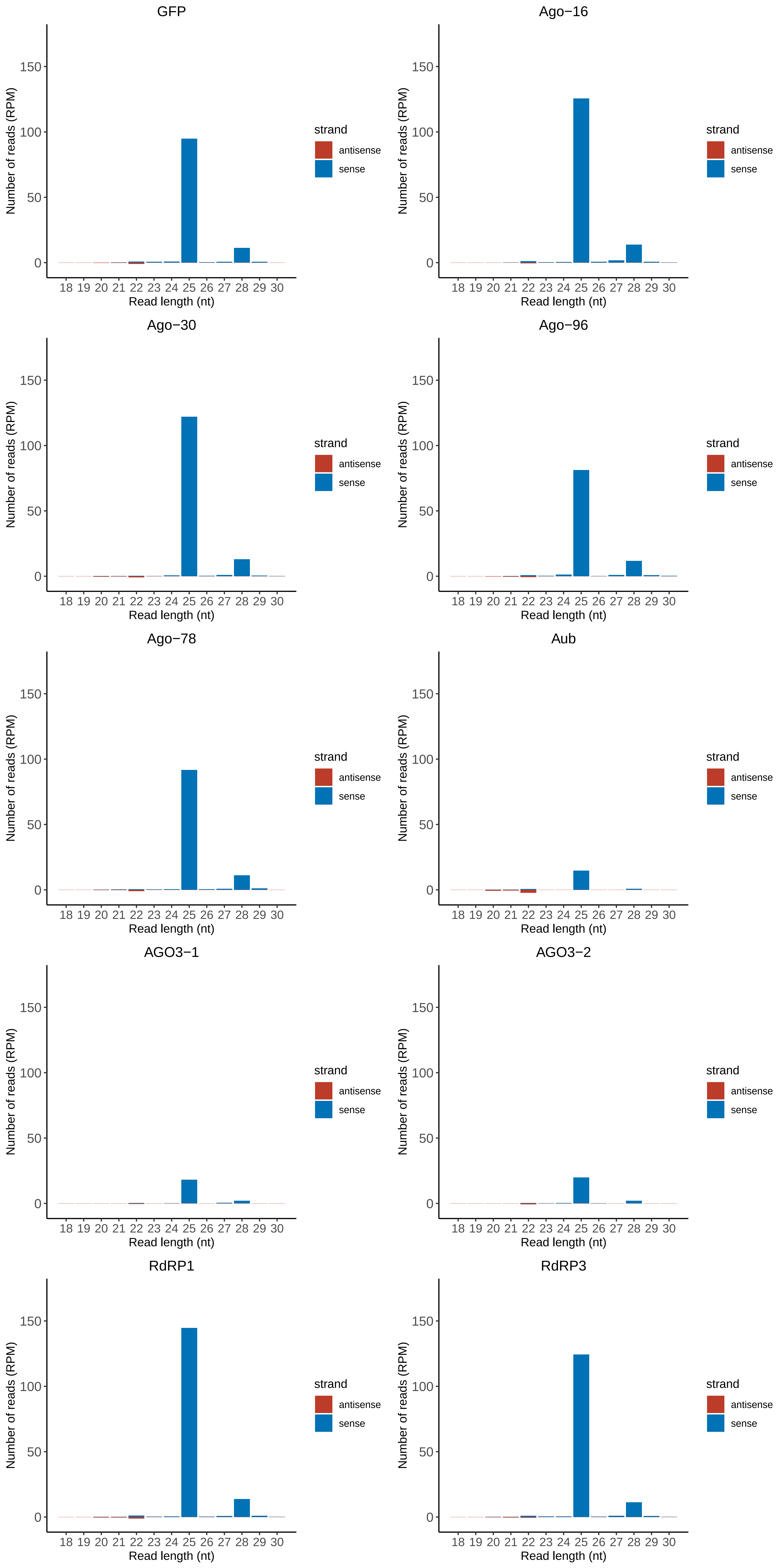

### Motif:rnd-1_family-1531TE_lengthdistribution.pdf

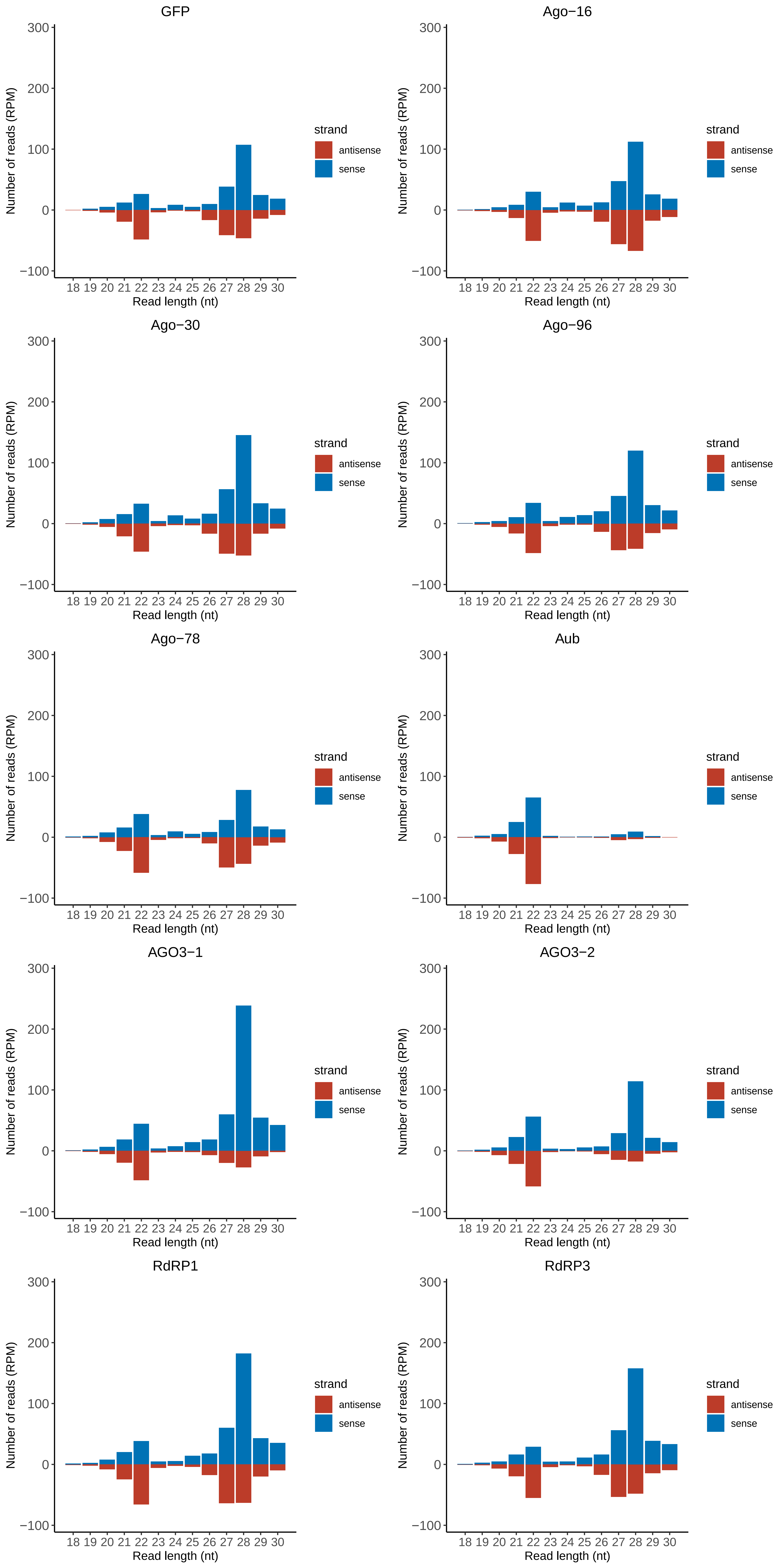

### Motif:rnd-1_family-1557TE_lengthdistribution.pdf

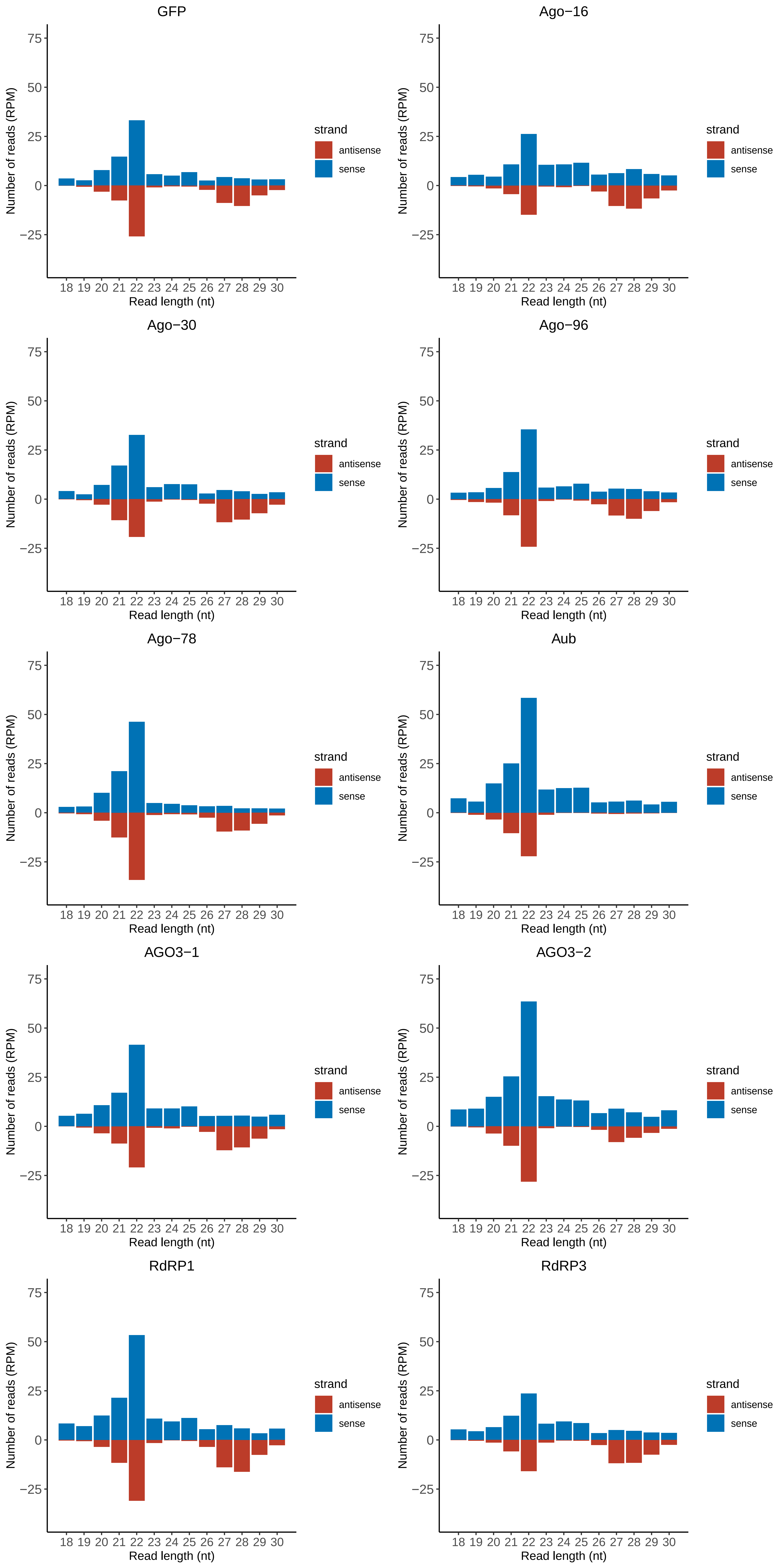

### Motif:rnd-1_family-1650TE_lengthdistribution.pdf

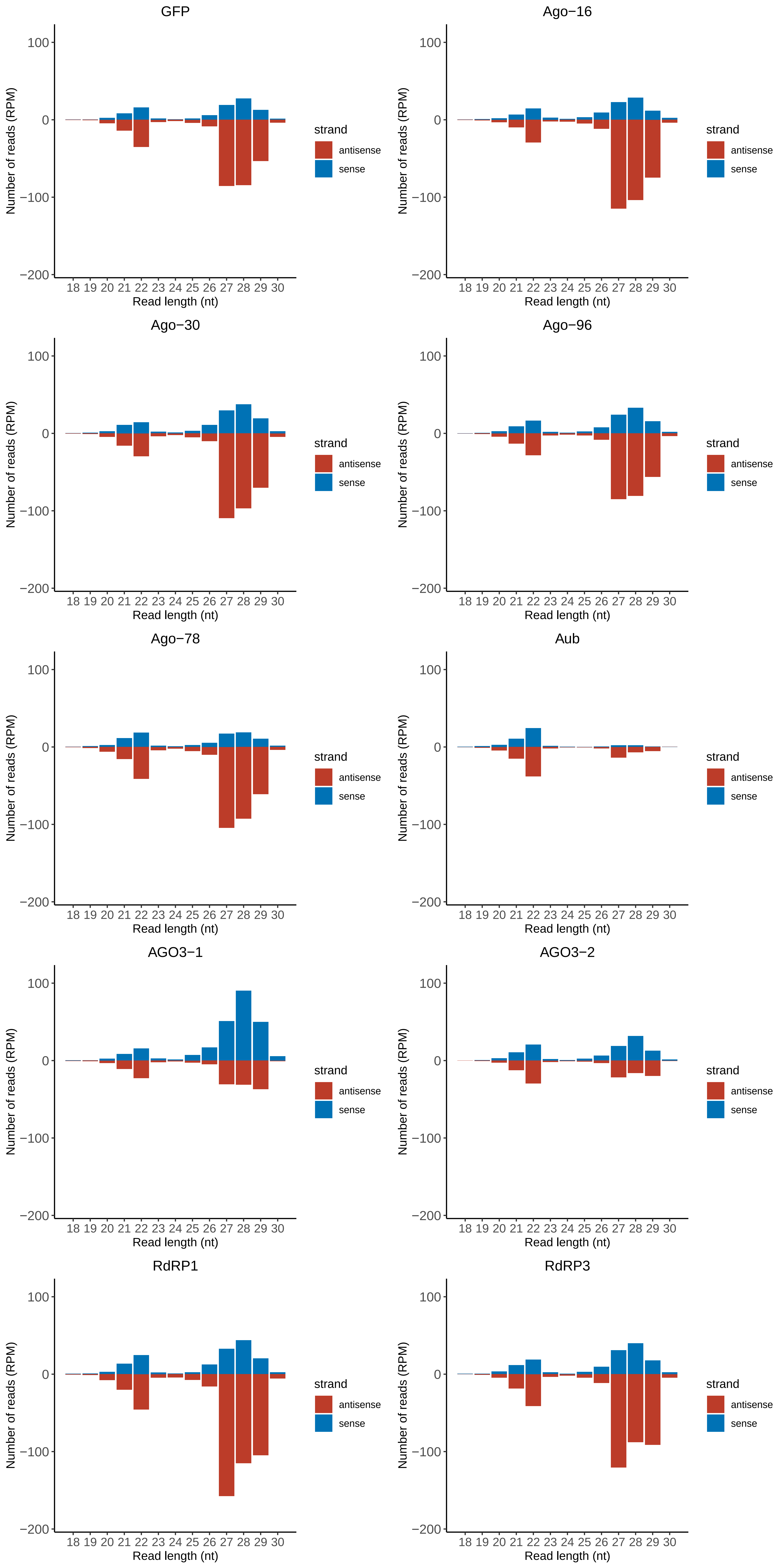

### Motif:rnd-1_family-1751TE_lengthdistribution.pdf

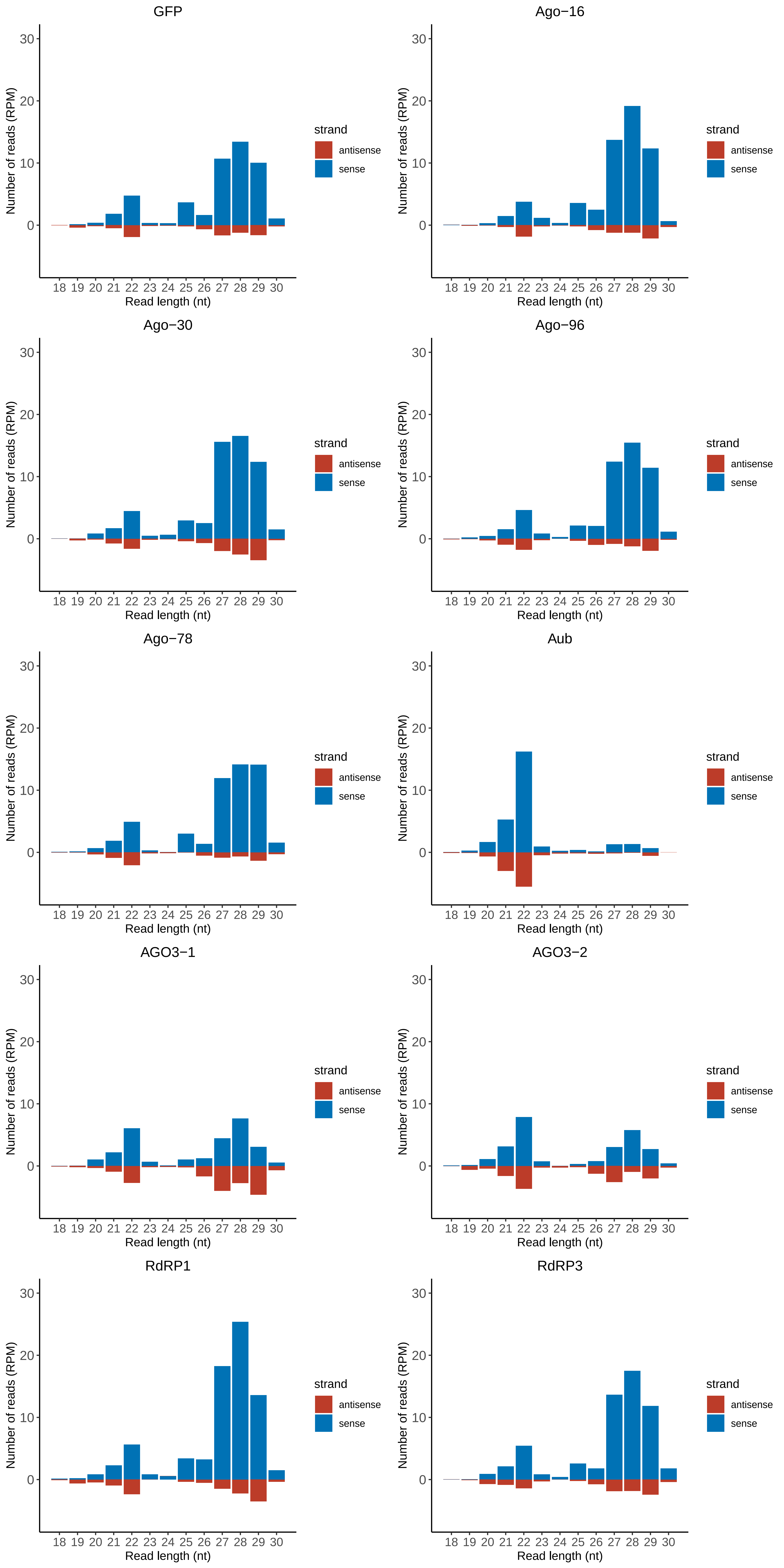

### Motif:rnd-1_family-1792TE_lengthdistribution.pdf

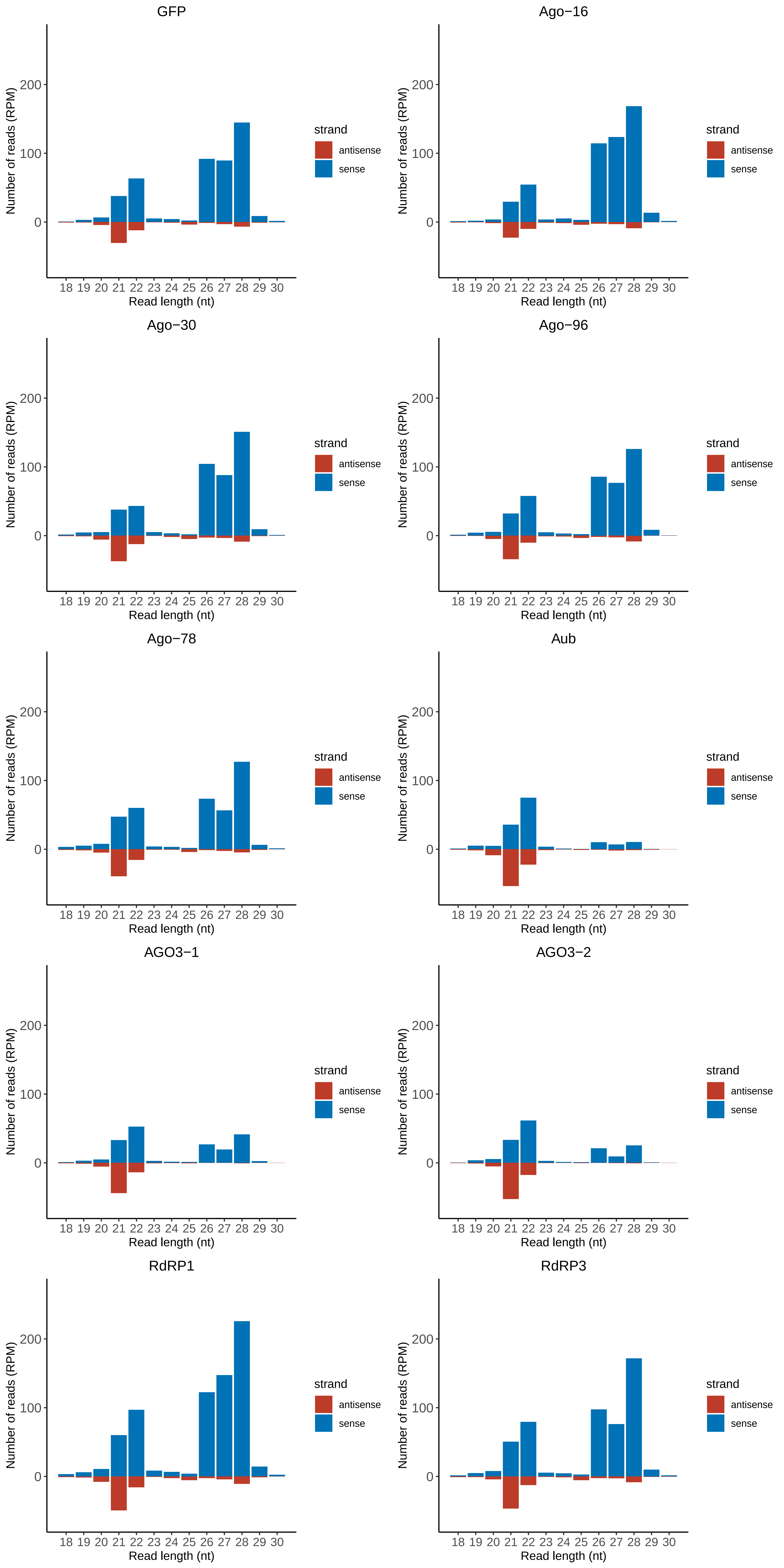

### Motif:rnd-1_family-1869TE_lengthdistribution.pdf

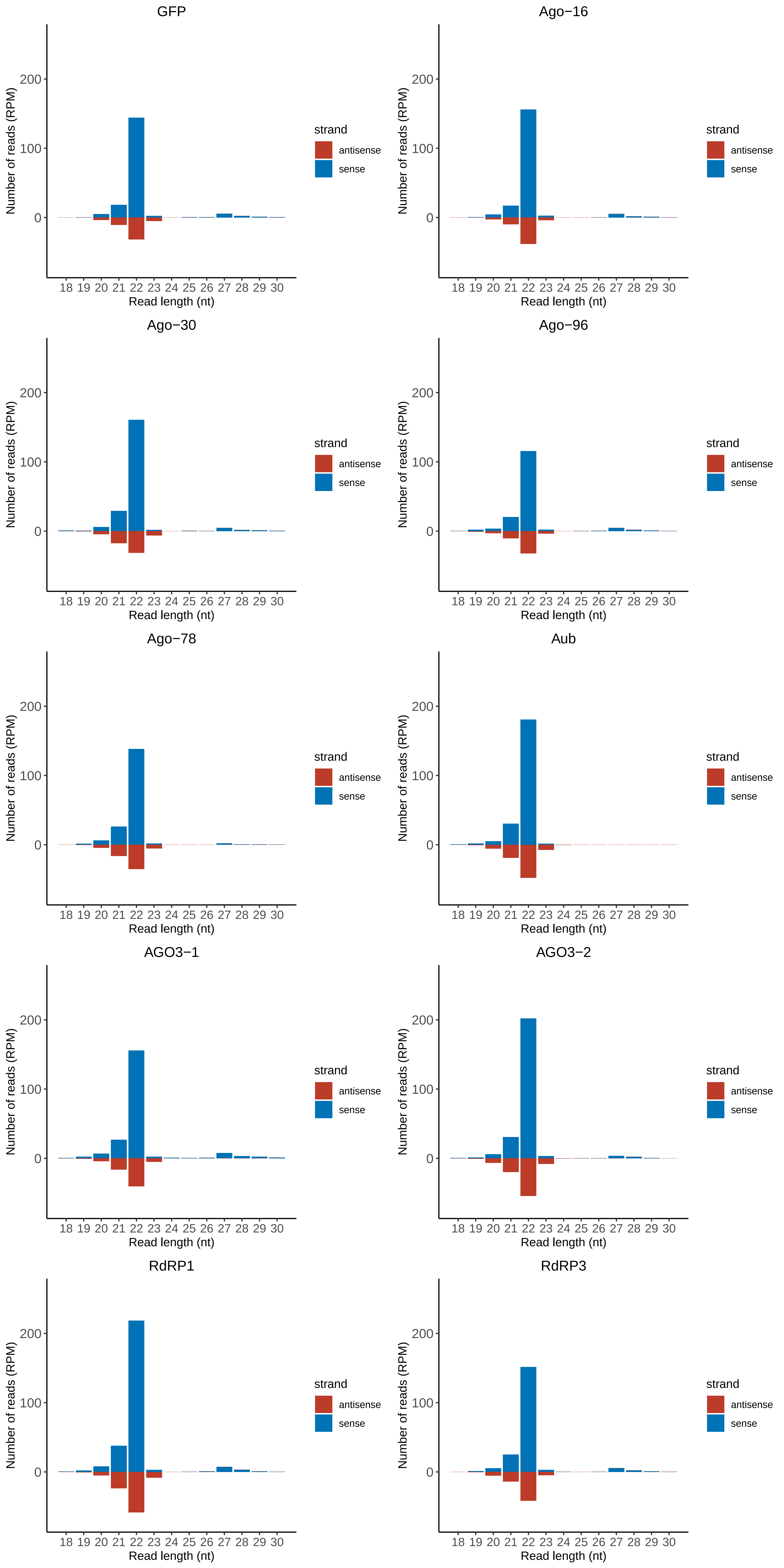

### Motif:rnd-3_family-857TE_lengthdistribution.pdf

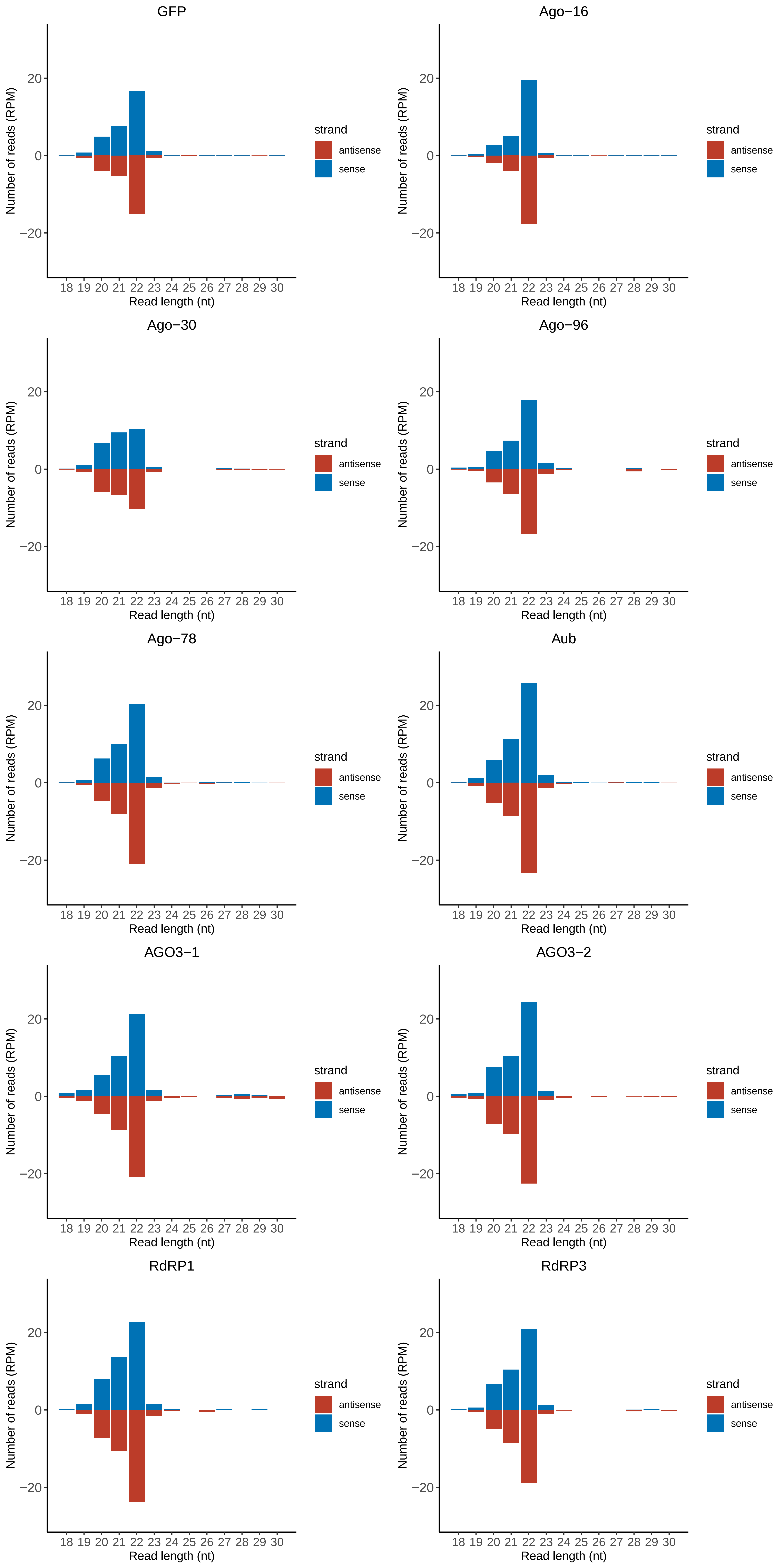

### Motif:rnd-4_family-281TE_lengthdistribution.pdf

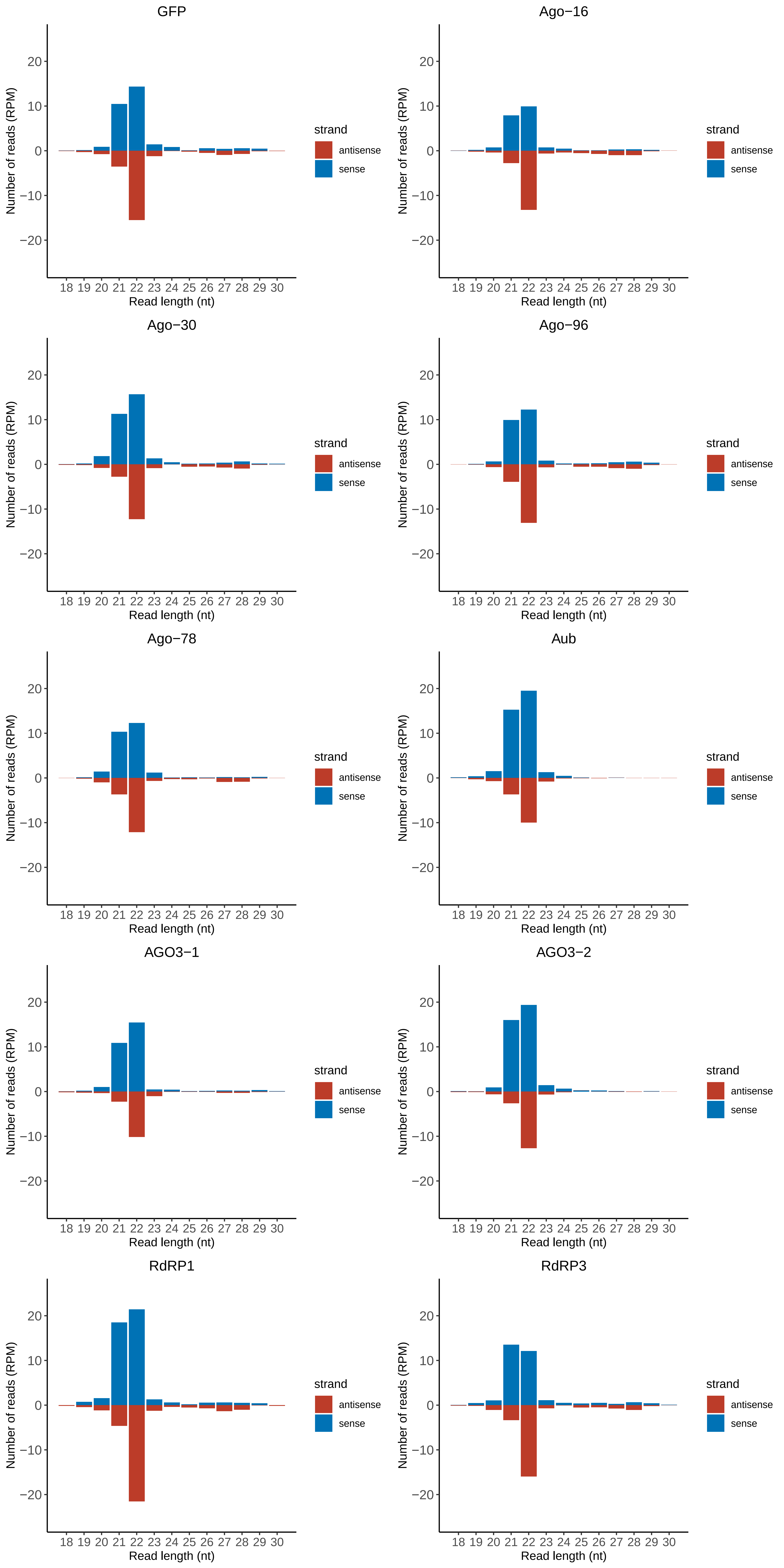

### Motif:rnd-4_family-408TE_lengthdistribution.pdf

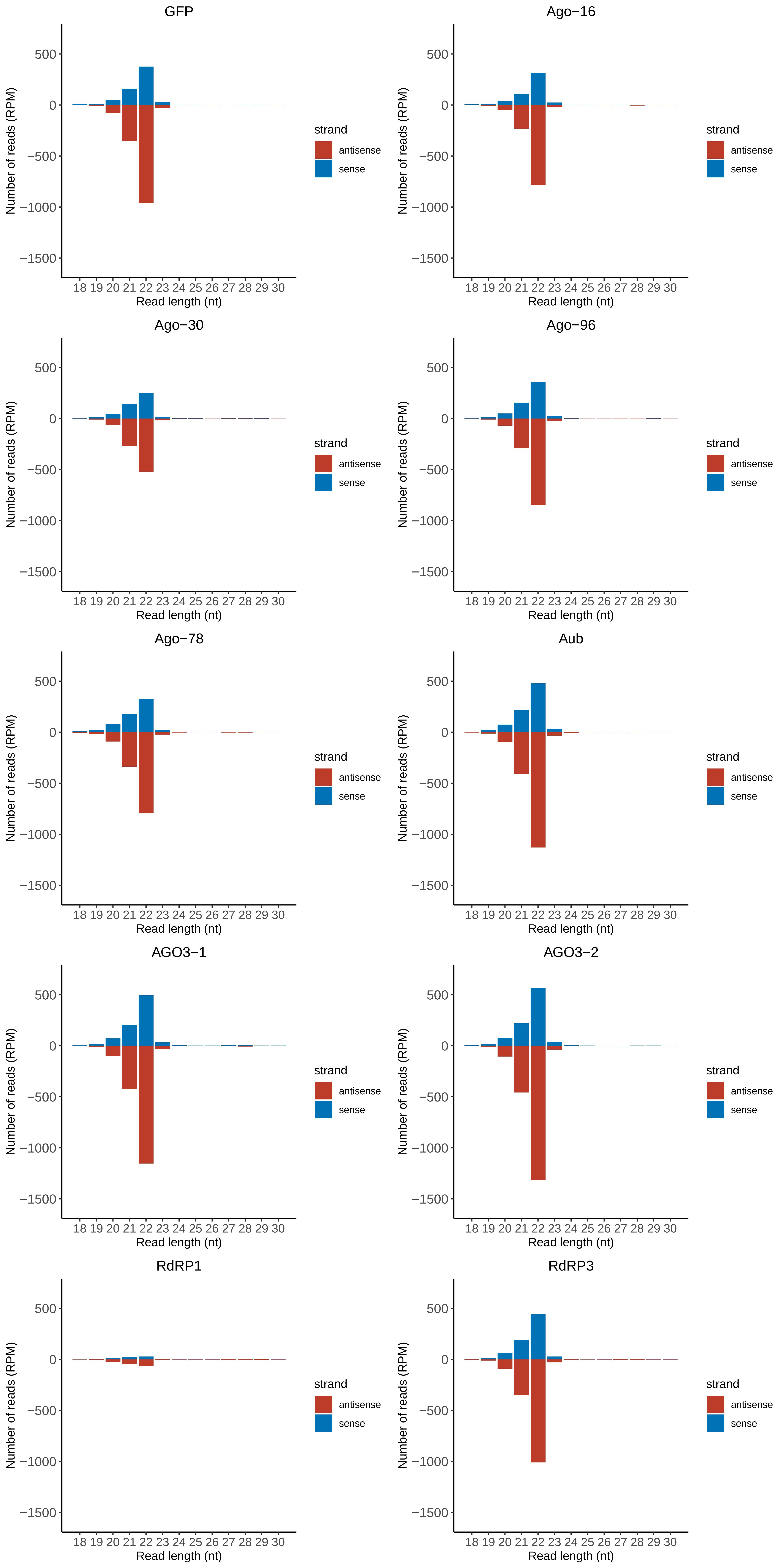

### Motif:rnd-5_family-150TE_lengthdistribution.pdf

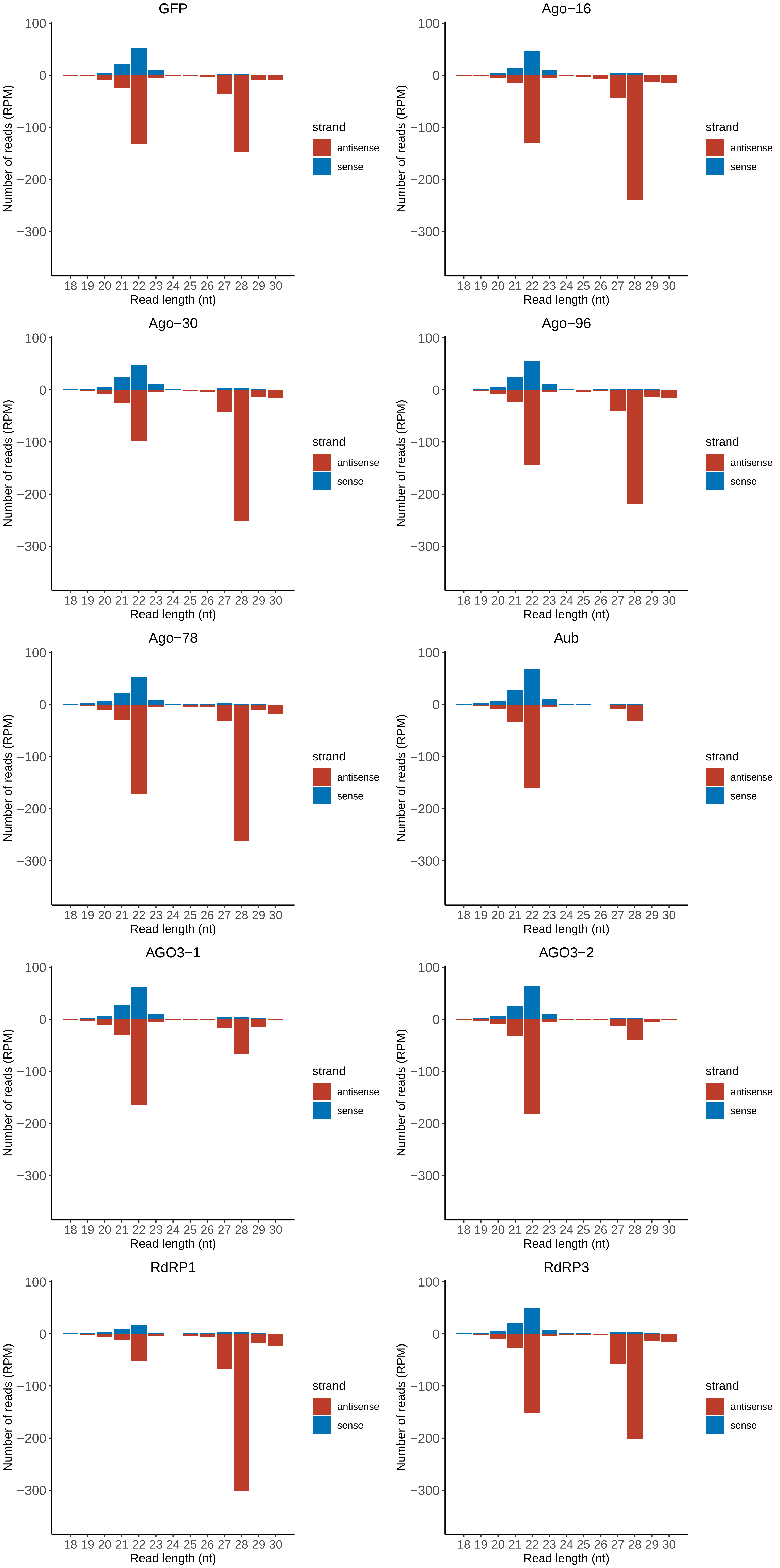

### Motif:rnd-5_family-154TE_lengthdistribution.pdf

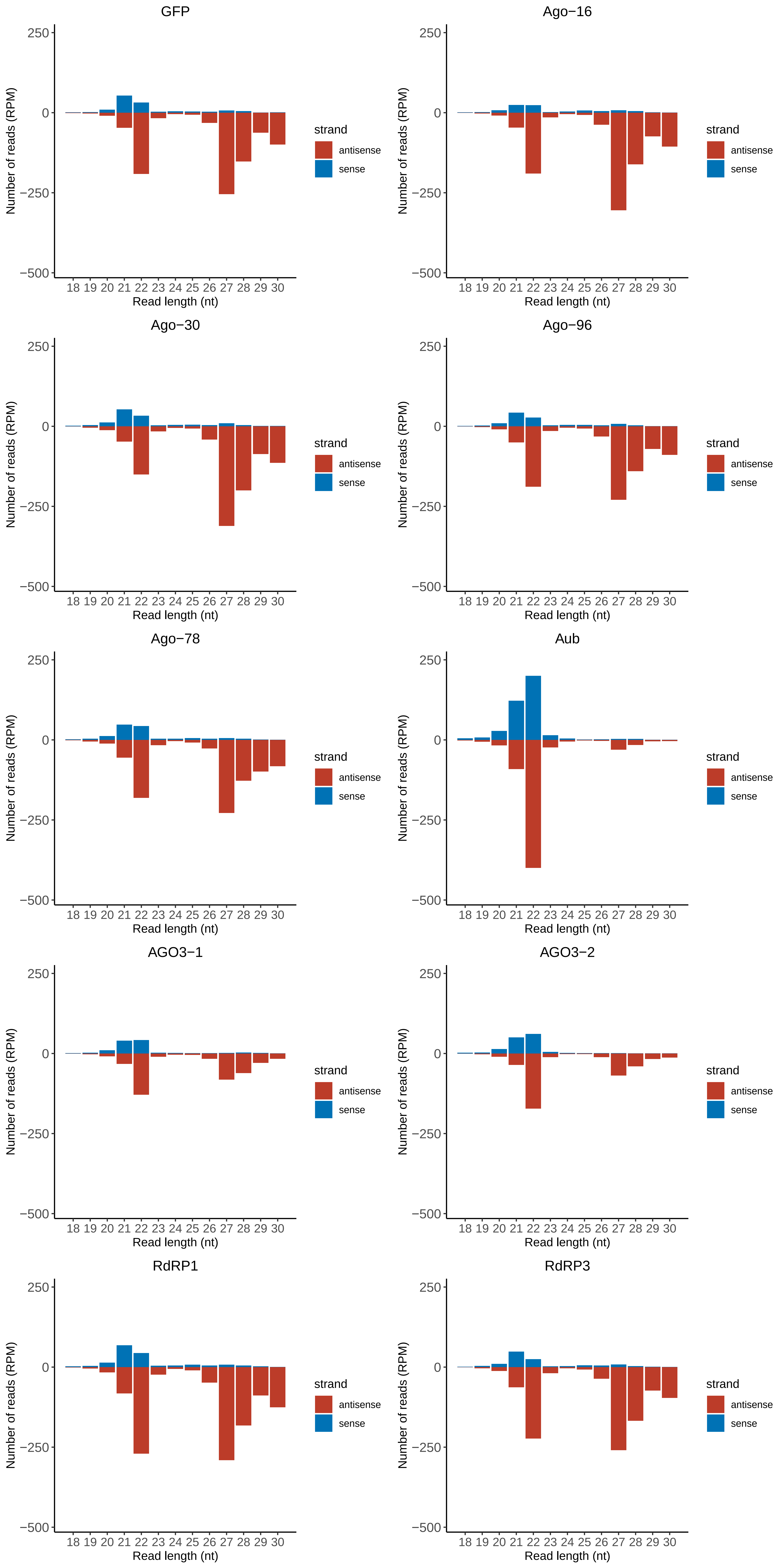

### Motif:rnd-5_family-990TE_lengthdistribution.pdf

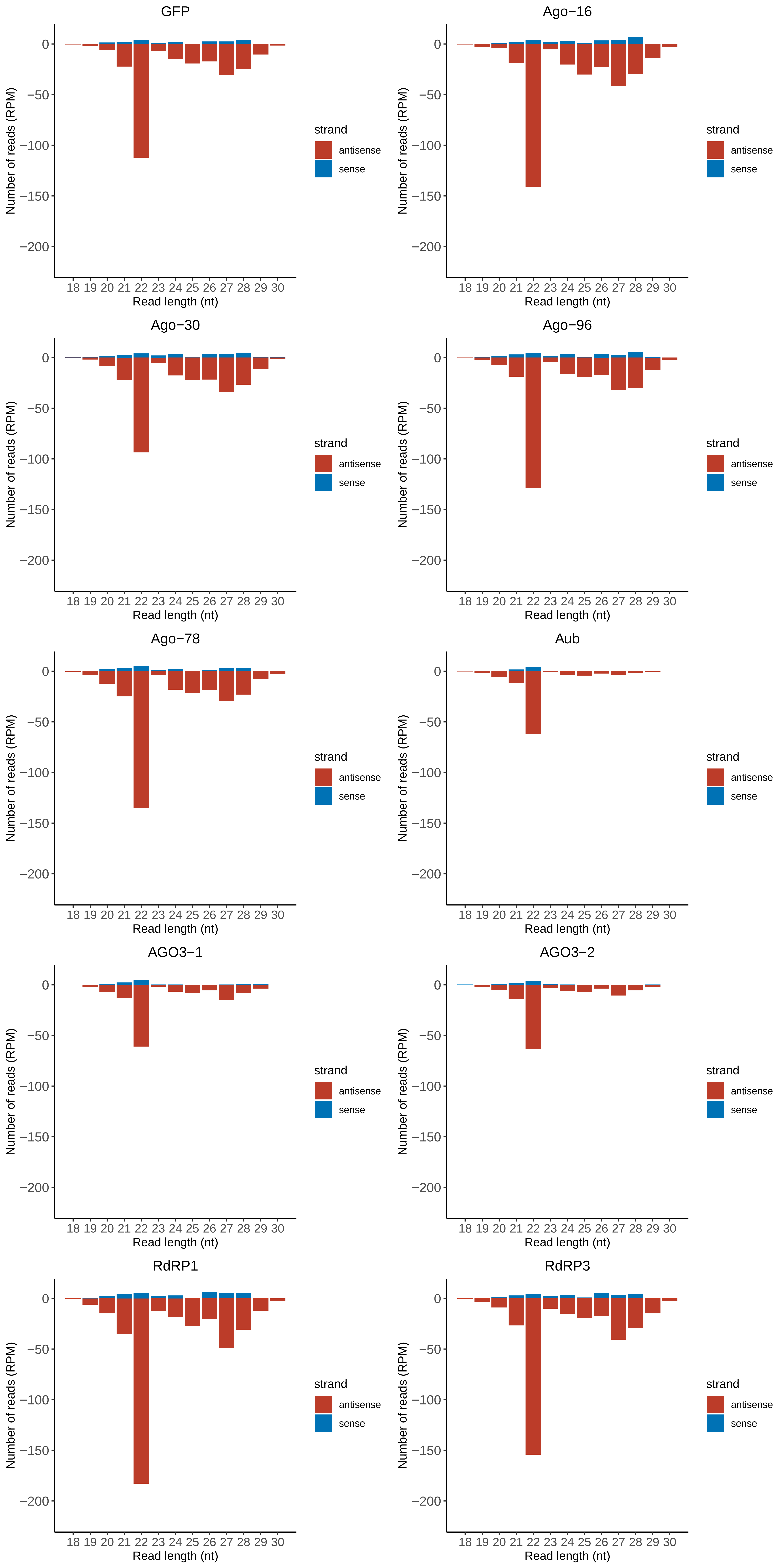

### Motif:rnd-5_family-1090TE_lengthdistribution.pdf

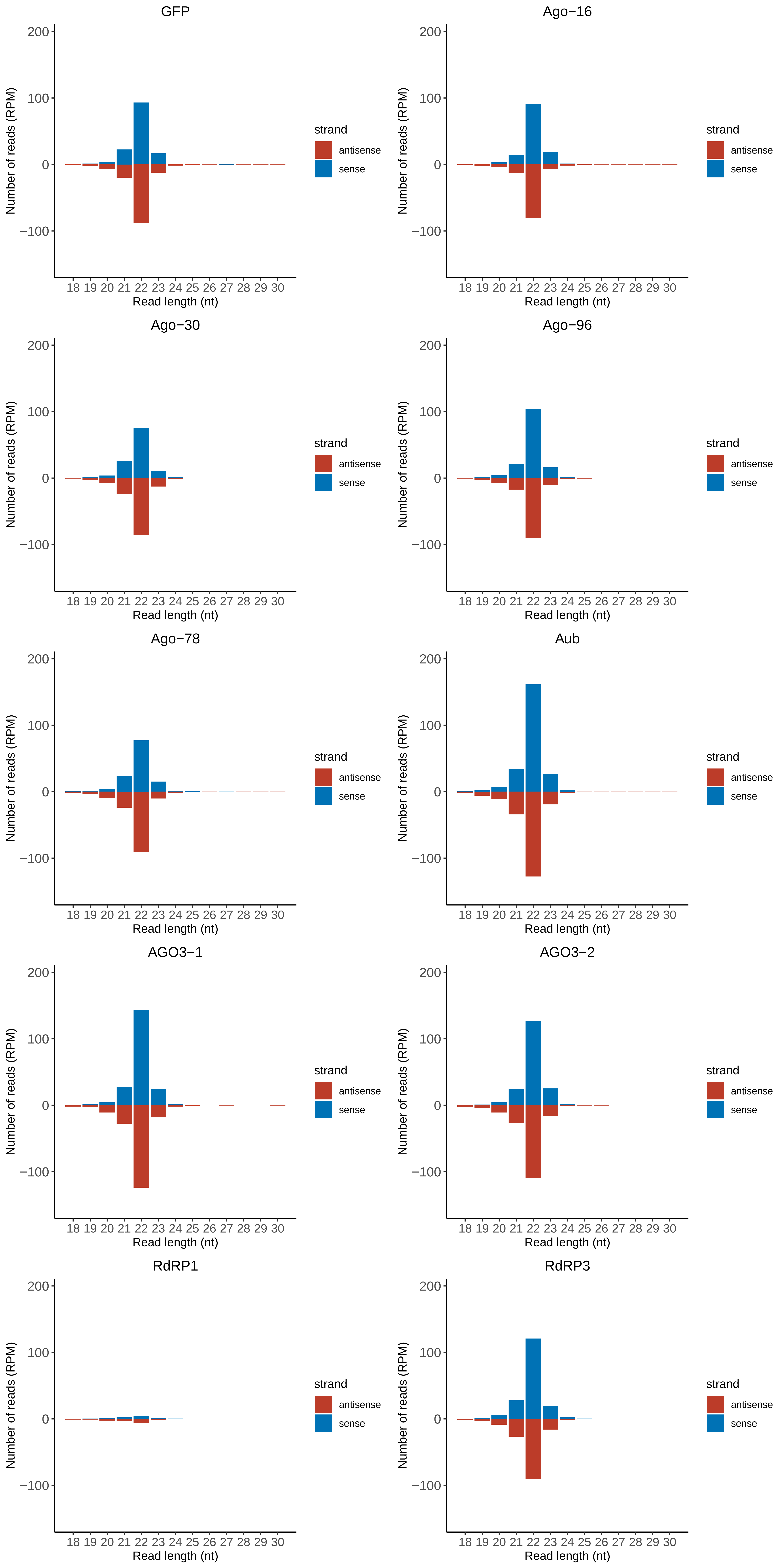

### Motif:rnd-5_family-1420TE_lengthdistribution.pdf

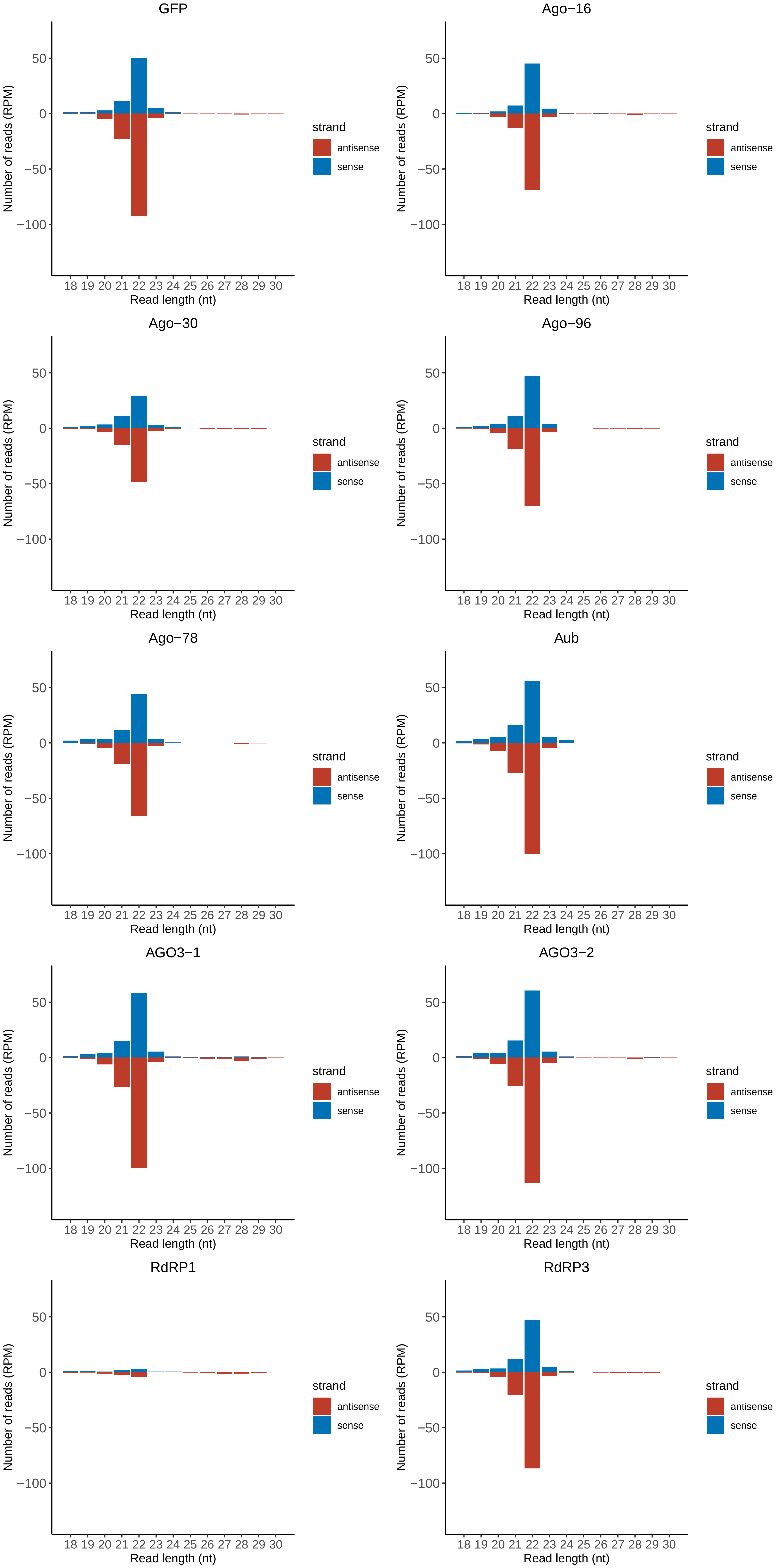

### Motif:rnd-5_family-1630TE_lengthdistribution.pdf

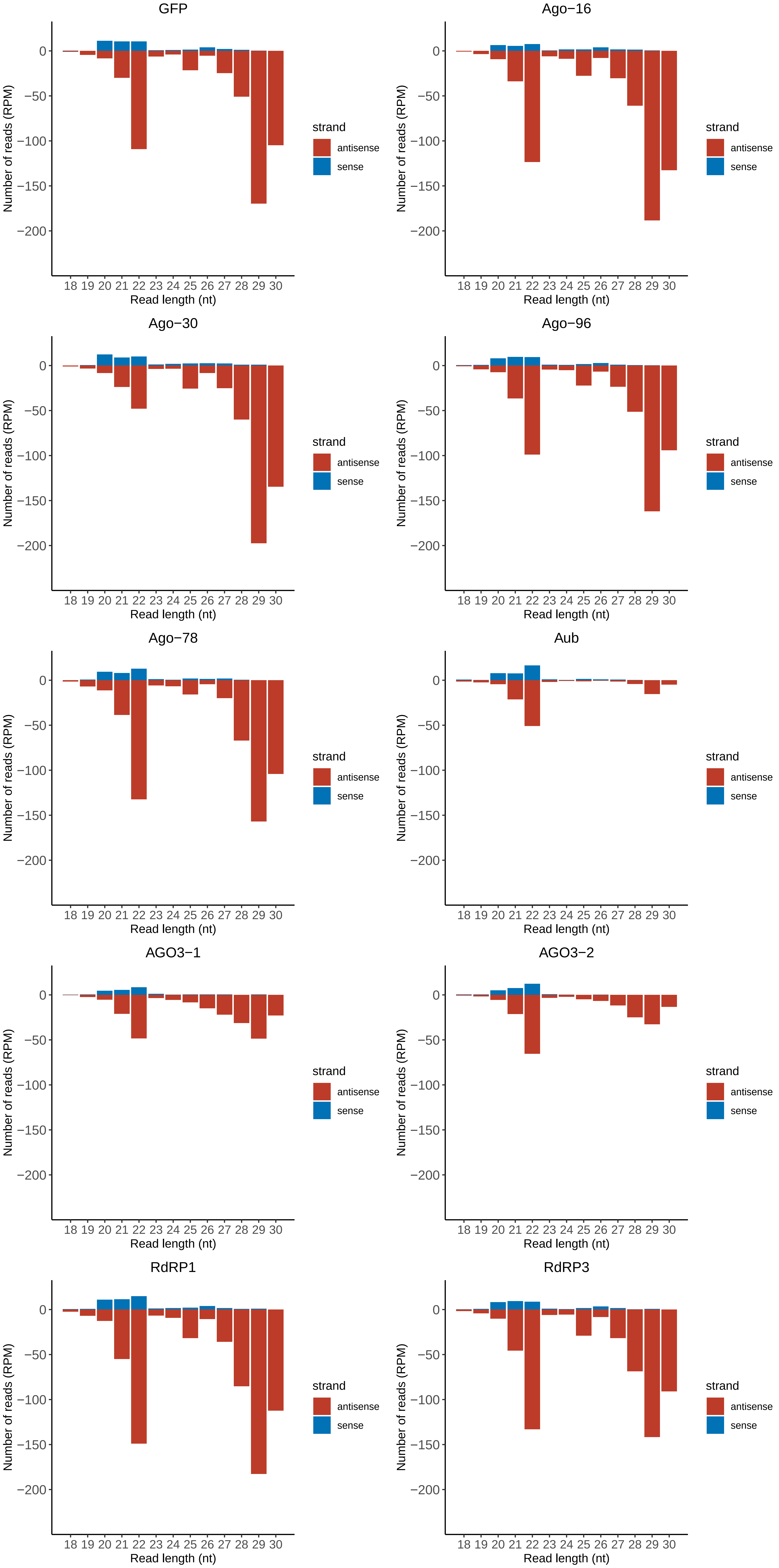

### Motif:rnd-5_family-1764TE_lengthdistribution.pdf

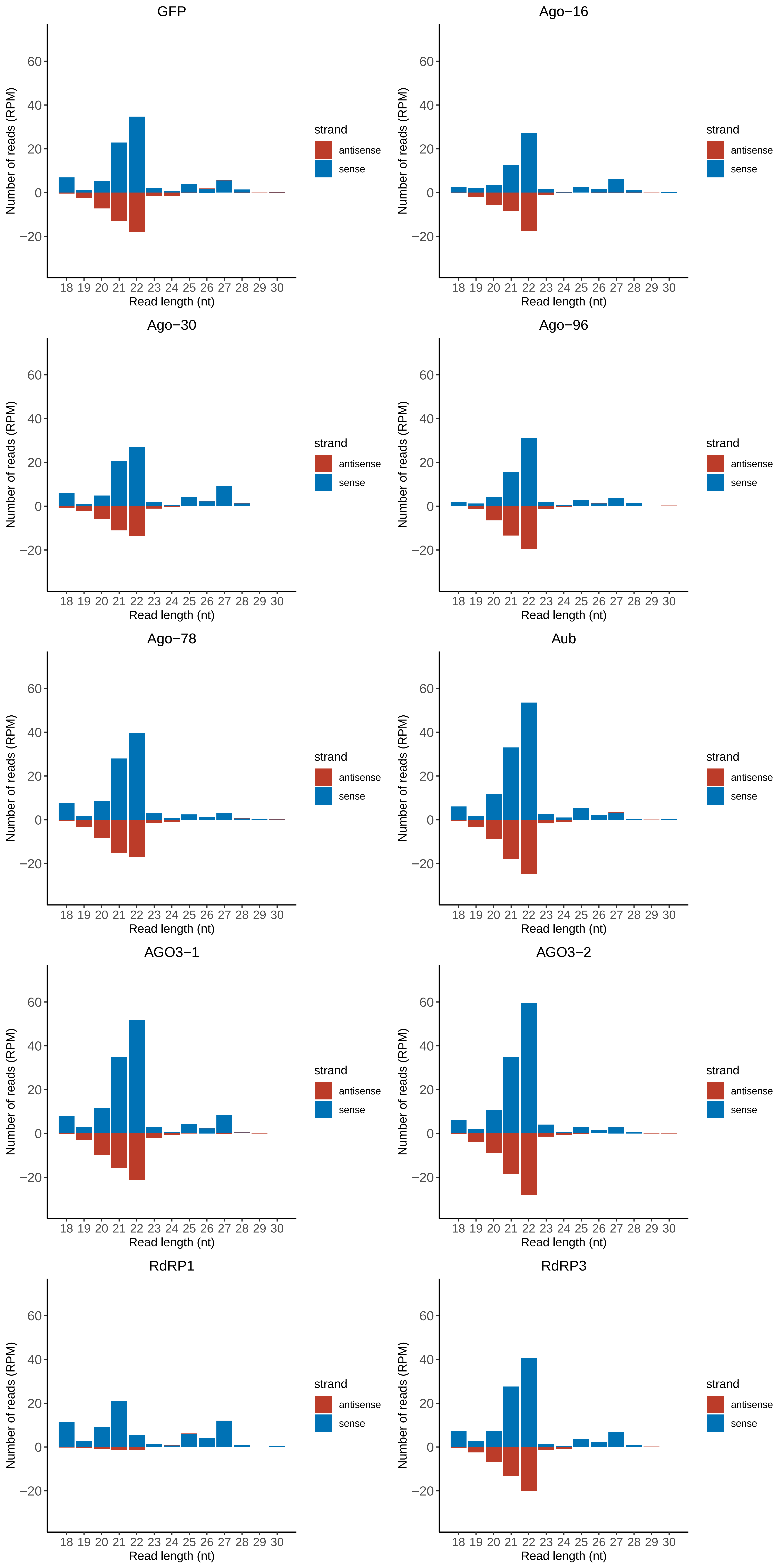

### Motif:rnd-5_family-2004TE_lengthdistribution.pdf

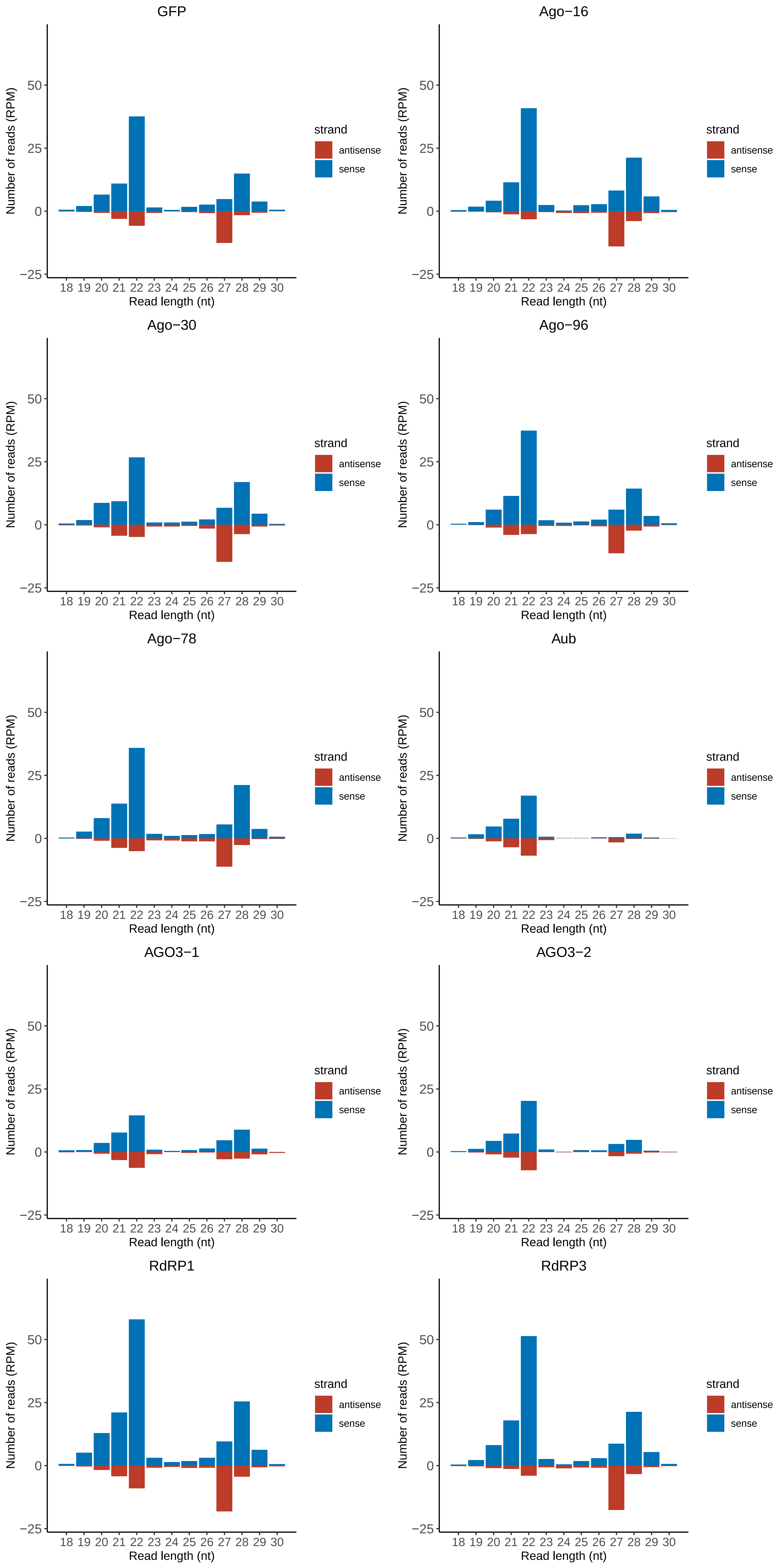

### Motif:rnd-5_family-2012TE_lengthdistribution.pdf

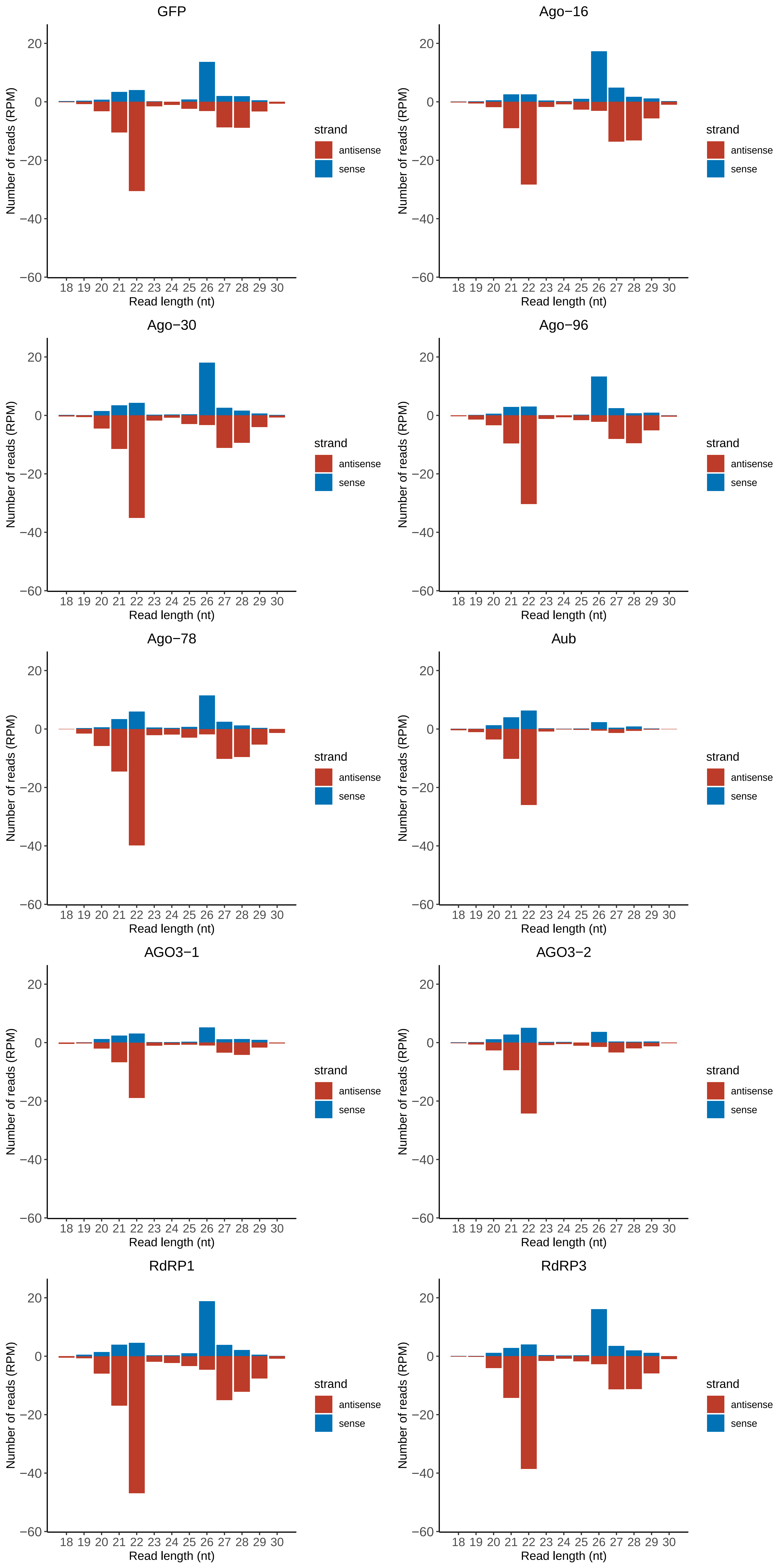

### Motif:rnd-5_family-2193TE_lengthdistribution.pdf

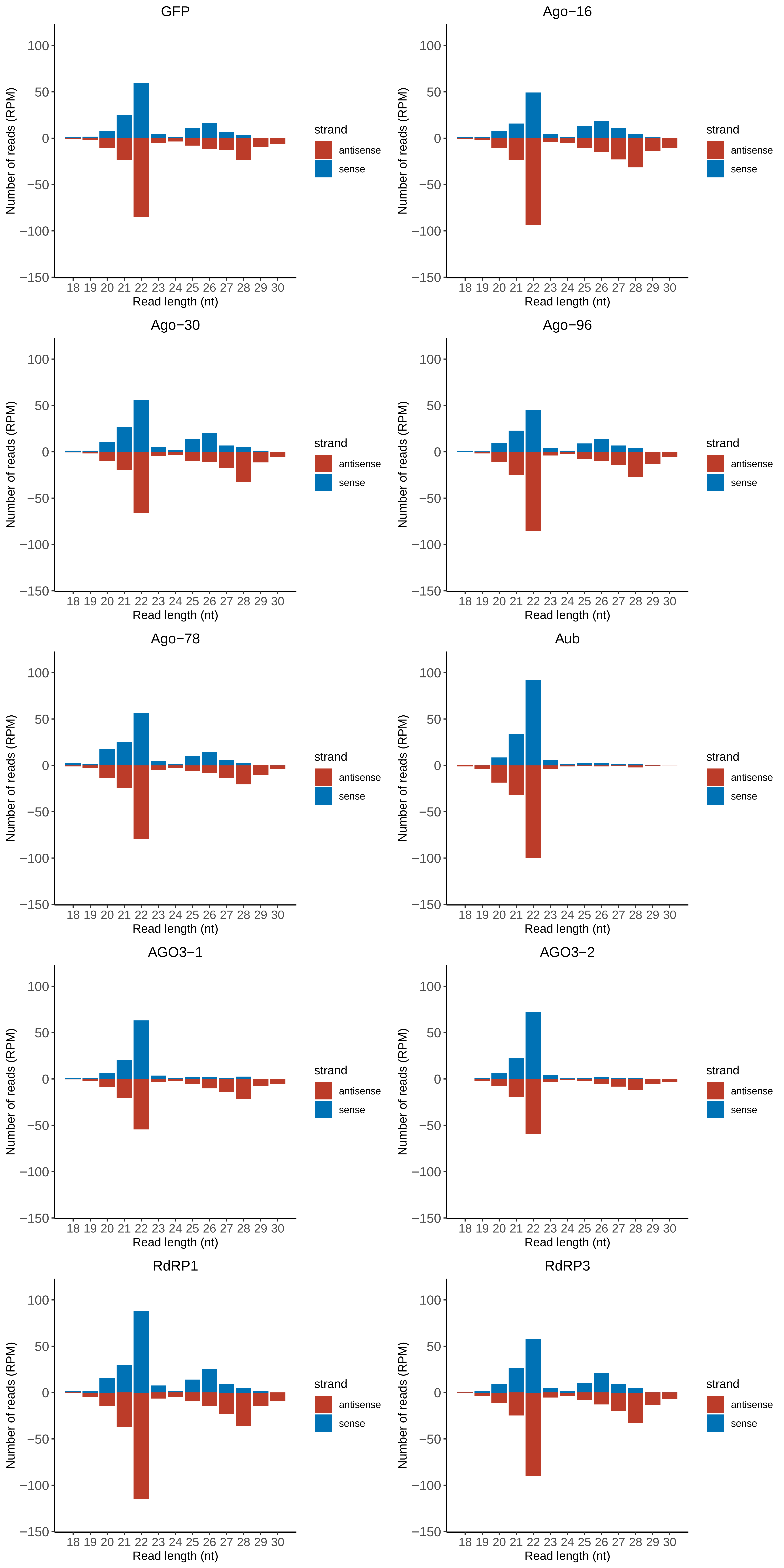

### Motif:rnd-5_family-2257TE_lengthdistribution.pdf

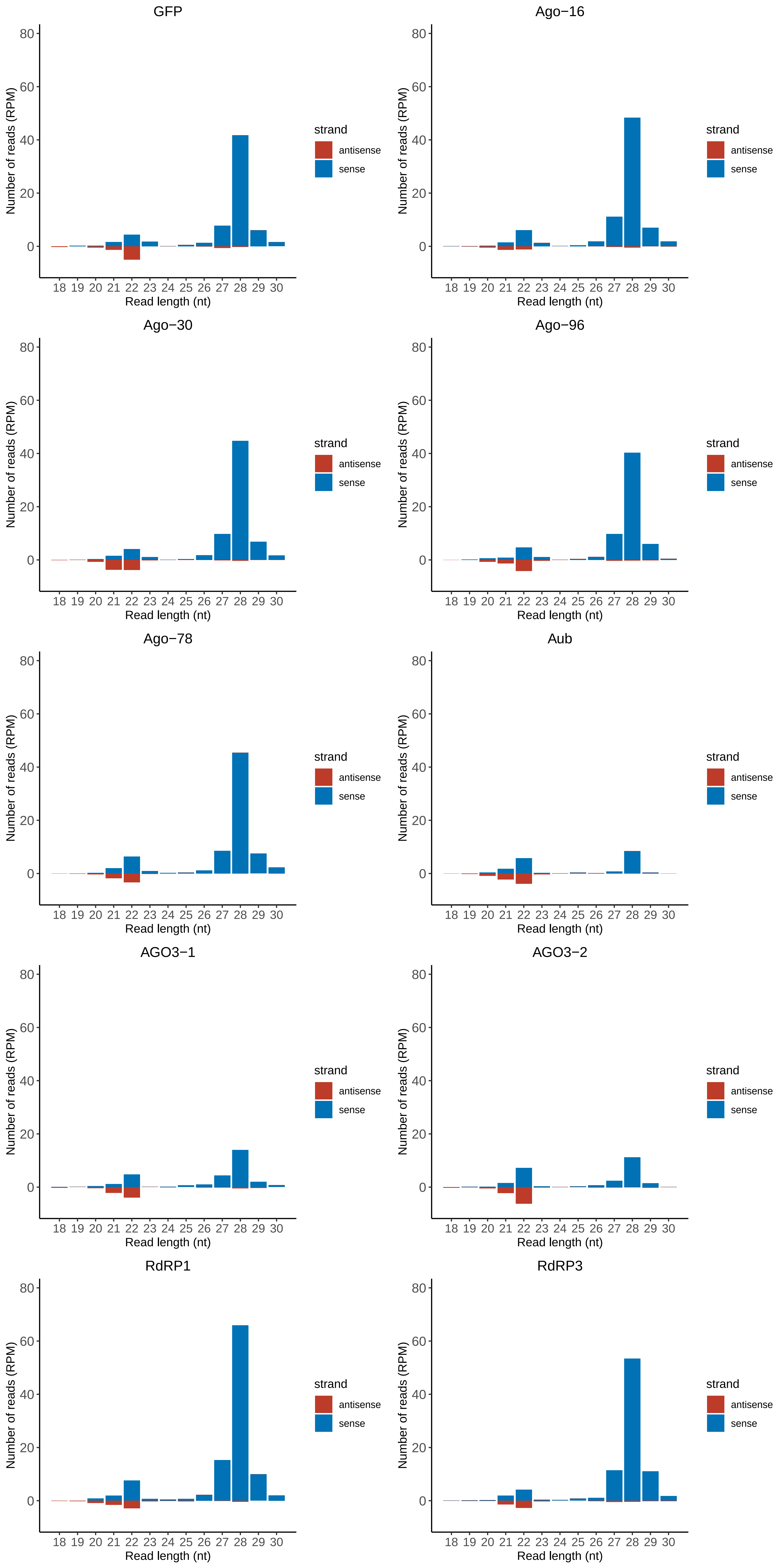

### Motif:rnd-5_family-2803TE_lengthdistribution.pdf

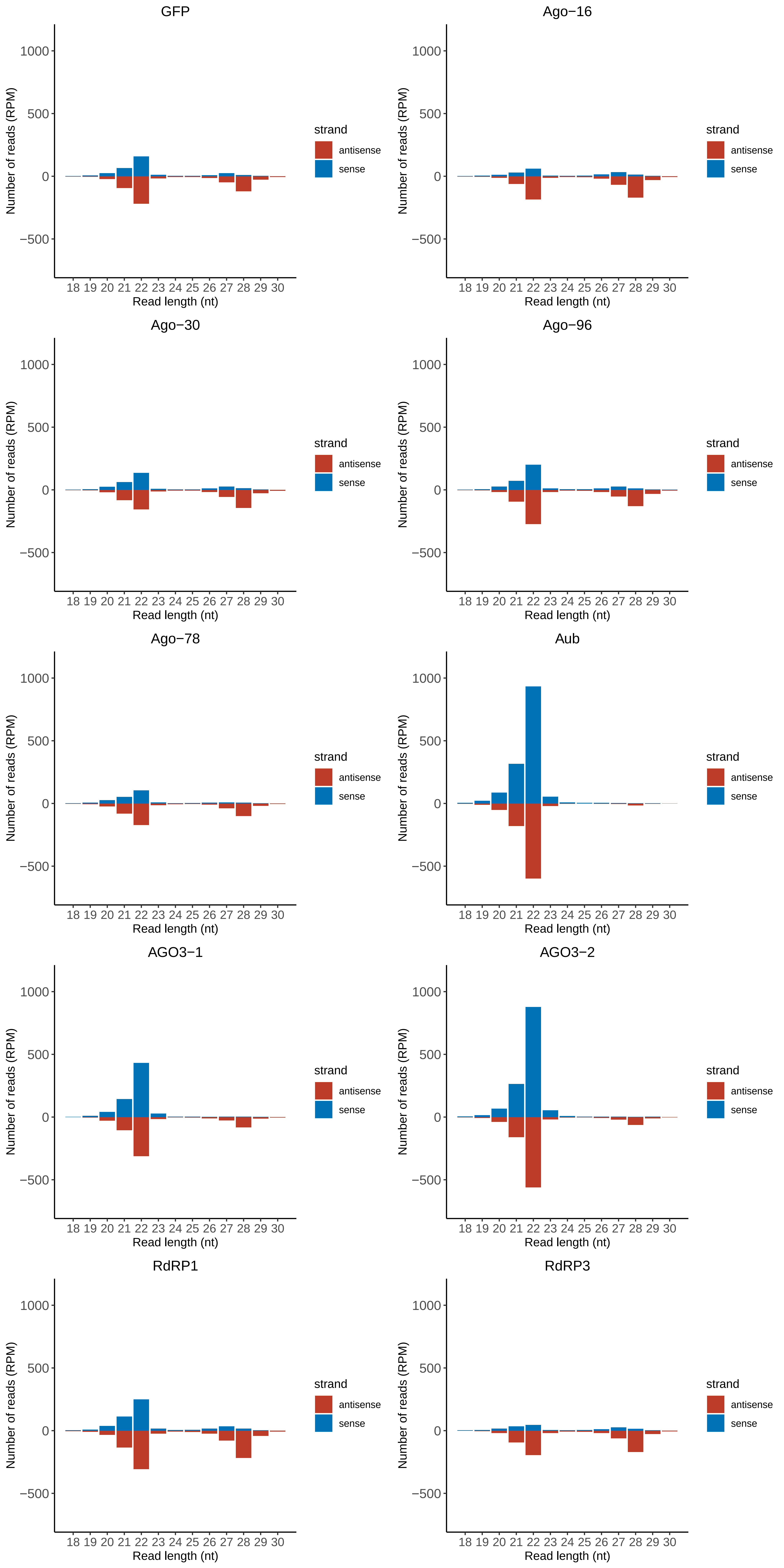

### Motif:rnd-5_family-2877TE_lengthdistribution.pdf

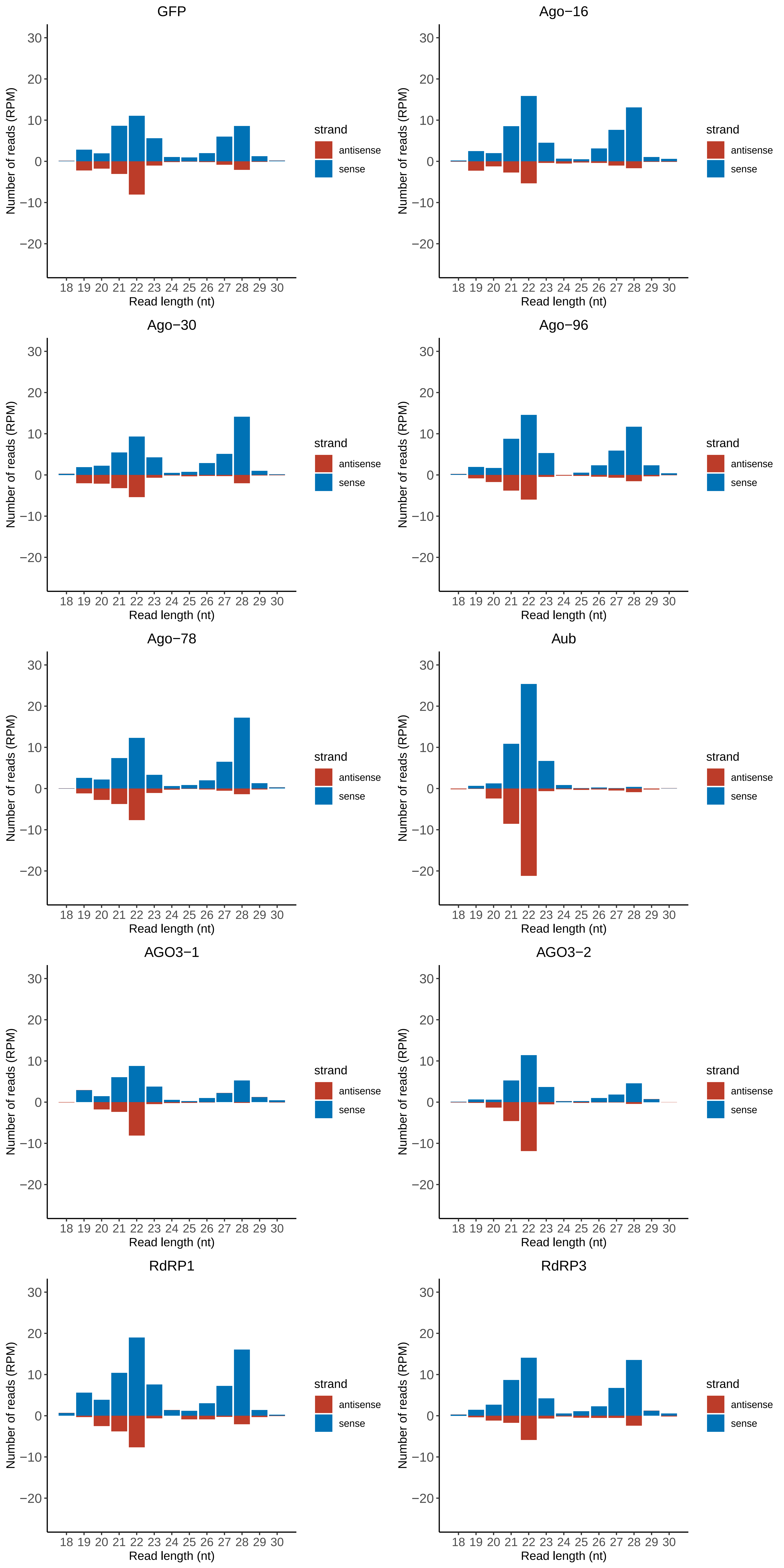

### Motif:rnd-5_family-3705TE_lengthdistribution.pdf

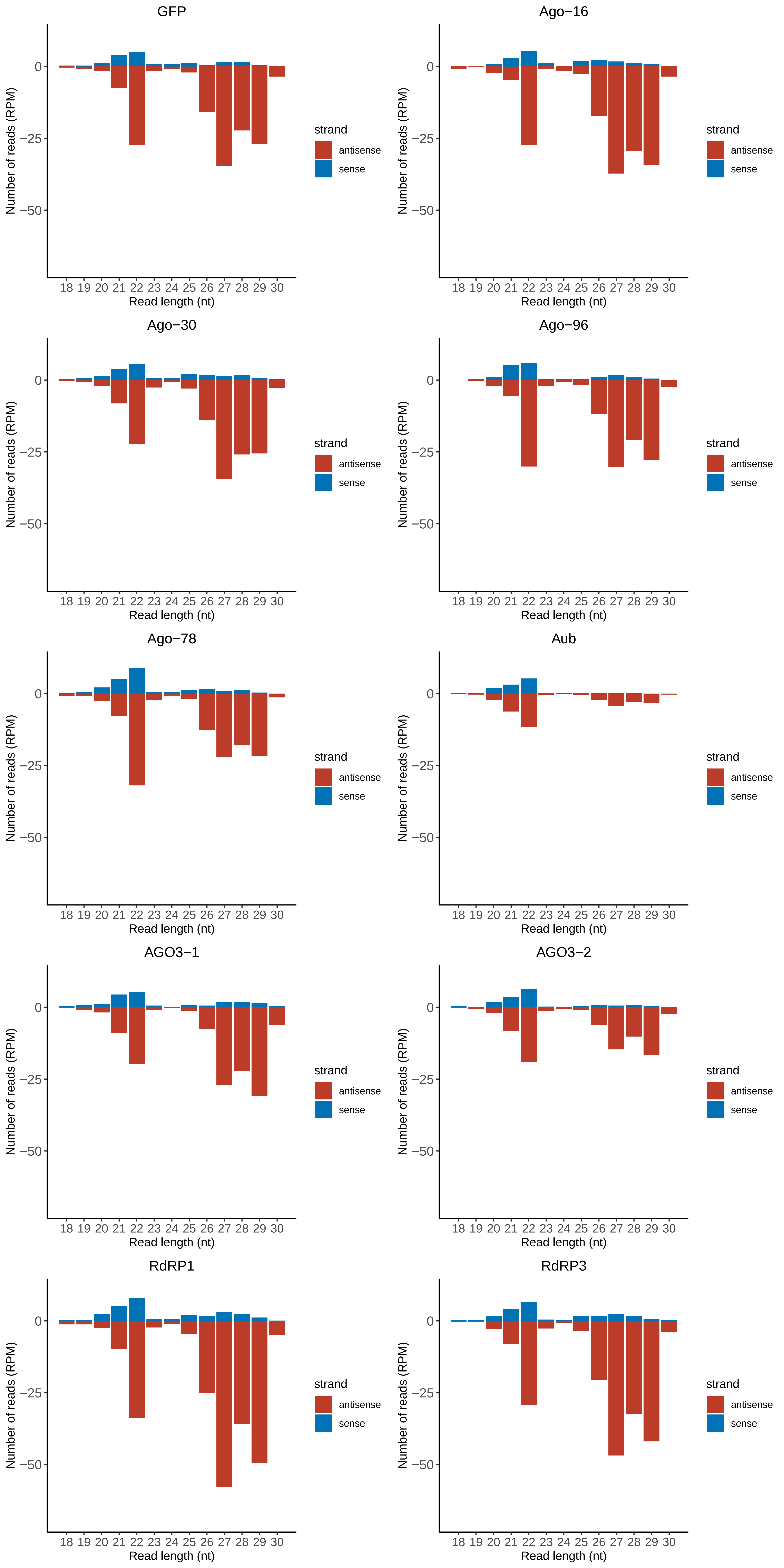

### Motif:rnd-5_family-4863TE_lengthdistribution.pdf

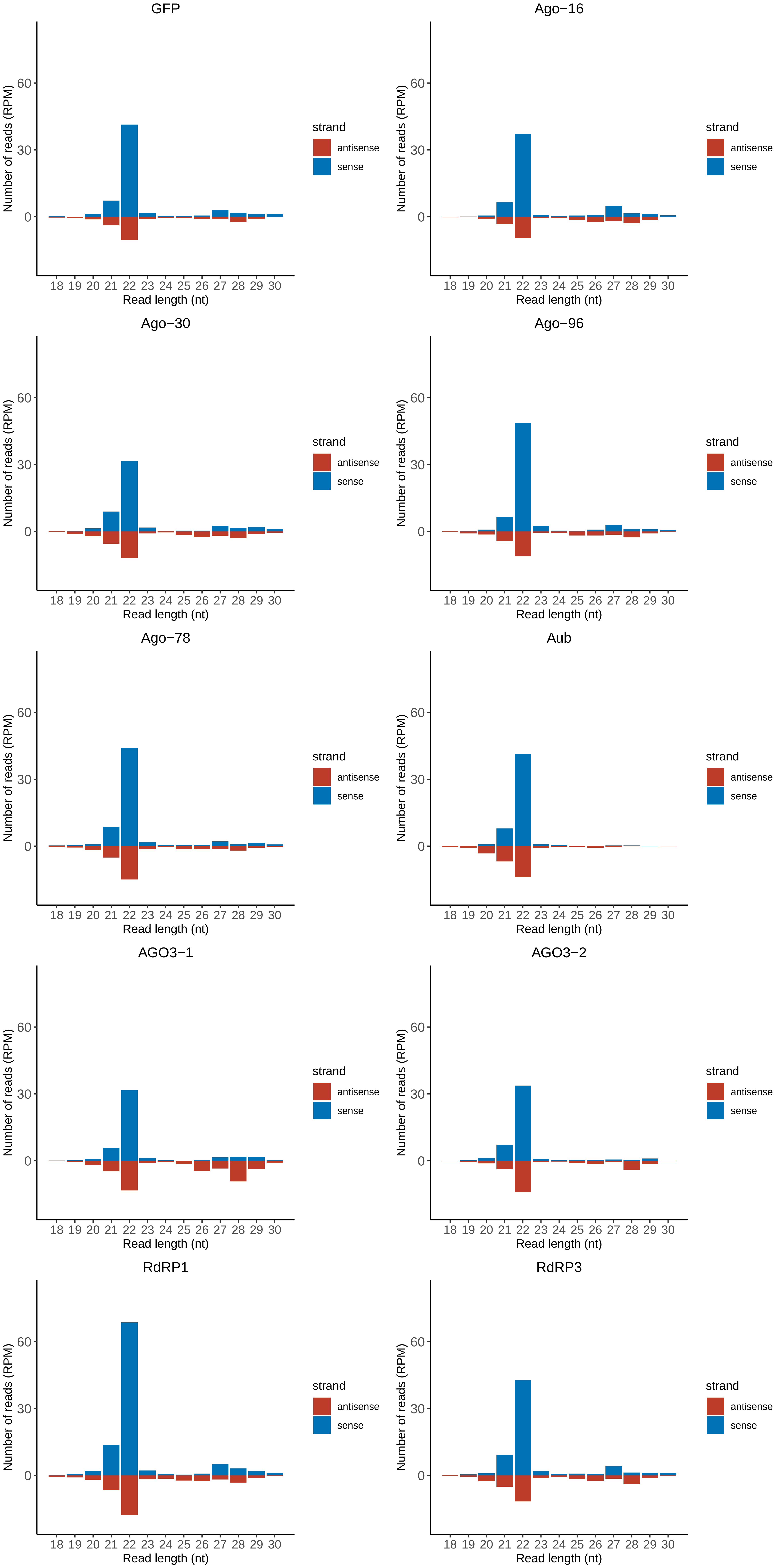

### Motif:rnd-5_family-5765TE_lengthdistribution.pdf

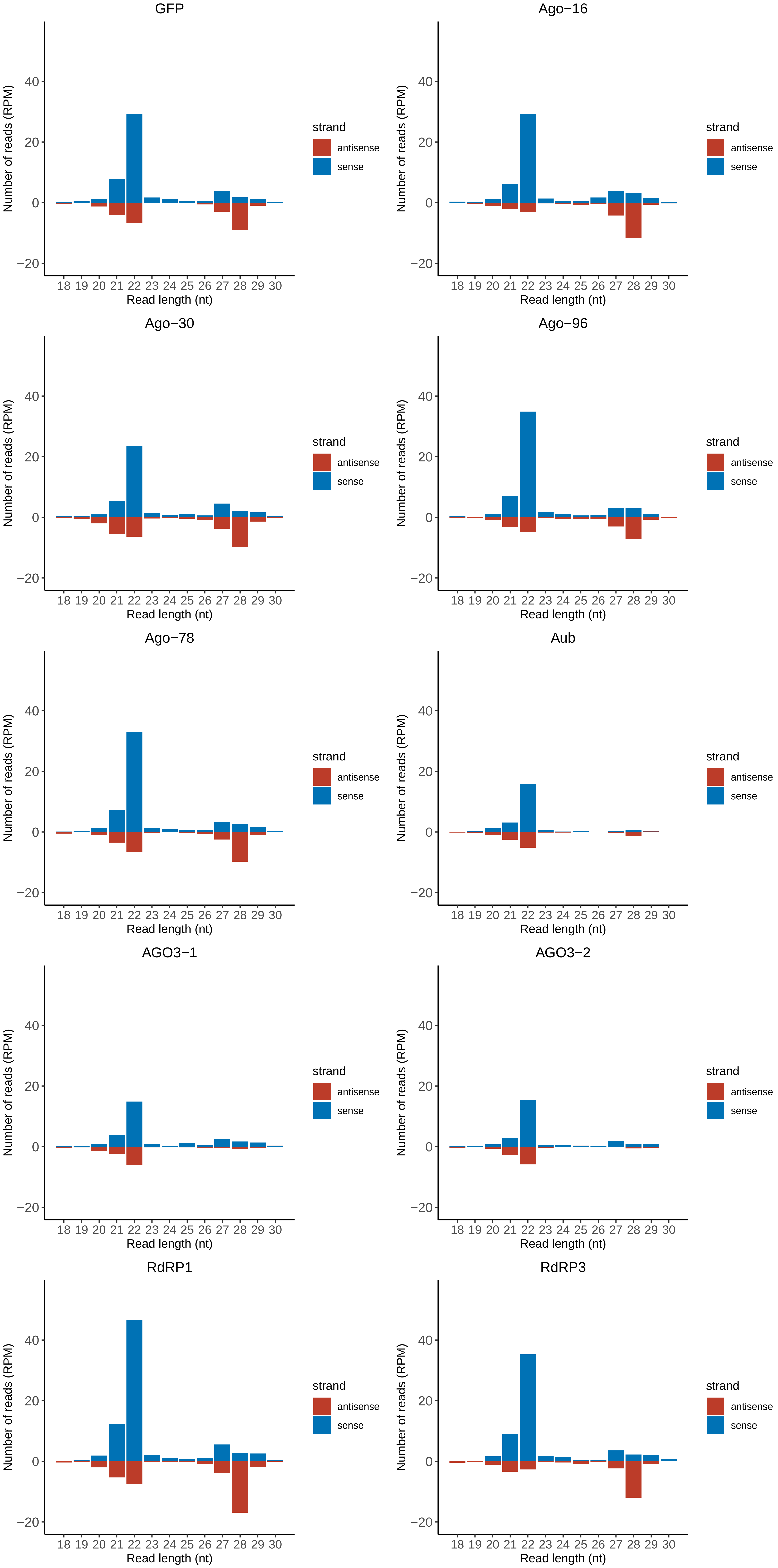

### Motif:rnd-5_family-5812TE_lengthdistribution.pdf

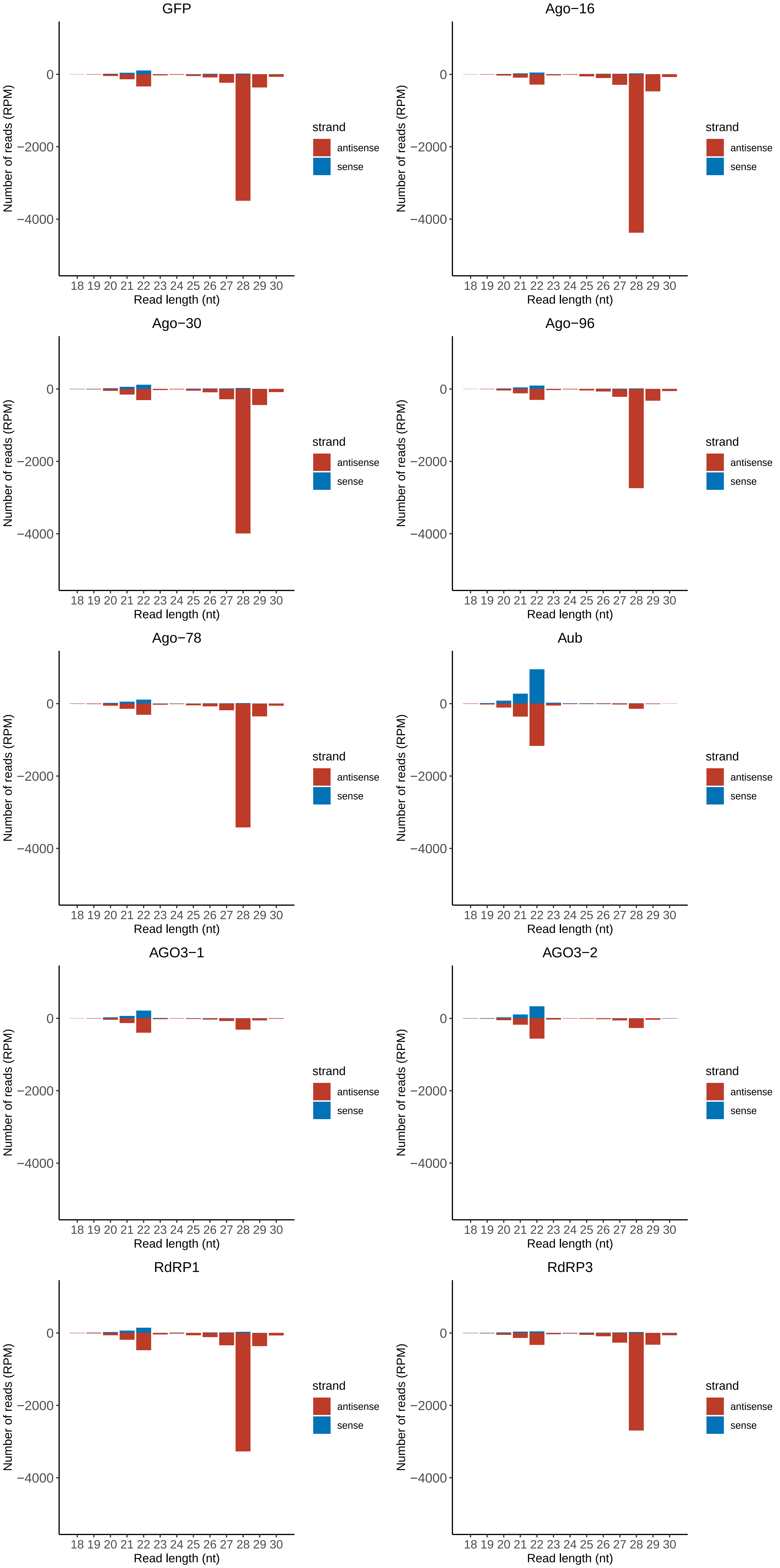

### Motif:rnd-5_family-7700TE_lengthdistribution.pdf

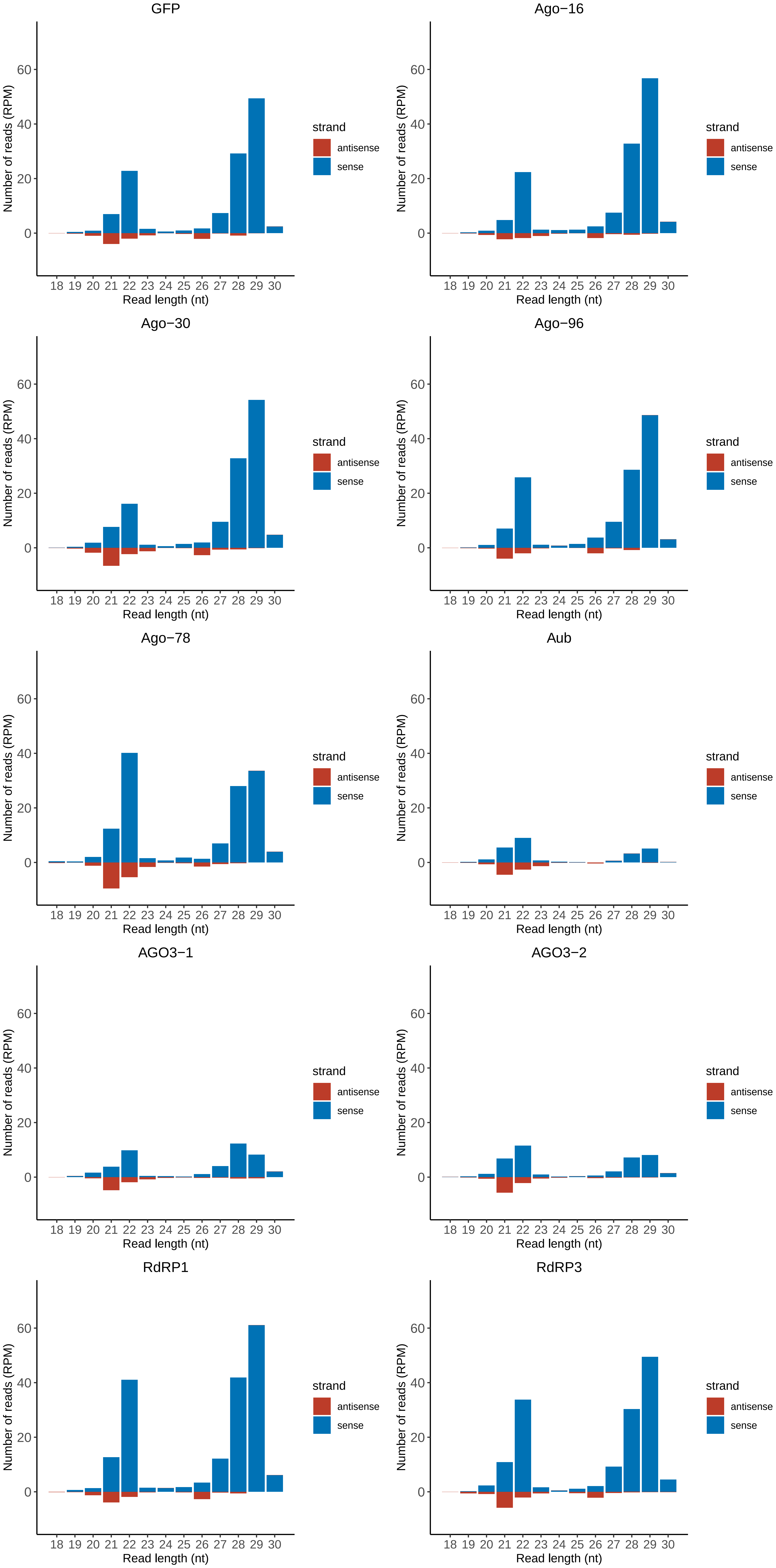

### Motif:rnd-6_family-46TE_lengthdistribution.pdf

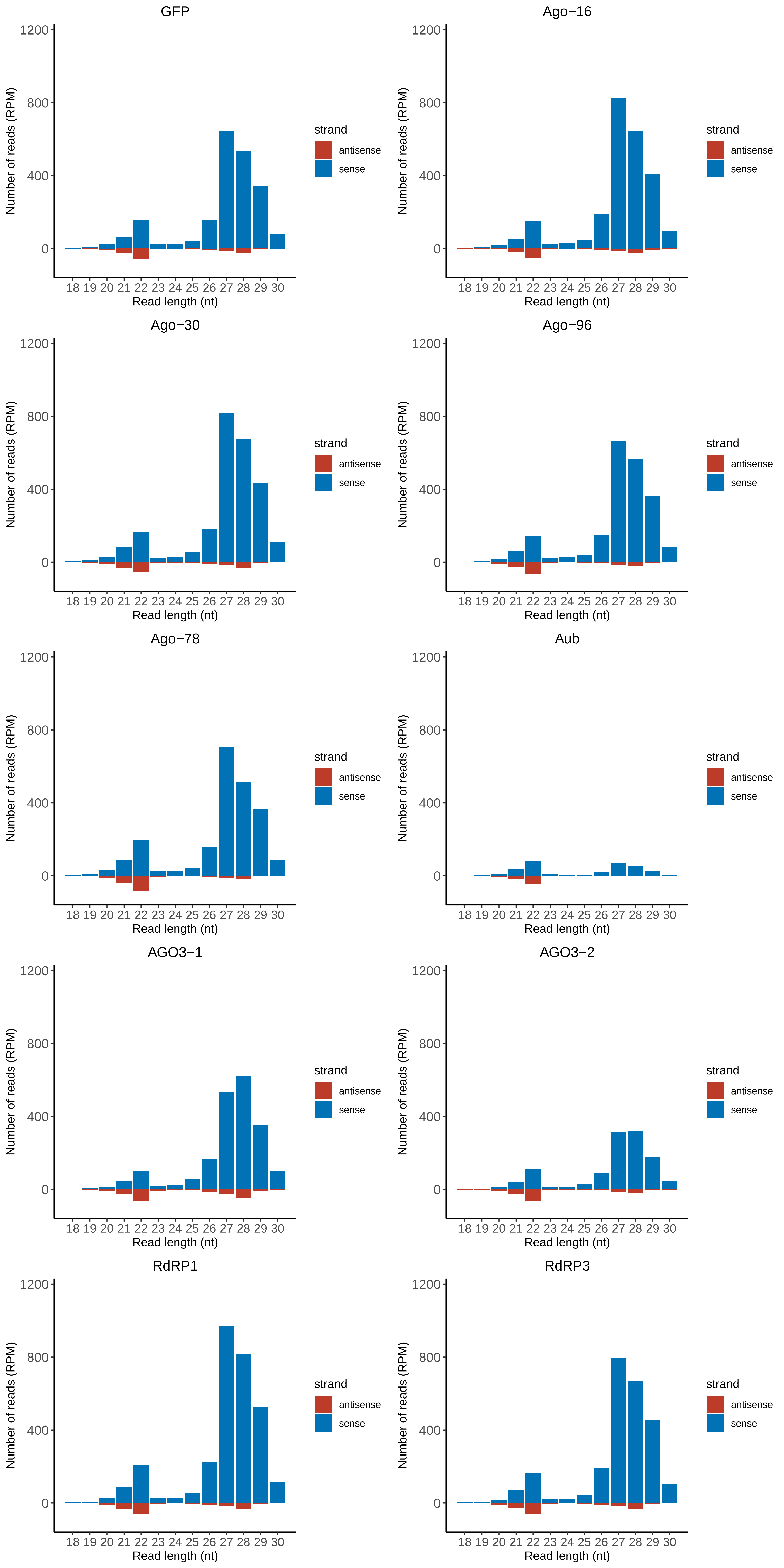

### Motif:rnd-6_family-58TE_lengthdistribution.pdf

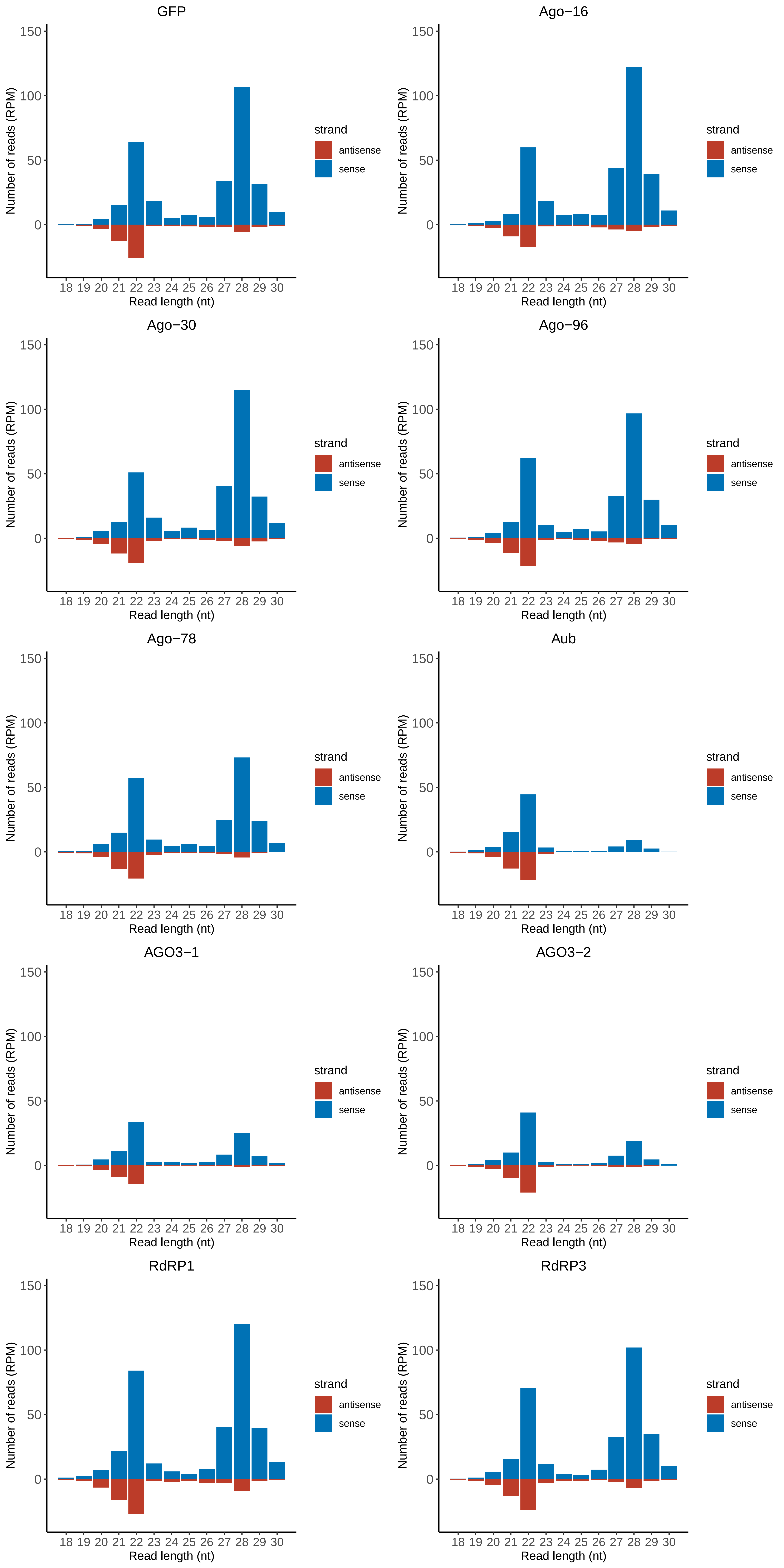

### Motif:rnd-6_family-71TE_lengthdistribution.pdf

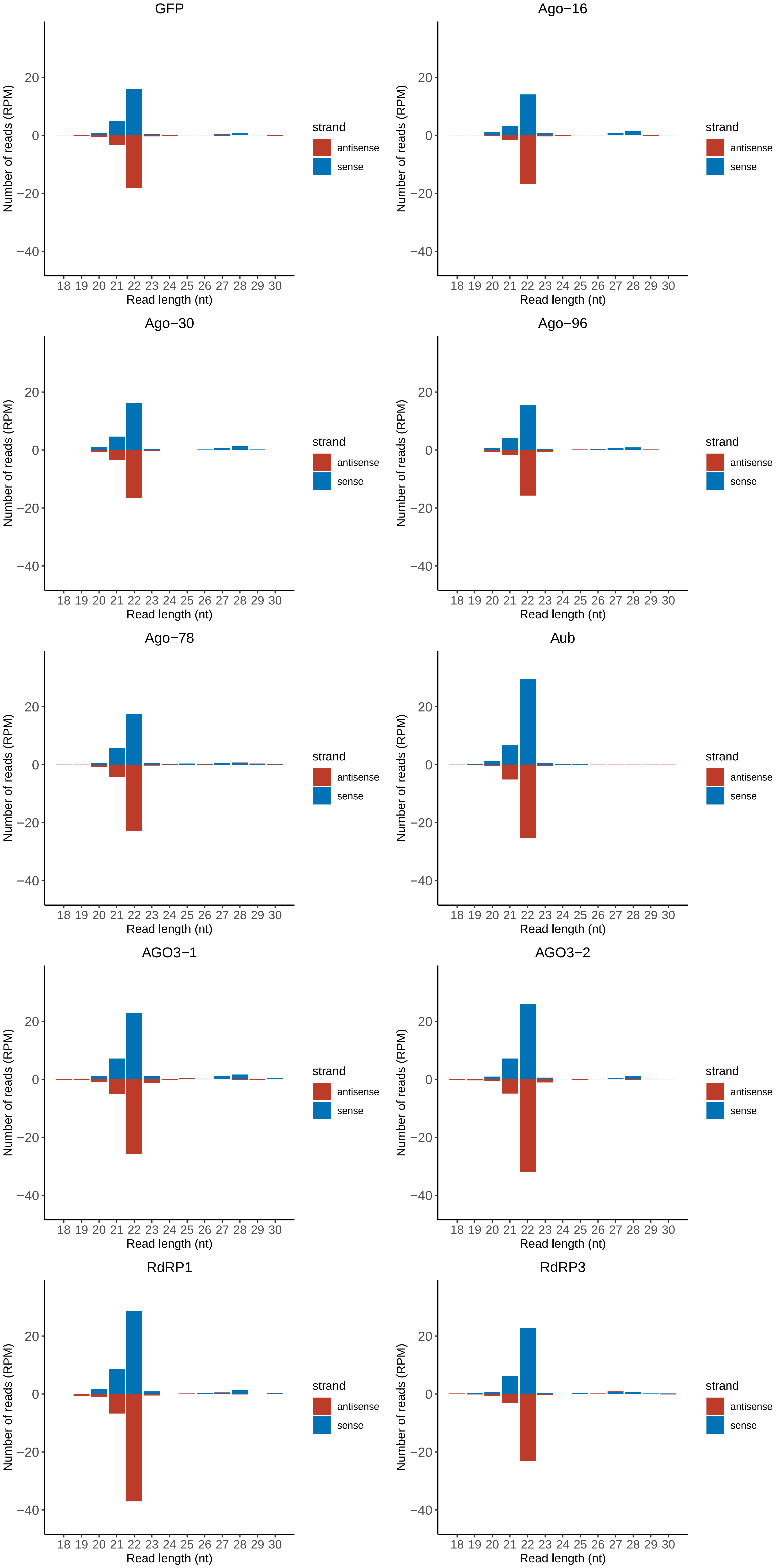

### Motif:rnd-6_family-75TE_lengthdistribution.pdf

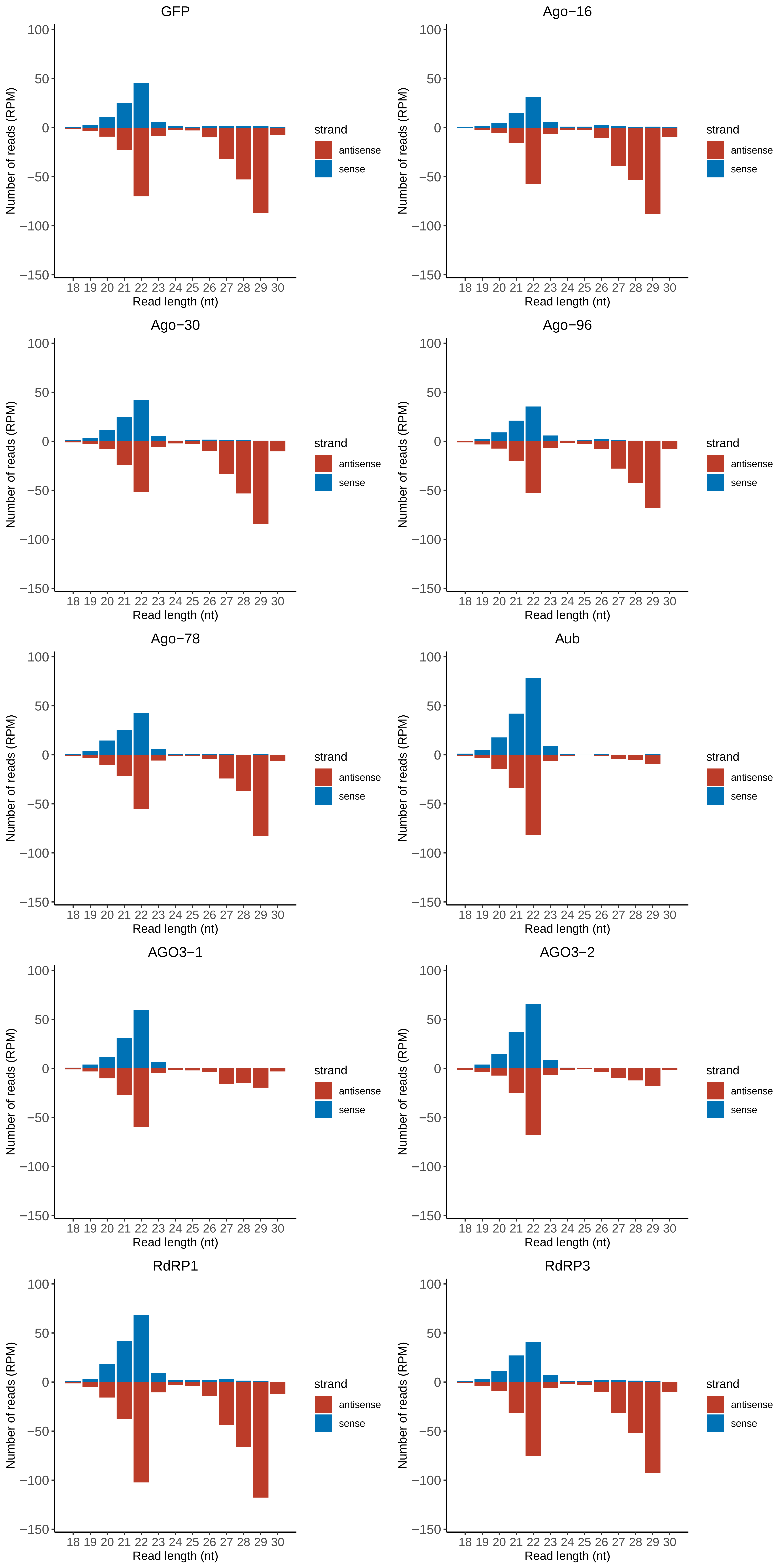

### Motif:rnd-6_family-902TE_lengthdistribution.pdf

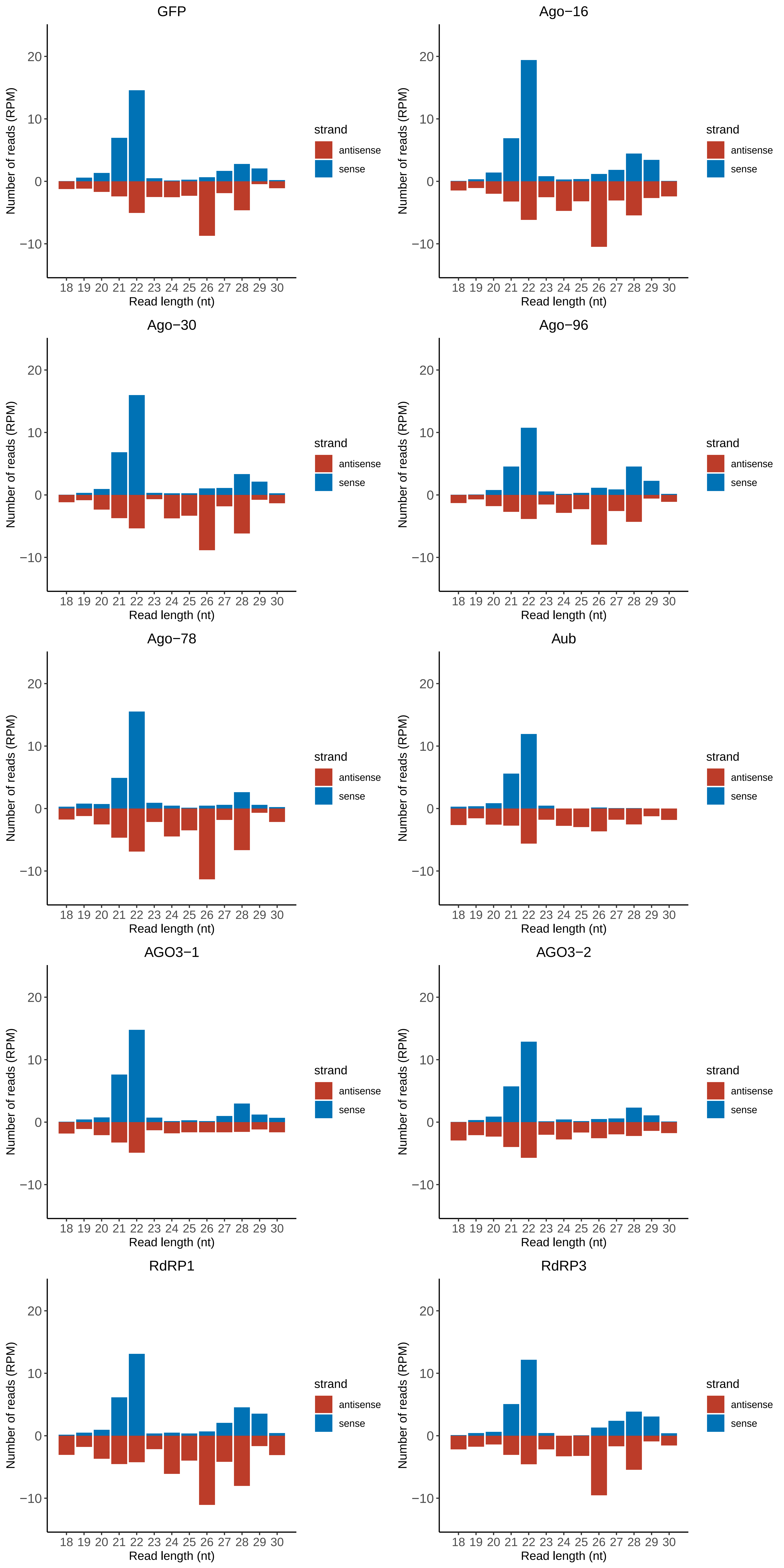

### Motif:rnd-6_family-906TE_lengthdistribution.pdf

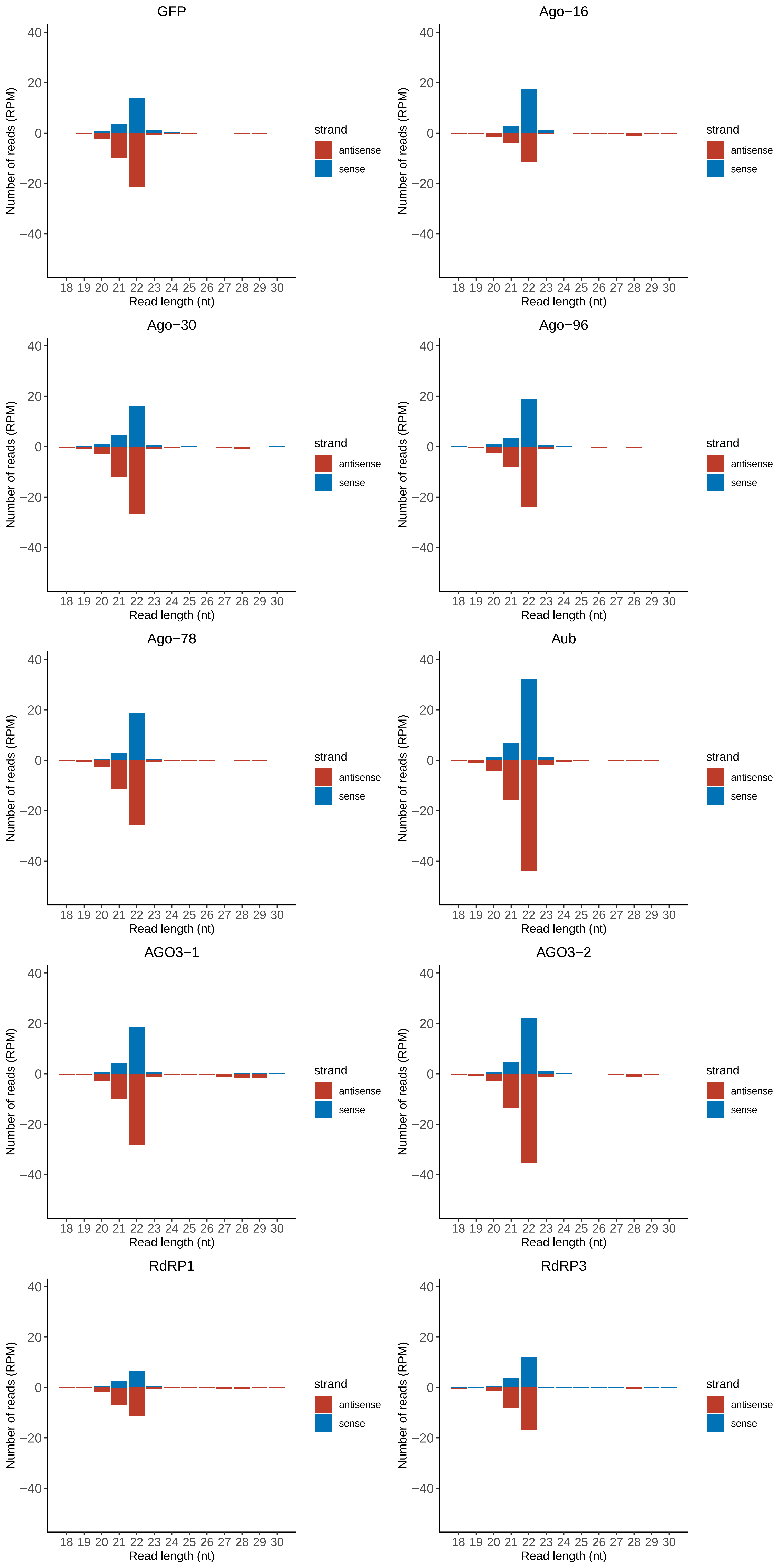

### Motif:rnd-6_family-1016TE_lengthdistribution.pdf

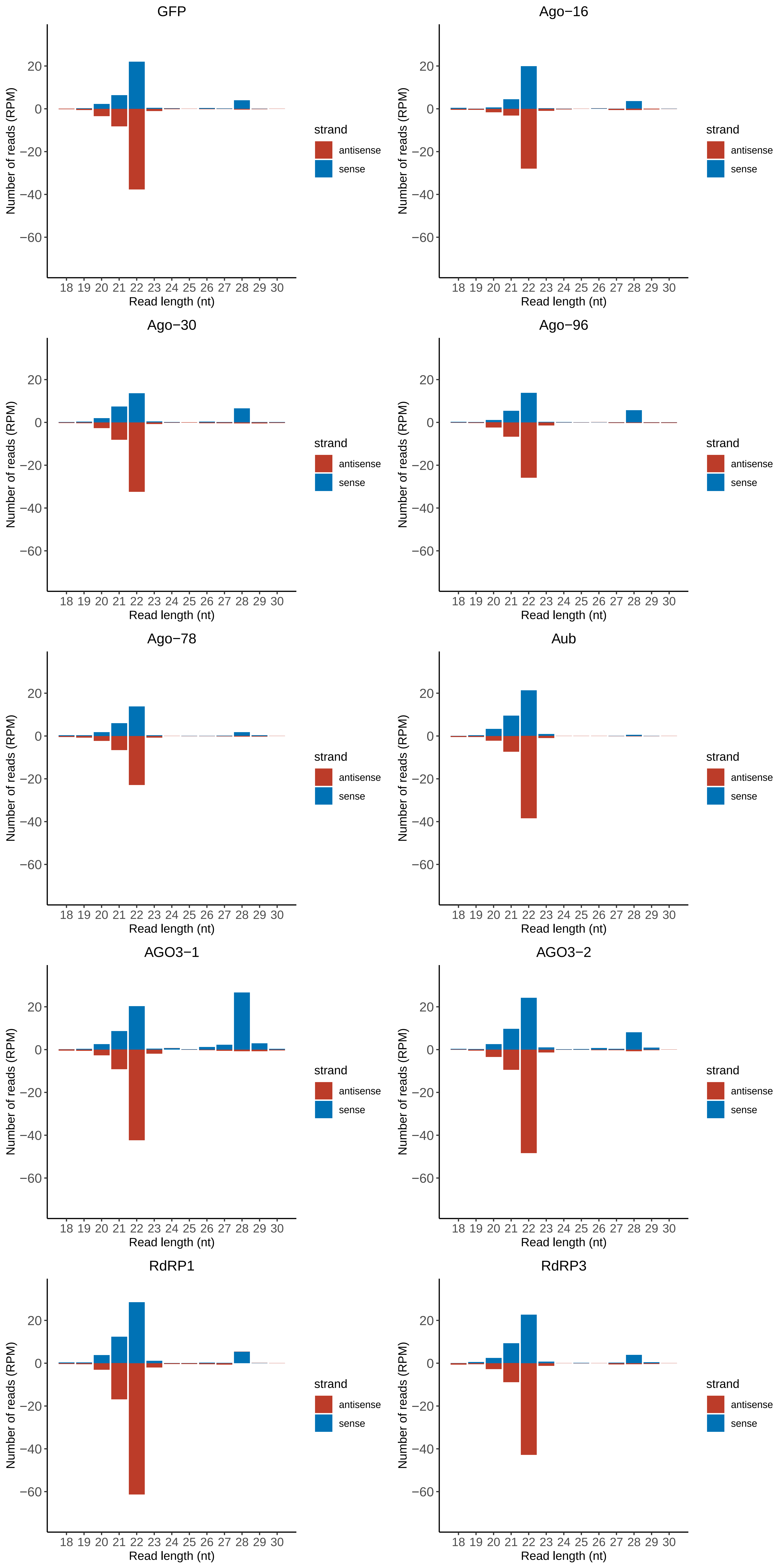

### Motif:rnd-6_family-1241TE_lengthdistribution.pdf

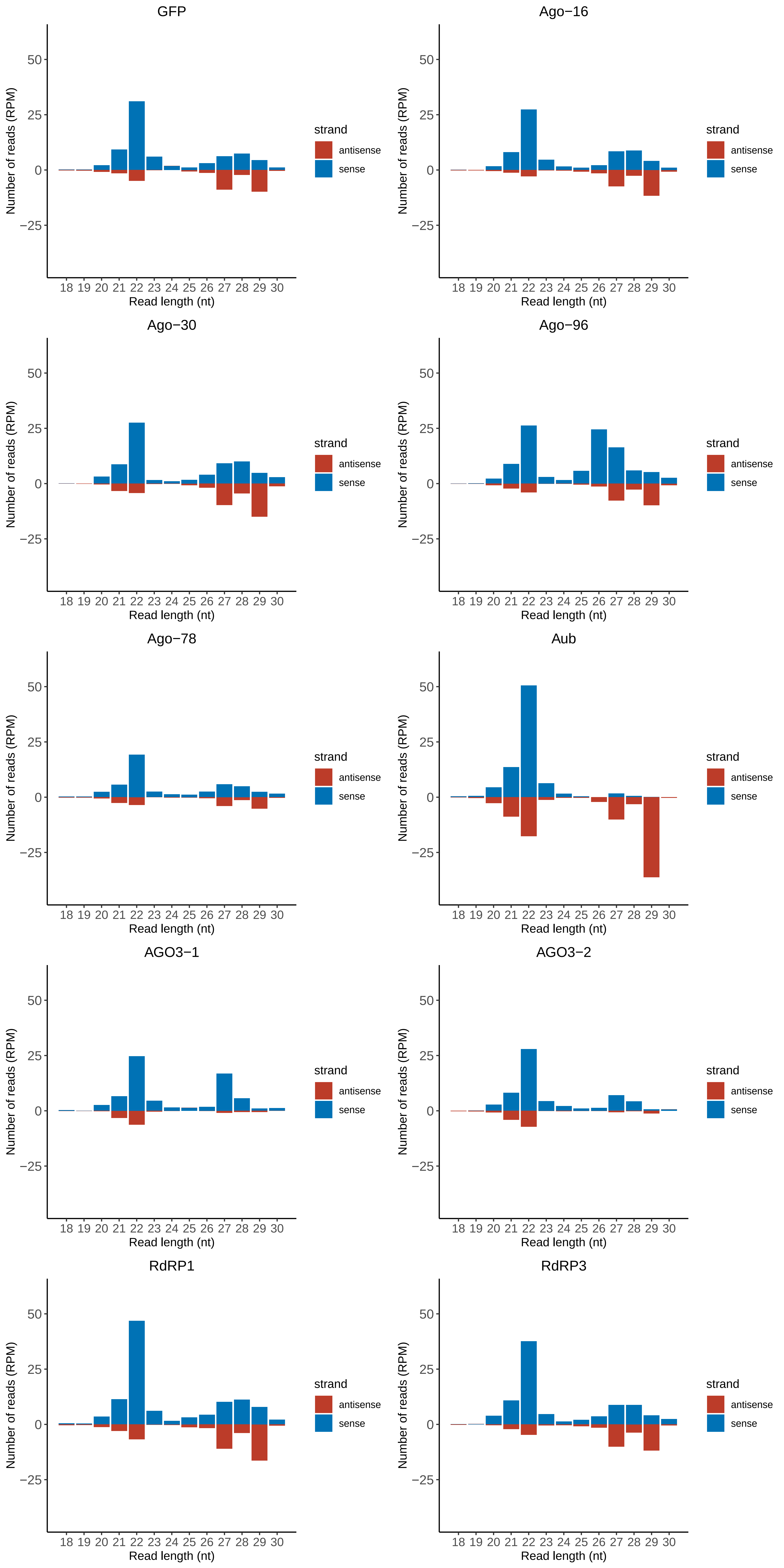

### Motif:rnd-6_family-1887TE_lengthdistribution.pdf

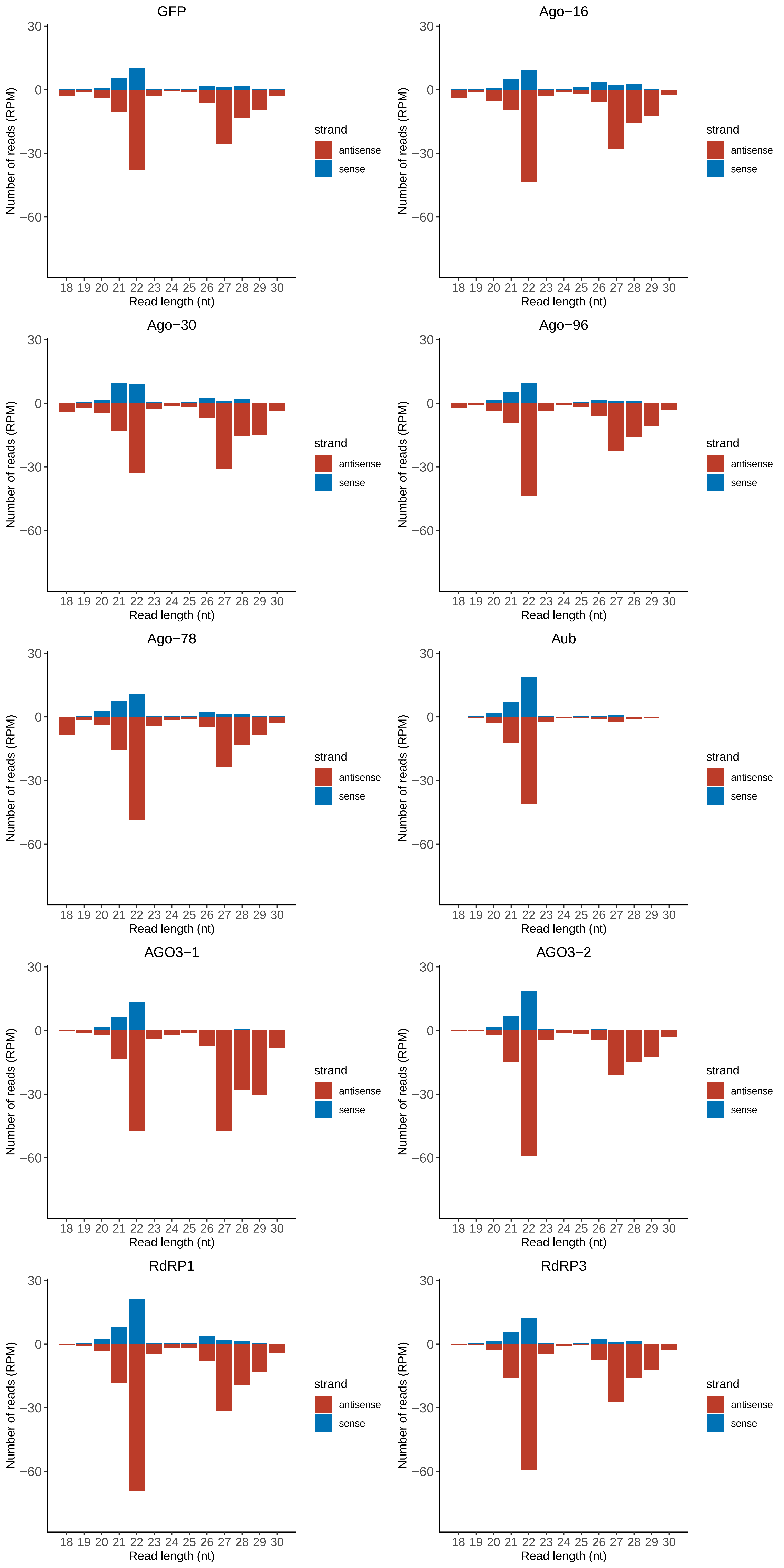

### Motif:rnd-6_family-2543TE_lengthdistribution.pdf

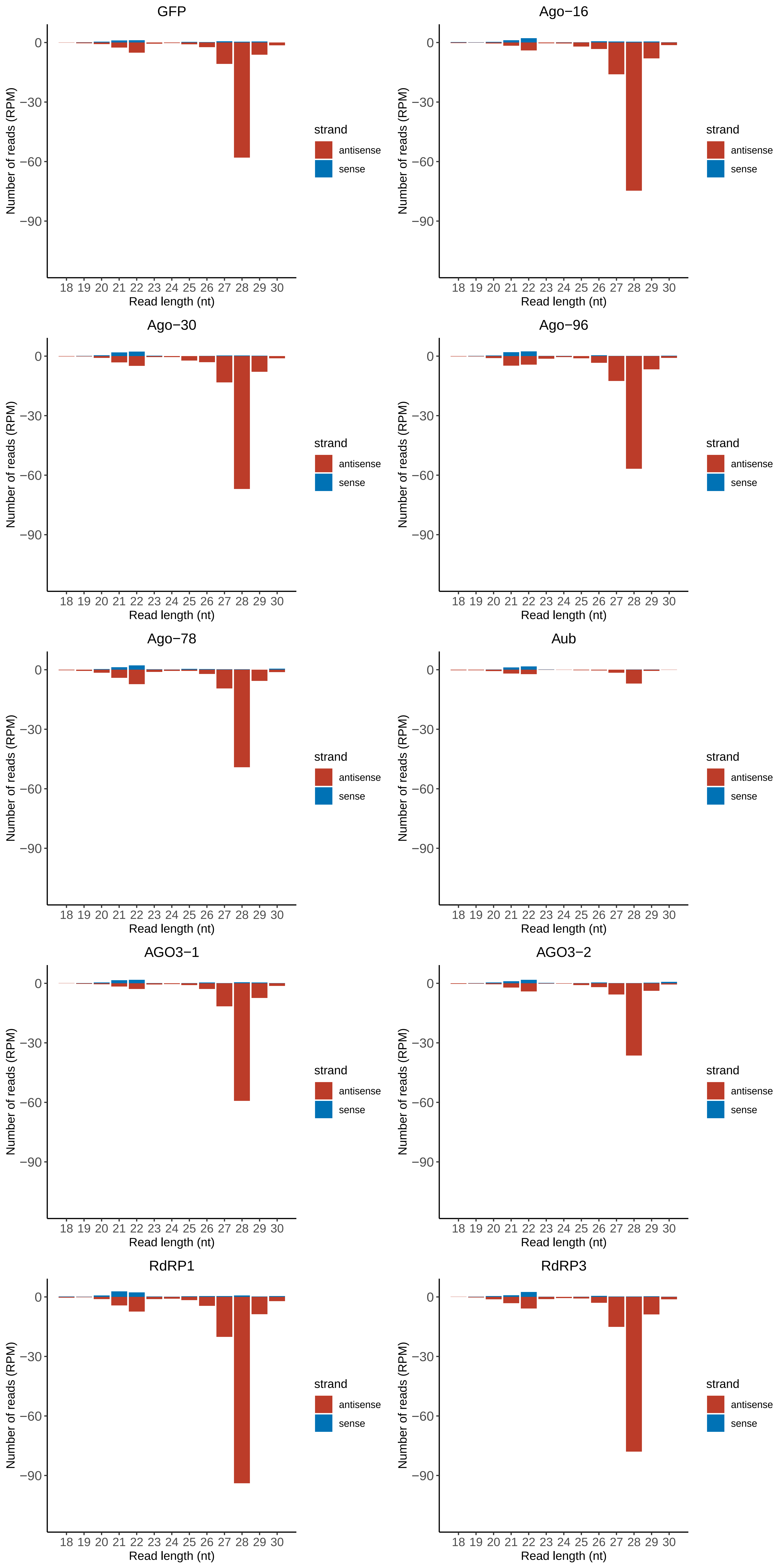

### Motif:rnd-6_family-2596TE_lengthdistribution.pdf

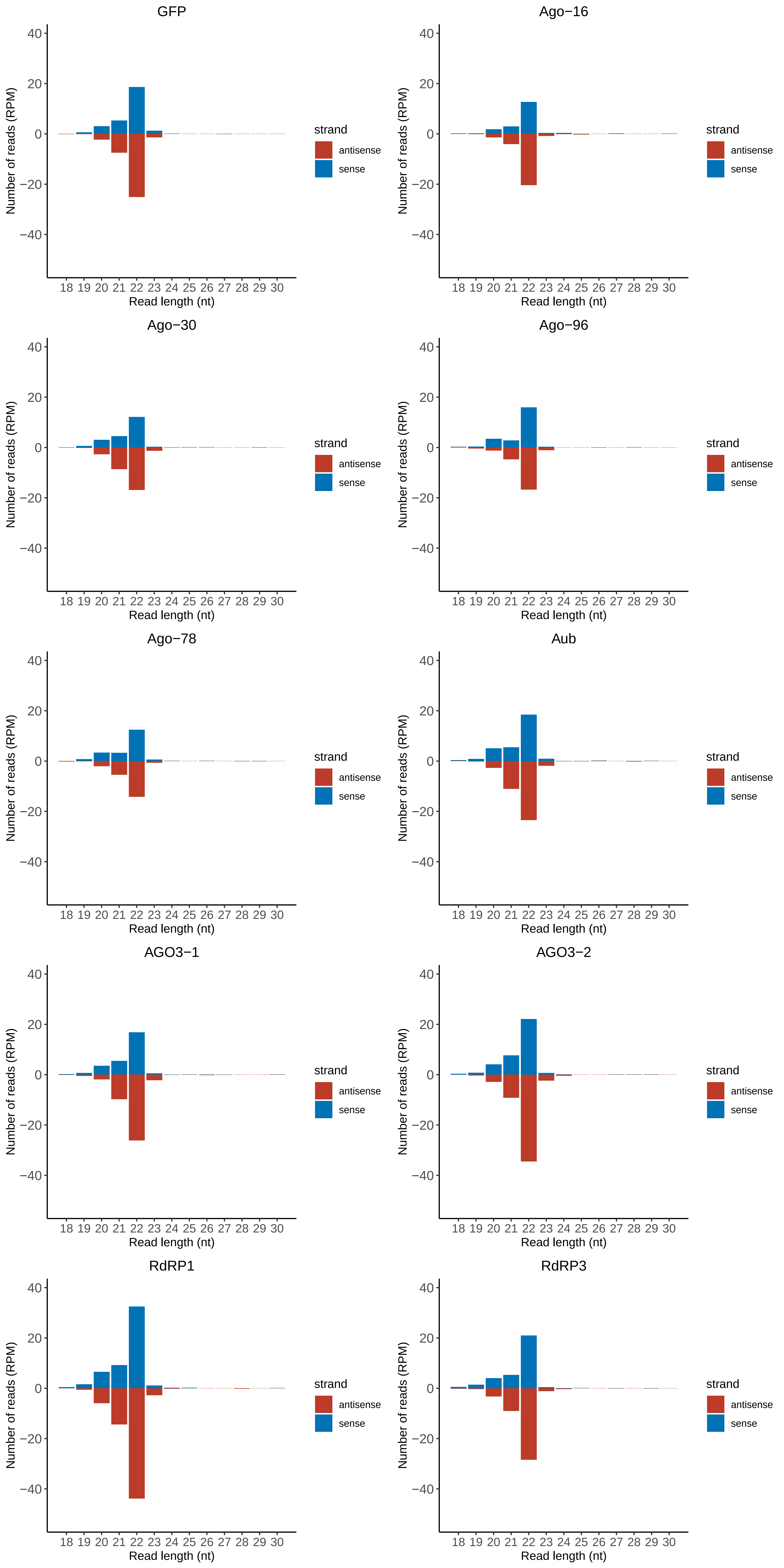

### Motif:rnd-6_family-2714TE_lengthdistribution.pdf

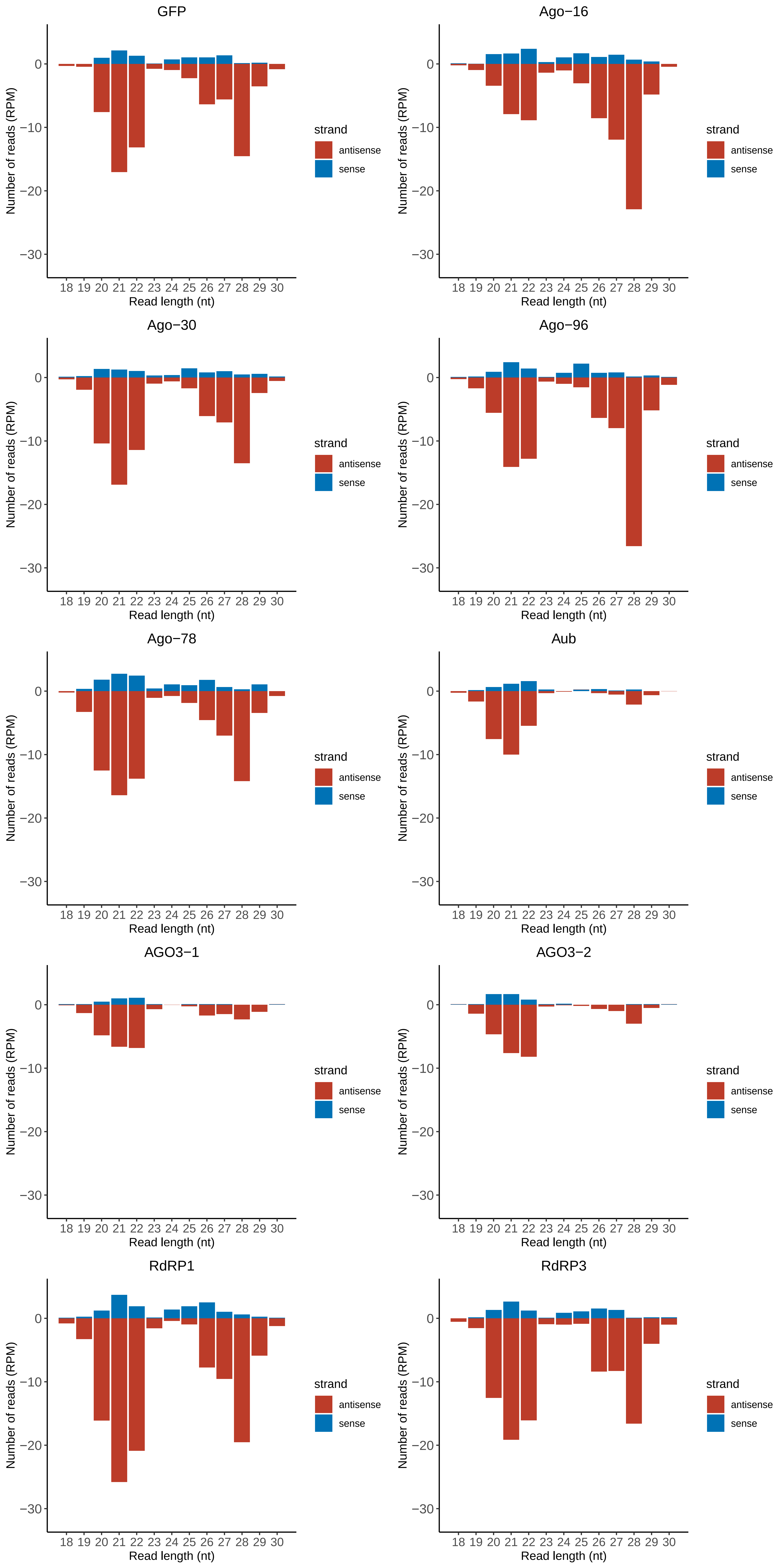

### Motif:rnd-6_family-2824TE_lengthdistribution.pdf

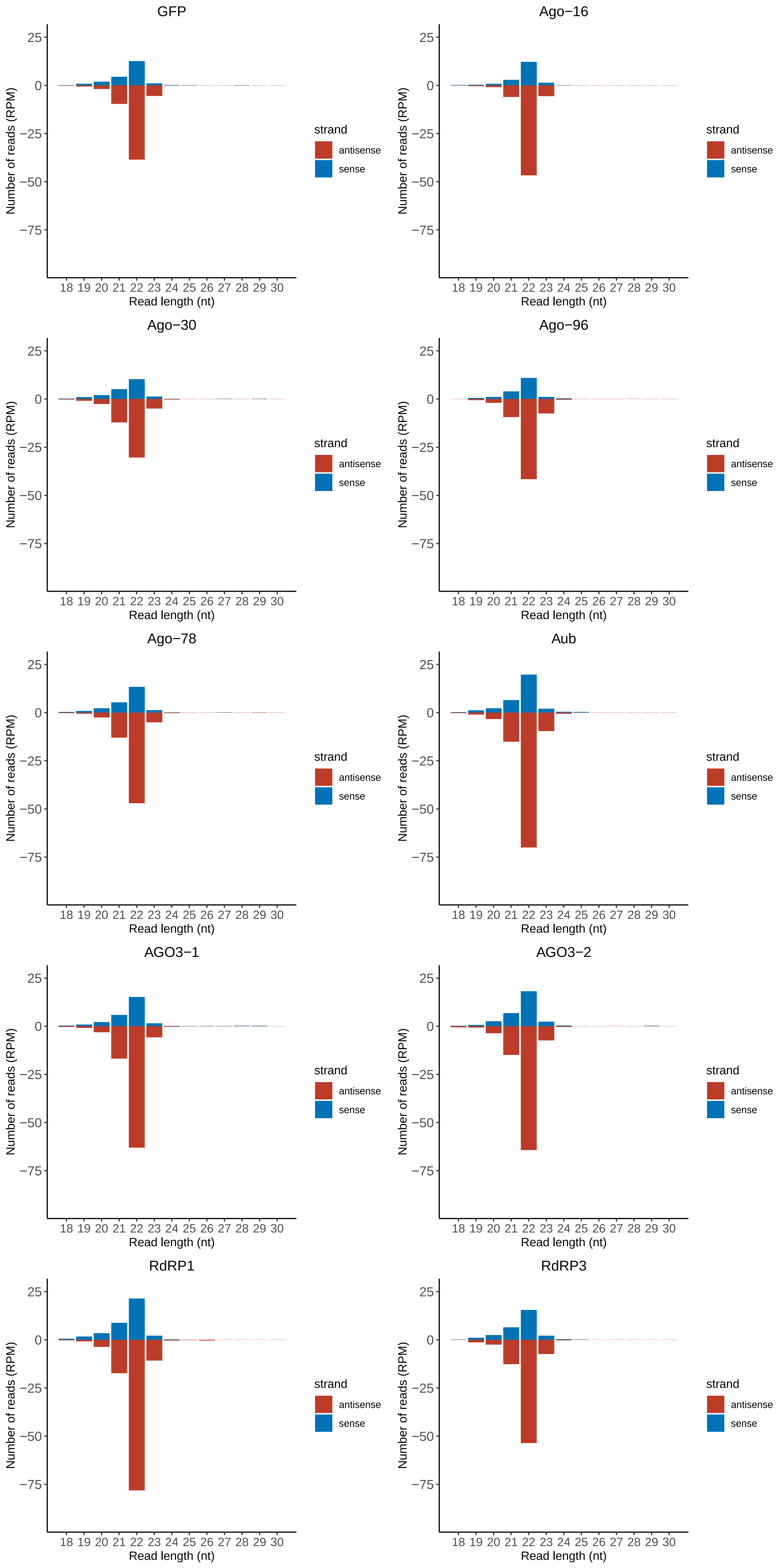

### Motif:rnd-6_family-2943TE_lengthdistribution.pdf

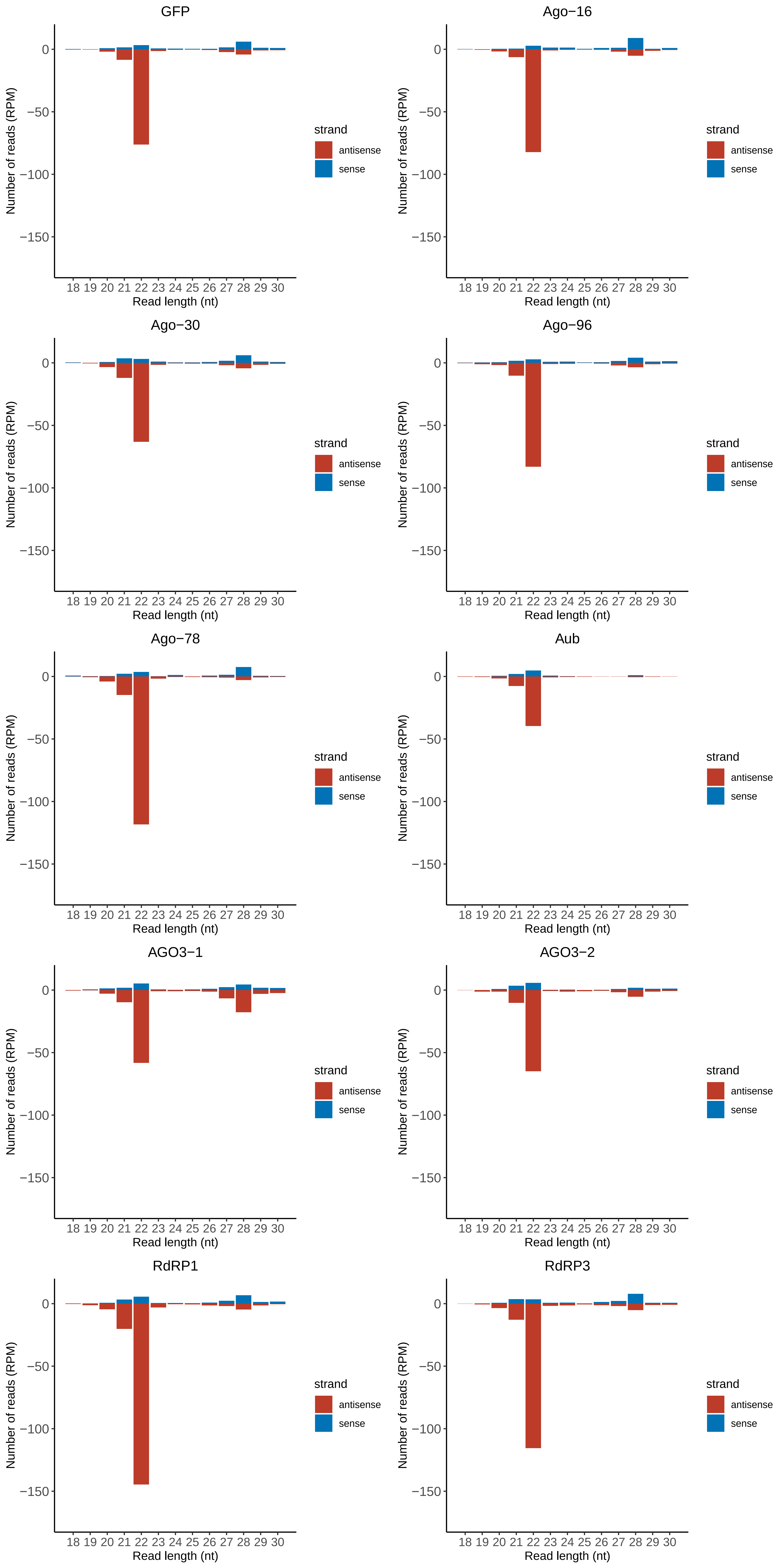

### Motif:rnd-6_family-3044TE_lengthdistribution.pdf

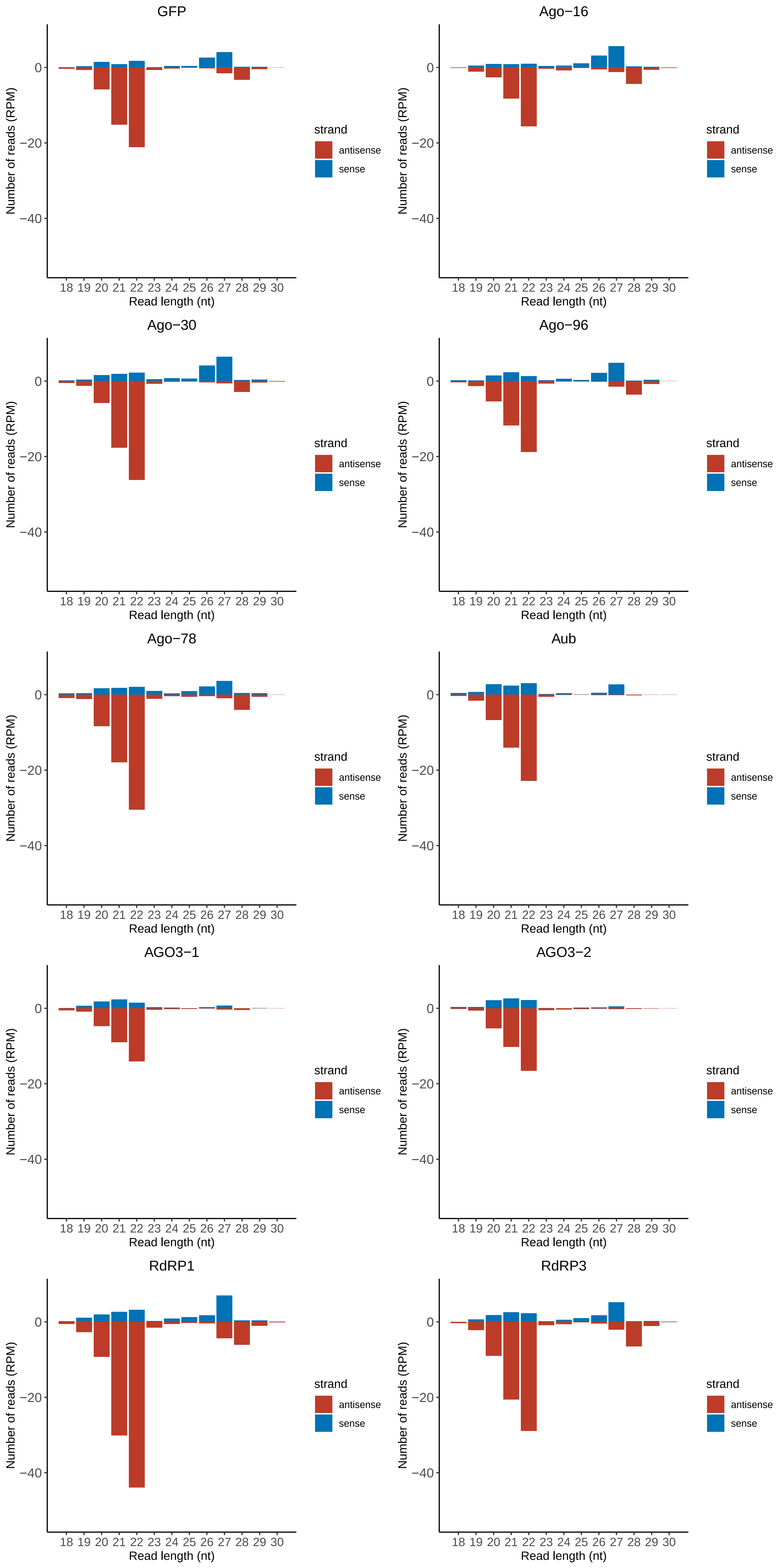

### Motif:rnd-6_family-3172TE_lengthdistribution.pdf

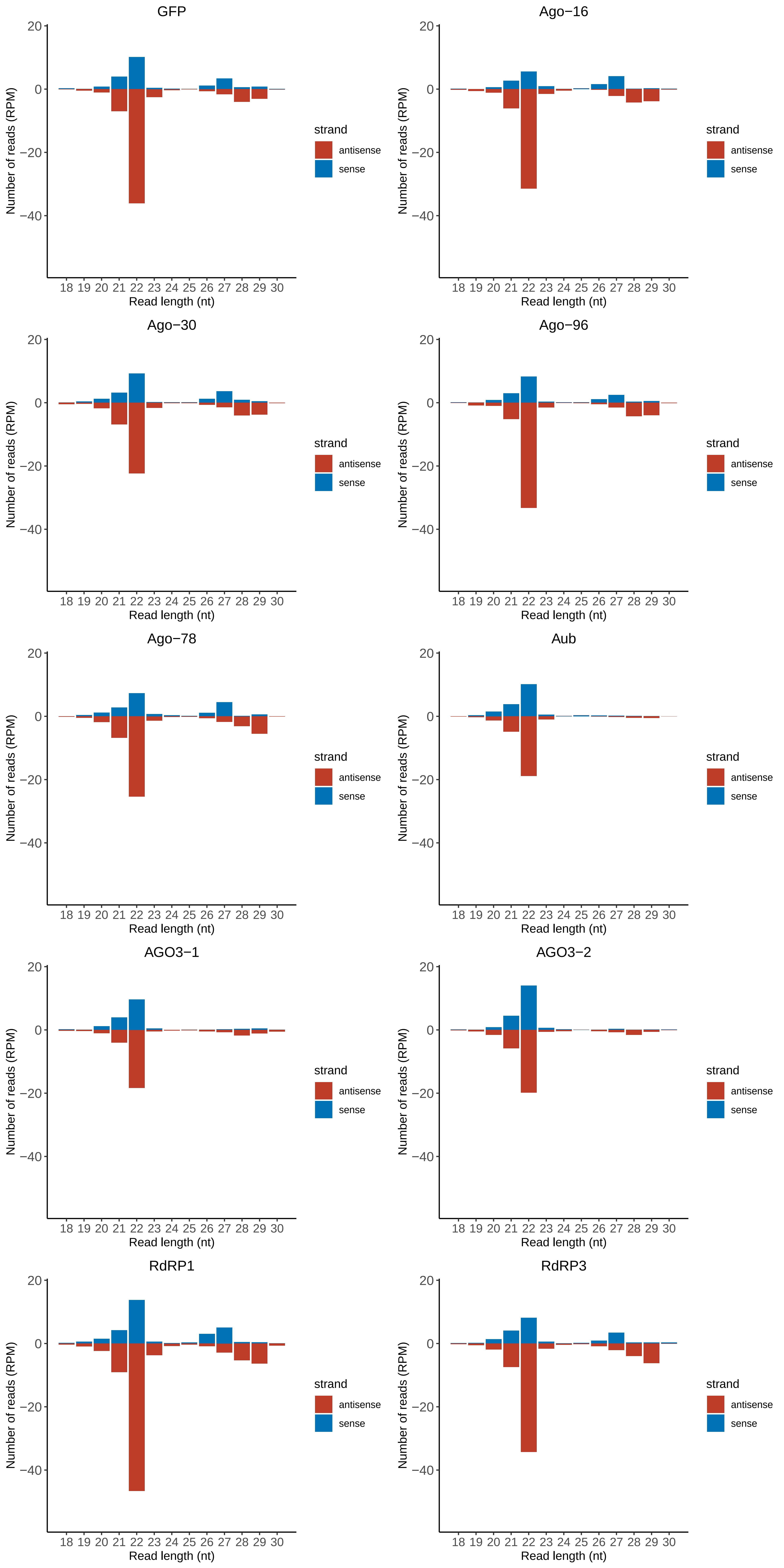

### Motif:rnd-6_family-3909TE_lengthdistribution.pdf

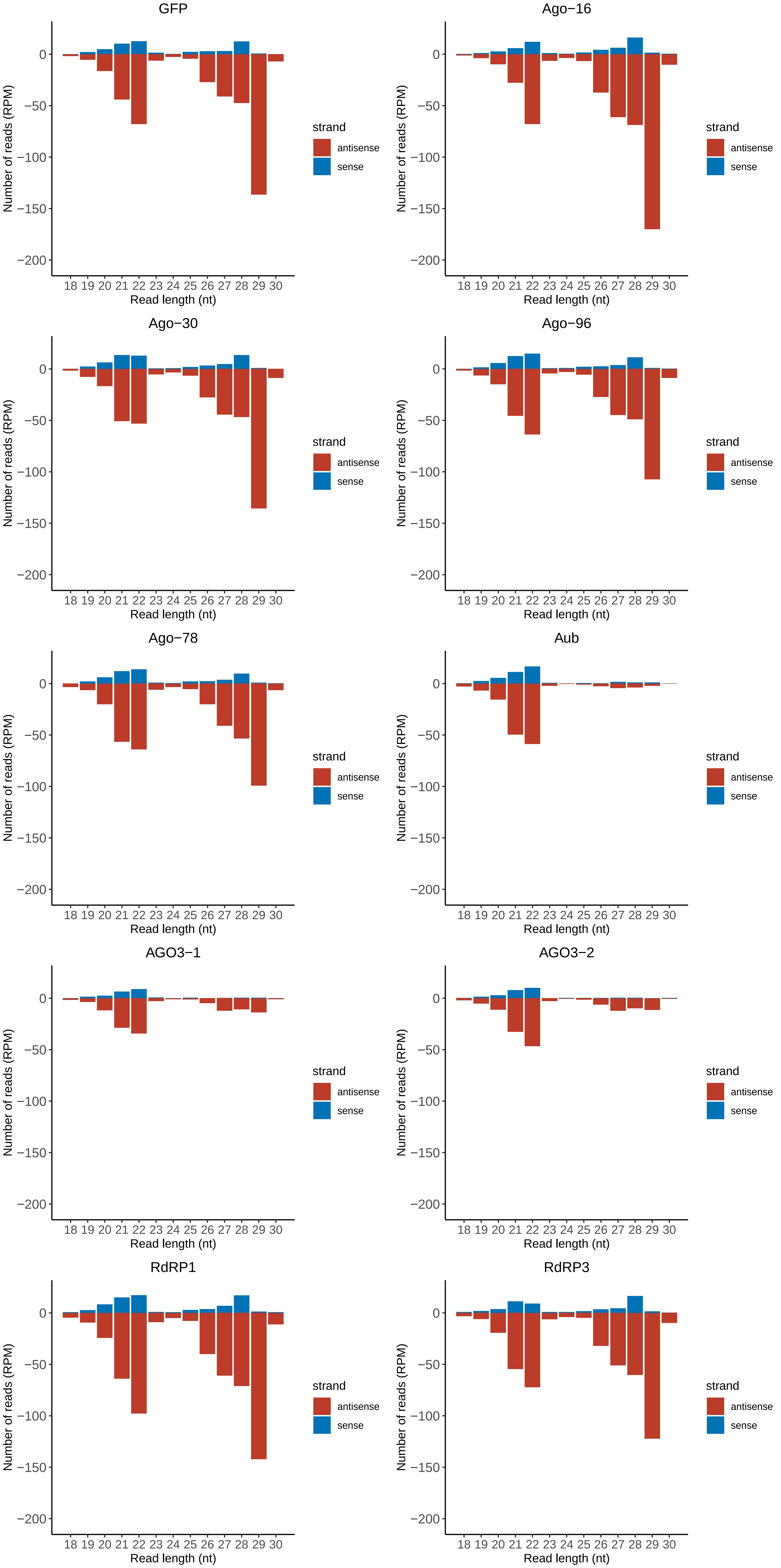

### Motif:rnd-6_family-4218TE_lengthdistribution.pdf

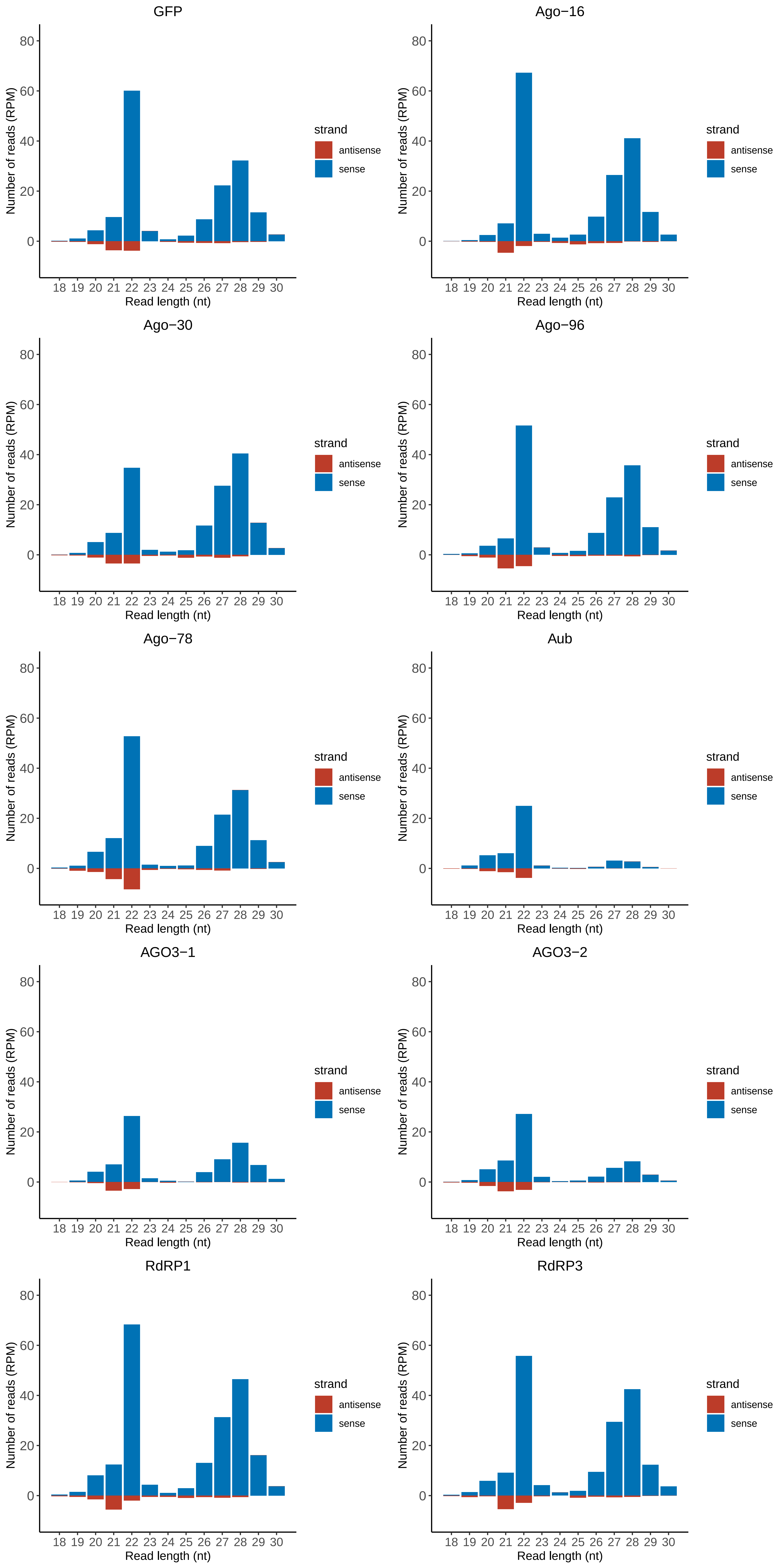

### Motif:rnd-6_family-4430TE_lengthdistribution.pdf

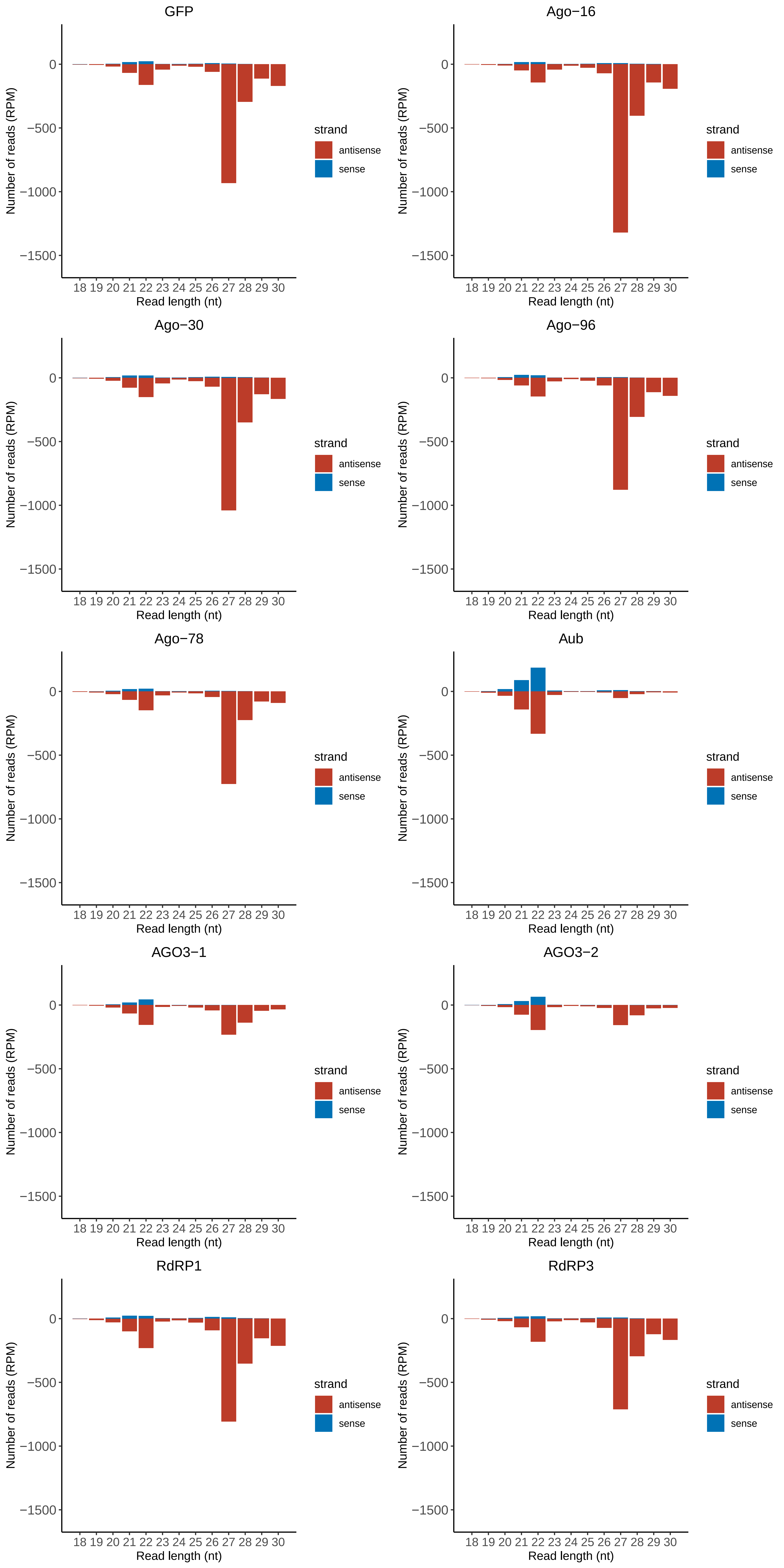

### Motif:rnd-6_family-4733TE_lengthdistribution.pdf

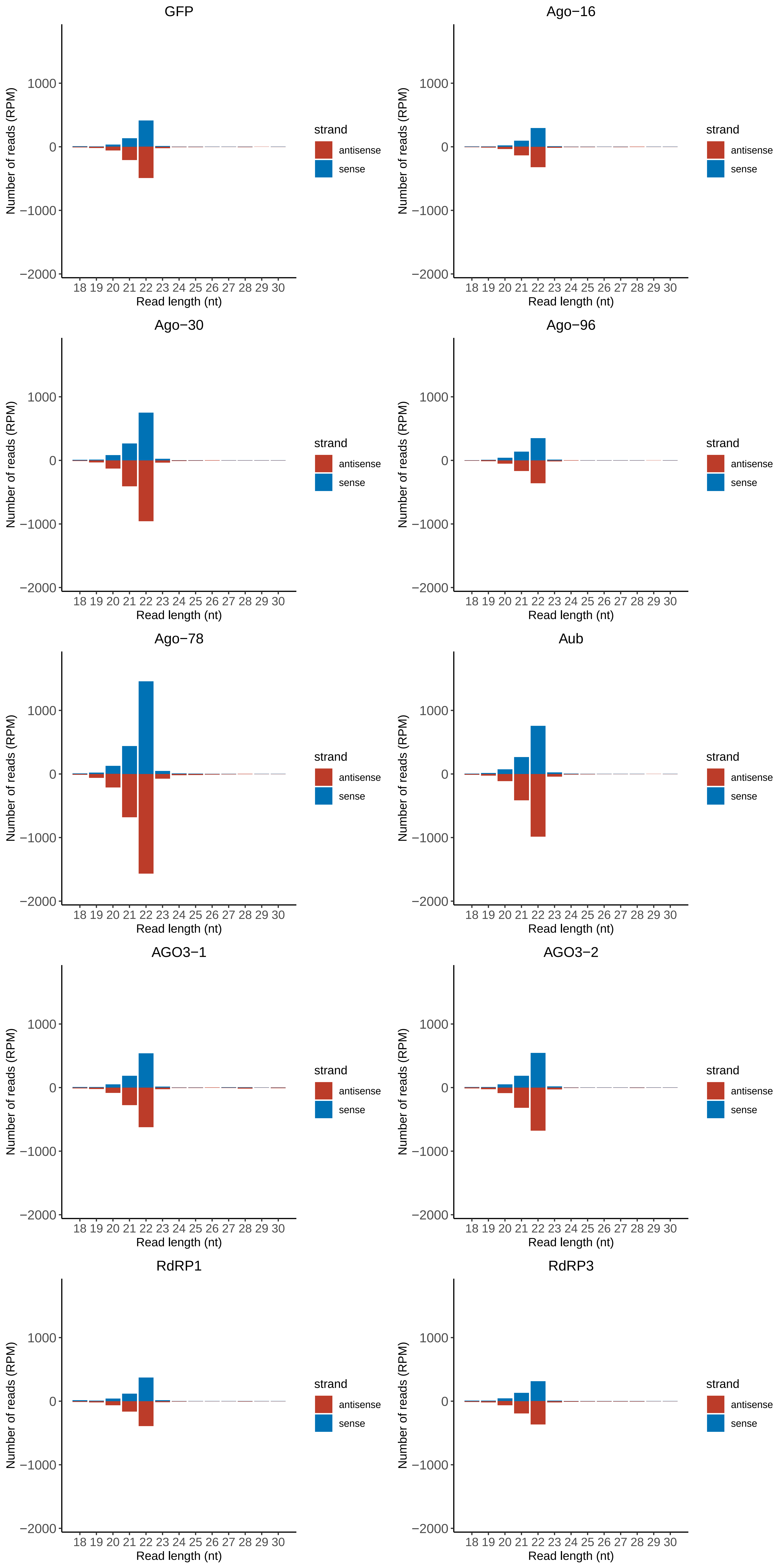

### Motif:rnd-6_family-4780TE_lengthdistribution.pdf

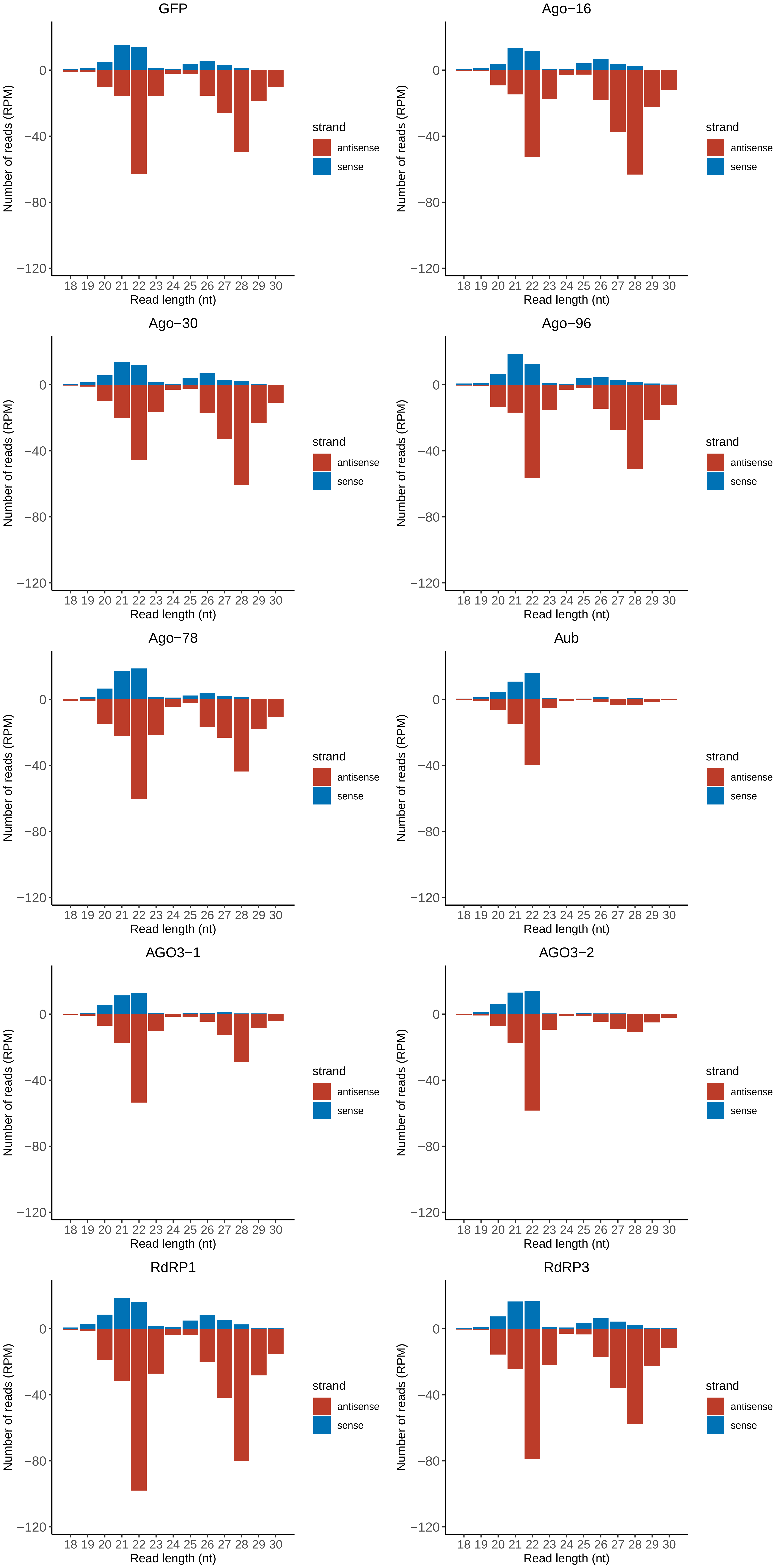

### Motif:rnd-6_family-5159TE_lengthdistribution.pdf

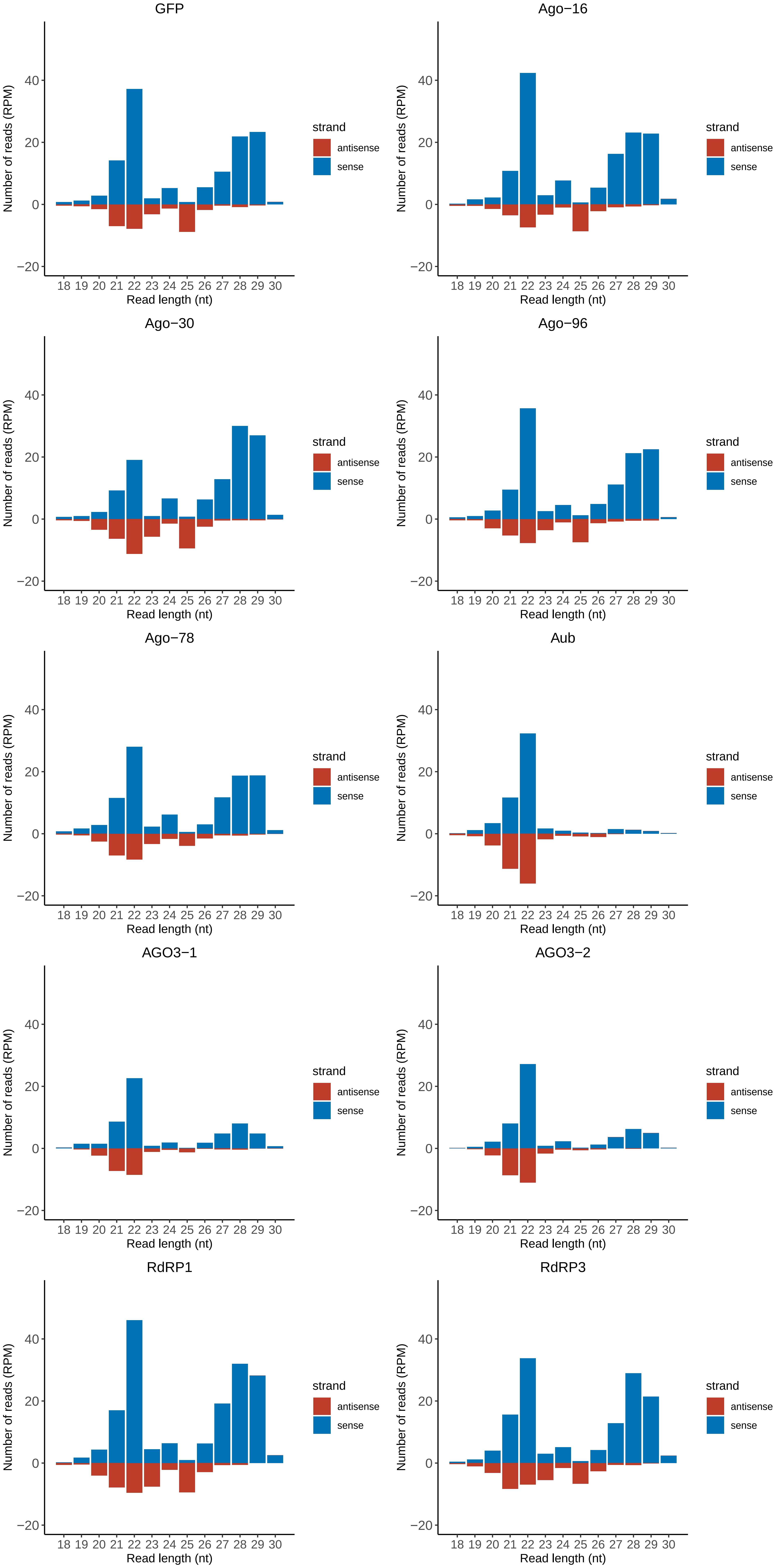

### Motif:rnd-6_family-5252TE_lengthdistribution.pdf

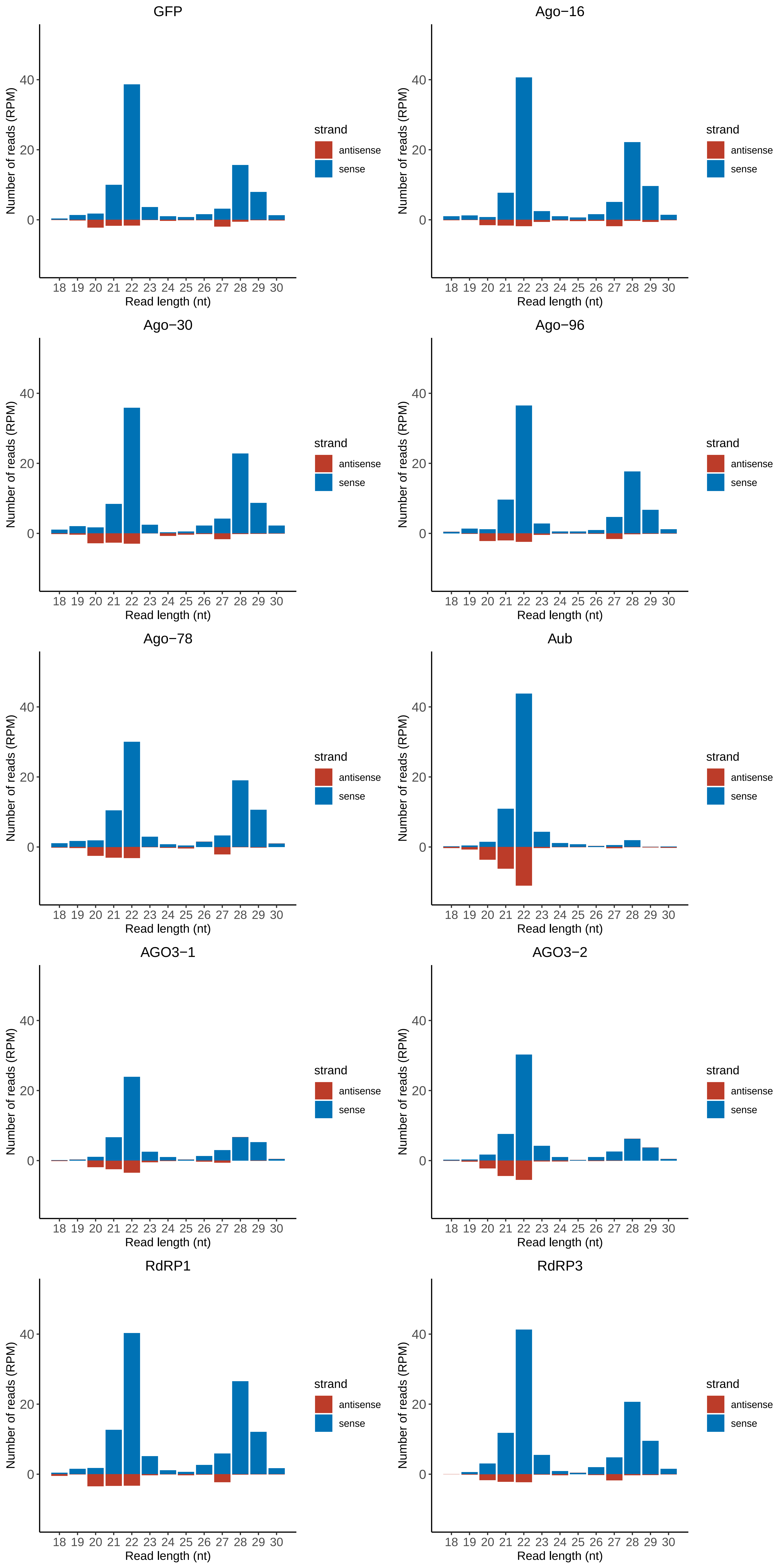

### Motif:rnd-6_family-5915TE_lengthdistribution.pdf

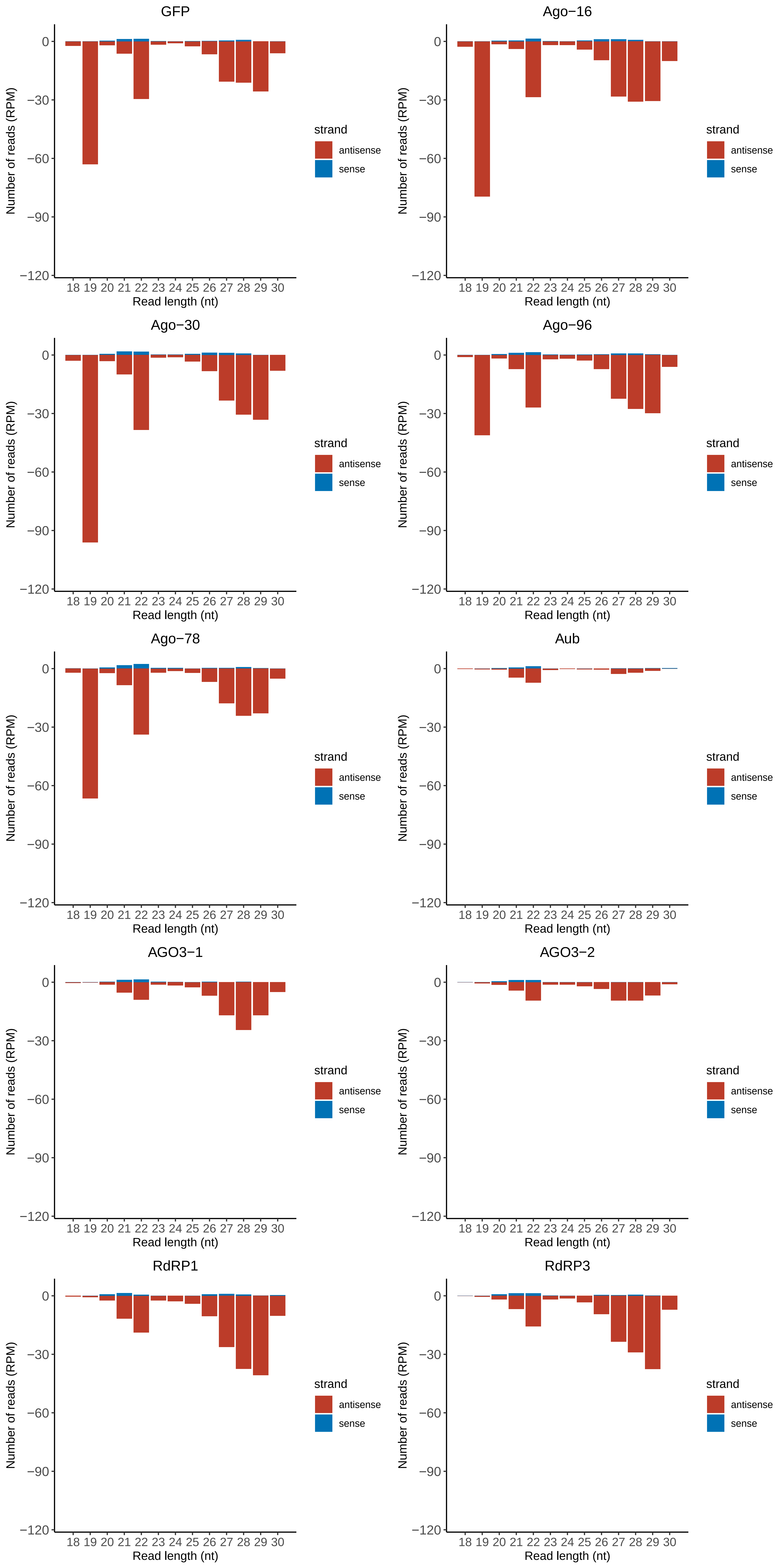

### Motif:rnd-6_family-6470TE_lengthdistribution.pdf

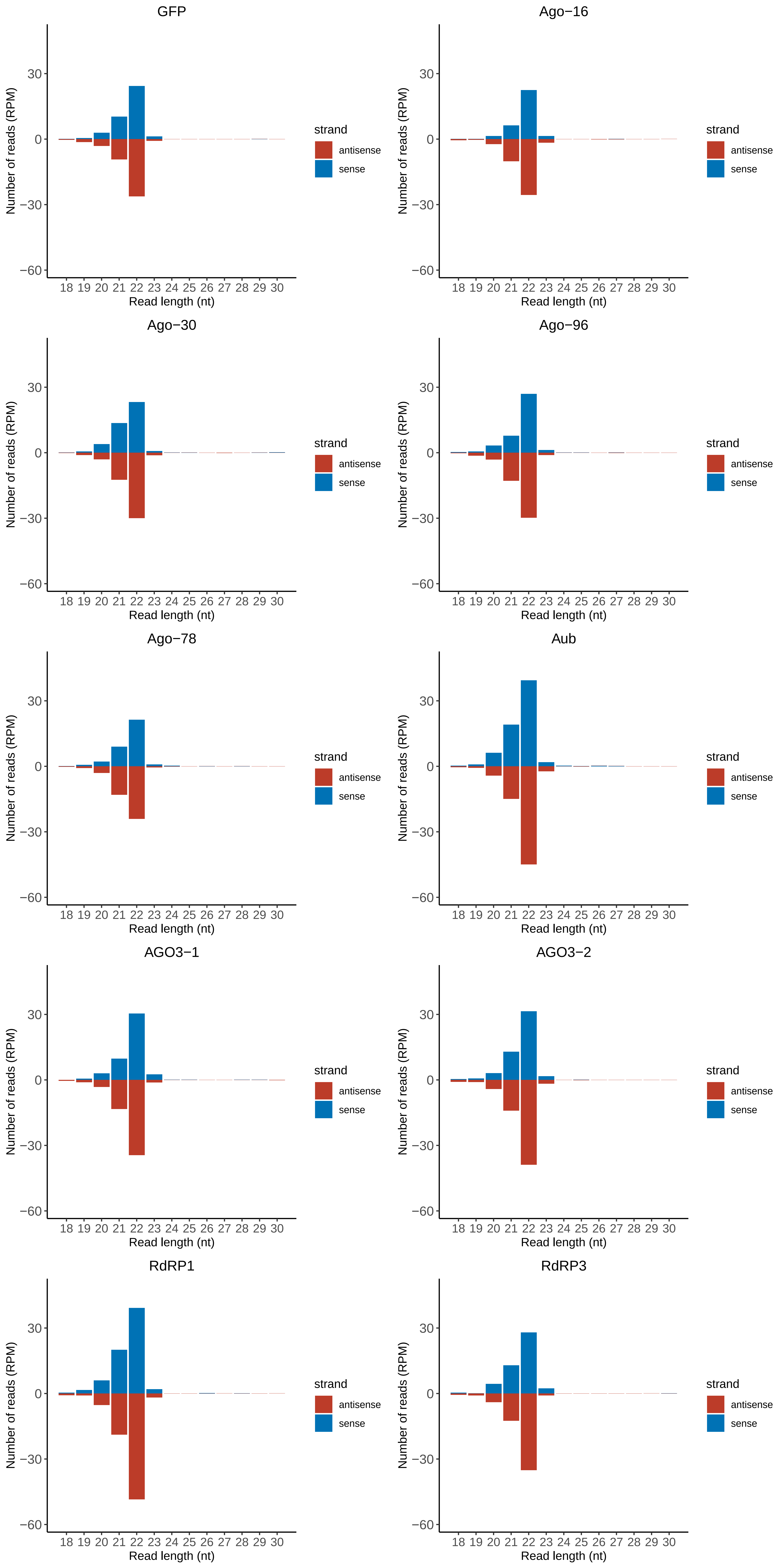

### Motif:rnd-6_family-7183TE_lengthdistribution.pdf

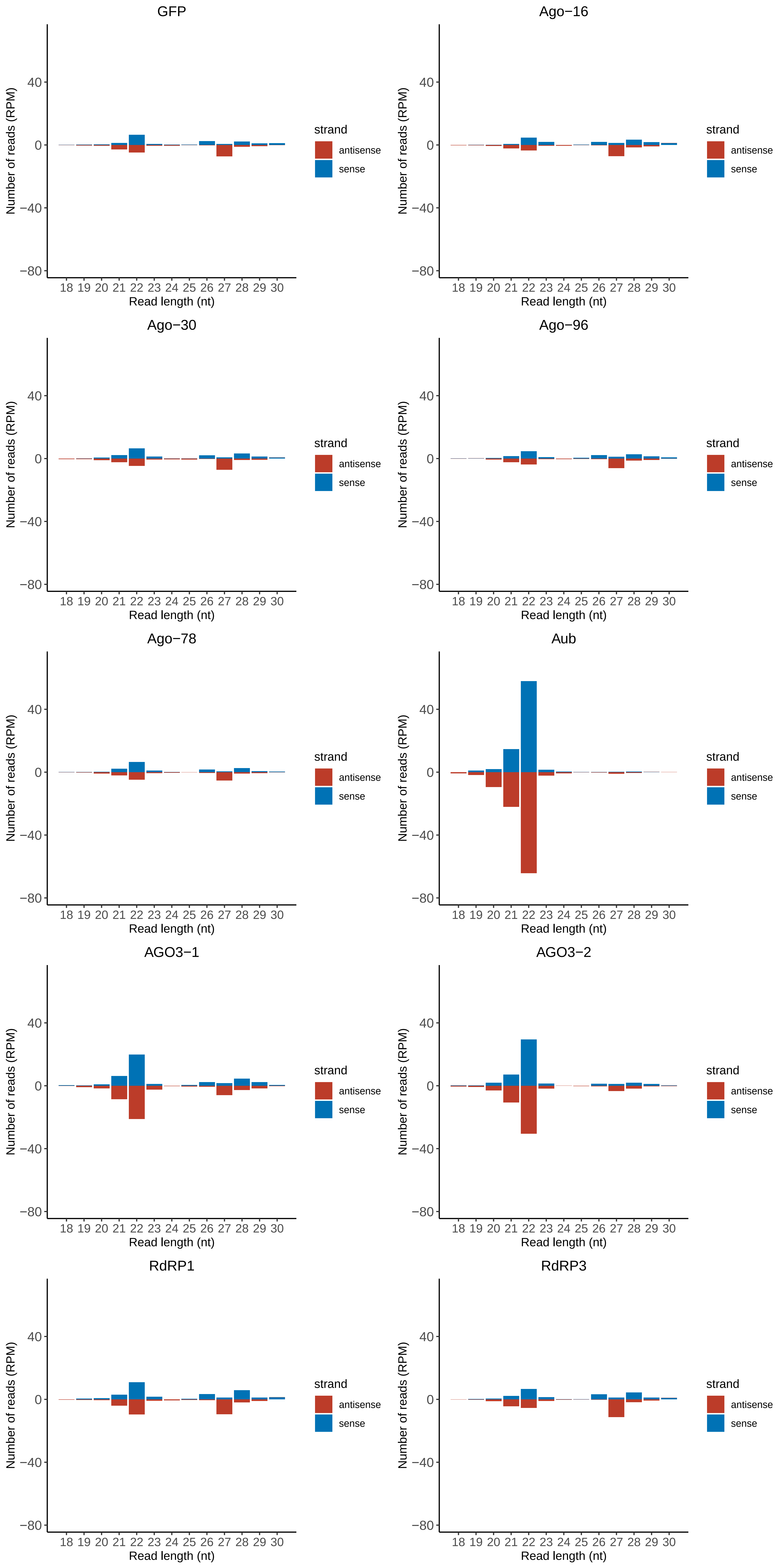

### Motif:rnd-6_family-7992TE_lengthdistribution.pdf

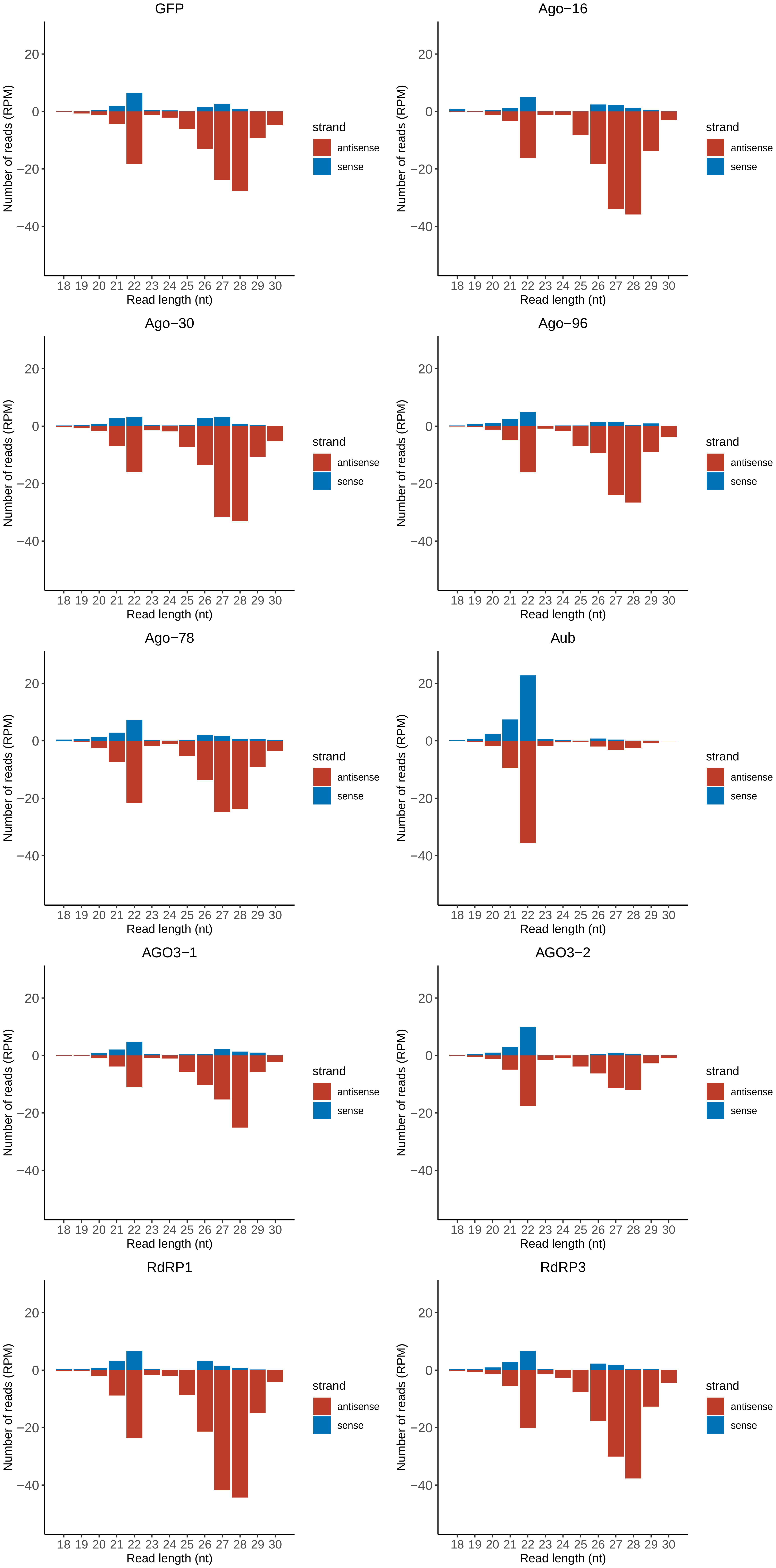

### Motif:rnd-6_family-8871TE_lengthdistribution.pdf

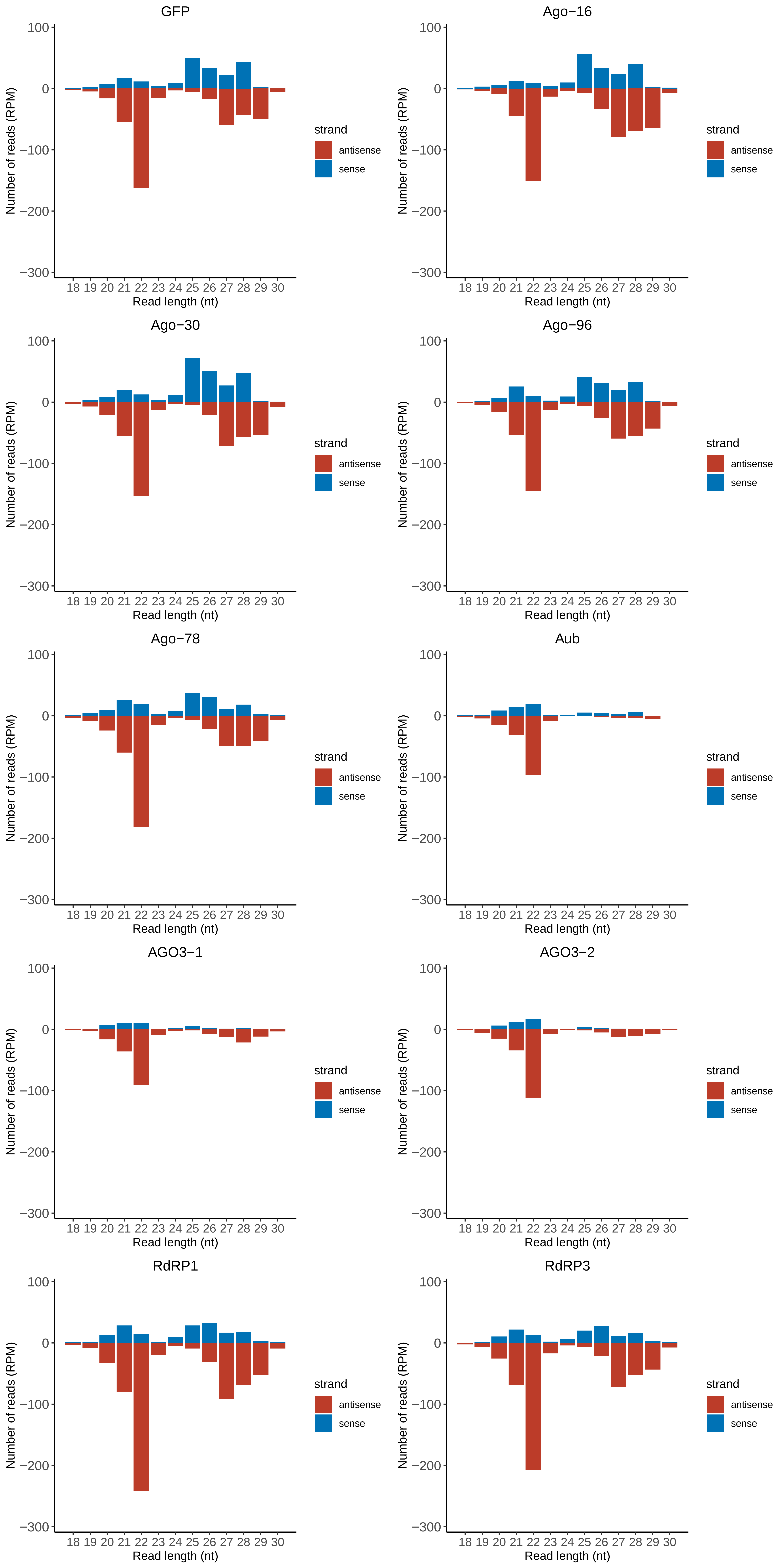

### Motif:rnd-6_family-9065TE_lengthdistribution.pdf

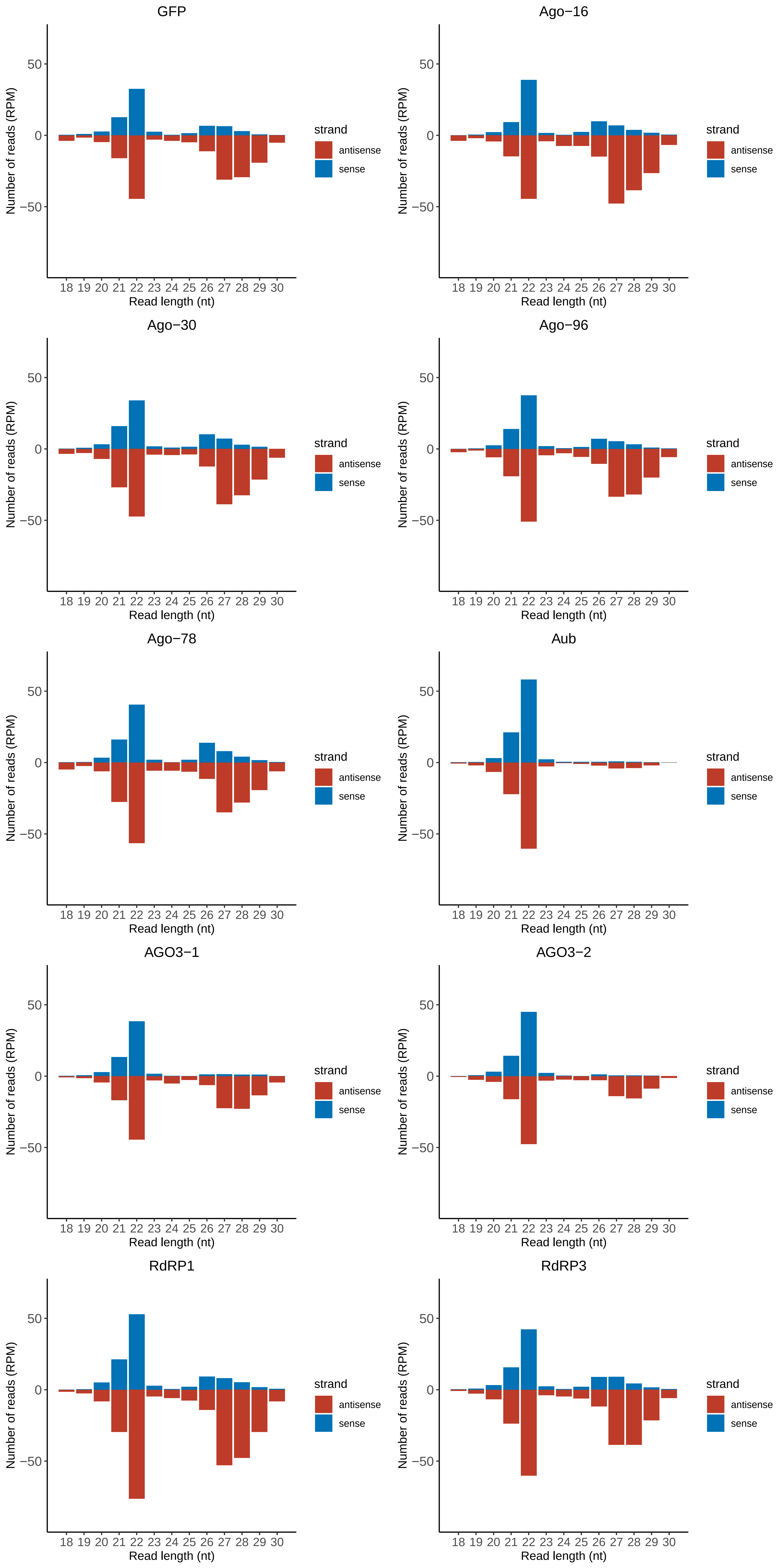

### Motif:rnd-6_family-9480TE_lengthdistribution.pdf

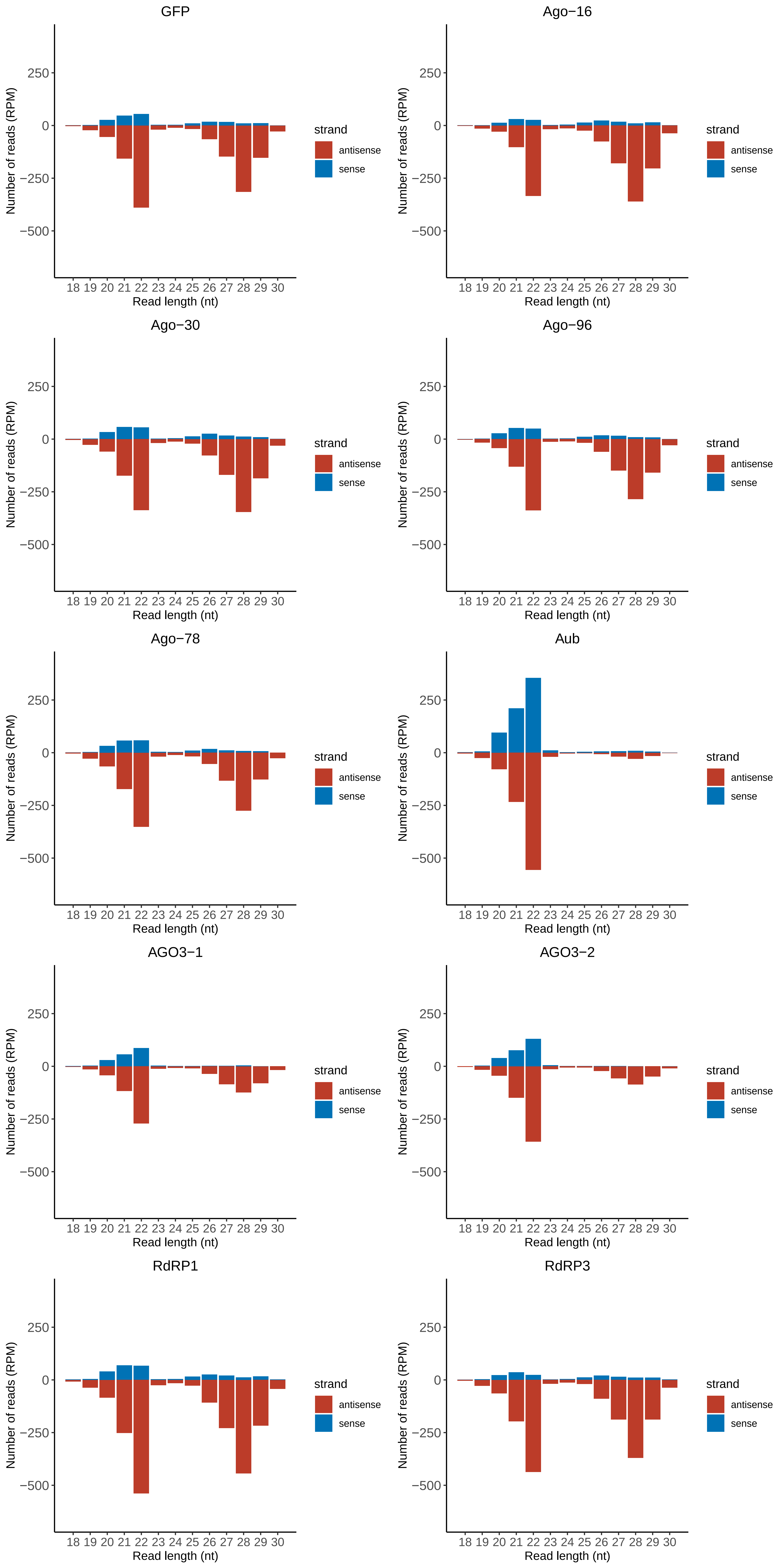

### Motif:rnd-6_family-10230TE_lengthdistribution.pdf

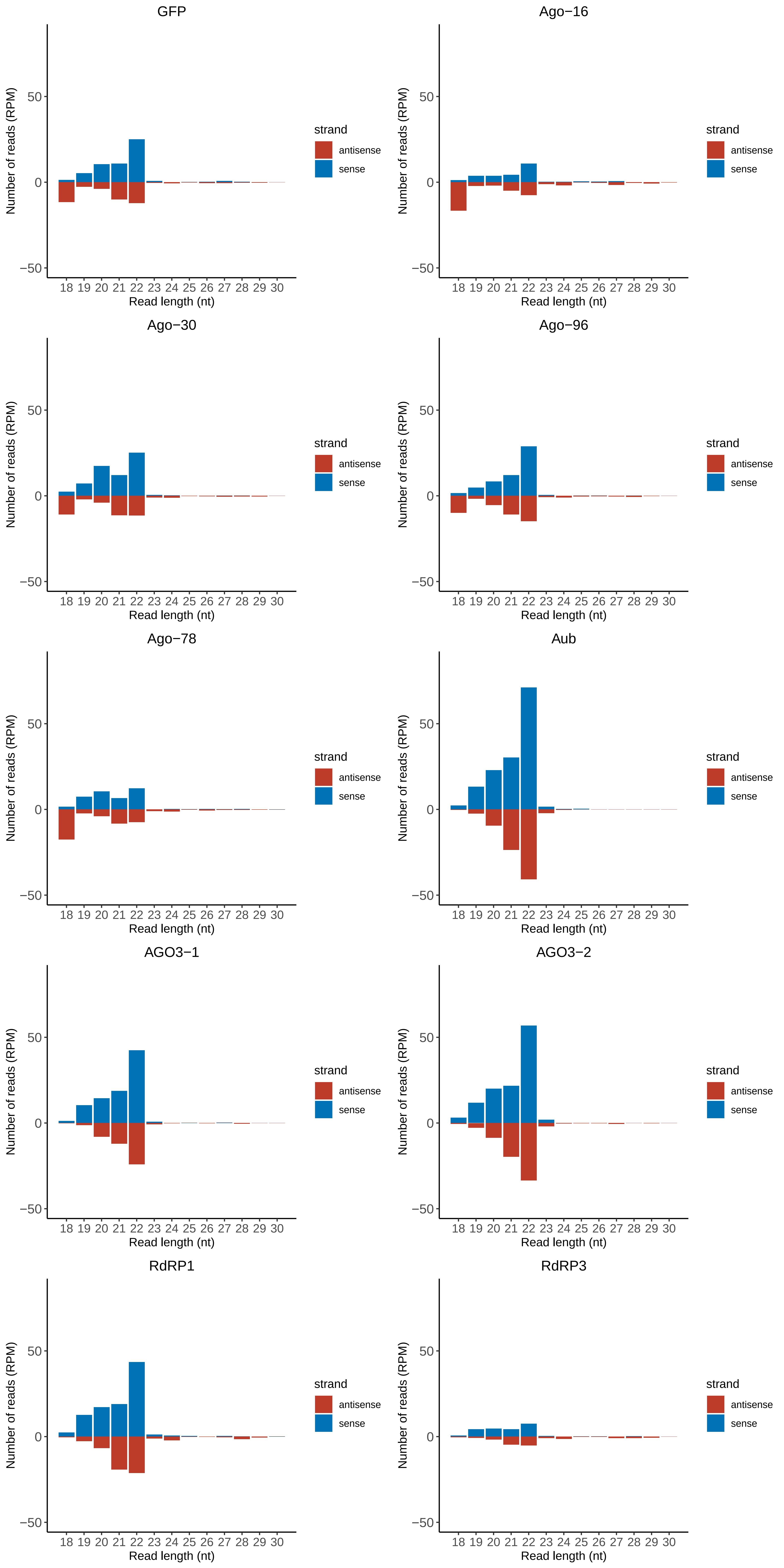

### Motif:rnd-6_family-10845TE_lengthdistribution.pdf

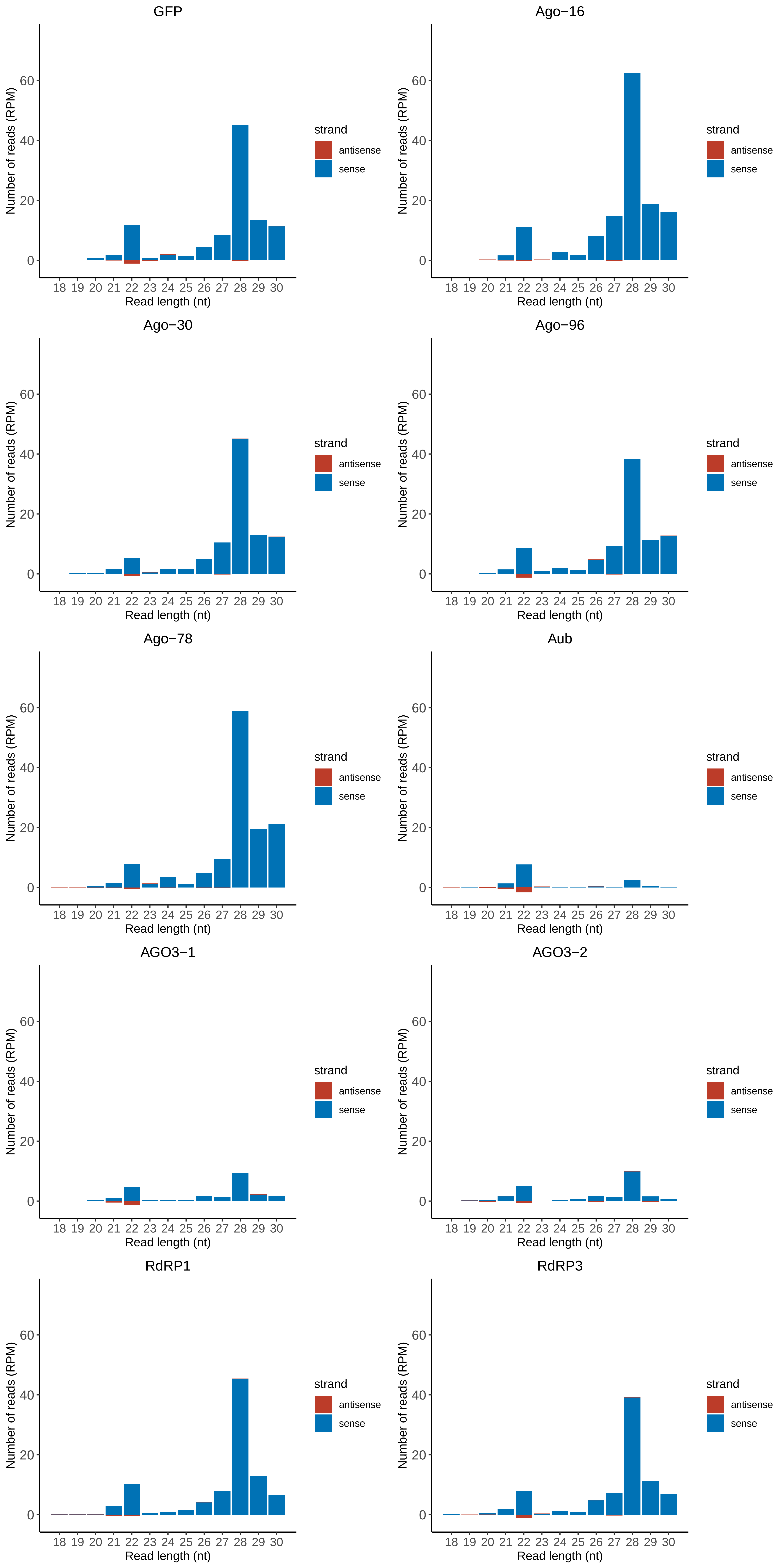

### Motif:rnd-6_family-12152TE_lengthdistribution.pdf

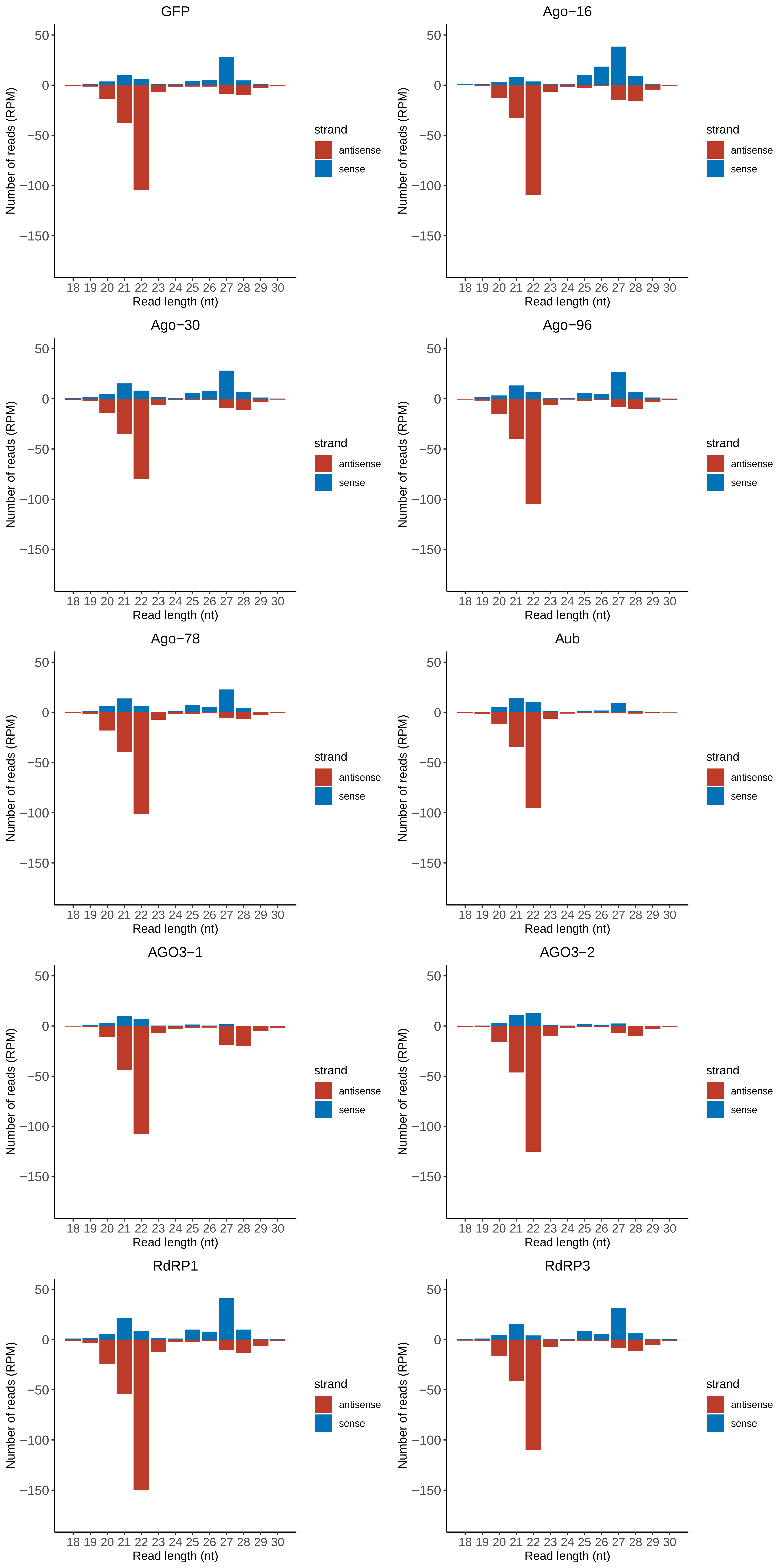

### Motif:rnd-6_family-12619TE_lengthdistribution.pdf

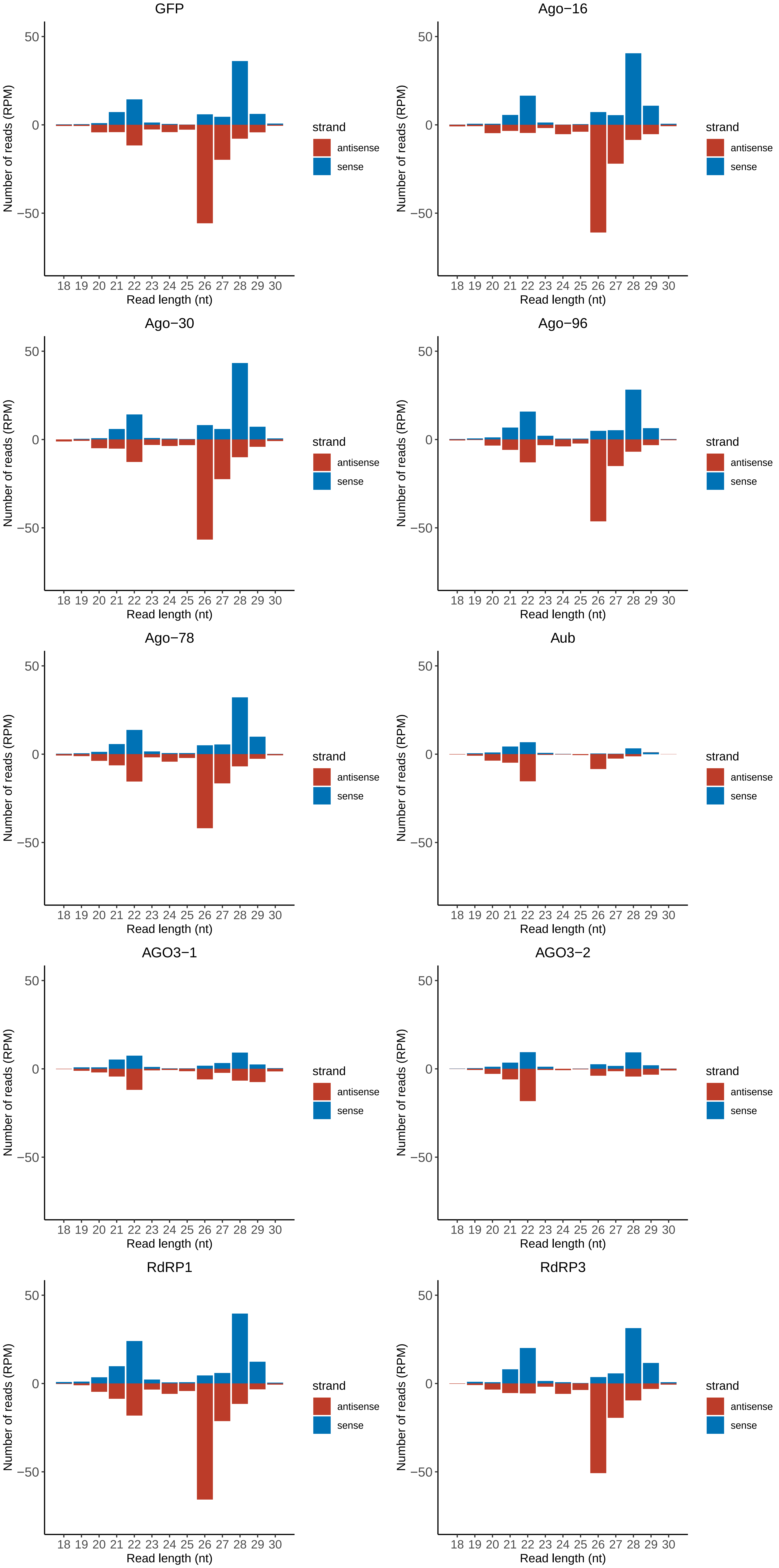

### Motif:rnd-6_family-12996TE_lengthdistribution.pdf

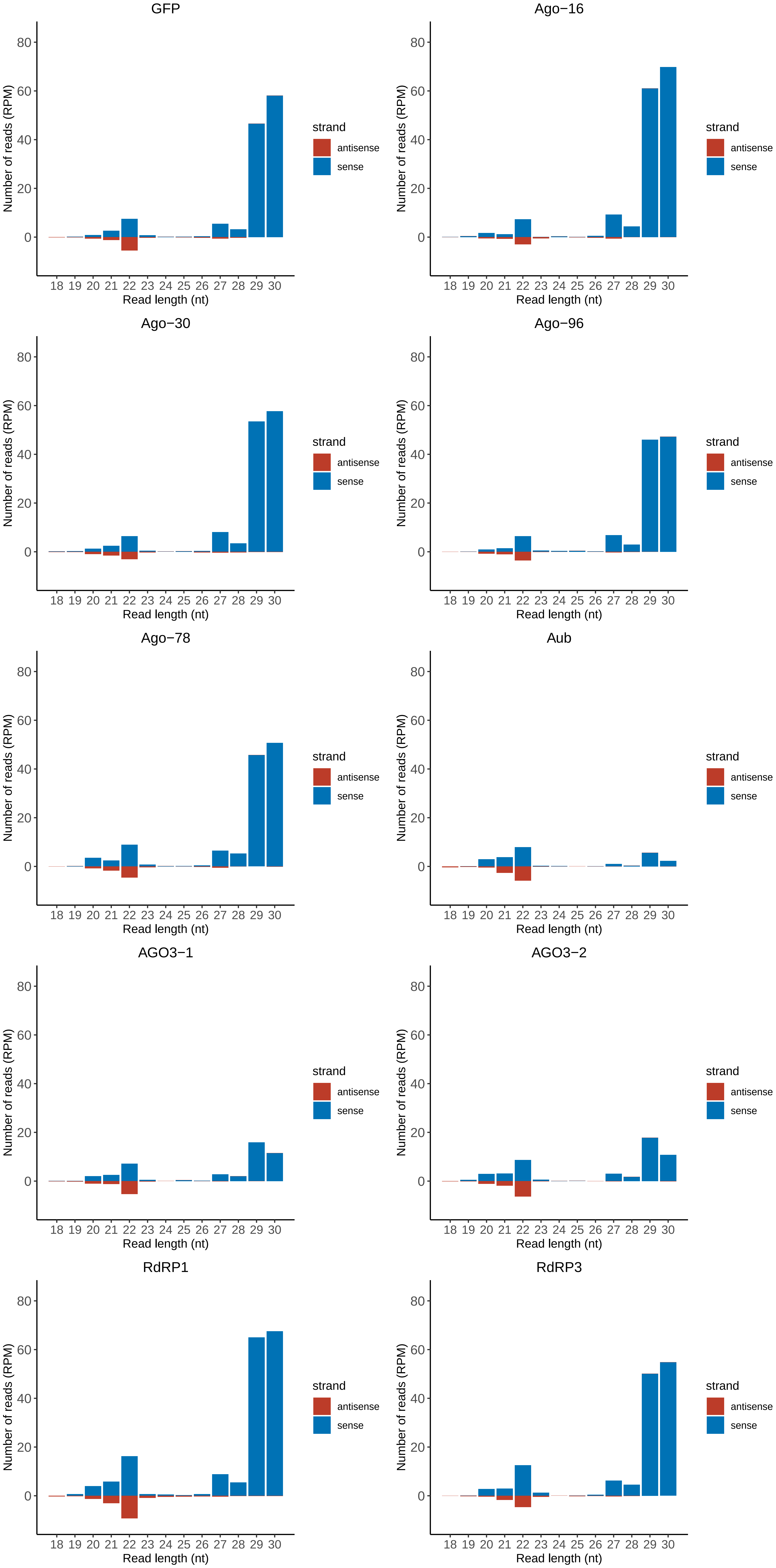
