## Supplemental PDF for "A novel eukaryotic RdRP-dependent small RNA pathway represses antiviral immunity by controlling an ERK pathway component in the black-legged tick"

### Gene annotation

- **Gene name**

    -- Uniprot (for protein coding gene)

    -- Vectorbase(for ncRNA gene)

    -- Description from EggNOG-mapper

- **GO term info**

    -- EggNOG-mapper(GO ids for protein coding gene)

    -- TRAPID(GO Ids for ncRNA gene)

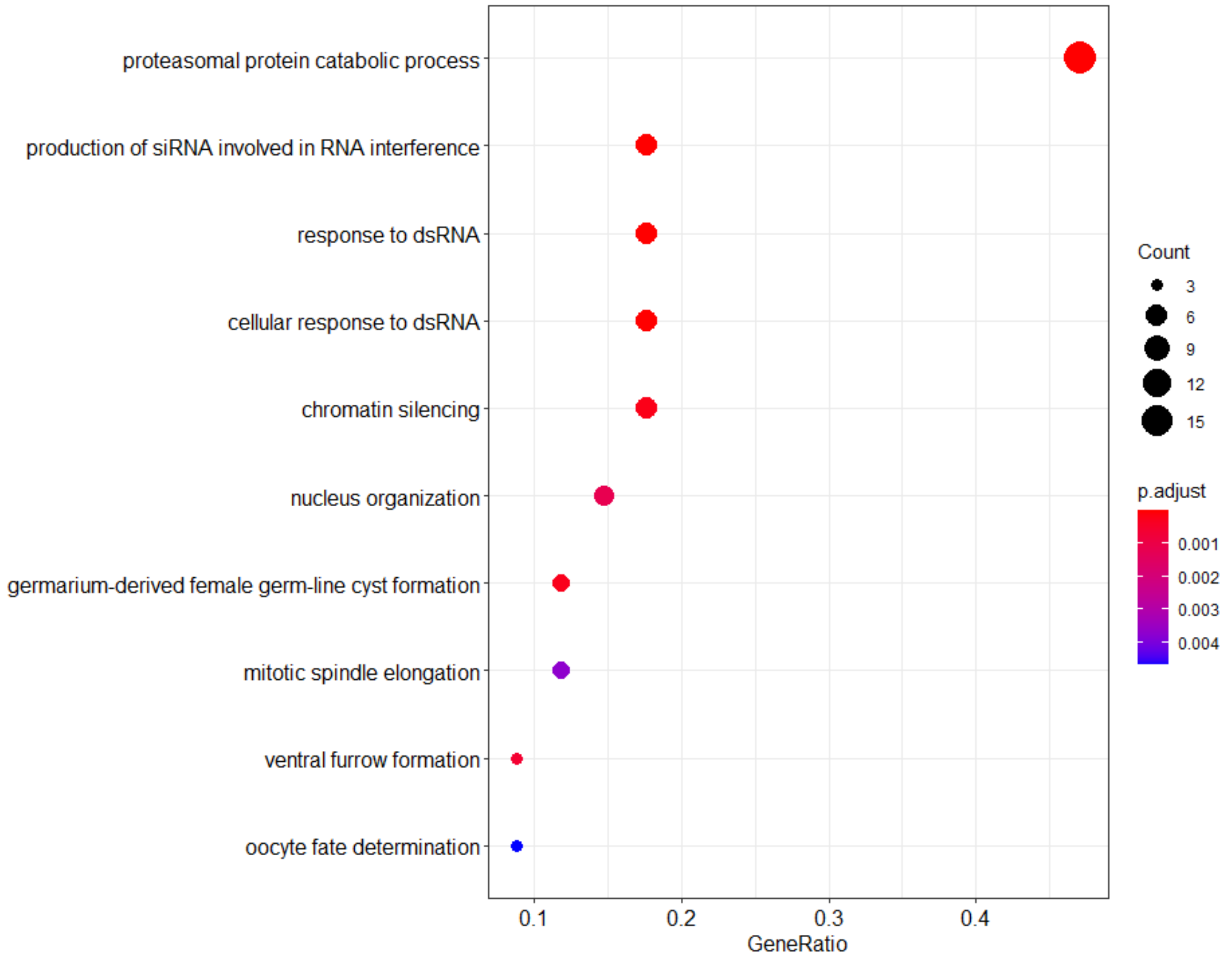


Dot plot showing the result of GO enrichment analysis (Biological Process) for misregulated genes upon Ago-16 knockdown.

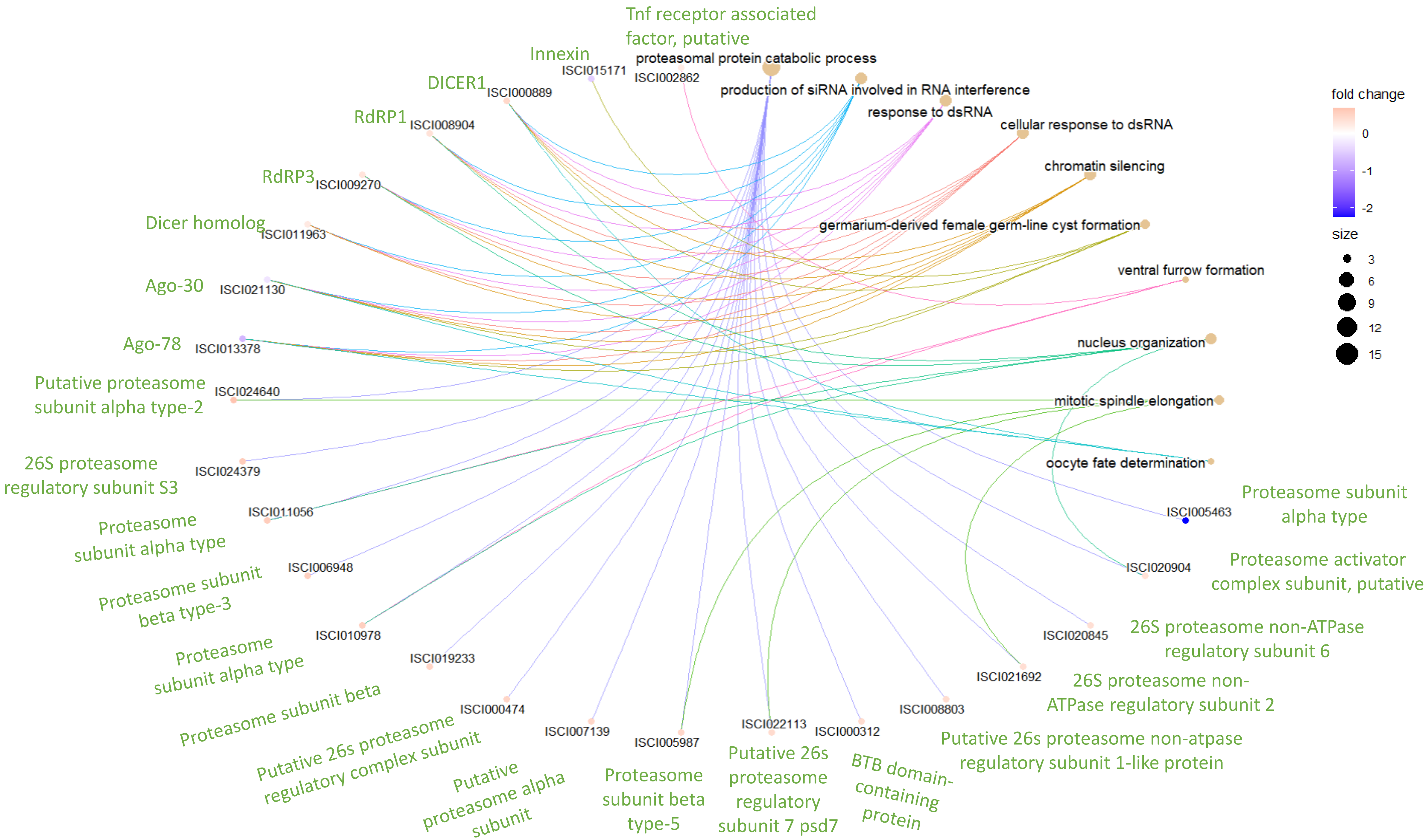


Linkage between GO terms(Biological Process) and genes misregulated upon Ago-16 knockdown.

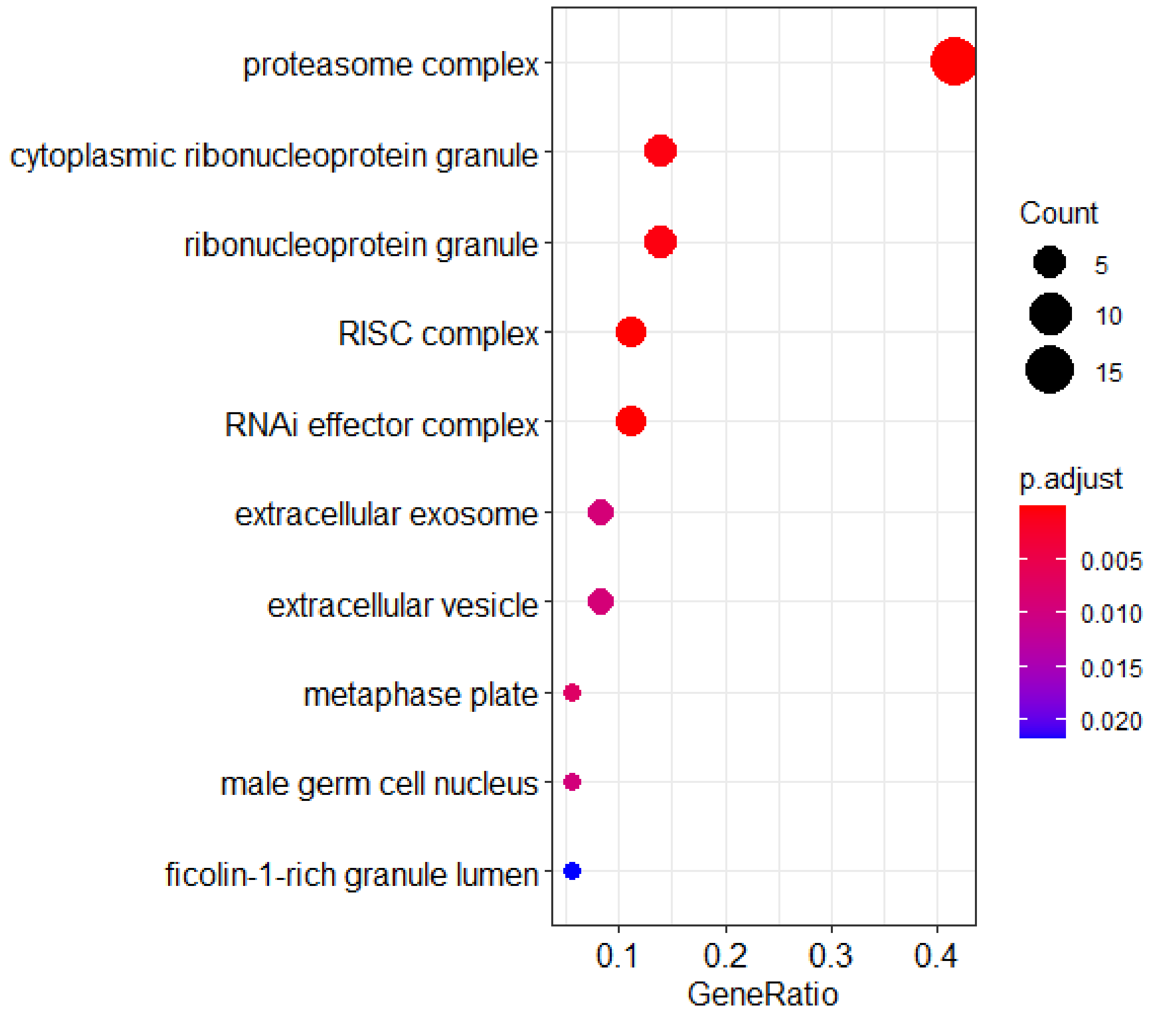


Dot plot showing the result of GO enrichment analysis (Cellular Component) for misregulated genes upon Ago-16 knockdown.

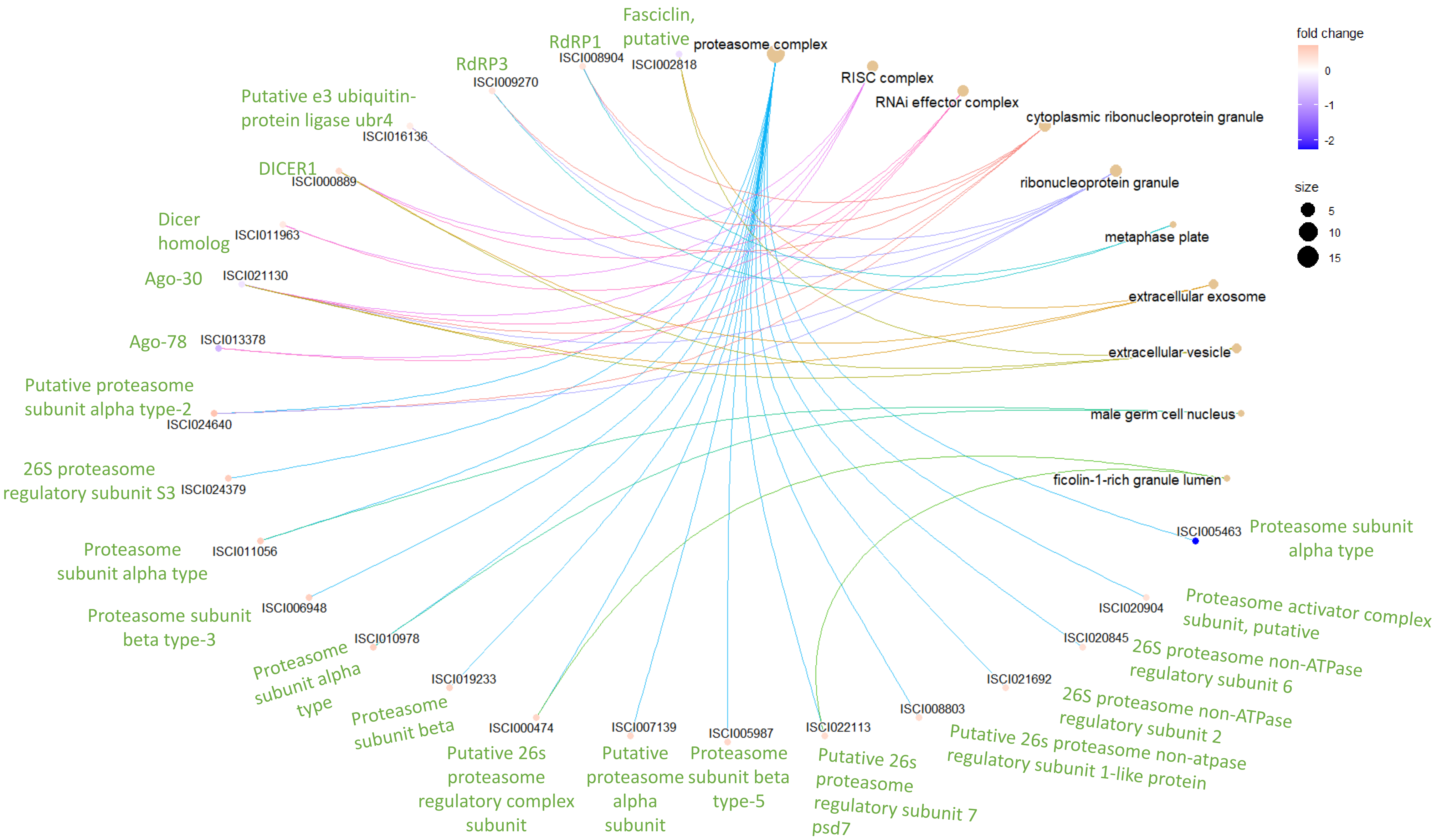


Linkage between GO terms(Cellular Component) and genes misregulated upon Ago-16 knockdown.

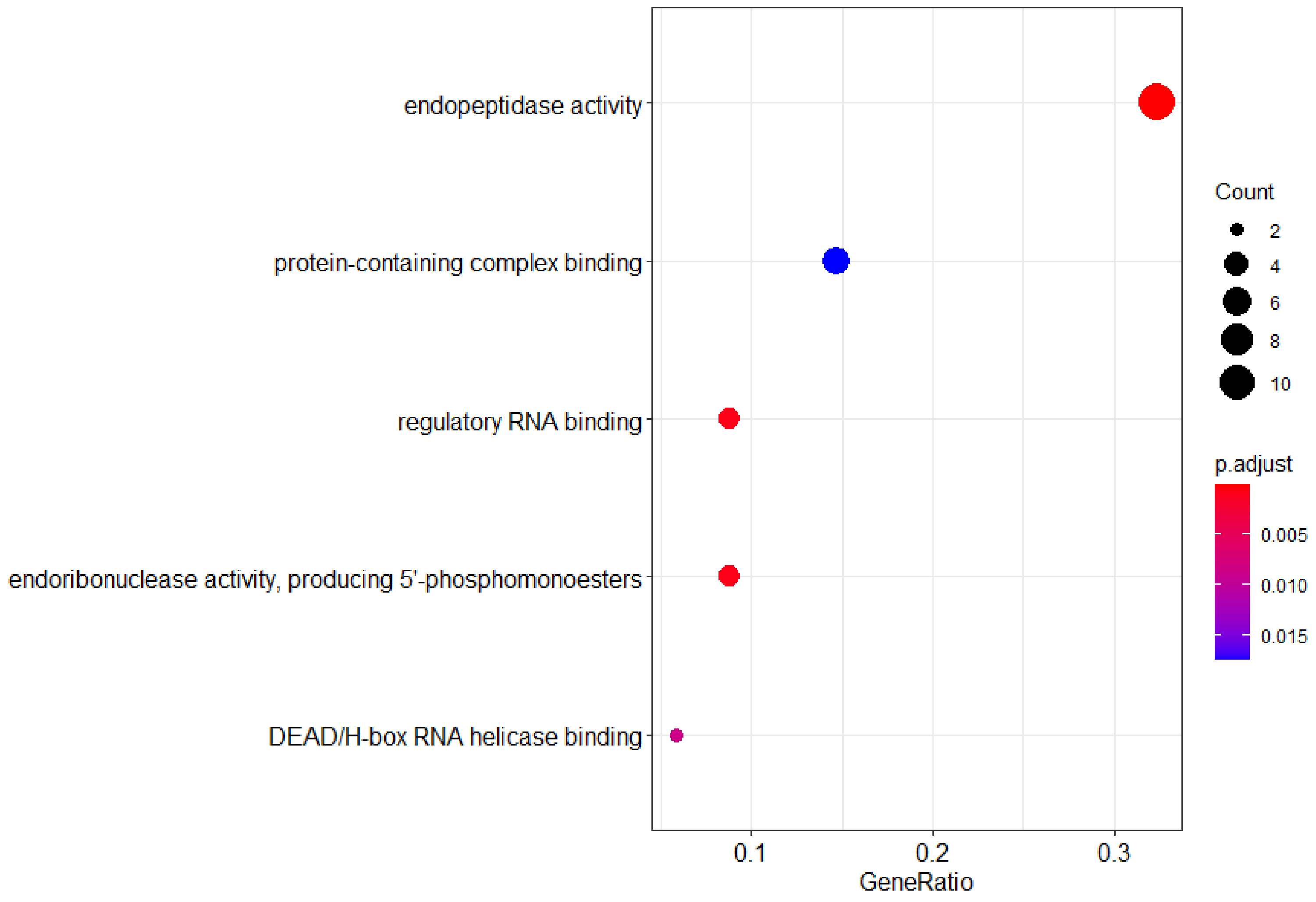


Dot plot showing the result of GO enrichment analysis (Molecular Function) for misregulated genes upon Ago-16 knockdown.

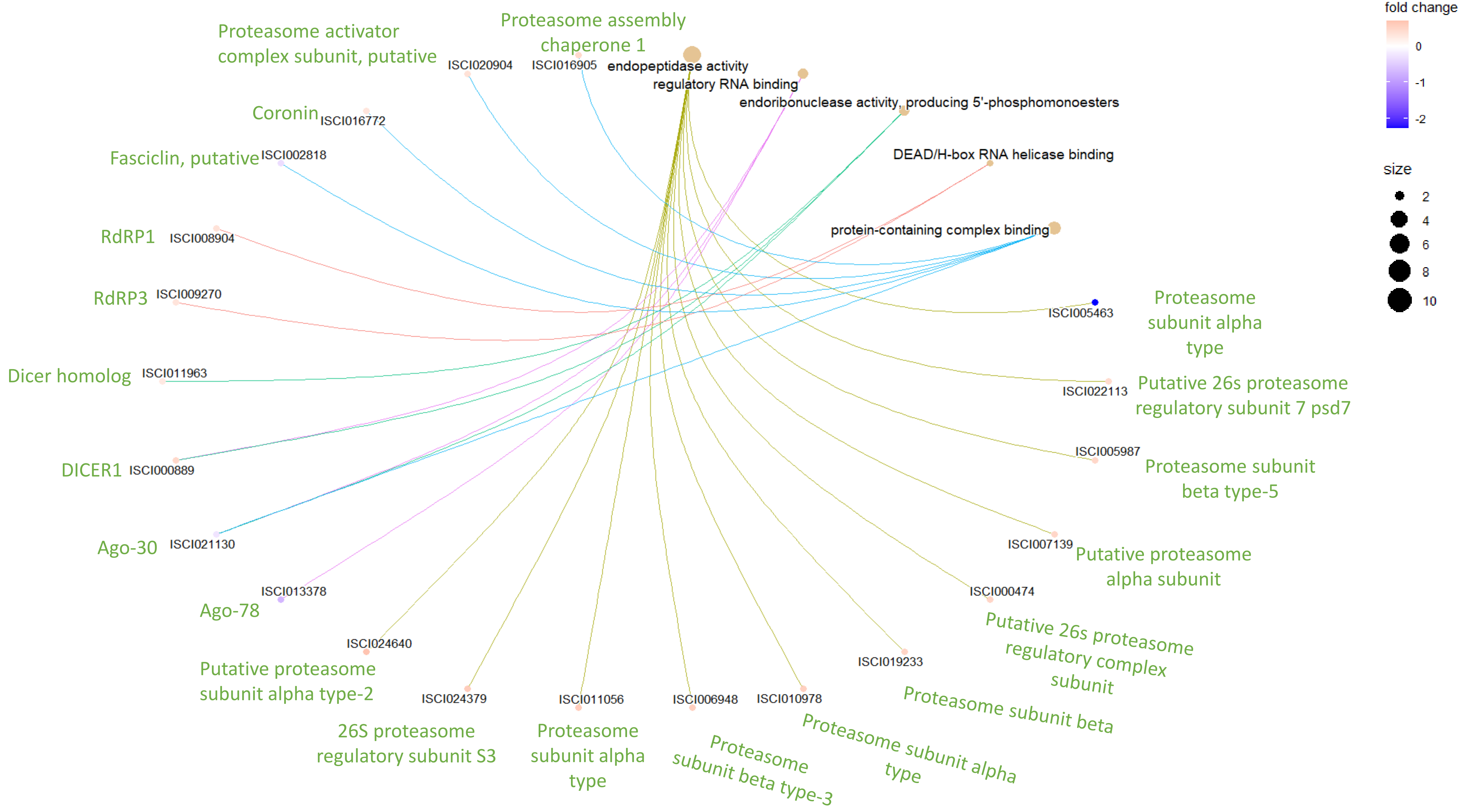


Linkage between GO terms(Molecular Function) and genes misregulated upon Ago-16 knockdown.

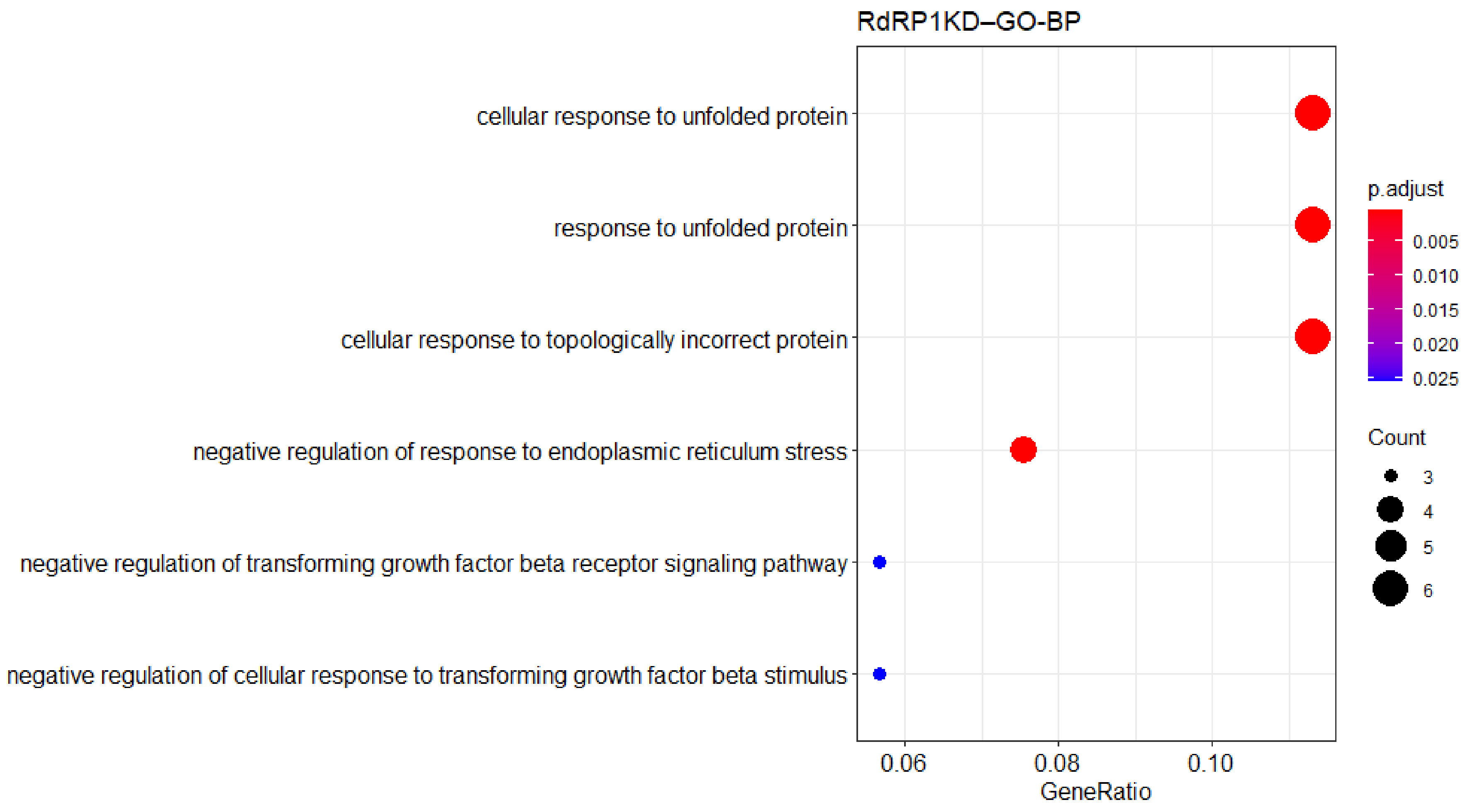


Dot plot showing the result of GO enrichment analysis (Biological Process) for misregulated genes upon RdRP1 knockdown.

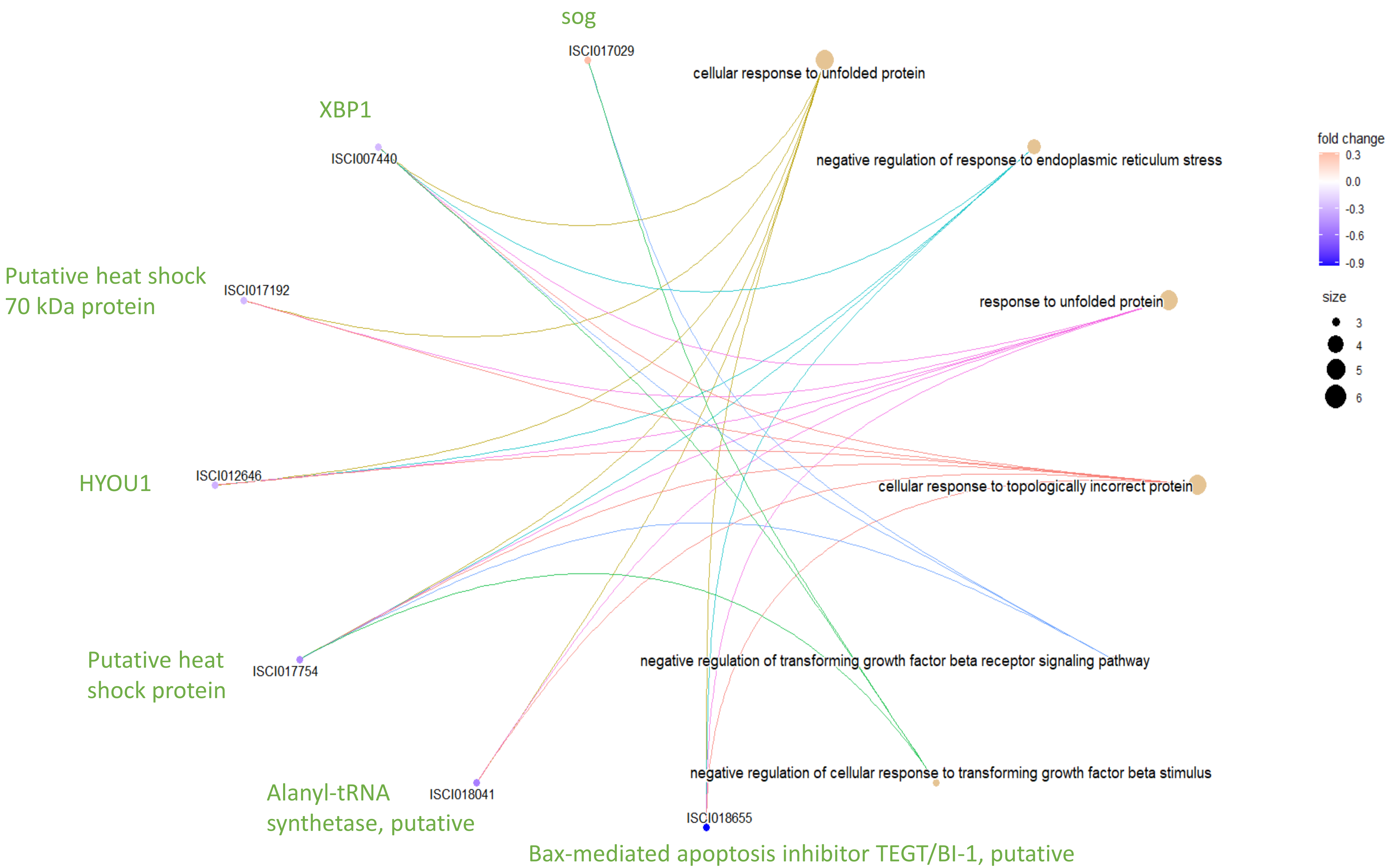


Linkage between GO terms(Biological Process) and genes misregulated upon RdRP1 knockdown.

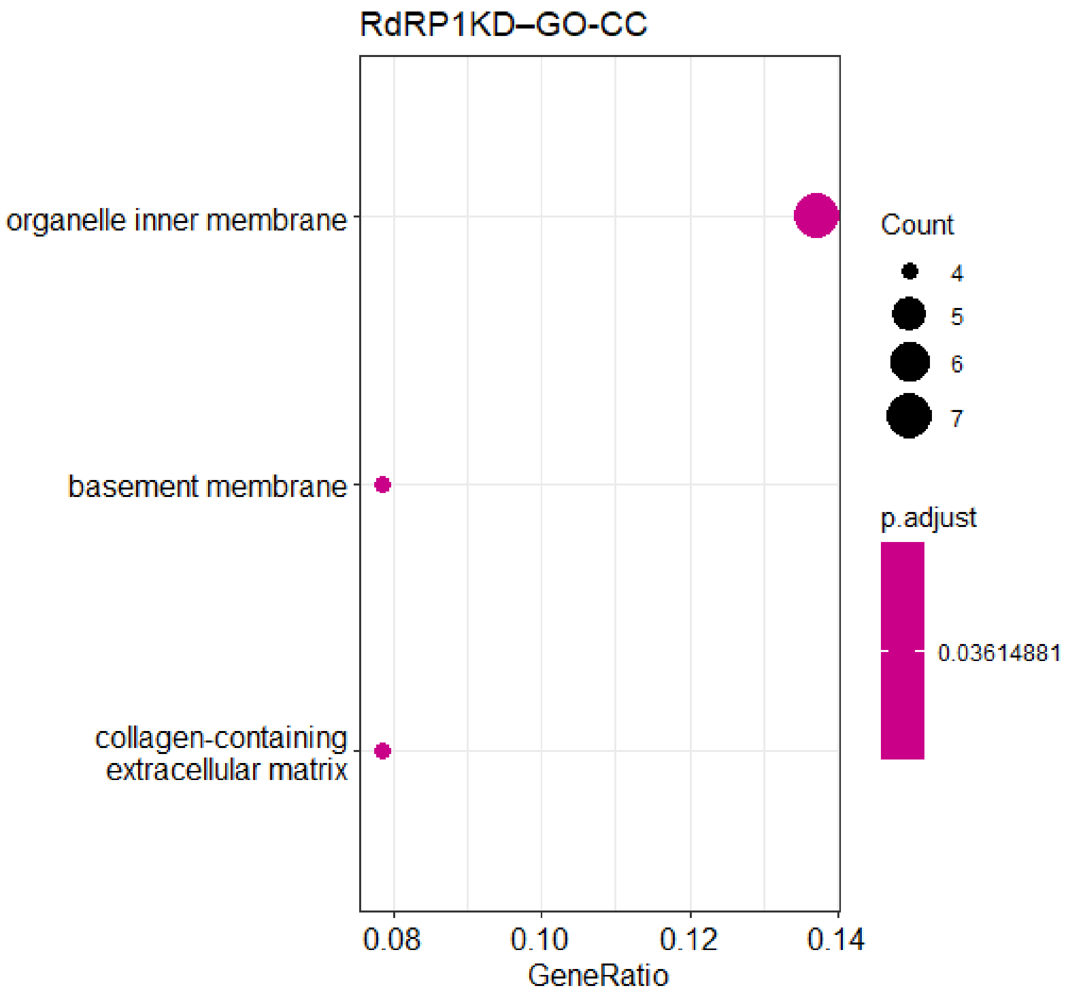


Dot plot showing the result of GO enrichment analysis (Cellular Component) for misregulated genes upon RdRP1 knockdown.

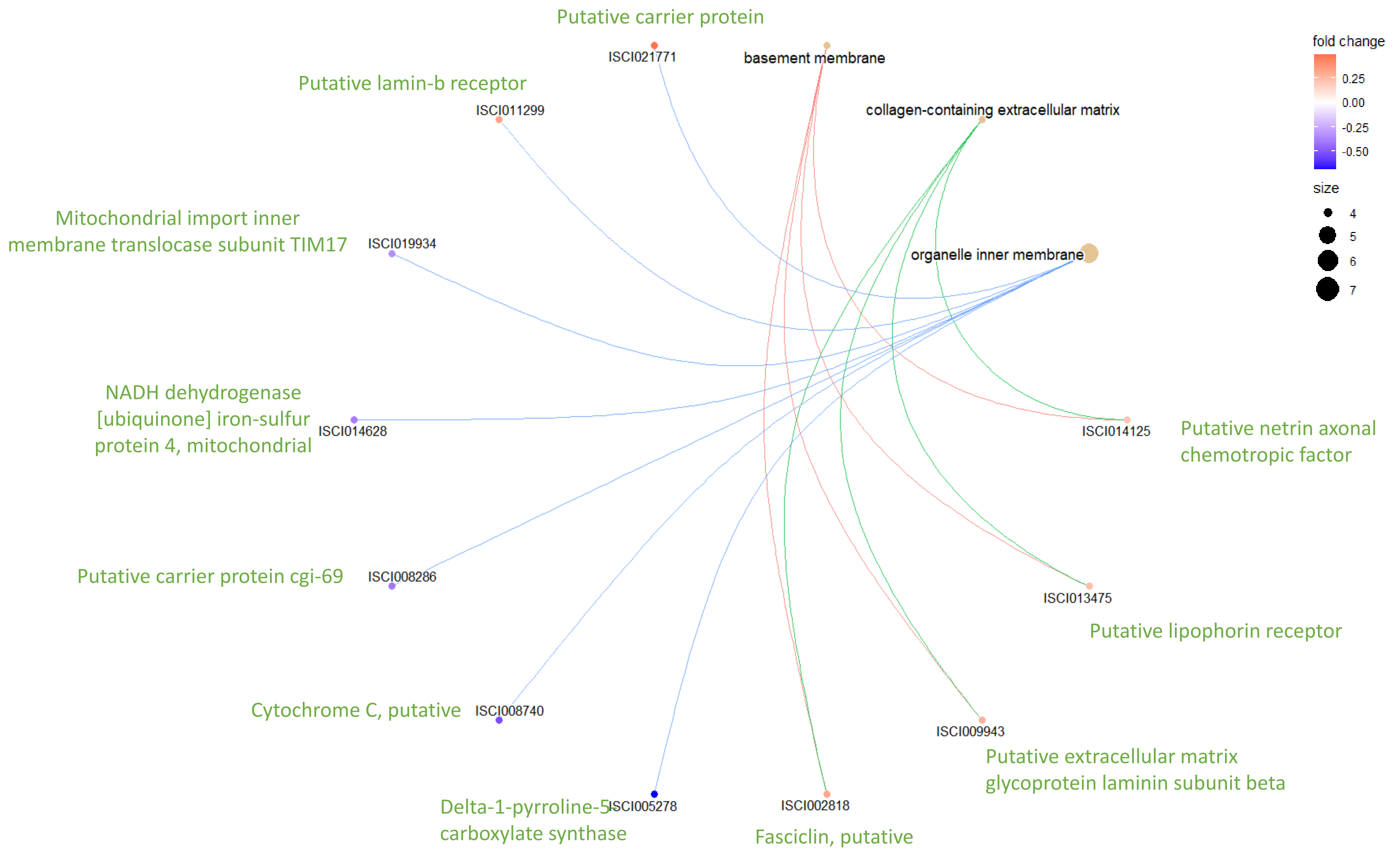


Linkage between GO terms(Cellular Component) and genes misregulated upon RdRP1 knockdown.

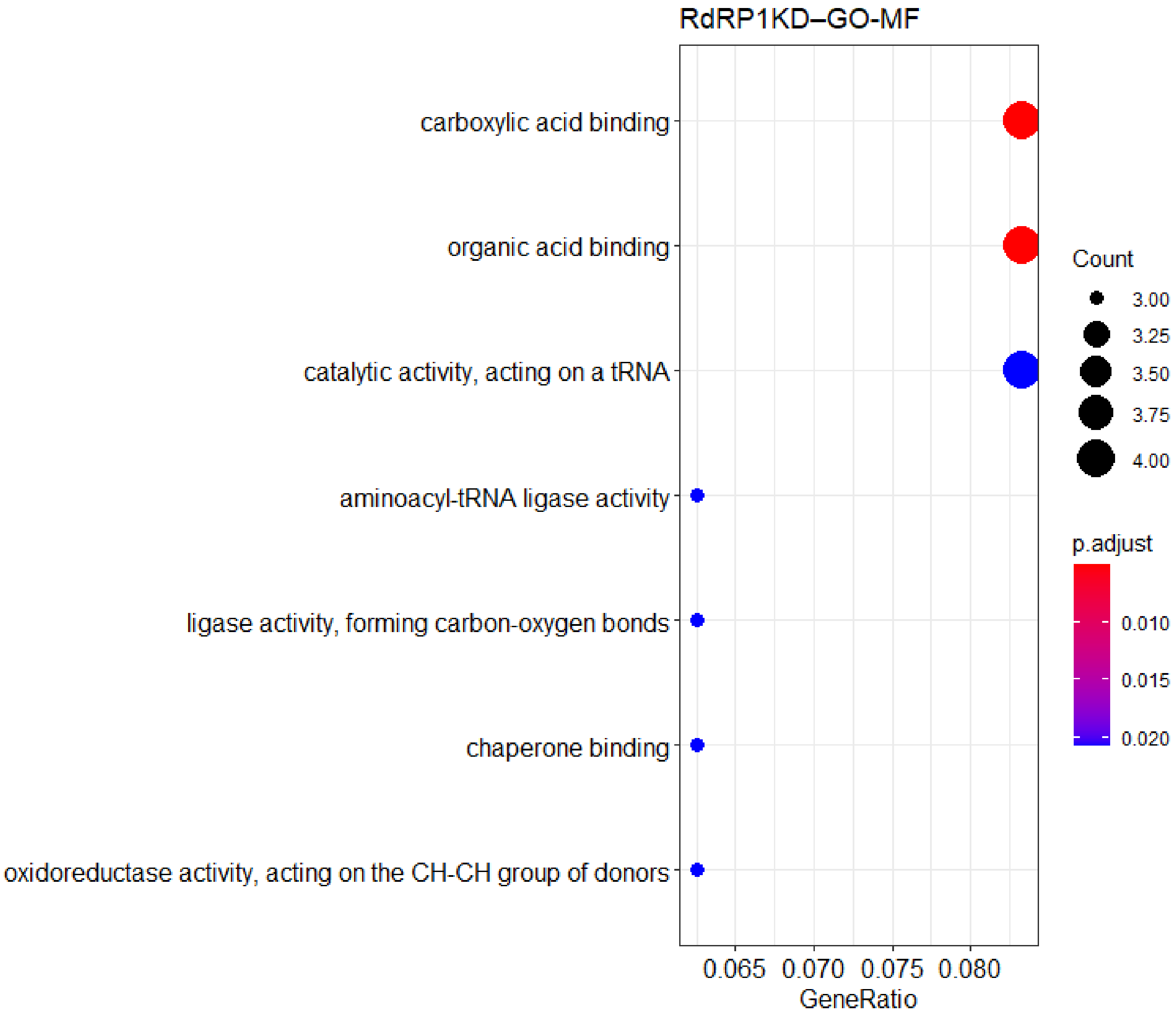


Dot plot showing the result of GO enrichment analysis (Molecular Function) for misregulated genes upon RdRP1 knockdown.

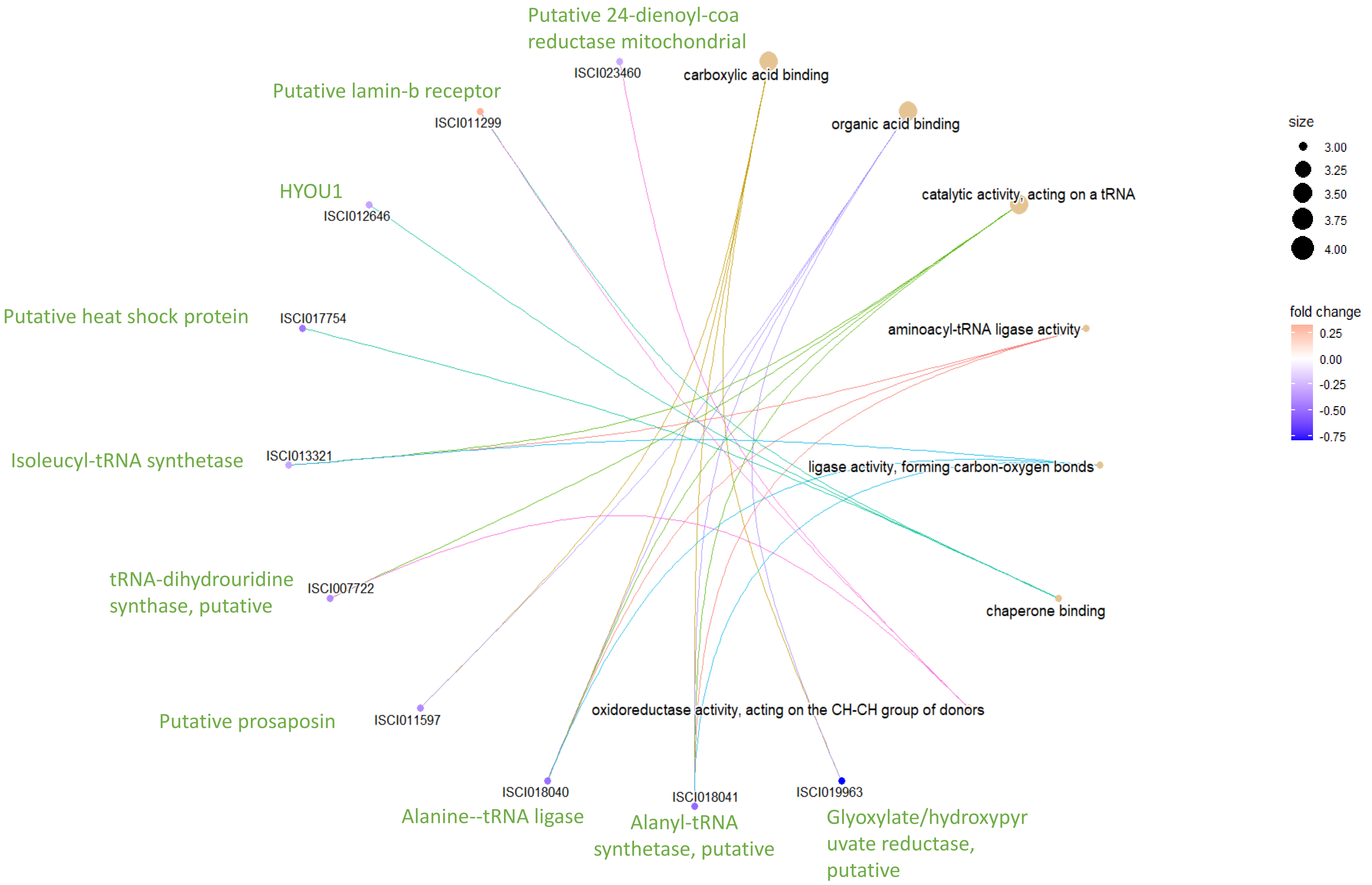


Linkage between GO terms(Molecular Function) and genes misregulated upon RdRP1 knockdown.

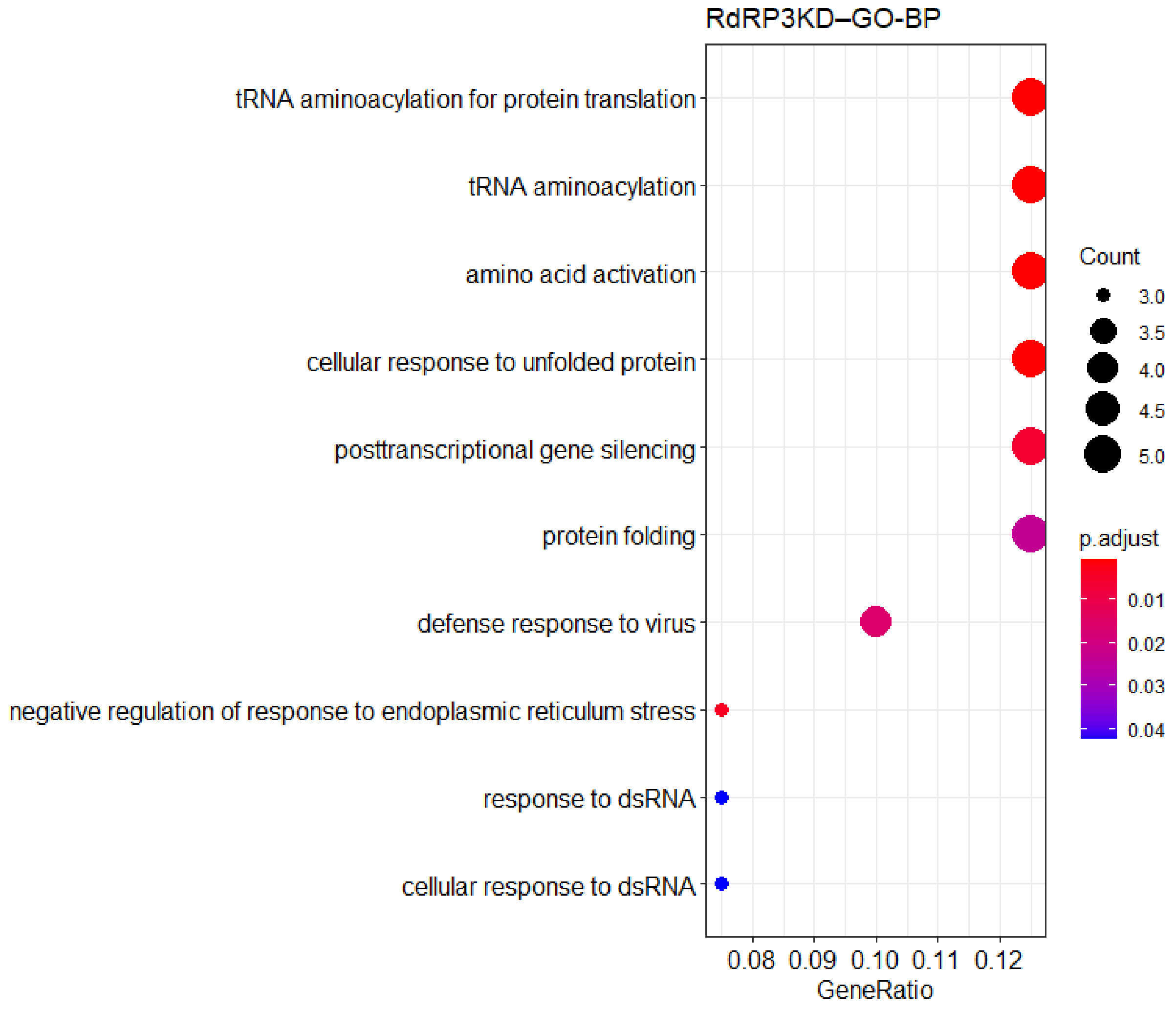


Dot plot showing the result of GO enrichment analysis (Biological Process) for misregulated genes upon RdRP3 knockdown.

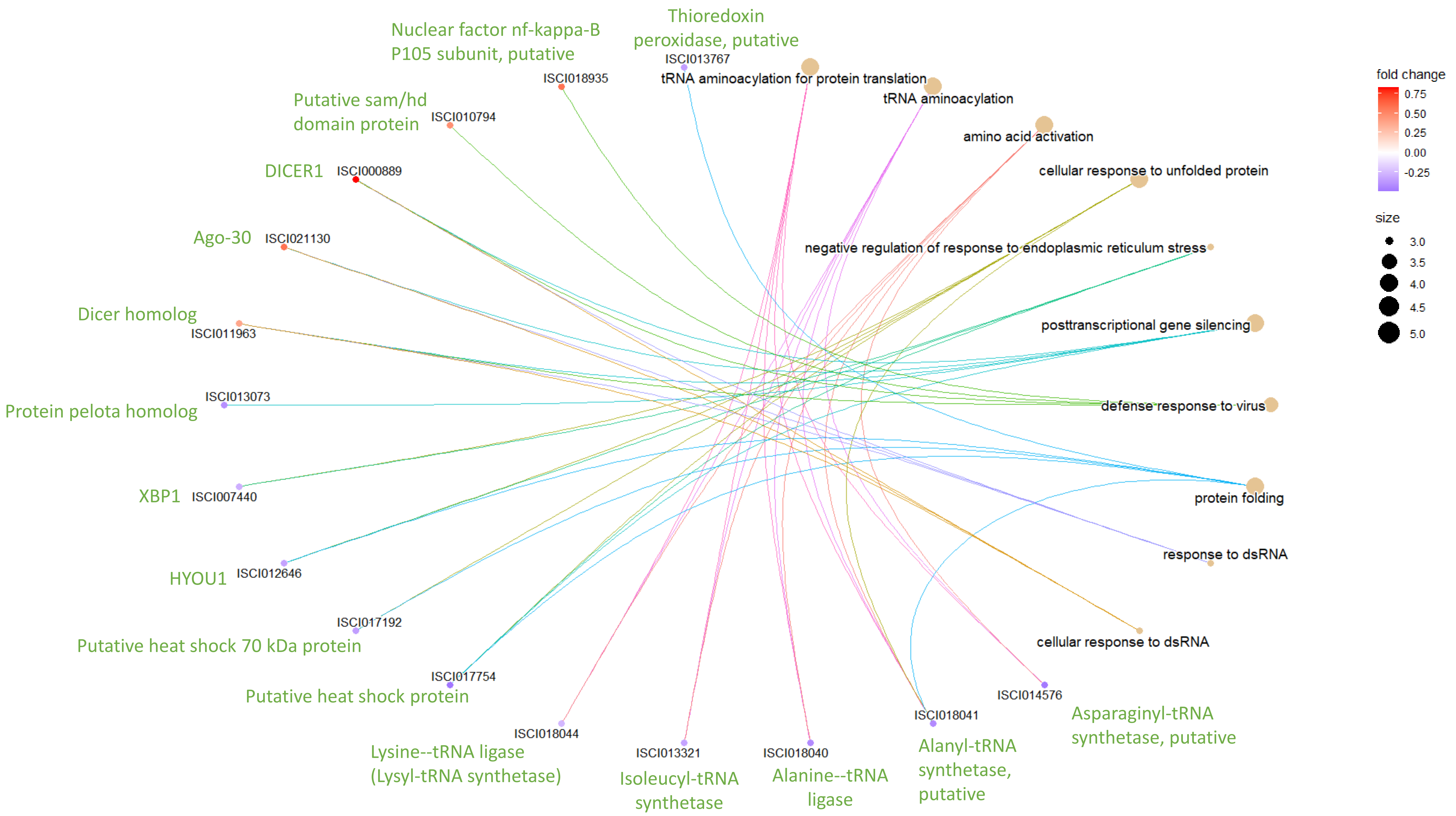


Linkage between GO terms(Biological Process) and genes misregulated upon RdRP3 knockdown.

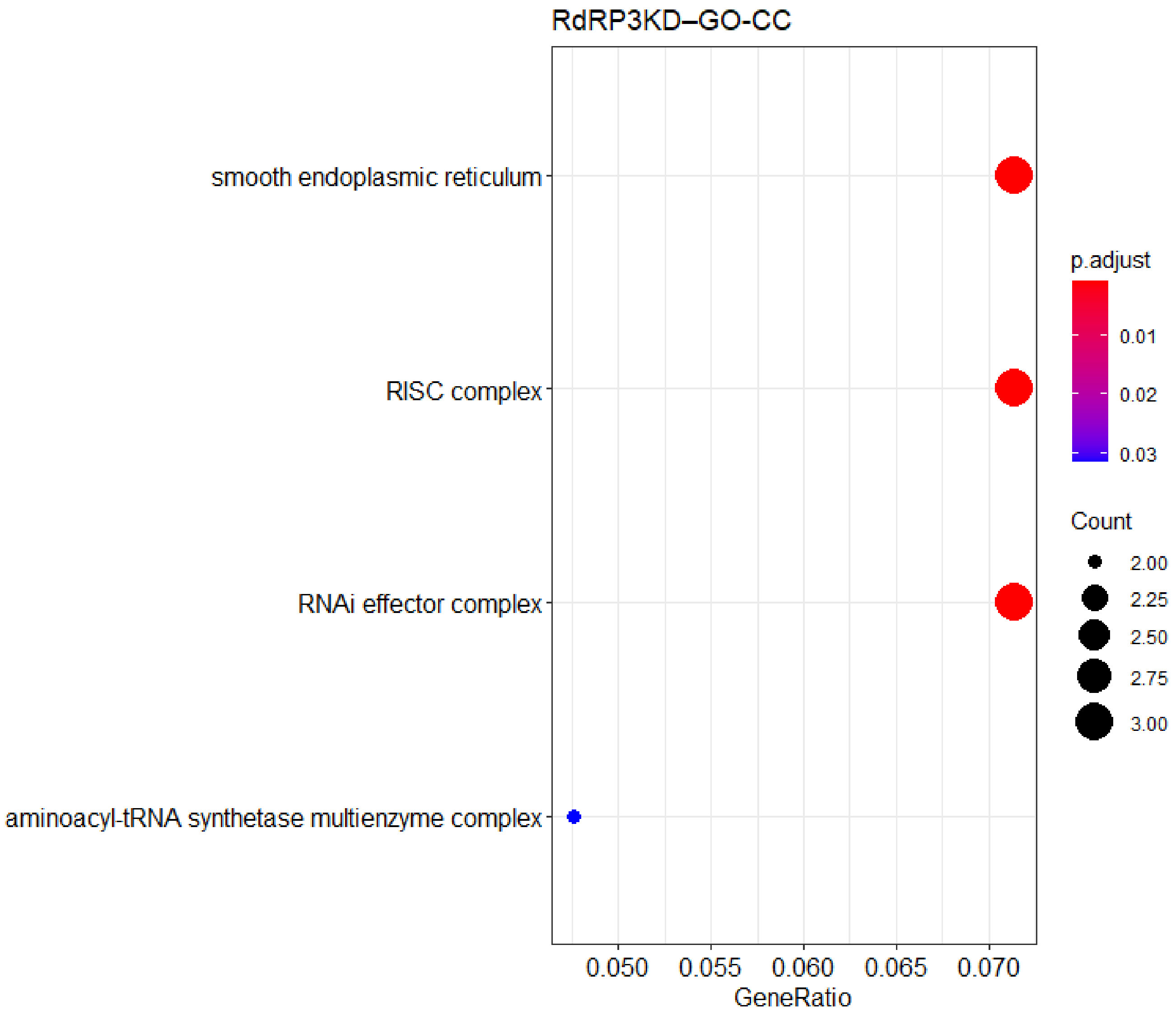


Dot plot showing the result of GO enrichment analysis (Cellular Component) for misregulated genes upon RdRP3 knockdown.

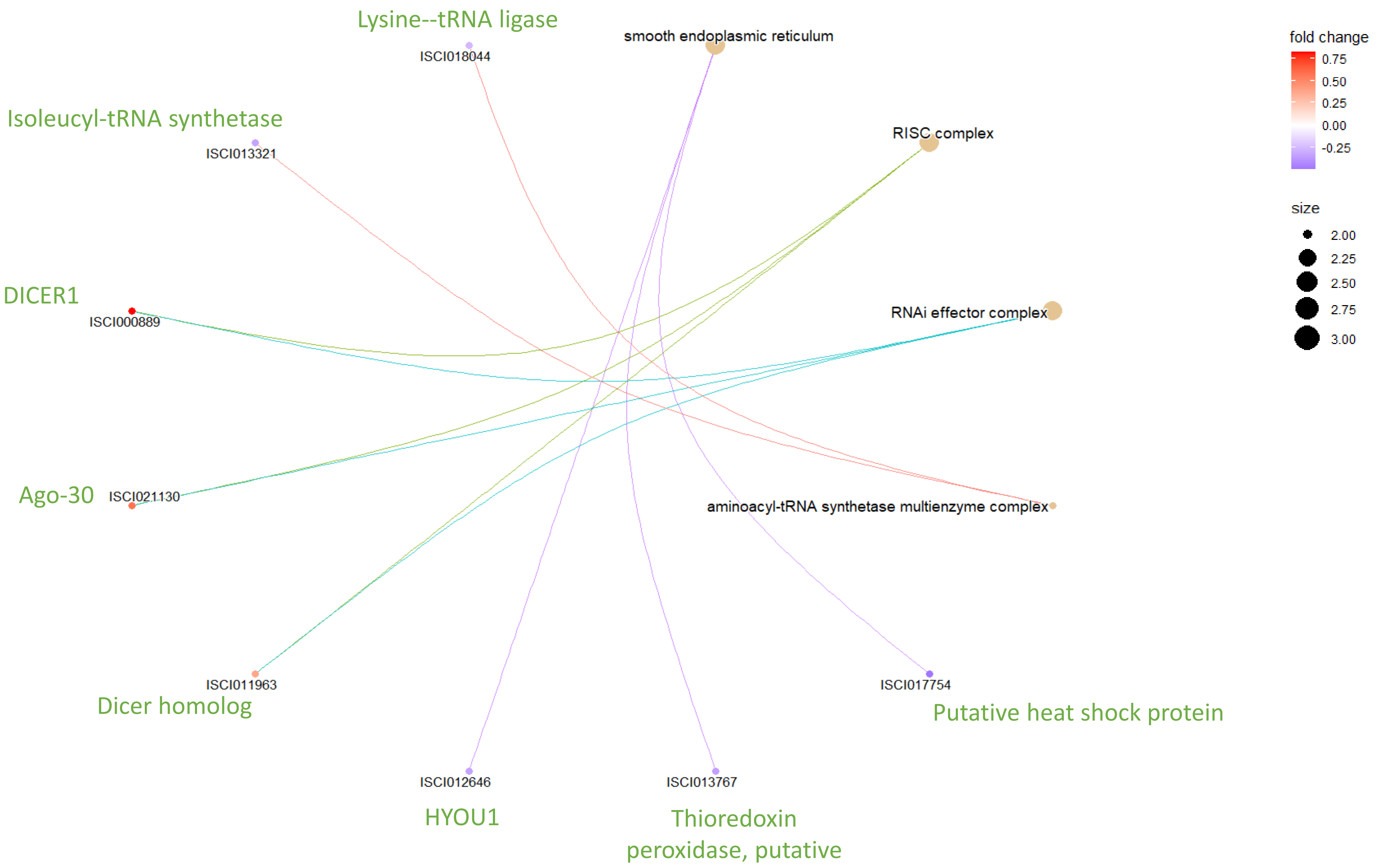


Linkage between GO terms(Cellular Component) and genes misregulated upon RdRP3 knockdown.

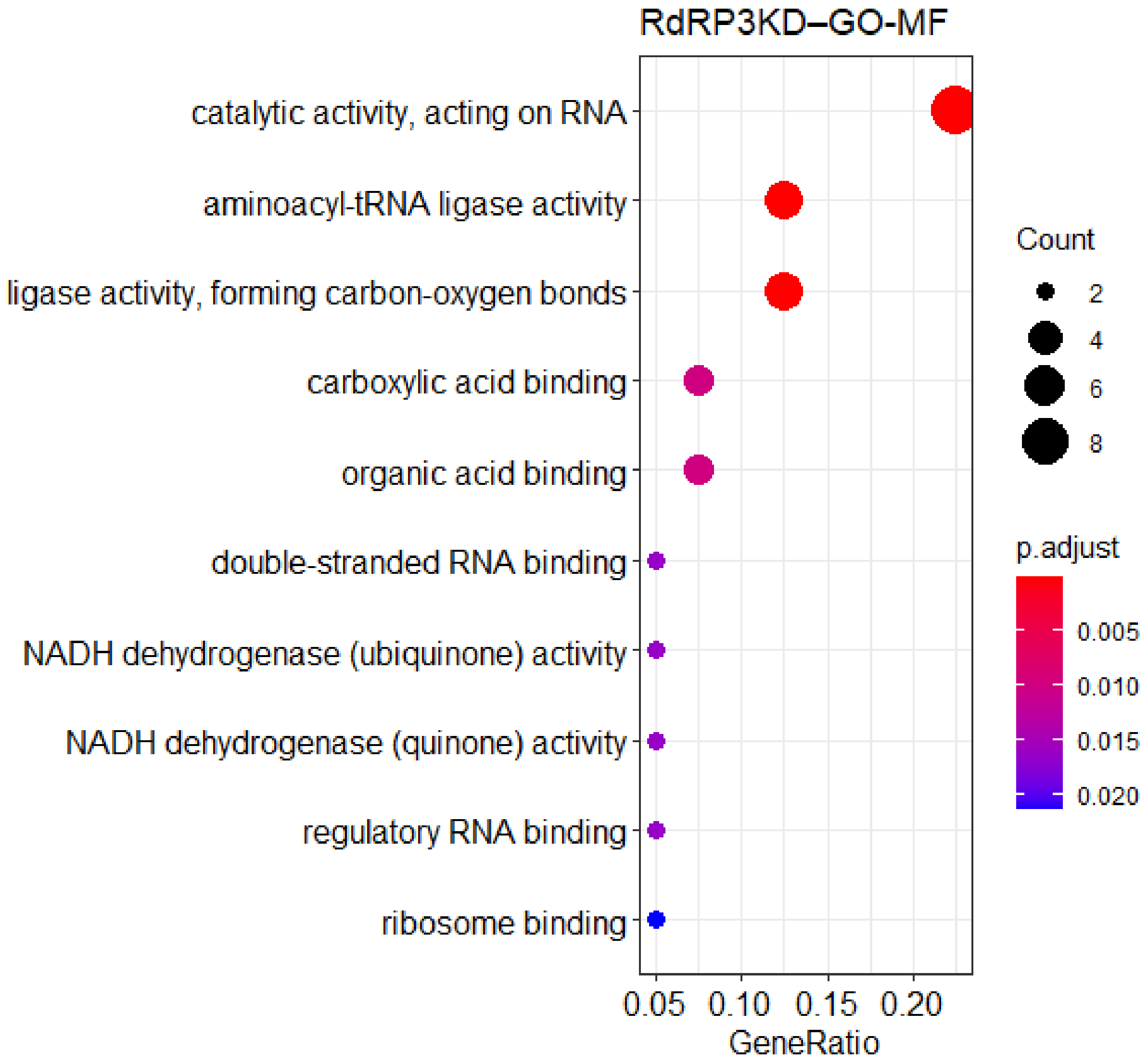


Dot plot showing the result of GO enrichment analysis (Molecular Function) for misregulated genes upon RdRP3 knockdown.

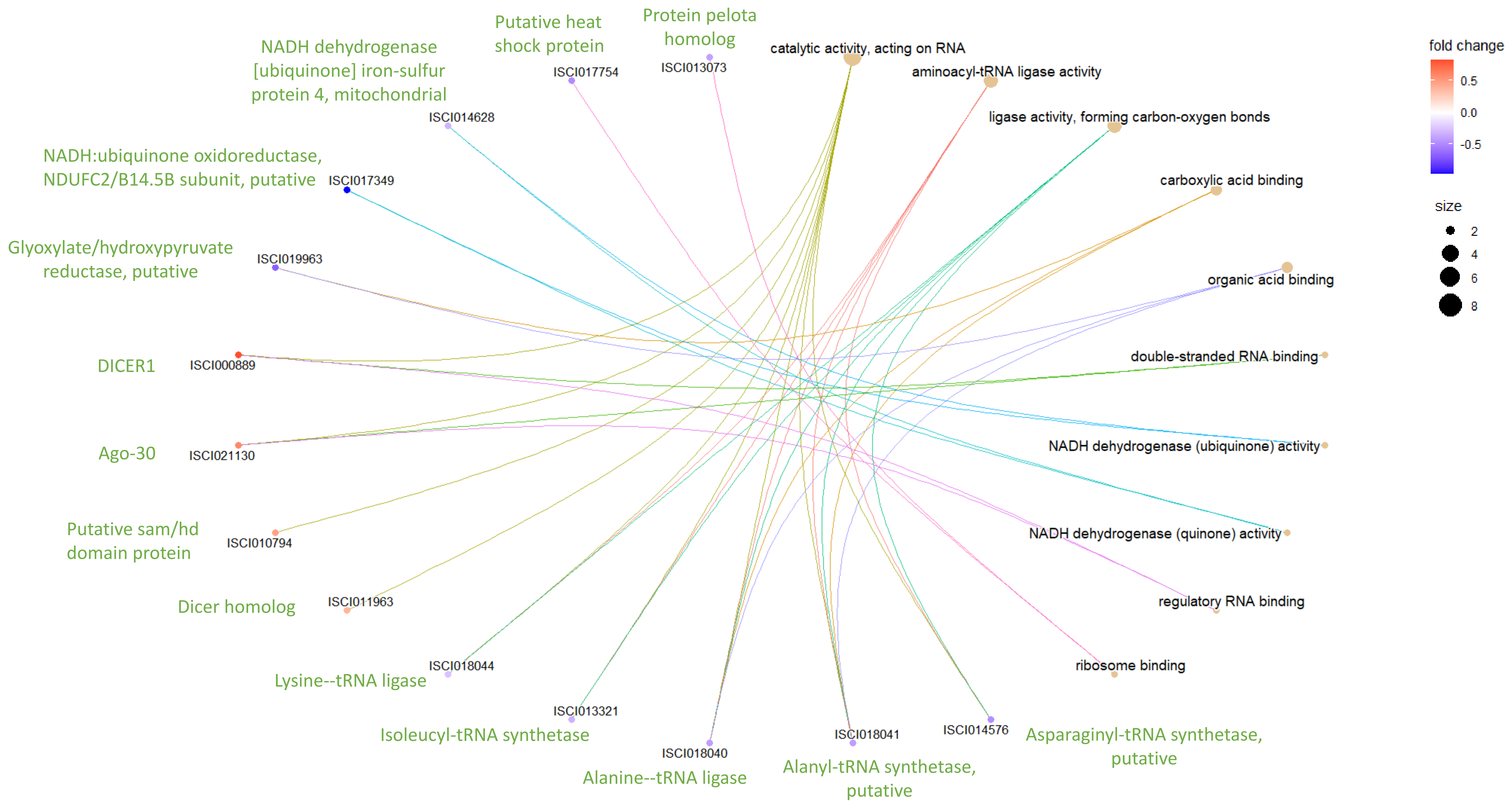


Linkage between GO terms(Molecular Function) and genes misregulated upon RdRP3 knockdown.
